# GPU Accelerated Adaptive Banded Event Alignment for Rapid Comparative Nanopore Signal Analysis

**DOI:** 10.1101/756122

**Authors:** Hasindu Gamaarachchi, Chun Wai Lam, Gihan Jayatilaka, Hiruna Samarakoon, Jared T. Simpson, Martin A. Smith, Sri Parameswaran

## Abstract

Nanopore sequencing has the potential to revolutionise genomics by realising portable, real-time sequencing applications, including point-of-care diagnostics and in-the-field genotyping. Achieving these applications requires efficient bioinformatic algorithms for the analysis of raw nanopore signal data. For instance, comparing raw nanopore signals to a biological reference sequence is a computationally complex task despite leveraging a dynamic programming algorithm for Adaptive Banded Event Alignment (ABEA)—a commonly used approach to polish sequencing data and identify non-standard nucleotides, such as measuring DNA methylation. Here, we parallelise and optimise an implementation of the ABEA algorithm (termed *f5c*) to efficiently run on heterogeneous CPU-GPU architectures. By optimising memory, compute and load balancing between CPU and GPU, we demonstrate how *f5c* can perform ~3-5× faster than the original implementation of ABEA in the *Nanopolish* software package. We also show that *f5c* enables DNA methylation detection on-the-fly using an embedded System on Chip (SoC) equipped with GPUs. Our work not only demonstrates that complex genomics analyses can be performed on lightweight computing systems, but also benefits High-Performance Computing (HPC). The associated source code for *f5c* along with GPU optimised ABEA is available at https://github.com/hasindu2008/f5c.

## 1 Introduction

Advances in genomic technologies have given rise to a new era in biomedical sciences, improving the feasibility and accessibility of rapid species identification, accurate clinical diagnostics, and specialised therapeutics, amongst other applications. Whole genome sequencing involves ‘reading’ the entire DNA sequence of a cell, revealing the genetic variation that underlies biological diversity and the onset of disease. A human genome encompasses two copies of ~3.2 billion DNA nucleotides, or ‘letters’. Therefore, analysing the data generated by contemporary high-throughput sequencing technologies typically requires high-performance computing support.

The latest generation (third generation) of sequencing technologies can generate ultra-long DNA ‘reads’ from single molecules in real-time. In particular, Oxford Nanopore Technologies (ONT) manufacture a pocket-sized sequencer called MinION (Fig. 1), a relatively inexpensive and portable sequencing device capable of sequencing in-the-field (e.g. remote area with no network connectivity) or at the point-of-care (e.g. hospital, clinic, pharmacy).

**Figure 1:**
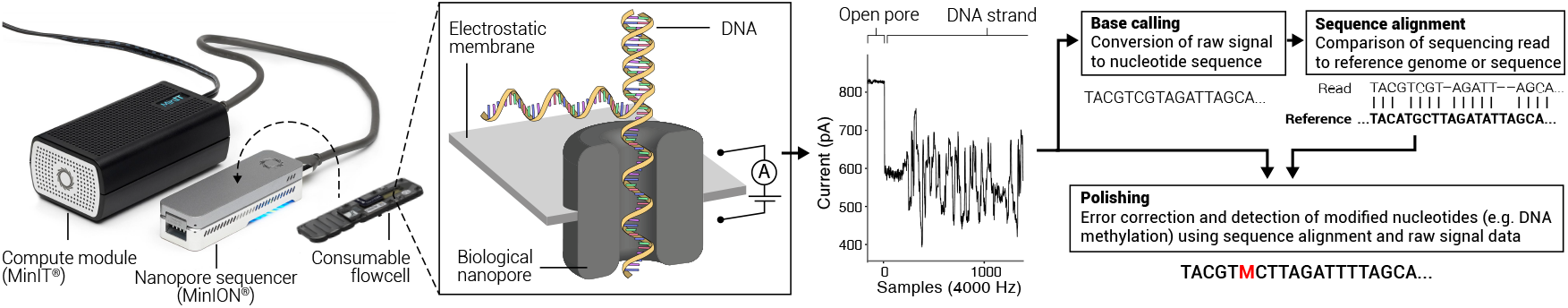
Nanopore portable sequencer and associated data analysis

In contrast to ‘second’ generation sequencers, which produce highly accurate short reads through enzymatic synthesis of the complementary strand of DNA, nanopore sequencing measures characteristic disruptions in the electric current (referred to hereafter as *raw signal*) when DNA passes through a biological nanopore (Fig. 1). A consumable flowcell containing an array of hundreds or thousands of such nanopores is loaded into the sequencing device (e.g. MinION), which is coupled to a generic (e.g. laptop) or dedicated (e.g. MinIT) compute module to acquire sequence data and base-call (the process of converting the raw signal to nucleotide characters) in parallel.

Nanopore sequencing offers several benefits over other technologies, including ultra-long reads (>1 Mbases), detection of non-standard DNA bases or biochemically modified DNA bases, and real-time analysis, at the expense of a higher error-rate, which is predominantly caused by the conversion of the raw signal into DNA bases via probabilistic models (referred to as ‘base-calling’). To overcome base-calling errors, the raw signal can be revisited *a posteriori* (see the polishing step in Fig. 1). Such *a posteriori* ‘polishing’ can correct for base-calling errors by aligning raw signal to a biological reference sequence, thus identifying idiosyncrasies in the raw signal by comparing observed signal levels to expected levels at all aligned positions. This process can also reveal base substitutions (i.e. mutations) or base modifications such as 5-methylcytosine (5mC), a dynamic biochemical modification of DNA that is associated with genetic activity and regulation. Detecting 5mC bases is important for the study of DNA methylation in the field of epigenetics.

A crucial algorithmic component of polishing is the alignment of raw signal–a time series of electric current to a biological reference sequence. One of the first raw nanopore signal alignment is implemented in the popular tool *Nanopolish* [1], which employs a dynamic programming strategy referred to as Adaptive Banded Event Alignment (ABEA). ABEA is one of the most time consuming steps when analysing raw nanopore data. For instance, when performing methylation detection with *Nanopolish*, the ABEA step consumes ~70% of the total CPU time. Consequently, it is important to investigate strategies to reduce the runtime of ABEA to improve the turnaround time of certain nanopore sequencing applications, such as real-time polishing or methylation detection.

In this study, we dissect the ABEA algorithm to optimise and parallelise its use on diverse hardware platforms, including Graphics Processing Units (GPUs). Adapting this ABEA algorithm for the GPU is not a straight forward task due to three main factors: (i) Read lengths vary significantly (from ~100 to >1M bases), thus requiring millions to billions of dynamic memory allocations—an expensive operation in GPUs. (ii) Inefficient memory access patterns which are not ideal for the GPUs having relatively less powerful and smaller caches (compared to CPUs) result in frequent instruction stalls. (iii) Varying read lengths cause irregular utilisation of the GPU cores.

We overcome the above mentioned challenges by: (i) employing a custom heuristic-based memory allocation scheme; (ii) tailoring the algorithm and the GPU user-managed cache to exploit cache-friendly memory access patterns; and, (iii) using a heuristic based work-partitioning and load-balancing scheme between the CPU and GPU.

We demonstrate the utility of our GPU optimised ABEA by incorporating to a completely reengineered version of the popular methylation detection tool *Nanopolish*. First, we re-engineered the original *Nanopolish* methylation detection tool to efficiently utilise existing CPU resources, which we refer to as *f5c*. Then, we incorporated our GPU optimised ABEA algorithm into the re-engineered *f5c*. We demonstrate how *f5c* enables DNA methylation detection using nanopore sequencers in real-time (i.e. on-the-fly processing of the output) by using an embedded System on Chip (SoC) equipped with a GPU. We also demonstrate how *f5c* benefits a wide range of computing systems from embedded systems and laptops to workstations and high performance servers.

The key contributions of this paper are: (i) the first example of GPU acceleration and optimisation of raw signal alignment algorithm; (ii) *f5c*, a re-engineered and optimised version of the popular DNA methylation detection tool *Nanopolish*; and, (iii) real-time detection of DNA methylation using a lightweight and portable embedded system (previously only possible on high-performance servers).

In the rest of the paper, we discuss the background of nanopore sequencing and ABEA algorithm in Section 2, related work in Section 3, methodology in Section 4, results in Section 5, followed by the discussion and future work in Section 6. The associated tool that includes the GPU-based acceleration is available at https://github.com/hasindu2008/f5c.

## 2 Background

Basic terms and concepts of DNA sequencing and data analysis are given in Section 2.1. Section 2.2 briefly explains methylation calling, an example nanopore data analysis workflow. Section 2.3 explains the Adaptive Banded Event Alignment (ABEA) algorithm, the algorithm which is optimised in this paper for execution on a CPU-GPU heterogeneous architecture. In Section 2.4, a brief account of GPU architectures and the programming methods for GPUs.

### 2.1 Nanopore sequencing and analysis

#### 2.1.1 Whole genome sequencing

The *genome* is a long sequence composed of four types of nucleotide bases: adenine (A), cytosine (C), guanine (G) and thymine (T). Nucleotide bases will be simply referred to as *bases* hereafter. The human genome is around 3.2 gigabases (Gbases) long and is composed of 23 pairs of chromosomes (46 chromosomes in total), where each chromosome is a single molecule of continuous deoxyribonucleic acid (DNA) polymer. The process of reading strings of contiguous bases is called *sequencing*, and the resulting strings of bases are called *reads*. In order to be sequenced, DNA molecules must be extracted and purified from cells before being biochemically prepared for sequencing. This *library preparation* process can fragment chromosomes (especially large ones) into smaller segments–either intentionally or incidentally–which are ‘read’ by the sequencer. Given that samples contain multiple cells, and thus several distinct DNA molecules, and that sequencing may introduce errors, it is desirable to generate enough reads to cover a particular position several times. The average number of reads at a given position is termed sequencing *coverage*. High coverage facilitates the characterisation of genetic variation and correct for errors. A human genome sequenced at around 20× average coverage corresponds to around 64 Gbases of sequencing reads.

#### 2.1.2 DNA methylation

DNA undergoes naturally regulated biochemical modification through the addition of a methyl group to certain bases. Methylation is reversible and can control the activity of a DNA segment, such as turning the expression of genes on or off, without modifying the genetic code itself—a process called *epigenetic* regulation. DNA methylation is dynamically regulated during normal biological development and in function of environmental factors; it plays an important role in disease aetiology and clinical diagnostics [2, 3, 4]. Methylation of cytosine (‘C’) bases is of particular interest in human biology, where CpG dinucleotides (‘C’ base followed by a ‘G’ base) are dynamically methylated in normal development and disease [5, 6, 7].

#### 2.1.3 Nanopore sequencing and the raw signal

Nanopore sequencing is a third generation sequencing technology that involves physical observation of atomic properties of DNA fragments using a nanometer scale biological pore coupled to an ammeter (see Fig. 1). The pore acts as a bottleneck to generate characteristic disruptions in ionic current (in the range of pico-amperes) that are indicative of the molecules passing through the pore. The size and nature of the pore influence the measured instantaneous current and how it is subsequently analysed. Oxford Nanopore Technologies (ONT) sequencing devices measure DNA strand passing through biological nanopores composed of recombinant (or ‘designer’) proteins at an average speed of ~450 bases/s while the current is sampled and digitised at ~4000 Hz^1^. The instantaneous current measured in ONT nanopore depends on 5-6 contiguous bases [8]. The measured signal also presents stochastic noise due to several factors, such as homopolymers (same base repeating multiple times) which produce constant current levels, contaminants in the sample, entanglement of long DNA strands, depletion of ions, etc [9]. Additionally, the movement speed of the DNA strand through the pore can vary, causing the signal to warp in the time domain [9]. The raw signal is converted into character representations of DNA bases (e.g. A,C,G,T) using artificial neural networks, generating a typical accuracy >90% for single reads [10]. This conversion process is referred to as *base-calling* and the software tools that perform this conversion are referred to as *base-callers*. Please refer to [8] for a detailed discussion of ONT sequencing.

An example of a raw nanopore signal is shown in Fig. 2a using blue coloured line. Assume that the signal is generated from the DNA sequence *GAATACGAAAATCATTA* which passed through the nanopore. In this example, the instantaneous current of the signal is affected by a string of 6 contiguous bases, known as a *6-mer* (or a *k-mer* in general). Let us assume that the annotation of the signal to the corresponding *k-mers* is known (the process of getting this annotation is detailed in Section 2.2). The *6-mers* in the sequence and the corresponding segments in the raw signal are marked using vertical grey lines in Fig. 2a. When the DNA sequence *GAATACGAAAATCATTA* moves through the pore, the first *6-mer* is *GAATAC*. Similarly, the subsequent *6-mers* are *AATACG*, *ATACGA*, *TACGAA*, …, *TCATTA*. *True annotation* (depicted by dotted green coloured step function in Fig. 2a) corresponds to the *ideal* average level of current for each *k-mer*. These ideal average values are obtained using the *pore-model* provided by ONT, which is elaborated in Section 2.2. The red coloured step function corresponds to an *event*—detailed in Section 2.2.

**Figure 2:**
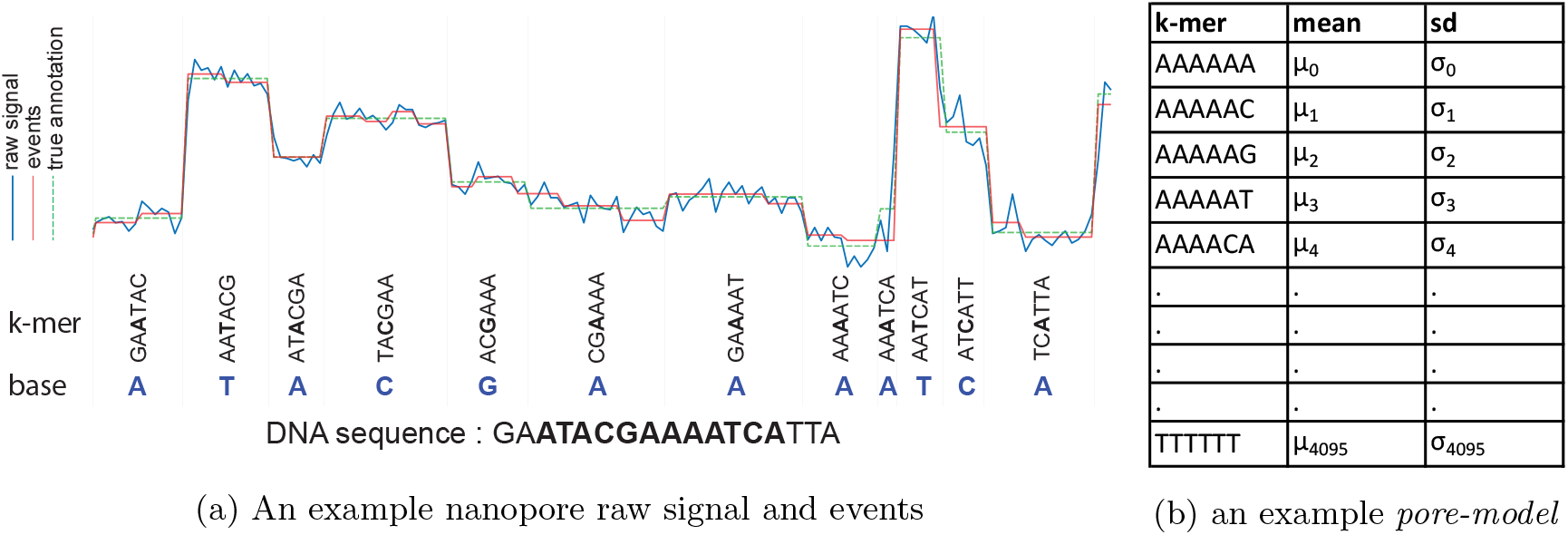
Illustration of a nanopore raw signal, events and *pore-model*

To deduce the sequence from the *k-mers*, the base at the centre (3rd base) of each k-mer is taken, as shown on the bottom of Fig. 2a. For instance, we take *A* from *GAATAC*, *T* from *AATACG*, *C* from *TACGAA* and etc. Hence, we obtain a sequence *ATACGAAAATCA* which is a part of the original sequence *GAATACGAAAATCATTA*. Note that the beginning and the end of the sequence (GA at the beginning and TTA at the end) are clipped.

#### 2.1.4 Nanopore read length distribution

The length of the reads generated from nanopore sequencers can vary from several hundred bases to even more than 2 million bases. A typical sequencing run of a particular sample (which completes after 48-64 hours) generates millions of such reads. The distribution of the read lengths varies in function of DNA integrity, extraction protocols, and sample preparation methods. Example distributions for three different samples are shown in Fig. 3, where both x and y axes are in logarithmic scale. The average read length of a sample typically falls between 8-20 Kilobases.

**Figure 3:**
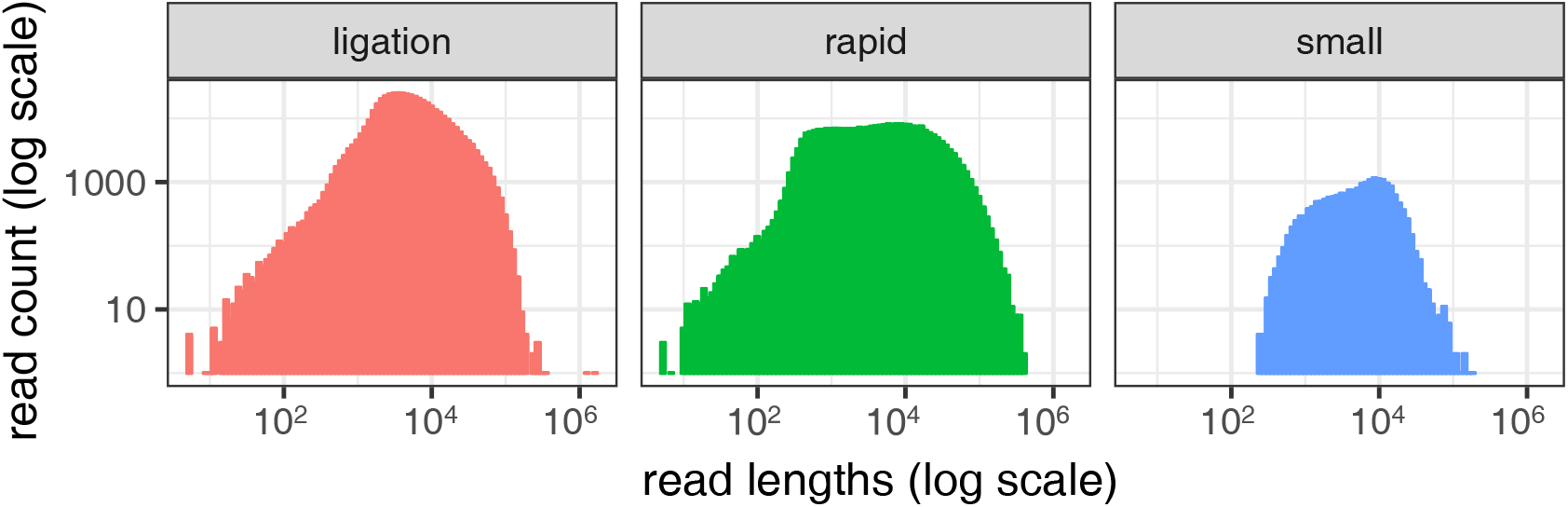
example nanopore read length distributions

#### 2.1.5 Sequence alignment/mapping in the base-space

Once a nanopore read is base-called, the sequence is aligned to a reference sequence (see Fig. 1). A reference sequence consists of a previously generated consensus sequence (such as the human genome reference). Sequence alignment involves global optimisation algorithms to identify the most similar target and to compare any differences between sequences. Compared to biologically occurring variation in individual genomes (<1% difference to the reference), the error-rate of nanopore sequencing is relatively high (5-10%). Thus, sequence alignments derived from nanopore reads are distinct in nature from previous sequencing technologies (such as highly accurate short reads). Consequently, unique analytic tools must be considered when aligning such reads. Alignment tools such as Minimap2 [11] that employ a hash table based genome index followed by a base-level dynamic programming alignment step can successfully align long and noisy reads.

#### 2.1.6 Polishing/Downstream processing using raw signal

The base-space alignment discussed previously in Section 2.1.5 is followed by ‘polishing’, a down-stream processing step which utilises both the base-space alignment results and the raw signal (see Fig. 1). The polishing step reuses the raw signal to recover the lost biological information during base-calling. This polishing step can be to correct errors during base-calling or to detect modified nucleotide bases (eg: DNA methylation).

Previous research has shown that identification of genetic variants can be improved up to an accuracy of more than 99% by using raw signal data from multiple overlapping reads [12, 13]. Thus, the downstream analysis that reuses raw signal data could correct for base-calling errors. It has also been shown that methylated C bases can be differentiated from non-methylated C bases by the use of signal data, using algorithms such as the one implemented in the software package *Nanopolish* [1]. Thus, the downstream analysis that reuses raw signal data could detect modified nucleotide bases.

Signal-space alignment is one of the crucial steps performed in these downstream analyses such as error correction and modified base detection. This signal alignment step is described in the context of modified base detection in the following sections.

### 2.2 Methylation calling

As discussed above, important biological information is lost during base-calling. Some base-calling models may not accommodate methylated data, either because they are trained on unmethylated sequences, or because they abstract away non-canonical bases. Therefore, these molecules may be erroneously classified as unmethylated bases. The process of identifying methylation is known as *methylation calling*.

As implemented in *Nanopolish*, methylation calling requires: 1, raw signals; 2, base-called reads; and 3, base-space alignment to a reference genome (output of the sequence alignment step described above). For a given read, the main steps for methylation calling are: 1, event detection; 2, signal-space alignment; and 3, Hidden Markov Model (HMM) profiling. These steps are performed for each individual read in the data set.

*Event detection* is the time series segmentation of the raw signal based on sudden signal level changes. Each segment is called an *event* and is typically denoted using the mean 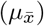, standard deviation 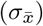 and the duration of the raw signal samples 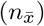 pertaining to the particular segment. The red step function in Fig. 2a denotes such detected events by plotting the mean value of the samples 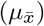 corresponding to the segment. Note that in Fig. 2a, events (red line) roughly match to the true annotation (dotted green line), nevertheless, are not exactly the same. Mostly, the signal has been over-segmented (eg: portion corresponding to k-mer *CGAAAA* has been segmented into 3 events) and seldom under-segmented (eg: k-mer *AAATCA*).

To obtain the true annotation in Fig. 2a, the events detected in the event detection step are aligned to a generic k-mer model signal. This generic k-mer model signal is derived from the base-called sequence and a *pore-model* provided by ONT. The *pore-model* corresponds to a table of all possible k-mers matched to their mean signal value and standard deviation (4^6^ k-mers if k is 6, as shown in Fig. 2b)^2^. For each 6-mer in the base-called read, the corresponding entry in the *pore model* (*mean,sd*) is obtained and these *mean,sd* pairs form the generic k-mer model signal. *Nanopolish* aligns the events from the event detection step to this generic k-mer model signal by using the algorithm named *Adaptive Banded Event Alignment (ABEA)* explained in Section 2.3.

ABEA above produces the alignment between the events and the k-mers in the base-called read. The base-space sequence alignment then is used to deduce which event corresponds to a given k-mer in the reference genome. Finally, this alignment between the events and the k-mers in the reference genome are subjected to Hidden Markov Model (HMM) profiling to identify if a given base is methylated or not.

### 2.3 Adaptive Banded Event Alignment (ABEA)

Algorithms to determine the optimal alignment between two biological sequences typically utilise dynamic programming (DP). Very first of such algorithms, Needleman–Wunsch (NW) algorithm dates back to the 1970s. NW and its variant, Smith–Waterman (SW) algorithm are of quadratic time and space complexity. Both NW and SW were used extensively to perform fine alignment of DNA sequences with high quality. However, due to its extended time consumption, several heuristic improvements were made to improve the speed of alignment without losing quality.

Fig. 4a exemplifies an original SW based alignment (no heuristic) between two sequences, *target sequence t*_*0*_*t*_*1*_*t*_*2*_*t*_*3*_*t*_*4*_*t*_*5*_ *(6 bases long), and query sequence q*_*0*_*q*_*1*_*q*_*2*_*q*_*3*_*q*_*4*_*q*_*5*_*q*_*6*_*q*_*7*_ (8 bases long). The DP table (scoring matrix) contains 6×8 cells as shown. First, the initial values are set (shown as 0 in the figure); second, the score for each cell (s_x,y_) is computed based on a scoring scheme; and third, the trace-back (backtracking denoted by red arrows on the figure) starting from the highest scoring cell and ending at a cell with 0 score, outputs the optimal alignment that yields the highest score (please refer [14] for a detailed explanation of SW).

**Figure 4:**
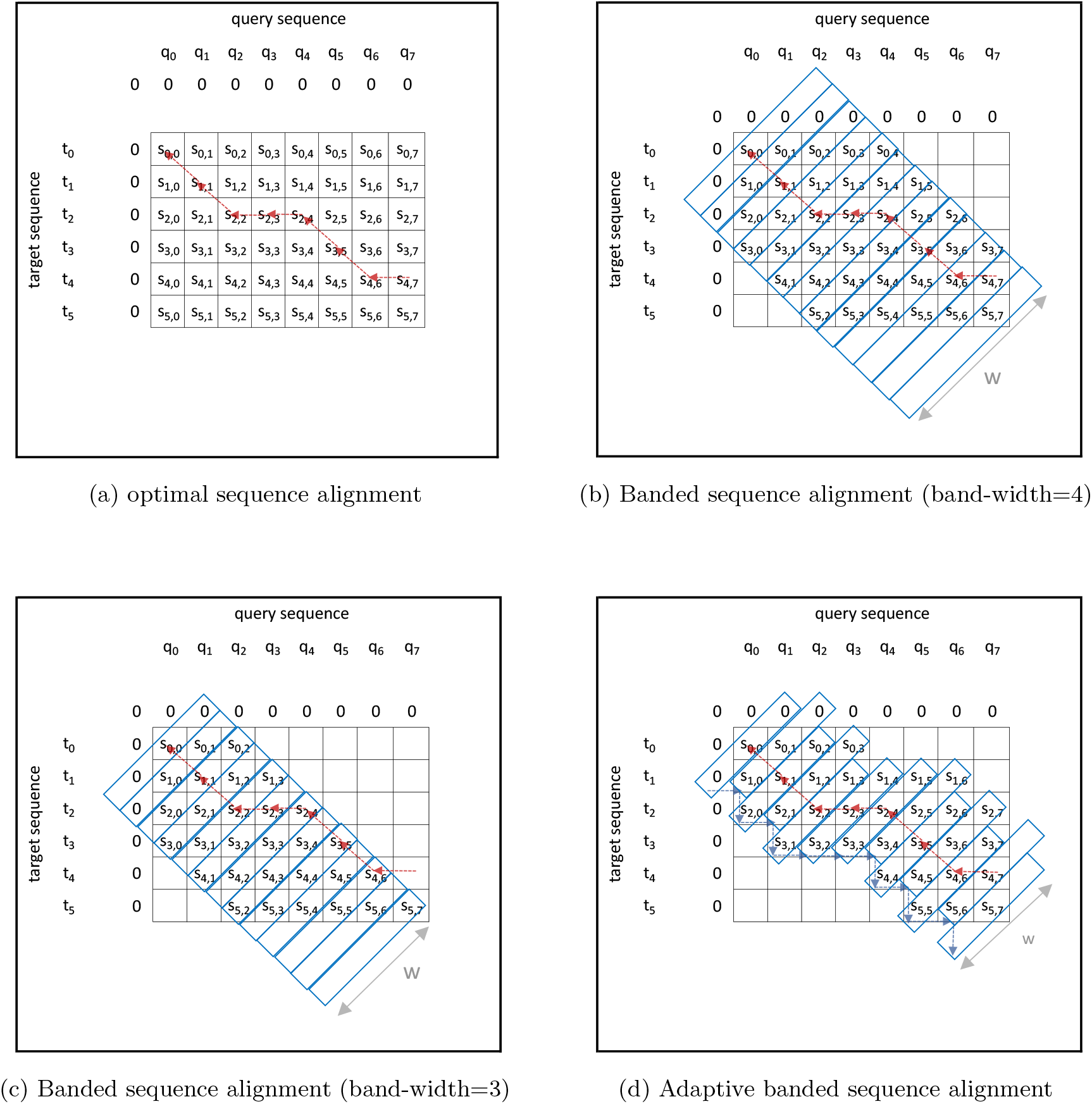
Evolution of dynamic programming based sequence alignment

In the case of short read alignment, the sequences to be aligned are small (typically 100-500 bases). Two sequences (each sequence ~100 bases long) can be aligned by filling ~10^4^ cells. While a single such alignment can be quickly handled by a modern computer, it is very computationally demanding when the number of alignments to be performed scales up to hundreds of millions and billions, which is the case for short reads. To reduce the number of computations, banded alignment approaches were introduced, where only the cells in the DP table along the left diagonal band are computed as shown in Fig. 4b. The underlying assumption is that, the sequences that are aligned to each other are essentially similar, thus the alignment (the trace-back arrows) should lie close to the left diagonal. Note that in the figure, only the cells in a band of width (W) four have been computed, yet has been sufficient to contain the alignment inside the band.

In contrast to short reads, the long reads which emanate from Nanopore, PacBio etc, have lengths which are 100 to 1000 orders of magnitude bigger than short reads, are noisier (with greater number of errors) and are typically not suitable for such small static bands. The 10% base-calling error rate would result in the alignment significantly deviating from the diagonal. A major advantage of long reads is the detection of long indels (insertions and deletions occasionally spanning lengths longer than short reads themselves). When aligning such reads, the alignment path deviates significantly from the diagonal. The high errors and the large indels require the bands to be of large width if they are to be static.

High band-width requirement causes processing times to be extremely high when aligning millions of reads. To improve the speed of this processing, Suzuki-Kasahara (SK) heuristic algorithm [15] was introduced in 2017. SK utilises an adaptive band scheme, letting a smaller band to contain such an alignment within the band, which is exemplified as below.

Consider the same example in Fig. 4b (performed previously with a static band of size 4) is now performed only with a band-width of size 3, as shown the Fig. 4c. Observe that the band is no longer sufficient to contain the whole alignment, i.e. the cell s_4,7_ which previously contained the maximum score is no longer computed, thus the trace-back would begin from the maximum value within the band, which leads to a non-optimal alignment. This is remedied using an *adaptive band* in Fig. 4d. The band moves either down or to the right (the band dynamically adapts) as determined by the Suzuki-Kasahara heuristic, which is illustrated by blue arrows. Observe how the alignment is possible to be contained inside a band of width 3 which was previously infeasible using a static band.

Modified versions of the SK algorithm are used for event-space alignment as exemplified in *Nanopolish* and is referred to as *Adaptive Banded Event Alignment (ABEA)*. In ABEA, the events are aligned to the k-mers of the base-called read (as stated in Section 2.2). As typically there are many more events than k-mers (usually by a factor 1.5-2) due to the frequent over-segmentation of events (discussed in Section 2.2), event alignment is even more difficult than base-space long read alignment if performed with static banding around the diagonal. Thus, an adaptive band is essential for event alignment.

The scoring function for signal alignment uses a 32 bit floating point data type, as opposed to 8-bit integer data type in sequence alignment. Furthermore, the signal alignment scoring function that computes the log-likelihood (which we elaborate shortly) is computationally expensive.

A simplified example of ABEA is shown in Fig. 5a. In Fig. 5a the horizontal axis represents the events (results of the event detection step) and the vertical axis represents the *ref* k-mers (k-mers of the base-called read). The dynamic programming table (DP table) in Fig. 5a is for 13 events, indexed from e_0_-e_12_ vertically, and the *ref* k-mers, indexed from k_0_-k_5_ horizontally. As mentioned previously for computational and memory efficiency, only the diagonal bands (marked using blue rectangles) with a band-width of *W* (typically *W* =100 for nanopore signals) are computed. The bands are computed along the diagonal from top-left (*b0*) to bottom-right (*b17*). Each cell score is computed in function of five factors: scores from the three neighbouring cells (up, left and diagonal); the corresponding *ref* k-mer; and, the event (shown for the cell e_6_, k_3_ via red arrows in Fig. 5b, details of the computation is explained later). Observe that all the cells in the *n*^th^ band can be computed in parallel as long as the *n* − 1^th^ and *n* − 2^th^ bands are computed beforehand. To contain the optimal alignment, the band adapts by moving down or to the right as shown using blue arrows in Fig. 5a. The adaptive band movement is determined by the Suzuki-Kasahara heuristic rule [15].

**Figure 5:**
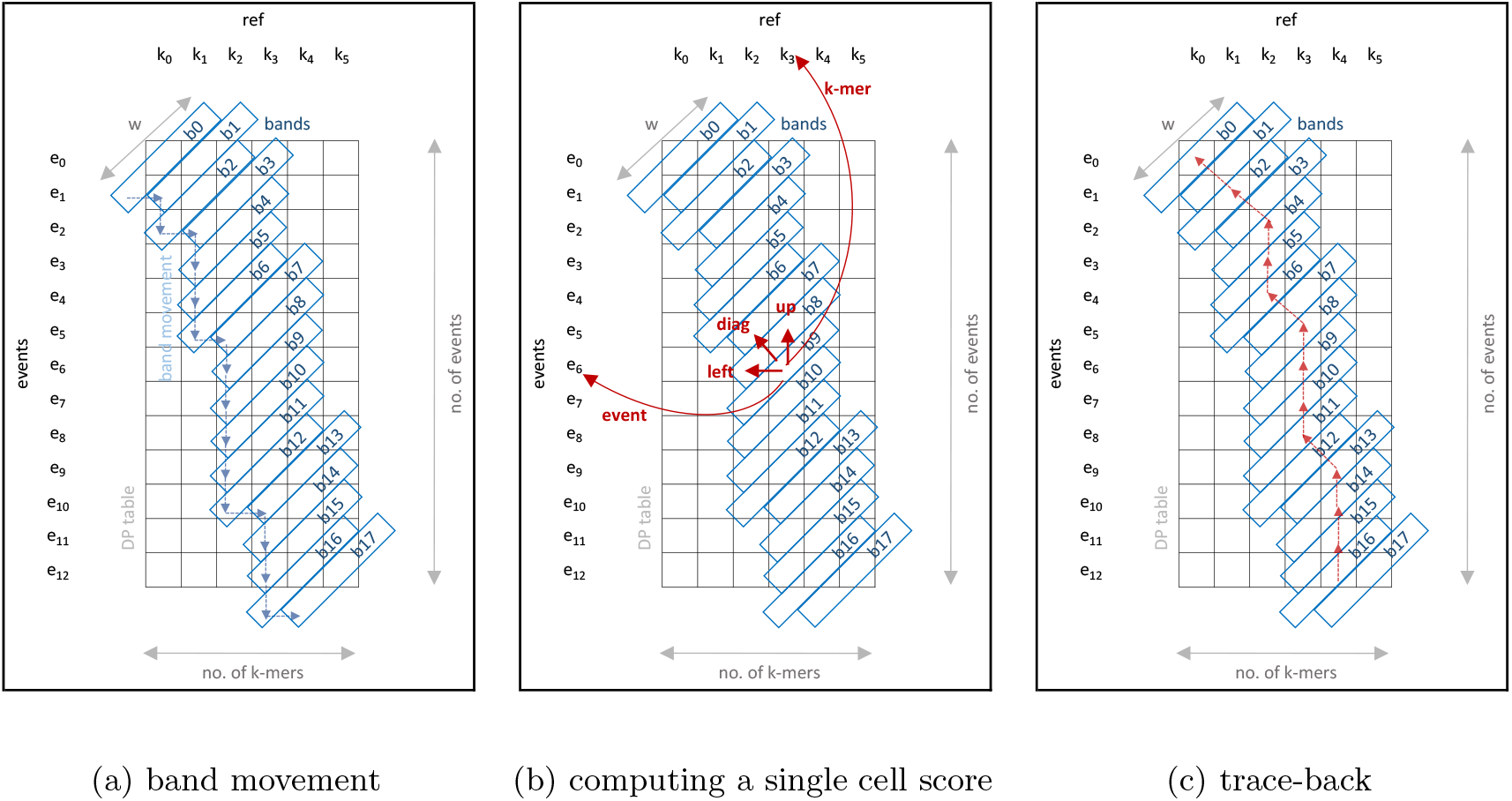
Adaptive Banded Event Alignment

Algorithm 1 summarises the ABEA algorithm used in *Nanopolish* [1] and is explained with the aid of the example in Fig. 5a.

The input to the Algorithm 1 are: 1, *ref* (the sequenced read in base-space—eg: *GAAT-ACG…*); 2, *events* (the output of the event detection step mentioned in Section 2.2); and 3, *model* (*pore-model*—Fig. 2b). As mentioned in Section 2.2, the ABEA algorithm (Algorithm 1) attempts to align the events to the generic signal model (produced with the use of *ref* and the *model*) and outputs the alignment as *event-ref* pairs. The algorithm requires three intermediate arrays, namely *score* (2D floating point array), *trace* (2D byte array) and *ll* (1D pointer array) to formulate the intermediate state during alignment computation, which is the DP table shown in Fig. 5a). Note that, *ll* stands for lower-left, which holds the coordinate of the start point of the band.

The initialisation of the first two bands (*b0* and *b1*) in Fig. 5a is performed by line 20 of Algorithm 1. Then, the outer loop (starting from line 3) iterates through rest of the bands from top-left to bottom-right of the DP table. The inner loop (lines 11-15) iterates through each cell in the current band *bi*. To ensure that only cells within the DP table are computed, the loop counter *j* iterates from *min_j* to *max_j*, instead of 0 to *W* − 1. Lines 4-9 of Algorithm 1 correspond to the movement of the band (corresponds to the blue arrows in Fig. 5a). Band movement is actuated by proper placement of the band in the static 2D arrays, *score* and *trace* via the array *ll* using the functions *move_band_right* and *move_band_down*.

Line 12 of the algorithm performs the cell score computation (explained in detail later) and generates a score and a direction flag for subsequent backtracking, which are henceforth stored in the arrays *score* and *trace*. When all the cells in the DP table are computed, the final operation is to find the actual alignment (*event-ref* pairs) through the backtracking operation (line 17 of Algorithm 1 and red trace-back arrows in Fig. 5c), which uses both the cell scores and the direction flags stored in *trace*.

The *compute* function (called at line 12 of Algorithm 5a) is elaborated in Algorithm 2. A number of heuristically determined constants suitable for Nanopore data, which are used during subsequent calculations are listed at the beginning of this algorithm. The first step of this algorithm is the computation of *lp_emission*, a log probability value (likelihood of the particular signal event being the particular *ref* k-mer), performed using the function elaborated in Algorithm 3. This computed *lp_emission* is used in lines 4-5 of Algorithm 2 along with the heuristically determined constants (*lp_skip,lp_stay,lp_step*) to compute three scores from the diagonal, left and up (*score_d, score_u, score_l*). The maximum of the three scores and direction from which the max score came (flags pertaining to diagonal, up or left) are returned as outputs from this function. The line 3 of Algorithm 2 refers to accessing the scores of the upward, left and diagonal cells which was previously mentioned with respect to cell e_6_,k_3_ and the red arrows in Fig. 5b.

**Algorithm 1.**
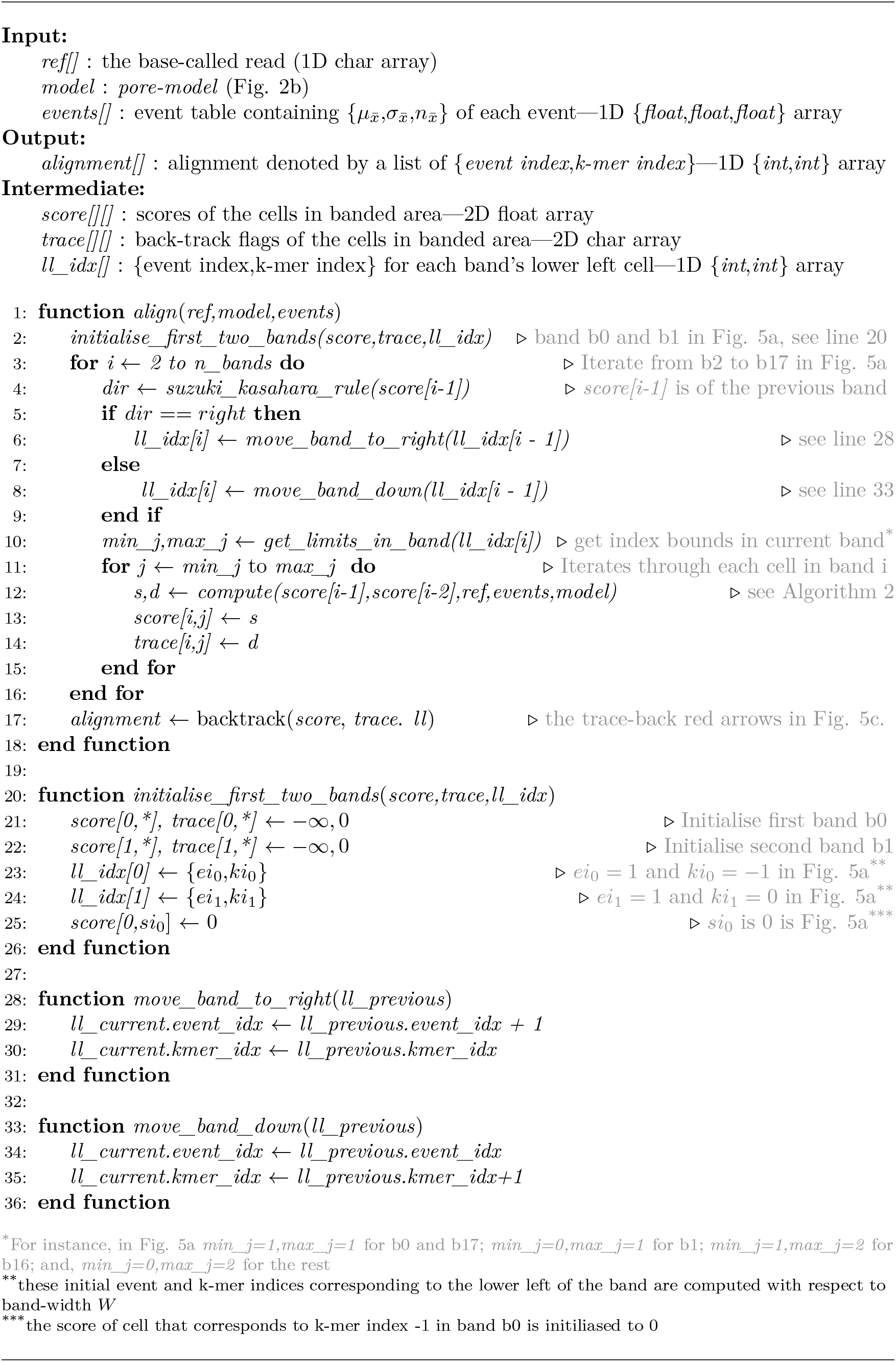
Adaptive Banded Event Alignment.

The log probability computation in Algorithm 3 involves floating point log probability computations. For the k-mer at the specific *ref* position, the *pore-model* table (Fig. 2b) is accessed to obtain the corresponding model values. This *model_kmer* (mean and the standard deviation of the particular model k-mer) and the mean value of the event is used for the log probability computation as shown in the Algorithm 3. Note that for event alignment neither the standard deviation or the duration of the event are used.

**Algorithm 2.**
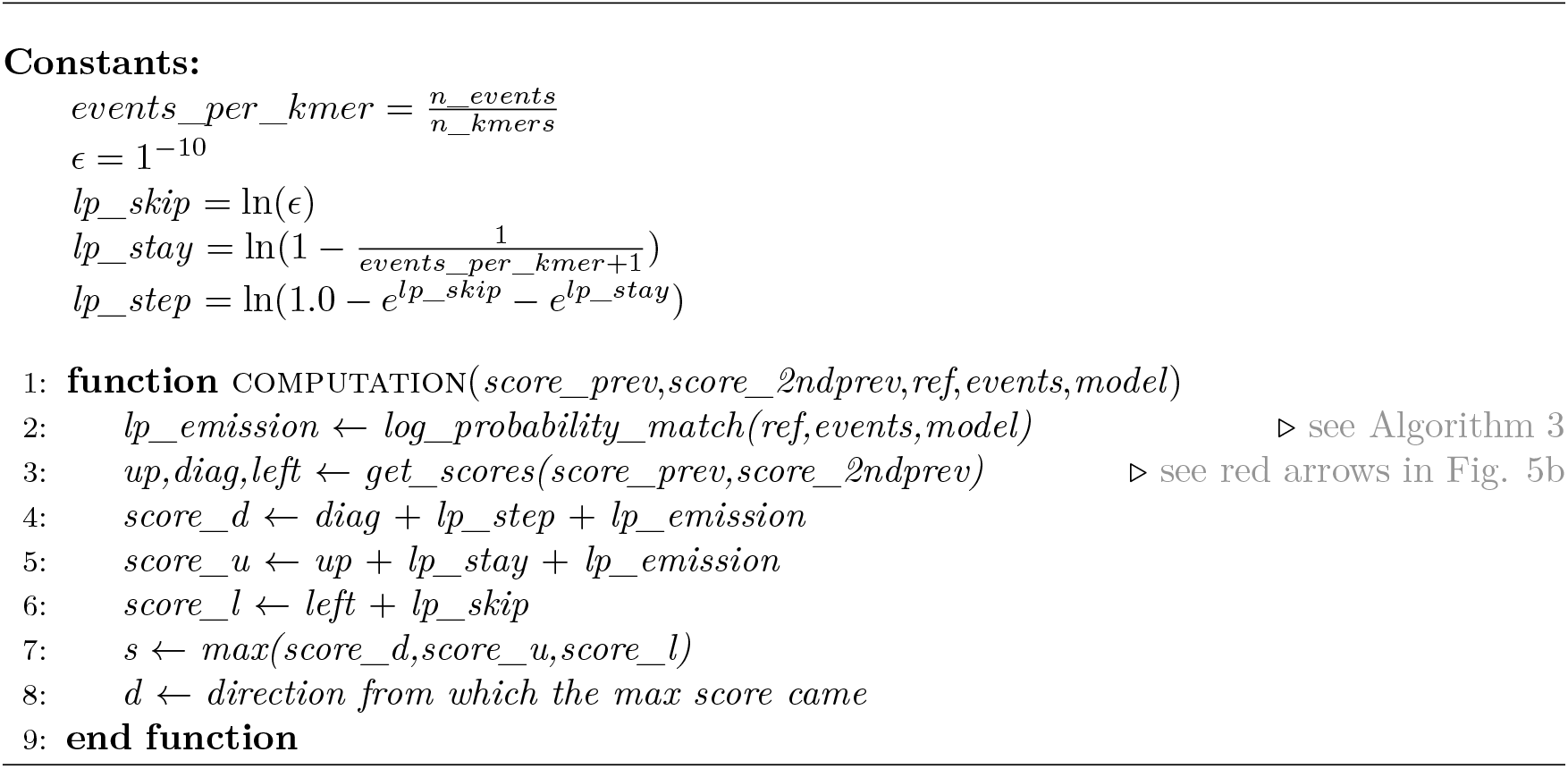
Adaptive Banded Event Alignment – cell score computation.

The above elaboration covers the ABEA algorithm to a sufficient enough level to explain our GPU implementation and optimisations. Therefore, implementation details of checking out of bound array accesses and the backtracking process were not discussed. Furthermore, the concept of the ‘trim state’ and ‘event scaling’ were not discussed as the control flow of the algorithm are not affected by them. Thus, those details not vital for the elaboration GPU implementation. However, for the sake of completeness, a brief account of this ‘trim state’ and ‘read-model scaling’ are given below.

**Algorithm 3.**
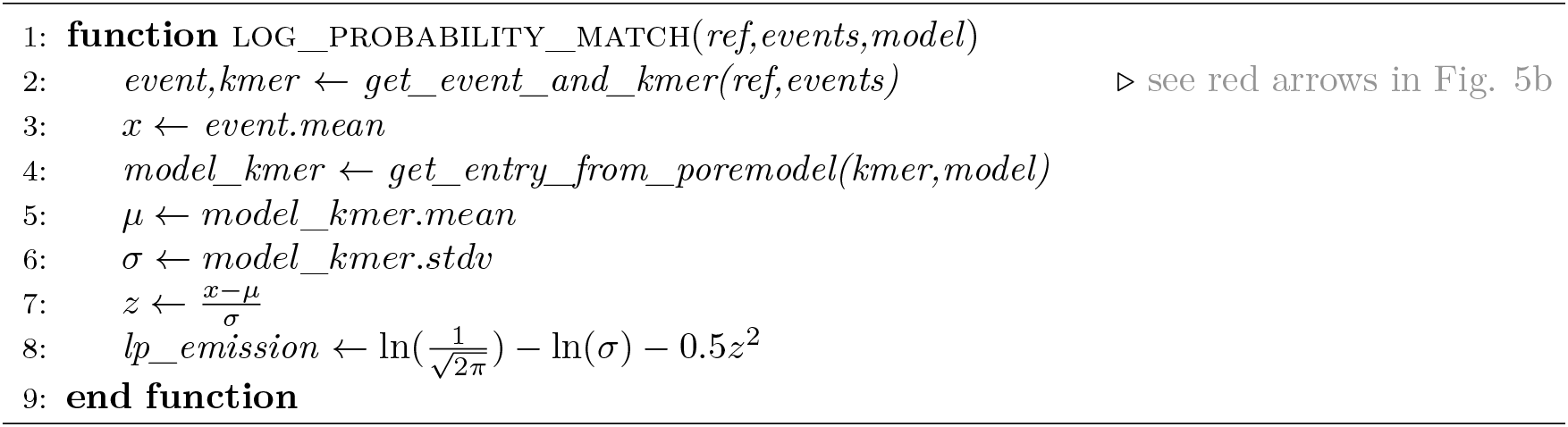
Adaptive Banded Event Alignment – log probability computation.

The raw signal may contain samples at the beginning/end that may be ignored by the base-caller and hence does not contribute to the base-called sequence. These samples may be open pore signal immediately before or after the DNA molecule is detected (i.e. the electric current when nothing is in the nanopore), or perhaps part of the adaptor (molecules bounds to the ends of the DNA molecules to enable sequencing). The ‘trim states’ allow the alignment to ignore these samples, since such samples should not be considered to be part of the base-called read.

Due to reasons such as slight variations between different nanopores and characteristic changes of the same nanopore with time, an event will not directly match the *pore-model* in Fig. 2b [16]. Therefore, to account for these variations either the events or the *pore-model* should be scaled on a per-read basis. In *Nanopolish*, two scaling parameters namely *shift* and *scale* are estimated on a per-read basis, prior to ABEA algorithm, using a ‘Method of Moments’ approach [16]. Then, during ABEA, the *pore-model* mean values are scaled using these two parameters. The scaling should be performed at line 5 of Algorithm 3 as *μ* ← *model*_*kmer.mean* × *scale* + *shif t* instead of directly assigning *model*_*kmer.mean* to *μ*.

### 2.4 GPU architecture and programming

Graphics Processing Units (GPUs) were originally designed as co-processors for graphics processing and rendering. Graphics processing and rendering algorithms involve pixel-wise operations which expose fine-grained parallelism, thus GPUs consists of hundreds of compute cores to perform parallel processing. Eventually, the concept of general purpose graphics processing units (GPGPU) emerged where the GPUs were exploited to accelerate compute intensive, yet highly parallelism portions of general purpose algorithms. GPUs are quite popular in scientific computations due to the significant speedup when used for common matrix manipulation which contains fine-grained parallelism. From around a decade ago, GPUs which are explicitly designed for high performance computers are available (e.g., Tesla GPUs from NVIDIA).

GPUs are of Single Instruction Multiple Data (SIMD) architecture (or more accurately Single Instruction Multiple thread, as stated by NVIDIA), where multiple threads run the same stream of instructions in parallel yet on different data. Conversely, CPUs are of Multiple Instruction Multiple Data (MIMD) architecture, where each thread runs its own instruction sequence and own data stream, independent of the others. GPUs have hundreds or even thousands of processing cores while a CPU would maximally have a few dozen cores. However, the GPU cores are relatively less complex (fewer instructions, smaller caches, no sophisticated branch prediction units etc.) and run at a lower clock speed when compared to a CPU. Due to these significant differences between CPU and GPU architectures, serial algorithms designed and developed for the CPUs are not suitable for execution on GPUs. Such algorithms have to be adapted and parallelised in a way that the GPU architectural features are efficiently used.

NVIDIA provides a programming model/framework for programming their GPUs for general purpose computations, called Compute Unified Device Architecture (CUDA). CUDA includes CUDA C/C++ (extended C/C++ syntax) and an Application Programming Interface (API) to provide a platform to write programs for the NVIDIA GPU. We used this CUDA C/C++ for our GPU implementation of the Adaptive Banded Event Alignment algorithm.

We will now briefly give GPU/CUDA related terms. Readers are advised to refer to [17] and [18] for further information.

A GPU *kernel* is a function that is executed on a GPU. A GPU kernel is written from the execution perspective of a single GPU thread. These GPU kernels will run in parallel, based on the parameters specified with the function call, known as the *thread configuration*. This thread configuration in CUDA is an abstraction which employs a hierarchy of threads. In the thread hierarchy, a group of threads are known as a *block*. A group of blocks form a *grid*. Instances of a single kernel are executed in a single grid. Blocks and grids can be 1 dimensional, 2 dimensional or 3 dimensional. The presence of this thread hierarchy lets the programmer organise and map the threads conveniently to a grid. These logical threads would be mapped to the hardware cores automatically by the underlying driver software and hardware.

A thread block consists of one or more *thread warps*. A warp is a group of threads sharing the same program counter. A data dependent conditional branch inside a warp causes the threads to execute each code path while disabling threads that are not in the path, known as *warp divergence*. The warp divergence affects the performance and should be minimised.

The *occupancy* is the percentage of the number of active warps to the maximally supported warps on the GPU. A lesser occupancy leads to under utilisation of GPU resources. Thus, a higher occupancy is preferable for better utilisation of GPU resources.

GPUs also employ a memory hierarchy. Relatively larger but slow Dynamic Random Access Memory (DRAM) that forms the lowest level in the memory hierarchy is known as *global memory*. Global memory is typically allocated using *cudaMalloc()* API function. Memory allocated in this global memory can be exclusively accessed by all the threads in the grid. The next level in the memory hierarchy which is made of relatively fast, yet smaller SRAM is called *shared memory*. Shared memory is allocated on a per-thread-block basis and is shared by all the threads in the block. Shared memory can be called user managed cache (more accurately a programmer managed cache) as the programmer is expected to identify and load frequently accessed data to the shared memory. In addition, there are one or more levels of SRAM caches managed by the hardware. The registers are the fastest and highest in the hierarchy and are allocated by the compiler on a per-thread basis.

The global memory can be easily saturated when hundreds of threads compete to access the memory at the same time. Thus, memory accesses should be batched such that contiguous threads access contiguous memory locations. This process is referred to as *memory coalescing* and reduces global memory requests thus reducing the impact on performance compared to scattered memory accesses. Additionally, the programmer could utilise the shared memory to load and store frequently accessed data, which also reduces global memory traffic.

## 3 Related work

An algorithm to call methylation using the raw signal from ONT sequencers was introduced by Simpson et al. [1]. The associated C++ based implementation of this algorithm is a sub-module under the open source tool *Nanopolish*. *Nanopolish* was designed to run on high-performance computers and is not lightweight or suitable for deployment on embedded systems.

The signal-space alignment algorithm, termed *Adaptive Banded Event Alignment (ABEA)*, used in *Nanopolish* is a customised version of the Suzuki-Kasahara alignment algorithm [15] for base-level sequence alignment. According to the best of our knowledge, neither of these algorithms (ABEA or Suzuki-Kasahara) have GPU accelerated versions. The root origins of these algorithms are dynamic programming sequence alignment algorithms, such as Smith-Waterman and Needleman-Wunsch. A number of GPU accelerated versions for Smith-Waterman exist in previous research [19, 20, 21] [20] [21]. However, the Smith-Waterman algorithm has a compute complexity of *O*(*n*^2^) and is most practical when the sequences are short, especially when millions of sequences need to be aligned. As nanopore sequencers can produce reads >1 million bases long, computing the full DP table for such reads using SW would require >10^12^ computations and hundreds of gigabytes of RAM—and even more if aligning raw nanopore signals.

**Algorithm 4.**
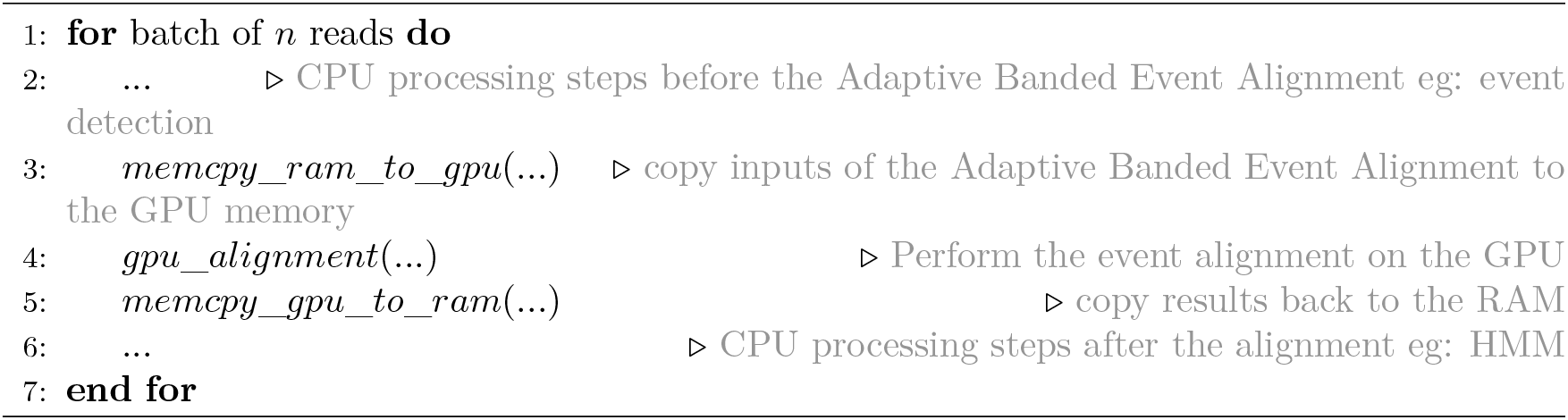
Outline of execution flow.

Heuristic approaches such as banded Smith-Waterman attempt to reduce the search space by limiting computation along the diagonal of the DP table. While the approach is suitable for Illumina short reads, it is less so for noisy long nanopore reads as substantial band-width is required to contain the alignment within the band. The Suzuki-Kasahara algorithm uses a heuristic that allows the band to adapt and move during the alignment, thus containing the optimal alignment within the band but allowing large gaps in the alignment. Modified versions of the adaptive banded alignment algorithm are used for signal-space alignment, as exemplified in *Nanopolish*. The band-width (width of the band) used for signal-space alignment is typically higher (~100) compared to other banded algorithms used for sequence alignment. In addition, the scoring function for signal alignment uses a 32 bit floating point data type, as opposed to 8-bit integers in sequence alignment. Furthermore, the signal alignment scoring function that computes the log-likelihood is computationally expensive. Taken together, these reasons motivated us to consider using GPUs to speedup the computation of signal-space alignment.

The portable compute module, MinIT, manufactured by ONT is composed of a NVIDIA SoC [22] that exploits GPUs for performing live base-calling, which can perform base-calling at a speed of ~150 Kbases per second, thus keeping up with the MinION sequencer’s output. In addition, our previous work has optimised the popular *Minimap2* [11] sequence alignment tool (which typically requires ~16GB memory) for reduced peak memory usage, enabling the software to be executed on embedded processors [23]. The data processing steps required for methylation calling are thus possible to run on embedded processors, therefore supporting the implementation of a portable, offline DNA methylation detection application that would facilitate such analyses in the field.

Load balancing between the CPU and GPU for heterogeneous processing has been explored for areas such as fluid dynamics [24] and conjugate gradient method [25]. However, nanopore data have different characteristics compared to aforementioned applications which are predominately based on matrices. Furthermore, the signal-space alignment algorithm is different from linear algebra algorithms used in these fields. We exploit characteristics of Nanopore data and algorithms to perform memory, compute and load balancing optimisations.

## 4 Methodology

To optimise the performance on GPUs, we process a batch of reads (original source code processes a read at a time) at a time. Such batch processing minimises data transfer initialisation overhead (between RAM and GPU memory); reduces the GPU kernel invocation overhead; and, allows parallelism which sufficiently occupies all available GPU cores. The execution flow follows the typical GPU programming paradigm, which is elaborated in Algorithm 4. In Algorithm 4, *gpu_alignment(…)* refers to the GPU implementation of the Adaptive Banded Event Alignment (CPU algorithm is elaborated in Algorithm 1). We present our methodology in three steps: parallelisation and compute optimisations in Section 4.1; memory optimisation in Section 4.2; and, the resource optimisation through heterogeneous processing in Section 4.3.

### 4.1 Parallelisation and compute optimisations

The GPU implementation of the Adaptive Banded Event Alignment (ABEA) algorithm is broken into three GPU kernels. Breaking down into the three GPU kernels allows for efficient thread assignment based on the workload type, synchronisation of all GPU threads (a GPU kernel execution is inherently a synchronisation barrier [17]) and minimising warp divergence compared to a big all-in-one GPU kernel.

The three GPU kernels are:

- *pre-kernel* – Initialising the first two bands of the dynamic programming table (corresponds to line 2 of algorithm 1) and pre-computing frequently accessed values by the next GPU kernel;
- *core-kernel* – The filling of dynamic programming table which is the compute intensive portion of the ABEA algorithm (corresponds to line 3-16 of Algorithm 1 composed of nested loop); and,
- *post-kernel* – Performs backtracking (corresponds to line 17 of algorithm 1)

#### 4.1.1 pre-kernel

The *pre-kernel* initialises the first two bands of the dynamic programming table (initialisation performed at line 2 of Algorithm 1 on CPU). The *pre-kernel* also pre-computes the values in a data structure called *kcache*, a newly introduced data structure in the GPU implementation that improves cache hits during the subsequent execution of the *core-kernel*.

A simplified version of the *pre-kernel* is in Algorithm 5 and thread configuration for the invocation of the *pre-kernel* is in Fig. 6. Note that the GPU kernel is presented (as is always the case) from the perspective of a single GPU thread in Fig. 6.

**Figure 6:**
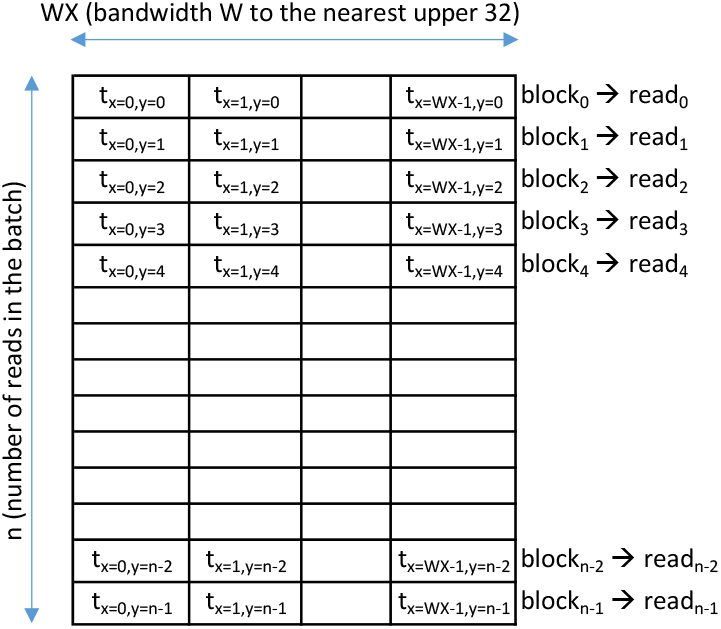
Thread configuration of *pre-kernel*

Each cell in Fig. 6 represents a GPU thread denoted as *t*, where the subscripts *x* and *y* denotes the thread index along the x-axis and the y-axis respectively. The thread grid in Fig. 6 is composed of *n* thread blocks, where *n* is the number of reads in the batch. Each thread block contains *WX* threads where *WX* is the nearest upper ceiling multiple of 32 to the band-width *W* (band-width of the ABEA algorithm); i.e. 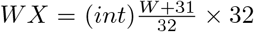 For instance, if *W=100*, *WX* is 128. The reason for taking a multiple of 32 is due to performance attributed by a thread block size that a multiple of the warp size (warp size is 32 currently) [18]. As shown in Fig. 6, a single thread block composed of *WX* threads is assigned to a single read.

In the Algorithm 5, lines 2-3 get the thread index of the thread being executed, i.e. the thread indices denoted as *x* and *y* in Fig. 6. Line 4 obtains the memory pointers of the input array *ref*; intermediate arrays *score* and *trace*; and the *kcache*, the use in which is explained in the memory optimisation Section (Section 4.2).

Lines 5-8 of Algorithm 5 initialises the first two bands of the dynamic programming table (which was performed at line 2 of original CPU Algorithm 1). The kernel is in from the perspective of a single thread and thus a single cell is initialised by a single thread. The collective execution of all the threads in Fig. 6, effectively sets a band for all the reads in the batch in parallel, which is illustrated in Fig. 7. Note that, only the first two reads are elaborated in Fig. 7, and in reality each thread block has a read assigned to it. In Fig. 7, each cell in band_0_ (marked as iteration 1) contains the index of the thread which performs the initialisation at line 6 of Algorithm 5. Similarly, iteration 2 corresponds to line 7 of Algorithm 5.

**Figure 7:**
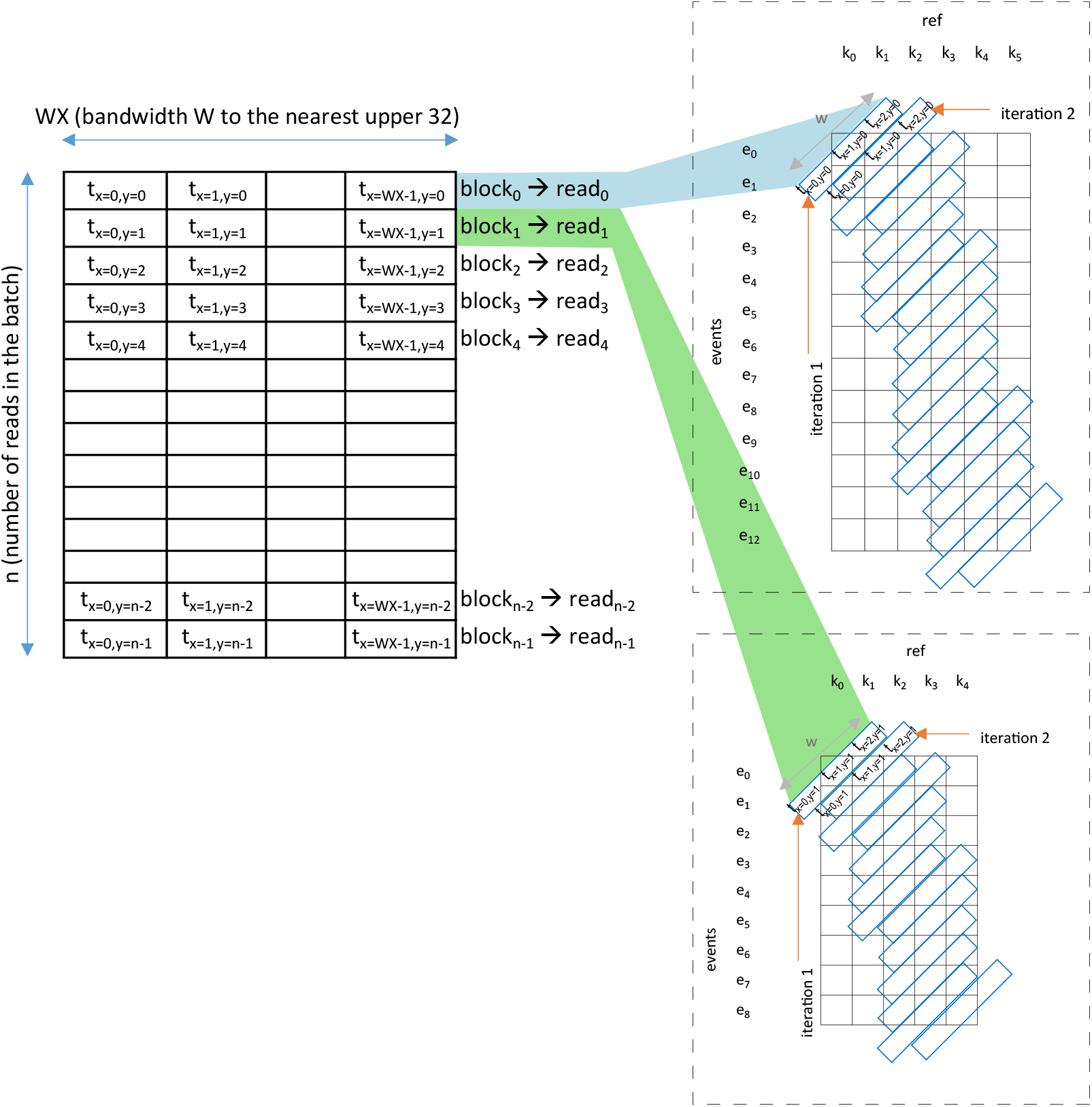
Thread assignment of *pre-kernel*. The assignment for the first two reads are shown. Each thread block has a read assigned to it (block_0_ refers to threads *t*_*X*=0,*y*=0_ to *t*_*x*=*W X* −1,*y*=0_, and read_0_ is processed by all threads in block_0_; similarly, block_1_ refers to *t*_*X*=0,*y*=1_ to *t*_*x*=*W X* −1,*y*=1_, and read_1_ is processed by threads in block_1_).

The *if* condition on line 5 of Algorithm 5 is to limit the threads to the width of the band *W*, a consequence of selecting *WX* which is a multiple of 32 (as stated previously). Note that there is a 1024 thread limit for a block [17] in current NVIDIA CUDA/GPU architecture, thus our implementation will only work for a maximum band-width of 1024. This limit is more than sufficient for a typical *W* of 100 in ABEA.

Line 10-11 of Algorithm 5 initialises the index of the lower left band which corresponds to line 23-24 of Algorithm 1. Note that this initialisation is executed by one thread per read (thread id 0 along y-axis). Lines 13-16 in Algorithm 5 initialises *kcache*. As stated previously *kcache* is a newly introduced array for the GPU implementation to minimise random accesses to the GPU memory during the *core-kernel* and will be explained in Section 4.1.2. Note that, this *kcache* initialisation in line 13-16 is also executed by one thread per read (thread id 0 along y-axis). The loop in 13-16 can be further parallelised; however, as the time spent on *pre-kernel* is comparatively negligible (see results), further parallelising this loop is superfluous.

**Algorithm 5.**
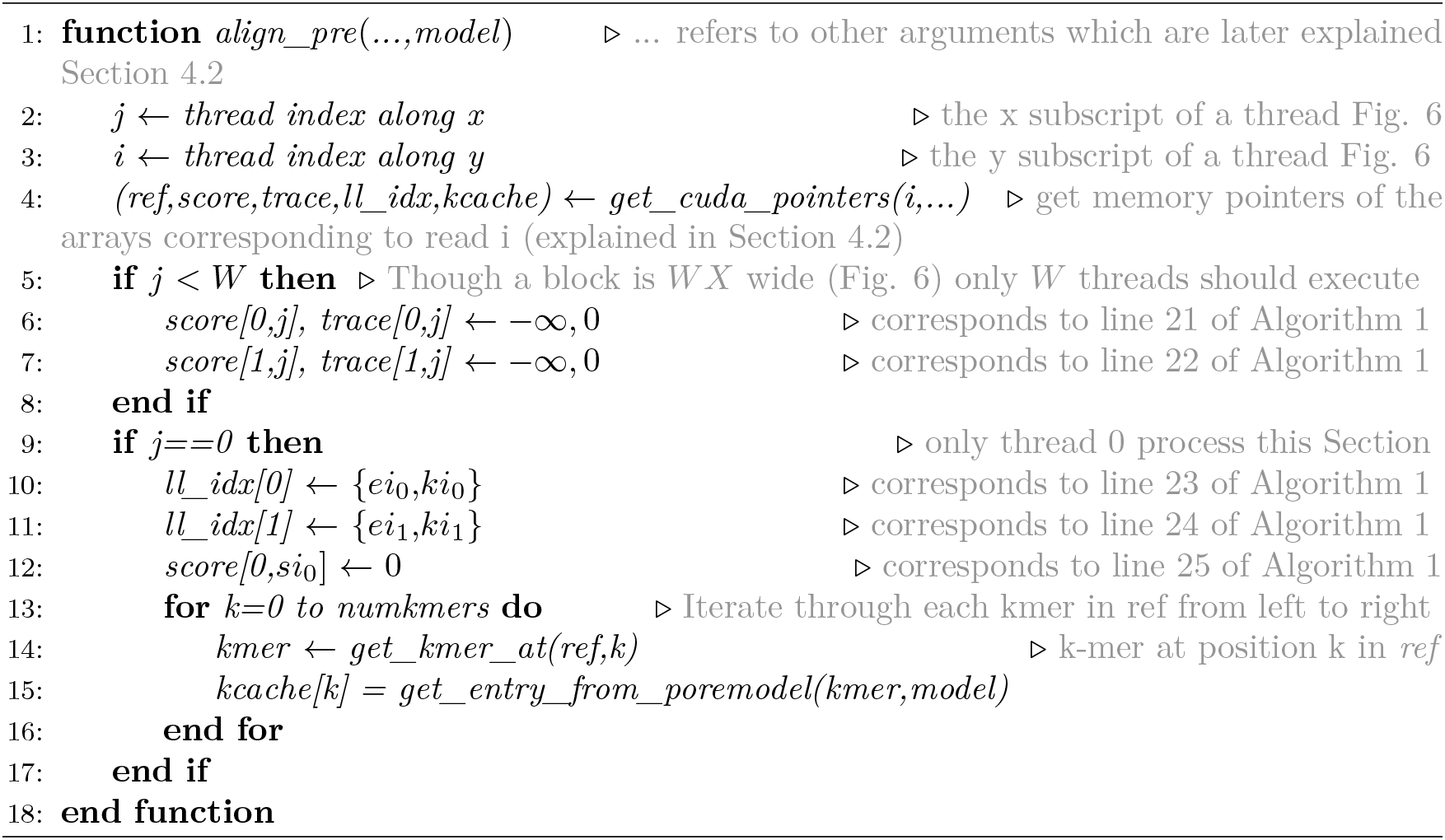
Adaptive Banded Event Alignment – *pre*-*kernel*.

#### 4.1.2 core-kernel

A simplified version of the *core-kernel* which fills the dynamic programming table in Fig. 5a (corresponds to line 3-16 of the original Algorithm 1) is in Algorithm 6. This kernel is executed with the same kernel thread configuration as *pre-kernel* in Fig. 6. Thus, a batch of reads are processed in parallel with a block of threads assigned to a single read in a similar way to that in *pre-kernel* (Fig. 7). The only difference in Fig. 7 for the *core-kernel* is that the third band to the last band are processed instead of the first two bands.

All the *W* cells in a given band (Fig. 5a) are computed by *W* number of GPU threads in parallel (lines 26-30 of Algorithm 6), thus the inner loop of Algorithm 1 (lines 11 and 15) is now no longer present. However, the outer loop of Algorithm 1 cannot be parallelised due to band *n* depending on *n* − 1 and *n* − 2 bands as explained the background. The movement/placement of the band (described in background) is performed by a single thread using the condition given on line 13 Algorithm 6 that limits the code segment to thread 0. In addition, synchronisation barriers per-thread-block basis (_*syncthreads*) in Algorithm 6 prevent any data hazards due to multiple threads assigned to a single read.

Another notable difference in the GPU implementation is the use of GPU shared memory [17] (user-managed cache or more accurately programmer-managed cache) for exploiting the temporal locality in the memory accesses to the dynamic programming table (*n*^*th*^ band in Fig. 5a is computed using bands *n-1* and *n-2*). Shared memory is allocated for three bands (current, previous band and second previous) by line 6-7 of Algorithm 6 which are then initialised at lines 9-10 of Algorithm 6. These initialised memory locations are used during band direction computation (lines 14-21 of Algorithm 6) and the cell score computation (lines 27-28 of Algorithm 6), eliminating any accesses to the slow GPU global memory (shared memory-SRAM vs global memory-DRAM). The cell score is written to the global memory at the end of the iteration (line 32 of of Algorithm 6) as scores are later required for backtracking. Finally, current, previous and second previous bands are set for the next iteration (lines 33-36 of Algorithm 6).

As stated under Section 4.1.1, the data structure *kcache* introduced to the GPU implementation facilitates memory coalescing by minimising random memory accesses to the *model* array (*pore-model* array in Fig. 2b). If *kcache* did not exist, access pattern by contiguous threads in the *core-kernel* (shown for the iteration 5 of read 0) would look like in Fig. 8a where accesses to the *ref* are shown in green colour arrows and the subsequent accesses to the *pore-model* are in red colour arrows. The green arrows (relates to getting the k-mer at line 2 of Algorithm 3 in the CPU version) are spatially local and would facilitate memory coalescing in the GPU. However, red arrows (relates to line 4 of Algorithm 3 in the CPU version) to the *model* array are random accesses. Note that such random accesses would occur during each iteration (iteration 3 to the last band iteration). Such multiple threads accessing random GPU memory locations degrade the performance due to smaller and less powerful GPU caches (compared to CPU), for instance, 32KB *pore model* array is larger than 8KB GPU constant cache [17].

**Figure 8:**
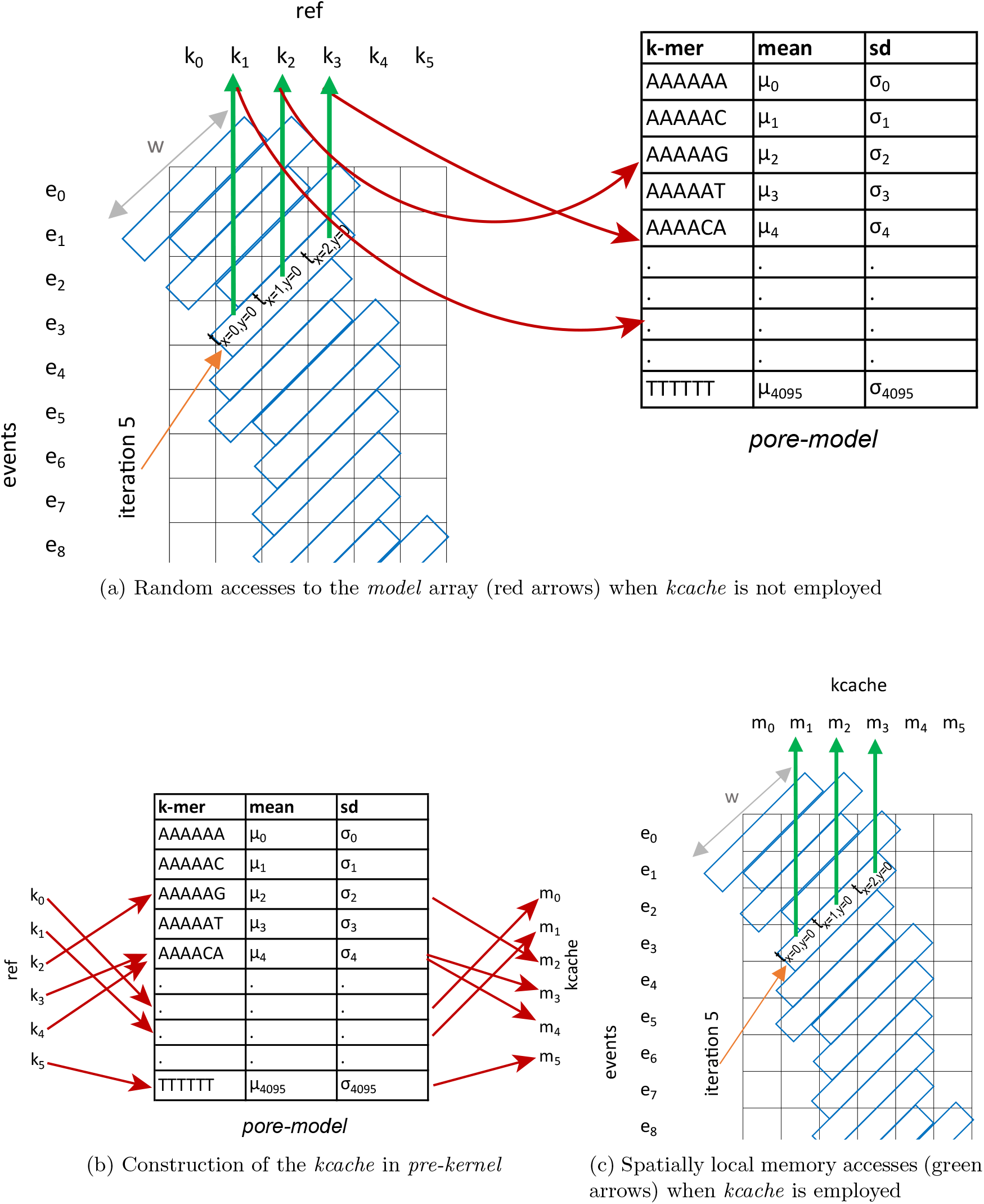
Utility of *kcache* in the *core-kernel* to impreo11ve memory coalescing

These random accesses are eliminated by the *kcache* constructed in *pre-kernel* (stated under Section 4.1.1) which is then passed as an argument to the *compute* function at line 27 in Algorithm 6). This *kcache* is then passed on to the *log_probability_match* function (at line 2 of Algorithm 7) which is then used at line 4 of Algorithm 8. The construction of the caches in the *pre-kernel* requires random accesses to the model as shown in Fig. 8b, which happens only once. However, this *kcache* is utilised by the *core-kernel* in every iteration and facilitates memory coalescing (see green arrows in Fig. 8c which are spatially local accesses to the *kcache* by contiguous threads in iteration 5).

It is noteworthy to mention that allocating one thread block per read is critical (in the kernel configuration) to: use lightweight block synchronisation primitives _*syncthreads* (instead of expensive kernel invocations as synchronisation barriers [17]); minimise warp divergence (otherwise the longest read in the thread block would consume the longest time which corresponds to the band filling loop); and, use shared memory per read (shared memory is allocated per block).

#### 4.1.3 post-kernel

The backtracking operation performed by this *post-kernel* (one thread assigned to one read) does not expose fine grained parallelism as in previous kernels and thus not ideal for the GPU. However, performing this on GPU is still advantageous when compared to transferring huge intermediate arrays (*scores* and *trace*—size in order of GB) from GPU to the RAM. In addition, no additional memory in the RAM is required, thus reducing peak RAM usage.

Allocating one thread block per read (as in *core-kernel* to reduce warp divergence) is not ideal for this *post-kernel* due to the lack of fine grained parallelism (i.e. 1 block having 1 thread), which results in reduced GPU occupancy (occupancy will be limited by the maximum thread blocks that can simultaneously reside in a GPU multi-processor). This is remedied without affecting the warp divergence by allocating a large number of threads per block (eg: 1024) and then limiting only the first thread in the warp (a warp is composed of 32 contiguous threads [17] and thus thread with indices 0, 32, 64, 96 … etc) to perform the actual computation (backtracking for a read).

**Algorithm 6.**
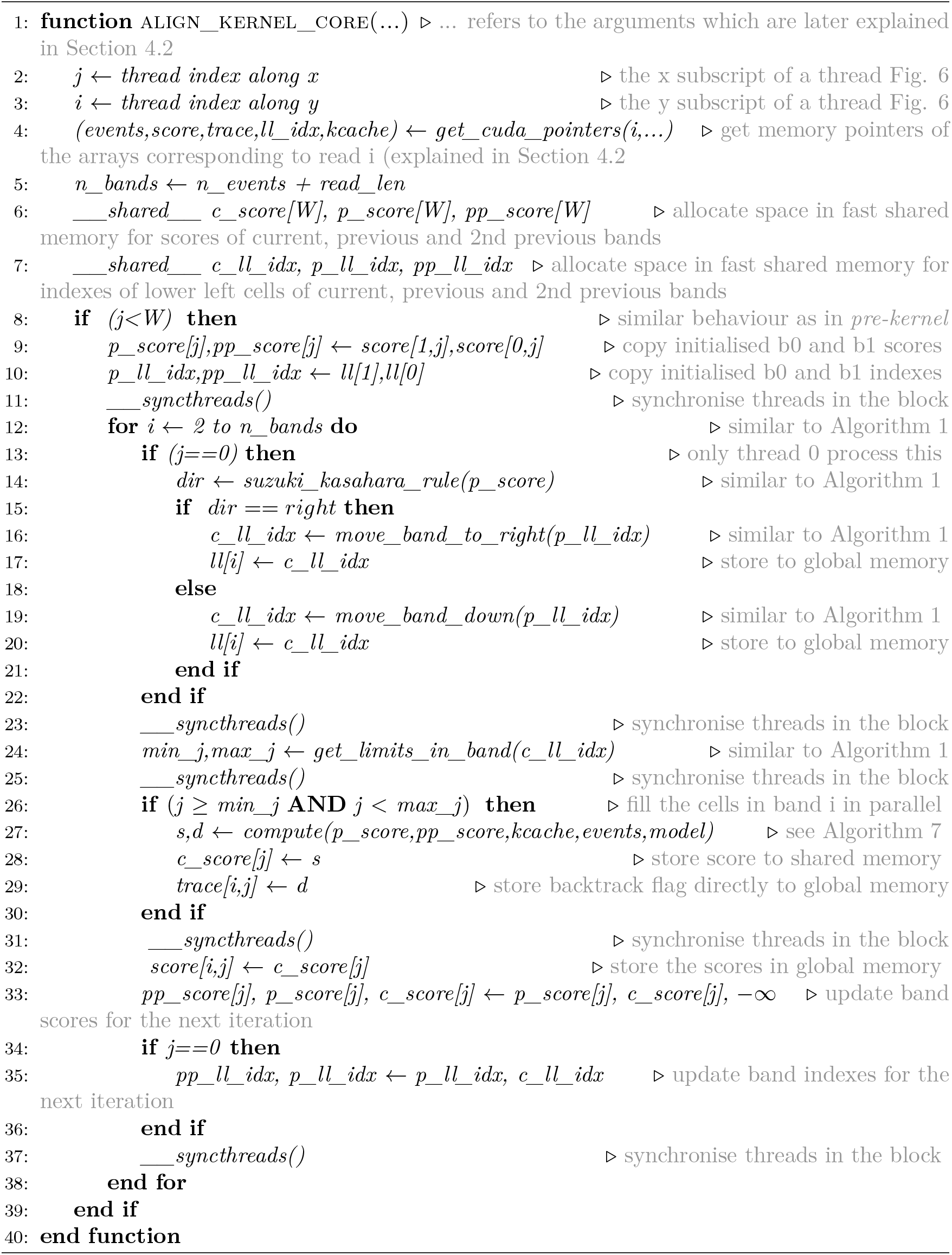
Adaptive Banded Event Alignment – *core*-*kernel*.

**Algorithm 7.**
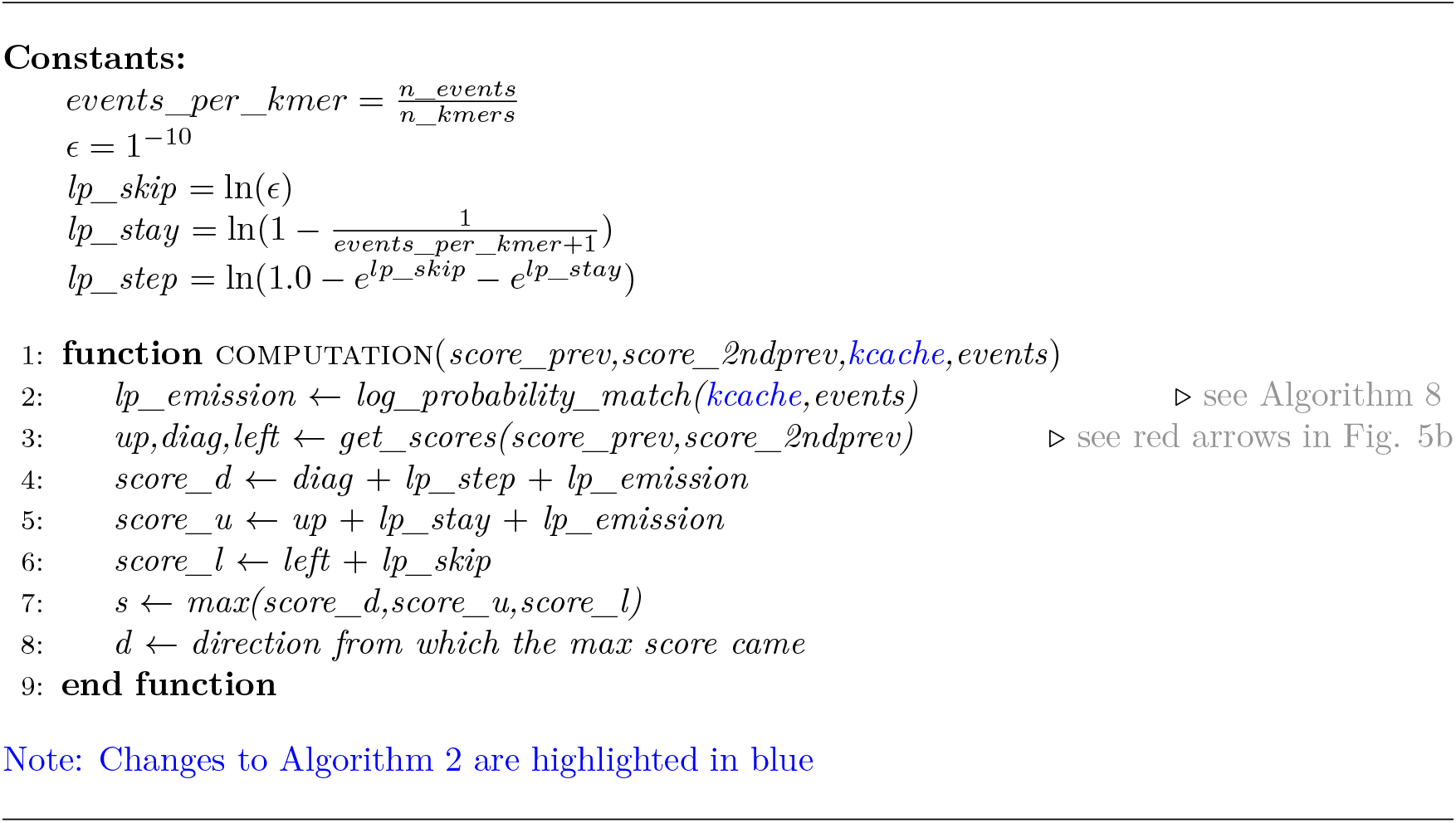
Adaptive Banded Event Alignment – *core*-*kernel* – cell score computation.

**Algorithm 8.**
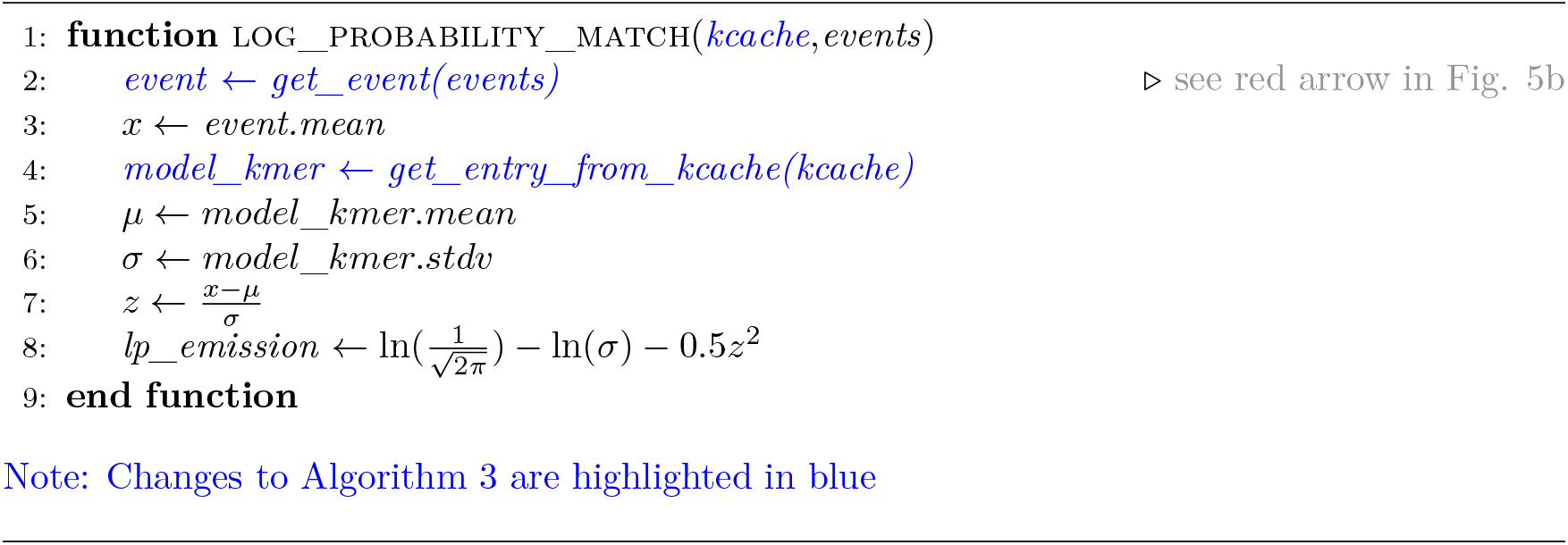
Adaptive Banded Event Alignment – *core*-*kernel* – log probability computation.

### 4.2 Memory optimisation

CPU version of the Adaptive Banded Event Alignment (ABEA) algorithm performs dynamic memory allocations (*malloc*) on a per read basis. The number of reads in a dataset is in the order of millions and thus incur millions of *malloc* calls. However, dynamic memory allocations (*malloc* performed inside GPU kernels) are extraordinarily expensive in terms of execution time [17]. In-fact, our initial GPU kernel implementation which performed such memory allocations was more than 100× slower than the CPU implementation. An intuitive approach of statically allocating memory at the compile time is not practical as nanopore read lengths vary significantly (~100 bases to >1 Mbases as explained previously) and thus the associated data structures vary from ~200 KB to >1.5 GB. We present a methodology that significantly reduces the number of memory allocations by pre-allocating large chunks of contiguous memory at the beginning of the program to accommodate a batch of reads, which are then reused throughout the life-time of the program. The sizes of these large chunks are determined by the available GPU memory and the average number of events per base (i.e. average value of the number of events divided the by the read length). For a given batch of reads, we assign reads to the GPU until the allocated GPU memory chunks saturate, and the rest of the reads are assigned to the CPU.

We describe the memory allocation technique in two steps: in Section 4.2.1 how the memory allocation for a batch of reads at a time is performed; and, in Section 4.2.2, how the method in Section 4.2.1 can be expanded to reuse large chunks of memory, allocated at the beginning of the program.

#### 4.2.1 Data array serialisation

In the three GPU kernels elaborated in Section 4.1, the associated data arrays per each read are *ref*, *kcache*, *events*, *score*, *trace*, *ll_idx* and *alignment* (final output from the *post-kernel*). If any of these arrays are allocated inside the GPU kernels on a per-read basis, for instance if *score* and *trace* arrays are allocated at line 4 of Algorithm 5 using *malloc*), the performance will be degraded.

We identified that the sizes of all the aforementioned data arrays are dependent only on the read length (known at run-time during file reading) and the number of events for the read (known after event detection described in Section 2). Thus, the sum of read lengths and the number of events for a batch of *n* reads (GPU processes a batch of *n* reads at a time) is used to calculate the sizes of memory allocations required for the particular batch according to the formulation below.

Let *n* be the number of reads loaded to the RAM (from the disk) at a time. Let *r*[] be the read length and *e*[] be the number of events for all the reads in batch of *n* reads. Column 1 of Table 1, lists the data arrays. The size of arrays *ref* and *kcache* depends only on read lengths *r*; *events* and *alignment* depend on number of events *e*; and, *score*, *trace* and *ll_idx* depend on both read length *r* and number of events *e*. Based on these dependencies, the arrays are categorised in Table 1 by horizontal separators. The second column of Table 1 states the data-type size of each array, denoted by constants of the form *c*_*x*_. Typical values of these constants (in our implementation) are given inside the brackets. For instance, the data type for *ref* is *char* and thus *C*_*r*_ is 1 byte. The data type for events is a *struct* of size *C*_*e*_ that is 20 bytes. Note that, the exact values may depend on the implementation and the underlying processor architecture, nevertheless are constants known at compile time. The third column of Table 1 shows the size required for the particular array for a single read, i.e. the size for the i^th^ read (assume 0 based index origin) in the batch of *n* reads. For instance, *ref* depends on the read length of the particular read and the datatype, thus the size is *C_r_r*[*i*]. *Score* depends on read length, number of events, data type size and band-width (*W*), thus *WC*_*s*_(*r*[*i*] + *e*[*i*]). The last column of Table 1 is the total size required for a batch of reads (based on sum of *r* and *e*). For instance, the sum of all the *ref* arrays for the batch is the product of data type size *C*_*r*_ and sum of all read lengths in the batch 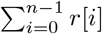.

**Table 1:**
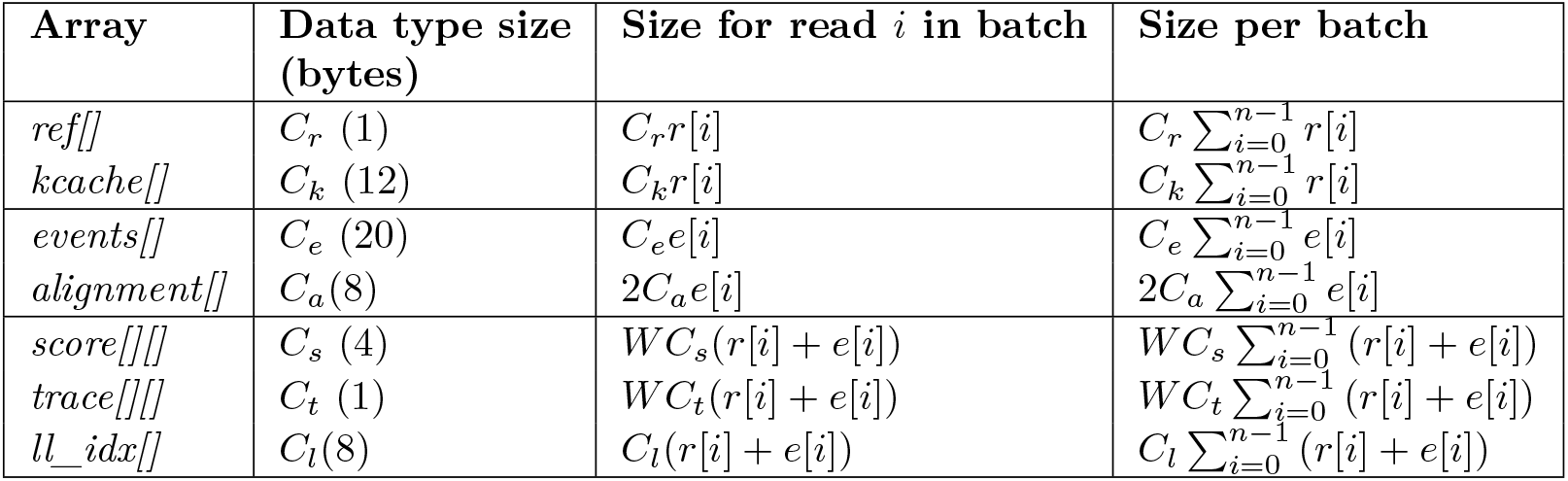
Data arrays associated with ABEA and their sizes

Based on the total array sizes in the last column Table 1, we can allocate seven big chunks of linear contiguous memory in the GPU. Let the base address of those chunks be represented by uppercase letters: *REF*; *KCACHE*; *EVENTS* etc. These memory allocations are performed using *cudaMalloc()* API calls, just before the kernel invocations and are deallocated after the kernels. Note that for now, we do these allocations and deallocations for each batch of reads.

The GPU arrays *REF*, *KCACHE*, *EVENTS* etc, allocated using *cudaMalloc* above are 1D arrays, thus multi-dimensional arrays in the RAM (eg: an array of pointers—each pointer pointing to a string/char array) must be serialised/flattened. One option is to save a series of pointers associated to each above array during the serialisation and then utilising those pointers for addressing a particular element later. However, this can be performed better by storing only two offset arrays of length *n* each: *read offset* array *p*[], which is the cumulative sum of read lengths in the batch 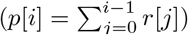; and, *event offset* array *q*, which is the the cumulative sum of events in the batch 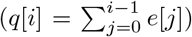. Note that, *r* and *e* have the same definitions as before. These two offset arrays *p* and *q* can be used to deduce the associated pointer to a given element when required, by computing the array offset as shown in Table 2a. The first column of Table 2a is the base address of the large GPU arrays we allocated above. The offset of the element pertaining to the i^th^ read (assume 0-indexing) in the particular array is given in the second column of Table 2a. The definition of constants *C*_*x*_ and *W* are the same as for the previous Table 1. These 1D array base addresses in the first column of Table 2a and the two associated offset arrays *p*[] and *q*[], are passed as arguments to the GPU kernels (Algorithm 5 and Algorithm 6). These arguments are used for the the memory pointer computation inside the GPU kernels (line 4 of Algorithm 5 and line 4 of Algorithm 6) based on the second column of Table 2a.

**Table 2:** GPU data arrays, pointer computation and heuristically determined sizes

(a) Computation of pointer for the read $i$
| 1D GPU array (base address) | Offset to element $i$ in the batch |
| --- | --- |
| $REF$ | $C_r p[i]$ |
| $KCACHE$ | $C_k p[i]$ |
| $EVENTS$ | $C_e q[i]$ |
| $ALIGNMENT$ | $2C_a q[i]$ |
| $SCORE$ | $WC_s(p[i] + q[i])$ |
| $TRACE$ | $WC_t(p[i] + q[i])$ |
| $LL\_IDX$ | $C_l(p[i] + q[i])$ |

(b) Heuristic allocation
| 1D GPU array (base address) | Allocated size per batch |
| --- | --- |
| $REF$ | $C_r X$ |
| $KCACHE$ | $C_k X$ |
| $EVENTS$ | $C_e Y$ |
| $ALIGNMENT$ | $2C_a Y$ |
| $SCORE$ | $WC_s(X + Y)$ |
| $TRACE$ | $WC_t(X + Y)$ |
| $LL\_IDX$ | $C_l(X + Y)$ |

Algorithm 9 elaborates how the above mentioned strategy is integrated into the previous execution flow depicted in Algorithm 4. Lines 3-7 of Algorithm 9 show how the offset arrays *p* and *q* are computed for each batch of reads. Line 8 of Algorithm 9 performs the serialisation of the multi-dimensional arrays with the use of offset arrays *p* and *q*. Line 9 of Algorithm 9 allocates GPU arrays based on sizes in last column of Table 1. Then, the serialised arrays are copied to allocated GPU memory (line 10 of Algorithm 9), GPU kernels (the three kernels discussed in Section 4.1) are executed (line 11) and the alignment result is copied back from the GPU (line 12). At the end, the alignment result is converted back to multi-dimesional arrays (line 13) and then the GPU memory (allocated at line 9) is deallocated (line 14).

The offset arrays *p* and *q* (and also REF, KCACHE, EVENTS, etc.) are passed onto the GPU kernels and are utilised inside the GPU kernels to compute the memory pointers (line 4 of Algorithms 5 and 6) through the equations listed on the second column of Table 2a.

The limitation of this strategy is the GPU memory allocation and de-allocation (line 9 and 14 of Algorithm 9) performed for each batch of reads (which is expensive on certain GPUs, see Section 5.2.2). This limitation is remedied by the heuristic based pre-allocation strategy explained in the next subsection.

**Algorithm 9.**
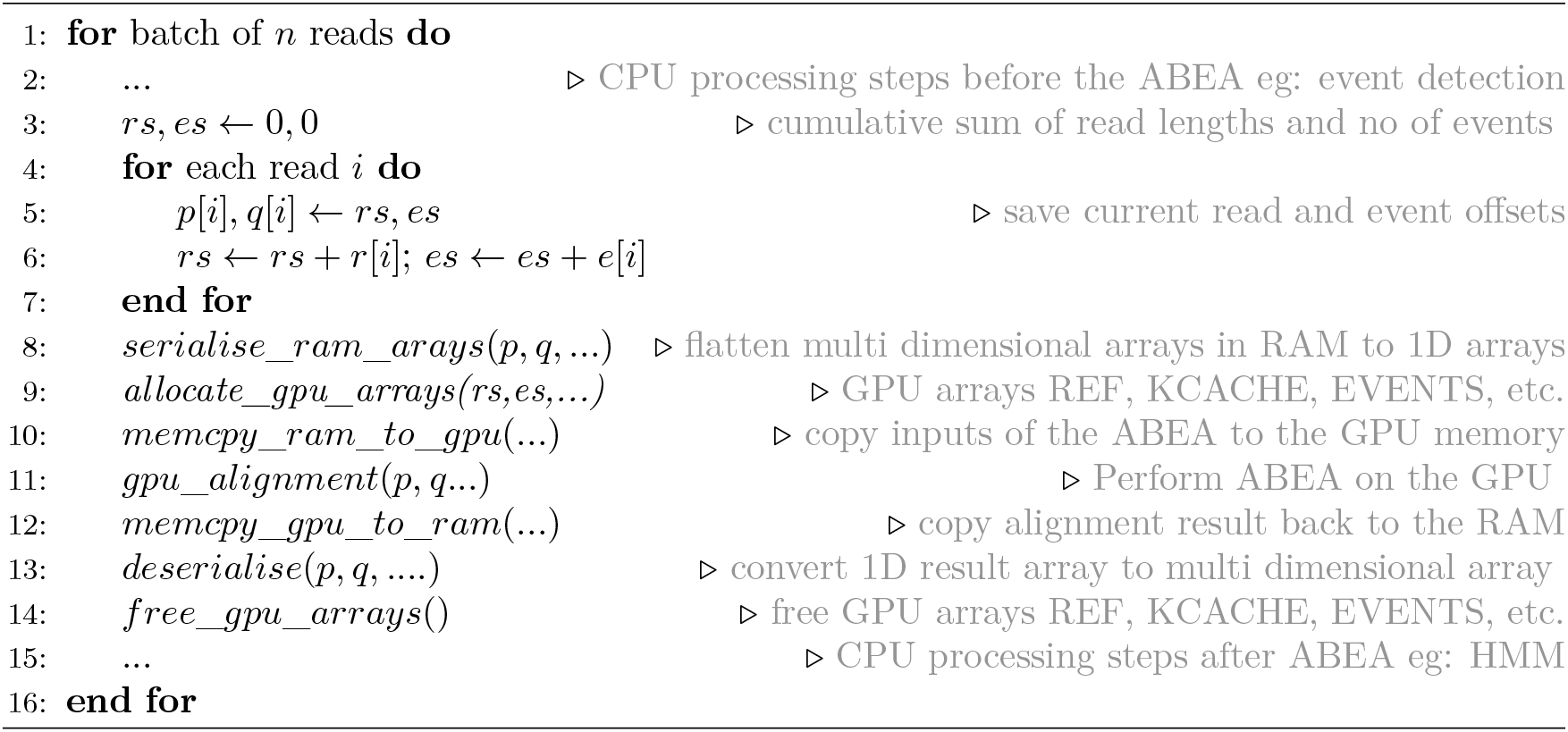
Memory allocation—data structure serialisation.

#### 4.2.2 Heuristic based memory pre-allocation

The GPU memory allocations in the previous section which were performed for each batch could be eliminated by pre-allocating all the available GPU memory at the startup of the program and then re-using for subsequent batches of reads). If the sizes of the arrays depended only on the read length, the total read length accommodable into the available GPU memory can be derived. Then, the available memory can be allocated among the seven large arrays (*REF*, *KCACHE*, *EV ENTS* etc) in correct proportion. However, these array sizes depend both on the read length and the number of events which are unknown at the beginning of the program; thus, memory cannot be partitioned among the data arrays. Therefore, We present a heuristic approach which exploits characteristic of nanopore data to estimate the proportion to maximally utilise the available GPU memory. In summary, we obtain the average number of events per base (average of the number of events divided by read length), use this average to determine the maximum read length that can be accommodated to the GPU, and proportionally allocate the GPU arrays. This approach is formulated as follows.

Sum of all the cells in column 4 of Table 1 is total memory required for a batch of *n* reads. This sum simplifies to equation 1 (due to the properties of constants) where *C*_*R*_ = *C*_*r*_ + *C*_*k*_ + *WC*_*s*_ + *WC*_*t*_ + *C*_*l*_ and *C*_*E*_ = *C*_*e*_ + 2*C*_*a*_ + *WC*_*s*_ + *WC*_*t*_ + *C*_*l*_. This sum represents the total size of all array (for adapted banded event alignment algorithm) for a batch of *n* reads.

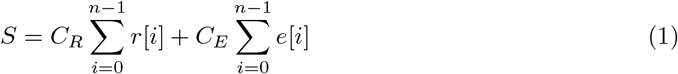

If 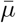 is the average number of events per base (total number of events divided by the total read length for all reads in the batch), we can write as 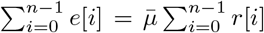. Now substituting this in equation 1 gives 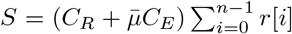. We observed that for a sufficient batch size (>64), 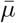; is stable ~2.5 (on more than 10 datasets we tested). Let this estimated value for 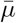 be represented by the constant *μ*. Thus, the total memory required for a batch of reads can be estimated using equation 2.

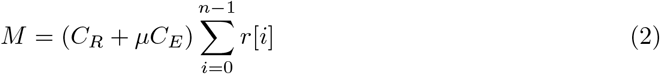

**Algorithm 10.**
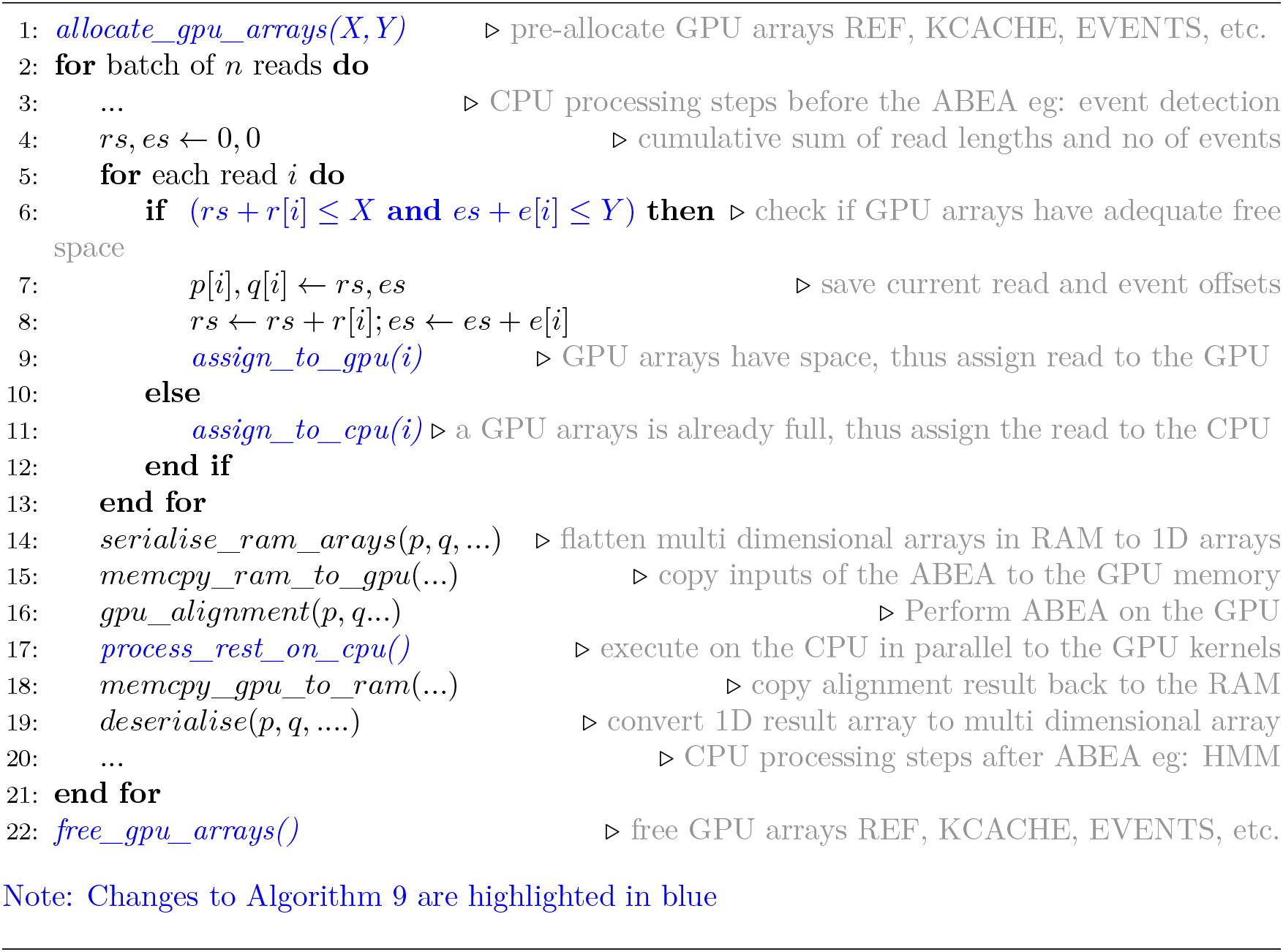
heuristic memory allocation scheme.

Equation 2 can be used to estimate the maximum number of bases (sum of read lengths) that a given amount of GPU memory can accommodate. Let *M* in equation 2 be the available GPU memory. Then, the approximate maximum number of bases *X* that fits available GPU memory *M* can be computed via equation 3. Then, the associated total number of total events *Y* which the GPU memory can accommodate, is found by equation 4.

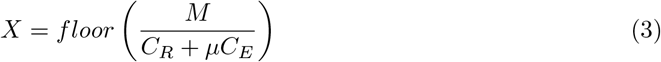

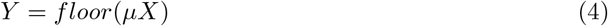

These *X* and *Y* allow the available GPU memory to be allocated among the seven large arrays (*REF*, *KCACHE*, *EV ENTS* etc) with approximately correct proportions, as shown in the second column of Table 2b. The values in the second column of Table 2b are obtained by substituting 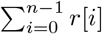 with *X* and 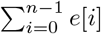 with *Y* in the last column of Table 1.

By incorporating the above heuristic based memory allocation strategy to Algorithm 9, we get the execution flow in Algorithm 10. The major changes to the previous Algorithm 9 are highlighted in blue text. Now the GPU memory is allocated at the beginning of the program based on the estimated *X* and *Y* on line 1 of Algorithm 10. As *X* and *Y* are approximations, the GPU arrays may saturate for certain batches of reads. Line 6 of Algorithm 10 checks if GPU arrays are saturated and assigns the read to either GPU (line 9) or CPU (line 11), accordingly. Only a few reads are assigned to the CPU and these few reads are processed on the CPU in parallel to the GPU kernel execution, and thus no additional execution time is incurred.

With the heuristic based memory pre-allocation strategy described in this section, *cudaMalloc* operations are invoked only at the beginning of the program and thus no additional memory allocation overhead during the processing. Note that, our implementation is future proof; i.e. *μ* is a user specified parameter (that is initialised to 2.5 by default) in case nanopore data characteristics change in future.

### 4.3 Heterogeneous processing

If all the reads were of similar length, GPU threads that process the reads would complete approximately at the same time, and thus GPU cores will be busy throughout the execution. However, as stated in Section 2, there can be a few reads which are significantly longer than the other reads (we will refer to them as *very long reads*). When the GPU threads process reads in parallel, presence of such *very long reads* will cause all other GPU threads to wait until the GPU threads processing the longest read complete. This thread waiting leads to under utilisation of GPU cores. Thus, we process these *very long reads* on the CPU while the GPU is processing the rest in parallel. However, there can be exceptionally long reads (we will refer to them as *ultra long reads*) which the CPU would take longer time than what the GPU took to process the whole batch. Such reads would lead the GPU to idle until the CPU completes. Thus, *ultra long reads* will be skipped and will be processed separately at the end by the CPU. Similarly, there can be a few *over segmented reads* (i.e. reads with a significantly higher events per base ratio than the others) which cause GPU under utilisation. These over-segmented reads will also be processed on the CPU.

We discuss these problems of *very long reads* and *ultra long reads* in detail with examples in Section 4.3.1, along with the solutions. Then, in this Section 4.3.2, we discuss the problem of over segmented reads and the respective solution. Then, in Section 4.3.3, we discuss another factor that affects performance, the batch size (number of reads loaded to the RAM at a time). Finally, in Section 4.3.4, we describe a method to detect and prompt the user of any drastic impacts on performance along with suggestions to tune parameters to minimise the impact.

#### 4.3.1 Very long reads and ultra long reads

Consider a batch of reads where ~90% of the reads are less than 30 Kbases in length. Assume the longest read in the batch is 90 Kbases. Assume that the GPU is processing all the reads (in the batch) in parallel. Suppose that GPU threads processing reads of length <30 Kbases (90% of the threads) would complete in <300ms while GPU threads processing the longest 90 Kbases read would take 900ms. As a result, the completed GPU threads will have to wait for additional 600ms. Similarly, the few *very long reads* consume a significant time to process on the GPU in comparison to other reads in the batch. Majority of the GPU threads will have to wait and this causes under-utilisation of GPU compute-cores. Furthermore, *very long reads* negatively affects the GPU occupancy by occupying a significant portion of GPU memory. For instance, a read of size ~10 Kbases requires only ~18 MB of GPU memory while a read with 90 Kbases requires ~160MB memory. Hence, *very long reads* occupy a significant portion of GPU memory, limits the number of reads that could be processed in parallel. This reduces the amount of parallelism and the occupancy of the GPU is reduced.

Fortunately, *very long reads* being few (see the typical read length distribution under results), the CPU (core frequency faster than on GPU) could process those reads while GPU is processing the rest of the reads. In the above example, selecting a static threshold (eg: processing reads of length <30Kbases on GPU and rest on CPU) would give reasonable performance. However, selecting such a static threshold is not ideal due to variations in the read length distributions based on the dataset (see background). Thus, we use the product of *max-lf* and the average read length in the batch to determine the threshold dynamically, where *max-lf* is a user-parameter that defaults to 5.0. This threshold was empirically determined.

Now assume amongst the *very long reads* processed on the CPU, a few *ultra long reads* (eg: read >100 Kbases in a dataset where >99% of the reads are <100 Kbases). Such *ultra long reads* could cause a severe load imbalance between the CPU and the GPU. For instance, assume that there exists a read which is 1 Mbases in a given read batch. Despite the high core frequency, the CPU will take a few seconds to process such an *ultra long read*. The GPU meanwhile would process the whole batch in less than 1s (see results for empirical evidence). Such *ultra long reads* being <1%, are skipped during the processing (while being written to a separate file) and are separately processed by the CPU at the end. In our implementation, the threshold for *ultra long reads* is a user defined parameter which defaults to 100 Kbases. There is an additional advantage of processing *ultra long reads* later. *Ultra long reads* usually require a significant amount of RAM (a few gigabytes) and may crash on limited memory systems. In the end, it is possible to process these reads with a limited amount of threads to reduce the peak memory consumption, particularly if the size of the RAM is limited.

#### 4.3.2 Over segmented reads

Once the *very long reads* and *ultra long reads* are processed as in Section 4.3.1, the performance impact due to the over-segmented events become prominent. While majority of the reads have a number of events per base that is close to the average *μ*(= 2.5), a few reads can have a very large value. For instance, a few reads with a number of events per base being more than eight times the average *μ*(= 2.5) can violate the suitability of our partitioning of GPU memory as *X* and *Y* (*X* and *Y* are derived in equations 3 and 4). These over-segmented reads lead to the GPU arrays that are proportional to *Y* be full, while the arrays proportional to *X* are left under-utilised. For instance, arrays proportional to *Y* can become 100% while arrays proportional to *X* are only filled to <70%. Hence, over segmented reads lead to under-utilisation of GPU memory and results in limiting the number of reads which are processed in parallel. We process the over-segmented reads on the CPU based on a user specifiable threshold *max-epk* which defaults to 5.0.

On rare occasions, reads with >100 events per base were observed. Such severely over-segmented reads can be processed separately at the end or ignored totally as such rare reads amongst millions of other reads are unlikely to affect the final polishing result.

#### 4.3.3 Batch size

Selection of proper batch size (reads loaded to RAM from the disk at a time) is another important parameter that affects performance. If the batch size is too small compared to what the GPU memory can accommodate, the number of reads to be processed in parallel is limited, thus leads to in-adequate occupancy. Conversely, if the batch is too large to fit the GPU, CPU will have to process many surplus reads that could not be accommodated into the GPU. The batch size in our implementation is determined by two user specified parameters: *K* which is the maximum number of reads; and, *B* which is the maximum number of total bases. When reading from the disk to RAM, the true batch size (*n*-number of reads and *b*-number of total bases are capped by *K* and *B*) is determined by the first value (*n* or *b*) reaching the cap (*K* or *B*) first. Having such a limit *B* allows to cap peak RAM due to adjacent *very long reads*. The suitable value for *B* is dependent on the available GPU memory, which can be estimated via the equation 3 discussed in Section 4.2.

#### 4.3.4 Detection of performance anomalies

While we have empirically determined typical parameters/thresholds (associated with above strategies), an unusual situation (for instance, a big gap between the CPU and GPU specifications or a data set that severely deviates from the heuristics we use) may cause performance anomalies. We employ the following method to detect a severe performance anomaly caused by such an unusual scenario.

We measure the quantities representing resource utilisation during run time, which are listed in Table 3. These quantities are measured per batch of reads loaded to the RAM at a time. We use those measured quantities to determine any severe performance issues and suggest suitable parameter adjustments to the user. The adjustable parameters (or thresholds) that can be tweaked to improve the resource utilisation are defined in Table 4. Determination of performance issues and suggestions are done via two decision trees, one that corresponds to GPU memory usage (Fig. 9a) and another which corresponds to balancing the load between CPU and GPU (Fig. 9b).

**Table 3:** measured quantities

| quantity | description |
| --- | --- |
| $t_{\text{CPU}}$ | processing time on CPU |
| $t_{\text{GPU}}$ | processing time on GPU |
| $X_{\text{util}}$ | utilisation percentage of the arrays proportional to $X$ ($rs$ as a percentage of $X$ in Algorithm 10) |
| $Y_{\text{util}}$ | utilisation percentage of the arrays proportional to $Y$ ($es$ as a percentage of $Y$ in Algorithm 10) |
| $N_{\text{memout}}$ | number of reads assigned to CPU due to GPU memory getting prematurely full (corresponds to line 11 of Algorithm 10) |
| $N_{\text{long}}$ | number of *very long reads* assigned on to the CPU (corresponds to user parameter *max-lf*) |
| $N_{\text{events}}$ | number of reads with too many events per read assigned onto the the CPU (corresponds to user parameter *max-epk*) |
| $n$ | the number of reads actually loaded to the RAM |
| $b$ | the number of bases actually loaded to the RAM |

**Table 4:**
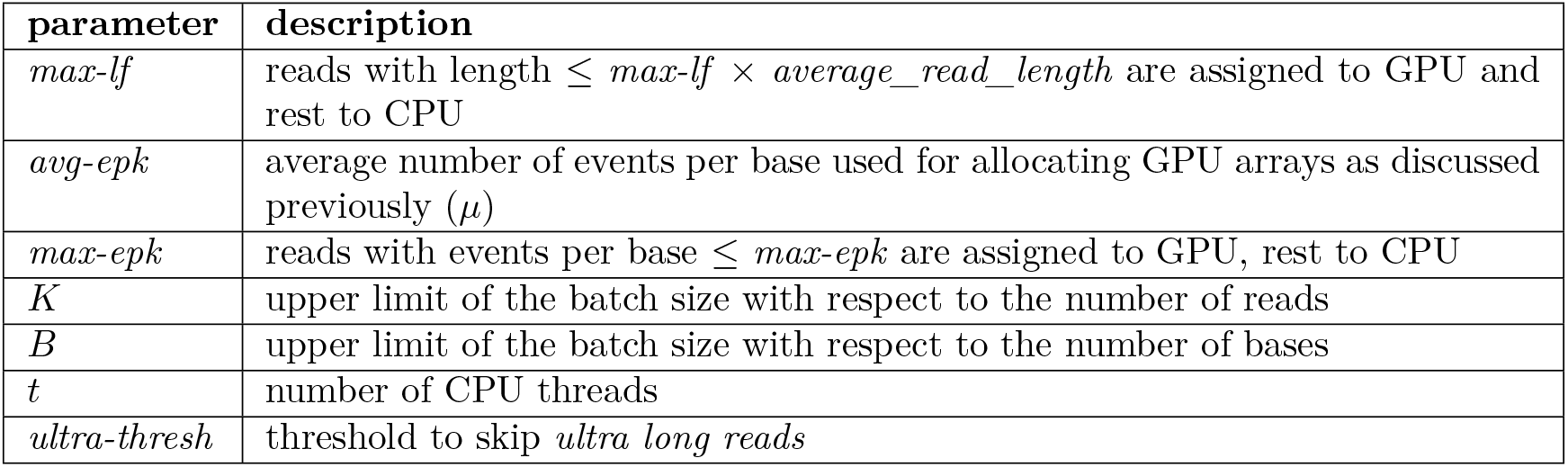
adjustable user parameters

**Figure 9:**
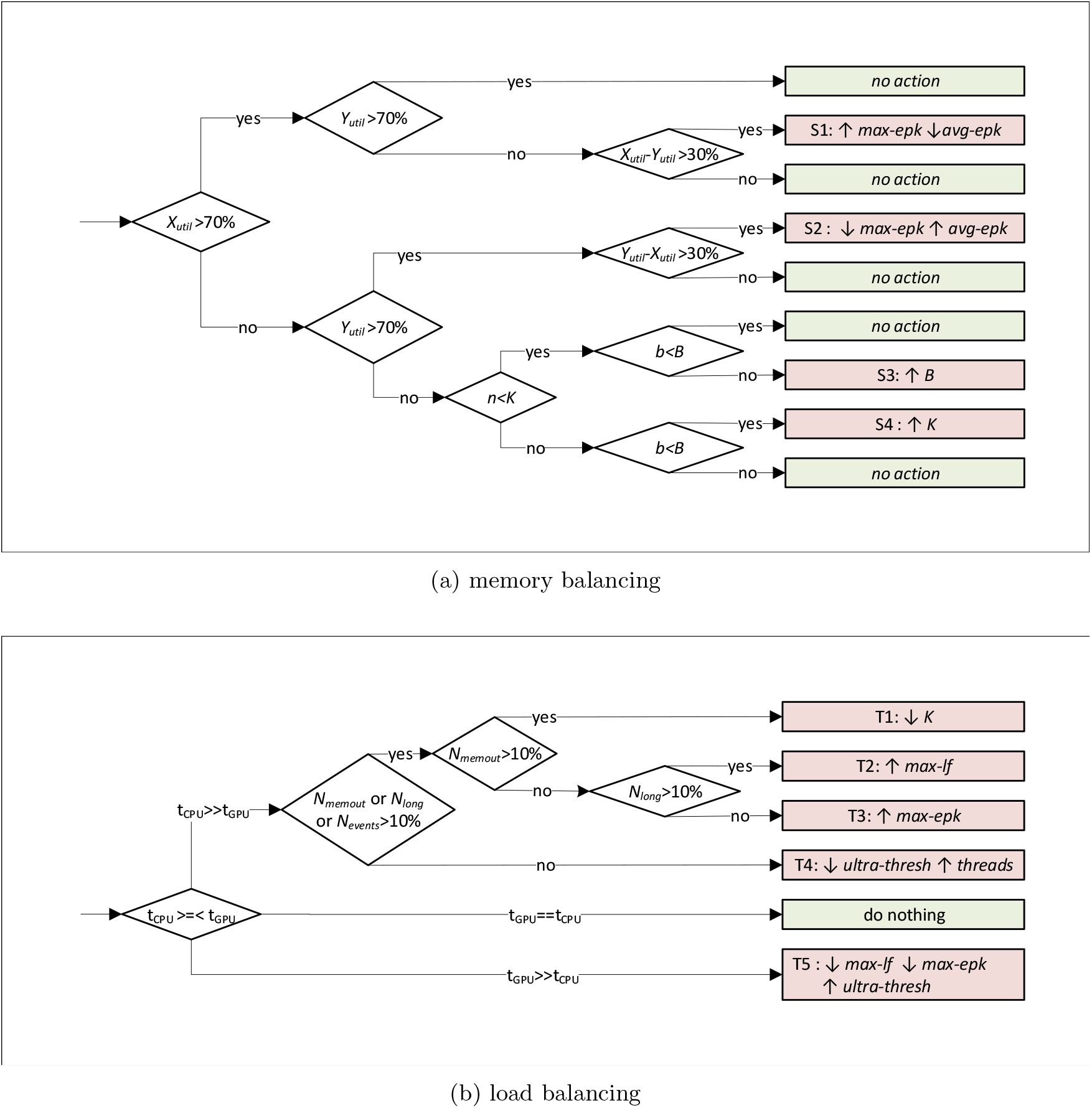
Decision trees for resource optimisation

Fig. 9a shows the decision tree that detects any imbalance in the proportions *X* and *Y* associated with GPU arrays allocation (*X* and *Y* derived in equations 3 and 4). The objective of this decision tree is to detect any GPU memory wastage and to increase the number of reads which the GPU gets to process in parallel.

As shown in Fig. 9a, if both *X*_*util*_ and *Y*_*util*_ (*rs* as a percentage of *X* and *es* as a percentage of *Y* in Algorithm 10) are more than 70%, the utilisation of GPU arrays is considered reasonable. Note that 70% is an empirically determined value that provides adequate performance. If *X*_*util*_ is reasonable (>70%) and *Y*_*util*_ is unreasonable (<70%), we inspect for any significant imbalance between *X*_*util*_ and *Y*_*util*_ (*X*_*util*_-*Y*_*util*_>30%). Such a significant gap suggests an under-utilisation, which should be remedied through the increase of *max-epk* (the threshold at which over-segmented reads are offloaded to the CPU) or reducing *Y* by decreasing *average-epk* (node S1 in Fig. 9a). In contrast, if *Y*_*util*_ is reasonable and the *X*_*util*_ is unreasonable, the strategy is the opposite, i.e, either decrease *max-epk* or increase *average-epk* (follow up to the node S2 in Fig. 9a).

If both *X*_*util*_ and *Y*_*util*_ are less than 70%, a likely cause is an inadequate batch size to fill the GPU memory. The actual batch size (*n*,*b*) is determined by both *K* and *B* as stated previously. As shown in Fig. 9a, we check which limit out of *K* and *B* was reached first. If both *n* < *K* and *b* < *K*, the currently processed batch being the last batch in the dataset (end of input data reached) is the likely cause. Thus, no parameter tuning action is necessary. If *B* was reached first (*n* < *K* and not *b* < *B*), *B* is the limitation and should be increased (S3 in Fig. 9a). If *K* was reached first (not *n* < *K* and *b* < *B*), *K* should be increased (S4 in Fig. 9a).

Fig. 9b shows the decision tree for CPU-GPU workload balancing. For a particular batch, if the CPU takes significantly more time than the GPU, the decision tree first inspects whether the CPU is assigned with an excessive workload. An excessive workload on the CPU can be attributed by: an extensively over-sized batch size (in comparison to the available GPU memory), which results in a majority of the reads being assigned to the CPU (N_memout_>10%); excessive number of *very long reads* assigned to the CPU (N_long_>10%); and, excessive number of over-segmented reads events assigned to the CPU (N_events_>10%). If N_memout_>10%, *K* is reduced (node T1 in Fig. 9b); if N_long_>10%, *max-lf* is increased (T2 in Fig. 9b); and, if N_events_>10%, *max-epk* is increased (T3 in Fig. 9b).

If the cause for higher CPU time is not the aforementioned excessive workload, a likely cause is *ultra long reads*, where a single *ultra long reads* processed on the CPU taking more time than the time taken by GPU for the whole batch. In such an event, *ultra-thresh* threshold is reduced so that more *ultra long reads* are skipped. Another likely cause is that the program was executed with inadequate threads (if the CPU had more hardware threads than the program was launched), which is to be remedied by increasing the number of CPU threads. Another cause might be that the CPU is not sufficiently powerful to match with the GPU and thus no action can be taken (except upgrading the CPU). These actions are denoted by T4 in Fig. 9b.

The ideal case is when the CPU and GPU take similar times which requires no intervention. Conversely, if the GPU takes significant time than the CPU, the likely causes are *very long reads* or over-segmented reads. In such event, the thresholds *max-lf* and *max-epk* are decreased so that more *very long reads* and over-segmented reads are assigned to the CPU. Another likely cause is the *ultra long read* which can be remedied by increasing *ultra-thresh* threshold. Another cause might be an insufficiently powerful GPU (less compute cores or less memory) compared to the CPU and no action is taken (except to upgrade the GPU).

To reduce false positives due to incidental under utilisation, a suggestion is provided to the user, only if the same condition (condition that led to the decision in the decision tree, S1 to S4 T1 in Fig. 9a and T1 to T4 in Fig. 9b) consecutively repeats more than a few times (eg: >3 times).

Note that the above mentioned strategy is to warn and suggest of potential parameter adjustments in the event of drastic performance degradation, rather than to obtain optimal performance or to determine the exact parameter values.

## 5 Results

Experimental setup is given in Section 5.1. In Section 5.2, we present experimental evidence that justify the selection of steps presented in Section 4. Next in Section 5.3, we compare the GPU implementation of the Adaptive Banded Event Alignment (ABEA) algorithm to its CPU implementation. Finally, we show the overall speedup of the GPU implementation when incorporated into an actual work-flow (i.e. detection of methylated bases).

### 5.1 Experimental setup

We re-engineered the *Nanopolish* methylation calling tool (existing methylation detection tool discussed in Section 2) to: one, load a batch of *n* reads from disk to RAM at a time, instead of on-demand loading; two, synchronise CPU threads prior to GPU kernel invocation (*Nanopolish* assigns a thread dynamically to a particular thread, thus each read follows its own code path); and three, optimise the CPU implementation which otherwise would result in an apparent unfair speedup (when the optimised GPU version is compared to an un-optimised CPU version). Re-engineered *Nanopolish* employs a fork-join multi-threading model (with work stealing) implemented using C POSIX threads. ABEA algorithm for the GPU was implemented using CUDA C. This re-engineered *Nanopolish* will be hitherto referred to as *f5c*.

We used publicly available NA12878 (human genome) Nanopore WGS Consortium sequencing data [12] for the experiments. The datasets used for the experiments, their statistics (number of reads, total bases, mean read length and maximum read length) and their source are listed in Table 5. D_small_ which is a small subset, is used for running on a wide range of systems (all systems in Table 6: embedded system, low-end and high-end laptops, workstation and high-performance server). Two complete MinION data sets (D_ligation_ and D_rapid_) are only tested on three systems due to large run-time and incidental access to the other two systems. D_ligation_ and D_rapid_ represent the two existing nanopore sample preparation methods (ligation and rapid [26]) that affects the read length distribution.

**Table 5:** Information of the datasets

| Dataset | Number of reads | Number of bases (Gbases) | Mean read length (Kbases) | Max read length (Kbases) | Source / SRA accession |
| --- | --- | --- | --- | --- | --- |
| D<sub>small</sub> | 19275 | 0.15 | 7.7 | 196 | [27] |
| D<sub>ligation</sub> | 451020 | 3.62 | 8.0 | 1500 | ERR2184733 |
| D<sub>rapid</sub> | 270189 | 2.73 | 10.0 | 386 | ERR2184734 |

**Table 6:** Different systems used for experiments

| System Name | Info | CPU | CPU cores / threads | RAM (GB) | GPU | GPU mem (GB) | GPU arch |
| --- | --- | --- | --- | --- | --- | --- | --- |
| SoC | NVIDIA Jetson TX2 embedded module | ARMv8 Cortex-A57 + NVIDIA Denver2 | 6 / 6 | 8 | Tegra | shared with RAM | Pascal / 6.2 |
| lapL | Acer F5-573G laptop | i7-7500U | 2/4 | 8 | Gefore 940M | 4 | Maxwell / 5.0 |
| lapH | Dell XPS 15 laptop | i7-8750H | 6/12 | 16 | Gefore 1050 Ti | 4 | Pascal / 6.1 |
| ws | HP Z640 workstation | Xeon E5-1630 | 4/8 | 32 | Tesla K40 | 12 | Kepler / 3.5 |
| HPC | Dell PowerEdge C4140 | Xeon Silver 4114 | 20/40 | 376 | Tesla V100 | 16 | Volta / 7.0 |

D_small_ dataset was used for experiments under Sections 5.2.2, 5.2.1 and 5.3.1. For experiments under Sections 5.3.2 and 5.4, the datasets D_rapid_ and D_ligation_ were used.

To obtain the results for Section 5.2.3, first we grouped the reads in dataset D_rapid_ based on their read lengths. We grouped the read into 10 Kbases bins (i.e., 0K-10K,10K-20K…90K-100K). Reads with >100 Kbases were grouped into larger bins (100K bin sizes; 100K-200K, 200K-300K and 200K-300K) as the read count is very little in the range that certain 10K bins would contain no reads at all. Then, we ran *f5c* with only CPU and *f5c* with GPU acceleration on each group of the reads separately. Then, we computed the speedup of ABEA for each group of reads: the kernel only speedup (*GPU kernel time / time on CPU*); and, the speedup with overheads (overheads such as memory copy, data structure serialisation). This experiment was performed on the system *lapH*.

For Sections 5.2 and 5.3, time measurements were obtained by inserting *gettimeofday* function invocations directly into the C source code. Total execution time and the peak RAM usage in Section 5.4 were measured by running the *GNU time* utility with the *verbose* option.

### 5.2 Effect of individual optimisations

#### 5.2.1 Compute optimisations

Fig. 10a shows the time consumed by the three GPU kernels after applying the compute optimisation techniques discussed in Section 4.1. Time taken by each of the three GPU kernels (*pre-kernel*, *core-kernel* and *post-kernel*) is plotted for each different GPU. It is observed that the *core-kernel*, which computes the dynamic programming table (compute-intensive portion), still consumes the majority of the GPU compute time. The *pre-kernel* which performs data structure initialisation consumes much lesser time and shows that there is no need to further parallelise the loop in Algorithm 5 (explained in Section 4.1). Despite the lack of fine-grained parallelism in *post-kernel* (which performs backtracking), the elapsed time is still considerably lesser than the *core-kernel*. Thus, any future optimisations should still mainly focus on the *core-kernel*, followed by the *post-kernel*.

**Figure 10:**
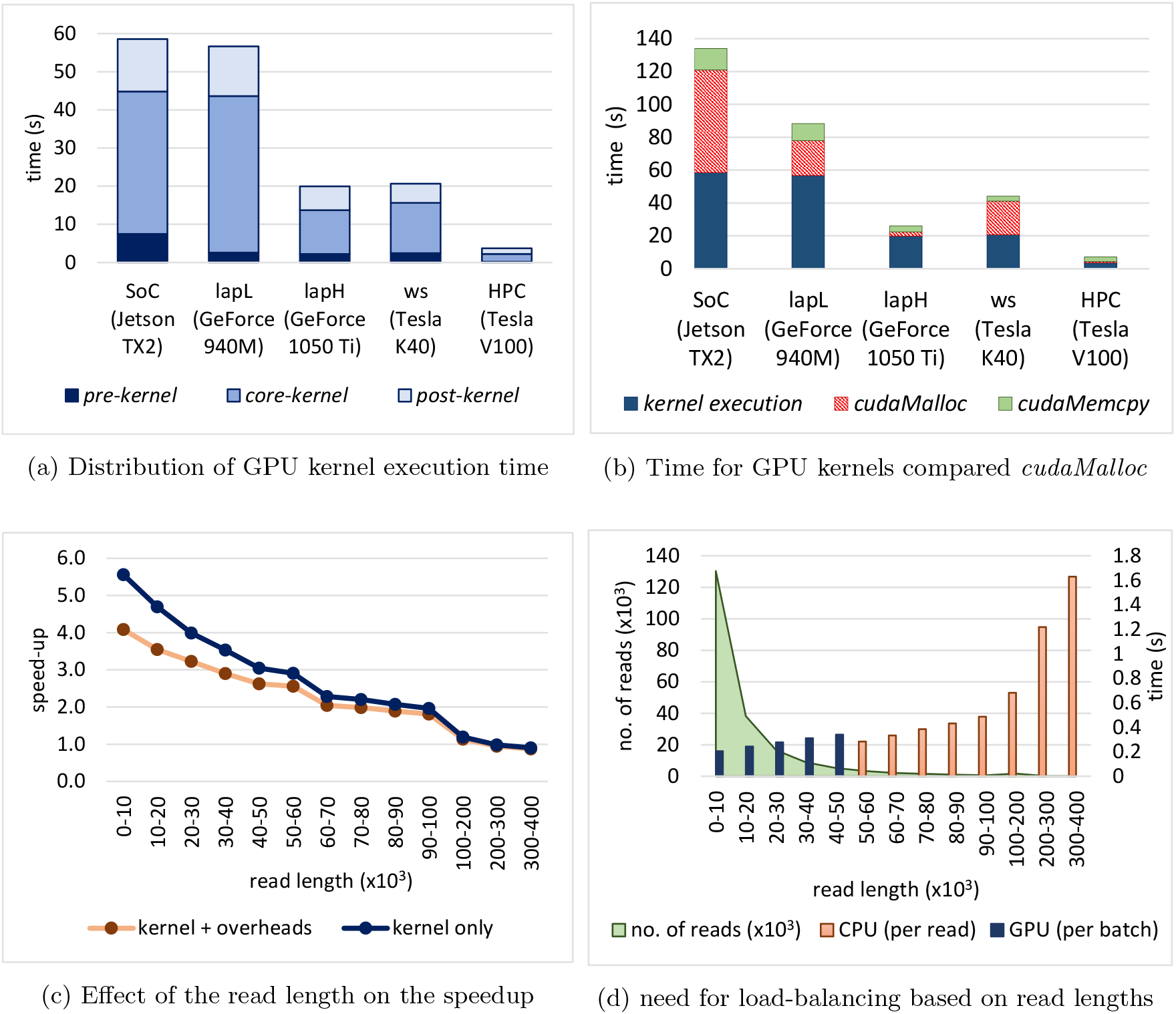
Effect of individual optimisations

The efficacy of our compute optimisations on the compute intensive *core-kernel* can be elaborated using the reported statistics from the NVIDIA profiler (instruction level profiling—PC sampling in NVIDIA visual profiler [28]). The profiler reports the percentage distribution of reasons that caused the thread warps to stall, based on the number of clock cycles. The percentage of the number of clock cycles that a warp was stalled due to a memory dependency (waiting for a previous memory accesses to complete), improved from 59.10% to 44.81% after the use of GPU shared memory. After exploiting the *kcache* for improving memory coalescing, this percentage further improved to 28.62%.

#### 5.2.2 Memory optimisations

As stated in Section 4.2.1, the data array serialisation technique eliminated all memory allocations inside GPU kernels (*malloc*); still, required memory allocations per each batch of reads (*cudaMalloc*). The overhead due to these *cudaMalloc* calls are plotted in Fig. 10b along with the time for kernel execution and data transfer to/from the GPU (using *cudaMemcpy*). Observe that on certain GPUs (Jetson TX2, GeForce 940M and Tesla K40), the overheads due to *cudaMalloc* operations are significant in comparison to the compute kernels (even higher than the compute kernels in Jetson TX2). Such significant overheads justify why we proposed a heuristic based memory pre-allocation technique (Section 4.2.2) which completely eliminates this overhead.

Interestingly, Tesla K40 and Gefore 940M which incurred high *cudaMalloc* overheads are of relatively older GPU architectures in comparison to GeForce 1050 and Tesla V100, where the overheads were minimal. This is probably due to hardware supported memory allocation in latest GPU architectures. However, the aforementioned observation seems to be valid only for GeForce GPUs (targeted for gaming on PC/laptops) and Tesla GPUs (targeted for high performance computing). On Tegra GPUs (SoC targeted for embedded devices) the overhead seems to be significant in spite of the latest architectures (Jetson TX2 is the same Pascal architecture as GeForce 1050). We additionally tested on a Jetson AGX Xavier (the most recent Tegra GPU based SoC — Volta architecture) and *cudaMalloc* was yet expensive (40s on GPU kernels and 44s on *cudaMalloc*, not shown in figure). Thus, our memory pre-allocation strategy (in Section 4.2.2) which totally eliminates this *cudaMalloc* overhead is specifically beneficial for GPU on SoCs.

#### 5.2.3 Heterogeneous processing

We stated in Section 4.3 that *very long reads* if processed on the GPU, limits the GPU occupancy. Fig. 10c provides experimental evidence and shows the need to process *very long reads* on CPU (explained in Section 4.3). Fig. 10c plots the variation of the speedup (GPU compared to CPU for ABEA) as the read length varies. The x-axis labels the range of the read length for which the speedup was computed (explained in the experimental setup). For instance, 0-10 on the x-axis refers to the group of reads with read length 0-10Kbases. Note that in Fig. 10c the bins are 100K wide from 100K-200K on-wards, due to less number of reads of those lengths (explained in the experimental setup). The speedup of *computations* (GPU kernel time / CPU time) and the speedup including *overheads* (GPU kernel time + overheads such as memory copy, data structure serialisation) are plotted in Fig. 10c. Speedup of more than 4X was observed for smaller read lengths (0-10K). speedup drops with increasing read-length and is less than 3X from 50K-60K. The longer the reads are, the lesser number of reads can be processed in the GPU in parallel (reduced occupancy), thus the reduced speedup. Hence, *very long reads* that significantly affects the performance should be performed on the CPU while the GPU is processing the rest.

Fig. 10d shows the need for processing *ultra long reads* separately (explained in Section 4.3). The x-axis in the figure is the read-length (similar to Fig. 10c). The blue bars (with reference to the right y-axis) denote the average time consumed by the GPU to process a batch of reads (1.5 Mbases), for each group of read lengths from 0 bases to 50Kbases. The orange bars (with reference to the right y-axis) denote the average time consumed by the CPU (1 thread) to process a single read in the particular group of reads. The read length distribution (left y-axis) is shown shaded in green colour to depict the abundance of reads in each read length. Observe that CPU takes >1.6s for a single read of 300K-400K length while the GPU completes a whole 40K-50K batch in <0.4s. Thus, the GPU would idle for >1.2s until the CPU completes processing. Hence, such *ultra long reads* (eg : >100 Kbases) must be skipped and processed separately at the end. Note that such *ultra long reads* are very few (green coloured read length distribution in Fig. 10d).

### 5.3 Speedup of Adaptive Banded Event Alignment

In this subsection, we present the performance of the GPU ABEA implementation when all the optimisations in Section 4 are applied together. Note that we compare this optimised GPU version with optimised CPU version in *f5c* (not the CPU version in original *Nanopolish*). The CPU version was run with maximum supported threads on the system. The optimised CPU version will be hitherto referred to as *CPU-opti* and the optimised GPU version will be referred to as *GPU-opti*. First, we compare the run-time of *CPU-opti* and *GPU-opti* on a wide range of different computer systems in Section 5.3.1, and then on the two big datasets in Section 5.3.2.

#### 5.3.1 Across different devices

Fig. 11a shows the time for *CPU-opti* (left bars) and the *GPU-opti* (right bars) for the Dataset D_small_, for each system listed in Table 6. The run-time for the GPU has been broken down in to: compute kernel time; different overheads (memory copying to/from the GPU, data serialisation time); and, the extra CPU time due to reads processed in the CPU. The compute kernel time includes the sum of time for all the three kernels (*pre-kernel*, *core-kernel* and *post-kernel*). The extra CPU time is the additional time spent by the CPU to process the reads assigned to the CPU (excluding the processing time that overlaps with the GPU execution, i.e. only the extra time which the GPU has to wait after the execution is included). Note that the *ultra long reads* were not separately processed on the CPU as the D_small_ contains a minuscule number of *ultra long reads*.

**Figure 11:**
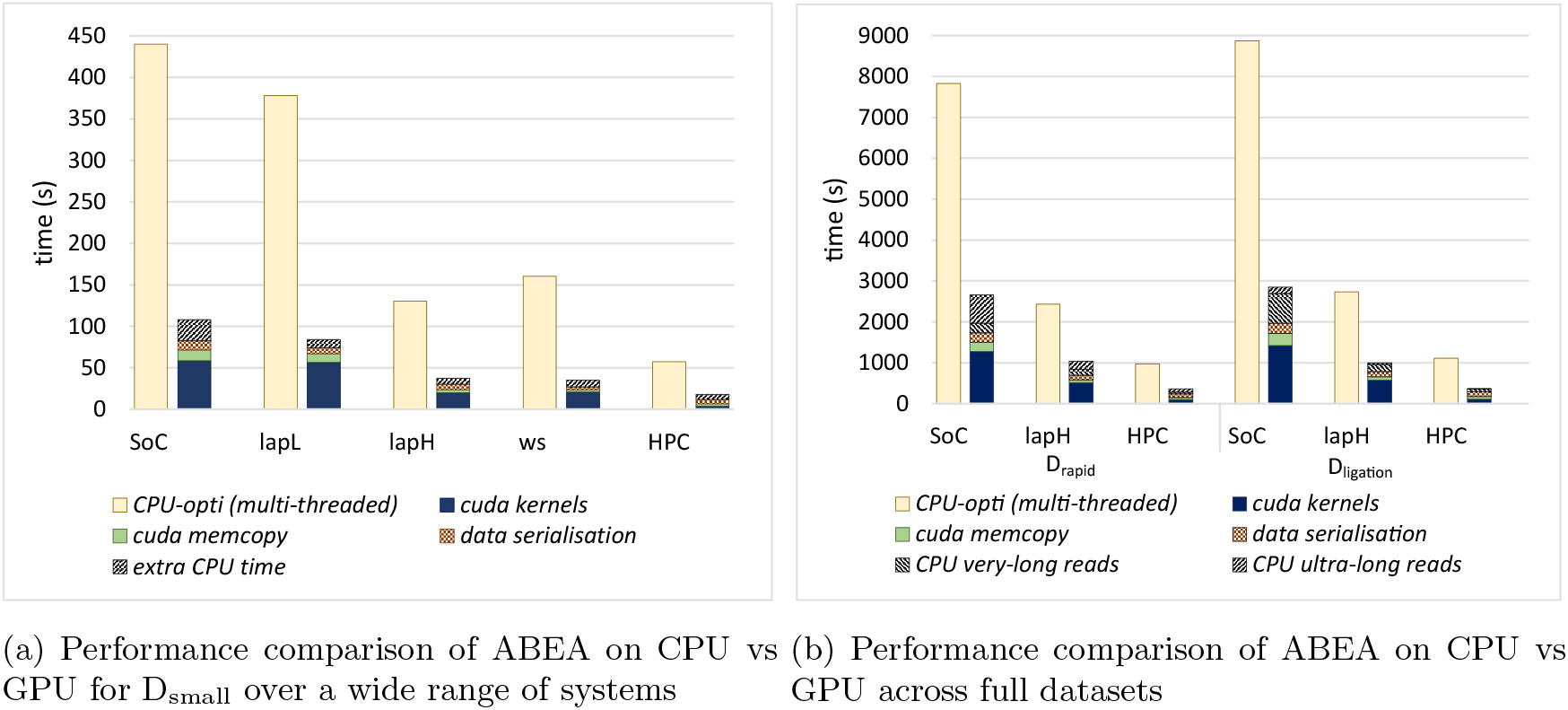
Speedup of ABEA on GPU compared to CPU

Speedups (including all the overheads) observed for *CPU-opti* compared to *GPU-opti* are: ~4.5× on the low-end-laptop and the workstation; ~4× on Jetson TX2 SoC; and ~3× on high-end-laptop and HPC. Note that only ~3× speedup on high-end-laptop and HPC (in comparison to >=4× on other systems) is due to the CPU on those particular systems having comparatively a higher number of CPU threads (12 and 40 respectively).

#### 5.3.2 Benchmark on big datasets

Time taken for *CPU-opti* compared to *GPU-opti* for the two big datasets (D_ligation_ and D_rapid_) is shown in Fig. 11b. Experiments were performed only on three systems due to the limited availability of other devices (mentioned previously). The graph is similar to the previous Fig. 11a, except the *extra CPU time* has been further broken down to: *CPU very-long reads*; and, *CPU ultra long reads*. *CPU very long reads* refers to the additional time spent by the CPU to process *very long reads* and, *CPU ultra long reads* refer to the *ultra long reads* (reads >100 Kbases) processing time performed separately on the CPU. A speedup up of ~3× was observed for all three systems. Due to more *ultra long reads* in D_ligation_ and D_rapid_ than in D_small_, the overall speedup for *SoC* is limited to around ~3× compared to ~4× for D_small_.

### 5.4 Total run-time of *f5c* compared with original *Nanopolish*

In this section, we demonstrate the overall performance when the GPU accelerated ABEA is incorporated into an actual methylation detection work-flow. As stated in the experimental setup, we re-engineered *Nanopolish* to overcome the limitations of original *Nanopolish*. We compare the total run-time for methylation calling using original *Nanopolish* against *f5c* (both CPU only and GPU accelerated versions).

We refer to original *Nanopolish* (version 0.9) as *nanopolish-unopti*, *f5c* run only on the CPU as *f5c-cpu-opti* and GPU accelerated *f5c* as *f5c-gpu-opti*. We executed *nanopolish-unopti*, *f5c-cpu-opti* and *f5c-gpu-opti* for the full datasets D_rapid_ and D_ligation_. Note that all the executions were performed with the maximum number of CPU threads supported on each system.

The run-time results are shown in Fig. 12. The reported run-times are for the whole methylation calling (all steps mentioned in Section 2.2) and also includes disk I/O time. As each read executes on its own code path in original *Nanopolish* (as mentioned in the experimental setup) the time for individual components (eg: ABEA) cannot be accurately measured, thus we only compare the total run-times.

**Figure 12:**
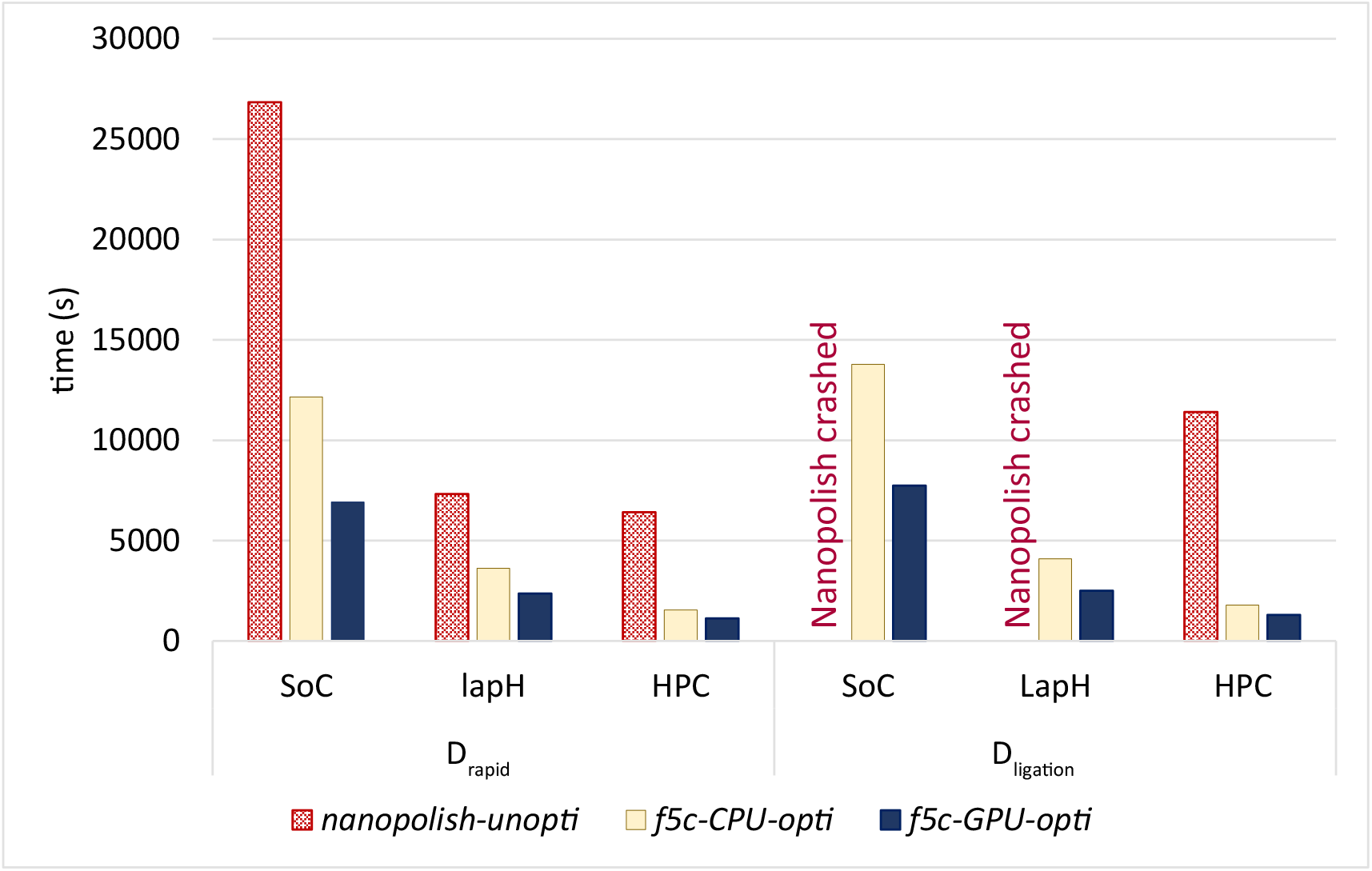
Comparison of *f5c* to *Nanopolish*

*f5c-cpu-opti* for D_rapid_ dataset was: ~2× faster on SoC and lapH; and, ~4× faster on HPC. *nanopolish-unopti* crashed on SoC (8GB RAM) and lapH (16GB RAM) when run for D_ligation_ dataset due to Linux Out Of Memory (OOM) killer [29]. When run for D_ligation_ on HPC, *f5c-cpu-opti* was not only 6× faster than original *Nanopolish*, but also consumed only ~15 GB RAM opposed to >100 GB by original *Nanopolish* (both run with 40 threads). Hence, it is evident that CPU optimisations alone can do significant improvements.

As per Fig. 12 for the whole methylation-calling process (including disk I/O), *f5c-gpu-opti* (only ABEA is performed on GPU) compared to *f5c-cpu-opti* was 1.7× faster on SoC, 1.5-1.6× on the lapH and <1.4× on HPC. On HPC the speedup was limited to <1.4× due to file I/O being the bottleneck.

When the execution time of *f5c-gpu-opti* for D_rapid_ is compared with original *Nanopolish*, it is ~4×, ~3× and ~6× faster on SoC, laptop and HPC, respectively. On HPC for D_ligation_, *f5c-gpu-opti* was ~9× faster.

Note that we used *Nanopolish* v0.9 for comparison as the re-engineering was done on this particular version. As we incorporated a number of CPU optimisations identified during the re-engineering into the subsequent version of *Nanopolish* (only those that did not require major code refactoring), latest *Nanopolish* v0.11 should be faster than v0.9 used in this paper.

## 6 Discussion

With the method discussed in this paper, the complete methylation calling of a human genome can now be performed on-the-fly (process in real-time while the nanopore sequencer is operating) on an embedded system (e.g., an SoC equipped with ARM processor and an NVIDIA GPU) as shown in Fig. 13 (four Oxford Nanopore MinION devices sequencing in parallel or a single Oxford Nanopore GridION, is capable of sequencing a human genome at an adequate coverage). *f5c* powered by GPU accelerated ABEA can process the output from the rest of the pipeline on a single NVIDIA TX2 SoC, at a speed of (>600 Kbases per second) to keep up with the sequencing output (~600 Kbases per second [22]) as shown in Fig. 13. Conversely, if the original *Nanopolish* was executed on the NVIDIA TX2 SoC, the processing speed is limited to ~256 Kbases per second. Our work will not only reduce the associated costs of Nanopore data processing and data transfer, but will also improve turnaround time of the final test outcome.

**Figure 13:**
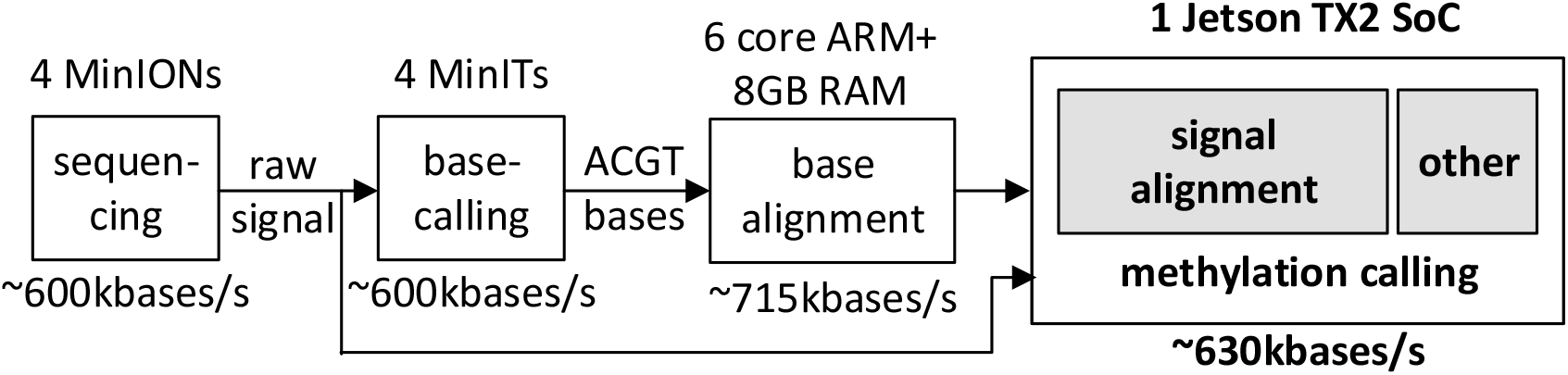
Human genome processing on-the-fly

In addition to embedded systems, our work benefits systems with or without GPU. Due to reduced peak memory usage, methylation calling can be performed on laptops with <16GB of RAM. Furthermore, post sequencing methylation calling execution on high performance computers also benefit from a significant speedup in processing.

A limitation of our implementation is that the parameter tuning cannot be performed automatically, which instead prompts the user when an un-optimal parameter is detected. This limitation is expected to be addressed in a future version by automatically tuning the parameters at run-time; or, by the use of pre-set parameter profiles for different types of datasets and/or computer systems.

## 7 Conclusion

Adaptive Banded Event Alignment algorithm is one of the key components in nanopore data analysis. Despite this algorithm being not embarrassingly parallel, we presented an approach that makes this algorithm efficiently execute on GPUs. The high variability of the read lengths was one of the main challenges, which was remedied through a number of memory optimisations and a heterogeneous processing strategy that uses both CPU and GPU. Our optimisations yield around 3-5× performance improvement on a CPU-GPU system when compared to a CPU. We incorporated the optimised Adaptive Banded Event Alignment algorithm into a methylation detection workflow and demonstrated that an embedded SoC equipped with an ARM processor (with six cores) and NVIDIA GPU (256 cores) is adequate to process data from a portable nanopore sequencer in real-time.

This work not only benefits embedded SoC, but also a wide range of systems equipped with GPUs from laptops to servers. The re-engineered version of the *Nanopolish* methylation detection module, *f5c* that employs the GPU accelerated Adaptive Banded Event Alignment was not only around 9× faster on an HPC, but also reduced the peak RAM by around 6× times. The source code of *f5c* is made available at https://github.com/hasindu2008/f5c.

## Supporting information

f5c-v0.2-beta-release

## Acknowledgements

We thank NVIDIA for providing the Jetson TX2 board (to University of New South Wales) and Tesla K40 GPU (to University of Peradeniya) through the GPU donation programme. We thank Dr. Roshan Ragel and Dr. Swarnalatha Radhakrishnan at University of Peradeniya who assisted the research. We thank our colleagues who provided support, especially, James Ferguson and Shaun Carswell at Garvan Institute of Medical Research; Thomas Daniell, Hassaan Saadat and Darshana Jayasinghe at University of New South Wales; Geesara Pratap at Innopolis University; and, Pim Schravendijk. We also thank the Data Intensive Computer Engineering (DICE) team at Data Sciences Platform, Garvan Institute of Medical Research.

## Author contributions

H.G. and S.P. conceived the study. H.G., C.W.L., H.S. and G.J. re-engineered, wrote, modified and optimised software. H.G. designed, developed and optimised the GPU implementation. H.G. and C.W.L. conducted the experiments and benchmarks. M.A.S., J.T.S and S.P. provided strategic oversight for the work. H.G, M.A.S. and S.P wrote and revised the manuscript. J.T.S. critically revised the manuscript. All authors read and approved the manuscript.

## Footnotes

1 these are typical values at present which may vary in the future

2 there can be other values in addition to mean and standard deviation, which are not required for our methylation calling

## References

[1] Simpson, J. T. et al. Detecting dna cytosine methylation using nanopore sequencing. Nature methods 14, 407 (2017).

[2] Liyanage, V. et al. Dna modifications: function and applications in normal and disease states. Biology 3, 670–723 (2014).

[3] Lewandowska, J. & Bartoszek, A. Dna methylation in cancer development, diagnosis and therapy—multiple opportunities for genotoxic agents to act as methylome disruptors or remediators. Mutagenesis 26, 475–487 (2011).

[4] Fraser, M. et al. Genomic hallmarks of localized, non-indolent prostate cancer. Nature 541, 359 (2017).

[5] Saxonov, S., Berg, P. & Brutlag, D. L. A genome-wide analysis of cpg dinucleotides in the human genome distinguishes two distinct classes of promoters. Proceedings of the National Academy of Sciences 103, 1412–1417 (2006).

[6] Bird, A. Dna methylation patterns and epigenetic memory. Genes & development 16, 6–21 (2002).

[7] Gonzalo, S. Epigenetic alterations in aging. Journal of applied physiology 109, 586–597 (2010).

[8] Lu, H., Giordano, F. & Ning, Z. Oxford nanopore minion sequencing and genome assembly. Genomics, proteomics & bioinformatics 14, 265–279 (2016).

[9] Rang, F. J., Kloosterman, W. P. & de Ridder, J. From squiggle to basepair: computational approaches for improving nanopore sequencing read accuracy. Genome biology 19, 90 (2018).

[10] Wick, R. R., Judd, L. M. & Holt, K. E. Performance of neural network basecalling tools for oxford nanopore sequencing. bioRxiv 543439 (2019).

[11] Li, H. Minimap2: pairwise alignment for nucleotide sequences. Bioinformatics bty191 (2018). URL http://dx.doi.org/10.1093/bioinformatics/bty191.

[12] Jain, M. et al. Nanopore sequencing and assembly of a human genome with ultra-long reads. Nature biotechnology 36, 338 (2018).

[13] Loman, N. J., Quick, J. & Simpson, J. T. A complete bacterial genome assembled de novo using only nanopore sequencing data. Nature methods 12, 733 (2015).

[14] Durbin, R., Eddy, S. R., Krogh, A. & Mitchison, G. Biological sequence analysis: probabilistic models of proteins and nucleic acids (Cambridge university press, 1998).

[15] Suzuki, H. & Kasahara, M. Introducing difference recurrence relations for faster semi-global alignment of long sequences. BMC bioinformatics 19, 45 (2018).

[16] David, M., Dursi, L. J., Yao, D., Boutros, P. C. & Simpson, J. T. Nanocall: an open source basecaller for oxford nanopore sequencing data. Bioinformatics 33, 49–55 (2016).

[17] NVIDIA. CUDA C Programming guide (2018). PG-02829-001_v10.0.

[18] NVIDIA. CUDA C best practices guide (2018). DG-05603-001_v10.0.

[19] Liu, Y., Maskell, D. L. & Schmidt, B. CUDASW++: optimizing Smith-Waterman sequence database searches for CUDA-enabled graphics processing units. BMC Research Notes 2, 73 (2009). URL http://bmcresnotes.biomedcentral.com/articles/10.1186/1756-0500-2-73.

[20] Liu, Y., Schmidt, B. & Maskell, D. L. CUDASW++2.0: enhanced Smith-Waterman protein database search on CUDA-enabled GPUs based on SIMT and virtualized SIMD abstractions. BMC Research Notes 3, 93 (2010). URL http://bmcresnotes.biomedcentral.com/articles/10.1186/1756-0500-3-93.

[21] Liu, Y., Wirawan, A. & Schmidt, B. CUDASW++ 3.0: accelerating Smith-Waterman protein database search by coupling CPU and GPU SIMD instructions. BMC Bioinformatics 14, 117 (2013). URL http://bmcbioinformatics.biomedcentral.com/articles/10.1186/1471-2105-14-117.

[22] Technologies, O. N. Minit is out – an analysis and device control accessory to enable powerful, real-time dna/rna sequencing by anyone, anywhere (2018). URL https://nanoporetech.com/about-us/news/minit-launch.

[23] Gamaarachchi, H., Parameswaran, S. & Smith, M. A. Featherweight long read alignment using partitioned reference indexes. Scientific Reports 9, 4318 (2019).

[24] Huismann, I., Lieber, M., Stiller, J. & Fröhlich, J. Load balancing for cpu-gpu coupling in computational fluid dynamics. In International Conference on Parallel Processing and Applied Mathematics, 337–347 (Springer, 2017).

[25] Lang, J. & Rünger, G. Dynamic distribution of workload between cpu and gpu for a parallel conjugate gradient method in an adaptive fem. Procedia Computer Science 18, 299–308 (2013).

[26] Technologies, O. N. Ligation sequencing kit 1d or rapid sequencing kit (2017). URL https://store.nanoporetech.com/media/Ligation_Sequencing_Kit_1D_or_Rapid_Sequencing_Kit_v5_Feb2017.pdf.

[27] Simpson, J. Stats and analysis (2017). URL https://nanopolish.readthedocs.io/en/latest/quickstart_call_methylation.html.

[28] NVIDIA. PROFILER USER’S GUIDE (2019). DU-05982-001_v10.1.

[29] Chase, R. How to configure the linux out-of-memory killer (2013). URL https://www.oracle.com/technical-resources/articles/it-infrastructure/dev-oom-killer.html.

