## Supplementary material for "GPU Accelerated Adaptive Banded Event Alignment for Rapid Comparative Nanopore Signal Analysis": f5c-v0.2-beta-release: configure

#! /bin/sh
# Guess values for system-dependent variables and create Makefiles.
# Generated by GNU Autoconf 2.69 for f5c 1.0.
#
# Report bugs to <>.
#
#
# Copyright (C) 1992-1996, 1998-2012 Free Software Foundation, Inc.
#
#
# This configure script is free software; the Free Software Foundation
# gives unlimited permission to copy, distribute and modify it.
## -------------------- ##
## M4sh Initialization. ##
## -------------------- ##

# Be more Bourne compatible
DUALCASE=1; export DUALCASE # for MKS sh
if test -n "${ZSH_VERSION+set}" && (emulate sh) >/dev/null 2>&1; then :
  emulate sh
  NULLCMD=:
  # Pre-4.2 versions of Zsh do word splitting on ${1+"$@"}, which
  # is contrary to our usage.  Disable this feature.
  alias -g '${1+"$@"}'='"$@"'
  setopt NO_GLOB_SUBST
else
  case `(set -o) 2>/dev/null` in #(
  *posix*) :
    set -o posix ;; #(
  *) :
     ;;
esac
fi


as_nl='
'
export as_nl
# Printing a long string crashes Solaris 7 /usr/bin/printf.
as_echo='\\\\\\\\\\\\\\\\\\\\\\\\\\\\\\\\\\\\\\\\\\\\\\\\\\\\\\\\\\\\\\\\\\\\\\\\\\\\\\\\\\\\\\\\\\\\\\\\\\\\\\\'
as_echo=$as_echo$as_echo$as_echo$as_echo$as_echo
as_echo=$as_echo$as_echo$as_echo$as_echo$as_echo$as_echo
# Prefer a ksh shell builtin over an external printf program on Solaris,
# but without wasting forks for bash or zsh.
if test -z "$BASH_VERSION$ZSH_VERSION" \
    && (test "X`print -r -- $as_echo`" = "X$as_echo") 2>/dev/null; then
  as_echo='print -r --'
  as_echo_n='print -rn --'
elif (test "X`printf %s $as_echo`" = "X$as_echo") 2>/dev/null; then
  as_echo='printf %s\n'
  as_echo_n='printf %s'
else
  if test "X`(/usr/ucb/echo -n -n $as_echo) 2>/dev/null`" = "X-n $as_echo"; then
    as_echo_body='eval /usr/ucb/echo -n "$1$as_nl"'
    as_echo_n='/usr/ucb/echo -n'
  else
    as_echo_body='eval expr "X$1" : "X\\(.*\\)"'
    as_echo_n_body='eval
      arg=$1;
      case $arg in #(
      *"$as_nl"*)
	expr "X$arg" : "X\\(.*\\)$as_nl";
	arg=`expr "X$arg" : ".*$as_nl\\(.*\\)"`;;
      esac;
      expr "X$arg" : "X\\(.*\\)" | tr -d "$as_nl"
    '
    export as_echo_n_body
    as_echo_n='sh -c $as_echo_n_body as_echo'
  fi
  export as_echo_body
  as_echo='sh -c $as_echo_body as_echo'
fi

# The user is always right.
if test "${PATH_SEPARATOR+set}" != set; then
  PATH_SEPARATOR=:
  (PATH='/bin;/bin'; FPATH=$PATH; sh -c :) >/dev/null 2>&1 && {
    (PATH='/bin:/bin'; FPATH=$PATH; sh -c :) >/dev/null 2>&1 ||
      PATH_SEPARATOR=';'
  }
fi


# IFS
# We need space, tab and new line, in precisely that order.  Quoting is
# there to prevent editors from complaining about space-tab.
# (If _AS_PATH_WALK were called with IFS unset, it would disable word
# splitting by setting IFS to empty value.)
IFS=" ""	$as_nl"

# Find who we are.  Look in the path if we contain no directory separator.
as_myself=
case $0 in #((
  *[\\/]* ) as_myself=$0 ;;
  *) as_save_IFS=$IFS; IFS=$PATH_SEPARATOR
for as_dir in $PATH
do
  IFS=$as_save_IFS
  test -z "$as_dir" && as_dir=.
    test -r "$as_dir/$0" && as_myself=$as_dir/$0 && break
  done
IFS=$as_save_IFS

     ;;
esac
# We did not find ourselves, most probably we were run as `sh COMMAND'
# in which case we are not to be found in the path.
if test "x$as_myself" = x; then
  as_myself=$0
fi
if test ! -f "$as_myself"; then
  $as_echo "$as_myself: error: cannot find myself; rerun with an absolute file name" >&2
  exit 1
fi

# Unset variables that we do not need and which cause bugs (e.g. in
# pre-3.0 UWIN ksh).  But do not cause bugs in bash 2.01; the "|| exit 1"
# suppresses any "Segmentation fault" message there.  '((' could
# trigger a bug in pdksh 5.2.14.
for as_var in BASH_ENV ENV MAIL MAILPATH
do eval test x\${$as_var+set} = xset \
  && ( (unset $as_var) || exit 1) >/dev/null 2>&1 && unset $as_var || :
done
PS1='$ '
PS2='> '
PS4='+ '

# NLS nuisances.
LC_ALL=C
export LC_ALL
LANGUAGE=C
export LANGUAGE

# CDPATH.
(unset CDPATH) >/dev/null 2>&1 && unset CDPATH

# Use a proper internal environment variable to ensure we don't fall
  # into an infinite loop, continuously re-executing ourselves.
  if test x"${_as_can_reexec}" != xno && test "x$CONFIG_SHELL" != x; then
    _as_can_reexec=no; export _as_can_reexec;
    # We cannot yet assume a decent shell, so we have to provide a
# neutralization value for shells without unset; and this also
# works around shells that cannot unset nonexistent variables.
# Preserve -v and -x to the replacement shell.
BASH_ENV=/dev/null
ENV=/dev/null
(unset BASH_ENV) >/dev/null 2>&1 && unset BASH_ENV ENV
case $- in # ((((
  *v*x* | *x*v* ) as_opts=-vx ;;
  *v* ) as_opts=-v ;;
  *x* ) as_opts=-x ;;
  * ) as_opts= ;;
esac
exec $CONFIG_SHELL $as_opts "$as_myself" ${1+"$@"}
# Admittedly, this is quite paranoid, since all the known shells bail
# out after a failed `exec'.
$as_echo "$0: could not re-execute with $CONFIG_SHELL" >&2
as_fn_exit 255
  fi
  # We don't want this to propagate to other subprocesses.
          { _as_can_reexec=; unset _as_can_reexec;}
if test "x$CONFIG_SHELL" = x; then
  as_bourne_compatible="if test -n \"\${ZSH_VERSION+set}\" && (emulate sh) >/dev/null 2>&1; then :
  emulate sh
  NULLCMD=:
  # Pre-4.2 versions of Zsh do word splitting on \${1+\"\$@\"}, which
  # is contrary to our usage.  Disable this feature.
  alias -g '\${1+\"\$@\"}'='\"\$@\"'
  setopt NO_GLOB_SUBST
else
  case \`(set -o) 2>/dev/null\` in #(
  *posix*) :
    set -o posix ;; #(
  *) :
     ;;
esac
fi
"
  as_required="as_fn_return () { (exit \$1); }
as_fn_success () { as_fn_return 0; }
as_fn_failure () { as_fn_return 1; }
as_fn_ret_success () { return 0; }
as_fn_ret_failure () { return 1; }

exitcode=0
as_fn_success || { exitcode=1; echo as_fn_success failed.; }
as_fn_failure && { exitcode=1; echo as_fn_failure succeeded.; }
as_fn_ret_success || { exitcode=1; echo as_fn_ret_success failed.; }
as_fn_ret_failure && { exitcode=1; echo as_fn_ret_failure succeeded.; }
if ( set x; as_fn_ret_success y && test x = \"\$1\" ); then :

else
  exitcode=1; echo positional parameters were not saved.
fi
test x\$exitcode = x0 || exit 1
test -x / || exit 1"
  as_suggested="  as_lineno_1=";as_suggested=$as_suggested$LINENO;as_suggested=$as_suggested" as_lineno_1a=\$LINENO
  as_lineno_2=";as_suggested=$as_suggested$LINENO;as_suggested=$as_suggested" as_lineno_2a=\$LINENO
  eval 'test \"x\$as_lineno_1'\$as_run'\" != \"x\$as_lineno_2'\$as_run'\" &&
  test \"x\`expr \$as_lineno_1'\$as_run' + 1\`\" = \"x\$as_lineno_2'\$as_run'\"' || exit 1
test \$(( 1 + 1 )) = 2 || exit 1"
  if (eval "$as_required") 2>/dev/null; then :
  as_have_required=yes
else
  as_have_required=no
fi
  if test x$as_have_required = xyes && (eval "$as_suggested") 2>/dev/null; then :

else
  as_save_IFS=$IFS; IFS=$PATH_SEPARATOR
as_found=false
for as_dir in /bin$PATH_SEPARATOR/usr/bin$PATH_SEPARATOR$PATH
do
  IFS=$as_save_IFS
  test -z "$as_dir" && as_dir=.
  as_found=:
  case $as_dir in #(
	 /*)
	   for as_base in sh bash ksh sh5; do
	     # Try only shells that exist, to save several forks.
	     as_shell=$as_dir/$as_base
	     if { test -f "$as_shell" || test -f "$as_shell.exe"; } &&
		    { $as_echo "$as_bourne_compatible""$as_required" | as_run=a "$as_shell"; } 2>/dev/null; then :
  CONFIG_SHELL=$as_shell as_have_required=yes
		   if { $as_echo "$as_bourne_compatible""$as_suggested" | as_run=a "$as_shell"; } 2>/dev/null; then :
  break 2
fi
fi
	   done;;
       esac
  as_found=false
done
$as_found || { if { test -f "$SHELL" || test -f "$SHELL.exe"; } &&
	      { $as_echo "$as_bourne_compatible""$as_required" | as_run=a "$SHELL"; } 2>/dev/null; then :
  CONFIG_SHELL=$SHELL as_have_required=yes
fi; }
IFS=$as_save_IFS


      if test "x$CONFIG_SHELL" != x; then :
  export CONFIG_SHELL
             # We cannot yet assume a decent shell, so we have to provide a
# neutralization value for shells without unset; and this also
# works around shells that cannot unset nonexistent variables.
# Preserve -v and -x to the replacement shell.
BASH_ENV=/dev/null
ENV=/dev/null
(unset BASH_ENV) >/dev/null 2>&1 && unset BASH_ENV ENV
case $- in # ((((
  *v*x* | *x*v* ) as_opts=-vx ;;
  *v* ) as_opts=-v ;;
  *x* ) as_opts=-x ;;
  * ) as_opts= ;;
esac
exec $CONFIG_SHELL $as_opts "$as_myself" ${1+"$@"}
# Admittedly, this is quite paranoid, since all the known shells bail
# out after a failed `exec'.
$as_echo "$0: could not re-execute with $CONFIG_SHELL" >&2
exit 255
fi

    if test x$as_have_required = xno; then :
  $as_echo "$0: This script requires a shell more modern than all"
  $as_echo "$0: the shells that I found on your system."
  if test x${ZSH_VERSION+set} = xset ; then
    $as_echo "$0: In particular, zsh $ZSH_VERSION has bugs and should"
    $as_echo "$0: be upgraded to zsh 4.3.4 or later."
  else
    $as_echo "$0: Please tell and
$0: about your system, including any
$0: error possibly output before this message. Then install
$0: a modern shell, or manually run the script under such a
$0: shell if you do have one."
  fi
  exit 1
fi
fi
fi
SHELL=${CONFIG_SHELL-/bin/sh}
export SHELL
# Unset more variables known to interfere with behavior of common tools.
CLICOLOR_FORCE= GREP_OPTIONS=
unset CLICOLOR_FORCE GREP_OPTIONS

## --------------------- ##
## M4sh Shell Functions. ##
## --------------------- ##
# as_fn_unset VAR
# ---------------
# Portably unset VAR.
as_fn_unset ()
{
  { eval $1=; unset $1;}
}
as_unset=as_fn_unset

# as_fn_set_status STATUS
# -----------------------
# Set $? to STATUS, without forking.
as_fn_set_status ()
{
  return $1
} # as_fn_set_status

# as_fn_exit STATUS
# -----------------
# Exit the shell with STATUS, even in a "trap 0" or "set -e" context.
as_fn_exit ()
{
  set +e
  as_fn_set_status $1
  exit $1
} # as_fn_exit

# as_fn_mkdir_p
# -------------
# Create "$as_dir" as a directory, including parents if necessary.
as_fn_mkdir_p ()
{

  case $as_dir in #(
  -*) as_dir=./$as_dir;;
  esac
  test -d "$as_dir" || eval $as_mkdir_p || {
    as_dirs=
    while :; do
      case $as_dir in #(
      *\'*) as_qdir=`$as_echo "$as_dir" | sed "s/'/'\\\\\\\\''/g"`;; #'(
      *) as_qdir=$as_dir;;
      esac
      as_dirs="'$as_qdir' $as_dirs"
      as_dir=`$as_dirname -- "$as_dir" ||
$as_expr X"$as_dir" : 'X\(.*[^/]\)//*[^/][^/]*/*$' \| \
	 X"$as_dir" : 'X\(//\)[^/]' \| \
	 X"$as_dir" : 'X\(//\)$' \| \
	 X"$as_dir" : 'X\(/\)' \| . 2>/dev/null ||
$as_echo X"$as_dir" |
    sed '/^X\(.*[^/]\)\/\/*[^/][^/]*\/*$/{
	    s//\1/
	    q
	  }
	  /^X\(\/\/\)[^/].*/{
	    s//\1/
	    q
	  }
	  /^X\(\/\/\)$/{
	    s//\1/
	    q
	  }
	  /^X\(\/\).*/{
	    s//\1/
	    q
	  }
	  s/.*/./; q'`
      test -d "$as_dir" && break
    done
    test -z "$as_dirs" || eval "mkdir $as_dirs"
  } || test -d "$as_dir" || as_fn_error $? "cannot create directory $as_dir"


} # as_fn_mkdir_p

# as_fn_executable_p FILE
# -----------------------
# Test if FILE is an executable regular file.
as_fn_executable_p ()
{
  test -f "$1" && test -x "$1"
} # as_fn_executable_p
# as_fn_append VAR VALUE
# ----------------------
# Append the text in VALUE to the end of the definition contained in VAR. Take
# advantage of any shell optimizations that allow amortized linear growth over
# repeated appends, instead of the typical quadratic growth present in naive
# implementations.
if (eval "as_var=1; as_var+=2; test x\$as_var = x12") 2>/dev/null; then :
  eval 'as_fn_append ()
  {
    eval $1+=\$2
  }'
else
  as_fn_append ()
  {
    eval $1=\$$1\$2
  }
fi # as_fn_append

# as_fn_arith ARG...
# ------------------
# Perform arithmetic evaluation on the ARGs, and store the result in the
# global $as_val. Take advantage of shells that can avoid forks. The arguments
# must be portable across $(()) and expr.
if (eval "test \$(( 1 + 1 )) = 2") 2>/dev/null; then :
  eval 'as_fn_arith ()
  {
    as_val=$(( $* ))
  }'
else
  as_fn_arith ()
  {
    as_val=`expr "$@" || test $? -eq 1`
  }
fi # as_fn_arith


# as_fn_error STATUS ERROR [LINENO LOG_FD]
# ----------------------------------------
# Output "`basename $0`: error: ERROR" to stderr. If LINENO and LOG_FD are
# provided, also output the error to LOG_FD, referencing LINENO. Then exit the
# script with STATUS, using 1 if that was 0.
as_fn_error ()
{
  as_status=$1; test $as_status -eq 0 && as_status=1
  if test "$4"; then
    as_lineno=${as_lineno-"$3"} as_lineno_stack=as_lineno_stack=$as_lineno_stack
    $as_echo "$as_me:${as_lineno-$LINENO}: error: $2" >&$4
  fi
  $as_echo "$as_me: error: $2" >&2
  as_fn_exit $as_status
} # as_fn_error

if expr a : '\(a\)' >/dev/null 2>&1 &&
   test "X`expr 00001 : '.*\(...\)'`" = X001; then
  as_expr=expr
else
  as_expr=false
fi

if (basename -- /) >/dev/null 2>&1 && test "X`basename -- / 2>&1`" = "X/"; then
  as_basename=basename
else
  as_basename=false
fi

if (as_dir=`dirname -- /` && test "X$as_dir" = X/) >/dev/null 2>&1; then
  as_dirname=dirname
else
  as_dirname=false
fi

as_me=`$as_basename -- "$0" ||
$as_expr X/"$0" : '.*/\([^/][^/]*\)/*$' \| \
	 X"$0" : 'X\(//\)$' \| \
	 X"$0" : 'X\(/\)' \| . 2>/dev/null ||
$as_echo X/"$0" |
    sed '/^.*\/\([^/][^/]*\)\/*$/{
	    s//\1/
	    q
	  }
	  /^X\/\(\/\/\)$/{
	    s//\1/
	    q
	  }
	  /^X\/\(\/\).*/{
	    s//\1/
	    q
	  }
	  s/.*/./; q'`

# Avoid depending upon Character Ranges.
as_cr_letters='abcdefghijklmnopqrstuvwxyz'
as_cr_LETTERS='ABCDEFGHIJKLMNOPQRSTUVWXYZ'
as_cr_Letters=$as_cr_letters$as_cr_LETTERS
as_cr_digits='0123456789'
as_cr_alnum=$as_cr_Letters$as_cr_digits


  as_lineno_1=$LINENO as_lineno_1a=$LINENO
  as_lineno_2=$LINENO as_lineno_2a=$LINENO
  eval 'test "x$as_lineno_1'$as_run'" != "x$as_lineno_2'$as_run'" &&
  test "x`expr $as_lineno_1'$as_run' + 1`" = "x$as_lineno_2'$as_run'"' || {
  # Blame Lee E. McMahon (1931-1989) for sed's syntax.  :-)
  sed -n '
    p
    /[$]LINENO/=
  ' <$as_myself |
    sed '
      s/[$]LINENO.*/&-/
      t lineno
      b
      :lineno
      N
      :loop
      s/[$]LINENO\([^'$as_cr_alnum'_].*\n\)\(.*\)/\2\1\2/
      t loop
      s/-\n.*//
    ' >$as_me.lineno &&
  chmod +x "$as_me.lineno" ||
    { $as_echo "$as_me: error: cannot create $as_me.lineno; rerun with a POSIX shell" >&2; as_fn_exit 1; }

  # If we had to re-execute with $CONFIG_SHELL, we're ensured to have
  # already done that, so ensure we don't try to do so again and fall
  # in an infinite loop.  This has already happened in practice.
  _as_can_reexec=no; export _as_can_reexec
  # Don't try to exec as it changes $[0], causing all sort of problems
  # (the dirname of $[0] is not the place where we might find the
  # original and so on.  Autoconf is especially sensitive to this).
  . "./$as_me.lineno"
  # Exit status is that of the last command.
  exit
}

ECHO_C= ECHO_N= ECHO_T=
case `echo -n x` in #(((((
-n*)
  case `echo 'xy\c'` in
  *c*) ECHO_T='	';;	# ECHO_T is single tab character.
  xy)  ECHO_C='\c';;
  *)   echo `echo ksh88 bug on AIX 6.1` > /dev/null
       ECHO_T='	';;
  esac;;
*)
  ECHO_N='-n';;
esac

rm -f conf$$ conf$$.exe conf$$.file
if test -d conf$$.dir; then
  rm -f conf$$.dir/conf$$.file
else
  rm -f conf$$.dir
  mkdir conf$$.dir 2>/dev/null
fi
if (echo >conf$$.file) 2>/dev/null; then
  if ln -s conf$$.file conf$$ 2>/dev/null; then
    as_ln_s='ln -s'
    # ... but there are two gotchas:
    # 1) On MSYS, both `ln -s file dir' and `ln file dir' fail.
    # 2) DJGPP < 2.04 has no symlinks; `ln -s' creates a wrapper executable.
    # In both cases, we have to default to `cp -pR'.
    ln -s conf$$.file conf$$.dir 2>/dev/null && test ! -f conf$$.exe ||
      as_ln_s='cp -pR'
  elif ln conf$$.file conf$$ 2>/dev/null; then
    as_ln_s=ln
  else
    as_ln_s='cp -pR'
  fi
else
  as_ln_s='cp -pR'
fi
rm -f conf$$ conf$$.exe conf$$.dir/conf$$.file conf$$.file
rmdir conf$$.dir 2>/dev/null

if mkdir -p . 2>/dev/null; then
  as_mkdir_p='mkdir -p "$as_dir"'
else
  test -d ./-p && rmdir ./-p
  as_mkdir_p=false
fi

as_test_x='test -x'
as_executable_p=as_fn_executable_p

# Sed expression to map a string onto a valid CPP name.
as_tr_cpp="eval sed 'y%*$as_cr_letters%P$as_cr_LETTERS%;s%[^_$as_cr_alnum]%_%g'"

# Sed expression to map a string onto a valid variable name.
as_tr_sh="eval sed 'y%*+%pp%;s%[^_$as_cr_alnum]%_%g'"


test -n "$DJDIR" || exec 7<&0 </dev/null
exec 6>&1

# Name of the host.
# hostname on some systems (SVR3.2, old GNU/Linux) returns a bogus exit status,
# so uname gets run too.
ac_hostname=`(hostname || uname -n) 2>/dev/null | sed 1q`

#
# Initializations.
#
ac_default_prefix=/usr/local
ac_clean_files=
ac_config_libobj_dir=.
LIBOBJS=
cross_compiling=no
subdirs=
MFLAGS=
MAKEFLAGS=

# Identity of this package.
PACKAGE_NAME='f5c'
PACKAGE_TARNAME='f5c'
PACKAGE_VERSION='1.0'
PACKAGE_STRING='f5c 1.0'
PACKAGE_BUGREPORT=''
PACKAGE_URL=''

ac_unique_file="src/f5c.c"
# Factoring default headers for most tests.
ac_includes_default="\
#include <stdio.h>
#ifdef HAVE_SYS_TYPES_H
# include <sys/types.h>
#endif
#ifdef HAVE_SYS_STAT_H
# include <sys/stat.h>
#endif
#ifdef STDC_HEADERS
# include <stdlib.h>
# include <stddef.h>
#else
# ifdef HAVE_STDLIB_H
#  include <stdlib.h>
# endif
#endif
#ifdef HAVE_STRING_H
# if !defined STDC_HEADERS && defined HAVE_MEMORY_H
#  include <memory.h>
# endif
# include <string.h>
#endif
#ifdef HAVE_STRINGS_H
# include <strings.h>
#endif
#ifdef HAVE_INTTYPES_H
# include <inttypes.h>
#endif
#ifdef HAVE_STDINT_H
# include <stdint.h>
#endif
#ifdef HAVE_UNISTD_H
# include <unistd.h>
#endif"

ac_subst_vars='LTLIBOBJS
hts_tar_install
hdf_tar_install
locallibhdf5
locallibhts
LIBOBJS
EGREP
GREP
CPP
ac_ct_CXX
CXXFLAGS
CXX
OBJEXT
EXEEXT
ac_ct_CC
CPPFLAGS
LDFLAGS
CFLAGS
CC
target_alias
host_alias
build_alias
LIBS
ECHO_T
ECHO_N
ECHO_C
DEFS
mandir
localedir
libdir
psdir
pdfdir
dvidir
htmldir
infodir
docdir
oldincludedir
includedir
localstatedir
sharedstatedir
sysconfdir
datadir
datarootdir
libexecdir
sbindir
bindir
program_transform_name
prefix
exec_prefix
PACKAGE_URL
PACKAGE_BUGREPORT
PACKAGE_STRING
PACKAGE_VERSION
PACKAGE_TARNAME
PACKAGE_NAME
PATH_SEPARATOR
SHELL'
ac_subst_files=''
ac_user_opts='
enable_option_checking
enable_systemhts
enable_localhdf5
enable_largefile
'
      ac_precious_vars='build_alias
host_alias
target_alias
CC
CFLAGS
LDFLAGS
LIBS
CPPFLAGS
CXX
CXXFLAGS
CCC
CPP'


# Initialize some variables set by options.
ac_init_help=
ac_init_version=false
ac_unrecognized_opts=
ac_unrecognized_sep=
# The variables have the same names as the options, with
# dashes changed to underlines.
cache_file=/dev/null
exec_prefix=NONE
no_create=
no_recursion=
prefix=NONE
program_prefix=NONE
program_suffix=NONE
program_transform_name=s,x,x,
silent=
site=
srcdir=
verbose=
x_includes=NONE
x_libraries=NONE

# Installation directory options.
# These are left unexpanded so users can "make install exec_prefix=/foo"
# and all the variables that are supposed to be based on exec_prefix
# by default will actually change.
# Use braces instead of parens because sh, perl, etc. also accept them.
# (The list follows the same order as the GNU Coding Standards.)
bindir='${exec_prefix}/bin'
sbindir='${exec_prefix}/sbin'
libexecdir='${exec_prefix}/libexec'
datarootdir='${prefix}/share'
datadir='${datarootdir}'
sysconfdir='${prefix}/etc'
sharedstatedir='${prefix}/com'
localstatedir='${prefix}/var'
includedir='${prefix}/include'
oldincludedir='/usr/include'
docdir='${datarootdir}/doc/${PACKAGE_TARNAME}'
infodir='${datarootdir}/info'
htmldir='${docdir}'
dvidir='${docdir}'
pdfdir='${docdir}'
psdir='${docdir}'
libdir='${exec_prefix}/lib'
localedir='${datarootdir}/locale'
mandir='${datarootdir}/man'

ac_prev=
ac_dashdash=
for ac_option
do
  # If the previous option needs an argument, assign it.
  if test -n "$ac_prev"; then
    eval $ac_prev=\$ac_option
    ac_prev=
    continue
  fi

  case $ac_option in
  *=?*) ac_optarg=`expr "X$ac_option" : '[^=]*=\(.*\)'` ;;
  *=)   ac_optarg= ;;
  *)    ac_optarg=yes ;;
  esac

  # Accept the important Cygnus configure options, so we can diagnose typos.

  case $ac_dashdash$ac_option in
  --)
    ac_dashdash=yes ;;

  -bindir | --bindir | --bindi | --bind | --bin | --bi)
    ac_prev=bindir ;;
  -bindir=* | --bindir=* | --bindi=* | --bind=* | --bin=* | --bi=*)
    bindir=$ac_optarg ;;

  -build | --build | --buil | --bui | --bu)
    ac_prev=build_alias ;;
  -build=* | --build=* | --buil=* | --bui=* | --bu=*)
    build_alias=$ac_optarg ;;

  -cache-file | --cache-file | --cache-fil | --cache-fi \
  | --cache-f | --cache- | --cache | --cach | --cac | --ca | --c)
    ac_prev=cache_file ;;
  -cache-file=* | --cache-file=* | --cache-fil=* | --cache-fi=* \
  | --cache-f=* | --cache-=* | --cache=* | --cach=* | --cac=* | --ca=* | --c=*)
    cache_file=$ac_optarg ;;

  --config-cache | -C)
    cache_file=config.cache ;;

  -datadir | --datadir | --datadi | --datad)
    ac_prev=datadir ;;
  -datadir=* | --datadir=* | --datadi=* | --datad=*)
    datadir=$ac_optarg ;;

  -datarootdir | --datarootdir | --datarootdi | --datarootd | --dataroot \
  | --dataroo | --dataro | --datar)
    ac_prev=datarootdir ;;
  -datarootdir=* | --datarootdir=* | --datarootdi=* | --datarootd=* \
  | --dataroot=* | --dataroo=* | --dataro=* | --datar=*)
    datarootdir=$ac_optarg ;;

  -disable-* | --disable-*)
    ac_useropt=`expr "x$ac_option" : 'x-*disable-\(.*\)'`
    # Reject names that are not valid shell variable names.
    expr "x$ac_useropt" : ".*[^-+._$as_cr_alnum]" >/dev/null &&
      as_fn_error $? "invalid feature name: $ac_useropt"
    ac_useropt_orig=$ac_useropt
    ac_useropt=`$as_echo "$ac_useropt" | sed 's/[-+.]/_/g'`
    case $ac_user_opts in
      *"
"enable_$ac_useropt"
"*) ;;
      *) ac_unrecognized_opts="$ac_unrecognized_opts$ac_unrecognized_sep--disable-$ac_useropt_orig"
	 ac_unrecognized_sep=', ';;
    esac
    eval enable_$ac_useropt=no ;;

  -docdir | --docdir | --docdi | --doc | --do)
    ac_prev=docdir ;;
  -docdir=* | --docdir=* | --docdi=* | --doc=* | --do=*)
    docdir=$ac_optarg ;;

  -dvidir | --dvidir | --dvidi | --dvid | --dvi | --dv)
    ac_prev=dvidir ;;
  -dvidir=* | --dvidir=* | --dvidi=* | --dvid=* | --dvi=* | --dv=*)
    dvidir=$ac_optarg ;;

  -enable-* | --enable-*)
    ac_useropt=`expr "x$ac_option" : 'x-*enable-\([^=]*\)'`
    # Reject names that are not valid shell variable names.
    expr "x$ac_useropt" : ".*[^-+._$as_cr_alnum]" >/dev/null &&
      as_fn_error $? "invalid feature name: $ac_useropt"
    ac_useropt_orig=$ac_useropt
    ac_useropt=`$as_echo "$ac_useropt" | sed 's/[-+.]/_/g'`
    case $ac_user_opts in
      *"
"enable_$ac_useropt"
"*) ;;
      *) ac_unrecognized_opts="$ac_unrecognized_opts$ac_unrecognized_sep--enable-$ac_useropt_orig"
	 ac_unrecognized_sep=', ';;
    esac
    eval enable_$ac_useropt=\$ac_optarg ;;

  -exec-prefix | --exec_prefix | --exec-prefix | --exec-prefi \
  | --exec-pref | --exec-pre | --exec-pr | --exec-p | --exec- \
  | --exec | --exe | --ex)
    ac_prev=exec_prefix ;;
  -exec-prefix=* | --exec_prefix=* | --exec-prefix=* | --exec-prefi=* \
  | --exec-pref=* | --exec-pre=* | --exec-pr=* | --exec-p=* | --exec-=* \
  | --exec=* | --exe=* | --ex=*)
    exec_prefix=$ac_optarg ;;

  -gas | --gas | --ga | --g)
    # Obsolete; use --with-gas.
    with_gas=yes ;;

  -help | --help | --hel | --he | -h)
    ac_init_help=long ;;
  -help=r* | --help=r* | --hel=r* | --he=r* | -hr*)
    ac_init_help=recursive ;;
  -help=s* | --help=s* | --hel=s* | --he=s* | -hs*)
    ac_init_help=short ;;

  -host | --host | --hos | --ho)
    ac_prev=host_alias ;;
  -host=* | --host=* | --hos=* | --ho=*)
    host_alias=$ac_optarg ;;

  -htmldir | --htmldir | --htmldi | --htmld | --html | --htm | --ht)
    ac_prev=htmldir ;;
  -htmldir=* | --htmldir=* | --htmldi=* | --htmld=* | --html=* | --htm=* \
  | --ht=*)
    htmldir=$ac_optarg ;;

  -includedir | --includedir | --includedi | --included | --include \
  | --includ | --inclu | --incl | --inc)
    ac_prev=includedir ;;
  -includedir=* | --includedir=* | --includedi=* | --included=* | --include=* \
  | --includ=* | --inclu=* | --incl=* | --inc=*)
    includedir=$ac_optarg ;;

  -infodir | --infodir | --infodi | --infod | --info | --inf)
    ac_prev=infodir ;;
  -infodir=* | --infodir=* | --infodi=* | --infod=* | --info=* | --inf=*)
    infodir=$ac_optarg ;;

  -libdir | --libdir | --libdi | --libd)
    ac_prev=libdir ;;
  -libdir=* | --libdir=* | --libdi=* | --libd=*)
    libdir=$ac_optarg ;;

  -libexecdir | --libexecdir | --libexecdi | --libexecd | --libexec \
  | --libexe | --libex | --libe)
    ac_prev=libexecdir ;;
  -libexecdir=* | --libexecdir=* | --libexecdi=* | --libexecd=* | --libexec=* \
  | --libexe=* | --libex=* | --libe=*)
    libexecdir=$ac_optarg ;;

  -localedir | --localedir | --localedi | --localed | --locale)
    ac_prev=localedir ;;
  -localedir=* | --localedir=* | --localedi=* | --localed=* | --locale=*)
    localedir=$ac_optarg ;;

  -localstatedir | --localstatedir | --localstatedi | --localstated \
  | --localstate | --localstat | --localsta | --localst | --locals)
    ac_prev=localstatedir ;;
  -localstatedir=* | --localstatedir=* | --localstatedi=* | --localstated=* \
  | --localstate=* | --localstat=* | --localsta=* | --localst=* | --locals=*)
    localstatedir=$ac_optarg ;;

  -mandir | --mandir | --mandi | --mand | --man | --ma | --m)
    ac_prev=mandir ;;
  -mandir=* | --mandir=* | --mandi=* | --mand=* | --man=* | --ma=* | --m=*)
    mandir=$ac_optarg ;;

  -nfp | --nfp | --nf)
    # Obsolete; use --without-fp.
    with_fp=no ;;

  -no-create | --no-create | --no-creat | --no-crea | --no-cre \
  | --no-cr | --no-c | -n)
    no_create=yes ;;

  -no-recursion | --no-recursion | --no-recursio | --no-recursi \
  | --no-recurs | --no-recur | --no-recu | --no-rec | --no-re | --no-r)
    no_recursion=yes ;;

  -oldincludedir | --oldincludedir | --oldincludedi | --oldincluded \
  | --oldinclude | --oldinclud | --oldinclu | --oldincl | --oldinc \
  | --oldin | --oldi | --old | --ol | --o)
    ac_prev=oldincludedir ;;
  -oldincludedir=* | --oldincludedir=* | --oldincludedi=* | --oldincluded=* \
  | --oldinclude=* | --oldinclud=* | --oldinclu=* | --oldincl=* | --oldinc=* \
  | --oldin=* | --oldi=* | --old=* | --ol=* | --o=*)
    oldincludedir=$ac_optarg ;;

  -prefix | --prefix | --prefi | --pref | --pre | --pr | --p)
    ac_prev=prefix ;;
  -prefix=* | --prefix=* | --prefi=* | --pref=* | --pre=* | --pr=* | --p=*)
    prefix=$ac_optarg ;;

  -program-prefix | --program-prefix | --program-prefi | --program-pref \
  | --program-pre | --program-pr | --program-p)
    ac_prev=program_prefix ;;
  -program-prefix=* | --program-prefix=* | --program-prefi=* \
  | --program-pref=* | --program-pre=* | --program-pr=* | --program-p=*)
    program_prefix=$ac_optarg ;;

  -program-suffix | --program-suffix | --program-suffi | --program-suff \
  | --program-suf | --program-su | --program-s)
    ac_prev=program_suffix ;;
  -program-suffix=* | --program-suffix=* | --program-suffi=* \
  | --program-suff=* | --program-suf=* | --program-su=* | --program-s=*)
    program_suffix=$ac_optarg ;;

  -program-transform-name | --program-transform-name \
  | --program-transform-nam | --program-transform-na \
  | --program-transform-n | --program-transform- \
  | --program-transform | --program-transfor \
  | --program-transfo | --program-transf \
  | --program-trans | --program-tran \
  | --progr-tra | --program-tr | --program-t)
    ac_prev=program_transform_name ;;
  -program-transform-name=* | --program-transform-name=* \
  | --program-transform-nam=* | --program-transform-na=* \
  | --program-transform-n=* | --program-transform-=* \
  | --program-transform=* | --program-transfor=* \
  | --program-transfo=* | --program-transf=* \
  | --program-trans=* | --program-tran=* \
  | --progr-tra=* | --program-tr=* | --program-t=*)
    program_transform_name=$ac_optarg ;;

  -pdfdir | --pdfdir | --pdfdi | --pdfd | --pdf | --pd)
    ac_prev=pdfdir ;;
  -pdfdir=* | --pdfdir=* | --pdfdi=* | --pdfd=* | --pdf=* | --pd=*)
    pdfdir=$ac_optarg ;;

  -psdir | --psdir | --psdi | --psd | --ps)
    ac_prev=psdir ;;
  -psdir=* | --psdir=* | --psdi=* | --psd=* | --ps=*)
    psdir=$ac_optarg ;;

  -q | -quiet | --quiet | --quie | --qui | --qu | --q \
  | -silent | --silent | --silen | --sile | --sil)
    silent=yes ;;

  -sbindir | --sbindir | --sbindi | --sbind | --sbin | --sbi | --sb)
    ac_prev=sbindir ;;
  -sbindir=* | --sbindir=* | --sbindi=* | --sbind=* | --sbin=* \
  | --sbi=* | --sb=*)
    sbindir=$ac_optarg ;;

  -sharedstatedir | --sharedstatedir | --sharedstatedi \
  | --sharedstated | --sharedstate | --sharedstat | --sharedsta \
  | --sharedst | --shareds | --shared | --share | --shar \
  | --sha | --sh)
    ac_prev=sharedstatedir ;;
  -sharedstatedir=* | --sharedstatedir=* | --sharedstatedi=* \
  | --sharedstated=* | --sharedstate=* | --sharedstat=* | --sharedsta=* \
  | --sharedst=* | --shareds=* | --shared=* | --share=* | --shar=* \
  | --sha=* | --sh=*)
    sharedstatedir=$ac_optarg ;;

  -site | --site | --sit)
    ac_prev=site ;;
  -site=* | --site=* | --sit=*)
    site=$ac_optarg ;;

  -srcdir | --srcdir | --srcdi | --srcd | --src | --sr)
    ac_prev=srcdir ;;
  -srcdir=* | --srcdir=* | --srcdi=* | --srcd=* | --src=* | --sr=*)
    srcdir=$ac_optarg ;;

  -sysconfdir | --sysconfdir | --sysconfdi | --sysconfd | --sysconf \
  | --syscon | --sysco | --sysc | --sys | --sy)
    ac_prev=sysconfdir ;;
  -sysconfdir=* | --sysconfdir=* | --sysconfdi=* | --sysconfd=* | --sysconf=* \
  | --syscon=* | --sysco=* | --sysc=* | --sys=* | --sy=*)
    sysconfdir=$ac_optarg ;;

  -target | --target | --targe | --targ | --tar | --ta | --t)
    ac_prev=target_alias ;;
  -target=* | --target=* | --targe=* | --targ=* | --tar=* | --ta=* | --t=*)
    target_alias=$ac_optarg ;;

  -v | -verbose | --verbose | --verbos | --verbo | --verb)
    verbose=yes ;;

  -version | --version | --versio | --versi | --vers | -V)
    ac_init_version=: ;;

  -with-* | --with-*)
    ac_useropt=`expr "x$ac_option" : 'x-*with-\([^=]*\)'`
    # Reject names that are not valid shell variable names.
    expr "x$ac_useropt" : ".*[^-+._$as_cr_alnum]" >/dev/null &&
      as_fn_error $? "invalid package name: $ac_useropt"
    ac_useropt_orig=$ac_useropt
    ac_useropt=`$as_echo "$ac_useropt" | sed 's/[-+.]/_/g'`
    case $ac_user_opts in
      *"
"with_$ac_useropt"
"*) ;;
      *) ac_unrecognized_opts="$ac_unrecognized_opts$ac_unrecognized_sep--with-$ac_useropt_orig"
	 ac_unrecognized_sep=', ';;
    esac
    eval with_$ac_useropt=\$ac_optarg ;;

  -without-* | --without-*)
    ac_useropt=`expr "x$ac_option" : 'x-*without-\(.*\)'`
    # Reject names that are not valid shell variable names.
    expr "x$ac_useropt" : ".*[^-+._$as_cr_alnum]" >/dev/null &&
      as_fn_error $? "invalid package name: $ac_useropt"
    ac_useropt_orig=$ac_useropt
    ac_useropt=`$as_echo "$ac_useropt" | sed 's/[-+.]/_/g'`
    case $ac_user_opts in
      *"
"with_$ac_useropt"
"*) ;;
      *) ac_unrecognized_opts="$ac_unrecognized_opts$ac_unrecognized_sep--without-$ac_useropt_orig"
	 ac_unrecognized_sep=', ';;
    esac
    eval with_$ac_useropt=no ;;

  --x)
    # Obsolete; use --with-x.
    with_x=yes ;;

  -x-includes | --x-includes | --x-include | --x-includ | --x-inclu \
  | --x-incl | --x-inc | --x-in | --x-i)
    ac_prev=x_includes ;;
  -x-includes=* | --x-includes=* | --x-include=* | --x-includ=* | --x-inclu=* \
  | --x-incl=* | --x-inc=* | --x-in=* | --x-i=*)
    x_includes=$ac_optarg ;;

  -x-libraries | --x-libraries | --x-librarie | --x-librari \
  | --x-librar | --x-libra | --x-libr | --x-lib | --x-li | --x-l)
    ac_prev=x_libraries ;;
  -x-libraries=* | --x-libraries=* | --x-librarie=* | --x-librari=* \
  | --x-librar=* | --x-libra=* | --x-libr=* | --x-lib=* | --x-li=* | --x-l=*)
    x_libraries=$ac_optarg ;;

  -*) as_fn_error $? "unrecognized option: \`$ac_option'
Try \`$0 --help' for more information"
    ;;

  *=*)
    ac_envvar=`expr "x$ac_option" : 'x\([^=]*\)='`
    # Reject names that are not valid shell variable names.
    case $ac_envvar in #(
      '' | [0-9]* | *[!_$as_cr_alnum]* )
      as_fn_error $? "invalid variable name: \`$ac_envvar'" ;;
    esac
    eval $ac_envvar=\$ac_optarg
    export $ac_envvar ;;

  *)
    # FIXME: should be removed in autoconf 3.0.
    $as_echo "$as_me: WARNING: you should use --build, --host, --target" >&2
    expr "x$ac_option" : ".*[^-._$as_cr_alnum]" >/dev/null &&
      $as_echo "$as_me: WARNING: invalid host type: $ac_option" >&2
    : "${build_alias=$ac_option} ${host_alias=$ac_option} ${target_alias=$ac_option}"
    ;;

  esac
done

if test -n "$ac_prev"; then
  ac_option=--`echo $ac_prev | sed 's/_/-/g'`
  as_fn_error $? "missing argument to $ac_option"
fi

if test -n "$ac_unrecognized_opts"; then
  case $enable_option_checking in
    no) ;;
    fatal) as_fn_error $? "unrecognized options: $ac_unrecognized_opts" ;;
    *)     $as_echo "$as_me: WARNING: unrecognized options: $ac_unrecognized_opts" >&2 ;;
  esac
fi

# Check all directory arguments for consistency.
for ac_var in	exec_prefix prefix bindir sbindir libexecdir datarootdir \
		datadir sysconfdir sharedstatedir localstatedir includedir \
		oldincludedir docdir infodir htmldir dvidir pdfdir psdir \
		libdir localedir mandir
do
  eval ac_val=\$$ac_var
  # Remove trailing slashes.
  case $ac_val in
    */ )
      ac_val=`expr "X$ac_val" : 'X\(.*[^/]\)' \| "X$ac_val" : 'X\(.*\)'`
      eval $ac_var=\$ac_val;;
  esac
  # Be sure to have absolute directory names.
  case $ac_val in
    [\\/$]* | ?:[\\/]* )  continue;;
    NONE | '' ) case $ac_var in *prefix ) continue;; esac;;
  esac
  as_fn_error $? "expected an absolute directory name for --$ac_var: $ac_val"
done

# There might be people who depend on the old broken behavior: `$host'
# used to hold the argument of --host etc.
# FIXME: To remove some day.
build=$build_alias
host=$host_alias
target=$target_alias

# FIXME: To remove some day.
if test "x$host_alias" != x; then
  if test "x$build_alias" = x; then
    cross_compiling=maybe
  elif test "x$build_alias" != "x$host_alias"; then
    cross_compiling=yes
  fi
fi

ac_tool_prefix=
test -n "$host_alias" && ac_tool_prefix=$host_alias-

test "$silent" = yes && exec 6>/dev/null


ac_pwd=`pwd` && test -n "$ac_pwd" &&
ac_ls_di=`ls -di .` &&
ac_pwd_ls_di=`cd "$ac_pwd" && ls -di .` ||
  as_fn_error $? "working directory cannot be determined"
test "X$ac_ls_di" = "X$ac_pwd_ls_di" ||
  as_fn_error $? "pwd does not report name of working directory"


# Find the source files, if location was not specified.
if test -z "$srcdir"; then
  ac_srcdir_defaulted=yes
  # Try the directory containing this script, then the parent directory.
  ac_confdir=`$as_dirname -- "$as_myself" ||
$as_expr X"$as_myself" : 'X\(.*[^/]\)//*[^/][^/]*/*$' \| \
	 X"$as_myself" : 'X\(//\)[^/]' \| \
	 X"$as_myself" : 'X\(//\)$' \| \
	 X"$as_myself" : 'X\(/\)' \| . 2>/dev/null ||
$as_echo X"$as_myself" |
    sed '/^X\(.*[^/]\)\/\/*[^/][^/]*\/*$/{
	    s//\1/
	    q
	  }
	  /^X\(\/\/\)[^/].*/{
	    s//\1/
	    q
	  }
	  /^X\(\/\/\)$/{
	    s//\1/
	    q
	  }
	  /^X\(\/\).*/{
	    s//\1/
	    q
	  }
	  s/.*/./; q'`
  srcdir=$ac_confdir
  if test ! -r "$srcdir/$ac_unique_file"; then
    srcdir=..
  fi
else
  ac_srcdir_defaulted=no
fi
if test ! -r "$srcdir/$ac_unique_file"; then
  test "$ac_srcdir_defaulted" = yes && srcdir="$ac_confdir or .."
  as_fn_error $? "cannot find sources ($ac_unique_file) in $srcdir"
fi
ac_msg="sources are in $srcdir, but \`cd $srcdir' does not work"
ac_abs_confdir=`(
	cd "$srcdir" && test -r "./$ac_unique_file" || as_fn_error $? "$ac_msg"
	pwd)`
# When building in place, set srcdir=.
if test "$ac_abs_confdir" = "$ac_pwd"; then
  srcdir=.
fi
# Remove unnecessary trailing slashes from srcdir.
# Double slashes in file names in object file debugging info
# mess up M-x gdb in Emacs.
case $srcdir in
*/) srcdir=`expr "X$srcdir" : 'X\(.*[^/]\)' \| "X$srcdir" : 'X\(.*\)'`;;
esac
for ac_var in $ac_precious_vars; do
  eval ac_env_${ac_var}_set=\${${ac_var}+set}
  eval ac_env_${ac_var}_value=\$${ac_var}
  eval ac_cv_env_${ac_var}_set=\${${ac_var}+set}
  eval ac_cv_env_${ac_var}_value=\$${ac_var}
done

#
# Report the --help message.
#
if test "$ac_init_help" = "long"; then
  # Omit some internal or obsolete options to make the list less imposing.
  # This message is too long to be a string in the A/UX 3.1 sh.
  cat <<_ACEOF
\`configure' configures f5c 1.0 to adapt to many kinds of systems.

Usage: $0 [OPTION]... [VAR=VALUE]...

To assign environment variables (e.g., CC, CFLAGS...), specify them as
VAR=VALUE.  See below for descriptions of some of the useful variables.

Defaults for the options are specified in brackets.

Configuration:
  -h, --help              display this help and exit
      --help=short        display options specific to this package
      --help=recursive    display the short help of all the included packages
  -V, --version           display version information and exit
  -q, --quiet, --silent   do not print \`checking ...' messages
      --cache-file=FILE   cache test results in FILE [disabled]
  -C, --config-cache      alias for \`--cache-file=config.cache'
  -n, --no-create         do not create output files
      --srcdir=DIR        find the sources in DIR [configure dir or \`..']

Installation directories:
  --prefix=PREFIX         install architecture-independent files in PREFIX
                          [$ac_default_prefix]
  --exec-prefix=EPREFIX   install architecture-dependent files in EPREFIX
                          [PREFIX]

By default, \`make install' will install all the files in
\`$ac_default_prefix/bin', \`$ac_default_prefix/lib' etc.  You can specify
an installation prefix other than \`$ac_default_prefix' using \`--prefix',
for instance \`--prefix=\$HOME'.

For better control, use the options below.

Fine tuning of the installation directories:
  --bindir=DIR            user executables [EPREFIX/bin]
  --sbindir=DIR           system admin executables [EPREFIX/sbin]
  --libexecdir=DIR        program executables [EPREFIX/libexec]
  --sysconfdir=DIR        read-only single-machine data [PREFIX/etc]
  --sharedstatedir=DIR    modifiable architecture-independent data [PREFIX/com]
  --localstatedir=DIR     modifiable single-machine data [PREFIX/var]
  --libdir=DIR            object code libraries [EPREFIX/lib]
  --includedir=DIR        C header files [PREFIX/include]
  --oldincludedir=DIR     C header files for non-gcc [/usr/include]
  --datarootdir=DIR       read-only arch.-independent data root [PREFIX/share]
  --datadir=DIR           read-only architecture-independent data [DATAROOTDIR]
  --infodir=DIR           info documentation [DATAROOTDIR/info]
  --localedir=DIR         locale-dependent data [DATAROOTDIR/locale]
  --mandir=DIR            man documentation [DATAROOTDIR/man]
  --docdir=DIR            documentation root [DATAROOTDIR/doc/f5c]
  --htmldir=DIR           html documentation [DOCDIR]
  --dvidir=DIR            dvi documentation [DOCDIR]
  --pdfdir=DIR            pdf documentation [DOCDIR]
  --psdir=DIR             ps documentation [DOCDIR]
_ACEOF

  cat <<\_ACEOF
_ACEOF
fi

if test -n "$ac_init_help"; then
  case $ac_init_help in
     short | recursive ) echo "Configuration of f5c 1.0:";;
   esac
  cat <<\_ACEOF

Optional Features:
  --disable-option-checking  ignore unrecognized --enable/--with options
  --disable-FEATURE       do not include FEATURE (same as --enable-FEATURE=no)
  --enable-FEATURE[=ARG]  include FEATURE [ARG=yes]
  --enable-systemhts=yes|no
                          Use system wide HTS. Default is disabled.
  --enable-localhdf5      Use local HDF5. Default is disabled.
  --disable-largefile     omit support for large files

Some influential environment variables:
  CC          C compiler command
  CFLAGS      C compiler flags
  LDFLAGS     linker flags, e.g. -L<lib dir> if you have libraries in a
              nonstandard directory <lib dir>
  LIBS        libraries to pass to the linker, e.g. -l<library>
  CPPFLAGS    (Objective) C/C++ preprocessor flags, e.g. -I<include dir> if
              you have headers in a nonstandard directory <include dir>
  CXX         C++ compiler command
  CXXFLAGS    C++ compiler flags
  CPP         C preprocessor

Use these variables to override the choices made by `configure' or to help
it to find libraries and programs with nonstandard names/locations.

Report bugs to <>.
_ACEOF
ac_status=$?
fi

if test "$ac_init_help" = "recursive"; then
  # If there are subdirs, report their specific --help.
  for ac_dir in : $ac_subdirs_all; do test "x$ac_dir" = x: && continue
    test -d "$ac_dir" ||
      { cd "$srcdir" && ac_pwd=`pwd` && srcdir=. && test -d "$ac_dir"; } ||
      continue
    ac_builddir=.

case "$ac_dir" in
.) ac_dir_suffix= ac_top_builddir_sub=. ac_top_build_prefix= ;;
*)
  ac_dir_suffix=/`$as_echo "$ac_dir" | sed 's|^\.[\\/]||'`
  # A ".." for each directory in $ac_dir_suffix.
  ac_top_builddir_sub=`$as_echo "$ac_dir_suffix" | sed 's|/[^\\/]*|/..|g;s|/||'`
  case $ac_top_builddir_sub in
  "") ac_top_builddir_sub=. ac_top_build_prefix= ;;
  *)  ac_top_build_prefix=$ac_top_builddir_sub/ ;;
  esac ;;
esac
ac_abs_top_builddir=$ac_pwd
ac_abs_builddir=$ac_pwd$ac_dir_suffix
# for backward compatibility:
ac_top_builddir=$ac_top_build_prefix

case $srcdir in
  .)  # We are building in place.
    ac_srcdir=.
    ac_top_srcdir=$ac_top_builddir_sub
    ac_abs_top_srcdir=$ac_pwd ;;
  [\\/]* | ?:[\\/]* )  # Absolute name.
    ac_srcdir=$srcdir$ac_dir_suffix;
    ac_top_srcdir=$srcdir
    ac_abs_top_srcdir=$srcdir ;;
  *) # Relative name.
    ac_srcdir=$ac_top_build_prefix$srcdir$ac_dir_suffix
    ac_top_srcdir=$ac_top_build_prefix$srcdir
    ac_abs_top_srcdir=$ac_pwd/$srcdir ;;
esac
ac_abs_srcdir=$ac_abs_top_srcdir$ac_dir_suffix

    cd "$ac_dir" || { ac_status=$?; continue; }
    # Check for guested configure.
    if test -f "$ac_srcdir/configure.gnu"; then
      echo &&
      $SHELL "$ac_srcdir/configure.gnu" --help=recursive
    elif test -f "$ac_srcdir/configure"; then
      echo &&
      $SHELL "$ac_srcdir/configure" --help=recursive
    else
      $as_echo "$as_me: WARNING: no configuration information is in $ac_dir" >&2
    fi || ac_status=$?
    cd "$ac_pwd" || { ac_status=$?; break; }
  done
fi

test -n "$ac_init_help" && exit $ac_status
if $ac_init_version; then
  cat <<\_ACEOF
f5c configure 1.0
generated by GNU Autoconf 2.69

Copyright (C) 2012 Free Software Foundation, Inc.
This configure script is free software; the Free Software Foundation
gives unlimited permission to copy, distribute and modify it.
_ACEOF
  exit
fi

## ------------------------ ##
## Autoconf initialization. ##
## ------------------------ ##

# ac_fn_c_try_compile LINENO
# --------------------------
# Try to compile conftest.$ac_ext, and return whether this succeeded.
ac_fn_c_try_compile ()
{
  as_lineno=${as_lineno-"$1"} as_lineno_stack=as_lineno_stack=$as_lineno_stack
  rm -f conftest.$ac_objext
  if { { ac_try="$ac_compile"
case "(($ac_try" in
  *\"* | *\`* | *\\*) ac_try_echo=\$ac_try;;
  *) ac_try_echo=$ac_try;;
esac
eval ac_try_echo="\"\$as_me:${as_lineno-$LINENO}: $ac_try_echo\""
$as_echo "$ac_try_echo"; } >&5
  (eval "$ac_compile") 2>conftest.err
  ac_status=$?
  if test -s conftest.err; then
    grep -v '^ *+' conftest.err >conftest.er1
    cat conftest.er1 >&5
    mv -f conftest.er1 conftest.err
  fi
  $as_echo "$as_me:${as_lineno-$LINENO}: \$? = $ac_status" >&5
  test $ac_status = 0; } && {
	 test -z "$ac_c_werror_flag" ||
	 test ! -s conftest.err
       } && test -s conftest.$ac_objext; then :
  ac_retval=0
else
  $as_echo "$as_me: failed program was:" >&5
sed 's/^/| /' conftest.$ac_ext >&5

	ac_retval=1
fi
  eval $as_lineno_stack; ${as_lineno_stack:+:} unset as_lineno
  as_fn_set_status $ac_retval

} # ac_fn_c_try_compile

# ac_fn_cxx_try_compile LINENO
# ----------------------------
# Try to compile conftest.$ac_ext, and return whether this succeeded.
ac_fn_cxx_try_compile ()
{
  as_lineno=${as_lineno-"$1"} as_lineno_stack=as_lineno_stack=$as_lineno_stack
  rm -f conftest.$ac_objext
  if { { ac_try="$ac_compile"
case "(($ac_try" in
  *\"* | *\`* | *\\*) ac_try_echo=\$ac_try;;
  *) ac_try_echo=$ac_try;;
esac
eval ac_try_echo="\"\$as_me:${as_lineno-$LINENO}: $ac_try_echo\""
$as_echo "$ac_try_echo"; } >&5
  (eval "$ac_compile") 2>conftest.err
  ac_status=$?
  if test -s conftest.err; then
    grep -v '^ *+' conftest.err >conftest.er1
    cat conftest.er1 >&5
    mv -f conftest.er1 conftest.err
  fi
  $as_echo "$as_me:${as_lineno-$LINENO}: \$? = $ac_status" >&5
  test $ac_status = 0; } && {
	 test -z "$ac_cxx_werror_flag" ||
	 test ! -s conftest.err
       } && test -s conftest.$ac_objext; then :
  ac_retval=0
else
  $as_echo "$as_me: failed program was:" >&5
sed 's/^/| /' conftest.$ac_ext >&5

	ac_retval=1
fi
  eval $as_lineno_stack; ${as_lineno_stack:+:} unset as_lineno
  as_fn_set_status $ac_retval

} # ac_fn_cxx_try_compile

# ac_fn_c_try_link LINENO
# -----------------------
# Try to link conftest.$ac_ext, and return whether this succeeded.
ac_fn_c_try_link ()
{
  as_lineno=${as_lineno-"$1"} as_lineno_stack=as_lineno_stack=$as_lineno_stack
  rm -f conftest.$ac_objext conftest$ac_exeext
  if { { ac_try="$ac_link"
case "(($ac_try" in
  *\"* | *\`* | *\\*) ac_try_echo=\$ac_try;;
  *) ac_try_echo=$ac_try;;
esac
eval ac_try_echo="\"\$as_me:${as_lineno-$LINENO}: $ac_try_echo\""
$as_echo "$ac_try_echo"; } >&5
  (eval "$ac_link") 2>conftest.err
  ac_status=$?
  if test -s conftest.err; then
    grep -v '^ *+' conftest.err >conftest.er1
    cat conftest.er1 >&5
    mv -f conftest.er1 conftest.err
  fi
  $as_echo "$as_me:${as_lineno-$LINENO}: \$? = $ac_status" >&5
  test $ac_status = 0; } && {
	 test -z "$ac_c_werror_flag" ||
	 test ! -s conftest.err
       } && test -s conftest$ac_exeext && {
	 test "$cross_compiling" = yes ||
	 test -x conftest$ac_exeext
       }; then :
  ac_retval=0
else
  $as_echo "$as_me: failed program was:" >&5
sed 's/^/| /' conftest.$ac_ext >&5

	ac_retval=1
fi
  # Delete the IPA/IPO (Inter Procedural Analysis/Optimization) information
  # created by the PGI compiler (conftest_ipa8_conftest.oo), as it would
  # interfere with the next link command; also delete a directory that is
  # left behind by Apple's compiler.  We do this before executing the actions.
  rm -rf conftest.dSYM conftest_ipa8_conftest.oo
  eval $as_lineno_stack; ${as_lineno_stack:+:} unset as_lineno
  as_fn_set_status $ac_retval

} # ac_fn_c_try_link

# ac_fn_c_try_cpp LINENO
# ----------------------
# Try to preprocess conftest.$ac_ext, and return whether this succeeded.
ac_fn_c_try_cpp ()
{
  as_lineno=${as_lineno-"$1"} as_lineno_stack=as_lineno_stack=$as_lineno_stack
  if { { ac_try="$ac_cpp conftest.$ac_ext"
case "(($ac_try" in
  *\"* | *\`* | *\\*) ac_try_echo=\$ac_try;;
  *) ac_try_echo=$ac_try;;
esac
eval ac_try_echo="\"\$as_me:${as_lineno-$LINENO}: $ac_try_echo\""
$as_echo "$ac_try_echo"; } >&5
  (eval "$ac_cpp conftest.$ac_ext") 2>conftest.err
  ac_status=$?
  if test -s conftest.err; then
    grep -v '^ *+' conftest.err >conftest.er1
    cat conftest.er1 >&5
    mv -f conftest.er1 conftest.err
  fi
  $as_echo "$as_me:${as_lineno-$LINENO}: \$? = $ac_status" >&5
  test $ac_status = 0; } > conftest.i && {
	 test -z "$ac_c_preproc_warn_flag$ac_c_werror_flag" ||
	 test ! -s conftest.err
       }; then :
  ac_retval=0
else
  $as_echo "$as_me: failed program was:" >&5
sed 's/^/| /' conftest.$ac_ext >&5

    ac_retval=1
fi
  eval $as_lineno_stack; ${as_lineno_stack:+:} unset as_lineno
  as_fn_set_status $ac_retval

} # ac_fn_c_try_cpp

# ac_fn_c_check_header_mongrel LINENO HEADER VAR INCLUDES
# -------------------------------------------------------
# Tests whether HEADER exists, giving a warning if it cannot be compiled using
# the include files in INCLUDES and setting the cache variable VAR
# accordingly.
ac_fn_c_check_header_mongrel ()
{
  as_lineno=${as_lineno-"$1"} as_lineno_stack=as_lineno_stack=$as_lineno_stack
  if eval \${$3+:} false; then :
  { $as_echo "$as_me:${as_lineno-$LINENO}: checking for $2" >&5
$as_echo_n "checking for $2... " >&6; }
if eval \${$3+:} false; then :
  $as_echo_n "(cached) " >&6
fi
eval ac_res=\$$3
	       { $as_echo "$as_me:${as_lineno-$LINENO}: result: $ac_res" >&5
$as_echo "$ac_res" >&6; }
else
  # Is the header compilable?
{ $as_echo "$as_me:${as_lineno-$LINENO}: checking $2 usability" >&5
$as_echo_n "checking $2 usability... " >&6; }
cat confdefs.h - <<_ACEOF >conftest.$ac_ext
/* end confdefs.h.  */
$4
#include <$2>
_ACEOF
if ac_fn_c_try_compile "$LINENO"; then :
  ac_header_compiler=yes
else
  ac_header_compiler=no
fi
rm -f core conftest.err conftest.$ac_objext conftest.$ac_ext
{ $as_echo "$as_me:${as_lineno-$LINENO}: result: $ac_header_compiler" >&5
$as_echo "$ac_header_compiler" >&6; }

# Is the header present?
{ $as_echo "$as_me:${as_lineno-$LINENO}: checking $2 presence" >&5
$as_echo_n "checking $2 presence... " >&6; }
cat confdefs.h - <<_ACEOF >conftest.$ac_ext
/* end confdefs.h.  */
#include <$2>
_ACEOF
if ac_fn_c_try_cpp "$LINENO"; then :
  ac_header_preproc=yes
else
  ac_header_preproc=no
fi
rm -f conftest.err conftest.i conftest.$ac_ext
{ $as_echo "$as_me:${as_lineno-$LINENO}: result: $ac_header_preproc" >&5
$as_echo "$ac_header_preproc" >&6; }

# So?  What about this header?
case $ac_header_compiler:$ac_header_preproc:$ac_c_preproc_warn_flag in #((
  yes:no: )
    { $as_echo "$as_me:${as_lineno-$LINENO}: WARNING: $2: accepted by the compiler, rejected by the preprocessor!" >&5
$as_echo "$as_me: WARNING: $2: accepted by the compiler, rejected by the preprocessor!" >&2;}
    { $as_echo "$as_me:${as_lineno-$LINENO}: WARNING: $2: proceeding with the compiler's result" >&5
$as_echo "$as_me: WARNING: $2: proceeding with the compiler's result" >&2;}
    ;;
  no:yes:* )
    { $as_echo "$as_me:${as_lineno-$LINENO}: WARNING: $2: present but cannot be compiled" >&5
$as_echo "$as_me: WARNING: $2: present but cannot be compiled" >&2;}
    { $as_echo "$as_me:${as_lineno-$LINENO}: WARNING: $2:     check for missing prerequisite headers?" >&5
$as_echo "$as_me: WARNING: $2:     check for missing prerequisite headers?" >&2;}
    { $as_echo "$as_me:${as_lineno-$LINENO}: WARNING: $2: see the Autoconf documentation" >&5
$as_echo "$as_me: WARNING: $2: see the Autoconf documentation" >&2;}
    { $as_echo "$as_me:${as_lineno-$LINENO}: WARNING: $2:     section \"Present But Cannot Be Compiled\"" >&5
$as_echo "$as_me: WARNING: $2:     section \"Present But Cannot Be Compiled\"" >&2;}
    { $as_echo "$as_me:${as_lineno-$LINENO}: WARNING: $2: proceeding with the compiler's result" >&5
$as_echo "$as_me: WARNING: $2: proceeding with the compiler's result" >&2;}
( $as_echo "## ---------------------------------- ##
## Report this to ##
## ---------------------------------- ##"
     ) | sed "s/^/$as_me: WARNING:     /" >&2
    ;;
esac
  { $as_echo "$as_me:${as_lineno-$LINENO}: checking for $2" >&5
$as_echo_n "checking for $2... " >&6; }
if eval \${$3+:} false; then :
  $as_echo_n "(cached) " >&6
else
  eval "$3=\$ac_header_compiler"
fi
eval ac_res=\$$3
	       { $as_echo "$as_me:${as_lineno-$LINENO}: result: $ac_res" >&5
$as_echo "$ac_res" >&6; }
fi
  eval $as_lineno_stack; ${as_lineno_stack:+:} unset as_lineno

} # ac_fn_c_check_header_mongrel

# ac_fn_c_try_run LINENO
# ----------------------
# Try to link conftest.$ac_ext, and return whether this succeeded. Assumes
# that executables *can* be run.
ac_fn_c_try_run ()
{
  as_lineno=${as_lineno-"$1"} as_lineno_stack=as_lineno_stack=$as_lineno_stack
  if { { ac_try="$ac_link"
case "(($ac_try" in
  *\"* | *\`* | *\\*) ac_try_echo=\$ac_try;;
  *) ac_try_echo=$ac_try;;
esac
eval ac_try_echo="\"\$as_me:${as_lineno-$LINENO}: $ac_try_echo\""
$as_echo "$ac_try_echo"; } >&5
  (eval "$ac_link") 2>&5
  ac_status=$?
  $as_echo "$as_me:${as_lineno-$LINENO}: \$? = $ac_status" >&5
  test $ac_status = 0; } && { ac_try='./conftest$ac_exeext'
  { { case "(($ac_try" in
  *\"* | *\`* | *\\*) ac_try_echo=\$ac_try;;
  *) ac_try_echo=$ac_try;;
esac
eval ac_try_echo="\"\$as_me:${as_lineno-$LINENO}: $ac_try_echo\""
$as_echo "$ac_try_echo"; } >&5
  (eval "$ac_try") 2>&5
  ac_status=$?
  $as_echo "$as_me:${as_lineno-$LINENO}: \$? = $ac_status" >&5
  test $ac_status = 0; }; }; then :
  ac_retval=0
else
  $as_echo "$as_me: program exited with status $ac_status" >&5
       $as_echo "$as_me: failed program was:" >&5
sed 's/^/| /' conftest.$ac_ext >&5

       ac_retval=$ac_status
fi
  rm -rf conftest.dSYM conftest_ipa8_conftest.oo
  eval $as_lineno_stack; ${as_lineno_stack:+:} unset as_lineno
  as_fn_set_status $ac_retval

} # ac_fn_c_try_run

# ac_fn_c_check_header_compile LINENO HEADER VAR INCLUDES
# -------------------------------------------------------
# Tests whether HEADER exists and can be compiled using the include files in
# INCLUDES, setting the cache variable VAR accordingly.
ac_fn_c_check_header_compile ()
{
  as_lineno=${as_lineno-"$1"} as_lineno_stack=as_lineno_stack=$as_lineno_stack
  { $as_echo "$as_me:${as_lineno-$LINENO}: checking for $2" >&5
$as_echo_n "checking for $2... " >&6; }
if eval \${$3+:} false; then :
  $as_echo_n "(cached) " >&6
else
  cat confdefs.h - <<_ACEOF >conftest.$ac_ext
/* end confdefs.h.  */
$4
#include <$2>
_ACEOF
if ac_fn_c_try_compile "$LINENO"; then :
  eval "$3=yes"
else
  eval "$3=no"
fi
rm -f core conftest.err conftest.$ac_objext conftest.$ac_ext
fi
eval ac_res=\$$3
	       { $as_echo "$as_me:${as_lineno-$LINENO}: result: $ac_res" >&5
$as_echo "$ac_res" >&6; }
  eval $as_lineno_stack; ${as_lineno_stack:+:} unset as_lineno

} # ac_fn_c_check_header_compile

# ac_fn_c_check_type LINENO TYPE VAR INCLUDES
# -------------------------------------------
# Tests whether TYPE exists after having included INCLUDES, setting cache
# variable VAR accordingly.
ac_fn_c_check_type ()
{
  as_lineno=${as_lineno-"$1"} as_lineno_stack=as_lineno_stack=$as_lineno_stack
  { $as_echo "$as_me:${as_lineno-$LINENO}: checking for $2" >&5
$as_echo_n "checking for $2... " >&6; }
if eval \${$3+:} false; then :
  $as_echo_n "(cached) " >&6
else
  eval "$3=no"
  cat confdefs.h - <<_ACEOF >conftest.$ac_ext
/* end confdefs.h.  */
$4
int
main ()
{
if (sizeof ($2))
	 return 0;
  ;
  return 0;
}
_ACEOF
if ac_fn_c_try_compile "$LINENO"; then :
  cat confdefs.h - <<_ACEOF >conftest.$ac_ext
/* end confdefs.h.  */
$4
int
main ()
{
if (sizeof (($2)))
	    return 0;
  ;
  return 0;
}
_ACEOF
if ac_fn_c_try_compile "$LINENO"; then :

else
  eval "$3=yes"
fi
rm -f core conftest.err conftest.$ac_objext conftest.$ac_ext
fi
rm -f core conftest.err conftest.$ac_objext conftest.$ac_ext
fi
eval ac_res=\$$3
	       { $as_echo "$as_me:${as_lineno-$LINENO}: result: $ac_res" >&5
$as_echo "$ac_res" >&6; }
  eval $as_lineno_stack; ${as_lineno_stack:+:} unset as_lineno

} # ac_fn_c_check_type

# ac_fn_c_find_intX_t LINENO BITS VAR
# -----------------------------------
# Finds a signed integer type with width BITS, setting cache variable VAR
# accordingly.
ac_fn_c_find_intX_t ()
{
  as_lineno=${as_lineno-"$1"} as_lineno_stack=as_lineno_stack=$as_lineno_stack
  { $as_echo "$as_me:${as_lineno-$LINENO}: checking for int$2_t" >&5
$as_echo_n "checking for int$2_t... " >&6; }
if eval \${$3+:} false; then :
  $as_echo_n "(cached) " >&6
else
  eval "$3=no"
     # Order is important - never check a type that is potentially smaller
     # than half of the expected target width.
     for ac_type in int$2_t 'int' 'long int' \
	 'long long int' 'short int' 'signed char'; do
       cat confdefs.h - <<_ACEOF >conftest.$ac_ext
/* end confdefs.h.  */
$ac_includes_default
	     enum { N = $2 / 2 - 1 };
int
main ()
{
static int test_array [1 - 2 * !(0 < ($ac_type) ((((($ac_type) 1 << N) << N) - 1) * 2 + 1))];
test_array [0] = 0;
return test_array [0];

  ;
  return 0;
}
_ACEOF
if ac_fn_c_try_compile "$LINENO"; then :
  cat confdefs.h - <<_ACEOF >conftest.$ac_ext
/* end confdefs.h.  */
$ac_includes_default
	        enum { N = $2 / 2 - 1 };
int
main ()
{
static int test_array [1 - 2 * !(($ac_type) ((((($ac_type) 1 << N) << N) - 1) * 2 + 1)
		 < ($ac_type) ((((($ac_type) 1 << N) << N) - 1) * 2 + 2))];
test_array [0] = 0;
return test_array [0];

  ;
  return 0;
}
_ACEOF
if ac_fn_c_try_compile "$LINENO"; then :

else
  case $ac_type in #(
  int$2_t) :
    eval "$3=yes" ;; #(
  *) :
    eval "$3=\$ac_type" ;;
esac
fi
rm -f core conftest.err conftest.$ac_objext conftest.$ac_ext
fi
rm -f core conftest.err conftest.$ac_objext conftest.$ac_ext
       if eval test \"x\$"$3"\" = x"no"; then :

else
  break
fi
     done
fi
eval ac_res=\$$3
	       { $as_echo "$as_me:${as_lineno-$LINENO}: result: $ac_res" >&5
$as_echo "$ac_res" >&6; }
  eval $as_lineno_stack; ${as_lineno_stack:+:} unset as_lineno

} # ac_fn_c_find_intX_t

# ac_fn_c_find_uintX_t LINENO BITS VAR
# ------------------------------------
# Finds an unsigned integer type with width BITS, setting cache variable VAR
# accordingly.
ac_fn_c_find_uintX_t ()
{
  as_lineno=${as_lineno-"$1"} as_lineno_stack=as_lineno_stack=$as_lineno_stack
  { $as_echo "$as_me:${as_lineno-$LINENO}: checking for uint$2_t" >&5
$as_echo_n "checking for uint$2_t... " >&6; }
if eval \${$3+:} false; then :
  $as_echo_n "(cached) " >&6
else
  eval "$3=no"
     # Order is important - never check a type that is potentially smaller
     # than half of the expected target width.
     for ac_type in uint$2_t 'unsigned int' 'unsigned long int' \
	 'unsigned long long int' 'unsigned short int' 'unsigned char'; do
       cat confdefs.h - <<_ACEOF >conftest.$ac_ext
/* end confdefs.h.  */
$ac_includes_default
int
main ()
{
static int test_array [1 - 2 * !((($ac_type) -1 >> ($2 / 2 - 1)) >> ($2 / 2 - 1) == 3)];
test_array [0] = 0;
return test_array [0];

  ;
  return 0;
}
_ACEOF
if ac_fn_c_try_compile "$LINENO"; then :
  case $ac_type in #(
  uint$2_t) :
    eval "$3=yes" ;; #(
  *) :
    eval "$3=\$ac_type" ;;
esac
fi
rm -f core conftest.err conftest.$ac_objext conftest.$ac_ext
       if eval test \"x\$"$3"\" = x"no"; then :

else
  break
fi
     done
fi
eval ac_res=\$$3
	       { $as_echo "$as_me:${as_lineno-$LINENO}: result: $ac_res" >&5
$as_echo "$ac_res" >&6; }
  eval $as_lineno_stack; ${as_lineno_stack:+:} unset as_lineno

} # ac_fn_c_find_uintX_t

# ac_fn_c_check_func LINENO FUNC VAR
# ----------------------------------
# Tests whether FUNC exists, setting the cache variable VAR accordingly
ac_fn_c_check_func ()
{
  as_lineno=${as_lineno-"$1"} as_lineno_stack=as_lineno_stack=$as_lineno_stack
  { $as_echo "$as_me:${as_lineno-$LINENO}: checking for $2" >&5
$as_echo_n "checking for $2... " >&6; }
if eval \${$3+:} false; then :
  $as_echo_n "(cached) " >&6
else
  cat confdefs.h - <<_ACEOF >conftest.$ac_ext
/* end confdefs.h.  */
/* Define $2 to an innocuous variant, in case <limits.h> declares $2.
   For example, HP-UX 11i <limits.h> declares gettimeofday.  */
#define $2 innocuous_$2

/* System header to define __stub macros and hopefully few prototypes,
    which can conflict with char $2 (); below.
    Prefer <limits.h> to <assert.h> if __STDC__ is defined, since
    <limits.h> exists even on freestanding compilers.  */

#ifdef __STDC__
# include <limits.h>
#else
# include <assert.h>
#endif

#undef $2

/* Override any GCC internal prototype to avoid an error.
   Use char because int might match the return type of a GCC
   builtin and then its argument prototype would still apply.  */
#ifdef __cplusplus
extern "C"
#endif
char $2 ();
/* The GNU C library defines this for functions which it implements
    to always fail with ENOSYS.  Some functions are actually named
    something starting with __ and the normal name is an alias.  */
#if defined __stub_$2 || defined __stub___$2
choke me
#endif

int
main ()
{
return $2 ();
  ;
  return 0;
}
_ACEOF
if ac_fn_c_try_link "$LINENO"; then :
  eval "$3=yes"
else
  eval "$3=no"
fi
rm -f core conftest.err conftest.$ac_objext \
    conftest$ac_exeext conftest.$ac_ext
fi
eval ac_res=\$$3
	       { $as_echo "$as_me:${as_lineno-$LINENO}: result: $ac_res" >&5
$as_echo "$ac_res" >&6; }
  eval $as_lineno_stack; ${as_lineno_stack:+:} unset as_lineno

} # ac_fn_c_check_func
cat >config.log <<_ACEOF
This file contains any messages produced by compilers while
running configure, to aid debugging if configure makes a mistake.

It was created by f5c $as_me 1.0, which was
generated by GNU Autoconf 2.69.  Invocation command line was

  $ $0 $@

_ACEOF
exec 5>>config.log
{
cat <<_ASUNAME
## --------- ##
## Platform. ##
## --------- ##

hostname = `(hostname || uname -n) 2>/dev/null | sed 1q`
uname -m = `(uname -m) 2>/dev/null || echo unknown`
uname -r = `(uname -r) 2>/dev/null || echo unknown`
uname -s = `(uname -s) 2>/dev/null || echo unknown`
uname -v = `(uname -v) 2>/dev/null || echo unknown`

/usr/bin/uname -p = `(/usr/bin/uname -p) 2>/dev/null || echo unknown`
/bin/uname -X     = `(/bin/uname -X) 2>/dev/null     || echo unknown`

/bin/arch              = `(/bin/arch) 2>/dev/null              || echo unknown`
/usr/bin/arch -k       = `(/usr/bin/arch -k) 2>/dev/null       || echo unknown`
/usr/convex/getsysinfo = `(/usr/convex/getsysinfo) 2>/dev/null || echo unknown`
/usr/bin/hostinfo      = `(/usr/bin/hostinfo) 2>/dev/null      || echo unknown`
/bin/machine           = `(/bin/machine) 2>/dev/null           || echo unknown`
/usr/bin/oslevel       = `(/usr/bin/oslevel) 2>/dev/null       || echo unknown`
/bin/universe          = `(/bin/universe) 2>/dev/null          || echo unknown`

_ASUNAME

as_save_IFS=$IFS; IFS=$PATH_SEPARATOR
for as_dir in $PATH
do
  IFS=$as_save_IFS
  test -z "$as_dir" && as_dir=.
    $as_echo "PATH: $as_dir"
  done
IFS=$as_save_IFS

} >&5

cat >&5 <<_ACEOF


## ----------- ##
## Core tests. ##
## ----------- ##

_ACEOF


# Keep a trace of the command line.
# Strip out --no-create and --no-recursion so they do not pile up.
# Strip out --silent because we don't want to record it for future runs.
# Also quote any args containing shell meta-characters.
# Make two passes to allow for proper duplicate-argument suppression.
ac_configure_args=
ac_configure_args0=
ac_configure_args1=
ac_must_keep_next=false
for ac_pass in 1 2
do
  for ac_arg
  do
    case $ac_arg in
    -no-create | --no-c* | -n | -no-recursion | --no-r*) continue ;;
    -q | -quiet | --quiet | --quie | --qui | --qu | --q \
    | -silent | --silent | --silen | --sile | --sil)
      continue ;;
    *\'*)
      ac_arg=`$as_echo "$ac_arg" | sed "s/'/'\\\\\\\\''/g"` ;;
    esac
    case $ac_pass in
    1) as_fn_append ac_configure_args0 " '$ac_arg'" ;;
    2)
      as_fn_append ac_configure_args1 " '$ac_arg'"
      if test $ac_must_keep_next = true; then
	ac_must_keep_next=false # Got value, back to normal.
      else
	case $ac_arg in
	  *=* | --config-cache | -C | -disable-* | --disable-* \
	  | -enable-* | --enable-* | -gas | --g* | -nfp | --nf* \
	  | -q | -quiet | --q* | -silent | --sil* | -v | -verb* \
	  | -with-* | --with-* | -without-* | --without-* | --x)
	    case "$ac_configure_args0 " in
	      "$ac_configure_args1"*" '$ac_arg' "* ) continue ;;
	    esac
	    ;;
	  -* ) ac_must_keep_next=true ;;
	esac
      fi
      as_fn_append ac_configure_args " '$ac_arg'"
      ;;
    esac
  done
done
{ ac_configure_args0=; unset ac_configure_args0;}
{ ac_configure_args1=; unset ac_configure_args1;}

# When interrupted or exit'd, cleanup temporary files, and complete
# config.log.  We remove comments because anyway the quotes in there
# would cause problems or look ugly.
# WARNING: Use '\'' to represent an apostrophe within the trap.
# WARNING: Do not start the trap code with a newline, due to a FreeBSD 4.0 bug.
trap 'exit_status=$?
  # Save into config.log some information that might help in debugging.
  {
    echo

    $as_echo "## ---------------- ##
## Cache variables. ##
## ---------------- ##"
    echo
    # The following way of writing the cache mishandles newlines in values,
(
  for ac_var in `(set) 2>&1 | sed -n '\''s/^\([a-zA-Z_][a-zA-Z0-9_]*\)=.*/\1/p'\''`; do
    eval ac_val=\$$ac_var
    case $ac_val in #(
    *${as_nl}*)
      case $ac_var in #(
      *_cv_*) { $as_echo "$as_me:${as_lineno-$LINENO}: WARNING: cache variable $ac_var contains a newline" >&5
$as_echo "$as_me: WARNING: cache variable $ac_var contains a newline" >&2;} ;;
      esac
      case $ac_var in #(
      _ | IFS | as_nl) ;; #(
      BASH_ARGV | BASH_SOURCE) eval $ac_var= ;; #(
      *) { eval $ac_var=; unset $ac_var;} ;;
      esac ;;
    esac
  done
  (set) 2>&1 |
    case $as_nl`(ac_space='\'' '\''; set) 2>&1` in #(
    *${as_nl}ac_space=\ *)
      sed -n \
	"s/'\''/'\''\\\\'\'''\''/g;
	  s/^\\([_$as_cr_alnum]*_cv_[_$as_cr_alnum]*\\)=\\(.*\\)/\\1='\''\\2'\''/p"
      ;; #(
    *)
      sed -n "/^[_$as_cr_alnum]*_cv_[_$as_cr_alnum]*=/p"
      ;;
    esac |
    sort
)
    echo

    $as_echo "## ----------------- ##
## Output variables. ##
## ----------------- ##"
    echo
    for ac_var in $ac_subst_vars
    do
      eval ac_val=\$$ac_var
      case $ac_val in
      *\'\''*) ac_val=`$as_echo "$ac_val" | sed "s/'\''/'\''\\\\\\\\'\'''\''/g"`;;
      esac
      $as_echo "$ac_var='\''$ac_val'\''"
    done | sort
    echo

    if test -n "$ac_subst_files"; then
      $as_echo "## ------------------- ##
## File substitutions. ##
## ------------------- ##"
      echo
      for ac_var in $ac_subst_files
      do
	eval ac_val=\$$ac_var
	case $ac_val in
	*\'\''*) ac_val=`$as_echo "$ac_val" | sed "s/'\''/'\''\\\\\\\\'\'''\''/g"`;;
	esac
	$as_echo "$ac_var='\''$ac_val'\''"
      done | sort
      echo
    fi

    if test -s confdefs.h; then
      $as_echo "## ----------- ##
## confdefs.h. ##
## ----------- ##"
      echo
      cat confdefs.h
      echo
    fi
    test "$ac_signal" != 0 &&
      $as_echo "$as_me: caught signal $ac_signal"
    $as_echo "$as_me: exit $exit_status"
  } >&5
  rm -f core *.core core.conftest.* &&
    rm -f -r conftest* confdefs* conf$$* $ac_clean_files &&
    exit $exit_status
' 0
for ac_signal in 1 2 13 15; do
  trap 'ac_signal='$ac_signal'; as_fn_exit 1' $ac_signal
done
ac_signal=0

# confdefs.h avoids OS command line length limits that DEFS can exceed.
rm -f -r conftest* confdefs.h

$as_echo "/* confdefs.h */" > confdefs.h

# Predefined preprocessor variables.

cat >>confdefs.h <<_ACEOF
#define PACKAGE_NAME "$PACKAGE_NAME"
_ACEOF

cat >>confdefs.h <<_ACEOF
#define PACKAGE_TARNAME "$PACKAGE_TARNAME"
_ACEOF

cat >>confdefs.h <<_ACEOF
#define PACKAGE_VERSION "$PACKAGE_VERSION"
_ACEOF

cat >>confdefs.h <<_ACEOF
#define PACKAGE_STRING "$PACKAGE_STRING"
_ACEOF

cat >>confdefs.h <<_ACEOF
#define PACKAGE_BUGREPORT "$PACKAGE_BUGREPORT"
_ACEOF

cat >>confdefs.h <<_ACEOF
#define PACKAGE_URL "$PACKAGE_URL"
_ACEOF


# Let the site file select an alternate cache file if it wants to.
# Prefer an explicitly selected file to automatically selected ones.
ac_site_file1=NONE
ac_site_file2=NONE
if test -n "$CONFIG_SITE"; then
  # We do not want a PATH search for config.site.
  case $CONFIG_SITE in #((
    -*)  ac_site_file1=./$CONFIG_SITE;;
    */*) ac_site_file1=$CONFIG_SITE;;
    *)   ac_site_file1=./$CONFIG_SITE;;
  esac
elif test "x$prefix" != xNONE; then
  ac_site_file1=$prefix/share/config.site
  ac_site_file2=$prefix/etc/config.site
else
  ac_site_file1=$ac_default_prefix/share/config.site
  ac_site_file2=$ac_default_prefix/etc/config.site
fi
for ac_site_file in "$ac_site_file1" "$ac_site_file2"
do
  test "x$ac_site_file" = xNONE && continue
  if test /dev/null != "$ac_site_file" && test -r "$ac_site_file"; then
    { $as_echo "$as_me:${as_lineno-$LINENO}: loading site script $ac_site_file" >&5
$as_echo "$as_me: loading site script $ac_site_file" >&6;}
    sed 's/^/| /' "$ac_site_file" >&5
    . "$ac_site_file" \
      || { { $as_echo "$as_me:${as_lineno-$LINENO}: error: in \`$ac_pwd':" >&5
$as_echo "$as_me: error: in \`$ac_pwd':" >&2;}
as_fn_error $? "failed to load site script $ac_site_file
See \`config.log' for more details" "$LINENO" 5; }
  fi
done

if test -r "$cache_file"; then
  # Some versions of bash will fail to source /dev/null (special files
  # actually), so we avoid doing that.  DJGPP emulates it as a regular file.
  if test /dev/null != "$cache_file" && test -f "$cache_file"; then
    { $as_echo "$as_me:${as_lineno-$LINENO}: loading cache $cache_file" >&5
$as_echo "$as_me: loading cache $cache_file" >&6;}
    case $cache_file in
      [\\/]* | ?:[\\/]* ) . "$cache_file";;
      *)                      . "./$cache_file";;
    esac
  fi
else
  { $as_echo "$as_me:${as_lineno-$LINENO}: creating cache $cache_file" >&5
$as_echo "$as_me: creating cache $cache_file" >&6;}
  >$cache_file
fi

# Check that the precious variables saved in the cache have kept the same
# value.
ac_cache_corrupted=false
for ac_var in $ac_precious_vars; do
  eval ac_old_set=\$ac_cv_env_${ac_var}_set
  eval ac_new_set=\$ac_env_${ac_var}_set
  eval ac_old_val=\$ac_cv_env_${ac_var}_value
  eval ac_new_val=\$ac_env_${ac_var}_value
  case $ac_old_set,$ac_new_set in
    set,)
      { $as_echo "$as_me:${as_lineno-$LINENO}: error: \`$ac_var' was set to \`$ac_old_val' in the previous run" >&5
$as_echo "$as_me: error: \`$ac_var' was set to \`$ac_old_val' in the previous run" >&2;}
      ac_cache_corrupted=: ;;
    ,set)
      { $as_echo "$as_me:${as_lineno-$LINENO}: error: \`$ac_var' was not set in the previous run" >&5
$as_echo "$as_me: error: \`$ac_var' was not set in the previous run" >&2;}
      ac_cache_corrupted=: ;;
    ,);;
    *)
      if test "x$ac_old_val" != "x$ac_new_val"; then
	# differences in whitespace do not lead to failure.
	ac_old_val_w=`echo x $ac_old_val`
	ac_new_val_w=`echo x $ac_new_val`
	if test "$ac_old_val_w" != "$ac_new_val_w"; then
	  { $as_echo "$as_me:${as_lineno-$LINENO}: error: \`$ac_var' has changed since the previous run:" >&5
$as_echo "$as_me: error: \`$ac_var' has changed since the previous run:" >&2;}
	  ac_cache_corrupted=:
	else
	  { $as_echo "$as_me:${as_lineno-$LINENO}: warning: ignoring whitespace changes in \`$ac_var' since the previous run:" >&5
$as_echo "$as_me: warning: ignoring whitespace changes in \`$ac_var' since the previous run:" >&2;}
	  eval $ac_var=\$ac_old_val
	fi
	{ $as_echo "$as_me:${as_lineno-$LINENO}:   former value:  \`$ac_old_val'" >&5
$as_echo "$as_me:   former value:  \`$ac_old_val'" >&2;}
	{ $as_echo "$as_me:${as_lineno-$LINENO}:   current value: \`$ac_new_val'" >&5
$as_echo "$as_me:   current value: \`$ac_new_val'" >&2;}
      fi;;
  esac
  # Pass precious variables to config.status.
  if test "$ac_new_set" = set; then
    case $ac_new_val in
    *\'*) ac_arg=$ac_var=`$as_echo "$ac_new_val" | sed "s/'/'\\\\\\\\''/g"` ;;
    *) ac_arg=$ac_var=$ac_new_val ;;
    esac
    case " $ac_configure_args " in
      *" '$ac_arg' "*) ;; # Avoid dups.  Use of quotes ensures accuracy.
      *) as_fn_append ac_configure_args " '$ac_arg'" ;;
    esac
  fi
done
if $ac_cache_corrupted; then
  { $as_echo "$as_me:${as_lineno-$LINENO}: error: in \`$ac_pwd':" >&5
$as_echo "$as_me: error: in \`$ac_pwd':" >&2;}
  { $as_echo "$as_me:${as_lineno-$LINENO}: error: changes in the environment can compromise the build" >&5
$as_echo "$as_me: error: changes in the environment can compromise the build" >&2;}
  as_fn_error $? "run \`make distclean' and/or \`rm $cache_file' and start over" "$LINENO" 5
fi
## -------------------- ##
## Main body of script. ##
## -------------------- ##

ac_ext=c
ac_cpp='$CPP $CPPFLAGS'
ac_compile='$CC -c $CFLAGS $CPPFLAGS conftest.$ac_ext >&5'
ac_link='$CC -o conftest$ac_exeext $CFLAGS $CPPFLAGS $LDFLAGS conftest.$ac_ext $LIBS >&5'
ac_compiler_gnu=$ac_cv_c_compiler_gnu


#AM_INIT_AUTOMAKE([foreign])

ac_config_headers="$ac_config_headers src/config.h"


# Check whether --enable-systemhts was given.
if test "${enable_systemhts+set}" = set; then :
  enableval=$enable_systemhts; enable_systemhts=$enableval
else
  enable_systemhts=no
fi


# Check whether --enable-localhdf5 was given.
if test "${enable_localhdf5+set}" = set; then :
  enableval=$enable_localhdf5; enable_localhdf5=$enableval
else
  enable_localhdf5=no
fi


# Checks for programs.
ac_ext=c
ac_cpp='$CPP $CPPFLAGS'
ac_compile='$CC -c $CFLAGS $CPPFLAGS conftest.$ac_ext >&5'
ac_link='$CC -o conftest$ac_exeext $CFLAGS $CPPFLAGS $LDFLAGS conftest.$ac_ext $LIBS >&5'
ac_compiler_gnu=$ac_cv_c_compiler_gnu
if test -n "$ac_tool_prefix"; then
  for ac_prog in gcc
  do
    # Extract the first word of "$ac_tool_prefix$ac_prog", so it can be a program name with args.
set dummy $ac_tool_prefix$ac_prog; ac_word=$2
{ $as_echo "$as_me:${as_lineno-$LINENO}: checking for $ac_word" >&5
$as_echo_n "checking for $ac_word... " >&6; }
if ${ac_cv_prog_CC+:} false; then :
  $as_echo_n "(cached) " >&6
else
  if test -n "$CC"; then
  ac_cv_prog_CC="$CC" # Let the user override the test.
else
as_save_IFS=$IFS; IFS=$PATH_SEPARATOR
for as_dir in $PATH
do
  IFS=$as_save_IFS
  test -z "$as_dir" && as_dir=.
    for ac_exec_ext in '' $ac_executable_extensions; do
  if as_fn_executable_p "$as_dir/$ac_word$ac_exec_ext"; then
    ac_cv_prog_CC="$ac_tool_prefix$ac_prog"
    $as_echo "$as_me:${as_lineno-$LINENO}: found $as_dir/$ac_word$ac_exec_ext" >&5
    break 2
  fi
done
  done
IFS=$as_save_IFS

fi
fi
CC=$ac_cv_prog_CC
if test -n "$CC"; then
  { $as_echo "$as_me:${as_lineno-$LINENO}: result: $CC" >&5
$as_echo "$CC" >&6; }
else
  { $as_echo "$as_me:${as_lineno-$LINENO}: result: no" >&5
$as_echo "no" >&6; }
fi


    test -n "$CC" && break
  done
fi
if test -z "$CC"; then
  ac_ct_CC=$CC
  for ac_prog in gcc
do
  # Extract the first word of "$ac_prog", so it can be a program name with args.
set dummy $ac_prog; ac_word=$2
{ $as_echo "$as_me:${as_lineno-$LINENO}: checking for $ac_word" >&5
$as_echo_n "checking for $ac_word... " >&6; }
if ${ac_cv_prog_ac_ct_CC+:} false; then :
  $as_echo_n "(cached) " >&6
else
  if test -n "$ac_ct_CC"; then
  ac_cv_prog_ac_ct_CC="$ac_ct_CC" # Let the user override the test.
else
as_save_IFS=$IFS; IFS=$PATH_SEPARATOR
for as_dir in $PATH
do
  IFS=$as_save_IFS
  test -z "$as_dir" && as_dir=.
    for ac_exec_ext in '' $ac_executable_extensions; do
  if as_fn_executable_p "$as_dir/$ac_word$ac_exec_ext"; then
    ac_cv_prog_ac_ct_CC="$ac_prog"
    $as_echo "$as_me:${as_lineno-$LINENO}: found $as_dir/$ac_word$ac_exec_ext" >&5
    break 2
  fi
done
  done
IFS=$as_save_IFS

fi
fi
ac_ct_CC=$ac_cv_prog_ac_ct_CC
if test -n "$ac_ct_CC"; then
  { $as_echo "$as_me:${as_lineno-$LINENO}: result: $ac_ct_CC" >&5
$as_echo "$ac_ct_CC" >&6; }
else
  { $as_echo "$as_me:${as_lineno-$LINENO}: result: no" >&5
$as_echo "no" >&6; }
fi


  test -n "$ac_ct_CC" && break
done

  if test "x$ac_ct_CC" = x; then
    CC=""
  else
    case $cross_compiling:$ac_tool_warned in
yes:)
{ $as_echo "$as_me:${as_lineno-$LINENO}: WARNING: using cross tools not prefixed with host triplet" >&5
$as_echo "$as_me: WARNING: using cross tools not prefixed with host triplet" >&2;}
ac_tool_warned=yes ;;
esac
    CC=$ac_ct_CC
  fi
fi


test -z "$CC" && { { $as_echo "$as_me:${as_lineno-$LINENO}: error: in \`$ac_pwd':" >&5
$as_echo "$as_me: error: in \`$ac_pwd':" >&2;}
as_fn_error $? "no acceptable C compiler found in \$PATH
See \`config.log' for more details" "$LINENO" 5; }

# Provide some information about the compiler.
$as_echo "$as_me:${as_lineno-$LINENO}: checking for C compiler version" >&5
set X $ac_compile
ac_compiler=$2
for ac_option in --version -v -V -qversion; do
  { { ac_try="$ac_compiler $ac_option >&5"
case "(($ac_try" in
  *\"* | *\`* | *\\*) ac_try_echo=\$ac_try;;
  *) ac_try_echo=$ac_try;;
esac
eval ac_try_echo="\"\$as_me:${as_lineno-$LINENO}: $ac_try_echo\""
$as_echo "$ac_try_echo"; } >&5
  (eval "$ac_compiler $ac_option >&5") 2>conftest.err
  ac_status=$?
  if test -s conftest.err; then
    sed '10a\
... rest of stderr output deleted ...
         10q' conftest.err >conftest.er1
    cat conftest.er1 >&5
  fi
  rm -f conftest.er1 conftest.err
  $as_echo "$as_me:${as_lineno-$LINENO}: \$? = $ac_status" >&5
  test $ac_status = 0; }
done

cat confdefs.h - <<_ACEOF >conftest.$ac_ext
/* end confdefs.h.  */

int
main ()
{

  ;
  return 0;
}
_ACEOF
ac_clean_files_save=$ac_clean_files
ac_clean_files="$ac_clean_files a.out a.out.dSYM a.exe b.out"
# Try to create an executable without -o first, disregard a.out.
# It will help us diagnose broken compilers, and finding out an intuition
# of exeext.
{ $as_echo "$as_me:${as_lineno-$LINENO}: checking whether the C compiler works" >&5
$as_echo_n "checking whether the C compiler works... " >&6; }
ac_link_default=`$as_echo "$ac_link" | sed 's/ -o *conftest[^ ]*//'`

# The possible output files:
ac_files="a.out conftest.exe conftest a.exe a_out.exe b.out conftest.*"

ac_rmfiles=
for ac_file in $ac_files
do
  case $ac_file in
    *.$ac_ext | *.xcoff | *.tds | *.d | *.pdb | *.xSYM | *.bb | *.bbg | *.map | *.inf | *.dSYM | *.o | *.obj ) ;;
    * ) ac_rmfiles="$ac_rmfiles $ac_file";;
  esac
done
rm -f $ac_rmfiles

if { { ac_try="$ac_link_default"
case "(($ac_try" in
  *\"* | *\`* | *\\*) ac_try_echo=\$ac_try;;
  *) ac_try_echo=$ac_try;;
esac
eval ac_try_echo="\"\$as_me:${as_lineno-$LINENO}: $ac_try_echo\""
$as_echo "$ac_try_echo"; } >&5
  (eval "$ac_link_default") 2>&5
  ac_status=$?
  $as_echo "$as_me:${as_lineno-$LINENO}: \$? = $ac_status" >&5
  test $ac_status = 0; }; then :
  # Autoconf-2.13 could set the ac_cv_exeext variable to `no'.
# So ignore a value of `no', otherwise this would lead to `EXEEXT = no'
# in a Makefile.  We should not override ac_cv_exeext if it was cached,
# so that the user can short-circuit this test for compilers unknown to
# Autoconf.
for ac_file in $ac_files ''
do
  test -f "$ac_file" || continue
  case $ac_file in
    *.$ac_ext | *.xcoff | *.tds | *.d | *.pdb | *.xSYM | *.bb | *.bbg | *.map | *.inf | *.dSYM | *.o | *.obj )
	;;
    [ab].out )
	# We found the default executable, but exeext='' is most
	# certainly right.
	break;;
    *.* )
	if test "${ac_cv_exeext+set}" = set && test "$ac_cv_exeext" != no;
	then :; else
	   ac_cv_exeext=`expr "$ac_file" : '[^.]*\(\..*\)'`
	fi
	# We set ac_cv_exeext here because the later test for it is not
	# safe: cross compilers may not add the suffix if given an `-o'
	# argument, so we may need to know it at that point already.
	# Even if this section looks crufty: it has the advantage of
	# actually working.
	break;;
    * )
	break;;
  esac
done
test "$ac_cv_exeext" = no && ac_cv_exeext=

else
  ac_file=''
fi
if test -z "$ac_file"; then :
  { $as_echo "$as_me:${as_lineno-$LINENO}: result: no" >&5
$as_echo "no" >&6; }
$as_echo "$as_me: failed program was:" >&5
sed 's/^/| /' conftest.$ac_ext >&5

{ { $as_echo "$as_me:${as_lineno-$LINENO}: error: in \`$ac_pwd':" >&5
$as_echo "$as_me: error: in \`$ac_pwd':" >&2;}
as_fn_error 77 "C compiler cannot create executables
See \`config.log' for more details" "$LINENO" 5; }
else
  { $as_echo "$as_me:${as_lineno-$LINENO}: result: yes" >&5
$as_echo "yes" >&6; }
fi
{ $as_echo "$as_me:${as_lineno-$LINENO}: checking for C compiler default output file name" >&5
$as_echo_n "checking for C compiler default output file name... " >&6; }
{ $as_echo "$as_me:${as_lineno-$LINENO}: result: $ac_file" >&5
$as_echo "$ac_file" >&6; }
ac_exeext=$ac_cv_exeext

rm -f -r a.out a.out.dSYM a.exe conftest$ac_cv_exeext b.out
ac_clean_files=$ac_clean_files_save
{ $as_echo "$as_me:${as_lineno-$LINENO}: checking for suffix of executables" >&5
$as_echo_n "checking for suffix of executables... " >&6; }
if { { ac_try="$ac_link"
case "(($ac_try" in
  *\"* | *\`* | *\\*) ac_try_echo=\$ac_try;;
  *) ac_try_echo=$ac_try;;
esac
eval ac_try_echo="\"\$as_me:${as_lineno-$LINENO}: $ac_try_echo\""
$as_echo "$ac_try_echo"; } >&5
  (eval "$ac_link") 2>&5
  ac_status=$?
  $as_echo "$as_me:${as_lineno-$LINENO}: \$? = $ac_status" >&5
  test $ac_status = 0; }; then :
  # If both `conftest.exe' and `conftest' are `present' (well, observable)
# catch `conftest.exe'.  For instance with Cygwin, `ls conftest' will
# work properly (i.e., refer to `conftest.exe'), while it won't with
# `rm'.
for ac_file in conftest.exe conftest conftest.*; do
  test -f "$ac_file" || continue
  case $ac_file in
    *.$ac_ext | *.xcoff | *.tds | *.d | *.pdb | *.xSYM | *.bb | *.bbg | *.map | *.inf | *.dSYM | *.o | *.obj ) ;;
    *.* ) ac_cv_exeext=`expr "$ac_file" : '[^.]*\(\..*\)'`
	  break;;
    * ) break;;
  esac
done
else
  { { $as_echo "$as_me:${as_lineno-$LINENO}: error: in \`$ac_pwd':" >&5
$as_echo "$as_me: error: in \`$ac_pwd':" >&2;}
as_fn_error $? "cannot compute suffix of executables: cannot compile and link
See \`config.log' for more details" "$LINENO" 5; }
fi
rm -f conftest conftest$ac_cv_exeext
{ $as_echo "$as_me:${as_lineno-$LINENO}: result: $ac_cv_exeext" >&5
$as_echo "$ac_cv_exeext" >&6; }

rm -f conftest.$ac_ext
EXEEXT=$ac_cv_exeext
ac_exeext=$EXEEXT
cat confdefs.h - <<_ACEOF >conftest.$ac_ext
/* end confdefs.h.  */
#include <stdio.h>
int
main ()
{
FILE *f = fopen ("conftest.out", "w");
 return ferror (f) || fclose (f) != 0;

  ;
  return 0;
}
_ACEOF
ac_clean_files="$ac_clean_files conftest.out"
# Check that the compiler produces executables we can run.  If not, either
# the compiler is broken, or we cross compile.
{ $as_echo "$as_me:${as_lineno-$LINENO}: checking whether we are cross compiling" >&5
$as_echo_n "checking whether we are cross compiling... " >&6; }
if test "$cross_compiling" != yes; then
  { { ac_try="$ac_link"
case "(($ac_try" in
  *\"* | *\`* | *\\*) ac_try_echo=\$ac_try;;
  *) ac_try_echo=$ac_try;;
esac
eval ac_try_echo="\"\$as_me:${as_lineno-$LINENO}: $ac_try_echo\""
$as_echo "$ac_try_echo"; } >&5
  (eval "$ac_link") 2>&5
  ac_status=$?
  $as_echo "$as_me:${as_lineno-$LINENO}: \$? = $ac_status" >&5
  test $ac_status = 0; }
  if { ac_try='./conftest$ac_cv_exeext'
  { { case "(($ac_try" in
  *\"* | *\`* | *\\*) ac_try_echo=\$ac_try;;
  *) ac_try_echo=$ac_try;;
esac
eval ac_try_echo="\"\$as_me:${as_lineno-$LINENO}: $ac_try_echo\""
$as_echo "$ac_try_echo"; } >&5
  (eval "$ac_try") 2>&5
  ac_status=$?
  $as_echo "$as_me:${as_lineno-$LINENO}: \$? = $ac_status" >&5
  test $ac_status = 0; }; }; then
    cross_compiling=no
  else
    if test "$cross_compiling" = maybe; then
	cross_compiling=yes
    else
	{ { $as_echo "$as_me:${as_lineno-$LINENO}: error: in \`$ac_pwd':" >&5
$as_echo "$as_me: error: in \`$ac_pwd':" >&2;}
as_fn_error $? "cannot run C compiled programs.
If you meant to cross compile, use \`--host'.
See \`config.log' for more details" "$LINENO" 5; }
    fi
  fi
fi
{ $as_echo "$as_me:${as_lineno-$LINENO}: result: $cross_compiling" >&5
$as_echo "$cross_compiling" >&6; }

rm -f conftest.$ac_ext conftest$ac_cv_exeext conftest.out
ac_clean_files=$ac_clean_files_save
{ $as_echo "$as_me:${as_lineno-$LINENO}: checking for suffix of object files" >&5
$as_echo_n "checking for suffix of object files... " >&6; }
if ${ac_cv_objext+:} false; then :
  $as_echo_n "(cached) " >&6
else
  cat confdefs.h - <<_ACEOF >conftest.$ac_ext
/* end confdefs.h.  */

int
main ()
{

  ;
  return 0;
}
_ACEOF
rm -f conftest.o conftest.obj
if { { ac_try="$ac_compile"
case "(($ac_try" in
  *\"* | *\`* | *\\*) ac_try_echo=\$ac_try;;
  *) ac_try_echo=$ac_try;;
esac
eval ac_try_echo="\"\$as_me:${as_lineno-$LINENO}: $ac_try_echo\""
$as_echo "$ac_try_echo"; } >&5
  (eval "$ac_compile") 2>&5
  ac_status=$?
  $as_echo "$as_me:${as_lineno-$LINENO}: \$? = $ac_status" >&5
  test $ac_status = 0; }; then :
  for ac_file in conftest.o conftest.obj conftest.*; do
  test -f "$ac_file" || continue;
  case $ac_file in
    *.$ac_ext | *.xcoff | *.tds | *.d | *.pdb | *.xSYM | *.bb | *.bbg | *.map | *.inf | *.dSYM ) ;;
    *) ac_cv_objext=`expr "$ac_file" : '.*\.\(.*\)'`
       break;;
  esac
done
else
  $as_echo "$as_me: failed program was:" >&5
sed 's/^/| /' conftest.$ac_ext >&5

{ { $as_echo "$as_me:${as_lineno-$LINENO}: error: in \`$ac_pwd':" >&5
$as_echo "$as_me: error: in \`$ac_pwd':" >&2;}
as_fn_error $? "cannot compute suffix of object files: cannot compile
See \`config.log' for more details" "$LINENO" 5; }
fi
rm -f conftest.$ac_cv_objext conftest.$ac_ext
fi
{ $as_echo "$as_me:${as_lineno-$LINENO}: result: $ac_cv_objext" >&5
$as_echo "$ac_cv_objext" >&6; }
OBJEXT=$ac_cv_objext
ac_objext=$OBJEXT
{ $as_echo "$as_me:${as_lineno-$LINENO}: checking whether we are using the GNU C compiler" >&5
$as_echo_n "checking whether we are using the GNU C compiler... " >&6; }
if ${ac_cv_c_compiler_gnu+:} false; then :
  $as_echo_n "(cached) " >&6
else
  cat confdefs.h - <<_ACEOF >conftest.$ac_ext
/* end confdefs.h.  */

int
main ()
{
#ifndef __GNUC__
       choke me
#endif

  ;
  return 0;
}
_ACEOF
if ac_fn_c_try_compile "$LINENO"; then :
  ac_compiler_gnu=yes
else
  ac_compiler_gnu=no
fi
rm -f core conftest.err conftest.$ac_objext conftest.$ac_ext
ac_cv_c_compiler_gnu=$ac_compiler_gnu

fi
{ $as_echo "$as_me:${as_lineno-$LINENO}: result: $ac_cv_c_compiler_gnu" >&5
$as_echo "$ac_cv_c_compiler_gnu" >&6; }
if test $ac_compiler_gnu = yes; then
  GCC=yes
else
  GCC=
fi
ac_test_CFLAGS=${CFLAGS+set}
ac_save_CFLAGS=$CFLAGS
{ $as_echo "$as_me:${as_lineno-$LINENO}: checking whether $CC accepts -g" >&5
$as_echo_n "checking whether $CC accepts -g... " >&6; }
if ${ac_cv_prog_cc_g+:} false; then :
  $as_echo_n "(cached) " >&6
else
  ac_save_c_werror_flag=$ac_c_werror_flag
   ac_c_werror_flag=yes
   ac_cv_prog_cc_g=no
   CFLAGS="-g"
   cat confdefs.h - <<_ACEOF >conftest.$ac_ext
/* end confdefs.h.  */

int
main ()
{

  ;
  return 0;
}
_ACEOF
if ac_fn_c_try_compile "$LINENO"; then :
  ac_cv_prog_cc_g=yes
else
  CFLAGS=""
      cat confdefs.h - <<_ACEOF >conftest.$ac_ext
/* end confdefs.h.  */

int
main ()
{

  ;
  return 0;
}
_ACEOF
if ac_fn_c_try_compile "$LINENO"; then :

else
  ac_c_werror_flag=$ac_save_c_werror_flag
	 CFLAGS="-g"
	 cat confdefs.h - <<_ACEOF >conftest.$ac_ext
/* end confdefs.h.  */

int
main ()
{

  ;
  return 0;
}
_ACEOF
if ac_fn_c_try_compile "$LINENO"; then :
  ac_cv_prog_cc_g=yes
fi
rm -f core conftest.err conftest.$ac_objext conftest.$ac_ext
fi
rm -f core conftest.err conftest.$ac_objext conftest.$ac_ext
fi
rm -f core conftest.err conftest.$ac_objext conftest.$ac_ext
   ac_c_werror_flag=$ac_save_c_werror_flag
fi
{ $as_echo "$as_me:${as_lineno-$LINENO}: result: $ac_cv_prog_cc_g" >&5
$as_echo "$ac_cv_prog_cc_g" >&6; }
if test "$ac_test_CFLAGS" = set; then
  CFLAGS=$ac_save_CFLAGS
elif test $ac_cv_prog_cc_g = yes; then
  if test "$GCC" = yes; then
    CFLAGS="-g -O2"
  else
    CFLAGS="-g"
  fi
else
  if test "$GCC" = yes; then
    CFLAGS="-O2"
  else
    CFLAGS=
  fi
fi
{ $as_echo "$as_me:${as_lineno-$LINENO}: checking for $CC option to accept ISO C89" >&5
$as_echo_n "checking for $CC option to accept ISO C89... " >&6; }
if ${ac_cv_prog_cc_c89+:} false; then :
  $as_echo_n "(cached) " >&6
else
  ac_cv_prog_cc_c89=no
ac_save_CC=$CC
cat confdefs.h - <<_ACEOF >conftest.$ac_ext
/* end confdefs.h.  */
#include <stdarg.h>
#include <stdio.h>
struct stat;
/* Most of the following tests are stolen from RCS 5.7's src/conf.sh.  */
struct buf { int x; };
FILE * (*rcsopen) (struct buf *, struct stat *, int);
static char *e (p, i)
     char **p;
     int i;
{
  return p[i];
}
static char *f (char * (*g) (char **, int), char **p, ...)
{
  char *s;
  va_list v;
  va_start (v,p);
  s = g (p, va_arg (v,int));
  va_end (v);
  return s;
}

/* OSF 4.0 Compaq cc is some sort of almost-ANSI by default.  It has
   function prototypes and stuff, but not '\xHH' hex character constants.
   These don't provoke an error unfortunately, instead are silently treated
   as 'x'.  The following induces an error, until -std is added to get
   proper ANSI mode.  Curiously '\x00'!='x' always comes out true, for an
   array size at least.  It's necessary to write '\x00'==0 to get something
   that's true only with -std.  */
int osf4_cc_array ['\x00' == 0 ? 1 : -1];

/* IBM C 6 for AIX is almost-ANSI by default, but it replaces macro parameters
   inside strings and character constants.  */
#define FOO(x) 'x'
int xlc6_cc_array[FOO(a) == 'x' ? 1 : -1];

int test (int i, double x);
struct s1 {int (*f) (int a);};
struct s2 {int (*f) (double a);};
int pairnames (int, char **, FILE *(*)(struct buf *, struct stat *, int), int, int);
int argc;
char **argv;
int
main ()
{
return f (e, argv, 0) != argv[0]  ||  f (e, argv, 1) != argv[1];
  ;
  return 0;
}
_ACEOF
for ac_arg in '' -qlanglvl=extc89 -qlanglvl=ansi -std \
	-Ae "-Aa -D_HPUX_SOURCE" "-Xc -D__EXTENSIONS__"
do
  CC="$ac_save_CC $ac_arg"
  if ac_fn_c_try_compile "$LINENO"; then :
  ac_cv_prog_cc_c89=$ac_arg
fi
rm -f core conftest.err conftest.$ac_objext
  test "x$ac_cv_prog_cc_c89" != "xno" && break
done
rm -f conftest.$ac_ext
CC=$ac_save_CC

fi
# AC_CACHE_VAL
case "x$ac_cv_prog_cc_c89" in
  x)
    { $as_echo "$as_me:${as_lineno-$LINENO}: result: none needed" >&5
$as_echo "none needed" >&6; } ;;
  xno)
    { $as_echo "$as_me:${as_lineno-$LINENO}: result: unsupported" >&5
$as_echo "unsupported" >&6; } ;;
  *)
    CC="$CC $ac_cv_prog_cc_c89"
    { $as_echo "$as_me:${as_lineno-$LINENO}: result: $ac_cv_prog_cc_c89" >&5
$as_echo "$ac_cv_prog_cc_c89" >&6; } ;;
esac
if test "x$ac_cv_prog_cc_c89" != xno; then :

fi

ac_ext=c
ac_cpp='$CPP $CPPFLAGS'
ac_compile='$CC -c $CFLAGS $CPPFLAGS conftest.$ac_ext >&5'
ac_link='$CC -o conftest$ac_exeext $CFLAGS $CPPFLAGS $LDFLAGS conftest.$ac_ext $LIBS >&5'
ac_compiler_gnu=$ac_cv_c_compiler_gnu

ac_ext=cpp
ac_cpp='$CXXCPP $CPPFLAGS'
ac_compile='$CXX -c $CXXFLAGS $CPPFLAGS conftest.$ac_ext >&5'
ac_link='$CXX -o conftest$ac_exeext $CXXFLAGS $CPPFLAGS $LDFLAGS conftest.$ac_ext $LIBS >&5'
ac_compiler_gnu=$ac_cv_cxx_compiler_gnu
if test -z "$CXX"; then
  if test -n "$CCC"; then
    CXX=$CCC
  else
    if test -n "$ac_tool_prefix"; then
  for ac_prog in g++
  do
    # Extract the first word of "$ac_tool_prefix$ac_prog", so it can be a program name with args.
set dummy $ac_tool_prefix$ac_prog; ac_word=$2
{ $as_echo "$as_me:${as_lineno-$LINENO}: checking for $ac_word" >&5
$as_echo_n "checking for $ac_word... " >&6; }
if ${ac_cv_prog_CXX+:} false; then :
  $as_echo_n "(cached) " >&6
else
  if test -n "$CXX"; then
  ac_cv_prog_CXX="$CXX" # Let the user override the test.
else
as_save_IFS=$IFS; IFS=$PATH_SEPARATOR
for as_dir in $PATH
do
  IFS=$as_save_IFS
  test -z "$as_dir" && as_dir=.
    for ac_exec_ext in '' $ac_executable_extensions; do
  if as_fn_executable_p "$as_dir/$ac_word$ac_exec_ext"; then
    ac_cv_prog_CXX="$ac_tool_prefix$ac_prog"
    $as_echo "$as_me:${as_lineno-$LINENO}: found $as_dir/$ac_word$ac_exec_ext" >&5
    break 2
  fi
done
  done
IFS=$as_save_IFS

fi
fi
CXX=$ac_cv_prog_CXX
if test -n "$CXX"; then
  { $as_echo "$as_me:${as_lineno-$LINENO}: result: $CXX" >&5
$as_echo "$CXX" >&6; }
else
  { $as_echo "$as_me:${as_lineno-$LINENO}: result: no" >&5
$as_echo "no" >&6; }
fi


    test -n "$CXX" && break
  done
fi
if test -z "$CXX"; then
  ac_ct_CXX=$CXX
  for ac_prog in g++
do
  # Extract the first word of "$ac_prog", so it can be a program name with args.
set dummy $ac_prog; ac_word=$2
{ $as_echo "$as_me:${as_lineno-$LINENO}: checking for $ac_word" >&5
$as_echo_n "checking for $ac_word... " >&6; }
if ${ac_cv_prog_ac_ct_CXX+:} false; then :
  $as_echo_n "(cached) " >&6
else
  if test -n "$ac_ct_CXX"; then
  ac_cv_prog_ac_ct_CXX="$ac_ct_CXX" # Let the user override the test.
else
as_save_IFS=$IFS; IFS=$PATH_SEPARATOR
for as_dir in $PATH
do
  IFS=$as_save_IFS
  test -z "$as_dir" && as_dir=.
    for ac_exec_ext in '' $ac_executable_extensions; do
  if as_fn_executable_p "$as_dir/$ac_word$ac_exec_ext"; then
    ac_cv_prog_ac_ct_CXX="$ac_prog"
    $as_echo "$as_me:${as_lineno-$LINENO}: found $as_dir/$ac_word$ac_exec_ext" >&5
    break 2
  fi
done
  done
IFS=$as_save_IFS

fi
fi
ac_ct_CXX=$ac_cv_prog_ac_ct_CXX
if test -n "$ac_ct_CXX"; then
  { $as_echo "$as_me:${as_lineno-$LINENO}: result: $ac_ct_CXX" >&5
$as_echo "$ac_ct_CXX" >&6; }
else
  { $as_echo "$as_me:${as_lineno-$LINENO}: result: no" >&5
$as_echo "no" >&6; }
fi


  test -n "$ac_ct_CXX" && break
done

  if test "x$ac_ct_CXX" = x; then
    CXX="g++"
  else
    case $cross_compiling:$ac_tool_warned in
yes:)
{ $as_echo "$as_me:${as_lineno-$LINENO}: WARNING: using cross tools not prefixed with host triplet" >&5
$as_echo "$as_me: WARNING: using cross tools not prefixed with host triplet" >&2;}
ac_tool_warned=yes ;;
esac
    CXX=$ac_ct_CXX
  fi
fi

  fi
fi
# Provide some information about the compiler.
$as_echo "$as_me:${as_lineno-$LINENO}: checking for C++ compiler version" >&5
set X $ac_compile
ac_compiler=$2
for ac_option in --version -v -V -qversion; do
  { { ac_try="$ac_compiler $ac_option >&5"
case "(($ac_try" in
  *\"* | *\`* | *\\*) ac_try_echo=\$ac_try;;
  *) ac_try_echo=$ac_try;;
esac
eval ac_try_echo="\"\$as_me:${as_lineno-$LINENO}: $ac_try_echo\""
$as_echo "$ac_try_echo"; } >&5
  (eval "$ac_compiler $ac_option >&5") 2>conftest.err
  ac_status=$?
  if test -s conftest.err; then
    sed '10a\
... rest of stderr output deleted ...
         10q' conftest.err >conftest.er1
    cat conftest.er1 >&5
  fi
  rm -f conftest.er1 conftest.err
  $as_echo "$as_me:${as_lineno-$LINENO}: \$? = $ac_status" >&5
  test $ac_status = 0; }
done

{ $as_echo "$as_me:${as_lineno-$LINENO}: checking whether we are using the GNU C++ compiler" >&5
$as_echo_n "checking whether we are using the GNU C++ compiler... " >&6; }
if ${ac_cv_cxx_compiler_gnu+:} false; then :
  $as_echo_n "(cached) " >&6
else
  cat confdefs.h - <<_ACEOF >conftest.$ac_ext
/* end confdefs.h.  */

int
main ()
{
#ifndef __GNUC__
       choke me
#endif

  ;
  return 0;
}
_ACEOF
if ac_fn_cxx_try_compile "$LINENO"; then :
  ac_compiler_gnu=yes
else
  ac_compiler_gnu=no
fi
rm -f core conftest.err conftest.$ac_objext conftest.$ac_ext
ac_cv_cxx_compiler_gnu=$ac_compiler_gnu

fi
{ $as_echo "$as_me:${as_lineno-$LINENO}: result: $ac_cv_cxx_compiler_gnu" >&5
$as_echo "$ac_cv_cxx_compiler_gnu" >&6; }
if test $ac_compiler_gnu = yes; then
  GXX=yes
else
  GXX=
fi
ac_test_CXXFLAGS=${CXXFLAGS+set}
ac_save_CXXFLAGS=$CXXFLAGS
{ $as_echo "$as_me:${as_lineno-$LINENO}: checking whether $CXX accepts -g" >&5
$as_echo_n "checking whether $CXX accepts -g... " >&6; }
if ${ac_cv_prog_cxx_g+:} false; then :
  $as_echo_n "(cached) " >&6
else
  ac_save_cxx_werror_flag=$ac_cxx_werror_flag
   ac_cxx_werror_flag=yes
   ac_cv_prog_cxx_g=no
   CXXFLAGS="-g"
   cat confdefs.h - <<_ACEOF >conftest.$ac_ext
/* end confdefs.h.  */

int
main ()
{

  ;
  return 0;
}
_ACEOF
if ac_fn_cxx_try_compile "$LINENO"; then :
  ac_cv_prog_cxx_g=yes
else
  CXXFLAGS=""
      cat confdefs.h - <<_ACEOF >conftest.$ac_ext
/* end confdefs.h.  */

int
main ()
{

  ;
  return 0;
}
_ACEOF
if ac_fn_cxx_try_compile "$LINENO"; then :

else
  ac_cxx_werror_flag=$ac_save_cxx_werror_flag
	 CXXFLAGS="-g"
	 cat confdefs.h - <<_ACEOF >conftest.$ac_ext
/* end confdefs.h.  */

int
main ()
{

  ;
  return 0;
}
_ACEOF
if ac_fn_cxx_try_compile "$LINENO"; then :
  ac_cv_prog_cxx_g=yes
fi
rm -f core conftest.err conftest.$ac_objext conftest.$ac_ext
fi
rm -f core conftest.err conftest.$ac_objext conftest.$ac_ext
fi
rm -f core conftest.err conftest.$ac_objext conftest.$ac_ext
   ac_cxx_werror_flag=$ac_save_cxx_werror_flag
fi
{ $as_echo "$as_me:${as_lineno-$LINENO}: result: $ac_cv_prog_cxx_g" >&5
$as_echo "$ac_cv_prog_cxx_g" >&6; }
if test "$ac_test_CXXFLAGS" = set; then
  CXXFLAGS=$ac_save_CXXFLAGS
elif test $ac_cv_prog_cxx_g = yes; then
  if test "$GXX" = yes; then
    CXXFLAGS="-g -O2"
  else
    CXXFLAGS="-g"
  fi
else
  if test "$GXX" = yes; then
    CXXFLAGS="-O2"
  else
    CXXFLAGS=
  fi
fi
ac_ext=c
ac_cpp='$CPP $CPPFLAGS'
ac_compile='$CC -c $CFLAGS $CPPFLAGS conftest.$ac_ext >&5'
ac_link='$CC -o conftest$ac_exeext $CFLAGS $CPPFLAGS $LDFLAGS conftest.$ac_ext $LIBS >&5'
ac_compiler_gnu=$ac_cv_c_compiler_gnu


# Check whether --enable-largefile was given.
if test "${enable_largefile+set}" = set; then :
  enableval=$enable_largefile;
fi

if test "$enable_largefile" != no; then

  { $as_echo "$as_me:${as_lineno-$LINENO}: checking for special C compiler options needed for large files" >&5
$as_echo_n "checking for special C compiler options needed for large files... " >&6; }
if ${ac_cv_sys_largefile_CC+:} false; then :
  $as_echo_n "(cached) " >&6
else
  ac_cv_sys_largefile_CC=no
     if test "$GCC" != yes; then
       ac_save_CC=$CC
       while :; do
	 # IRIX 6.2 and later do not support large files by default,
	 # so use the C compiler's -n32 option if that helps.
	 cat confdefs.h - <<_ACEOF >conftest.$ac_ext
/* end confdefs.h.  */
#include <sys/types.h>
 /* Check that off_t can represent 2**63 - 1 correctly.
    We can't simply define LARGE_OFF_T to be 9223372036854775807,
    since some C++ compilers masquerading as C compilers
    incorrectly reject 9223372036854775807.  */
#define LARGE_OFF_T ((((off_t) 1 << 31) << 31) - 1 + (((off_t) 1 << 31) << 31))
  int off_t_is_large[(LARGE_OFF_T % 2147483629 == 721
		       && LARGE_OFF_T % 2147483647 == 1)
		      ? 1 : -1];
int
main ()
{

  ;
  return 0;
}
_ACEOF
	 if ac_fn_c_try_compile "$LINENO"; then :
  break
fi
rm -f core conftest.err conftest.$ac_objext
	 CC="$CC -n32"
	 if ac_fn_c_try_compile "$LINENO"; then :
  ac_cv_sys_largefile_CC=' -n32'; break
fi
rm -f core conftest.err conftest.$ac_objext
	 break
       done
       CC=$ac_save_CC
       rm -f conftest.$ac_ext
    fi
fi
{ $as_echo "$as_me:${as_lineno-$LINENO}: result: $ac_cv_sys_largefile_CC" >&5
$as_echo "$ac_cv_sys_largefile_CC" >&6; }
  if test "$ac_cv_sys_largefile_CC" != no; then
    CC=$CC$ac_cv_sys_largefile_CC
  fi

  { $as_echo "$as_me:${as_lineno-$LINENO}: checking for _FILE_OFFSET_BITS value needed for large files" >&5
$as_echo_n "checking for _FILE_OFFSET_BITS value needed for large files... " >&6; }
if ${ac_cv_sys_file_offset_bits+:} false; then :
  $as_echo_n "(cached) " >&6
else
  while :; do
  cat confdefs.h - <<_ACEOF >conftest.$ac_ext
/* end confdefs.h.  */
#include <sys/types.h>
 /* Check that off_t can represent 2**63 - 1 correctly.
    We can't simply define LARGE_OFF_T to be 9223372036854775807,
    since some C++ compilers masquerading as C compilers
    incorrectly reject 9223372036854775807.  */
#define LARGE_OFF_T ((((off_t) 1 << 31) << 31) - 1 + (((off_t) 1 << 31) << 31))
  int off_t_is_large[(LARGE_OFF_T % 2147483629 == 721
		       && LARGE_OFF_T % 2147483647 == 1)
		      ? 1 : -1];
int
main ()
{

  ;
  return 0;
}
_ACEOF
if ac_fn_c_try_compile "$LINENO"; then :
  ac_cv_sys_file_offset_bits=no; break
fi
rm -f core conftest.err conftest.$ac_objext conftest.$ac_ext
  cat confdefs.h - <<_ACEOF >conftest.$ac_ext
/* end confdefs.h.  */
#define _FILE_OFFSET_BITS 64
#include <sys/types.h>
 /* Check that off_t can represent 2**63 - 1 correctly.
    We can't simply define LARGE_OFF_T to be 9223372036854775807,
    since some C++ compilers masquerading as C compilers
    incorrectly reject 9223372036854775807.  */
#define LARGE_OFF_T ((((off_t) 1 << 31) << 31) - 1 + (((off_t) 1 << 31) << 31))
  int off_t_is_large[(LARGE_OFF_T % 2147483629 == 721
		       && LARGE_OFF_T % 2147483647 == 1)
		      ? 1 : -1];
int
main ()
{

  ;
  return 0;
}
_ACEOF
if ac_fn_c_try_compile "$LINENO"; then :
  ac_cv_sys_file_offset_bits=64; break
fi
rm -f core conftest.err conftest.$ac_objext conftest.$ac_ext
  ac_cv_sys_file_offset_bits=unknown
  break
done
fi
{ $as_echo "$as_me:${as_lineno-$LINENO}: result: $ac_cv_sys_file_offset_bits" >&5
$as_echo "$ac_cv_sys_file_offset_bits" >&6; }
case $ac_cv_sys_file_offset_bits in #(
  no | unknown) ;;
  *)
cat >>confdefs.h <<_ACEOF
#define _FILE_OFFSET_BITS $ac_cv_sys_file_offset_bits
_ACEOF
;;
esac
rm -rf conftest*
  if test $ac_cv_sys_file_offset_bits = unknown; then
    { $as_echo "$as_me:${as_lineno-$LINENO}: checking for _LARGE_FILES value needed for large files" >&5
$as_echo_n "checking for _LARGE_FILES value needed for large files... " >&6; }
if ${ac_cv_sys_large_files+:} false; then :
  $as_echo_n "(cached) " >&6
else
  while :; do
  cat confdefs.h - <<_ACEOF >conftest.$ac_ext
/* end confdefs.h.  */
#include <sys/types.h>
 /* Check that off_t can represent 2**63 - 1 correctly.
    We can't simply define LARGE_OFF_T to be 9223372036854775807,
    since some C++ compilers masquerading as C compilers
    incorrectly reject 9223372036854775807.  */
#define LARGE_OFF_T ((((off_t) 1 << 31) << 31) - 1 + (((off_t) 1 << 31) << 31))
  int off_t_is_large[(LARGE_OFF_T % 2147483629 == 721
		       && LARGE_OFF_T % 2147483647 == 1)
		      ? 1 : -1];
int
main ()
{

  ;
  return 0;
}
_ACEOF
if ac_fn_c_try_compile "$LINENO"; then :
  ac_cv_sys_large_files=no; break
fi
rm -f core conftest.err conftest.$ac_objext conftest.$ac_ext
  cat confdefs.h - <<_ACEOF >conftest.$ac_ext
/* end confdefs.h.  */
#define _LARGE_FILES 1
#include <sys/types.h>
 /* Check that off_t can represent 2**63 - 1 correctly.
    We can't simply define LARGE_OFF_T to be 9223372036854775807,
    since some C++ compilers masquerading as C compilers
    incorrectly reject 9223372036854775807.  */
#define LARGE_OFF_T ((((off_t) 1 << 31) << 31) - 1 + (((off_t) 1 << 31) << 31))
  int off_t_is_large[(LARGE_OFF_T % 2147483629 == 721
		       && LARGE_OFF_T % 2147483647 == 1)
		      ? 1 : -1];
int
main ()
{

  ;
  return 0;
}
_ACEOF
if ac_fn_c_try_compile "$LINENO"; then :
  ac_cv_sys_large_files=1; break
fi
rm -f core conftest.err conftest.$ac_objext conftest.$ac_ext
  ac_cv_sys_large_files=unknown
  break
done
fi
{ $as_echo "$as_me:${as_lineno-$LINENO}: result: $ac_cv_sys_large_files" >&5
$as_echo "$ac_cv_sys_large_files" >&6; }
case $ac_cv_sys_large_files in #(
  no | unknown) ;;
  *)
cat >>confdefs.h <<_ACEOF
#define _LARGE_FILES $ac_cv_sys_large_files
_ACEOF
;;
esac
rm -rf conftest*
  fi


fi

{ $as_echo "$as_me:${as_lineno-$LINENO}: checking for _LARGEFILE_SOURCE value needed for large files" >&5
$as_echo_n "checking for _LARGEFILE_SOURCE value needed for large files... " >&6; }
if ${ac_cv_sys_largefile_source+:} false; then :
  $as_echo_n "(cached) " >&6
else
  while :; do
  cat confdefs.h - <<_ACEOF >conftest.$ac_ext
/* end confdefs.h.  */
#include <sys/types.h> /* for off_t */
     #include <stdio.h>
int
main ()
{
int (*fp) (FILE *, off_t, int) = fseeko;
     return fseeko (stdin, 0, 0) && fp (stdin, 0, 0);
  ;
  return 0;
}
_ACEOF
if ac_fn_c_try_link "$LINENO"; then :
  ac_cv_sys_largefile_source=no; break
fi
rm -f core conftest.err conftest.$ac_objext \
    conftest$ac_exeext conftest.$ac_ext
  cat confdefs.h - <<_ACEOF >conftest.$ac_ext
/* end confdefs.h.  */
#define _LARGEFILE_SOURCE 1
#include <sys/types.h> /* for off_t */
     #include <stdio.h>
int
main ()
{
int (*fp) (FILE *, off_t, int) = fseeko;
     return fseeko (stdin, 0, 0) && fp (stdin, 0, 0);
  ;
  return 0;
}
_ACEOF
if ac_fn_c_try_link "$LINENO"; then :
  ac_cv_sys_largefile_source=1; break
fi
rm -f core conftest.err conftest.$ac_objext \
    conftest$ac_exeext conftest.$ac_ext
  ac_cv_sys_largefile_source=unknown
  break
done
fi
{ $as_echo "$as_me:${as_lineno-$LINENO}: result: $ac_cv_sys_largefile_source" >&5
$as_echo "$ac_cv_sys_largefile_source" >&6; }
case $ac_cv_sys_largefile_source in #(
  no | unknown) ;;
  *)
cat >>confdefs.h <<_ACEOF
#define _LARGEFILE_SOURCE $ac_cv_sys_largefile_source
_ACEOF
;;
esac
rm -rf conftest*

# We used to try defining _XOPEN_SOURCE=500 too, to work around a bug
# in glibc 2.1.3, but that breaks too many other things.
# If you want fseeko and ftello with glibc, upgrade to a fixed glibc.
if test $ac_cv_sys_largefile_source != unknown; then

$as_echo "#define HAVE_FSEEKO 1" >>confdefs.h

fi


# Checks for header files.
ac_ext=c
ac_cpp='$CPP $CPPFLAGS'
ac_compile='$CC -c $CFLAGS $CPPFLAGS conftest.$ac_ext >&5'
ac_link='$CC -o conftest$ac_exeext $CFLAGS $CPPFLAGS $LDFLAGS conftest.$ac_ext $LIBS >&5'
ac_compiler_gnu=$ac_cv_c_compiler_gnu
{ $as_echo "$as_me:${as_lineno-$LINENO}: checking how to run the C preprocessor" >&5
$as_echo_n "checking how to run the C preprocessor... " >&6; }
# On Suns, sometimes $CPP names a directory.
if test -n "$CPP" && test -d "$CPP"; then
  CPP=
fi
if test -z "$CPP"; then
  if ${ac_cv_prog_CPP+:} false; then :
  $as_echo_n "(cached) " >&6
else
      # Double quotes because CPP needs to be expanded
    for CPP in "$CC -E" "$CC -E -traditional-cpp" "/lib/cpp"
    do
      ac_preproc_ok=false
for ac_c_preproc_warn_flag in '' yes
do
  # Use a header file that comes with gcc, so configuring glibc
  # with a fresh cross-compiler works.
  # Prefer <limits.h> to <assert.h> if __STDC__ is defined, since
  # <limits.h> exists even on freestanding compilers.
  # On the NeXT, cc -E runs the code through the compiler's parser,
  # not just through cpp. "Syntax error" is here to catch this case.
  cat confdefs.h - <<_ACEOF >conftest.$ac_ext
/* end confdefs.h.  */
#ifdef __STDC__
# include <limits.h>
#else
# include <assert.h>
#endif
		     Syntax error
_ACEOF
if ac_fn_c_try_cpp "$LINENO"; then :

else
  # Broken: fails on valid input.
continue
fi
rm -f conftest.err conftest.i conftest.$ac_ext

  # OK, works on sane cases.  Now check whether nonexistent headers
  # can be detected and how.
  cat confdefs.h - <<_ACEOF >conftest.$ac_ext
/* end confdefs.h.  */
#include <ac_nonexistent.h>
_ACEOF
if ac_fn_c_try_cpp "$LINENO"; then :
  # Broken: success on invalid input.
continue
else
  # Passes both tests.
ac_preproc_ok=:
break
fi
rm -f conftest.err conftest.i conftest.$ac_ext

done
# Because of `break', _AC_PREPROC_IFELSE's cleaning code was skipped.
rm -f conftest.i conftest.err conftest.$ac_ext
if $ac_preproc_ok; then :
  break
fi

    done
    ac_cv_prog_CPP=$CPP

fi
  CPP=$ac_cv_prog_CPP
else
  ac_cv_prog_CPP=$CPP
fi
{ $as_echo "$as_me:${as_lineno-$LINENO}: result: $CPP" >&5
$as_echo "$CPP" >&6; }
ac_preproc_ok=false
for ac_c_preproc_warn_flag in '' yes
do
  # Use a header file that comes with gcc, so configuring glibc
  # with a fresh cross-compiler works.
  # Prefer <limits.h> to <assert.h> if __STDC__ is defined, since
  # <limits.h> exists even on freestanding compilers.
  # On the NeXT, cc -E runs the code through the compiler's parser,
  # not just through cpp. "Syntax error" is here to catch this case.
  cat confdefs.h - <<_ACEOF >conftest.$ac_ext
/* end confdefs.h.  */
#ifdef __STDC__
# include <limits.h>
#else
# include <assert.h>
#endif
		     Syntax error
_ACEOF
if ac_fn_c_try_cpp "$LINENO"; then :

else
  # Broken: fails on valid input.
continue
fi
rm -f conftest.err conftest.i conftest.$ac_ext

  # OK, works on sane cases.  Now check whether nonexistent headers
  # can be detected and how.
  cat confdefs.h - <<_ACEOF >conftest.$ac_ext
/* end confdefs.h.  */
#include <ac_nonexistent.h>
_ACEOF
if ac_fn_c_try_cpp "$LINENO"; then :
  # Broken: success on invalid input.
continue
else
  # Passes both tests.
ac_preproc_ok=:
break
fi
rm -f conftest.err conftest.i conftest.$ac_ext

done
# Because of `break', _AC_PREPROC_IFELSE's cleaning code was skipped.
rm -f conftest.i conftest.err conftest.$ac_ext
if $ac_preproc_ok; then :

else
  { { $as_echo "$as_me:${as_lineno-$LINENO}: error: in \`$ac_pwd':" >&5
$as_echo "$as_me: error: in \`$ac_pwd':" >&2;}
as_fn_error $? "C preprocessor \"$CPP\" fails sanity check
See \`config.log' for more details" "$LINENO" 5; }
fi

ac_ext=c
ac_cpp='$CPP $CPPFLAGS'
ac_compile='$CC -c $CFLAGS $CPPFLAGS conftest.$ac_ext >&5'
ac_link='$CC -o conftest$ac_exeext $CFLAGS $CPPFLAGS $LDFLAGS conftest.$ac_ext $LIBS >&5'
ac_compiler_gnu=$ac_cv_c_compiler_gnu


{ $as_echo "$as_me:${as_lineno-$LINENO}: checking for grep that handles long lines and -e" >&5
$as_echo_n "checking for grep that handles long lines and -e... " >&6; }
if ${ac_cv_path_GREP+:} false; then :
  $as_echo_n "(cached) " >&6
else
  if test -z "$GREP"; then
  ac_path_GREP_found=false
  # Loop through the user's path and test for each of PROGNAME-LIST
  as_save_IFS=$IFS; IFS=$PATH_SEPARATOR
for as_dir in $PATH$PATH_SEPARATOR/usr/xpg4/bin
do
  IFS=$as_save_IFS
  test -z "$as_dir" && as_dir=.
    for ac_prog in grep ggrep; do
    for ac_exec_ext in '' $ac_executable_extensions; do
      ac_path_GREP="$as_dir/$ac_prog$ac_exec_ext"
      as_fn_executable_p "$ac_path_GREP" || continue
# Check for GNU ac_path_GREP and select it if it is found.
  # Check for GNU $ac_path_GREP
case `"$ac_path_GREP" --version 2>&1` in
*GNU*)
  ac_cv_path_GREP="$ac_path_GREP" ac_path_GREP_found=:;;
*)
  ac_count=0
  $as_echo_n 0123456789 >"conftest.in"
  while :
  do
    cat "conftest.in" "conftest.in" >"conftest.tmp"
    mv "conftest.tmp" "conftest.in"
    cp "conftest.in" "conftest.nl"
    $as_echo 'GREP' >> "conftest.nl"
    "$ac_path_GREP" -e 'GREP$' -e '-(cannot match)-' < "conftest.nl" >"conftest.out" 2>/dev/null || break
    diff "conftest.out" "conftest.nl" >/dev/null 2>&1 || break
    as_fn_arith $ac_count + 1 && ac_count=$as_val
    if test $ac_count -gt ${ac_path_GREP_max-0}; then
      # Best one so far, save it but keep looking for a better one
      ac_cv_path_GREP="$ac_path_GREP"
      ac_path_GREP_max=$ac_count
    fi
    # 10*(2^10) chars as input seems more than enough
    test $ac_count -gt 10 && break
  done
  rm -f conftest.in conftest.tmp conftest.nl conftest.out;;
esac

      $ac_path_GREP_found && break 3
    done
  done
  done
IFS=$as_save_IFS
  if test -z "$ac_cv_path_GREP"; then
    as_fn_error $? "no acceptable grep could be found in $PATH$PATH_SEPARATOR/usr/xpg4/bin" "$LINENO" 5
  fi
else
  ac_cv_path_GREP=$GREP
fi

fi
{ $as_echo "$as_me:${as_lineno-$LINENO}: result: $ac_cv_path_GREP" >&5
$as_echo "$ac_cv_path_GREP" >&6; }
 GREP="$ac_cv_path_GREP"


{ $as_echo "$as_me:${as_lineno-$LINENO}: checking for egrep" >&5
$as_echo_n "checking for egrep... " >&6; }
if ${ac_cv_path_EGREP+:} false; then :
  $as_echo_n "(cached) " >&6
else
  if echo a | $GREP -E '(a|b)' >/dev/null 2>&1
   then ac_cv_path_EGREP="$GREP -E"
   else
     if test -z "$EGREP"; then
  ac_path_EGREP_found=false
  # Loop through the user's path and test for each of PROGNAME-LIST
  as_save_IFS=$IFS; IFS=$PATH_SEPARATOR
for as_dir in $PATH$PATH_SEPARATOR/usr/xpg4/bin
do
  IFS=$as_save_IFS
  test -z "$as_dir" && as_dir=.
    for ac_prog in egrep; do
    for ac_exec_ext in '' $ac_executable_extensions; do
      ac_path_EGREP="$as_dir/$ac_prog$ac_exec_ext"
      as_fn_executable_p "$ac_path_EGREP" || continue
# Check for GNU ac_path_EGREP and select it if it is found.
  # Check for GNU $ac_path_EGREP
case `"$ac_path_EGREP" --version 2>&1` in
*GNU*)
  ac_cv_path_EGREP="$ac_path_EGREP" ac_path_EGREP_found=:;;
*)
  ac_count=0
  $as_echo_n 0123456789 >"conftest.in"
  while :
  do
    cat "conftest.in" "conftest.in" >"conftest.tmp"
    mv "conftest.tmp" "conftest.in"
    cp "conftest.in" "conftest.nl"
    $as_echo 'EGREP' >> "conftest.nl"
    "$ac_path_EGREP" 'EGREP$' < "conftest.nl" >"conftest.out" 2>/dev/null || break
    diff "conftest.out" "conftest.nl" >/dev/null 2>&1 || break
    as_fn_arith $ac_count + 1 && ac_count=$as_val
    if test $ac_count -gt ${ac_path_EGREP_max-0}; then
      # Best one so far, save it but keep looking for a better one
      ac_cv_path_EGREP="$ac_path_EGREP"
      ac_path_EGREP_max=$ac_count
    fi
    # 10*(2^10) chars as input seems more than enough
    test $ac_count -gt 10 && break
  done
  rm -f conftest.in conftest.tmp conftest.nl conftest.out;;
esac

      $ac_path_EGREP_found && break 3
    done
  done
  done
IFS=$as_save_IFS
  if test -z "$ac_cv_path_EGREP"; then
    as_fn_error $? "no acceptable egrep could be found in $PATH$PATH_SEPARATOR/usr/xpg4/bin" "$LINENO" 5
  fi
else
  ac_cv_path_EGREP=$EGREP
fi

   fi
fi
{ $as_echo "$as_me:${as_lineno-$LINENO}: result: $ac_cv_path_EGREP" >&5
$as_echo "$ac_cv_path_EGREP" >&6; }
 EGREP="$ac_cv_path_EGREP"


{ $as_echo "$as_me:${as_lineno-$LINENO}: checking for ANSI C header files" >&5
$as_echo_n "checking for ANSI C header files... " >&6; }
if ${ac_cv_header_stdc+:} false; then :
  $as_echo_n "(cached) " >&6
else
  cat confdefs.h - <<_ACEOF >conftest.$ac_ext
/* end confdefs.h.  */
#include <stdlib.h>
#include <stdarg.h>
#include <string.h>
#include <float.h>

int
main ()
{

  ;
  return 0;
}
_ACEOF
if ac_fn_c_try_compile "$LINENO"; then :
  ac_cv_header_stdc=yes
else
  ac_cv_header_stdc=no
fi
rm -f core conftest.err conftest.$ac_objext conftest.$ac_ext

if test $ac_cv_header_stdc = yes; then
  # SunOS 4.x string.h does not declare mem*, contrary to ANSI.
  cat confdefs.h - <<_ACEOF >conftest.$ac_ext
/* end confdefs.h.  */
#include <string.h>

_ACEOF
if (eval "$ac_cpp conftest.$ac_ext") 2>&5 |
  $EGREP "memchr" >/dev/null 2>&1; then :

else
  ac_cv_header_stdc=no
fi
rm -f conftest*

fi

if test $ac_cv_header_stdc = yes; then
  # ISC 2.0.2 stdlib.h does not declare free, contrary to ANSI.
  cat confdefs.h - <<_ACEOF >conftest.$ac_ext
/* end confdefs.h.  */
#include <stdlib.h>

_ACEOF
if (eval "$ac_cpp conftest.$ac_ext") 2>&5 |
  $EGREP "free" >/dev/null 2>&1; then :

else
  ac_cv_header_stdc=no
fi
rm -f conftest*

fi

if test $ac_cv_header_stdc = yes; then
  # /bin/cc in Irix-4.0.5 gets non-ANSI ctype macros unless using -ansi.
  if test "$cross_compiling" = yes; then :
  :
else
  cat confdefs.h - <<_ACEOF >conftest.$ac_ext
/* end confdefs.h.  */
#include <ctype.h>
#include <stdlib.h>
#if ((' ' & 0x0FF) == 0x020)
# define ISLOWER(c) ('a' <= (c) && (c) <= 'z')
# define TOUPPER(c) (ISLOWER(c) ? 'A' + ((c) - 'a') : (c))
#else
# define ISLOWER(c) \
		   (('a' <= (c) && (c) <= 'i') \
		     || ('j' <= (c) && (c) <= 'r') \
		     || ('s' <= (c) && (c) <= 'z'))
# define TOUPPER(c) (ISLOWER(c) ? ((c) | 0x40) : (c))
#endif

#define XOR(e, f) (((e) && !(f)) || (!(e) && (f)))
int
main ()
{
  int i;
  for (i = 0; i < 256; i++)
    if (XOR (islower (i), ISLOWER (i))
	|| toupper (i) != TOUPPER (i))
      return 2;
  return 0;
}
_ACEOF
if ac_fn_c_try_run "$LINENO"; then :

else
  ac_cv_header_stdc=no
fi
rm -f core *.core core.conftest.* gmon.out bb.out conftest$ac_exeext \
  conftest.$ac_objext conftest.beam conftest.$ac_ext
fi

fi
fi
{ $as_echo "$as_me:${as_lineno-$LINENO}: result: $ac_cv_header_stdc" >&5
$as_echo "$ac_cv_header_stdc" >&6; }
if test $ac_cv_header_stdc = yes; then

$as_echo "#define STDC_HEADERS 1" >>confdefs.h

fi

# On IRIX 5.3, sys/types and inttypes.h are conflicting.
for ac_header in sys/types.h sys/stat.h stdlib.h string.h memory.h strings.h \
		  inttypes.h stdint.h unistd.h
do :
  as_ac_Header=`$as_echo "ac_cv_header_$ac_header" | $as_tr_sh`
ac_fn_c_check_header_compile "$LINENO" "$ac_header" "$as_ac_Header" "$ac_includes_default
"
if eval test \"x\$"$as_ac_Header"\" = x"yes"; then :
  cat >>confdefs.h <<_ACEOF
#define `$as_echo "HAVE_$ac_header" | $as_tr_cpp` 1
_ACEOF

fi

done


for ac_header in float.h inttypes.h stdint.h stdlib.h string.h sys/time.h unistd.h execinfo.h
do :
  as_ac_Header=`$as_echo "ac_cv_header_$ac_header" | $as_tr_sh`
ac_fn_c_check_header_mongrel "$LINENO" "$ac_header" "$as_ac_Header" "$ac_includes_default"
if eval test \"x\$"$as_ac_Header"\" = x"yes"; then :
  cat >>confdefs.h <<_ACEOF
#define `$as_echo "HAVE_$ac_header" | $as_tr_cpp` 1
_ACEOF

fi

done


# Checks for typedefs, structures, and compiler characteristics.
{ $as_echo "$as_me:${as_lineno-$LINENO}: checking for stdbool.h that conforms to C99" >&5
$as_echo_n "checking for stdbool.h that conforms to C99... " >&6; }
if ${ac_cv_header_stdbool_h+:} false; then :
  $as_echo_n "(cached) " >&6
else
  cat confdefs.h - <<_ACEOF >conftest.$ac_ext
/* end confdefs.h.  */

             #include <stdbool.h>
             #ifndef bool
              "error: bool is not defined"
             #endif
             #ifndef false
              "error: false is not defined"
             #endif
             #if false
              "error: false is not 0"
             #endif
             #ifndef true
              "error: true is not defined"
             #endif
             #if true != 1
              "error: true is not 1"
             #endif
             #ifndef __bool_true_false_are_defined
              "error: __bool_true_false_are_defined is not defined"
             #endif

             struct s { _Bool s: 1; _Bool t; } s;

             char a[true == 1 ? 1 : -1];
             char b[false == 0 ? 1 : -1];
             char c[__bool_true_false_are_defined == 1 ? 1 : -1];
             char d[(bool) 0.5 == true ? 1 : -1];
             /* See body of main program for 'e'.  */
             char f[(_Bool) 0.0 == false ? 1 : -1];
             char g[true];
             char h[sizeof (_Bool)];
             char i[sizeof s.t];
             enum { j = false, k = true, l = false * true, m = true * 256 };
             /* The following fails for
                HP aC++/ANSI C B3910B A.05.55 [Dec 04 2003]. */
             _Bool n[m];
             char o[sizeof n == m * sizeof n[0] ? 1 : -1];
             char p[-1 - (_Bool) 0 < 0 && -1 - (bool) 0 < 0 ? 1 : -1];
             /* Catch a bug in an HP-UX C compiler.  See
                http://gcc.gnu.org/ml/gcc-patches/2003-12/msg02303.html
                http://lists.gnu.org/archive/html/bug-coreutils/2005-11/msg00161.html
              */
             _Bool q = true;
             _Bool *pq = &q;

int
main ()
{

             bool e = &s;
             *pq |= q;
             *pq |= ! q;
             /* Refer to every declared value, to avoid compiler optimizations.  */
             return (!a + !b + !c + !d + !e + !f + !g + !h + !i + !!j + !k + !!l
                     + !m + !n + !o + !p + !q + !pq);

  ;
  return 0;
}
_ACEOF
if ac_fn_c_try_compile "$LINENO"; then :
  ac_cv_header_stdbool_h=yes
else
  ac_cv_header_stdbool_h=no
fi
rm -f core conftest.err conftest.$ac_objext conftest.$ac_ext
fi
{ $as_echo "$as_me:${as_lineno-$LINENO}: result: $ac_cv_header_stdbool_h" >&5
$as_echo "$ac_cv_header_stdbool_h" >&6; }
   ac_fn_c_check_type "$LINENO" "_Bool" "ac_cv_type__Bool" "$ac_includes_default"
if test "x$ac_cv_type__Bool" = xyes; then :

cat >>confdefs.h <<_ACEOF
#define HAVE__BOOL 1
_ACEOF


fi


{ $as_echo "$as_me:${as_lineno-$LINENO}: checking for inline" >&5
$as_echo_n "checking for inline... " >&6; }
if ${ac_cv_c_inline+:} false; then :
  $as_echo_n "(cached) " >&6
else
  ac_cv_c_inline=no
for ac_kw in inline __inline__ __inline; do
  cat confdefs.h - <<_ACEOF >conftest.$ac_ext
/* end confdefs.h.  */
#ifndef __cplusplus
typedef int foo_t;
static $ac_kw foo_t static_foo () {return 0; }
$ac_kw foo_t foo () {return 0; }
#endif

_ACEOF
if ac_fn_c_try_compile "$LINENO"; then :
  ac_cv_c_inline=$ac_kw
fi
rm -f core conftest.err conftest.$ac_objext conftest.$ac_ext
  test "$ac_cv_c_inline" != no && break
done

fi
{ $as_echo "$as_me:${as_lineno-$LINENO}: result: $ac_cv_c_inline" >&5
$as_echo "$ac_cv_c_inline" >&6; }

case $ac_cv_c_inline in
  inline | yes) ;;
  *)
    case $ac_cv_c_inline in
      no) ac_val=;;
      *) ac_val=$ac_cv_c_inline;;
    esac
    cat >>confdefs.h <<_ACEOF
#ifndef __cplusplus
#define inline $ac_val
#endif
_ACEOF
    ;;
esac

ac_fn_c_find_intX_t "$LINENO" "32" "ac_cv_c_int32_t"
case $ac_cv_c_int32_t in #(
  no|yes) ;; #(
  *)

cat >>confdefs.h <<_ACEOF
#define int32_t $ac_cv_c_int32_t
_ACEOF
;;
esac

ac_fn_c_check_type "$LINENO" "size_t" "ac_cv_type_size_t" "$ac_includes_default"
if test "x$ac_cv_type_size_t" = xyes; then :

else

cat >>confdefs.h <<_ACEOF
#define size_t unsigned int
_ACEOF

fi

ac_fn_c_check_type "$LINENO" "ssize_t" "ac_cv_type_ssize_t" "$ac_includes_default"
if test "x$ac_cv_type_ssize_t" = xyes; then :

else

cat >>confdefs.h <<_ACEOF
#define ssize_t int
_ACEOF

fi

ac_fn_c_find_uintX_t "$LINENO" "32" "ac_cv_c_uint32_t"
case $ac_cv_c_uint32_t in #(
  no|yes) ;; #(
  *)

$as_echo "#define _UINT32_T 1" >>confdefs.h


cat >>confdefs.h <<_ACEOF
#define uint32_t $ac_cv_c_uint32_t
_ACEOF
;;
  esac

ac_fn_c_find_uintX_t "$LINENO" "64" "ac_cv_c_uint64_t"
case $ac_cv_c_uint64_t in #(
  no|yes) ;; #(
  *)

$as_echo "#define _UINT64_T 1" >>confdefs.h


cat >>confdefs.h <<_ACEOF
#define uint64_t $ac_cv_c_uint64_t
_ACEOF
;;
  esac

ac_fn_c_find_uintX_t "$LINENO" "8" "ac_cv_c_uint8_t"
case $ac_cv_c_uint8_t in #(
  no|yes) ;; #(
  *)

$as_echo "#define _UINT8_T 1" >>confdefs.h


cat >>confdefs.h <<_ACEOF
#define uint8_t $ac_cv_c_uint8_t
_ACEOF
;;
  esac


# Checks for library functions.
for ac_header in stdlib.h
do :
  ac_fn_c_check_header_mongrel "$LINENO" "stdlib.h" "ac_cv_header_stdlib_h" "$ac_includes_default"
if test "x$ac_cv_header_stdlib_h" = xyes; then :
  cat >>confdefs.h <<_ACEOF
#define HAVE_STDLIB_H 1
_ACEOF

fi

done

{ $as_echo "$as_me:${as_lineno-$LINENO}: checking for GNU libc compatible malloc" >&5
$as_echo_n "checking for GNU libc compatible malloc... " >&6; }
if ${ac_cv_func_malloc_0_nonnull+:} false; then :
  $as_echo_n "(cached) " >&6
else
  if test "$cross_compiling" = yes; then :
  ac_cv_func_malloc_0_nonnull=no
else
  cat confdefs.h - <<_ACEOF >conftest.$ac_ext
/* end confdefs.h.  */
#if defined STDC_HEADERS || defined HAVE_STDLIB_H
# include <stdlib.h>
#else
char *malloc ();
#endif

int
main ()
{
return ! malloc (0);
  ;
  return 0;
}
_ACEOF
if ac_fn_c_try_run "$LINENO"; then :
  ac_cv_func_malloc_0_nonnull=yes
else
  ac_cv_func_malloc_0_nonnull=no
fi
rm -f core *.core core.conftest.* gmon.out bb.out conftest$ac_exeext \
  conftest.$ac_objext conftest.beam conftest.$ac_ext
fi

fi
{ $as_echo "$as_me:${as_lineno-$LINENO}: result: $ac_cv_func_malloc_0_nonnull" >&5
$as_echo "$ac_cv_func_malloc_0_nonnull" >&6; }
if test $ac_cv_func_malloc_0_nonnull = yes; then :

$as_echo "#define HAVE_MALLOC 1" >>confdefs.h

else
  $as_echo "#define HAVE_MALLOC 0" >>confdefs.h

   case " $LIBOBJS " in
  *" malloc.$ac_objext "* ) ;;
  *) LIBOBJS="$LIBOBJS malloc.$ac_objext"
 ;;
esac


$as_echo "#define malloc rpl_malloc" >>confdefs.h

fi


for ac_func in gettimeofday memset sqrt strerror
do :
  as_ac_var=`$as_echo "ac_cv_func_$ac_func" | $as_tr_sh`
ac_fn_c_check_func "$LINENO" "$ac_func" "$as_ac_var"
if eval test \"x\$"$as_ac_var"\" = x"yes"; then :
  cat >>confdefs.h <<_ACEOF
#define `$as_echo "HAVE_$ac_func" | $as_tr_cpp` 1
_ACEOF

fi
done


# Checks for libraries.

zlib_devel=ok
ac_fn_c_check_header_compile "$LINENO" "zlib.h" "ac_cv_header_zlib_h" ";
"
if test "x$ac_cv_header_zlib_h" = xyes; then :

else
  zlib_devel=missing
fi


{ $as_echo "$as_me:${as_lineno-$LINENO}: checking for inflate in -lz" >&5
$as_echo_n "checking for inflate in -lz... " >&6; }
if ${ac_cv_lib_z_inflate+:} false; then :
  $as_echo_n "(cached) " >&6
else
  ac_check_lib_save_LIBS=$LIBS
LIBS="-lz  $LIBS"
cat confdefs.h - <<_ACEOF >conftest.$ac_ext
/* end confdefs.h.  */

/* Override any GCC internal prototype to avoid an error.
   Use char because int might match the return type of a GCC
   builtin and then its argument prototype would still apply.  */
#ifdef __cplusplus
extern "C"
#endif
char inflate ();
int
main ()
{
return inflate ();
  ;
  return 0;
}
_ACEOF
if ac_fn_c_try_link "$LINENO"; then :
  ac_cv_lib_z_inflate=yes
else
  ac_cv_lib_z_inflate=no
fi
rm -f core conftest.err conftest.$ac_objext \
    conftest$ac_exeext conftest.$ac_ext
LIBS=$ac_check_lib_save_LIBS
fi
{ $as_echo "$as_me:${as_lineno-$LINENO}: result: $ac_cv_lib_z_inflate" >&5
$as_echo "$ac_cv_lib_z_inflate" >&6; }
if test "x$ac_cv_lib_z_inflate" = xyes; then :
  cat >>confdefs.h <<_ACEOF
#define HAVE_LIBZ 1
_ACEOF

  LIBS="-lz $LIBS"

else
  zlib_devel=missing
fi


if test $zlib_devel != ok; then
  cat > config.mk <<'EOF'
ifneq ($(MAKECMDGOALS),distclean)
$(error Resolve configure error first)
endif
EOF
   as_fn_error $? "zlib development files not found
	you may need to ensure a package such as zlib1g-dev (on Debian or
Ubuntu Linux) or zlib-devel (on RPM-based Linux distributions or Cygwin)
is installed.
" "$LINENO" 5
fi

hdf_tar_install=autoconf
hts_tar_install=autoconf

locallibhts=no
if test "$enable_systemhts" = yes; then
	libhts=yes
	for ac_header in htslib/hts.h
do :
  ac_fn_c_check_header_mongrel "$LINENO" "htslib/hts.h" "ac_cv_header_htslib_hts_h" "$ac_includes_default"
if test "x$ac_cv_header_htslib_hts_h" = xyes; then :
  cat >>confdefs.h <<_ACEOF
#define HAVE_HTSLIB_HTS_H 1
_ACEOF

else
  libhts=no
fi

done

	{ $as_echo "$as_me:${as_lineno-$LINENO}: checking for faidx_fetch_seq in -lhts" >&5
$as_echo_n "checking for faidx_fetch_seq in -lhts... " >&6; }
if ${ac_cv_lib_hts_faidx_fetch_seq+:} false; then :
  $as_echo_n "(cached) " >&6
else
  ac_check_lib_save_LIBS=$LIBS
LIBS="-lhts  $LIBS"
cat confdefs.h - <<_ACEOF >conftest.$ac_ext
/* end confdefs.h.  */

/* Override any GCC internal prototype to avoid an error.
   Use char because int might match the return type of a GCC
   builtin and then its argument prototype would still apply.  */
#ifdef __cplusplus
extern "C"
#endif
char faidx_fetch_seq ();
int
main ()
{
return faidx_fetch_seq ();
  ;
  return 0;
}
_ACEOF
if ac_fn_c_try_link "$LINENO"; then :
  ac_cv_lib_hts_faidx_fetch_seq=yes
else
  ac_cv_lib_hts_faidx_fetch_seq=no
fi
rm -f core conftest.err conftest.$ac_objext \
    conftest$ac_exeext conftest.$ac_ext
LIBS=$ac_check_lib_save_LIBS
fi
{ $as_echo "$as_me:${as_lineno-$LINENO}: result: $ac_cv_lib_hts_faidx_fetch_seq" >&5
$as_echo "$ac_cv_lib_hts_faidx_fetch_seq" >&6; }
if test "x$ac_cv_lib_hts_faidx_fetch_seq" = xyes; then :
  cat >>confdefs.h <<_ACEOF
#define HAVE_LIBHTS 1
_ACEOF

  LIBS="-lhts $LIBS"

else
  libhts=no
fi

	if test $libhts != yes; then
		cat > config.mk <<'EOF'
ifneq ($(MAKECMDGOALS),distclean)
$(error Resolve configure error first)
endif
EOF
   as_fn_error $? "System wide HTS library development files not found!
		  On Debian based systems such as Ubuntu you may use sudo apt-get install libhts-dev.
		  You can also download and install the latest release of htslib from http://www.htslib.org/download/.
		  Or else locally install through ./scripts/install-hts.sh and run ./configure without --enable_systemhts.
		  " "$LINENO" 5
	fi
	libhtsnew=yes
	{ $as_echo "$as_me:${as_lineno-$LINENO}: checking for fai_load3 in -lhts" >&5
$as_echo_n "checking for fai_load3 in -lhts... " >&6; }
if ${ac_cv_lib_hts_fai_load3+:} false; then :
  $as_echo_n "(cached) " >&6
else
  ac_check_lib_save_LIBS=$LIBS
LIBS="-lhts  $LIBS"
cat confdefs.h - <<_ACEOF >conftest.$ac_ext
/* end confdefs.h.  */

/* Override any GCC internal prototype to avoid an error.
   Use char because int might match the return type of a GCC
   builtin and then its argument prototype would still apply.  */
#ifdef __cplusplus
extern "C"
#endif
char fai_load3 ();
int
main ()
{
return fai_load3 ();
  ;
  return 0;
}
_ACEOF
if ac_fn_c_try_link "$LINENO"; then :
  ac_cv_lib_hts_fai_load3=yes
else
  ac_cv_lib_hts_fai_load3=no
fi
rm -f core conftest.err conftest.$ac_objext \
    conftest$ac_exeext conftest.$ac_ext
LIBS=$ac_check_lib_save_LIBS
fi
{ $as_echo "$as_me:${as_lineno-$LINENO}: result: $ac_cv_lib_hts_fai_load3" >&5
$as_echo "$ac_cv_lib_hts_fai_load3" >&6; }
if test "x$ac_cv_lib_hts_fai_load3" = xyes; then :
  cat >>confdefs.h <<_ACEOF
#define HAVE_LIBHTS 1
_ACEOF

  LIBS="-lhts $LIBS"

else
  libhtsnew=no
fi

	if test $libhtsnew != yes; then
		cat > config.mk <<'EOF'
ifneq ($(MAKECMDGOALS),distclean)
$(error Resolve configure error first)
endif
EOF
   as_fn_error $? "System wide HTS library development files are old and does not support fai_load3!
		  You can also download and install the latest release of htslib from http://www.htslib.org/download/.
		  Or else locally install through ./scripts/install-hts.sh and run ./configure without --enable_systemhts.
		  " "$LINENO" 5
	fi

else
	libhts=yes
	{ $as_echo "$as_me:${as_lineno-$LINENO}: checking for ./htslib/htslib/hts.h" >&5
$as_echo_n "checking for ./htslib/htslib/hts.h... " >&6; }
if ${ac_cv_file___htslib_htslib_hts_h+:} false; then :
  $as_echo_n "(cached) " >&6
else
  test "$cross_compiling" = yes &&
  as_fn_error $? "cannot check for file existence when cross compiling" "$LINENO" 5
if test -r "./htslib/htslib/hts.h"; then
  ac_cv_file___htslib_htslib_hts_h=yes
else
  ac_cv_file___htslib_htslib_hts_h=no
fi
fi
{ $as_echo "$as_me:${as_lineno-$LINENO}: result: $ac_cv_file___htslib_htslib_hts_h" >&5
$as_echo "$ac_cv_file___htslib_htslib_hts_h" >&6; }
if test "x$ac_cv_file___htslib_htslib_hts_h" = xyes; then :

else
  libhts=no
fi

	{ $as_echo "$as_me:${as_lineno-$LINENO}: checking for ./htslib/libhts.a" >&5
$as_echo_n "checking for ./htslib/libhts.a... " >&6; }
if ${ac_cv_file___htslib_libhts_a+:} false; then :
  $as_echo_n "(cached) " >&6
else
  test "$cross_compiling" = yes &&
  as_fn_error $? "cannot check for file existence when cross compiling" "$LINENO" 5
if test -r "./htslib/libhts.a"; then
  ac_cv_file___htslib_libhts_a=yes
else
  ac_cv_file___htslib_libhts_a=no
fi
fi
{ $as_echo "$as_me:${as_lineno-$LINENO}: result: $ac_cv_file___htslib_libhts_a" >&5
$as_echo "$ac_cv_file___htslib_libhts_a" >&6; }
if test "x$ac_cv_file___htslib_libhts_a" = xyes; then :

else
  libhts=no
fi

	if test $libhts != yes; then
		cat > config.mk <<'EOF'
ifneq ($(MAKECMDGOALS),distclean)
$(error Resolve configure error first)
endif
EOF
   as_fn_error $? "Could not find a working local hts installation in ./htslib!
		Make sure you run ./scripts/install-hts.sh
		" "$LINENO" 5
	fi
	locallibhts=yes
fi


locallibhdf5=no
if test "$enable_localhdf5" = yes; then
	libhdf=yes
	{ $as_echo "$as_me:${as_lineno-$LINENO}: checking for ./hdf5/include/hdf5.h" >&5
$as_echo_n "checking for ./hdf5/include/hdf5.h... " >&6; }
if ${ac_cv_file___hdf5_include_hdf5_h+:} false; then :
  $as_echo_n "(cached) " >&6
else
  test "$cross_compiling" = yes &&
  as_fn_error $? "cannot check for file existence when cross compiling" "$LINENO" 5
if test -r "./hdf5/include/hdf5.h"; then
  ac_cv_file___hdf5_include_hdf5_h=yes
else
  ac_cv_file___hdf5_include_hdf5_h=no
fi
fi
{ $as_echo "$as_me:${as_lineno-$LINENO}: result: $ac_cv_file___hdf5_include_hdf5_h" >&5
$as_echo "$ac_cv_file___hdf5_include_hdf5_h" >&6; }
if test "x$ac_cv_file___hdf5_include_hdf5_h" = xyes; then :

cat >>confdefs.h <<_ACEOF
#define HAVE___HDF5_INCLUDE_HDF5_H 1
_ACEOF

else
  libhdf=no
fi

	{ $as_echo "$as_me:${as_lineno-$LINENO}: checking for ./hdf5/lib/libhdf5.a" >&5
$as_echo_n "checking for ./hdf5/lib/libhdf5.a... " >&6; }
if ${ac_cv_file___hdf5_lib_libhdf5_a+:} false; then :
  $as_echo_n "(cached) " >&6
else
  test "$cross_compiling" = yes &&
  as_fn_error $? "cannot check for file existence when cross compiling" "$LINENO" 5
if test -r "./hdf5/lib/libhdf5.a"; then
  ac_cv_file___hdf5_lib_libhdf5_a=yes
else
  ac_cv_file___hdf5_lib_libhdf5_a=no
fi
fi
{ $as_echo "$as_me:${as_lineno-$LINENO}: result: $ac_cv_file___hdf5_lib_libhdf5_a" >&5
$as_echo "$ac_cv_file___hdf5_lib_libhdf5_a" >&6; }
if test "x$ac_cv_file___hdf5_lib_libhdf5_a" = xyes; then :

else
  libhdf=no
fi

	if test $libhdf != yes; then
	  cat > config.mk <<'EOF'
ifneq ($(MAKECMDGOALS),distclean)
$(error Resolve configure error first)
endif
EOF
   as_fn_error $? "local HDF5 library development files not found!
		Make sure you run ./scripts/install-hdf5.sh
	  " "$LINENO" 5
	fi
	locallibhdf5=yes
else
	libhdf=yes
	{ $as_echo "$as_me:${as_lineno-$LINENO}: checking for library containing H5Fopen" >&5
$as_echo_n "checking for library containing H5Fopen... " >&6; }
if ${ac_cv_search_H5Fopen+:} false; then :
  $as_echo_n "(cached) " >&6
else
  ac_func_search_save_LIBS=$LIBS
cat confdefs.h - <<_ACEOF >conftest.$ac_ext
/* end confdefs.h.  */

/* Override any GCC internal prototype to avoid an error.
   Use char because int might match the return type of a GCC
   builtin and then its argument prototype would still apply.  */
#ifdef __cplusplus
extern "C"
#endif
char H5Fopen ();
int
main ()
{
return H5Fopen ();
  ;
  return 0;
}
_ACEOF
for ac_lib in '' hdf5 hdf5_serial; do
  if test -z "$ac_lib"; then
    ac_res="none required"
  else
    ac_res=-l$ac_lib
    LIBS="-l$ac_lib  $ac_func_search_save_LIBS"
  fi
  if ac_fn_c_try_link "$LINENO"; then :
  ac_cv_search_H5Fopen=$ac_res
fi
rm -f core conftest.err conftest.$ac_objext \
    conftest$ac_exeext
  if ${ac_cv_search_H5Fopen+:} false; then :
  break
fi
done
if ${ac_cv_search_H5Fopen+:} false; then :

else
  ac_cv_search_H5Fopen=no
fi
rm conftest.$ac_ext
LIBS=$ac_func_search_save_LIBS
fi
{ $as_echo "$as_me:${as_lineno-$LINENO}: result: $ac_cv_search_H5Fopen" >&5
$as_echo "$ac_cv_search_H5Fopen" >&6; }
ac_res=$ac_cv_search_H5Fopen
if test "$ac_res" != no; then :
  test "$ac_res" = "none required" || LIBS="$ac_res $LIBS"

else
  libhdf=no
fi

	if test $libhdf != yes; then
	  cat > config.mk <<'EOF'
ifneq ($(MAKECMDGOALS),distclean)
$(error Resolve configure error first)
endif
EOF
   as_fn_error $? "HDF5 library development files not found!
	  On Debian based systems such as Ubuntu you may use sudo apt-get install libhdf5-dev.
	  On RPM based systems such as Fedora you may use sudo dnf install hdf5-devel.
	  " "$LINENO" 5
	fi

	libhdfheader=no
	for ac_header in hdf5.h hdf5/hdf5.h hdf5/serial/hdf5.h
do :
  as_ac_Header=`$as_echo "ac_cv_header_$ac_header" | $as_tr_sh`
ac_fn_c_check_header_mongrel "$LINENO" "$ac_header" "$as_ac_Header" "$ac_includes_default"
if eval test \"x\$"$as_ac_Header"\" = x"yes"; then :
  cat >>confdefs.h <<_ACEOF
#define `$as_echo "HAVE_$ac_header" | $as_tr_cpp` 1
_ACEOF
 libhdfheader=yes;break;
fi

done

	if test $libhdfheader != yes; then
	  cat > config.mk <<'EOF'
ifneq ($(MAKECMDGOALS),distclean)
$(error Resolve configure error first)
endif
EOF
   as_fn_error $? "HDF5 header files not found!
	  On Debian based systems such as Ubuntu you may use sudo apt-get install libhdf5-dev.
	  On RPM based systems such as Fedora you may use sudo dnf install hdf5-devel.
	  " "$LINENO" 5
	fi


fi


ac_config_files="$ac_config_files config.mk"

cat >confcache <<\_ACEOF
# This file is a shell script that caches the results of configure
# tests run on this system so they can be shared between configure
# scripts and configure runs, see configure's option --config-cache.
# It is not useful on other systems.  If it contains results you don't
# want to keep, you may remove or edit it.
#
# config.status only pays attention to the cache file if you give it
# the --recheck option to rerun configure.
#
# `ac_cv_env_foo' variables (set or unset) will be overridden when
# loading this file, other *unset* `ac_cv_foo' will be assigned the
# following values.

_ACEOF

# The following way of writing the cache mishandles newlines in values,
# but we know of no workaround that is simple, portable, and efficient.
# So, we kill variables containing newlines.
# Ultrix sh set writes to stderr and can't be redirected directly,
# and sets the high bit in the cache file unless we assign to the vars.
(
  for ac_var in `(set) 2>&1 | sed -n 's/^\([a-zA-Z_][a-zA-Z0-9_]*\)=.*/\1/p'`; do
    eval ac_val=\$$ac_var
    case $ac_val in #(
    *${as_nl}*)
      case $ac_var in #(
      *_cv_*) { $as_echo "$as_me:${as_lineno-$LINENO}: WARNING: cache variable $ac_var contains a newline" >&5
$as_echo "$as_me: WARNING: cache variable $ac_var contains a newline" >&2;} ;;
      esac
      case $ac_var in #(
      _ | IFS | as_nl) ;; #(
      BASH_ARGV | BASH_SOURCE) eval $ac_var= ;; #(
      *) { eval $ac_var=; unset $ac_var;} ;;
      esac ;;
    esac
  done

  (set) 2>&1 |
    case $as_nl`(ac_space=' '; set) 2>&1` in #(
    *${as_nl}ac_space=\ *)
      # `set' does not quote correctly, so add quotes: double-quote
      # substitution turns \\\\ into \\, and sed turns \\ into \.
      sed -n \
	"s/'/'\\\\''/g;
	  s/^\\([_$as_cr_alnum]*_cv_[_$as_cr_alnum]*\\)=\\(.*\\)/\\1='\\2'/p"
      ;; #(
    *)
      # `set' quotes correctly as required by POSIX, so do not add quotes.
      sed -n "/^[_$as_cr_alnum]*_cv_[_$as_cr_alnum]*=/p"
      ;;
    esac |
    sort
) |
  sed '
     /^ac_cv_env_/b end
     t clear
     :clear
     s/^\([^=]*\)=\(.*[{}].*\)$/test "${\1+set}" = set || &/
     t end
     s/^\([^=]*\)=\(.*\)$/\1=${\1=\2}/
     :end' >>confcache
if diff "$cache_file" confcache >/dev/null 2>&1; then :; else
  if test -w "$cache_file"; then
    if test "x$cache_file" != "x/dev/null"; then
      { $as_echo "$as_me:${as_lineno-$LINENO}: updating cache $cache_file" >&5
$as_echo "$as_me: updating cache $cache_file" >&6;}
      if test ! -f "$cache_file" || test -h "$cache_file"; then
	cat confcache >"$cache_file"
      else
        case $cache_file in #(
        */* | ?:*)
	  mv -f confcache "$cache_file"$$ &&
	  mv -f "$cache_file"$$ "$cache_file" ;; #(
        *)
	  mv -f confcache "$cache_file" ;;
	esac
      fi
    fi
  else
    { $as_echo "$as_me:${as_lineno-$LINENO}: not updating unwritable cache $cache_file" >&5
$as_echo "$as_me: not updating unwritable cache $cache_file" >&6;}
  fi
fi
rm -f confcache

test "x$prefix" = xNONE && prefix=$ac_default_prefix
# Let make expand exec_prefix.
test "x$exec_prefix" = xNONE && exec_prefix='${prefix}'

DEFS=-DHAVE_CONFIG_H

ac_libobjs=
ac_ltlibobjs=
U=
for ac_i in : $LIBOBJS; do test "x$ac_i" = x: && continue
  # 1. Remove the extension, and $U if already installed.
  ac_script='s/\$U\././;s/\.o$//;s/\.obj$//'
  ac_i=`$as_echo "$ac_i" | sed "$ac_script"`
  # 2. Prepend LIBOBJDIR.  When used with automake>=1.10 LIBOBJDIR
  #    will be set to the directory where LIBOBJS objects are built.
  as_fn_append ac_libobjs " \${LIBOBJDIR}$ac_i\$U.$ac_objext"
  as_fn_append ac_ltlibobjs " \${LIBOBJDIR}$ac_i"'$U.lo'
done
LIBOBJS=$ac_libobjs

LTLIBOBJS=$ac_ltlibobjs


: "${CONFIG_STATUS=./config.status}"
ac_write_fail=0
ac_clean_files_save=$ac_clean_files
ac_clean_files="$ac_clean_files $CONFIG_STATUS"
{ $as_echo "$as_me:${as_lineno-$LINENO}: creating $CONFIG_STATUS" >&5
$as_echo "$as_me: creating $CONFIG_STATUS" >&6;}
as_write_fail=0
cat >$CONFIG_STATUS <<_ASEOF || as_write_fail=1
#! $SHELL
# Generated by $as_me.
# Run this file to recreate the current configuration.
# Compiler output produced by configure, useful for debugging
# configure, is in config.log if it exists.

debug=false
ac_cs_recheck=false
ac_cs_silent=false

SHELL=\${CONFIG_SHELL-$SHELL}
export SHELL
_ASEOF
cat >>$CONFIG_STATUS <<\_ASEOF || as_write_fail=1
## -------------------- ##
## M4sh Initialization. ##
## -------------------- ##

# Be more Bourne compatible
DUALCASE=1; export DUALCASE # for MKS sh
if test -n "${ZSH_VERSION+set}" && (emulate sh) >/dev/null 2>&1; then :
  emulate sh
  NULLCMD=:
  # Pre-4.2 versions of Zsh do word splitting on ${1+"$@"}, which
  # is contrary to our usage.  Disable this feature.
  alias -g '${1+"$@"}'='"$@"'
  setopt NO_GLOB_SUBST
else
  case `(set -o) 2>/dev/null` in #(
  *posix*) :
    set -o posix ;; #(
  *) :
     ;;
esac
fi


as_nl='
'
export as_nl
# Printing a long string crashes Solaris 7 /usr/bin/printf.
as_echo='\\\\\\\\\\\\\\\\\\\\\\\\\\\\\\\\\\\\\\\\\\\\\\\\\\\\\\\\\\\\\\\\\\\\\\\\\\\\\\\\\\\\\\\\\\\\\\\\\\\\\\\'
as_echo=$as_echo$as_echo$as_echo$as_echo$as_echo
as_echo=$as_echo$as_echo$as_echo$as_echo$as_echo$as_echo
# Prefer a ksh shell builtin over an external printf program on Solaris,
# but without wasting forks for bash or zsh.
if test -z "$BASH_VERSION$ZSH_VERSION" \
    && (test "X`print -r -- $as_echo`" = "X$as_echo") 2>/dev/null; then
  as_echo='print -r --'
  as_echo_n='print -rn --'
elif (test "X`printf %s $as_echo`" = "X$as_echo") 2>/dev/null; then
  as_echo='printf %s\n'
  as_echo_n='printf %s'
else
  if test "X`(/usr/ucb/echo -n -n $as_echo) 2>/dev/null`" = "X-n $as_echo"; then
    as_echo_body='eval /usr/ucb/echo -n "$1$as_nl"'
    as_echo_n='/usr/ucb/echo -n'
  else
    as_echo_body='eval expr "X$1" : "X\\(.*\\)"'
    as_echo_n_body='eval
      arg=$1;
      case $arg in #(
      *"$as_nl"*)
	expr "X$arg" : "X\\(.*\\)$as_nl";
	arg=`expr "X$arg" : ".*$as_nl\\(.*\\)"`;;
      esac;
      expr "X$arg" : "X\\(.*\\)" | tr -d "$as_nl"
    '
    export as_echo_n_body
    as_echo_n='sh -c $as_echo_n_body as_echo'
  fi
  export as_echo_body
  as_echo='sh -c $as_echo_body as_echo'
fi

# The user is always right.
if test "${PATH_SEPARATOR+set}" != set; then
  PATH_SEPARATOR=:
  (PATH='/bin;/bin'; FPATH=$PATH; sh -c :) >/dev/null 2>&1 && {
    (PATH='/bin:/bin'; FPATH=$PATH; sh -c :) >/dev/null 2>&1 ||
      PATH_SEPARATOR=';'
  }
fi


# IFS
# We need space, tab and new line, in precisely that order.  Quoting is
# there to prevent editors from complaining about space-tab.
# (If _AS_PATH_WALK were called with IFS unset, it would disable word
# splitting by setting IFS to empty value.)
IFS=" ""	$as_nl"

# Find who we are.  Look in the path if we contain no directory separator.
as_myself=
case $0 in #((
  *[\\/]* ) as_myself=$0 ;;
  *) as_save_IFS=$IFS; IFS=$PATH_SEPARATOR
for as_dir in $PATH
do
  IFS=$as_save_IFS
  test -z "$as_dir" && as_dir=.
    test -r "$as_dir/$0" && as_myself=$as_dir/$0 && break
  done
IFS=$as_save_IFS

     ;;
esac
# We did not find ourselves, most probably we were run as `sh COMMAND'
# in which case we are not to be found in the path.
if test "x$as_myself" = x; then
  as_myself=$0
fi
if test ! -f "$as_myself"; then
  $as_echo "$as_myself: error: cannot find myself; rerun with an absolute file name" >&2
  exit 1
fi

# Unset variables that we do not need and which cause bugs (e.g. in
# pre-3.0 UWIN ksh).  But do not cause bugs in bash 2.01; the "|| exit 1"
# suppresses any "Segmentation fault" message there.  '((' could
# trigger a bug in pdksh 5.2.14.
for as_var in BASH_ENV ENV MAIL MAILPATH
do eval test x\${$as_var+set} = xset \
  && ( (unset $as_var) || exit 1) >/dev/null 2>&1 && unset $as_var || :
done
PS1='$ '
PS2='> '
PS4='+ '

# NLS nuisances.
LC_ALL=C
export LC_ALL
LANGUAGE=C
export LANGUAGE

# CDPATH.
(unset CDPATH) >/dev/null 2>&1 && unset CDPATH


# as_fn_error STATUS ERROR [LINENO LOG_FD]
# ----------------------------------------
# Output "`basename $0`: error: ERROR" to stderr. If LINENO and LOG_FD are
# provided, also output the error to LOG_FD, referencing LINENO. Then exit the
# script with STATUS, using 1 if that was 0.
as_fn_error ()
{
  as_status=$1; test $as_status -eq 0 && as_status=1
  if test "$4"; then
    as_lineno=${as_lineno-"$3"} as_lineno_stack=as_lineno_stack=$as_lineno_stack
    $as_echo "$as_me:${as_lineno-$LINENO}: error: $2" >&$4
  fi
  $as_echo "$as_me: error: $2" >&2
  as_fn_exit $as_status
} # as_fn_error


# as_fn_set_status STATUS
# -----------------------
# Set $? to STATUS, without forking.
as_fn_set_status ()
{
  return $1
} # as_fn_set_status

# as_fn_exit STATUS
# -----------------
# Exit the shell with STATUS, even in a "trap 0" or "set -e" context.
as_fn_exit ()
{
  set +e
  as_fn_set_status $1
  exit $1
} # as_fn_exit

# as_fn_unset VAR
# ---------------
# Portably unset VAR.
as_fn_unset ()
{
  { eval $1=; unset $1;}
}
as_unset=as_fn_unset
# as_fn_append VAR VALUE
# ----------------------
# Append the text in VALUE to the end of the definition contained in VAR. Take
# advantage of any shell optimizations that allow amortized linear growth over
# repeated appends, instead of the typical quadratic growth present in naive
# implementations.
if (eval "as_var=1; as_var+=2; test x\$as_var = x12") 2>/dev/null; then :
  eval 'as_fn_append ()
  {
    eval $1+=\$2
  }'
else
  as_fn_append ()
  {
    eval $1=\$$1\$2
  }
fi # as_fn_append

# as_fn_arith ARG...
# ------------------
# Perform arithmetic evaluation on the ARGs, and store the result in the
# global $as_val. Take advantage of shells that can avoid forks. The arguments
# must be portable across $(()) and expr.
if (eval "test \$(( 1 + 1 )) = 2") 2>/dev/null; then :
  eval 'as_fn_arith ()
  {
    as_val=$(( $* ))
  }'
else
  as_fn_arith ()
  {
    as_val=`expr "$@" || test $? -eq 1`
  }
fi # as_fn_arith


if expr a : '\(a\)' >/dev/null 2>&1 &&
   test "X`expr 00001 : '.*\(...\)'`" = X001; then
  as_expr=expr
else
  as_expr=false
fi

if (basename -- /) >/dev/null 2>&1 && test "X`basename -- / 2>&1`" = "X/"; then
  as_basename=basename
else
  as_basename=false
fi

if (as_dir=`dirname -- /` && test "X$as_dir" = X/) >/dev/null 2>&1; then
  as_dirname=dirname
else
  as_dirname=false
fi

as_me=`$as_basename -- "$0" ||
$as_expr X/"$0" : '.*/\([^/][^/]*\)/*$' \| \
	 X"$0" : 'X\(//\)$' \| \
	 X"$0" : 'X\(/\)' \| . 2>/dev/null ||
$as_echo X/"$0" |
    sed '/^.*\/\([^/][^/]*\)\/*$/{
	    s//\1/
	    q
	  }
	  /^X\/\(\/\/\)$/{
	    s//\1/
	    q
	  }
	  /^X\/\(\/\).*/{
	    s//\1/
	    q
	  }
	  s/.*/./; q'`

# Avoid depending upon Character Ranges.
as_cr_letters='abcdefghijklmnopqrstuvwxyz'
as_cr_LETTERS='ABCDEFGHIJKLMNOPQRSTUVWXYZ'
as_cr_Letters=$as_cr_letters$as_cr_LETTERS
as_cr_digits='0123456789'
as_cr_alnum=$as_cr_Letters$as_cr_digits

ECHO_C= ECHO_N= ECHO_T=
case `echo -n x` in #(((((
-n*)
  case `echo 'xy\c'` in
  *c*) ECHO_T='	';;	# ECHO_T is single tab character.
  xy)  ECHO_C='\c';;
  *)   echo `echo ksh88 bug on AIX 6.1` > /dev/null
       ECHO_T='	';;
  esac;;
*)
  ECHO_N='-n';;
esac

rm -f conf$$ conf$$.exe conf$$.file
if test -d conf$$.dir; then
  rm -f conf$$.dir/conf$$.file
else
  rm -f conf$$.dir
  mkdir conf$$.dir 2>/dev/null
fi
if (echo >conf$$.file) 2>/dev/null; then
  if ln -s conf$$.file conf$$ 2>/dev/null; then
    as_ln_s='ln -s'
    # ... but there are two gotchas:
    # 1) On MSYS, both `ln -s file dir' and `ln file dir' fail.
    # 2) DJGPP < 2.04 has no symlinks; `ln -s' creates a wrapper executable.
    # In both cases, we have to default to `cp -pR'.
    ln -s conf$$.file conf$$.dir 2>/dev/null && test ! -f conf$$.exe ||
      as_ln_s='cp -pR'
  elif ln conf$$.file conf$$ 2>/dev/null; then
    as_ln_s=ln
  else
    as_ln_s='cp -pR'
  fi
else
  as_ln_s='cp -pR'
fi
rm -f conf$$ conf$$.exe conf$$.dir/conf$$.file conf$$.file
rmdir conf$$.dir 2>/dev/null


# as_fn_mkdir_p
# -------------
# Create "$as_dir" as a directory, including parents if necessary.
as_fn_mkdir_p ()
{

  case $as_dir in #(
  -*) as_dir=./$as_dir;;
  esac
  test -d "$as_dir" || eval $as_mkdir_p || {
    as_dirs=
    while :; do
      case $as_dir in #(
      *\'*) as_qdir=`$as_echo "$as_dir" | sed "s/'/'\\\\\\\\''/g"`;; #'(
      *) as_qdir=$as_dir;;
      esac
      as_dirs="'$as_qdir' $as_dirs"
      as_dir=`$as_dirname -- "$as_dir" ||
$as_expr X"$as_dir" : 'X\(.*[^/]\)//*[^/][^/]*/*$' \| \
	 X"$as_dir" : 'X\(//\)[^/]' \| \
	 X"$as_dir" : 'X\(//\)$' \| \
	 X"$as_dir" : 'X\(/\)' \| . 2>/dev/null ||
$as_echo X"$as_dir" |
    sed '/^X\(.*[^/]\)\/\/*[^/][^/]*\/*$/{
	    s//\1/
	    q
	  }
	  /^X\(\/\/\)[^/].*/{
	    s//\1/
	    q
	  }
	  /^X\(\/\/\)$/{
	    s//\1/
	    q
	  }
	  /^X\(\/\).*/{
	    s//\1/
	    q
	  }
	  s/.*/./; q'`
      test -d "$as_dir" && break
    done
    test -z "$as_dirs" || eval "mkdir $as_dirs"
  } || test -d "$as_dir" || as_fn_error $? "cannot create directory $as_dir"


} # as_fn_mkdir_p
if mkdir -p . 2>/dev/null; then
  as_mkdir_p='mkdir -p "$as_dir"'
else
  test -d ./-p && rmdir ./-p
  as_mkdir_p=false
fi


# as_fn_executable_p FILE
# -----------------------
# Test if FILE is an executable regular file.
as_fn_executable_p ()
{
  test -f "$1" && test -x "$1"
} # as_fn_executable_p
as_test_x='test -x'
as_executable_p=as_fn_executable_p

# Sed expression to map a string onto a valid CPP name.
as_tr_cpp="eval sed 'y%*$as_cr_letters%P$as_cr_LETTERS%;s%[^_$as_cr_alnum]%_%g'"

# Sed expression to map a string onto a valid variable name.
as_tr_sh="eval sed 'y%*+%pp%;s%[^_$as_cr_alnum]%_%g'"


exec 6>&1
## ----------------------------------- ##
## Main body of $CONFIG_STATUS script. ##
## ----------------------------------- ##
_ASEOF
test $as_write_fail = 0 && chmod +x $CONFIG_STATUS || ac_write_fail=1

cat >>$CONFIG_STATUS <<\_ACEOF || ac_write_fail=1
# Save the log message, to keep $0 and so on meaningful, and to
# report actual input values of CONFIG_FILES etc. instead of their
# values after options handling.
ac_log="
This file was extended by f5c $as_me 1.0, which was
generated by GNU Autoconf 2.69.  Invocation command line was

  CONFIG_FILES    = $CONFIG_FILES
  CONFIG_HEADERS  = $CONFIG_HEADERS
  CONFIG_LINKS    = $CONFIG_LINKS
  CONFIG_COMMANDS = $CONFIG_COMMANDS
  $ $0 $@

on `(hostname || uname -n) 2>/dev/null | sed 1q`
"

_ACEOF

case $ac_config_files in *"
"*) set x $ac_config_files; shift; ac_config_files=$*;;
esac

case $ac_config_headers in *"
"*) set x $ac_config_headers; shift; ac_config_headers=$*;;
esac


cat >>$CONFIG_STATUS <<_ACEOF || ac_write_fail=1
# Files that config.status was made for.
config_files="$ac_config_files"
config_headers="$ac_config_headers"

_ACEOF

cat >>$CONFIG_STATUS <<\_ACEOF || ac_write_fail=1
ac_cs_usage="\
\`$as_me' instantiates files and other configuration actions
from templates according to the current configuration.  Unless the files
and actions are specified as TAGs, all are instantiated by default.

Usage: $0 [OPTION]... [TAG]...

  -h, --help       print this help, then exit
  -V, --version    print version number and configuration settings, then exit
      --config     print configuration, then exit
  -q, --quiet, --silent
                   do not print progress messages
  -d, --debug      don't remove temporary files
      --recheck    update $as_me by reconfiguring in the same conditions
      --file=FILE[:TEMPLATE]
                   instantiate the configuration file FILE
      --header=FILE[:TEMPLATE]
                   instantiate the configuration header FILE

Configuration files:
$config_files

Configuration headers:
$config_headers

Report bugs to <>."

_ACEOF
cat >>$CONFIG_STATUS <<_ACEOF || ac_write_fail=1
ac_cs_config="`$as_echo "$ac_configure_args" | sed 's/^ //; s/[\\""\`\$]/\\\\&/g'`"
ac_cs_version="\\
f5c config.status 1.0
configured by $0, generated by GNU Autoconf 2.69,
  with options \\"\$ac_cs_config\\"

Copyright (C) 2012 Free Software Foundation, Inc.
This config.status script is free software; the Free Software Foundation
gives unlimited permission to copy, distribute and modify it."

ac_pwd='$ac_pwd'
srcdir='$srcdir'
test -n "\$AWK" || AWK=awk
_ACEOF

cat >>$CONFIG_STATUS <<\_ACEOF || ac_write_fail=1
# The default lists apply if the user does not specify any file.
ac_need_defaults=:
while test $# != 0
do
  case $1 in
  --*=?*)
    ac_option=`expr "X$1" : 'X\([^=]*\)='`
    ac_optarg=`expr "X$1" : 'X[^=]*=\(.*\)'`
    ac_shift=:
    ;;
  --*=)
    ac_option=`expr "X$1" : 'X\([^=]*\)='`
    ac_optarg=
    ac_shift=:
    ;;
  *)
    ac_option=$1
    ac_optarg=$2
    ac_shift=shift
    ;;
  esac

  case $ac_option in
  # Handling of the options.
  -recheck | --recheck | --rechec | --reche | --rech | --rec | --re | --r)
    ac_cs_recheck=: ;;
  --version | --versio | --versi | --vers | --ver | --ve | --v | -V )
    $as_echo "$ac_cs_version"; exit ;;
  --config | --confi | --conf | --con | --co | --c )
    $as_echo "$ac_cs_config"; exit ;;
  --debug | --debu | --deb | --de | --d | -d )
    debug=: ;;
  --file | --fil | --fi | --f )
    $ac_shift
    case $ac_optarg in
    *\'*) ac_optarg=`$as_echo "$ac_optarg" | sed "s/'/'\\\\\\\\''/g"` ;;
    '') as_fn_error $? "missing file argument" ;;
    esac
    as_fn_append CONFIG_FILES " '$ac_optarg'"
    ac_need_defaults=false;;
  --header | --heade | --head | --hea )
    $ac_shift
    case $ac_optarg in
    *\'*) ac_optarg=`$as_echo "$ac_optarg" | sed "s/'/'\\\\\\\\''/g"` ;;
    esac
    as_fn_append CONFIG_HEADERS " '$ac_optarg'"
    ac_need_defaults=false;;
  --he | --h)
    # Conflict between --help and --header
    as_fn_error $? "ambiguous option: \`$1'
Try \`$0 --help' for more information.";;
  --help | --hel | -h )
    $as_echo "$ac_cs_usage"; exit ;;
  -q | -quiet | --quiet | --quie | --qui | --qu | --q \
  | -silent | --silent | --silen | --sile | --sil | --si | --s)
    ac_cs_silent=: ;;

  # This is an error.
  -*) as_fn_error $? "unrecognized option: \`$1'
Try \`$0 --help' for more information." ;;

  *) as_fn_append ac_config_targets " $1"
     ac_need_defaults=false ;;

  esac
  shift
done

ac_configure_extra_args=

if $ac_cs_silent; then
  exec 6>/dev/null
  ac_configure_extra_args="$ac_configure_extra_args --silent"
fi

_ACEOF
cat >>$CONFIG_STATUS <<_ACEOF || ac_write_fail=1
if \$ac_cs_recheck; then
  set X $SHELL '$0' $ac_configure_args \$ac_configure_extra_args --no-create --no-recursion
  shift
  \$as_echo "running CONFIG_SHELL=$SHELL \$*" >&6
  CONFIG_SHELL='$SHELL'
  export CONFIG_SHELL
  exec "\$@"
fi

_ACEOF
cat >>$CONFIG_STATUS <<\_ACEOF || ac_write_fail=1
exec 5>>config.log
{
  echo
  sed 'h;s/./-/g;s/^.../## /;s/...$/ ##/;p;x;p;x' <<_ASBOX
## Running $as_me. ##
_ASBOX
  $as_echo "$ac_log"
} >&5

_ACEOF
cat >>$CONFIG_STATUS <<_ACEOF || ac_write_fail=1
_ACEOF

cat >>$CONFIG_STATUS <<\_ACEOF || ac_write_fail=1

# Handling of arguments.
for ac_config_target in $ac_config_targets
do
  case $ac_config_target in
    "src/config.h") CONFIG_HEADERS="$CONFIG_HEADERS src/config.h" ;;
    "config.mk") CONFIG_FILES="$CONFIG_FILES config.mk" ;;

  *) as_fn_error $? "invalid argument: \`$ac_config_target'" "$LINENO" 5;;
  esac
done


# If the user did not use the arguments to specify the items to instantiate,
# then the envvar interface is used.  Set only those that are not.
# We use the long form for the default assignment because of an extremely
# bizarre bug on SunOS 4.1.3.
if $ac_need_defaults; then
  test "${CONFIG_FILES+set}" = set || CONFIG_FILES=$config_files
  test "${CONFIG_HEADERS+set}" = set || CONFIG_HEADERS=$config_headers
fi

# Have a temporary directory for convenience.  Make it in the build tree
# simply because there is no reason against having it here, and in addition,
# creating and moving files from /tmp can sometimes cause problems.
# Hook for its removal unless debugging.
# Note that there is a small window in which the directory will not be cleaned:
# after its creation but before its name has been assigned to `$tmp'.
$debug ||
{
  tmp= ac_tmp=
  trap 'exit_status=$?
  : "${ac_tmp:=$tmp}"
  { test ! -d "$ac_tmp" || rm -fr "$ac_tmp"; } && exit $exit_status
' 0
  trap 'as_fn_exit 1' 1 2 13 15
}
# Create a (secure) tmp directory for tmp files.

{
  tmp=`(umask 077 && mktemp -d "./confXXXXXX") 2>/dev/null` &&
  test -d "$tmp"
}  ||
{
  tmp=./conf$$-$RANDOM
  (umask 077 && mkdir "$tmp")
} || as_fn_error $? "cannot create a temporary directory in ." "$LINENO" 5
ac_tmp=$tmp

# Set up the scripts for CONFIG_FILES section.
# No need to generate them if there are no CONFIG_FILES.
# This happens for instance with `./config.status config.h'.
if test -n "$CONFIG_FILES"; then


ac_cr=`echo X | tr X '\015'`
# On cygwin, bash can eat \r inside `` if the user requested igncr.
# But we know of no other shell where ac_cr would be empty at this
# point, so we can use a bashism as a fallback.
if test "x$ac_cr" = x; then
  eval ac_cr=\$\'\\r\'
fi
ac_cs_awk_cr=`$AWK 'BEGIN { print "a\rb" }' </dev/null 2>/dev/null`
if test "$ac_cs_awk_cr" = "a${ac_cr}b"; then
  ac_cs_awk_cr='\\r'
else
  ac_cs_awk_cr=$ac_cr
fi

echo 'BEGIN {' >"$ac_tmp/subs1.awk" &&
_ACEOF


{
  echo "cat >conf$$subs.awk <<_ACEOF" &&
  echo "$ac_subst_vars" | sed 's/.*/&!$&$ac_delim/' &&
  echo "_ACEOF"
} >conf$$subs.sh ||
  as_fn_error $? "could not make $CONFIG_STATUS" "$LINENO" 5
ac_delim_num=`echo "$ac_subst_vars" | grep -c '^'`
ac_delim='%!_!# '
for ac_last_try in false false false false false :; do
  . ./conf$$subs.sh ||
    as_fn_error $? "could not make $CONFIG_STATUS" "$LINENO" 5

  ac_delim_n=`sed -n "s/.*$ac_delim\$/X/p" conf$$subs.awk | grep -c X`
  if test $ac_delim_n = $ac_delim_num; then
    break
  elif $ac_last_try; then
    as_fn_error $? "could not make $CONFIG_STATUS" "$LINENO" 5
  else
    ac_delim="$ac_delim!$ac_delim _$ac_delim!! "
  fi
done
rm -f conf$$subs.sh

cat >>$CONFIG_STATUS <<_ACEOF || ac_write_fail=1
cat >>"\$ac_tmp/subs1.awk" <<\\_ACAWK &&
_ACEOF
sed -n '
h
s/^/S["/; s/!.*/"]=/
p
g
s/^[^!]*!//
:repl
t repl
s/'"$ac_delim"'$//
t delim
:nl
h
s/\(.\{148\}\)..*/\1/
t more1
s/["\\]/\\&/g; s/^/"/; s/$/\\n"\\/
p
n
b repl
:more1
s/["\\]/\\&/g; s/^/"/; s/$/"\\/
p
g
s/.\{148\}//
t nl
:delim
h
s/\(.\{148\}\)..*/\1/
t more2
s/["\\]/\\&/g; s/^/"/; s/$/"/
p
b
:more2
s/["\\]/\\&/g; s/^/"/; s/$/"\\/
p
g
s/.\{148\}//
t delim
' <conf$$subs.awk | sed '
/^[^""]/{
  N
  s/\n//
}
' >>$CONFIG_STATUS || ac_write_fail=1
rm -f conf$$subs.awk
cat >>$CONFIG_STATUS <<_ACEOF || ac_write_fail=1
_ACAWK
cat >>"\$ac_tmp/subs1.awk" <<_ACAWK &&
  for (key in S) S_is_set[key] = 1
  FS = ""

}
{
  line = $ 0
  nfields = split(line, field, "@")
  substed = 0
  len = length(field[1])
  for (i = 2; i < nfields; i++) {
    key = field[i]
    keylen = length(key)
    if (S_is_set[key]) {
      value = S[key]
      line = substr(line, 1, len) "" value "" substr(line, len + keylen + 3)
      len += length(value) + length(field[++i])
      substed = 1
    } else
      len += 1 + keylen
  }

  print line
}

_ACAWK
_ACEOF
cat >>$CONFIG_STATUS <<\_ACEOF || ac_write_fail=1
if sed "s/$ac_cr//" < /dev/null > /dev/null 2>&1; then
  sed "s/$ac_cr\$//; s/$ac_cr/$ac_cs_awk_cr/g"
else
  cat
fi < "$ac_tmp/subs1.awk" > "$ac_tmp/subs.awk" \
  || as_fn_error $? "could not setup config files machinery" "$LINENO" 5
_ACEOF

# VPATH may cause trouble with some makes, so we remove sole $(srcdir),
# ${srcdir} and @srcdir@ entries from VPATH if srcdir is ".", strip leading and
# trailing colons and then remove the whole line if VPATH becomes empty
# (actually we leave an empty line to preserve line numbers).
if test "x$srcdir" = x.; then
  ac_vpsub='/^[	 ]*VPATH[	 ]*=[	 ]*/{
h
s///
s/^/:/
s/[	 ]*$/:/
s/:\$(srcdir):/:/g
s/:\${srcdir}:/:/g
s/:@srcdir@:/:/g
s/^:*//
s/:*$//
x
s/\(=[	 ]*\).*/\1/
G
s/\n//
s/^[^=]*=[	 ]*$//
}'
fi

cat >>$CONFIG_STATUS <<\_ACEOF || ac_write_fail=1
fi # test -n "$CONFIG_FILES"

# Set up the scripts for CONFIG_HEADERS section.
# No need to generate them if there are no CONFIG_HEADERS.
# This happens for instance with `./config.status Makefile'.
if test -n "$CONFIG_HEADERS"; then
cat >"$ac_tmp/defines.awk" <<\_ACAWK ||
BEGIN {
_ACEOF

# Transform confdefs.h into an awk script `defines.awk', embedded as
# here-document in config.status, that substitutes the proper values into
# config.h.in to produce config.h.

# Create a delimiter string that does not exist in confdefs.h, to ease
# handling of long lines.
ac_delim='%!_!# '
for ac_last_try in false false :; do
  ac_tt=`sed -n "/$ac_delim/p" confdefs.h`
  if test -z "$ac_tt"; then
    break
  elif $ac_last_try; then
    as_fn_error $? "could not make $CONFIG_HEADERS" "$LINENO" 5
  else
    ac_delim="$ac_delim!$ac_delim _$ac_delim!! "
  fi
done

# For the awk script, D is an array of macro values keyed by name,
# likewise P contains macro parameters if any.  Preserve backslash
# newline sequences.

ac_word_re=[_$as_cr_Letters][_$as_cr_alnum]*
sed -n '
s/.\{148\}/&'"$ac_delim"'/g
t rset
:rset
s/^[	 ]*#[	 ]*define[	 ][	 ]*/ /
t def
d
:def
s/\\$//
t bsnl
s/["\\]/\\&/g
s/^ \('"$ac_word_re"'\)\(([^()]*)\)[	 ]*\(.*\)/P["\1"]="\2"\
D["\1"]=" \3"/p
s/^ \('"$ac_word_re"'\)[	 ]*\(.*\)/D["\1"]=" \2"/p
d
:bsnl
s/["\\]/\\&/g
s/^ \('"$ac_word_re"'\)\(([^()]*)\)[	 ]*\(.*\)/P["\1"]="\2"\
D["\1"]=" \3\\\\\\n"\\/p
t cont
s/^ \('"$ac_word_re"'\)[	 ]*\(.*\)/D["\1"]=" \2\\\\\\n"\\/p
t cont
d
:cont
n
s/.\{148\}/&'"$ac_delim"'/g
t clear
:clear
s/\\$//
t bsnlc
s/["\\]/\\&/g; s/^/"/; s/$/"/p
d
:bsnlc
s/["\\]/\\&/g; s/^/"/; s/$/\\\\\\n"\\/p
b cont
' <confdefs.h | sed '
s/'"$ac_delim"'/"\\\
"/g' >>$CONFIG_STATUS || ac_write_fail=1

cat >>$CONFIG_STATUS <<_ACEOF || ac_write_fail=1
  for (key in D) D_is_set[key] = 1
  FS = ""
}
/^[\t ]*#[\t ]*(define|undef)[\t ]+$ac_word_re([\t (]|\$)/ {
  line = \$ 0
  split(line, arg, " ")
  if (arg[1] == "#") {
    defundef = arg[2]
    mac1 = arg[3]
  } else {
    defundef = substr(arg[1], 2)
    mac1 = arg[2]
  }
  split(mac1, mac2, "(") #)
  macro = mac2[1]
  prefix = substr(line, 1, index(line, defundef) - 1)
  if (D_is_set[macro]) {
    # Preserve the white space surrounding the "#".
    print prefix "define", macro P[macro] D[macro]
    next
  } else {
    # Replace #undef with comments.  This is necessary, for example,
    # in the case of _POSIX_SOURCE, which is predefined and required
    # on some systems where configure will not decide to define it.
    if (defundef == "undef") {
      print "/*", prefix defundef, macro, "*/"
      next
    }
  }
}
{ print }
_ACAWK
_ACEOF
cat >>$CONFIG_STATUS <<\_ACEOF || ac_write_fail=1
  as_fn_error $? "could not setup config headers machinery" "$LINENO" 5
fi # test -n "$CONFIG_HEADERS"


eval set X "  :F $CONFIG_FILES  :H $CONFIG_HEADERS    "
shift
for ac_tag
do
  case $ac_tag in
  :[FHLC]) ac_mode=$ac_tag; continue;;
  esac
  case $ac_mode$ac_tag in
  :[FHL]*:*);;
  :L* | :C*:*) as_fn_error $? "invalid tag \`$ac_tag'" "$LINENO" 5;;
  :[FH]-) ac_tag=-:-;;
  :[FH]*) ac_tag=$ac_tag:$ac_tag.in;;
  esac
  ac_save_IFS=$IFS
  IFS=:
  set x $ac_tag
  IFS=$ac_save_IFS
  shift
  ac_file=$1
  shift

  case $ac_mode in
  :L) ac_source=$1;;
  :[FH])
    ac_file_inputs=
    for ac_f
    do
      case $ac_f in
      -) ac_f="$ac_tmp/stdin";;
      *) # Look for the file first in the build tree, then in the source tree
	 # (if the path is not absolute).  The absolute path cannot be DOS-style,
	 # because $ac_f cannot contain `:'.
	 test -f "$ac_f" ||
	   case $ac_f in
	   [\\/$]*) false;;
	   *) test -f "$srcdir/$ac_f" && ac_f="$srcdir/$ac_f";;
	   esac ||
	   as_fn_error 1 "cannot find input file: \`$ac_f'" "$LINENO" 5;;
      esac
      case $ac_f in *\'*) ac_f=`$as_echo "$ac_f" | sed "s/'/'\\\\\\\\''/g"`;; esac
      as_fn_append ac_file_inputs " '$ac_f'"
    done

    # Let's still pretend it is `configure' which instantiates (i.e., don't
    # use $as_me), people would be surprised to read:
    #    /* config.h.  Generated by config.status.  */
    configure_input='Generated from '`
	  $as_echo "$*" | sed 's|^[^:]*/||;s|:[^:]*/|, |g'
	`' by configure.'
    if test x"$ac_file" != x-; then
      configure_input="$ac_file.  $configure_input"
      { $as_echo "$as_me:${as_lineno-$LINENO}: creating $ac_file" >&5
$as_echo "$as_me: creating $ac_file" >&6;}
    fi
    # Neutralize special characters interpreted by sed in replacement strings.
    case $configure_input in #(
    *\&* | *\|* | *\\* )
       ac_sed_conf_input=`$as_echo "$configure_input" |
       sed 's/[\\\\&|]/\\\\&/g'`;; #(
    *) ac_sed_conf_input=$configure_input;;
    esac

    case $ac_tag in
    *:-:* | *:-) cat >"$ac_tmp/stdin" \
      || as_fn_error $? "could not create $ac_file" "$LINENO" 5 ;;
    esac
    ;;
  esac

  ac_dir=`$as_dirname -- "$ac_file" ||
$as_expr X"$ac_file" : 'X\(.*[^/]\)//*[^/][^/]*/*$' \| \
	 X"$ac_file" : 'X\(//\)[^/]' \| \
	 X"$ac_file" : 'X\(//\)$' \| \
	 X"$ac_file" : 'X\(/\)' \| . 2>/dev/null ||
$as_echo X"$ac_file" |
    sed '/^X\(.*[^/]\)\/\/*[^/][^/]*\/*$/{
	    s//\1/
	    q
	  }
	  /^X\(\/\/\)[^/].*/{
	    s//\1/
	    q
	  }
	  /^X\(\/\/\)$/{
	    s//\1/
	    q
	  }
	  /^X\(\/\).*/{
	    s//\1/
	    q
	  }
	  s/.*/./; q'`
  as_dir="$ac_dir"; as_fn_mkdir_p
  ac_builddir=.

case "$ac_dir" in
.) ac_dir_suffix= ac_top_builddir_sub=. ac_top_build_prefix= ;;
*)
  ac_dir_suffix=/`$as_echo "$ac_dir" | sed 's|^\.[\\/]||'`
  # A ".." for each directory in $ac_dir_suffix.
  ac_top_builddir_sub=`$as_echo "$ac_dir_suffix" | sed 's|/[^\\/]*|/..|g;s|/||'`
  case $ac_top_builddir_sub in
  "") ac_top_builddir_sub=. ac_top_build_prefix= ;;
  *)  ac_top_build_prefix=$ac_top_builddir_sub/ ;;
  esac ;;
esac
ac_abs_top_builddir=$ac_pwd
ac_abs_builddir=$ac_pwd$ac_dir_suffix
# for backward compatibility:
ac_top_builddir=$ac_top_build_prefix

case $srcdir in
  .)  # We are building in place.
    ac_srcdir=.
    ac_top_srcdir=$ac_top_builddir_sub
    ac_abs_top_srcdir=$ac_pwd ;;
  [\\/]* | ?:[\\/]* )  # Absolute name.
    ac_srcdir=$srcdir$ac_dir_suffix;
    ac_top_srcdir=$srcdir
    ac_abs_top_srcdir=$srcdir ;;
  *) # Relative name.
    ac_srcdir=$ac_top_build_prefix$srcdir$ac_dir_suffix
    ac_top_srcdir=$ac_top_build_prefix$srcdir
    ac_abs_top_srcdir=$ac_pwd/$srcdir ;;
esac
ac_abs_srcdir=$ac_abs_top_srcdir$ac_dir_suffix


  case $ac_mode in
  :F)
  #
  # CONFIG_FILE
  #

_ACEOF

cat >>$CONFIG_STATUS <<\_ACEOF || ac_write_fail=1
# If the template does not know about datarootdir, expand it.
# FIXME: This hack should be removed a few years after 2.60.
ac_datarootdir_hack=; ac_datarootdir_seen=
ac_sed_dataroot='
/datarootdir/ {
  p
  q
}
/@datadir@/p
/@docdir@/p
/@infodir@/p
/@localedir@/p
/@mandir@/p'
case `eval "sed -n \"\$ac_sed_dataroot\" $ac_file_inputs"` in
*datarootdir*) ac_datarootdir_seen=yes;;
*@datadir@*|*@docdir@*|*@infodir@*|*@localedir@*|*@mandir@*)
  { $as_echo "$as_me:${as_lineno-$LINENO}: WARNING: $ac_file_inputs seems to ignore the --datarootdir setting" >&5
$as_echo "$as_me: WARNING: $ac_file_inputs seems to ignore the --datarootdir setting" >&2;}
_ACEOF
cat >>$CONFIG_STATUS <<_ACEOF || ac_write_fail=1
  ac_datarootdir_hack='
  s&@datadir@&$datadir&g
  s&@docdir@&$docdir&g
  s&@infodir@&$infodir&g
  s&@localedir@&$localedir&g
  s&@mandir@&$mandir&g
  s&\\\${datarootdir}&$datarootdir&g' ;;
esac
_ACEOF

# Neutralize VPATH when `$srcdir' = `.'.
# Shell code in configure.ac might set extrasub.
# FIXME: do we really want to maintain this feature?
cat >>$CONFIG_STATUS <<_ACEOF || ac_write_fail=1
ac_sed_extra="$ac_vpsub
$extrasub
_ACEOF
cat >>$CONFIG_STATUS <<\_ACEOF || ac_write_fail=1
:t
/@[a-zA-Z_][a-zA-Z_0-9]*@/!b
s|@configure_input@|$ac_sed_conf_input|;t t
s&@top_builddir@&$ac_top_builddir_sub&;t t
s&@top_build_prefix@&$ac_top_build_prefix&;t t
s&@srcdir@&$ac_srcdir&;t t
s&@abs_srcdir@&$ac_abs_srcdir&;t t
s&@top_srcdir@&$ac_top_srcdir&;t t
s&@abs_top_srcdir@&$ac_abs_top_srcdir&;t t
s&@builddir@&$ac_builddir&;t t
s&@abs_builddir@&$ac_abs_builddir&;t t
s&@abs_top_builddir@&$ac_abs_top_builddir&;t t
$ac_datarootdir_hack
"
eval sed \"\$ac_sed_extra\" "$ac_file_inputs" | $AWK -f "$ac_tmp/subs.awk" \
  >$ac_tmp/out || as_fn_error $? "could not create $ac_file" "$LINENO" 5

test -z "$ac_datarootdir_hack$ac_datarootdir_seen" &&
  { ac_out=`sed -n '/\${datarootdir}/p' "$ac_tmp/out"`; test -n "$ac_out"; } &&
  { ac_out=`sed -n '/^[	 ]*datarootdir[	 ]*:*=/p' \
      "$ac_tmp/out"`; test -z "$ac_out"; } &&
  { $as_echo "$as_me:${as_lineno-$LINENO}: WARNING: $ac_file contains a reference to the variable \`datarootdir'
which seems to be undefined.  Please make sure it is defined" >&5
$as_echo "$as_me: WARNING: $ac_file contains a reference to the variable \`datarootdir'
which seems to be undefined.  Please make sure it is defined" >&2;}

  rm -f "$ac_tmp/stdin"
  case $ac_file in
  -) cat "$ac_tmp/out" && rm -f "$ac_tmp/out";;
  *) rm -f "$ac_file" && mv "$ac_tmp/out" "$ac_file";;
  esac \
  || as_fn_error $? "could not create $ac_file" "$LINENO" 5
 ;;
  :H)
  #
  # CONFIG_HEADER
  #
  if test x"$ac_file" != x-; then
    {
      $as_echo "/* $configure_input  */" \
      && eval '$AWK -f "$ac_tmp/defines.awk"' "$ac_file_inputs"
    } >"$ac_tmp/config.h" \
      || as_fn_error $? "could not create $ac_file" "$LINENO" 5
    if diff "$ac_file" "$ac_tmp/config.h" >/dev/null 2>&1; then
      { $as_echo "$as_me:${as_lineno-$LINENO}: $ac_file is unchanged" >&5
$as_echo "$as_me: $ac_file is unchanged" >&6;}
    else
      rm -f "$ac_file"
      mv "$ac_tmp/config.h" "$ac_file" \
	|| as_fn_error $? "could not create $ac_file" "$LINENO" 5
    fi
  else
    $as_echo "/* $configure_input  */" \
      && eval '$AWK -f "$ac_tmp/defines.awk"' "$ac_file_inputs" \
      || as_fn_error $? "could not create -" "$LINENO" 5
  fi
 ;;


  esac

done # for ac_tag


as_fn_exit 0
_ACEOF
ac_clean_files=$ac_clean_files_save

test $ac_write_fail = 0 ||
  as_fn_error $? "write failure creating $CONFIG_STATUS" "$LINENO" 5


# configure is writing to config.log, and then calls config.status.
# config.status does its own redirection, appending to config.log.
# Unfortunately, on DOS this fails, as config.log is still kept open
# by configure, so config.status won't be able to write to it; its
# output is simply discarded.  So we exec the FD to /dev/null,
# effectively closing config.log, so it can be properly (re)opened and
# appended to by config.status.  When coming back to configure, we
# need to make the FD available again.
if test "$no_create" != yes; then
  ac_cs_success=:
  ac_config_status_args=
  test "$silent" = yes &&
    ac_config_status_args="$ac_config_status_args --quiet"
  exec 5>/dev/null
  $SHELL $CONFIG_STATUS $ac_config_status_args || ac_cs_success=false
  exec 5>>config.log
  # Use ||, not &&, to avoid exiting from the if with $? = 1, which
  # would make configure fail if this is the last instruction.
  $ac_cs_success || as_fn_exit 1
fi
if test -n "$ac_unrecognized_opts" && test "$enable_option_checking" != no; then
  { $as_echo "$as_me:${as_lineno-$LINENO}: WARNING: unrecognized options: $ac_unrecognized_opts" >&5
$as_echo "$as_me: WARNING: unrecognized options: $ac_unrecognized_opts" >&2;}
fi
