## Supplementary material for "GPU Accelerated Adaptive Banded Event Alignment for Rapid Comparative Nanopore Signal Analysis": f5c-v0.2-beta-release: Dockerfile

```
FROM ubuntu:16.04
WORKDIR /
RUN apt-get update && apt-get install libhdf5-dev zlib1g-dev git wget tar gcc g++ make autoconf bash bzip2 -y
RUN git clone https://github.com/hasindu2008/f5c.git
WORKDIR /f5c
RUN autoreconf && ./scripts/install-hts.sh && ./configure && make && make test
CMD ./f5c
```
