## Supplementary material for "GPU Accelerated Adaptive Banded Event Alignment for Rapid Comparative Nanopore Signal Analysis": f5c-v0.2-beta-release: Makefile

```
-include config.mk
-include installdeps.mk

CC       = gcc
CXX      = g++
CFLAGS   += -g -rdynamic -Wall -O2 -std=c++11 
LDFLAGS  += $(LIBS) -lpthread -lz
BUILD_DIR = build

BINARY = f5c
OBJ = $(BUILD_DIR)/main.o \
      $(BUILD_DIR)/meth_main.o \
      $(BUILD_DIR)/f5c.o \
      $(BUILD_DIR)/events.o \
      $(BUILD_DIR)/nanopolish_read_db.o \
      $(BUILD_DIR)/nanopolish_index.o \
      $(BUILD_DIR)/model.o \
      $(BUILD_DIR)/align.o \
      $(BUILD_DIR)/meth.o \
      $(BUILD_DIR)/hmm.o \
      $(BUILD_DIR)/freq.o \
      $(BUILD_DIR)/eventalign.o

PREFIX = /usr/local
VERSION = `git describe --tags`

ifdef cuda
    CUDA_ROOT = /usr/local/cuda
    CUDA_LIB ?= $(CUDA_ROOT)/lib64
    CUDA_OBJ = $(BUILD_DIR)/f5c_cuda.o $(BUILD_DIR)/align_cuda.o
    NVCC ?= nvcc
    CUDA_CFLAGS += -g  -O2 -std=c++11 -lineinfo $(CUDA_ARCH) -Xcompiler -Wall
    CUDA_LDFLAGS = -L$(CUDA_LIB) -lcudart_static -lrt -ldl
    OBJ += $(BUILD_DIR)/gpucode.o $(CUDA_OBJ)
    CPPFLAGS += -DHAVE_CUDA=1
endif

.PHONY: clean distclean format test install uninstall

$(BINARY): src/config.h $(HTS_LIB) $(HDF5_LIB) $(OBJ)
	$(CXX) $(CFLAGS) $(OBJ) $(LDFLAGS) $(CUDA_LDFLAGS) -o $@

$(BUILD_DIR)/main.o: src/main.c src/f5cmisc.h src/error.h
	$(CXX) $(CFLAGS) $(CPPFLAGS) $< -c -o $@

$(BUILD_DIR)/meth_main.o: src/meth_main.c src/f5c.h src/fast5lite.h src/f5cmisc.h src/logsum.h
	$(CXX) $(CFLAGS) $(CPPFLAGS) $< -c -o $@

$(BUILD_DIR)/f5c.o: src/f5c.c src/f5c.h src/fast5lite.h src/f5cmisc.h
	$(CXX) $(CFLAGS) $(CPPFLAGS) $< -c -o $@

$(BUILD_DIR)/events.o: src/events.c src/f5c.h src/fast5lite.h src/f5cmisc.h src/fast5lite.h src/nanopolish_read_db.h src/ksort.h
	$(CXX) $(CFLAGS) $(CPPFLAGS) $< -c -o $@

$(BUILD_DIR)/nanopolish_read_db.o: src/nanopolish_read_db.c src/nanopolish_read_db.h
	$(CXX) $(CFLAGS) $(CPPFLAGS) $< -c -o $@

$(BUILD_DIR)/nanopolish_index.o: src/nanopolish_index.c src/nanopolish_read_db.h src/fast5lite.h
	$(CXX) $(CFLAGS) $(CPPFLAGS) $< -c -o $@

$(BUILD_DIR)/model.o: src/model.c src/model.h src/f5c.h src/fast5lite.h src/f5cmisc.h
	$(CXX) $(CFLAGS) $(CPPFLAGS) $< -c -o $@

$(BUILD_DIR)/align.o: src/align.c src/f5c.h src/fast5lite.h
	$(CXX) $(CFLAGS) $(CPPFLAGS) $< -c -o $@

$(BUILD_DIR)/meth.o: src/meth.c src/f5c.h src/fast5lite.h src/f5cmisc.h
	$(CXX) $(CFLAGS) $(CPPFLAGS) $< -c -o $@

$(BUILD_DIR)/hmm.o: src/hmm.c src/f5c.h src/fast5lite.h src/f5cmisc.h src/matrix.h src/logsum.h
	$(CXX) $(CFLAGS) $(CPPFLAGS) $< -c -o $@

$(BUILD_DIR)/freq.o: src/freq.c src/khash.h
	$(CXX) $(CFLAGS) $(CPPFLAGS) $< -c -o $@	

$(BUILD_DIR)/eventalign.o: src/eventalign.c
	$(CXX) $(CFLAGS) $(CPPFLAGS) $< -c -o $@
	
# cuda stuff
$(BUILD_DIR)/gpucode.o: $(CUDA_OBJ)
	$(NVCC) $(CUDA_CFLAGS) -dlink $^ -o $@ 

$(BUILD_DIR)/f5c_cuda.o: src/f5c.cu src/error.h src/f5c.h src/fast5lite.h src/f5cmisc.cuh src/f5cmisc.h
	$(NVCC) -x cu $(CUDA_CFLAGS) $(CPPFLAGS) -rdc=true -c $< -o $@

$(BUILD_DIR)/align_cuda.o: src/align.cu src/f5c.h src/fast5lite.h src/f5cmisc.cuh
	$(NVCC) -x cu $(CUDA_CFLAGS) $(CPPFLAGS) -rdc=true -c $< -o $@

src/config.h:
	echo "/* Default config.h generated by Makefile */" >> $@
	echo "#define HAVE_HDF5_H 1" >> $@

$(BUILD_DIR)/lib/libhts.a:
	@if command -v curl; then \
		curl -o $(BUILD_DIR)/htslib.tar.bz2 -L https://github.com/samtools/htslib/releases/download/$(HTS_VERSION)/htslib-$(HTS_VERSION).tar.bz2; \
	else \
		wget -O $(BUILD_DIR)/htslib.tar.bz2 https://github.com/samtools/htslib/releases/download/$(HTS_VERSION)/htslib-$(HTS_VERSION).tar.bz2; \
	fi
	tar -xf $(BUILD_DIR)/htslib.tar.bz2 -C $(BUILD_DIR)
	mv $(BUILD_DIR)/htslib-$(HTS_VERSION) $(BUILD_DIR)/htslib
	rm -f $(BUILD_DIR)/htslib.tar.bz2
	cd $(BUILD_DIR)/htslib && \
	./configure --prefix=`pwd`/../ --enable-bz2=no --enable-lzma=no --with-libdeflate=no --enable-libcurl=no  --enable-gcs=no --enable-s3=no && \
	make -j8 && \
	make install

$(BUILD_DIR)/lib/libhdf5.a:
	if command -v curl; then \
		curl -o $(BUILD_DIR)/hdf5.tar.bz2 https://support.hdfgroup.org/ftp/HDF5/releases/hdf5-$(HDF5_MAJOR_MINOR)/hdf5-$(HDF5_VERSION)/src/hdf5-$(HDF5_VERSION).tar.bz2; \
	else \
		wget -O $(BUILD_DIR)/hdf5.tar.bz2 https://support.hdfgroup.org/ftp/HDF5/releases/hdf5-$(HDF5_MAJOR_MINOR)/hdf5-$(HDF5_VERSION)/src/hdf5-$(HDF5_VERSION).tar.bz2; \
	fi
	tar -xf $(BUILD_DIR)/hdf5.tar.bz2 -C $(BUILD_DIR)
	mv $(BUILD_DIR)/hdf5-$(HDF5_VERSION) $(BUILD_DIR)/hdf5
	rm -f $(BUILD_DIR)/hdf5.tar.bz2
	cd $(BUILD_DIR)/hdf5 && \
	./configure --prefix=`pwd`/../ && \
	make -j8 && \
	make install

clean: 
	rm -rf $(BINARY) $(BUILD_DIR)/*.o calculate_methylation_frequency

# Delete all gitignored files (but not directories)
distclean: clean
	git clean -f -X 
	rm -rf $(BUILD_DIR)/* autom4te.cache

dist: distclean
	mkdir -p f5c-$(VERSION)
	autoreconf
	cp -r README.md LICENSE Dockerfile Makefile configure.ac config.mk.in \
		installdeps.mk src docs build .dockerignore configure f5c-$(VERSION)
	mkdir -p f5c-$(VERSION)/scripts
	cp scripts/install-hdf5.sh scripts/install-hts.sh scripts/test.sh scripts/common.sh scripts/test.awk f5c-$(VERSION)/scripts
	tar -zcf f5c-$(VERSION)-release.tar.gz f5c-$(VERSION)
	rm -rf f5c-$(VERSION)

binary: 
	mkdir -p f5c-$(VERSION)
	make clean
	make cuda=1 && mv f5c f5c-$(VERSION)/f5c_x86_64_linux_cuda && make clean
	make && mv f5c f5c-$(VERSION)/f5c_x86_64_linux
	cp -r README.md LICENSE docs f5c-$(VERSION)/
	mkdir -p f5c-$(VERSION)/scripts
	cp scripts/test.sh scripts/common.sh scripts/test.awk f5c-$(VERSION)/scripts
	tar -zcf f5c-$(VERSION)-binaries.tar.gz f5c-$(VERSION)
	rm -rf f5c-$(VERSION)

install: $(BINARY)
	mkdir -p $(DESTDIR)$(PREFIX)/bin
	mkdir -p $(DESTDIR)$(PREFIX)/share/man/man1
	cp -f $(BINARY) $(DESTDIR)$(PREFIX)/bin
	gzip < docs/f5c.1 > $(DESTDIR)$(PREFIX)/share/man/man1/f5c.1.gz

uninstall:
	rm -f $(DESTDIR)$(PREFIX)/bin/$(BINARY) \
		$(DESTDIR)$(PREFIX)/share/man/man1/f5c.1.gz

test: $(BINARY)
	./scripts/test.sh

test_eventalign: $(BINARY)
	./scripts/test_eventalign.sh
```
