## Supplementary material for "GPU Accelerated Adaptive Banded Event Alignment for Rapid Comparative Nanopore Signal Analysis": f5c-v0.2-beta-release: README.md

# f5c

An optimised re-implementation of the call-methylation module in [Nanopolish](https://github.com/jts/nanopolish). Given a set of basecalled Nanopore reads and the raw signals, f5c detects the methylated cytosine bases. f5c can optionally utilise NVIDIA graphics cards for acceleration.

First the reads have to be indexed using `f5c index`. Then invoke `f5c call-methylation` to detect methylated cytosine bases. Finally, you may use `f5c meth-freq` to obtain methylation frequencies. The results are almost the same as from nanopolish except a few differences due to floating point approximations.

[![Build Status](https://travis-ci.org/hasindu2008/f5c.svg?branch=master)](https://travis-ci.org/hasindu2008/f5c)

## Quick start

If you are a Linux user and want to quickly try out download the compiled binaries from the [latest release](https://github.com/hasindu2008/f5c/releases). For example:
```sh
VERSION=v0.2-beta
wget "https://github.com/hasindu2008/f5c/releases/download/$VERSION/f5c-$VERSION-binaries.tar.gz" && tar xvf f5c-$VERSION-binaries.tar.gz && cd f5c-$VERSION/
./f5c_x86_64_linux        # CPU version
./f5c_x86_64_linux_cuda   # cuda supported version
```
Binaries should work on most Linux distributions and the only dependency is `zlib` which is available by default on most distros.

## Building

Users are recommended to build from the  [latest release](https://github.com/hasindu2008/f5c/releases) tar ball. You need a compiler that supports C++11. Quick example for Ubuntu :
```sh
sudo apt-get install libhdf5-dev zlib1g-dev   #install HDF5 and zlib development libraries
VERSION=v0.2-beta
wget "https://github.com/hasindu2008/f5c/releases/download/$VERSION/f5c-$VERSION-release.tar.gz" && tar xvf f5c-$VERSION-release.tar.gz && cd f5c-$VERSION/
scripts/install-hts.sh  # download and compile the htslib
./configure             
make                    # make cuda=1 to enable CUDA support
```
The commands to install hdf5 (and zlib) __development libraries__ on some popular distributions :
```sh
On Debian/Ubuntu : sudo apt-get install libhdf5-dev zlib1g-dev
On Fedora/CentOS : sudo dnf/yum install hdf5-devel zlib-devel
On Arch Linux: sudo pacman -S hdf5
On OS X : brew install hdf5
```
If you skip `scripts/install-hts.sh` and `./configure` hdf5 will be compiled locally. It is a good option if you cannot install hdf5 library system wide. However, building hdf5 takes ages.

Building from the Github repository additionally requires `autoreconf` which can be installed on Ubuntu using `sudo apt-get install autoconf automake`.

Other building options are detailed [here](https://hasindu2008.github.io/f5c/docs/building).
Instruction to build a docker image is detailed [here](https://hasindu2008.github.io/f5c/docs/docker).

### NVIDIA CUDA support

To build for the GPU, you need to have the CUDA toolkit installed. Make nvcc (NVIDIA C Compiler) is in your PATH.  

The building instructions are the same as above except that you should call make as :
```
make cuda=1
```
Optionally you can provide the CUDA architecture as :
```
make cuda=1 CUDA_ARCH=-arch=sm_xy
```
If your CUDA library is not in the default location /usr/local/cuda/lib64, point to the correct location as:
```
make cuda=1 CUDA_LIB=/path/to/cuda/library/
```
Visit [here](https://hasindu2008.github.io/f5c/docs/cuda-troubleshoot) for troubleshooting CUDA related problems.

## Usage

```sh
f5c index -d [fast5_folder] [read.fastq|fasta]
f5c call-methylation -b [reads.sorted.bam] -g [ref.fa] -r [reads.fastq|fasta] > [meth.tsv]
f5c meth-freq -i [meth.tsv] > [freq.tsv]
```

Visit the [man page](https://hasindu2008.github.io/f5c/docs/commands) for all the commands and options.

### Example

Follow the same steps as in [Nanopolish tutorial](https://nanopolish.readthedocs.io/en/latest/quickstart_call_methylation.html) while replacing `nanopolish` with `f5c`. If you only want to perform a quick test of f5c :
```sh
#download and extract the dataset including sorted alignments
wget -O f5c_na12878_test.tgz "http://genome.cse.unsw.edu.au/tmp/f5c_na12878_test.tgz"
tar xf f5c_na12878_test.tgz

#index, call methylation and get methylation frequencies
f5c index -d chr22_meth_example/fast5_files chr22_meth_example/reads.fastq
f5c call-methylation -b chr22_meth_example/reads.sorted.bam -g chr22_meth_example/humangenome.fa -r chr22_meth_example/reads.fastq > chr22_meth_example/result.tsv
f5c meth-freq -i chr22_meth_example/result.tsv > chr22_meth_example/freq.tsv
```

## Acknowledgement
This reuses code and methods from [Nanopolish](https://github.com/jts/nanopolish).
The event detection code is from Oxford Nanopore's [Scrappie basecaller](https://github.com/nanoporetech/scrappie).
Some code snippets have been taken from [Minimap2](https://github.com/lh3/minimap2) and [Samtools](http://samtools.sourceforge.net/).
