## Supplementary material for "GPU Accelerated Adaptive Banded Event Alignment for Rapid Comparative Nanopore Signal Analysis": f5c-v0.2-beta-release: common.sh

```
#!/bin/sh

set -e

clean_cache=false

# terminate script
die() {
	echo "$1" >&2
	echo
	exit 1
}

clear_fscache() {
	sync
	echo 3 | tee /proc/sys/vm/drop_caches
}

# echo command before running it
run() {
	echo "$1"
	eval "$1"
	if [ $clean_cache = true ]
	then
		clear_fscache
	fi
	return 0
}
```
