## Supplementary material for "GPU Accelerated Adaptive Banded Event Alignment for Rapid Comparative Nanopore Signal Analysis": f5c-v0.2-beta-release: install-hdf5.sh

```
#!/bin/bash

#TODO : better error checking, exit with proper error message and state save
#Todo : use curl as a failsafe 

test -e hdf5-1.10.4.tar.gz && rm hdf5-1.10.4.tar.gz 
test -d hdf5 && rm -r hdf5
wget https://support.hdfgroup.org/ftp/HDF5/releases/hdf5-1.10/hdf5-1.10.4/src/hdf5-1.10.4.tar.gz || exit 1
tar -xzf hdf5-1.10.4.tar.gz || exit
rm hdf5-1.10.4.tar.gz 
mv hdf5-1.10.4 hdf5 || exit 1
cd hdf5 || exit 1
./configure --prefix=`pwd`
make -j8 || exit 1
make install || exit 1
echo "Successfully installed HDF5 to ./hdf5."
```
