## Supplementary material for "GPU Accelerated Adaptive Banded Event Alignment for Rapid Comparative Nanopore Signal Analysis": f5c-v0.2-beta-release: install-hts.sh

```
#!/bin/bash

# terminate script
die() {
	echo "install-hts.sh: $1" >&2
	exit 1
}

LINK=https://github.com/samtools/htslib/releases/download/1.9/htslib-1.9.tar.bz2
TARBALL=htslib-1.9.tar.bz2
FOLDER=htslib

test -e  $TARBALL && rm $TARBALL
test -d $FOLDER && rm -r $FOLDER
wget -O $TARBALL $LINK || curl -o $TARBALL $LINK || die "Downloading htslib from $LINK failed."
tar -xf $TARBALL  || die "Extracting $TARBALL failed"
rm $TARBALL
mv htslib-1.9 $FOLDER || die "moving htslib-1.9 to $FOLDER failed"
cd $FOLDER || die "changing directory to $FOLDER failed"
chmod +x ./configure || die "changing permission failed"
./configure --enable-bz2=no --enable-lzma=no --with-libdeflate=no --enable-libcurl=no  --enable-gcs=no --enable-s3=no || die "Fix the issue and run scripts/install-hts.sh again"
make -j8 || die "Fix the issue and run scripts/install-hts.sh again"
echo "Successfully installed htslib to ./htslib."
echo 'Now run ./configure (again)!'
```
