## Supplementary material for "GPU Accelerated Adaptive Banded Event Alignment for Rapid Comparative Nanopore Signal Analysis": f5c-v0.2-beta-release: test.awk

```
#!/bin/awk -f
function abs(x) {
	return (((x) < 0.0) ? -(x) : (x))
}

BEGIN {status=0}
{if (abs($2-$5)>abs(thresh*$5)+0.02 \
	|| abs($3-$6)>abs(thresh*$6)+0.02 \
	|| abs($4-$7)>abs(thresh*$7)+0.02)
	{print $0,abs($2-$5)">"abs(thresh*$5)+0.02, \
		abs($3-$6)">"abs(thresh*$6)+0.02, \
		abs($4-$7)">"abs(thresh*$7)+0.02; status=1}}
END {if (status > 0) {exit 1}}
```
