## Supplementary material for "GPU Accelerated Adaptive Banded Event Alignment for Rapid Comparative Nanopore Signal Analysis": f5c-v0.2-beta-release: test.sh

```
#!/bin/sh

# exit when command fails
set -e

. scripts/common.sh

# defaults
exepath=./f5c
testdir=test/ecoli_2kb_region

bamfile=${testdir}/reads.sorted.bam
ref=${testdir}/draft.fa
reads=${testdir}/reads.fasta
batchsize=256
max_bases=2M
if command -v nproc > /dev/null; then
	threads=$(nproc --all)
else
	threads=8
fi
# execution mode (valgrind/gdb/cpu/cuda/echo)
mode=
testset_url="http://genome.cse.unsw.edu.au/tmp/f5c_ecoli_2kb_region_test.tgz"
fallback_url="https://ndownloader.figshare.com/files/13784075?private_link=b04e3976eaed2225b848"

# download test set given url
#
# $1 - URL
# $2 - Fallback URL (optional)
download_test_set() {
	# data set exists
	if [ -d ${testdir} ]; then
		return
	fi

	mkdir -p test
	tar_path=test/data.tgz
	wget -O $tar_path "$1" || wget -O $tar_path "$2" || rm -rf $tar_path ${testdir}
	echo "Extracting. Please wait."
	tar -xf $tar_path -C test || rm -rf $tar_path ${testdir}
	rm -f $tar_path
}


handle_tests() {
	numfailed=$(wc -l < ${testdir}/floatdiff.txt)
	numcases=$(wc -l < ${testdir}/meth_float.txt)
	numres=$(wc -l < ${testdir}/result_float.txt)
	echo "$numfailed of $numcases test cases deviated."
	missing=$(echo "$numcases-$numres" | bc)
	echo "$missing entries in the truthset are missing in the testset"
	failp=$(echo "$numfailed/$numcases" | bc)
	[ "$failp" -gt 0 ] && die "${1}: Validation failed"
}

execute_test() {

	if [ $testdir = test/chr22_meth_example ]; then
		grep -w "chr20" ${testdir}/result.txt | awk '{print $1$2$3$4$8$9$10"\t"$5"\t"$6"\t"$7}' > ${testdir}/result_float.txt
		grep -w "chr20" ${testdir}/meth.exp | awk '{print $1$2$3$4$8$9$10"\t"$5"\t"$6"\t"$7}'  > ${testdir}/meth_float.txt

		join  ${testdir}/result_float.txt ${testdir}/meth_float.txt | awk -v thresh=0.1 -f scripts/test.awk > ${testdir}/floatdiff.txt || handle_tests "${file}"
	else	
		tail -n +2 ${testdir}/result.txt | awk '{print $1,$2,$3,$4,$8,$9,$10}' > ${testdir}/result_exact.txt
		awk '{print $1,$2,$3,$4,$8,$9,$10}' ${testdir}/meth.exp > ${testdir}/meth_exact.txt
		diff -q ${testdir}/meth_exact.txt ${testdir}/result_exact.txt || die "diff ${testdir}/result_exact.txt ${testdir}/meth_exact.txt failed" 

		tail -n +2  ${testdir}/result.txt | awk '{print $1$2$3$4$8$9$10"\t"$5"\t"$6"\t"$7}' > ${testdir}/result_float.txt
		awk '{print $1$2$3$4$8$9$10"\t"$5"\t"$6"\t"$7}' ${testdir}/meth.exp > ${testdir}/meth_float.txt	

		join -a 1 -a 2 ${testdir}/result_float.txt ${testdir}/meth_float.txt | awk -v thresh=0.1 -f scripts/test.awk > ${testdir}/floatdiff.txt || die "${file}: Validation failed" 
	fi	
}

mode_test() {
	cmd="${exepath} call-methylation -b ${bamfile} -g ${ref} -r ${reads} -t ${threads} -K $batchsize -B $max_bases"

	case $1 in
		valgrind) valgrind $cmd > /dev/null;;
		gdb) gdb --args "$cmd";;
		cpu) $cmd --disable-cuda=yes > ${testdir}/result.txt; execute_test;;
		cuda) $cmd --disable-cuda=no > ${testdir}/result.txt; execute_test;;
		echo) echo "$cmd -t $threads > ${testdir}/result.txt";;
		nvprof) nvprof  -f --analysis-metrics -o profile.nvprof "$cmd" --disable-cuda=no --debug-break=5 > /dev/null;;
		custom) shift; $cmd "$@" --secondary=yes --min-mapq=0 > ${testdir}/result.txt; execute_test;;
		*) die "Unknown mode: $1";;
	esac
}

help_msg() {
	echo "Test script for f5c."
	echo "Usage: f5c_dir/script/test.sh [-c] [-b bam file] [-g reference genome] [-r fastq/fasta read] mode"
	echo
	echo "mode                 one of: valgrind/gdb/cpu/cuda/echo"
	echo
	echo "-c                   Uses chr22_meth_example test set."
	echo "-b [bam file]        Same as f5c -b."
	echo "-K [n]               Same as f5c -K."
	echo "-B [n]               Same as f5c -B."
	echo "-r [read file]       Same as f5c -r."
	echo "-g [ref genome]      Same as f5c -g."
	echo "-t [n]               Number of threads."
	echo "-d                   Download chr22_meth_example data set and exit."
	echo "-h                   Show this help message."
}

# parse options
while getopts b:g:r:t:K:B:cdh opt
do
	case $opt in
		b) bamfile="$OPTARG";;
		g) ref="$OPTARG";;
		r) reads="$OPTARG";;
		t) threads="$OPTARG";;
		c) testdir=test/chr22_meth_example
		   bamfile=${testdir}/reads.sorted.bam
		   ref=${testdir}/humangenome.fa
		   reads=${testdir}/reads.fastq
		   testset_url="http://genome.cse.unsw.edu.au/tmp/f5c_na12878_test.tgz"
		   fallback_url="https://ndownloader.figshare.com/files/13784792?private_link=5dd2077f1041412a9518";;
		K) batchsize="$OPTARG";;
		B) max_bases="$OPTARG";;
		d) download_test_set "http://genome.cse.unsw.edu.au/tmp/f5c_na12878_test.tgz" "https://ndownloader.figshare.com/files/13784792?private_link=5dd2077f1041412a9518"
		   exit 0;;
		h) help_msg
		   exit 0;;
		?) printf "Usage: %s [-c] [-b bam file] [-g reference genome] [-r fastq/fasta read] args" "$0"
		   exit 2;;
	esac
done
shift $(($OPTIND - 1))
mode=$1

download_test_set $testset_url $fallback_url

# validate files
for file in ${bamfile} ${ref} ${reads}; do
	[ -f ${file} ] || die "${file}: File does not exist"
done

if [ -z "$mode" ]; then
	if [ $testdir = test/chr22_meth_example ]; then
		${exepath} call-methylation -b ${bamfile} -g ${ref} -r ${reads} -t "$threads" -K "$batchsize" -B "$max_bases" > ${testdir}/result.txt
		execute_test
	else
		${exepath} index -d ${testdir}/fast5_files ${testdir}/reads.fasta
		${exepath} call-methylation -b ${bamfile} -g ${ref} -r ${reads} --secondary=yes --min-mapq=0 -B "$max_bases" > ${testdir}/result.txt
		execute_test
	fi
else
	mode_test "$@"
fi
```
