## Supplementary material for "GPU Accelerated Adaptive Banded Event Alignment for Rapid Comparative Nanopore Signal Analysis": f5c-v0.2-beta-release: align.c

```
#include "f5c.h"
#include <assert.h>
//#define DEBUG_ESTIMATED_SCALING 1
//#define DEBUG_RECALIB_SCALING 1
//#define DEBUG_ADAPTIVE 1

//Code was adapted from Nanopolish : nanopolish_raw_loader.cpp

//todo : can make more efficient using bit encoding
static inline uint32_t get_rank(char base) {
    if (base == 'A') { //todo: do we neeed simple alpha?
        return 0;
    } else if (base == 'C') {
        return 1;
    } else if (base == 'G') {
        return 2;
    } else if (base == 'T') {
        return 3;
    } else {
        WARNING("A None ACGT base found : %c", base);
        return 0;
    }
}

// return the lexicographic rank of the kmer amongst all strings of
// length k for this alphabet
static inline uint32_t get_kmer_rank(const char* str, uint32_t k) {
    //uint32_t p = 1;
    uint32_t r = 0;

    // from last base to first
    for (uint32_t i = 0; i < k; ++i) {
        //r += rank(str[k - i - 1]) * p;
        //p *= size();
        r += get_rank(str[k - i - 1]) << (i << 1);
    }
    return r;
}

//copy a kmer from a reference
static inline void kmer_cpy(char* dest, char* src, uint32_t k) {
    uint32_t i = 0;
    for (i = 0; i < k; i++) {
        dest[i] = src[i];
    }
    dest[i] = '\0';
}

scalings_t estimate_scalings_using_mom(char* sequence, int32_t sequence_len,
                                       model_t* pore_model, event_table et) {
    scalings_t out;
    int32_t n_kmers =
        sequence_len - KMER_SIZE + 1; //todo :strlen can be pre-calculated

    //const Alphabet* alphabet = pore_model.pmalphabet;

    // Calculate summary statistics over the events and
    // the model implied by the read
    double event_level_sum = 0.0f; //do we need double?
    for (size_t i = 0; i < et.n; ++i) {
        event_level_sum += et.event[i].mean;
    }

    double kmer_level_sum = 0.0f;
    double kmer_level_sq_sum = 0.0f;
    for (int32_t i = 0; i < n_kmers; ++i) {
        int32_t kr = get_kmer_rank(&sequence[i], KMER_SIZE);
        double l = pore_model[kr].level_mean;
        //fprintf(stderr,"Kmer : %c%c%c%c%c%c, kmer_rank : %d , kmer_mean : %f \n",sequence[i],sequence[i+1],sequence[i+2],sequence[i+3],sequence[i+4],sequence[i+5],kr,l);
        kmer_level_sum += l;
        kmer_level_sq_sum += l * l;
    }

    double shift = event_level_sum / et.n - kmer_level_sum / n_kmers;

    // estimate scale
    double event_level_sq_sum = 0.0f;
    for (size_t i = 0; i < et.n; ++i) {
        event_level_sq_sum +=
            (et.event[i].mean - shift) * (et.event[i].mean - shift);
    }

    double scale = (event_level_sq_sum / et.n) / (kmer_level_sq_sum / n_kmers);

    //out.set4(shift, scale, 0.0, 1.0);
    out.shift = (float)shift;
    out.scale = (float)scale;

#ifdef DEBUG_ESTIMATED_SCALING
    fprintf(stderr, "event mean: %.2lf kmer mean: %.2lf shift: %.2lf\n",
            event_level_sum / et.n, kmer_level_sum / n_kmers, out.shift);
    fprintf(stderr, "event sq-mean: %.2lf kmer sq-mean: %.2lf scale: %.2lf\n",
            event_level_sq_sum / et.n, kmer_level_sq_sum / n_kmers, out.scale);
    //fprintf(stderr, "truth shift: %.2lf scale: %.2lf\n", pore_model.shift, pore_model.scale);
#endif
    return out;
}

static inline float log_normal_pdf(float x, float gp_mean, float gp_stdv,
                                   float gp_log_stdv) {
    /*INCOMPLETE*/
    float log_inv_sqrt_2pi = -0.918938f; // Natural logarithm
    float a = (x - gp_mean) / gp_stdv;
    return log_inv_sqrt_2pi - gp_log_stdv + (-0.5f * a * a);
    // return 1;
}

static inline float log_probability_match_r9(scalings_t scaling,
                                             model_t* models,
                                             event_table events, int event_idx,
                                             uint32_t kmer_rank, uint8_t strand,
                                             float sample_rate) {
    // event level mean, scaled with the drift value
    strand = 0;
    assert(kmer_rank < 4096);
    //float level = read.get_drift_scaled_level(event_idx, strand);

    //float time =
    //    (events.event[event_idx].start - events.event[0].start) / sample_rate;
    float unscaledLevel = events.event[event_idx].mean;
    float scaledLevel = unscaledLevel;
    //float scaledLevel = unscaledLevel - time * scaling.shift;

    //fprintf(stderr, "level %f\n",scaledLevel);
    //GaussianParameters gp = read.get_scaled_gaussian_from_pore_model_state(pore_model, strand, kmer_rank);
    float gp_mean =
        scaling.scale * models[kmer_rank].level_mean + scaling.shift;
    float gp_stdv = models[kmer_rank].level_stdv * 1; //scaling.var = 1;
    // float gp_stdv = 0;
    // float gp_log_stdv = models[kmer_rank].level_log_stdv + scaling.log_var;
    // if(models[kmer_rank].level_stdv <0.01 ){
    // 	fprintf(stderr,"very small std dev %f\n",models[kmer_rank].level_stdv);
    // }
    #ifdef CACHED_LOG
        float gp_log_stdv = models[kmer_rank].level_log_stdv;
    #else
        float gp_log_stdv =
        log(models[kmer_rank].level_stdv); // scaling.log_var = log(1)=0;
    #endif

    float lp = log_normal_pdf(scaledLevel, gp_mean, gp_stdv, gp_log_stdv);
    return lp;
}

#define event_kmer_to_band(ei, ki) (ei + 1) + (ki + 1)
#define band_event_to_offset(bi, ei) band_lower_left[bi].event_idx - (ei)
#define band_kmer_to_offset(bi, ki) (ki) - band_lower_left[bi].kmer_idx
#define is_offset_valid(offset) (offset) >= 0 && (offset) < bandwidth
#define event_at_offset(bi, offset) band_lower_left[(bi)].event_idx - (offset)
#define kmer_at_offset(bi, offset) band_lower_left[(bi)].kmer_idx + (offset)

#define move_down(curr_band)                                                   \
    { curr_band.event_idx + 1, curr_band.kmer_idx }
#define move_right(curr_band)                                                  \
    { curr_band.event_idx, curr_band.kmer_idx + 1 }

#define MIN(a, b) (((a) < (b)) ? (a) : (b))
#define MAX(a, b) (((a) > (b)) ? (a) : (b))


#ifdef ALIGN_2D_ARRAY
    #define BAND_ARRAY(r, c) ( bands[(r)][(c)] )
    #define TRACE_ARRAY(r, c) ( trace[(r)][(c)] )
#else
    #define BAND_ARRAY(r, c) ( bands[((r)*(ALN_BANDWIDTH)+(c))] )
    #define TRACE_ARRAY(r, c) ( trace[((r)*(ALN_BANDWIDTH)+(c))] )
#endif

int32_t align(AlignedPair* out_2, char* sequence, int32_t sequence_len,
              event_table events, model_t* models, scalings_t scaling,
              float sample_rate) {
    //fprintf(stderr, "%s\n", sequence);
    //fprintf(stderr, "Scaling %f %f", scaling.scale, scaling.shift);

    size_t strand_idx = 0;
    size_t k = 6;

    // size_t n_events = events[strand_idx].n;
    size_t n_events = events.n;
    size_t n_kmers = sequence_len - k + 1;
    //fprintf(stderr,"n_kmers : %d\n",n_kmers);
    // backtrack markers
    const uint8_t FROM_D = 0;
    const uint8_t FROM_U = 1;
    const uint8_t FROM_L = 2;

    // qc
    double min_average_log_emission = -5.0;
    int max_gap_threshold = 50;

    // banding
    int bandwidth = ALN_BANDWIDTH;
    int half_bandwidth = ALN_BANDWIDTH / 2;

    // transition penalties
    double events_per_kmer = (double)n_events / n_kmers;
    double p_stay = 1 - (1 / (events_per_kmer + 1));

    // setting a tiny skip penalty helps keep the true alignment within the adaptive band
    // this was empirically determined
    double epsilon = 1e-10;
    double lp_skip = log(epsilon);
    double lp_stay = log(p_stay);
    double lp_step = log(1.0 - exp(lp_skip) - exp(lp_stay));
    double lp_trim = log(0.01);

    // dp matrix
    size_t n_rows = n_events + 1;
    size_t n_cols = n_kmers + 1;
    size_t n_bands = n_rows + n_cols;

    // Initialize

    // Precompute k-mer ranks to avoid doing this in the inner loop
    size_t* kmer_ranks = (size_t*)malloc(sizeof(size_t) * n_kmers);
    MALLOC_CHK(kmer_ranks);

    for (size_t i = 0; i < n_kmers; ++i) {
        //>>>>>>>>> New replacement begin
        char* substring = &sequence[i];
        kmer_ranks[i] = get_kmer_rank(substring, k);
        //<<<<<<<<< New replacement over
    }

#ifdef ALIGN_2D_ARRAY
    float** bands = (float**)malloc(sizeof(float*) * n_bands);
    MALLOC_CHK(bands);
    uint8_t** trace = (uint8_t**)malloc(sizeof(uint8_t*) * n_bands);
    MALLOC_CHK(trace);
#else
    float* bands = (float*)malloc(sizeof(float) * n_bands * bandwidth);
    MALLOC_CHK(bands);
    uint8_t* trace = (uint8_t*)malloc(sizeof(uint8_t) * n_bands * bandwidth);
    MALLOC_CHK(trace);
#endif
    for (size_t i = 0; i < n_bands; i++) {
    #ifdef ALIGN_2D_ARRAY
        bands[i] = (float*)malloc(sizeof(float) * bandwidth);
        MALLOC_CHK(bands[i]);
        trace[i] = (uint8_t*)malloc(sizeof(uint8_t) * bandwidth);
        MALLOC_CHK(trace[i]);
    #endif

        for (int j = 0; j < bandwidth; j++) {
            BAND_ARRAY(i,j) = -INFINITY;
            TRACE_ARRAY(i,j) = 0;
        }
    }

    // Keep track of the event/kmer index for the lower left corner of the band
    // these indices are updated at every iteration to perform the adaptive banding
    // Only the first two  have their coordinates initialized, the rest are computed adaptively

    struct EventKmerPair {
        int event_idx;
        int kmer_idx;
    };
    //>>>>>>>>>>>>>>>>>New Replacement Begin
    struct EventKmerPair* band_lower_left =
        (struct EventKmerPair*)malloc(sizeof(struct EventKmerPair) * n_bands);
    MALLOC_CHK(band_lower_left);
    //std::vector<EventKmerPair> band_lower_left(n_);
    //<<<<<<<<<<<<<<<<<New Replacement over

    // initialize range of first two
    band_lower_left[0].event_idx = half_bandwidth - 1;
    band_lower_left[0].kmer_idx = -1 - half_bandwidth;
    band_lower_left[1] = move_down(band_lower_left[0]);

    int start_cell_offset = band_kmer_to_offset(0, -1);
    assert(is_offset_valid(start_cell_offset));
    assert(band_event_to_offset(0, -1) == start_cell_offset);
    BAND_ARRAY(0,start_cell_offset) = 0.0f;

    // band 1: first event is trimmed
    int first_trim_offset = band_event_to_offset(1, 0);
    assert(kmer_at_offset(1, first_trim_offset) == -1);
    assert(is_offset_valid(first_trim_offset));
    BAND_ARRAY(1,first_trim_offset) = lp_trim;
    TRACE_ARRAY(1,first_trim_offset) = FROM_U;

    int fills = 0;
#ifdef DEBUG_ADAPTIVE
    fprintf(stderr, "[trim] bi: %d o: %d e: %d k: %d s: %.2lf\n", 1,
            first_trim_offset, 0, -1, bands[1][first_trim_offset]);
#endif

    // fill in remaining bands
    for (size_t band_idx = 2; band_idx < n_bands; ++band_idx) {
        // Determine placement of this band according to Suzuki's adaptive algorithm
        // When both ll and ur are out-of-band (ob) we alternate movements
        // otherwise we decide based on scores
        float ll = BAND_ARRAY(band_idx - 1,0);
        float ur = BAND_ARRAY(band_idx - 1,bandwidth - 1);
        bool ll_ob = ll == -INFINITY;
        bool ur_ob = ur == -INFINITY;

        bool right = false;
        if (ll_ob && ur_ob) {
            right = band_idx % 2 == 1;
        } else {
            right = ll < ur; // Suzuki's rule
        }

        if (right) {
            band_lower_left[band_idx] =
                move_right(band_lower_left[band_idx - 1]);
        } else {
            band_lower_left[band_idx] =
                move_down(band_lower_left[band_idx - 1]);
        }
        // If the trim state is within the band, fill it in here
        int trim_offset = band_kmer_to_offset(band_idx, -1);
        if (is_offset_valid(trim_offset)) {
            size_t event_idx = event_at_offset(band_idx, trim_offset);
            if (event_idx >= 0 && event_idx < n_events) {
                BAND_ARRAY(band_idx,trim_offset) = lp_trim * (event_idx + 1);
                TRACE_ARRAY(band_idx,trim_offset) = FROM_U;
            } else {
                BAND_ARRAY(band_idx,trim_offset) = -INFINITY;
            }
        }

        // Get the offsets for the first and last event and kmer
        // We restrict the inner loop to only these values
        int kmer_min_offset = band_kmer_to_offset(band_idx, 0);
        int kmer_max_offset = band_kmer_to_offset(band_idx, n_kmers);
        int event_min_offset = band_event_to_offset(band_idx, n_events - 1);
        int event_max_offset = band_event_to_offset(band_idx, -1);

        int min_offset = MAX(kmer_min_offset, event_min_offset);
        min_offset = MAX(min_offset, 0);

        int max_offset = MIN(kmer_max_offset, event_max_offset);
        max_offset = MIN(max_offset, bandwidth);

        for (int offset = min_offset; offset < max_offset; ++offset) {
            int event_idx = event_at_offset(band_idx, offset);
            int kmer_idx = kmer_at_offset(band_idx, offset);

            size_t kmer_rank = kmer_ranks[kmer_idx];

            int offset_up = band_event_to_offset(band_idx - 1, event_idx - 1);
            int offset_left = band_kmer_to_offset(band_idx - 1, kmer_idx - 1);
            int offset_diag = band_kmer_to_offset(band_idx - 2, kmer_idx - 1);

#ifdef DEBUG_ADAPTIVE
            // verify loop conditions
            assert(kmer_idx >= 0 && kmer_idx < n_kmers);
            assert(event_idx >= 0 && event_idx < n_events);
            assert(offset_diag ==
                   band_event_to_offset(band_idx - 2, event_idx - 1));
            assert(offset_up - offset_left == 1);
            assert(offset >= 0 && offset < bandwidth);
#endif

            float up = is_offset_valid(offset_up)
                           ? BAND_ARRAY(band_idx - 1,offset_up)
                           : -INFINITY;
            float left = is_offset_valid(offset_left)
                             ? BAND_ARRAY(band_idx - 1,offset_left)
                             : -INFINITY;
            float diag = is_offset_valid(offset_diag)
                             ? BAND_ARRAY(band_idx - 2,offset_diag)
                             : -INFINITY;

            float lp_emission =
                log_probability_match_r9(scaling, models, events, event_idx,
                                         kmer_rank, strand_idx, sample_rate);
            //fprintf(stderr, "lp emiision : %f , event idx %d, kmer rank %d\n", lp_emission,event_idx,kmer_rank);
            float score_d = diag + lp_step + lp_emission;
            float score_u = up + lp_stay + lp_emission;
            float score_l = left + lp_skip;

            float max_score = score_d;
            uint8_t from = FROM_D;

            max_score = score_u > max_score ? score_u : max_score;
            from = max_score == score_u ? FROM_U : from;
            max_score = score_l > max_score ? score_l : max_score;
            from = max_score == score_l ? FROM_L : from;

#ifdef DEBUG_ADAPTIVE
            fprintf(stderr,
                    "[adafill] offset-up: %d offset-diag: %d offset-left: %d\n",
                    offset_up, offset_diag, offset_left);
            fprintf(stderr, "[adafill] up: %.2lf diag: %.2lf left: %.2lf\n", up,
                    diag, left);
            fprintf(stderr,
                    "[adafill] bi: %d o: %d e: %d k: %d s: %.2lf f: %d emit: "
                    "%.2lf\n",
                    band_idx, offset, event_idx, kmer_idx, max_score, from,
                    lp_emission);
#endif
            BAND_ARRAY(band_idx,offset) = max_score;
            TRACE_ARRAY(band_idx,offset) = from;
            fills += 1;
        }
    }

    //
    // Backtrack to compute alignment
    //
    double sum_emission = 0;
    double n_aligned_events = 0;

    //>>>>>>>>>>>>>> New replacement begin
    // std::vector<AlignedPair> out;

    int outIndex = 0;
    //<<<<<<<<<<<<<<<<New Replacement over

    float max_score = -INFINITY;
    int curr_event_idx = 0;
    int curr_kmer_idx = n_kmers - 1;

    // Find best score between an event and the last k-mer. after trimming the remaining evnets
    for (size_t event_idx = 0; event_idx < n_events; ++event_idx) {
        int band_idx = event_kmer_to_band(event_idx, curr_kmer_idx);

        //>>>>>>>New  replacement begin
        /*assert(band_idx < bands.size());*/
        assert((size_t)band_idx < n_bands);
        //<<<<<<<<New Replacement over
        int offset = band_event_to_offset(band_idx, event_idx);
        if (is_offset_valid(offset)) {
            float s =
                BAND_ARRAY(band_idx,offset) + (n_events - event_idx) * lp_trim;
            if (s > max_score) {
                max_score = s;
                curr_event_idx = event_idx;
            }
        }
    }

#ifdef DEBUG_ADAPTIVE
    fprintf(stderr, "[adaback] ei: %d ki: %d s: %.2f\n", curr_event_idx,
            curr_kmer_idx, max_score);
#endif

    int curr_gap = 0;
    int max_gap = 0;
    while (curr_kmer_idx >= 0 && curr_event_idx >= 0) {
        // emit alignment
        //>>>>>>>New Repalcement begin
        assert(outIndex < (int)(n_events * 2));
        out_2[outIndex].ref_pos = curr_kmer_idx;
        out_2[outIndex].read_pos = curr_event_idx;
        outIndex++;
        // out.push_back({curr_kmer_idx, curr_event_idx});
        //<<<<<<<<<New Replacement over

#ifdef DEBUG_ADAPTIVE
        fprintf(stderr, "[adaback] ei: %d ki: %d\n", curr_event_idx,
                curr_kmer_idx);
#endif
        // qc stats
        //>>>>>>>>>>>>>>New Replacement begin
        char* substring = &sequence[curr_kmer_idx];
        size_t kmer_rank = get_kmer_rank(substring, k);
        //<<<<<<<<<<<<<New Replacement over
        float tempLogProb = log_probability_match_r9(
            scaling, models, events, curr_event_idx, kmer_rank, 0, sample_rate);

        sum_emission += tempLogProb;
        //fprintf(stderr, "lp_emission %f \n", tempLogProb);
        //fprintf(stderr,"lp_emission %f, sum_emission %f, n_aligned_events %d\n",tempLogProb,sum_emission,outIndex);

        n_aligned_events += 1;

        int band_idx = event_kmer_to_band(curr_event_idx, curr_kmer_idx);
        int offset = band_event_to_offset(band_idx, curr_event_idx);
        assert(band_kmer_to_offset(band_idx, curr_kmer_idx) == offset);

        uint8_t from = TRACE_ARRAY(band_idx,offset);
        if (from == FROM_D) {
            curr_kmer_idx -= 1;
            curr_event_idx -= 1;
            curr_gap = 0;
        } else if (from == FROM_U) {
            curr_event_idx -= 1;
            curr_gap = 0;
        } else {
            curr_kmer_idx -= 1;
            curr_gap += 1;
            max_gap = MAX(curr_gap, max_gap);
        }
    }

    //>>>>>>>>New replacement begin
    // std::reverse(out.begin(), out.end());
    int c;
    int end = outIndex - 1;
    for (c = 0; c < outIndex / 2; c++) {
        int ref_pos_temp = out_2[c].ref_pos;
        int read_pos_temp = out_2[c].read_pos;
        out_2[c].ref_pos = out_2[end].ref_pos;
        out_2[c].read_pos = out_2[end].read_pos;
        out_2[end].ref_pos = ref_pos_temp;
        out_2[end].read_pos = read_pos_temp;
        end--;
    }

    // if(outIndex>1){
    //   AlignedPair temp={out_2[0].ref_pos,out[0].read_pos};
    //   int i;
    //   for(i=0;i<outIndex-1;i++){
    //     out_2[i]={out_2[outIndex-1-i].ref_pos,out[outIndex-1-i].read_pos};
    //   }
    //   out[outIndex-1]={temp.ref_pos,temp.read_pos};
    // }
    //<<<<<<<<<New replacement over

    // QC results
    double avg_log_emission = sum_emission / n_aligned_events;
    //fprintf(stderr,"sum_emission %f, n_aligned_events %f, avg_log_emission %f\n",sum_emission,n_aligned_events,avg_log_emission);
    //>>>>>>>>>>>>>New replacement begin
    bool spanned = out_2[0].ref_pos == 0 &&
                   out_2[outIndex - 1].ref_pos == int(n_kmers - 1);
    // bool spanned = out.front().ref_pos == 0 && out.back().ref_pos == n_kmers - 1;
    //<<<<<<<<<<<<<New replacement over
    //bool failed = false;
    if (avg_log_emission < min_average_log_emission || !spanned ||
        max_gap > max_gap_threshold) {
        //failed = true;
        //>>>>>>>>>>>>>New replacement begin
        outIndex = 0;
        // out.clear();
        //free(out_2);
        //out_2 = NULL;
        //<<<<<<<<<<<<<New replacement over
    }

    free(kmer_ranks);
#ifdef ALIGN_2D_ARRAY
    for (size_t i = 0; i < n_bands; i++) {
        free(bands[i]);
        free(trace[i]);
    }
#endif
    free(bands);
    free(trace);
    free(band_lower_left);
    //fprintf(stderr, "ada\t%s\t%s\t%.2lf\t%zu\t%.2lf\t%d\t%d\t%d\n", read.read_name.substr(0, 6).c_str(), failed ? "FAILED" : "OK", events_per_kmer, sequence.size(), avg_log_emission, curr_event_idx, max_gap, fills);
    //outSize=outIndex;
    //if(outIndex>500000)fprintf(stderr, "Max outSize %d\n", outIndex);
    return outIndex;
}

int32_t postalign(event_alignment_t* alignment, index_pair_t* base_to_event_map,double* events_per_base,
                  char* sequence, int32_t n_kmers, AlignedPair* event_alignment,
                  int32_t n_events) {
    /* transform alignment into the base-to-event map*/
    // create base-to-event map
    // index_pair_t* base_to_event_map =
    //     (index_pair_t*)(malloc(sizeof(index_pair_t) * n_kmers));
    // MALLOC_CHK(base_to_event_map);

    //initialisesing (todo : check if really required)
    int32_t i = 0;
    for (i = 0; i < n_kmers; i++) {
        base_to_event_map[i].start = -1;
        base_to_event_map[i].stop = -1;
    }

    int32_t max_event = 0;
    int32_t min_event = INT32_MAX;

    int32_t prev_event_idx = -1;

    for (i = 0; i < n_events; ++i) {
        int32_t k_idx = event_alignment[i].ref_pos;
        int32_t event_idx = event_alignment[i].read_pos;
        index_pair_t* elem = &base_to_event_map[k_idx];
        //fprintf(stderr, "eventpar %d %d k_idx %d event_idx %d\n",elem.start, elem.stop,k_idx,event_idx);
        if (event_idx != prev_event_idx) {
            if (elem->start == -1) {
                elem->start = event_idx;
            }
            elem->stop = event_idx;
        }
        max_event = max_event > event_idx ? max_event : event_idx;
        min_event = min_event < event_idx ? min_event : event_idx;
        prev_event_idx = event_idx;
    }

    // for (i = 0; i < n_kmers; ++i) {
    //     fprintf(stderr,"base_to_event_map - start %d stop %d\n", base_to_event_map[i].start,base_to_event_map[i].stop);
    // }

    *events_per_base = (double)(max_event - min_event) / n_kmers;

    /*prepare data structures for the final calibration*/

    int32_t alignment_index = 0;
    int32_t prev_kmer_rank = -1;

    int32_t ki;
    for (ki = 0; ki < n_kmers; ++ki) {
        index_pair_t event_range_for_kmer = base_to_event_map[ki];

        //fprintf(stderr, "kindex %d base_to_event_map - start %d stop %d\n",ki,event_range_for_kmer.start, event_range_for_kmer.stop);

        // skip kmers without events
        if (event_range_for_kmer.start == -1) {
            continue;
        }

        //skip k-mers that cannot be shifted to a valid position
        // int32_t shift_offset=0;
        // if(ki + shift_offset < 0 || ki + shift_offset >= n_kmers) {
        //     continue;
        // }

        for (int32_t event_idx = event_range_for_kmer.start;
             event_idx <= event_range_for_kmer.stop; event_idx++) {
            //fprintf(stderr,"event idx %d n events %d\n",event_idx,n_events);
            // assert(event_idx < n_events);

            // since we use the 1D read seqence here we never have to reverse complement
            int32_t kmer_rank = get_kmer_rank(&sequence[ki], KMER_SIZE);

            event_alignment_t ea;
            // ref data
            //ea.ref_name = "read";
            ea.read_idx = -1; // not needed
            kmer_cpy(ea.ref_kmer, &sequence[ki], KMER_SIZE);
            ea.ref_position = ki;
            //ea.strand_idx = strand_idx;
            ea.event_idx = event_idx;
            ea.rc = false;
            kmer_cpy(ea.model_kmer, &sequence[ki], KMER_SIZE);
            ea.hmm_state = prev_kmer_rank != kmer_rank ? 'M' : 'E';
            if (alignment_index >
                n_events) { //todo : this is ugly. check and fix.
                ERROR("We have run out of space in event_alignment_t* "
                      "alignment. Assumption fialed. Current size %d",
                      n_events);
                exit(EXIT_FAILURE);
            }
            alignment[alignment_index] = ea;
            alignment_index++;
            prev_kmer_rank = kmer_rank;
        }
        //fprintf(stderr,"event idx : %d\n",alignment_index);
    }

    //free(base_to_event_map);
    return alignment_index;
}

// recalculate shift, scale, drift, scale_sd from an alignment and the read
// returns true if the recalibration was performed
// in either case, sets residual to the L1 norm of the residual
bool recalibrate_model(model_t* pore_model, event_table et,
                       scalings_t* scallings,
                       const event_alignment_t* alignment_output,
                       int32_t num_alignments, bool scale_var) {
    //std::vector<double> raw_events, times, level_means, level_stdvs;
    //std::cout << "Previous pore model parameters: " << sr.pore_model[strand_idx].shift << ", "
    //                                                << sr.pore_model[strand_idx].scale << ", "
    //                                                << sr.pore_model[strand_idx].drift << ", "
    //                                                << sr.pore_model[strand_idx].var   << std::endl;

    // extract necessary vectors from the read and the pore model; note do not want scaled values
    int32_t num_M_state = 0;
    for (int32_t ei = 0; ei < num_alignments; ++ei) {
        event_alignment_t ea = alignment_output[ei];
        if (ea.hmm_state == 'M') {
            num_M_state++;
            //
            //fprintf(stdout, "recalibrate ei: %zu level: %.2lf kmer: %s model: %.2lf\n",
            //        ei, sr.get_uncorrected_level(ea.event_idx, strand_idx), model_kmer.c_str(),
            //        sr.pore_model[strand_idx].states[rank].level_mean);
            //
        }
    }

    const int32_t minNumEventsToRescale = 200;
    bool recalibrated = false;
    if (num_M_state >= minNumEventsToRescale) {
        // Assemble linear system corresponding to weighted least squares problem
        // Can just directly call a weighted least squares solver, but there's enough
        // structure in our problem it's a little faster just to build the normal eqn
        // matrices ourselves

        double A00 = 0, A01 = 0, A10 = 0, A11 = 0;
        double b0 = 0, b1 = 0;
        double x0 = 0, x1 = 0;

        for (int32_t ei = 0; ei < num_alignments; ++ei) {
            event_alignment_t ea = alignment_output[ei];
            if (ea.hmm_state == 'M') {
                //std::string model_kmer = ea.rc ? pore_model.pmalphabet->reverse_complement(ea.ref_kmer) : ea.ref_kmer;
                uint32_t rank = get_kmer_rank(ea.ref_kmer, KMER_SIZE);

                double raw_event = et.event[ea.event_idx].mean;
                double level_mean = pore_model[rank].level_mean;
                double level_stdv = pore_model[rank].level_stdv;

                double inv_var = 1. / (level_stdv * level_stdv);
                double mu = level_mean;
                double e = raw_event;

                A00 += inv_var;
                A01 += mu * inv_var;
                A11 += mu * mu * inv_var;

                b0 += e * inv_var;
                b1 += mu * e * inv_var;
            }
        }

        A10 = A01;

        // perform the linear solve
        //Eigen::VectorXd x = A.fullPivLu().solve(b);
        double div = A00 * A11 - A01 * A10;
        x0 = -(A01 * b1 - A11 * b0) / div;
        x1 = (A00 * b1 - A10 * b0) / div;

        double shift = x0;
        double scale = x1;
        //double drift = 0.;
        double var = 1.0;

        if (scale_var) {
            var = 0.;
            for (int32_t ei = 0; ei < num_alignments; ++ei) {
                event_alignment_t ea = alignment_output[ei];
                if (ea.hmm_state == 'M') {
                    uint32_t rank = get_kmer_rank(ea.ref_kmer, KMER_SIZE);
                    double raw_event = et.event[ea.event_idx].mean;
                    double level_mean = pore_model[rank].level_mean;
                    double level_stdv = pore_model[rank].level_stdv;
                    double yi = (raw_event - shift - scale * level_mean);
                    var += yi * yi / (level_stdv * level_stdv);
                }
            }
            var /= num_M_state;
            var = sqrt(var);
        }

        scallings->shift = shift;
        scallings->scale = scale;
        //scallings->drift=drift;
        scallings->var = var;
#ifdef CACHED_LOG
        scallings->log_var = log(var);
#endif

        recalibrated = true;

#ifdef DEBUG_RECALIB_SCALING
        fprintf(stderr, "shift: %.2lf scale: %.2lf var: %.2lf\n",
                scallings->shift, scallings->scale, scallings->var);
        //fprintf(stderr, "truth shift: %.2lf scale: %.2lf\n", pore_model.shift, pore_model.scale);
#endif
    }

    return recalibrated;
}
```
