## Supplementary material for "GPU Accelerated Adaptive Banded Event Alignment for Rapid Comparative Nanopore Signal Analysis": f5c-v0.2-beta-release: align.cu

```
#include "f5c.h"
#include <assert.h>
#include "f5cmisc.cuh"

//#define DEBUG_ESTIMATED_SCALING 1
//#define DEBUG_RECALIB_SCALING 1
//#define DEBUG_ADAPTIVE 1


//todo : can make more efficient using bit encoding
//todo : is inlining correct?
__forceinline__ __device__  uint32_t get_rank(char base) {
    if (base == 'A') { //todo: do we neeed simple alpha?
        return 0;
    } else if (base == 'C') {
        return 1;
    } else if (base == 'G') {
        return 2;
    } else if (base == 'T') {
        return 3;
    } else {
        //WARNING("A None ACGT base found : %c", base); //todo : fix this in gpu code
        return 0;
    }
}

// return the lexicographic rank of the kmer amongst all strings of
// length k for this alphabet
__forceinline__ __device__  uint32_t get_kmer_rank(const char* str, uint32_t k) {
    //uint32_t p = 1;
    uint32_t r = 0;

    // from last base to first
    for (uint32_t i = 0; i < k; ++i) {
        //r += rank(str[k - i - 1]) * p;
        //p *= size();
        r += get_rank(str[k - i - 1]) << (i << 1);
    }
    return r;
}

//copy a kmer from a reference
__forceinline__ __device__ void kmer_cpy(char* dest, char* src, uint32_t k) {
    uint32_t i = 0;
    for (i = 0; i < k; i++) {
        dest[i] = src[i];
    }
    dest[i] = '\0';
}

#define log_inv_sqrt_2pi  -0.918938f // Natural logarithm

__forceinline__ __device__ float
log_normal_pdf(float x, float gp_mean, float gp_stdv, float gp_log_stdv) {
    /*INCOMPLETE*/
    //float log_inv_sqrt_2pi = -0.918938f; // Natural logarithm
    float a = (x - gp_mean) / gp_stdv;
    return log_inv_sqrt_2pi - gp_log_stdv + (-0.5f * a * a);
    // return 1;
}

__forceinline__ __device__ float
log_probability_match_r9(scalings_t scaling, model_t* models, event_t* event,
                         int event_idx, uint32_t kmer_rank) {
    // event level mean, scaled with the drift value
    //strand = 0;
 #ifdef DEBUG_ADAPTIVE   
    assert(kmer_rank < 4096);
 #endif   
    //float level = read.get_drift_scaled_level(event_idx, strand);

    //float time =
    //    (events.event[event_idx].start - events.event[0].start) / sample_rate;
    float unscaledLevel = event[event_idx].mean;
    float scaledLevel = unscaledLevel;
    //float scaledLevel = unscaledLevel - time * scaling.shift;

    //fprintf(stderr, "level %f\n",scaledLevel);
    //GaussianParameters gp = read.get_scaled_gaussian_from_pore_model_state(pore_model, strand, kmer_rank);
    float gp_mean =
        scaling.scale * models[kmer_rank].level_mean + scaling.shift;
    float gp_stdv = models[kmer_rank].level_stdv * 1; //scaling.var = 1;
    // float gp_stdv = 0;
    // float gp_log_stdv = models[kmer_rank].level_log_stdv + scaling.log_var;
    // if(models[kmer_rank].level_stdv <0.01 ){
    // 	fprintf(stderr,"very small std dev %f\n",models[kmer_rank].level_stdv);
    // }
    #ifdef CACHED_LOG
        float gp_log_stdv = models[kmer_rank].level_log_stdv;
    #else
        float gp_log_stdv =
        log(models[kmer_rank].level_stdv); // scaling.log_var = log(1)=0;
    #endif

    float lp = log_normal_pdf(scaledLevel, gp_mean, gp_stdv, gp_log_stdv);
    return lp;
}

#define event_kmer_to_band(ei, ki) (ei + 1) + (ki + 1)
#define band_event_to_offset(bi, ei) band_lower_left[bi].event_idx - (ei)
#define band_kmer_to_offset(bi, ki) (ki) - band_lower_left[bi].kmer_idx
#define is_offset_valid(offset) (offset) >= 0 && (offset) < bandwidth
#define event_at_offset(bi, offset) band_lower_left[(bi)].event_idx - (offset)
#define kmer_at_offset(bi, offset) band_lower_left[(bi)].kmer_idx + (offset)

#define move_down(curr_band)                                                   \
    { curr_band.event_idx + 1, curr_band.kmer_idx }
#define move_right(curr_band)                                                  \
    { curr_band.event_idx, curr_band.kmer_idx + 1 }

#define MIN(a, b) (((a) < (b)) ? (a) : (b))
#define MAX(a, b) (((a) > (b)) ? (a) : (b))

#define BAND_ARRAY(r, c) ( bands[((r)*(ALN_BANDWIDTH)+(c))] )
#define TRACE_ARRAY(r, c) ( trace[((r)*(ALN_BANDWIDTH)+(c))] )

#define FROM_D  0
#define FROM_U  1
#define FROM_L  2


#define max_gap_threshold  50
#define bandwidth  ALN_BANDWIDTH
#define half_bandwidth  ALN_BANDWIDTH/2

#ifndef ALIGN_KERNEL_FLOAT
    #define min_average_log_emission  -5.0
    #define epsilon 1e-10
#else
    #define min_average_log_emission -5.0f
    #define epsilon 1e-10f
#endif    


/************************kernels with 2D thread models*****************/


__global__ void align_kernel_pre_2d(char* read,
    int32_t* read_len, ptr_t* read_ptr,
    int32_t* n_events,
    ptr_t* event_ptr, model_t* models,
    int32_t n_bam_rec,model_t* model_kmer_caches,float *bands1,uint8_t *trace1, EventKmerPair* band_lower_left1) {


    int i = blockDim.y * blockIdx.y + threadIdx.y;
    int tid=blockIdx.x*blockDim.x+threadIdx.x;


    if (i < n_bam_rec) {
        char* sequence = &read[read_ptr[i]];
        int32_t sequence_len = read_len[i];
        //int32_t n_event = n_events[i];
        model_t* model_kmer_cache = &model_kmer_caches[read_ptr[i]];
        float *bands = &bands1[(read_ptr[i]+event_ptr[i])*ALN_BANDWIDTH];
        uint8_t *trace = &trace1[(read_ptr[i]+event_ptr[i])*ALN_BANDWIDTH];
        EventKmerPair* band_lower_left = &band_lower_left1[read_ptr[i]+event_ptr[i]];    

        //int32_t n_events = n_event;
        int32_t n_kmers = sequence_len - KMER_SIZE + 1;
        //fprintf(stderr,"n_kmers : %d\n",n_kmers);

        // transition penalties
        // float events_per_kmer = (float)n_events / n_kmers;
        // float p_stay = 1 - (1 / (events_per_kmer + 1));

        // setting a tiny skip penalty helps keep the true alignment within the adaptive band
        // this was empirically determined
        //double epsilon = 1e-10;
        // double lp_skip = log(epsilon);
        // double lp_stay = log(p_stay);
        // double lp_step = log(1.0 - exp(lp_skip) - exp(lp_stay));
        #ifndef ALIGN_KERNEL_FLOAT
            double lp_trim = log(0.01);
        #else
            float lp_trim = logf(0.01f);
        #endif

        // dp matrix
        //int32_t n_rows = n_events + 1;
        //int32_t n_cols = n_kmers + 1;
        //int32_t n_bands = n_rows + n_cols;

        // Initialize
        // Precompute k-mer ranks to avoid doing this in the inner loop

    // #ifdef  PRE_3D    
    //     if(band_i<n_kmers && band_j==0){
    // #else
    //     if(band_i<n_kmers){
    // #endif

        if(tid==0){ //todo : can be optimised
            for (int32_t i = 0; i < n_kmers; ++i) {
                char* substring = &sequence[i];
                //kmer_ranks[i] = get_kmer_rank(substring, KMER_SIZE);
                uint32_t kmer_ranks = get_kmer_rank(substring, KMER_SIZE);
                model_kmer_cache[i] = models[kmer_ranks];            }
        }

        if(tid<bandwidth){
            for (int32_t i = 0; i < 3; i++) {
                    BAND_ARRAY(i,tid) = -INFINITY;
                    //TRACE_ARRAY(i,tid) = 0;
            }
        }

        if(tid==0){
            // initialize range of first two bands
            band_lower_left[0].event_idx = half_bandwidth - 1;
            band_lower_left[0].kmer_idx = -1 - half_bandwidth;
            band_lower_left[1] = move_down(band_lower_left[0]);

            int start_cell_offset = band_kmer_to_offset(0, -1);
            assert(is_offset_valid(start_cell_offset));
            assert(band_event_to_offset(0, -1) == start_cell_offset);
            BAND_ARRAY(0,start_cell_offset) = 0.0f;

            // band 1: first event is trimmed
            int first_trim_offset = band_event_to_offset(1, 0);
            assert(kmer_at_offset(1, first_trim_offset) == -1);
            assert(is_offset_valid(first_trim_offset));
            BAND_ARRAY(1,first_trim_offset) = lp_trim;
            TRACE_ARRAY(1,first_trim_offset) = FROM_U;

            //int fills = 0;
        #ifdef DEBUG_ADAPTIVE
            fprintf(stderr, "[trim] bi: %d o: %d e: %d k: %d s: %.2lf\n", 1,
                    first_trim_offset, 0, -1, BAND_ARRAY(1,first_trim_offset);
        #endif

        }
    }
}


#define PROFILE 1

#define band_event_to_offset_shm(bi, ei) band_lower_left_shm[bi].event_idx - (ei)
#define band_kmer_to_offset_shm(bi, ki) (ki) - band_lower_left_shm[bi].kmer_idx

#define event_at_offset_shm(bi, offset) band_lower_left_shm[(bi)].event_idx - (offset)
#define kmer_at_offset_shm(bi, offset) band_lower_left_shm[(bi)].kmer_idx + (offset)

#define BAND_ARRAY_SHM(r, c) ( bands_shm[(r)][(c)] )
__global__ void 
//__launch_bounds__(MY_KERNEL_MAX_THREADS, MY_KERNEL_MIN_BLOCKS)
align_kernel_core_2d_shm(int32_t* read_len, ptr_t* read_ptr,
    event_t* event_table, int32_t* n_events1,
    ptr_t* event_ptr, 
    scalings_t* scalings, int32_t n_bam_rec,model_t* model_kmer_caches,float *band,uint8_t *traces, EventKmerPair* band_lower_lefts) {
   
    int i = blockDim.y * blockIdx.y + threadIdx.y;
    int offset=blockIdx.x*blockDim.x+threadIdx.x;

    __shared__ float  bands_shm[3][ALN_BANDWIDTH];
    __shared__ EventKmerPair  band_lower_left_shm[3];

    if (i < n_bam_rec && offset<ALN_BANDWIDTH) {   

        int32_t sequence_len = read_len[i];
        event_t* events = &event_table[event_ptr[i]];
        int32_t n_event = n_events1[i];
        scalings_t scaling = scalings[i];
        model_t* model_kmer_cache = &model_kmer_caches[read_ptr[i]];
        float *bands = &band[(read_ptr[i]+event_ptr[i])*ALN_BANDWIDTH];
        uint8_t *trace = &traces[(read_ptr[i]+event_ptr[i])*ALN_BANDWIDTH];
        EventKmerPair* band_lower_left = &band_lower_lefts[read_ptr[i]+event_ptr[i]];;

        // size_t n_events = events[strand_idx].n;
        int32_t n_events = n_event;
        int32_t n_kmers = sequence_len - KMER_SIZE + 1;
        //fprintf(stderr,"n_kmers : %d\n",n_kmers);

        // transition penalties
        float events_per_kmer = (float)n_events / n_kmers;
        float p_stay = 1 - (1 / (events_per_kmer + 1));

        // setting a tiny skip penalty helps keep the true alignment within the adaptive band
        // this was empirically determined
        //double epsilon = 1e-10;

#ifndef ALIGN_KERNEL_FLOAT       
        double lp_skip = log(epsilon);
        double lp_stay = log(p_stay);
        double lp_step = log(1.0 - exp(lp_skip) - exp(lp_stay));
        double lp_trim = log(0.01);
#else
        float lp_skip = logf(epsilon);
        float lp_stay = logf(p_stay);
        float lp_step = logf(1.0f - expf(lp_skip) - expf(lp_stay));
        float lp_trim = logf(0.01f);
#endif
        // dp matrix
        int32_t n_rows = n_events + 1;
        int32_t n_cols = n_kmers + 1;
        int32_t n_bands = n_rows + n_cols;                                    


        BAND_ARRAY_SHM(0,offset) = BAND_ARRAY(2,offset);
        BAND_ARRAY_SHM(1,offset) = BAND_ARRAY(1,offset);
        BAND_ARRAY_SHM(2,offset) = BAND_ARRAY(0,offset);

        band_lower_left_shm[0] = band_lower_left[2];
        band_lower_left_shm[1] = band_lower_left[1];
        band_lower_left_shm[2] = band_lower_left[0];

        __syncthreads();

        // fill in remaining bands
        for (int32_t band_idx = 2; band_idx < n_bands; ++band_idx) {
            
            if(offset==0){
                // Determine placement of this band according to Suzuki's adaptive algorithm
                // When both ll and ur are out-of-band (ob) we alternate movements
                // otherwise we decide based on scores
                //float ll = BAND_ARRAY((band_idx - 1), 0);
                float ll = BAND_ARRAY_SHM((1), 0);
                //float ur = BAND_ARRAY((band_idx - 1),(bandwidth - 1));
                float ur = BAND_ARRAY_SHM((1),(bandwidth - 1));
                bool ll_ob = ll == -INFINITY;
                bool ur_ob = ur == -INFINITY;

                bool right = false;
                if (ll_ob && ur_ob) {
                    right = band_idx % 2 == 1;
                } else {
                    right = ll < ur; // Suzuki's rule
                }

                if (right) {
                    band_lower_left[band_idx] = band_lower_left_shm[0] =
                        move_right(band_lower_left_shm[1]);                        
                } else {
                    band_lower_left[band_idx] = band_lower_left_shm[0] =
                        move_down(band_lower_left_shm[1]);                        
                }
                // If the trim state is within the band, fill it in here
                int trim_offset = band_kmer_to_offset_shm(0, -1);
                if (is_offset_valid(trim_offset)) {
                    int32_t event_idx = event_at_offset_shm(0, trim_offset);
                    if (event_idx >= 0 && event_idx < n_events) {
                        //BAND_ARRAY(band_idx,trim_offset) = lp_trim * (event_idx + 1);
                        BAND_ARRAY_SHM(0,trim_offset) = lp_trim * (event_idx + 1);
                        TRACE_ARRAY(band_idx,trim_offset) = FROM_U;
                    } else {
                        //BAND_ARRAY(band_idx,trim_offset) = -INFINITY;
                        BAND_ARRAY_SHM(0,trim_offset) = -INFINITY;
                    }
                }
            }
            __syncthreads();

            // Get the offsets for the first and last event and kmer
            // We restrict the inner loop to only these values
            int kmer_min_offset = band_kmer_to_offset_shm(0, 0);
            int kmer_max_offset = band_kmer_to_offset_shm(0, n_kmers);
            int event_min_offset = band_event_to_offset_shm(0, n_events - 1);
            int event_max_offset = band_event_to_offset_shm(0, -1);

            int min_offset = MAX(kmer_min_offset, event_min_offset);
            min_offset = MAX(min_offset, 0);

            int max_offset = MIN(kmer_max_offset, event_max_offset);
            max_offset = MIN(max_offset, bandwidth);

            __syncthreads();    
   
            if(offset>=min_offset && offset< max_offset) {

                int event_idx = event_at_offset_shm(0, offset);
                int kmer_idx = kmer_at_offset_shm(0, offset);

                //int32_t kmer_rank = kmer_ranks[kmer_idx];

                int offset_up = band_event_to_offset_shm(1, event_idx - 1);
                int offset_left = band_kmer_to_offset_shm(1, kmer_idx - 1);
                int offset_diag = band_kmer_to_offset_shm(2, kmer_idx - 1);

    #ifdef DEBUG_ADAPTIVE
                // verify loop conditions
                assert(kmer_idx >= 0 && kmer_idx < n_kmers);
                assert(event_idx >= 0 && event_idx < n_events);
                assert(offset_diag ==
                       band_event_to_offset_shm(2, event_idx - 1));
                assert(offset_up - offset_left == 1);
                assert(offset >= 0 && offset < bandwidth);
    #endif //DEBUG_ADAPTIVE

                float up = is_offset_valid(offset_up)
                               ? BAND_ARRAY_SHM(1,offset_up)
                               : -INFINITY;
                float left = is_offset_valid(offset_left)
                                 ? BAND_ARRAY_SHM(1,offset_left)
                                 : -INFINITY;
                float diag = is_offset_valid(offset_diag)
                                 ? BAND_ARRAY_SHM(2,offset_diag)
                                 : -INFINITY;

            #ifndef PROFILE
                float lp_emission = log_probability_match_r9(
                    scaling, model_kmer_cache, events, event_idx,kmer_idx);
                //fprintf(stderr, "lp emiision : %f , event idx %d, kmer rank %d\n", lp_emission,event_idx,kmer_rank);
            #else
                float unscaledLevel = events[event_idx].mean;
                float scaledLevel = unscaledLevel;
                model_t model = model_kmer_cache[kmer_idx];
                float gp_mean =
                    scaling.scale * model.level_mean + scaling.shift;
                float gp_stdv = model.level_stdv ; //scaling.var = 1;
                
                #ifdef  CACHED_LOG
                    float gp_log_stdv = model.level_log_stdv;
                #else
                    #ifndef ALIGN_KERNEL_FLOAT  
                        float gp_log_stdv = log(gp_stdv); // scaling.log_var = log(1)=0;
                    #else
                        float gp_log_stdv = logf(gp_stdv); // scaling.log_var = log(1)=0;
                    #endif  
                #endif

                float a = (scaledLevel - gp_mean) / gp_stdv;
                float lp_emission  = log_inv_sqrt_2pi - gp_log_stdv + (-0.5f * a * a);

            #endif    


                float score_d = diag + lp_step + lp_emission;
                float score_u = up + lp_stay + lp_emission;
                float score_l = left + lp_skip;

                float max_score = score_d;
                uint8_t from = FROM_D;

                max_score = score_u > max_score ? score_u : max_score;
                from = max_score == score_u ? FROM_U : from;
                max_score = score_l > max_score ? score_l : max_score;
                from = max_score == score_l ? FROM_L : from;

    #ifdef DEBUG_ADAPTIVE
                fprintf(stderr,
                        "[adafill] offset-up: %d offset-diag: %d offset-left: %d\n",
                        offset_up, offset_diag, offset_left);
                fprintf(stderr, "[adafill] up: %.2lf diag: %.2lf left: %.2lf\n", up,
                        diag, left);
                fprintf(stderr,
                        "[adafill] bi: %d o: %d e: %d k: %d s: %.2lf f: %d emit: "
                        "%.2lf\n",
                        band_idx, offset, event_idx, kmer_idx, max_score, from,
                        lp_emission);
    #endif //DEBUG_ADAPTIVE
                //BAND_ARRAY(band_idx,offset) = max_score;
                BAND_ARRAY_SHM(0,offset) = max_score;
                TRACE_ARRAY(band_idx,offset) = from;
                //fills += 1;
            }

            

            __syncthreads();
            BAND_ARRAY(band_idx,offset) = BAND_ARRAY_SHM(0,offset);
            
            BAND_ARRAY_SHM(2,offset) = BAND_ARRAY_SHM(1,offset);
            BAND_ARRAY_SHM(1,offset) = BAND_ARRAY_SHM(0,offset);
            BAND_ARRAY_SHM(0,offset) = -INFINITY;

            if(offset==0){
                band_lower_left_shm[2]=band_lower_left_shm[1];
                band_lower_left_shm[1]=band_lower_left_shm[0];
            }


            __syncthreads();  

        }
    }
}


//align post kernel
__global__ void align_kernel_post(AlignedPair* event_align_pairs,
    int32_t* n_event_align_pairs, 
    int32_t* read_len, ptr_t* read_ptr,
    event_t* event_table, int32_t* n_events,
    ptr_t* event_ptr, 
    scalings_t* scalings, int32_t n_bam_rec,model_t* model_kmer_caches,float *bands1,uint8_t *trace1, EventKmerPair* band_lower_left1) {

    #ifndef WARP_HACK        
        int i = blockDim.x * blockIdx.x + threadIdx.x;
        if (i < n_bam_rec) {
    #else
        int tid = blockDim.x * blockIdx.x + threadIdx.x;
        int i = tid/32;    
        if (i < n_bam_rec && tid%32==0) {
    #endif  
        AlignedPair* out_2 = &event_align_pairs[event_ptr[i] * 2];
        int32_t sequence_len = read_len[i];
        event_t* events = &event_table[event_ptr[i]];
        int32_t n_event = n_events[i];
        scalings_t scaling = scalings[i];
        model_t* model_kmer_cache = &model_kmer_caches[read_ptr[i]];
        float *bands = &bands1[(read_ptr[i]+event_ptr[i])*ALN_BANDWIDTH];
        uint8_t *trace = &trace1[(read_ptr[i]+event_ptr[i])*ALN_BANDWIDTH];
        EventKmerPair* band_lower_left = &band_lower_left1[read_ptr[i]+event_ptr[i]];;
 
        //fprintf(stderr, "%s\n", sequence);
        //fprintf(stderr, "Scaling %f %f", scaling.scale, scaling.shift);

        //size_t strand_idx = 0;
        //size_t k = 6;

        // size_t n_events = events[strand_idx].n;
        int32_t n_events = n_event;
        int32_t n_kmers = sequence_len - KMER_SIZE + 1;
        //fprintf(stderr,"n_kmers : %d\n",n_kmers);
        // backtrack markers
        //const uint8_t FROM_D = 0;
        //const uint8_t FROM_U = 1;
        //const uint8_t FROM_L = 2;

        // qc
        //double min_average_log_emission = -5.0;
        //int max_gap_threshold = 50;

        // banding
        //int bandwidth = ALN_BANDWIDTH;
        //half_bandwidth = bandwidth / 2;

        // transition penalties
        float events_per_kmer = (float)n_events / n_kmers;
        float p_stay = 1 - (1 / (events_per_kmer + 1));

        // setting a tiny skip penalty helps keep the true alignment within the adaptive band
        // this was empirically determined
        //double epsilon = 1e-10;
#ifndef ALIGN_KERNEL_FLOAT       
        double lp_skip = log(epsilon);
        double lp_stay = log(p_stay);
        double lp_step = log(1.0 - exp(lp_skip) - exp(lp_stay));
        double lp_trim = log(0.01);
#else
        float lp_skip = logf(epsilon);
        float lp_stay = logf(p_stay);
        float lp_step = logf(1.0f - expf(lp_skip) - expf(lp_stay));
        float lp_trim = logf(0.01f);
#endif        
        // dp matrix
        int32_t n_rows = n_events + 1;
        int32_t n_cols = n_kmers + 1;
        int32_t n_bands = n_rows + n_cols;
        //
        // Backtrack to compute alignment
        //
        double sum_emission = 0;
        double n_aligned_events = 0;

        //>>>>>>>>>>>>>> New replacement begin
        // std::vector<AlignedPair> out;

        int outIndex = 0;
        //<<<<<<<<<<<<<<<<New Replacement over

        float max_score = -INFINITY;
        int curr_event_idx = 0;
        int curr_kmer_idx = n_kmers - 1;

        // Find best score between an event and the last k-mer. after trimming the remaining evnets
        for (int32_t event_idx = 0; event_idx < n_events; ++event_idx) {
            int band_idx = event_kmer_to_band(event_idx, curr_kmer_idx);

            //>>>>>>>New  replacement begin
            /*assert(band_idx < bands.size());*/

            assert(band_idx < n_bands);

            //<<<<<<<<New Replacement over
            int offset = band_event_to_offset(band_idx, event_idx);
            if (is_offset_valid(offset)) {
                float s =
                    BAND_ARRAY(band_idx,offset) + (n_events - event_idx) * lp_trim;
                if (s > max_score) {
                    max_score = s;
                    curr_event_idx = event_idx;
                }
            }
        }

    #ifdef DEBUG_ADAPTIVE
        fprintf(stderr, "[adaback] ei: %d ki: %d s: %.2f\n", curr_event_idx,
                curr_kmer_idx, max_score);
    #endif

        int curr_gap = 0;
        int max_gap = 0;
        while (curr_kmer_idx >= 0 && curr_event_idx >= 0) {
            // emit alignment
            //>>>>>>>New Repalcement begin
            assert(outIndex < n_events * 2);
            out_2[outIndex].ref_pos = curr_kmer_idx;
            out_2[outIndex].read_pos = curr_event_idx;
            outIndex++;
            // out.push_back({curr_kmer_idx, curr_event_idx});
            //<<<<<<<<<New Replacement over

    #ifdef DEBUG_ADAPTIVE
            fprintf(stderr, "[adaback] ei: %d ki: %d\n", curr_event_idx,
                    curr_kmer_idx);
    #endif
            // qc stats
            //>>>>>>>>>>>>>>New Replacement begin
            // char* substring = &sequence[curr_kmer_idx];
            // int32_t kmer_rank = get_kmer_rank(substring, KMER_SIZE);
            // //<<<<<<<<<<<<<New Replacement over
            // float tempLogProb = log_probability_match_r9(
            //     scaling, models, events, curr_event_idx, kmer_rank);

            #ifndef PROFILE
                float tempLogProb = log_probability_match_r9(
                    scaling, model_kmer_cache, events, curr_event_idx,curr_kmer_idx);
                //fprintf(stderr, "lp emiision : %f , event idx %d, kmer rank %d\n", lp_emission,event_idx,kmer_rank);
            #else
                float unscaledLevel = events[curr_event_idx].mean;
                float scaledLevel = unscaledLevel;
                model_t model = model_kmer_cache[curr_kmer_idx];
                float gp_mean =
                    scaling.scale * model.level_mean + scaling.shift;
                float gp_stdv = model.level_stdv ; //scaling.var = 1;
                
                #ifdef  CACHED_LOG
                    float gp_log_stdv = model.level_log_stdv;
                #else
                    #ifndef ALIGN_KERNEL_FLOAT  
                        float gp_log_stdv = log(gp_stdv); // scaling.log_var = log(1)=0;
                    #else
                        float gp_log_stdv = logf(gp_stdv); // scaling.log_var = log(1)=0;
                    #endif  
                #endif

                float a = (scaledLevel - gp_mean) / gp_stdv;
                float tempLogProb  = log_inv_sqrt_2pi - gp_log_stdv + (-0.5f * a * a);

            #endif 
                

            sum_emission += tempLogProb;
            //fprintf(stderr, "lp_emission %f \n", tempLogProb);
            //fprintf(stderr,"lp_emission %f, sum_emission %f, n_aligned_events %d\n",tempLogProb,sum_emission,outIndex);

            n_aligned_events += 1;

            int band_idx = event_kmer_to_band(curr_event_idx, curr_kmer_idx);
            int offset = band_event_to_offset(band_idx, curr_event_idx);
            assert(band_kmer_to_offset(band_idx, curr_kmer_idx) == offset);

            uint8_t from = TRACE_ARRAY(band_idx,offset);
            if (from == FROM_D) {
                curr_kmer_idx -= 1;
                curr_event_idx -= 1;
                curr_gap = 0;
            } else if (from == FROM_U) {
                curr_event_idx -= 1;
                curr_gap = 0;
            } else {
                curr_kmer_idx -= 1;
                curr_gap += 1;
                max_gap = MAX(curr_gap, max_gap);
            }
        }

    
#ifndef REVERSAL_ON_CPU
        //>>>>>>>>New replacement begin
        // std::reverse(out.begin(), out.end());
        int c;
        int end = outIndex - 1;
        for (c = 0; c < outIndex / 2; c++) {
            int ref_pos_temp = out_2[c].ref_pos;
            int read_pos_temp = out_2[c].read_pos;
            out_2[c].ref_pos = out_2[end].ref_pos;
            out_2[c].read_pos = out_2[end].read_pos;
            out_2[end].ref_pos = ref_pos_temp;
            out_2[end].read_pos = read_pos_temp;
            end--;
        }

        // if(outIndex>1){
        //   AlignedPair temp={out_2[0].ref_pos,out[0].read_pos};
        //   int i;
        //   for(i=0;i<outIndex-1;i++){
        //     out_2[i]={out_2[outIndex-1-i].ref_pos,out[outIndex-1-i].read_pos};
        //   }
        //   out[outIndex-1]={temp.ref_pos,temp.read_pos};
        // }
        //<<<<<<<<<New replacement over

        //>>>>>>>>>>>>>New replacement begin
        bool spanned = out_2[0].ref_pos == 0 &&
                    out_2[outIndex - 1].ref_pos == int(n_kmers - 1);

        //assert(spanned==spanned_before_rev);            
        // bool spanned = out.front().ref_pos == 0 && out.back().ref_pos == n_kmers - 1;
        //<<<<<<<<<<<<<New replacement over        
#else
        bool spanned = out_2[outIndex - 1].ref_pos == 0 &&
                    out_2[0].ref_pos == int(n_kmers - 1);
#endif
        // QC results
        double avg_log_emission = sum_emission / n_aligned_events;
        //fprintf(stderr,"sum_emission %f, n_aligned_events %f, avg_log_emission %f\n",sum_emission,n_aligned_events,avg_log_emission);

        //bool failed = false;
        if (avg_log_emission < min_average_log_emission || !spanned ||
            max_gap > max_gap_threshold) {
            //failed = true;
            //>>>>>>>>>>>>>New replacement begin
            outIndex = 0;
            // out.clear();
            //free(out_2);
            //out_2 = NULL;
            //<<<<<<<<<<<<<New replacement over
        }

        // free(kmer_ranks);
        // for (size_t i = 0; i < n_bands; i++) {
        //     free(bands[i]);
        //     free(trace[i]);
        // }
        // free(bands);
        // free(trace);
        // free(band_lower_left);
        //fprintf(stderr, "ada\t%s\t%s\t%.2lf\t%zu\t%.2lf\t%d\t%d\t%d\n", read.read_name.substr(0, 6).c_str(), failed ? "FAILED" : "OK", events_per_kmer, sequence.size(), avg_log_emission, curr_event_idx, max_gap, fills);
        //outSize=outIndex;
        //if(outIndex>500000)fprintf(stderr, "Max outSize %d\n", outIndex);
        n_event_align_pairs[i] = outIndex;
    
    }
}
```
