## Supplementary material for "GPU Accelerated Adaptive Banded Event Alignment for Rapid Comparative Nanopore Signal Analysis": f5c-v0.2-beta-release: error.h

```
#ifndef ERROR_H
#define ERROR_H

#include <errno.h>

#define WARN "[%s::WARNING]\033[1;33m "
#define ERR "[%s::ERROR]\033[1;31m "
#define CEND "\033[0m\n"

#define STDERR(arg, ...)                                                      \
    fprintf(stderr, "[%s] " arg "\n", __func__,                        \
            __VA_ARGS__)
#define WARNING(arg, ...)                                                      \
    fprintf(stderr, "[%s::WARNING]\033[1;33m " arg "\033[0m\n", __func__,      \
            __VA_ARGS__)
#define ERROR(arg, ...)                                                        \
    fprintf(stderr, "[%s::ERROR]\033[1;31m " arg "\033[0m\n", __func__,        \
            __VA_ARGS__)
#define INFO(arg, ...)                                                         \
    fprintf(stderr, "[%s::INFO]\033[1;34m " arg "\033[0m\n", __func__,         \
            __VA_ARGS__)
#define SUCCESS(arg, ...)                                                      \
    fprintf(stderr, "[%s::SUCCESS]\033[1;32m " arg "\033[0m\n", __func__,      \
            __VA_ARGS__)
#define DEBUG(arg, ...)                                                        \
    fprintf(stderr,                                                            \
            "[%s::DEBUG]\033[1;35m Error occured at %s:%d. " arg "\033[0m\n",  \
            __func__, __FILE__, __LINE__ - 2, __VA_ARGS__)

#define MALLOC_CHK(ret) malloc_chk((void*)ret, __func__, __FILE__, __LINE__ - 1)
#define F_CHK(ret, filename)                                                   \
    f_chk((void*)ret, __func__, __FILE__, __LINE__ - 1, filename);
#define NULL_CHK(ret) null_chk((void*)ret, __func__, __FILE__, __LINE__ - 1)
#define NEG_CHK(ret) neg_chk(ret, __func__, __FILE__, __LINE__ - 1)

static inline void malloc_chk(void* ret, const char* func, const char* file,
                              int line) {
    if (ret != NULL)
        return;
    fprintf(
        stderr,
        "[%s::ERROR]\033[1;31m Failed to allocate memory : "
        "%s.\033[0m\n[%s::DEBUG]\033[1;35m Error occured at %s:%d. Try with a small batchsize (-K and/or -B options) to reduce the peak memory\033[0m\n\n",
        func, strerror(errno), func, file, line);
    exit(EXIT_FAILURE);
}

static inline void f_chk(void* ret, const char* func, const char* file,
                         int line, const char* fopen_f) {
    if (ret != NULL)
        return;
    fprintf(
        stderr,
        "[%s::ERROR]\033[1;31m Failed to open %s : "
        "%s.\033[0m\n[%s::DEBUG]\033[1;35m Error occured at %s:%d.\033[0m\n\n",
        func, fopen_f, strerror(errno), func, file, line);
    exit(EXIT_FAILURE);
}

// Die on error. Print the error and exit if the return value of the previous function NULL
static inline void null_chk(void* ret, const char* func, const char* file,
                            int line) {
    if (ret != NULL)
        return;
    fprintf(stderr,
            "[%s::ERROR]\033[1;31m %s.\033[0m\n[%s::DEBUG]\033[1;35m Error "
            "occured at %s:%d.\033[0m\n\n",
            func, strerror(errno), func, file, line);
    exit(EXIT_FAILURE);
}

// Die on error. Print the error and exit if the return value of the previous function is -1
static inline void neg_chk(int ret, const char* func, const char* file,
                           int line) {
    if (ret >= 0)
        return;
    fprintf(stderr,
            "[%s::ERROR]\033[1;31m %s.\033[0m\n[%s::DEBUG]\033[1;35m Error "
            "occured at %s:%d.\033[0m\n\n",
            func, strerror(errno), func, file, line);
    exit(EXIT_FAILURE);
}

#endif
```
