## Supplementary material for "GPU Accelerated Adaptive Banded Event Alignment for Rapid Comparative Nanopore Signal Analysis": f5c-v0.2-beta-release: eventalign.c

```
#include <assert.h>
#include <stdlib.h>
#include <stdio.h>
#include <stdint.h>
#include "matrix.h"
#include <algorithm>
#include <vector>
#include <string>

#include <fstream>
#include <string>
#include <iostream>

#include "f5c.h"
#include "f5cmisc.h"

/*
Code is adapted from Nanopolish eventalign module
Contains redundant code (duplicated code from meth.c and hmm.c) at the moment
Requires a thorough cleanup when everything is optimised
*/

#define METHYLATED_SYMBOL 'M'

#define TRANS_START_TO_CLIP 0.5
#define TRANS_CLIP_SELF 0.9

//copy a kmer from a reference
static inline void kmer_cpy(char* dest, const char* src, uint32_t k) {
    uint32_t i = 0;
    for (i = 0; i < k; i++) {
        dest[i] = src[i];
    }
    dest[i] = '\0';
}

typedef std::vector<AlignedPair> AlignedSegment;


enum ProfileStateR9
{
    PSR9_KMER_SKIP = 0,
    PSR9_BAD_EVENT,
    PSR9_MATCH,
    PSR9_NUM_STATES = 3,
    PSR9_PRE_SOFT // intentionally after PS_NUM_STATES
};

enum HMMMovementType
{
    HMT_FROM_SAME_M = 0,
    HMT_FROM_PREV_M,
    HMT_FROM_SAME_B,
    HMT_FROM_PREV_B,
    HMT_FROM_PREV_K,
    HMT_FROM_SOFT,
    HMT_NUM_MOVEMENT_TYPES
};
typedef struct { float x[HMT_NUM_MOVEMENT_TYPES]; } HMMUpdateScores;

// from https://github.com/jts/nanopolish/blob/3180474dc1a79c076a21e70bb4d19d2705b39b56/src/common/nanopolish_common.h
// A representation of an event->kmer alignment
struct HMMAlignmentState
{
    uint32_t event_idx;
    uint32_t kmer_idx;
    double l_posterior;
    double l_fm;
    double log_transition_probability;
    char state;
};

// Allocate a vector with the model probabilities of skipping the remaining
// events after the alignment of event i
inline std::vector<float> make_post_flanking(const uint32_t e_start,
                                             const uint32_t num_events,
                                             int8_t event_stride,
                                             uint32_t event_stop_idx)
{
    // post_flank[i] means that the i-th event was the last one
    // aligned and the remainder should be emitted from the background model
    std::vector<float> post_flank(num_events, 0.0f);

    // base case, all events aligned
    post_flank[num_events - 1] = log(1 - TRANS_START_TO_CLIP);

    if(num_events > 1) {
        // base case, all events aligned but 1
        {
            uint32_t event_idx = e_start + (num_events - 1) * event_stride; // last event
            assert(event_idx == event_stop_idx);
            // post_flank[num_events - 2] = log(TRANS_START_TO_CLIP) + // transition from pre to background state
            //                              log_probability_background(*data.read, event_idx, data.strand) + // emit from background
            //                              log(1 - TRANS_CLIP_SELF); // transition to silent pre state
            post_flank[num_events - 2] = log(TRANS_START_TO_CLIP) + // transition from pre to background state
                                         -3.0f + // emit from background
                                         log(1 - TRANS_CLIP_SELF); // transition to silent pre state
        }

        for(int i = num_events - 3; i >= 0; --i) {
            //uint32_t event_idx = e_start + (i + 1) * event_stride;
            // post_flank[i] = log(TRANS_CLIP_SELF) +
            //                 log_probability_background(*data.read, event_idx, data.strand) + // emit from background
            //                 post_flank[i + 1]; // this accounts for the transition from start, and to silent pre
            post_flank[i] = log(TRANS_CLIP_SELF) +
                            -3.0f + // emit from background
                            post_flank[i + 1]; // this accounts for the transition from start, and to silent pre
        }
    }
    return post_flank;
}


// Allocate a vector with the model probabilities of skipping the first i events
inline std::vector<float> make_pre_flanking(const uint32_t e_start,
                                            const uint32_t num_events,
                                            int8_t event_stride)
{
    std::vector<float> pre_flank(num_events + 1, 0.0f);

    // base cases

    // no skipping
    pre_flank[0] = log(1 - TRANS_START_TO_CLIP);

    // skipping the first event
    // this includes the transition probability into and out of the skip state
    // pre_flank[1] = log(TRANS_START_TO_CLIP) + // transition from start to the background state
    //                log_probability_background(*data.read, e_start, data.strand) + // emit from background
    //                log(1 - TRANS_CLIP_SELF); // transition to silent pre state
    pre_flank[1] = log(TRANS_START_TO_CLIP) + // transition from start to the background state
                   -3.0f + // emit from background
                   log(1 - TRANS_CLIP_SELF); // transition to silent pre state

    // skip the remaining events
    for(size_t i = 2; i < pre_flank.size(); ++i) {
        //uint32_t event_idx = e_start + (i - 1) * event_stride;
        // pre_flank[i] = log(TRANS_CLIP_SELF) +
        //                log_probability_background(*data.read, event_idx, data.strand) + // emit from background
        //                pre_flank[i - 1]; // this accounts for the transition from the start & to the silent pre
        pre_flank[i] = log(TRANS_CLIP_SELF) +
                       -3.0f + // emit from background
                       pre_flank[i - 1]; // this accounts for the transition from the start & to the silent pre

    }

    return pre_flank;
}

// Pre-computed transitions from the previous block
// into the current block of states. Log-scaled.
struct BlockTransitions
{
    // Transition from m state (match event to k-mer)
    float lp_mm_self;
    float lp_mb;
    float lp_mk;
    float lp_mm_next;

    // Transitions from b state (bad event that should be ignored)
    float lp_bb;
    float lp_bk;
    float lp_bm_next; // movement to next k-mer
    float lp_bm_self; // movement to k-mer that we came from

    // Transitions from k state (no observation from k-mer)
    float lp_kk;
    float lp_km;
};

inline std::vector<BlockTransitions> calculate_transitions(uint32_t num_kmers, const char *m_seq,
                                                                                const char *m_rc_seq,
                                                                                event_t* event,
                                                                                scalings_t scaling,
                                                                                model_t* cpgmodel,
                                                                                uint32_t event_start_idx,
                                                                                uint32_t event_stop_idx,
                                                                                uint8_t strand,
                                                                                int8_t event_stride,
                                                                                uint8_t rc,
                                                                                double events_per_base)
{
    std::vector<BlockTransitions> transitions(num_kmers);

    //double read_events_per_base = data.read->events_per_base[data.strand];
    double read_events_per_base = events_per_base;
    for(uint32_t ki = 0; ki < num_kmers; ++ki) {
        // if(ki == 0) fprintf(stderr, "double read_events_per_base = %f\n",read_events_per_base );
        // probability of skipping k_i from k_(i - 1)
        //float p_stay = 0.4;
        float p_stay = 1 - (1 / read_events_per_base);
        // if(ki == 0) fprintf(stderr, "float p_stay = %f\n",p_stay );
#ifndef USE_EXTERNAL_PARAMS
        // fprintf(stderr, "ifndef USE_EXTERNAL_PARAMS float p_stay = %f\n",p_stay);
        float p_skip = 0.0025;
        float p_bad = 0.001;
        float p_bad_self = p_bad;
        float p_skip_self = 0.3;
#else
        extern float g_p_skip, g_p_skip_self, g_p_bad, g_p_bad_self;
        float p_skip = g_p_skip;
        float p_skip_self = g_p_skip_self;
        float p_bad = g_p_bad;
        float p_bad_self = g_p_bad_self;
#endif
        // transitions from match state in previous block
        float p_mk = p_skip; // probability of not observing an event at all
        float p_mb = p_bad; // probabilty of observing a bad event
        float p_mm_self = p_stay; // probability of observing additional events from this k-mer
        float p_mm_next = 1.0f - p_mm_self - p_mk - p_mb; // normal movement from state to state

        // transitions from event split state in previous block
        float p_bb = p_bad_self;
        float p_bk, p_bm_next, p_bm_self;
        p_bk = p_bm_next = p_bm_self = (1.0f - p_bb) / 3;

        // transitions from kmer skip state in previous block
        float p_kk = p_skip_self;
        float p_km = 1.0f - p_kk;
        // p_kb not needed, equivalent to B->K

        // log-transform and store
        BlockTransitions& bt = transitions[ki];

        bt.lp_mm_self = log(p_mm_self);
        bt.lp_mb = log(p_mb);
        bt.lp_mk = log(p_mk);
        bt.lp_mm_next = log(p_mm_next);

        bt.lp_bb = log(p_bb);
        bt.lp_bk = log(p_bk);
        bt.lp_bm_next = log(p_bm_next);
        bt.lp_bm_self = log(p_bm_self);

        bt.lp_kk = log(p_kk);
        bt.lp_km = log(p_km);
    }

    return transitions;
}


//todo : can make more efficient using bit encoding
static inline uint32_t get_rank(char base) {
    if (base == 'A') { //todo: do we neeed simple alpha?
        return 0;
    } else if (base == 'C') {
        return 1;
    } else if (base == 'G') {
        return 2;
    } else if (base == 'T') {
        return 3;
    } else if (base == 'N') {
        return 0;        
    } else {
        WARNING("A None ACGTN base found : %c", base);
        return 0;
    }
}

// // return the lexicographic rank of the kmer amongst all strings of
// // length k for this alphabet
// static inline uint32_t get_kmer_rank(const char* str, uint32_t k) {
//     uint32_t p = 1;
//     uint32_t r = 0;

//     // from last base to first
//     for (uint32_t i = 0; i < k; ++i) {
//         //r += rank(str[k - i - 1]) * p;
//         //p *= size();
//         r += get_rank(str[k - i - 1]) * p;
//         p *= 5;
//     }
//     return r;
// }


// return the lexicographic rank of the kmer amongst all strings of
// length k for this alphabet
static inline uint32_t get_kmer_rank(const char* str, uint32_t k) {
    //uint32_t p = 1;
    uint32_t r = 0;

    // from last base to first
    for (uint32_t i = 0; i < k; ++i) {
        //r += rank(str[k - i - 1]) * p;
        //p *= size();
        r += get_rank(str[k - i - 1]) << (i << 1);
    }
    return r;
}

static inline float log_normal_pdf(float x, float gp_mean, float gp_stdv,
                                   float gp_log_stdv) {
    /*INCOMPLETE*/
    float log_inv_sqrt_2pi = -0.918938f; // Natural logarithm
    float a = (x - gp_mean) / gp_stdv;
    return log_inv_sqrt_2pi - gp_log_stdv + (-0.5f * a * a);
    // return 1;
}

static inline float log_probability_match_r9(scalings_t scaling,
                                             model_t* models,
                                             event_t* event, int event_idx,
                                             uint32_t kmer_rank, uint8_t strand,
                                             float sample_rate) {
    // event level mean, scaled with the drift value
    strand = 0;
    assert(kmer_rank < 15625);
    //float level = read.get_drift_scaled_level(event_idx, strand);

    //float time =
    //    (events.event[event_idx].start - events.event[0].start) / sample_rate;
    float unscaledLevel = event[event_idx].mean;
    float scaledLevel = unscaledLevel;
    //float scaledLevel = unscaledLevel - time * scaling.shift;

    //fprintf(stderr, "level %f\n",scaledLevel);
    //GaussianParameters gp = read.get_scaled_gaussian_from_pore_model_state(pore_model, strand, kmer_rank);
    float gp_mean =
        scaling.scale * models[kmer_rank].level_mean + scaling.shift;
    float gp_stdv = models[kmer_rank].level_stdv * scaling.var;
    // float gp_stdv = 0;
    // float gp_log_stdv = models[kmer_rank].level_log_stdv + scaling.log_var;
    // if(models[kmer_rank].level_stdv <0.01 ){
    //  fprintf(stderr,"very small std dev %f\n",models[kmer_rank].level_stdv);
    // }
#ifdef CACHED_LOG
    float gp_log_stdv =
        models[kmer_rank].level_log_stdv + scaling.log_var;
#else
    float gp_log_stdv =
        log(models[kmer_rank].level_stdv) + log(scaling.var);
#endif

    float lp = log_normal_pdf(scaledLevel, gp_mean, gp_stdv, gp_log_stdv);
    return lp;
}


// This function fills in a matrix with the result of running the HMM.
// The templated ProfileHMMOutput class allows one to run either Viterbi
// or the Forward algorithm.
template<class ProfileHMMOutput>
inline float profile_hmm_fill_generic_r9(const char *m_seq,
                                         const char *m_rc_seq,
                                        event_t* event,
                                        scalings_t scaling,
                                        model_t* cpgmodel,
                                        uint32_t event_start_idx,
                                        uint32_t event_stop_idx,
                                        uint8_t strand,
                                        int8_t event_stride,
                                        uint8_t rc,
                                        const uint32_t e_start,
                                        double events_per_base,
                                        uint32_t hmm_flags,
                                        ProfileHMMOutput& output){
    // PROFILE_FUNC("profile_hmm_fill_generic")
    // HMMInputSequence sequence = _sequence;
    // HMMInputData data = _data;

    assert( (rc && event_stride == -1) || (!rc && event_stride == 1));

#if HMM_REVERSE_FIX
    fprintf(stderr, "HMM_REVERSE_FIX\n");
    if(event_stride == -1) {
        // sequence.swap();
        char *temp = m_seq;
        m_seq = m_rc_seq;
        m_rc_seq = temp;
        uint32_t tmp = event_stop_idx;
        event_stop_idx = event_start_idx;
        event_start_idx = tmp;
        event_stride = 1;
        rc = false;
    }
#endif

    //e_start = event_start_idx;

    // Calculate number of blocks
    // A block of the HMM is a set of states for one kmer
    uint32_t num_blocks = output.get_num_columns() / PSR9_NUM_STATES;
    //fprintf(stderr,"%d %d\n",output.get_num_columns(),PSR9_NUM_STATES);
    uint32_t last_event_row_idx = output.get_num_rows() - 1;

    // Precompute the transition probabilites for each kmer block
    uint32_t num_kmers = num_blocks - 2; // two terminal blocks
    uint32_t last_kmer_idx = num_kmers - 1;


    std::vector<BlockTransitions> transitions = calculate_transitions(num_kmers,
                                                                        m_seq,
                                                                        m_rc_seq,
                                                                        event,
                                                                        scaling,
                                                                        cpgmodel,
                                                                        event_start_idx,
                                                                        event_stop_idx,
                                                                        strand,
                                                                        event_stride,
                                                                        rc,
                                                                        events_per_base);

    // fprintf(stderr, "std::vector<BlockTransitions> transitions len = %d\n",transitions.size());

    // std::ofstream wBlockTransitions("wBlockTransitions");
    // for(size_t i = 0; i < transitions.size(); i++){
    //     BlockTransitions b = transitions[i];
    //     wBlockTransitions << b.lp_mm_self << " " << b.lp_mb << " " << b.lp_mk << " " << b.lp_mm_next << "\n";
    //     wBlockTransitions << b.lp_bm_self << " " << b.lp_bb << " " << b.lp_bk << " " << b.lp_bm_next << "\n";
    //     wBlockTransitions << b.lp_km << " " << b.lp_kk << "\n";
    // }
    // Precompute kmer ranks
    // const uint32_t k = data.pore_model->k;
    //const uint32_t k = KMER_SIZE;
    // Make sure the HMMInputSequence's alphabet matches the state space of the read


    ////change start

    // assert( data.pore_model->states.size() == sequence.get_num_kmer_ranks(k) );

    std::vector<uint32_t> kmer_ranks(num_kmers);

    //todo : pre-calculate
    int32_t seq_len = strlen(m_seq);

    // std::ofstream wkmer_ranks("wkmer_ranks");
    //check : this might be reverse cmplement kmer rnak
    for(size_t ki = 0; ki < num_kmers; ++ki){
        const char* substring = 0;
        if(rc==0){
            substring=m_seq+ki;
        }
        else{
            substring=m_rc_seq+seq_len-ki-KMER_SIZE;
        }

        // kmer_ranks[ki] = sequence.get_kmer_rank(ki, k, data.rc);
        kmer_ranks[ki] = get_kmer_rank(substring,KMER_SIZE);
        // wkmer_ranks << kmer_ranks[ki] << " ";
    }


    ///change over

    size_t num_events = output.get_num_rows() - 1;

    std::vector<float> pre_flank = make_pre_flanking(e_start, num_events,event_stride);
    std::vector<float> post_flank = make_post_flanking(e_start, num_events,event_stride,event_stop_idx);


    // std::ofstream wpre_flank("wpre_flank");
    // for(size_t i = 0; i < pre_flank.size(); i++){
    //     wpre_flank << pre_flank[i] << " ";
    // }
    // std::ofstream wpost_flank("wpost_flank");
    // for(size_t i = 0; i < post_flank.size(); i++){
    //     wpost_flank << post_flank[i] << " ";
    // }

    // The model is currently constrainted to always transition
    // from the terminal/clipped state to the first kmer (and from the
    // last kmer to the terminal/clipping state so these are log(1.0).
    // They are kept as variables as it might be relaxed later.
    float lp_sm, lp_ms;
    lp_sm = lp_ms = 0.0f;

    // the penalty is controlled by the transition probability
    float BAD_EVENT_PENALTY = 0.0f;
    // std::ofstream wlp_emission_m("wlp_emission_m");
    // fprintf(stderr, "num_blocks = %d\n",num_blocks);
    // Fill in matrix
    size_t tester_inside = 0;

    for(uint32_t row = 1; row < output.get_num_rows(); row++) {
    // for(uint32_t row = 1; row < 2; row++) {
        // Skip the first block which is the start state, it was initialized above
        // Similarily skip the last block, which is calculated in the terminate() function
        for(uint32_t block = 1; block < num_blocks - 1; block++) {
// for(uint32_t block = 1; block < 3; block++) {
            // retrieve transitions
            uint32_t kmer_idx = block - 1;
            BlockTransitions& bt = transitions[kmer_idx];

            uint32_t prev_block = block - 1;
            uint32_t prev_block_offset = PSR9_NUM_STATES * prev_block;
            uint32_t curr_block_offset = PSR9_NUM_STATES * block;
            // fprintf(stderr, "prev_block = %d prev_block_offset= %d curr_block_offset= %d \n",prev_block,prev_block_offset,curr_block_offset);
            // Emission probabilities
            uint32_t event_idx = e_start + (row - 1) * event_stride;
            uint32_t rank = kmer_ranks[kmer_idx];
            // fprintf(stderr, "event_idx = %d rank= %d \n",event_idx,rank);
            // float lp_emission_m = log_probability_match_r9(*data.read, *data.pore_model, rank, event_idx, data.strand);
            float lp_emission_m =
                log_probability_match_r9(scaling, cpgmodel, event, event_idx,rank, strand, 0);
            // wlp_emission_m << lp_emission_m << " ";
            // fprintf(stderr, "float lp_emission_m = %f\n",lp_emission_m );

            //fprintf(stderr,"m_seq %s, event_idx %d, kmer_rank %d, log prob : %f\n",m_seq,event_idx,rank,lp_emission_m);
            //fprintf(stderr,"e_start %d, row %d, event_stride %d, block %d, num_block %d\n",e_start,row,event_stride,block,num_blocks);
            float lp_emission_b = BAD_EVENT_PENALTY;
            HMMUpdateScores scores;

            // state PSR9_MATCH
            scores.x[HMT_FROM_SAME_M] = bt.lp_mm_self + output.get(row - 1, curr_block_offset + PSR9_MATCH);
            scores.x[HMT_FROM_PREV_M] = bt.lp_mm_next + output.get(row - 1, prev_block_offset + PSR9_MATCH);
            scores.x[HMT_FROM_SAME_B] = bt.lp_bm_self + output.get(row - 1, curr_block_offset + PSR9_BAD_EVENT);
            scores.x[HMT_FROM_PREV_B] = bt.lp_bm_next + output.get(row - 1, prev_block_offset + PSR9_BAD_EVENT);
            scores.x[HMT_FROM_PREV_K] = bt.lp_km + output.get(row - 1, prev_block_offset + PSR9_KMER_SKIP);

            // m_s is the probability of going from the start state
            // to this kmer. The start state is (currently) only
            // allowed to go to the first kmer. If ALLOW_PRE_CLIP
            // is defined, we allow all events before this one to be skipped,
            // with a penalty;
            scores.x[HMT_FROM_SOFT] = (kmer_idx == 0 &&
                                        (event_idx == e_start ||
                                             (hmm_flags & HAF_ALLOW_PRE_CLIP))) ? lp_sm + pre_flank[row - 1] : -INFINITY;
            // fprintf(stderr, "%f %f %f %f %f %f %f \n",scores.x[0],scores.x[1],scores.x[2],scores.x[3],scores.x[4],scores.x[5]);                           

            output.update_cell(row, curr_block_offset + PSR9_MATCH, scores, lp_emission_m);
            // fprintf(stderr, "cell value = %f\n",output.get(row, curr_block_offset + PSR9_MATCH));
    
            // state PSR9_BAD_EVENT
            scores.x[HMT_FROM_SAME_M] = bt.lp_mb + output.get(row - 1, curr_block_offset + PSR9_MATCH);
            scores.x[HMT_FROM_PREV_M] = -INFINITY; // not allowed
            scores.x[HMT_FROM_SAME_B] = bt.lp_bb + output.get(row - 1, curr_block_offset + PSR9_BAD_EVENT);
            scores.x[HMT_FROM_PREV_B] = -INFINITY;
            scores.x[HMT_FROM_PREV_K] = -INFINITY;
            scores.x[HMT_FROM_SOFT] = -INFINITY;
            // fprintf(stderr, "%f %f %f %f %f %f %f \n",scores.x[0],scores.x[1],scores.x[2],scores.x[3],scores.x[4],scores.x[5]);                           
            output.update_cell(row, curr_block_offset + PSR9_BAD_EVENT, scores, lp_emission_b);
            // fprintf(stderr, "cell value = %f\n",output.get(row, curr_block_offset + PSR9_BAD_EVENT));


            // state PSR9_KMER_SKIP
            scores.x[HMT_FROM_SAME_M] = -INFINITY;
            scores.x[HMT_FROM_PREV_M] = bt.lp_mk + output.get(row, prev_block_offset + PSR9_MATCH);
            scores.x[HMT_FROM_SAME_B] = -INFINITY;
            scores.x[HMT_FROM_PREV_B] = bt.lp_bk + output.get(row, prev_block_offset + PSR9_BAD_EVENT);
            scores.x[HMT_FROM_PREV_K] = bt.lp_kk + output.get(row, prev_block_offset + PSR9_KMER_SKIP);
            scores.x[HMT_FROM_SOFT] = -INFINITY;
            output.update_cell(row, curr_block_offset + PSR9_KMER_SKIP, scores, 0.0f); // no emission

            // If POST_CLIP is enabled we allow the last kmer to transition directly
            // to the end after any event. Otherwise we only allow it from the
            // last kmer/event match.
                tester_inside++;
            if(kmer_idx == last_kmer_idx && ( (hmm_flags & HAF_ALLOW_POST_CLIP) || row == last_event_row_idx)) {
                // fprintf(stderr, "inside\n");    
                float lp1 = lp_ms + output.get(row, curr_block_offset + PSR9_MATCH) + post_flank[row - 1];
                float lp2 = lp_ms + output.get(row, curr_block_offset + PSR9_BAD_EVENT) + post_flank[row - 1];
                float lp3 = lp_ms + output.get(row, curr_block_offset + PSR9_KMER_SKIP) + post_flank[row - 1];

                output.update_end(lp1, row, curr_block_offset + PSR9_MATCH);
                output.update_end(lp2, row, curr_block_offset + PSR9_BAD_EVENT);
                output.update_end(lp3, row, curr_block_offset + PSR9_KMER_SKIP);
            }

#ifdef DEBUG_LOCAL_ALIGNMENT
            fprintf(stderr, "%s\n","DEBUG_LOCAL_ALIGNMENT" );
            printf("[%d %d] start: %.2lf  pre: %.2lf fm: %.2lf\n", event_idx, kmer_idx, m_s + lp_emission_m, pre_flank[row - 1], output.get(row, curr_block_offset + PSR9_MATCH));
            printf("[%d %d]   end: %.2lf post: %.2lf\n", event_idx, kmer_idx, lp_end, post_flank[row - 1]);
#endif

#ifdef DEBUG_FILL
            fprintf(stderr, "%s\n","DEBUG_FILL" );
            printf("Row %u block %u\n", row, block);

            printf("\tPSR9_MATCH -- Transitions: [%.3lf %.3lf %.3lf %.3lf %.3lf] Prev: [%.2lf %.2lf %.2lf %.2lf %.2lf] out: %.2lf\n",
                    bt.lp_mm_self, bt.lp_mm_next, bt.lp_bm_self, bt.lp_bm_next, bt.lp_km,
                    output.get(row - 1, prev_block_offset + PSR9_MATCH),
                    output.get(row - 1, curr_block_offset + PSR9_MATCH),
                    output.get(row - 1, prev_block_offset + PSR9_BAD_EVENT),
                    output.get(row - 1, curr_block_offset + PSR9_BAD_EVENT),
                    output.get(row - 1, prev_block_offset + PSR9_KMER_SKIP),
                    output.get(row, curr_block_offset + PSR9_MATCH));
            printf("\tPSR9_BAD_EVENT -- Transitions: [%.3lf %.3lf] Prev: [%.2lf %.2lf] out: %.2lf\n",
                    bt.lp_mb, bt.lp_bb,
                    output.get(row - 1, curr_block_offset + PSR9_MATCH),
                    output.get(row - 1, curr_block_offset + PSR9_BAD_EVENT),
                    output.get(row, curr_block_offset + PSR9_BAD_EVENT));

            printf("\tPSR9_KMER_SKIP -- Transitions: [%.3lf %.3lf %.3lf] Prev: [%.2lf %.2lf %.2lf] sum: %.2lf\n",
                    bt.lp_mk, bt.lp_bk, bt.lp_kk,
                    output.get(row, prev_block_offset + PSR9_MATCH),
                    output.get(row, prev_block_offset + PSR9_BAD_EVENT),
                    output.get(row, prev_block_offset + PSR9_KMER_SKIP),
                    output.get(row, curr_block_offset + PSR9_KMER_SKIP));

            printf("\tEMISSION: %.2lf %.2lf\n", lp_emission_m, lp_emission_b);
#endif
        }
    }
    // fprintf(stderr, "output.get_end() = %f\n",output.get_end() );
        // fprintf(stderr, "tester_inside = %d\n",tester_inside );

    // std::ofstream woutput_mat("woutput_mat");
    // for(size_t i = 0 ; i < output.get_num_rows(); i++){
    //     for(size_t j = 0 ; j < output.get_num_columns(); j++){
    //         woutput_mat << output.get(i,j) << " ";
    //     }
    // }    
    return output.get_end();
}


// Output writer for the Viterbi Algorithm
class ProfileHMMViterbiOutputR9
{
    public:
        ProfileHMMViterbiOutputR9(FloatMatrix* pf, UInt8Matrix* pb) : p_fm(pf), p_bm(pb), lp_end(-INFINITY) {}
        
        inline void update_cell(uint32_t row, uint32_t col, const HMMUpdateScores& scores, float lp_emission)
        {
            // probability update
            float max = scores.x[0];
            uint8_t from = 0;
            for(auto i = 1; i < HMT_NUM_MOVEMENT_TYPES; ++i) {
                max = scores.x[i] > max ? scores.x[i] : max;
                from = max == scores.x[i] ? i : from;
            }

            set(*p_fm, row, col, max + lp_emission);
            set(*p_bm, row, col, from);
        }
        
        // add in the probability of ending the alignment at row,col
        inline void update_end(float v, uint32_t row, uint32_t col)
        {
            if(v > lp_end) {
                lp_end = v;
                end_row = row;
                end_col = col;
            }
        }

        // get the log probability stored at a particular row/column
        inline float get(uint32_t row, uint32_t col) const
        {
            return ::get(*p_fm, row, col);
        }

        // get the log probability for the end state
        inline float get_end() const
        {
            return lp_end;
        }
        
        // get the row/col that lead to the end state
        inline void get_end_cell(uint32_t& row, uint32_t& col)
        {
            row = end_row;
            col = end_col;
        }

        inline size_t get_num_columns() const
        {
            return p_fm->n_cols;
        }

        inline size_t get_num_rows() const
        {
            return p_fm->n_rows;
        }
    
    private:
        ProfileHMMViterbiOutputR9(); // not allowed

        FloatMatrix* p_fm;
        UInt8Matrix* p_bm;

        float lp_end;
        uint32_t end_row;
        uint32_t end_col;
};

static inline void profile_hmm_forward_initialize_r9(FloatMatrix& fm)
{
    // initialize forward calculation
    for(uint32_t si = 0; si < fm.n_cols; si++) {
        set(fm, 0, si, -INFINITY);
    }

    for(uint32_t ri = 0; ri < fm.n_rows; ri++) {
        set(fm, ri, PSR9_KMER_SKIP, -INFINITY);
        set(fm, ri, PSR9_BAD_EVENT, -INFINITY);
        set(fm, ri, PSR9_MATCH, -INFINITY);
    }
}

// Convert an enumerated state into a symbol
static inline char ps2char(ProfileStateR9 ps) { return "KBMNS"[ps]; }


static inline std::vector<HMMAlignmentState> profile_hmm_align(
    std::string fwd_subseq,
    std::string rc_subseq,
    event_t* event,
    scalings_t scaling,
    model_t* cpgmodel,
    double events_per_base,
    uint8_t strand,
    uint8_t rc,
    uint32_t k, //KMER_SIZE
    uint32_t e_start, //curr_start_event;
    uint32_t e_end, //input_event_stop_idx
    int8_t event_stride) // event_stride )
{
    std::vector<HMMAlignmentState> alignment;
    uint32_t n_kmers = fwd_subseq.length() - k + 1;
    uint32_t n_states = PSR9_NUM_STATES * (n_kmers + 2); // + 2 for explicit terminal states
    uint32_t n_events = 0;
    if(e_end > e_start)
        n_events = e_end - e_start + 1;
    else
        n_events = e_start - e_end + 1;
    assert(n_events >= 2);
    uint32_t n_rows = n_events + 1;
    // Allocate a matrix to hold the HMM result
    // fprintf(stderr, "n_states = %d\n",n_states);
    FloatMatrix vm;
    allocate_matrix(vm, n_rows, n_states);
    UInt8Matrix bm;
    allocate_matrix(bm, n_rows, n_states);
    // fprintf(stderr, "n_kmers = %d n_states = %d\n",n_kmers,n_states);
    // std::ofstream wfile_bm("wbfile_bm");
    // for(size_t i = 0; i < n_rows; i++){
    //     for(size_t j = 0; j < n_states; j++){
    //         wfile_bm << get(bm, i, j);
    //     }
    // }
    ProfileHMMViterbiOutputR9 output(&vm, &bm);
    //printing output
    // std::ofstream woutput_mat("woutput_mat");
    // for(size_t i = 0 ; i < n_rows; i++){
    //     for(size_t j = 0 ; j < n_states; j++){
    //         woutput_mat << output.get(i,j) << " ";
    //     }
    // }

    profile_hmm_forward_initialize_r9(vm);//profile_hmm_viterbi_initialize_r9(vm);
    uint32_t hmm_flags = 0; // as in nanopolish

    // std::ofstream wfile_vm("wvfile_vm");
    // for(size_t i = 0; i < n_rows; i++){
    //     for(size_t j = 0; j < n_states; j++){
    //         wfile_vm << get(vm, i, j);
    //     }
    // }


    // std::ofstream woutput_mat_2("woutput_mat_2");
    // for(size_t i = 0 ; i < n_rows; i++){
    //     for(size_t j = 0 ; j < n_states; j++){
    //         woutput_mat_2 << output.get(i,j) << " ";
    //     }
    // }

    profile_hmm_fill_generic_r9(fwd_subseq.c_str(),
                                rc_subseq.c_str(),
                                event,
                                scaling,
                                cpgmodel,
                                e_start,
                                e_end,
                                strand,     
                                event_stride,
                                rc,
                                e_start,
                                events_per_base,
                                hmm_flags,
                                output);
    // std::ofstream woutput_mat_3("woutput_mat_3");
    // for(size_t i = 0 ; i < n_rows; i++){
    //     for(size_t j = 0 ; j < n_states; j++){
    //         woutput_mat_3 << output.get(i,j) << " ";
    //     }
    // }
   


   //? profile_hmm_fill_generic_r9(sequence, data, e_start, flags, output);
     // Traverse the backtrack matrix to compute the results
    int traversal_stride = event_stride;


    // checking if the vars are the same

#if HMM_REVERSE_FIX
    // Hack to support the fixed HMM
    // TODO: clean up
    fprintf(stderr, "HMM_REVERSE_FIX\n");
    traversal_stride = 1;
    if(event_stride == -1) {
        e_start = event_stop_idx;
    }
#endif

    // start from the last event matched to the last kmer
    uint32_t row = n_rows - 1;
    uint32_t col = PSR9_NUM_STATES * n_kmers + PSR9_MATCH;
    // fprintf(stderr, "row = %d\n",row);
    size_t tester_j = 0;
    // std::ofstream wrow("wrow");

    while(row > 0) {
        tester_j++;
        uint32_t event_idx = e_start + (row - 1) * traversal_stride;
        uint32_t block = col / PSR9_NUM_STATES;
        uint32_t kmer_idx = block - 1;
        ProfileStateR9 curr_ps = (ProfileStateR9) (col % PSR9_NUM_STATES);

#if DEBUG_BACKTRACK
        fprintf(stderr, "DEBUG_BACKTRACK\n");
        printf("backtrace %zu %zu coord: (%zu, %zu, %zu) state: %d\n", event_idx, kmer_idx, row, col, block, curr_ps);
#endif

        assert(block > 0);
        assert(get(vm, row, col) != -INFINITY);

        HMMAlignmentState as;
        as.event_idx = event_idx;
        as.kmer_idx = kmer_idx;
        as.l_posterior = -INFINITY; // not computed
        as.l_fm = get(vm, row, col);
        as.log_transition_probability = -INFINITY; // not computed
        as.state = ps2char(curr_ps);
        alignment.push_back(as);

        // Update the event (row) and k-mer using the backtrack matrix
        HMMMovementType movement = (HMMMovementType)get(bm, row, col);
        if(movement == HMT_FROM_SOFT) {
            // fprintf(stderr, "HMT_FROM_SOFT , row = %d \n", row);
            break;
        }
        
        // update kmer_idx and state
        ProfileStateR9 next_ps;
        switch(movement) {
            case HMT_FROM_SAME_M: 
                next_ps = PSR9_MATCH;
                break;
            case HMT_FROM_PREV_M: 
                kmer_idx -= 1;
                next_ps = PSR9_MATCH;
                break;
            case HMT_FROM_SAME_B:
                next_ps = PSR9_BAD_EVENT;
                break;
            case HMT_FROM_PREV_B:
                kmer_idx -= 1;
                next_ps = PSR9_BAD_EVENT;
                break;
            case HMT_FROM_PREV_K:
                kmer_idx -= 1;
                next_ps = PSR9_KMER_SKIP;
                break;
            case HMT_FROM_SOFT:
                assert(false);
                break;
            case HMT_NUM_MOVEMENT_TYPES:
                assert(0);
                break;    
            default:
                assert(0);
                break;    
        }

        // update row (event) idx only if this isn't a kmer skip, which is silent
        if(curr_ps != PSR9_KMER_SKIP) {
            row -= 1;
            // fprintf(stderr, "inside row = %d\n",row);
        }
            // wrow << row << " ";

        col = PSR9_NUM_STATES * (kmer_idx + 1) + next_ps;
    }
    // fprintf(stderr, "row = %d\n",row);
    // fprintf(stderr, "tester_j = %d\n",tester_j);    

#if HMM_REVERSE_FIX
// change the strand of the kmer indices if we aligned to the reverse strand
if(event_stride == -1) {
    for(size_t ai = 0; ai < alignment.size(); ++ai) {
        size_t k_idx = alignment[ai].kmer_idx;
        alignment[ai].kmer_idx = fwd_subseq.length() - k_idx - k;
    }
} else {
    std::reverse(alignment.begin(), alignment.end());
}
#else
    //
    std::reverse(alignment.begin(), alignment.end());
#endif

 //
    free_matrix(vm);
    free_matrix(bm);

    return alignment;

}


//from nanopolish eventalign.c
// Returns the index into the aligned_pairs vector that has the highest ref_pos
// that is not greater than ref_pos_max. It starts the search at pair_idx
static int get_end_pair(const std::vector<AlignedPair>& aligned_pairs, int ref_pos_max, int pair_idx)
{
    while(pair_idx < (int)aligned_pairs.size()) {
        if(aligned_pairs[pair_idx].ref_pos > ref_pos_max)
            return pair_idx - 1;
        pair_idx += 1;
    }

    return aligned_pairs.size() - 1;
}
//from nanopolish eventalign.c
// Modify the aligned_pairs vector to ensure there are no alignments
// outside of the given reference coordinates
static void trim_aligned_pairs_to_ref_region(std::vector<AlignedPair>& aligned_pairs, int ref_start, int ref_end)
{
    std::vector<AlignedPair> trimmed;
    for(size_t i = 0; i < aligned_pairs.size(); ++i) {
        if(aligned_pairs[i].ref_pos >= ref_start && 
           aligned_pairs[i].ref_pos <= ref_end) {
            trimmed.push_back(aligned_pairs[i]);
        }
    }
    
    aligned_pairs.swap(trimmed);
}
//from nanopolish eventalign.c
// Modify the aligned_pairs vector to ensure the highest read position
// does not exceed max_kmer
static void trim_aligned_pairs_to_kmer(std::vector<AlignedPair>& aligned_pairs, int max_kmer_idx)
{
    int idx = aligned_pairs.size() - 1;
    while(idx >= 0 && aligned_pairs[idx].read_pos > max_kmer_idx)
        idx -= 1;

    if(idx < 0)
        aligned_pairs.clear(); // no valid data
    else
        aligned_pairs.resize(idx + 1);
}


//from https://github.com/hasindu2008/f5c/blob/64d9d51cd773e46e1a72dbf2e2bf333be24df7c4/meth.c
//helper for get_closest_event_to
static inline int get_next_event(int start, int stop, int stride,index_pair_t* base_to_event_map) 
{
    while(start != stop) {

        int ei = base_to_event_map[start].start;
        if(ei != -1)
            return ei;
        start += stride;
    }
    return -1;
}

//from https://github.com/hasindu2008/f5c/blob/64d9d51cd773e46e1a72dbf2e2bf333be24df7c4/meth.c
static inline int get_closest_event_to(int k_idx, index_pair_t* base_to_event_map, int base_to_event_map_size)
{
    int stop_before = std::max(0, k_idx - 1000);
    int stop_after = std::min(k_idx + 1000, base_to_event_map_size - 1);

    int event_before = get_next_event(k_idx, stop_before, -1, base_to_event_map);
    int event_after = get_next_event(k_idx, stop_after, 1, base_to_event_map);

    // TODO: better selection of "best" event to return
    if(event_before == -1)
        return event_after;
    return event_before;
}

//from https://github.com/hasindu2008/f5c/blob/64d9d51cd773e46e1a72dbf2e2bf333be24df7c4/meth.c
//Returns true if c is a valid ambiguity code
static inline bool isAmbiguous(char c) {
    switch (c) {
        case 'M':
        case 'R':
        case 'W':
        case 'S':
        case 'Y':
        case 'K':
        case 'V':
        case 'H':
        case 'D':
        case 'B':
        case 'N':
            return true;
        default:
            return false;
    }
}
//from https://github.com/hasindu2008/f5c/blob/64d9d51cd773e46e1a72dbf2e2bf333be24df7c4/meth.c
//IUPAC alphabet
static inline bool isUnambiguous(char c) {
    switch (c) {
        case 'A':
        case 'C':
        case 'G':
        case 'T':
            return true;
        default:
            return false;
    }
}

//from https://github.com/hasindu2008/f5c/blob/64d9d51cd773e46e1a72dbf2e2bf333be24df7c4/meth.c
//Returns true if c is a valid symbol in this alphabet
static inline bool isValid(char c) { return isUnambiguous(c) || isAmbiguous(c); }
static const char* complement_dna = "TGCA";
static const uint8_t rank_dna[256] = {
    0, 0, 0, 0, 0, 0, 0, 0, 0, 0, 0, 0, 0, 0, 0, 0, 0, 0, 0, 0, 0, 0, 0, 0,
    0, 0, 0, 0, 0, 0, 0, 0, 0, 0, 0, 0, 0, 0, 0, 0, 0, 0, 0, 0, 0, 0, 0, 0,
    0, 0, 0, 0, 0, 0, 0, 0, 0, 0, 0, 0, 0, 0, 0, 0, 0, 0, 0, 1, 0, 0, 0, 2,
    0, 0, 0, 0, 0, 0, 0, 0, 0, 0, 0, 0, 3, 0, 0, 0, 0, 0, 0, 0, 0, 0, 0, 0,
    0, 0, 0, 0, 0, 0, 0, 0, 0, 0, 0, 0, 0, 0, 0, 0, 0, 0, 0, 0, 0, 0, 0, 0,
    0, 0, 0, 0, 0, 0, 0, 0, 0, 0, 0, 0, 0, 0, 0, 0, 0, 0, 0, 0, 0, 0, 0, 0,
    0, 0, 0, 0, 0, 0, 0, 0, 0, 0, 0, 0, 0, 0, 0, 0, 0, 0, 0, 0, 0, 0, 0, 0,
    0, 0, 0, 0, 0, 0, 0, 0, 0, 0, 0, 0, 0, 0, 0, 0, 0, 0, 0, 0, 0, 0, 0, 0,
    0, 0, 0, 0, 0, 0, 0, 0, 0, 0, 0, 0, 0, 0, 0, 0, 0, 0, 0, 0, 0, 0, 0, 0,
    0, 0, 0, 0, 0, 0, 0, 0, 0, 0, 0, 0, 0, 0, 0, 0, 0, 0, 0, 0, 0, 0, 0, 0,
    0, 0, 0, 0, 0, 0, 0, 0, 0, 0, 0, 0, 0, 0, 0, 0};

//from https://github.com/hasindu2008/f5c/blob/64d9d51cd773e46e1a72dbf2e2bf333be24df7c4/meth.c
//reverse-complement a DNA string
static inline std::string reverse_complement(const std::string& str) {
    std::string out(str.length(), 'A');
    size_t i = 0;             // input
    int j = str.length() - 1; // output
    while (i < str.length()) {
        // complement a single base
        assert(str[i] != METHYLATED_SYMBOL);
        out[j--] = complement_dna[rank_dna[(int)str[i++]]];
    }
    return out;
}
//from https://github.com/hasindu2008/f5c/blob/64d9d51cd773e46e1a72dbf2e2bf333be24df7c4/meth.c
static inline std::string getPossibleSymbols(char c) {
    switch (c) {
        case 'A':
            return "A";
        case 'C':
            return "C";
        case 'G':
            return "G";
        case 'T':
            return "T";
        case 'M':
            return "AC";
        case 'R':
            return "AG";
        case 'W':
            return "AT";
        case 'S':
            return "CG";
        case 'Y':
            return "CT";
        case 'K':
            return "GT";
        case 'V':
            return "ACG";
        case 'H':
            return "ACT";
        case 'D':
            return "AGT";
        case 'B':
            return "CGT";
        case 'N':
            return "ACGT";
        default:
            return "";
    }
}
//from https://github.com/hasindu2008/f5c/blob/64d9d51cd773e46e1a72dbf2e2bf333be24df7c4/meth.c
static inline std::string disambiguate(const std::string& str) {
    // create output and convert lower case to upper case
    std::string out(str);
    std::transform(out.begin(), out.end(), out.begin(), ::toupper);

    size_t i = 0;
    while (i < out.length()) {
        size_t stride = 1;
        //bool is_recognition_site = false;

        assert(isValid(out[i]));
        out[i] = getPossibleSymbols(out[i])[0];
        stride = 1;


        i += stride;
    }
    return out;
}

// from https://github.com/hasindu2008/f5c/blob/64d9d51cd773e46e1a72dbf2e2bf333be24df7c4/meth.c
static std::vector<AlignedSegment> get_aligned_segments_two_params(const bam1_t* record, int read_stride)
{
    std::vector<AlignedSegment> out;
    // Initialize first segment
    out.push_back(AlignedSegment());

    // This code is derived from bam_fillmd1_core
    //uint8_t *ref = NULL;
    //uint8_t *seq = bam_get_seq(record);
    uint32_t *cigar = bam_get_cigar(record);
    const bam1_core_t *c = &record->core;

    // read pos is an index into the original sequence that is present in the FASTQ
    // on the strand matching the reference
    int read_pos = 0;

    // query pos is an index in the query string that is recorded in the bam
    // we record this as a sanity check
    //int query_pos = 0;
    
    int ref_pos = c->pos;

    for (uint32_t ci = 0; ci < c->n_cigar; ++ci) {
        
        int cigar_len = cigar[ci] >> 4;
        int cigar_op = cigar[ci] & 0xf;

        // Set the amount that the ref/read positions should be incremented
        // based on the cigar operation
        int read_inc = 0;
        int ref_inc = 0;
 
        // Process match between the read and the reference
        bool is_aligned = false;
        if(cigar_op == BAM_CMATCH || cigar_op == BAM_CEQUAL || cigar_op == BAM_CDIFF) {
            is_aligned = true;
            read_inc = read_stride;
            ref_inc = 1;
        } else if(cigar_op == BAM_CDEL) {
            ref_inc = 1;   
        } else if(cigar_op == BAM_CREF_SKIP) {
            // end the current segment and start a new one
            out.push_back(AlignedSegment());
            ref_inc = 1;
        } else if(cigar_op == BAM_CINS) {
            read_inc = read_stride;
        } else if(cigar_op == BAM_CSOFT_CLIP) {
            read_inc = 1; // special case, do not use read_stride
        } else if(cigar_op == BAM_CHARD_CLIP) {
            read_inc = 0;
        } else {
            printf("Cigar: %d\n", cigar_op);
            assert(false && "Unhandled cigar operation");
        }

        // Iterate over the pairs of aligned bases
        for(int j = 0; j < cigar_len; ++j) {
            if(is_aligned) {
                out.back().push_back({ref_pos, read_pos});
            }

            // increment
            read_pos += read_inc;
            ref_pos += ref_inc;
        }
    }
    return out;
}


// from https://github.com/hasindu2008/f5c/blob/64d9d51cd773e46e1a72dbf2e2bf333be24df7c4/meth.c
static inline int32_t flip_k_strand(int32_t read_length,int32_t k_idx, uint32_t k)
{
    return read_length - k_idx - k;
}

// from https://github.com/hasindu2008/f5c/blob/64d9d51cd773e46e1a72dbf2e2bf333be24df7c4/meth.c
struct EventAlignment
{
    // ref data
    // std::string ref_name;
    std::string ref_kmer;
    int ref_position;

    // event data
    size_t read_idx;
    // int strand_idx;
    int event_idx;
    bool rc;

    // hmm data
    std::string model_kmer;
    char hmm_state;
};


//-------------
struct EventAlignmentParameters
{
    EventAlignmentParameters()
    {
        //sr = NULL;
        et = NULL;
        //fai = NULL;
        //hdr = NULL;
        record = NULL;
        model = NULL;
        //strand_idx = NUM_STRANDS;
        
        alphabet = "";
        read_idx = -1;
        region_start = -1;
        region_end = -1;
    }


    // Mandatory
    //SquiggleRead* sr;
    //necessary parameters inside sr
    index_pair_t* base_to_event_map;
    int32_t read_length;
    scalings_t scalings;


    std::string ref_name;

    ///
    const event_table* et;
    //const faidx_t* fai;
    //const bam_hdr_t* hdr;
    const bam1_t* record;
    model_t* model;
    //size_t strand_idx;

    double events_per_base;
    
    // optional
    std::string alphabet;
    int read_idx;
    int region_start;
    int region_end;
};


 std::vector<event_alignment_t> align_read_to_ref(const EventAlignmentParameters& params, char *ref)
{


    // Sanity check input parameters
    //assert(params.sr != NULL);
    assert(params.et != NULL);
    //assert(params.fai != NULL);
    //assert(params.hdr != NULL);
    assert(params.record != NULL);
    assert(params.model != NULL);
    //assert(params.strand_idx < NUM_STRANDS);
    assert( (params.region_start == -1 && params.region_end == -1) || (params.region_start <= params.region_end));
    //const PoreModel* pore_model = params.get_model(); // --hasindu this is model now

    std::vector<event_alignment_t> alignment_output;
    

    // Extract the reference subsequence for the entire alignment
    // int fetched_len = 0;
    int ref_offset = params.record->core.pos;
    // std::string ref_name(params.hdr->target_name[params.record->core.tid]);
    // std::string ref_seq = get_reference_region_ts(params.fai, ref_name.c_str(), ref_offset, 
    //                                               bam_endpos(params.record), &fetched_len);
    // hasindu - a hack to get the reference sequence
    std::string ref_seq = ref;
    // fprintf(stderr, "std::string ref_seq len = %d\n",ref_seq.length());

    

    // convert to upper case
    std::transform(ref_seq.begin(), ref_seq.end(), ref_seq.begin(), ::toupper);
    
    // k from read pore model
    const uint32_t k = KMER_SIZE;
    // If the reference sequence contains ambiguity codes
    // switch them to the lexicographically lowest base
    ref_seq =disambiguate(ref_seq);
    std::string rc_ref_seq = reverse_complement(ref_seq);

    // --hasindu : this is already done outside of this function
    // Skip unmapped
    // if((params.record->core.flag & BAM_FUNMAP) != 0) {
    //     return alignment_output;
    // }

    // Get the read-to-reference aligned segments
    std::vector<AlignedSegment> aligned_segments = get_aligned_segments_two_params(params.record,1);
    // fprintf(stderr, "std::vector<AlignedSegment> aligned_segments len= %d\n",aligned_segments.size());
    // fprintf(stderr, "std::vector<AlignedPair> aligned_segments[0] len= %d\n",aligned_segments[0].size());

    // std::ofstream file_wrong("wfile_worng");
    // for(size_t i = 0; i < aligned_segments[0].size(); i++){
    //     int ref_pos_0 = aligned_segments[0][i].ref_pos;
    //     int read_pos_0 = aligned_segments[0][i].read_pos;
    //     file_wrong  << ref_pos_0 << " " << read_pos_0 << "\n" ;
    // }


    for(size_t segment_idx = 0; segment_idx < aligned_segments.size(); ++segment_idx) {

        AlignedSegment& aligned_pairs = aligned_segments[segment_idx];

        if(params.region_start != -1 && params.region_end != -1) {
            fprintf(stderr, "params.region_start = %d params.region_end = %d\n",params.region_start,params.region_end);
            trim_aligned_pairs_to_ref_region(aligned_pairs, params.region_start, params.region_end);
        }

        // Trim the aligned pairs to be within the range of the maximum kmer index
        int max_kmer_idx = params.read_length - k;
        trim_aligned_pairs_to_kmer(aligned_pairs, max_kmer_idx);

        if(aligned_pairs.empty()) {
            return alignment_output;
        }

        bool do_base_rc = bam_is_rev(params.record);
        bool rc_flags[2] = { do_base_rc, !do_base_rc }; // indexed by strand
        const int align_stride = 100; // approximately how many reference bases to align to at once
        const int output_stride = 50; // approximately how many event alignments to output at once

        // get the event range of the read to re-align
        int read_kidx_start = aligned_pairs.front().read_pos;
        int read_kidx_end = aligned_pairs.back().read_pos;
        
        if(do_base_rc) {
            read_kidx_start = flip_k_strand(params.read_length,read_kidx_start, k);
            read_kidx_end = flip_k_strand(params.read_length,read_kidx_end, k);
        }
        
        assert(read_kidx_start >= 0);
        assert(read_kidx_end >= 0);

        int first_event = get_closest_event_to(read_kidx_start, params.base_to_event_map, params.read_length-KMER_SIZE + 1);
        int last_event = get_closest_event_to(read_kidx_end,params.base_to_event_map, params.read_length-KMER_SIZE + 1);
        bool forward = first_event < last_event;

        int curr_start_event = first_event;
        int curr_start_ref = aligned_pairs.front().ref_pos;

//before while starts let's check all the vars
        // fprintf(stderr, "do_base_rc =  %s\n", do_base_rc ? "true" : "false");
        // fprintf(stderr, "first_event = %d last_event = %d\n",first_event,last_event);
        // fprintf(stderr, "forward =  %s\n", forward ? "true" : "false");
        // fprintf(stderr, "curr_start_event = %d curr_start_ref = %d\n",curr_start_event,curr_start_ref);


        int curr_pair_idx = 0;
        size_t tester_i = 0;
        while( (forward && curr_start_event < last_event) ||
               (!forward && curr_start_event > last_event)) {
            
        // while(tester_i == 0 ){
            // Get the index of the aligned pair approximately align_stride away
            int end_pair_idx = get_end_pair(aligned_pairs, curr_start_ref + align_stride, curr_pair_idx);
        
            int curr_end_ref = aligned_pairs[end_pair_idx].ref_pos;
            int curr_end_read = aligned_pairs[end_pair_idx].read_pos;

            if(do_base_rc) {
                curr_end_read = flip_k_strand(params.read_length,curr_end_read, k);
            }
            assert(curr_end_read >= 0);

            int s = curr_start_ref - ref_offset;
            int l = curr_end_ref - curr_start_ref + 1;

            std::string fwd_subseq = ref_seq.substr(s, l);
            std::string rc_subseq = rc_ref_seq.substr(ref_seq.length() - s - l, l);
            assert(fwd_subseq.length() == rc_subseq.length());

     //?       HMMInputSequence hmm_sequence(fwd_subseq, rc_subseq, pore_model->pmalphabet);

            //lets print the things used to build hmm_sequence
            // std::ofstream wfile("wfile");
            // wfile << fwd_subseq << "\n";
            // wfile << rc_subseq << "\n";
            
            // Require a minimum amount of sequence to align to
            if(fwd_subseq.length() < 2 * k) {
                break;
            }


            //printf("[SEGMENT_START] read: %s event start: %zu event end: %zu\n", params.sr->read_name.c_str(), input.event_start_idx, input.event_stop_idx);

            // A limitation of the segment-by-segment alignment is that we can't jump
            // over very large deletions wrt to the reference. The effect of this
            // is that we can get segments that have very few alignable events. We
            // just stop processing them for now
            int input_event_stop_idx = get_closest_event_to(curr_end_read, params.base_to_event_map, params.read_length-KMER_SIZE + 1);
            
            // fprintf(stderr, "input_event_stop_idx = %d curr_start_event = %d\n",input_event_stop_idx, curr_start_event);
            if(abs((int)curr_start_event - input_event_stop_idx) < 2)
                break;
            uint8_t input_strand = 0;


            int8_t event_stride = curr_start_event < input_event_stop_idx ? 1 : -1;  
            
            // fprintf(stderr, "event_stride =  %d\n",event_stride);
            uint8_t input_rc = rc_flags[input_strand];
            std::vector<HMMAlignmentState> event_alignment = profile_hmm_align(
                fwd_subseq, //std::string fwd_subseq,
                rc_subseq,  //std::string rc_subseq,
                params.et->event,  //event_t* event,
                params.scalings,     //scalings_t scaling,
                params.model,    //model_t* cpgmodel,
                params.events_per_base, // double events_per_base
                input_strand,  //uint8_t strand,
                input_rc,   //uint8_t rc,
                k,  //uint32_t k, //KMER_SIZE
                curr_start_event,   //uint32_t e_start, //curr_start_event;
                input_event_stop_idx,   //uint32_t e_end, //input_event_stop_idx
                event_stride);  //int8_t event_stride) // event_stride 
            


            // fprintf(stderr, "std::vector<HMMAlignmentState> event_alignment len= %d\n",event_alignment.size());

            // std::ofstream wfile("wfile");
            // for(size_t i = 0; i < event_alignment.size(); i++){
            //     char state = event_alignment[i].state;
            //     wfile << state ;//<< " " << read_pos_0 << "\n" ;
            // }


            // Output alignment
            size_t num_output = 0;
            size_t event_align_idx = 0;

            // If we aligned to the last event, output everything and stop
            bool last_section = end_pair_idx == (int)aligned_pairs.size() - 1;

           

            int last_event_output = 0;
            int last_ref_kmer_output = 0;
            
            for(; event_align_idx < event_alignment.size() && 
                  (num_output < output_stride || last_section); event_align_idx++) {


                HMMAlignmentState& as = event_alignment[event_align_idx];
                if(as.state != 'K' && (int)as.event_idx != curr_start_event) {

                    event_alignment_t ea;
                    
                    
                    ea.ref_position = curr_start_ref + as.kmer_idx;
                    std::string ref__kmer = ref_seq.substr(ea.ref_position - ref_offset, k);
                    kmer_cpy(ea.ref_kmer, ref__kmer.c_str(),KMER_SIZE); 

                    // event
                    ea.read_idx = params.read_idx;
                    // ea.strand_idx = params.strand_idx;
                    ea.event_idx = as.event_idx;
                    ea.rc = rc_flags[0];

                    // hmm
                    ea.hmm_state = as.state;

                    if(ea.hmm_state != 'B') {
                        // hiruna
                        // since we are using one strand, the rc flag should be rc_flags[0] and removed calling get_kmer and replaced it with code inside
                        // from https://github.com/jts/nanopolish/blob/b9dc627e73816a415e4b96b14e8a8bf53622a6c9/src/hmm/nanopolish_hmm_input_sequence.h
                        // ea.model_kmer = ! rc_flags[0] ? fwd_subseq.substr(as.kmer_idx, k) : rc_subseq.substr(rc_subseq.length() - as.kmer_idx - k, k);
                        //hasindu : strcpy is wrong, use kmer_cpy instead
                        if(rc_flags[0]){
                            std::string kmer_one = rc_subseq.substr(rc_subseq.length() - as.kmer_idx - k, k);
                            kmer_cpy(ea.model_kmer, kmer_one.c_str(), KMER_SIZE);
                        }
                        else{
                            std::string kmer_one = fwd_subseq.substr(as.kmer_idx, k);
                            kmer_cpy(ea.model_kmer, kmer_one.c_str(), KMER_SIZE);
                        }
                    } else {
                        kmer_cpy(ea.model_kmer, std::string(k, 'N').c_str(), KMER_SIZE);
                        // ea.model_kmer = std::string(k, 'N');
                    }

                    // store
                    alignment_output.push_back(ea);

                    // update
                    last_event_output = as.event_idx;
                    last_ref_kmer_output = curr_start_ref + as.kmer_idx;

                    num_output += 1;
                }
            }
            // Advance the pair iterator to the ref base
            curr_start_event = last_event_output;
            curr_start_ref = last_ref_kmer_output;
            //printf("[SEGMENT_END] read: %s last event output: %zu ref pos: %zu (%s)\n", params.sr->read_name.c_str(), last_event_output, last_ref_kmer_output, ref_seq.substr(last_ref_kmer_output - ref_offset, k).c_str());

            curr_pair_idx = get_end_pair(aligned_pairs, curr_start_ref, curr_pair_idx);

//hiruna
// right now we don't use this to train
// #if EVENTALIGN_TRAIN
//             // update training data for read
//             params.sr->parameters[params.strand_idx].add_training_from_alignment(hmm_sequence, input, event_alignment);
//             global_training[params.strand_idx].add_training_from_alignment(hmm_sequence, input, event_alignment);
// #endif

            if(num_output == 0) {
                break;
            }
            tester_i++;
        } // for realignmentsegment
        // fprintf(stderr, "tester_i = %d\n",tester_i);
    } // for bam aligned segment
    // fprintf(stderr, "alignment_output len = %d\n",alignment_output.size());
    // assert(alignment_output.size() == 4451);
    return alignment_output;
}


// Return the duration of the specified event for one strand
inline float get_duration(const event_table* events, uint32_t event_idx)
{
    assert(event_idx < events->n);
    return (events->event)[event_idx].length;
}


inline float z_score(const event_table* events, model_t* models, scalings_t scaling,
                     uint32_t kmer_rank,
                     uint32_t event_idx,
                     uint8_t strand)
{

    float unscaledLevel = (events->event)[event_idx].mean;
    float level = unscaledLevel;
    //float scaledLevel = unscaledLevel - time * scaling.shift;

    //fprintf(stderr, "level %f\n",scaledLevel);
    //GaussianParameters gp = read.get_scaled_gaussian_from_pore_model_state(pore_model, strand, kmer_rank);
    float gp_mean =
        scaling.scale * models[kmer_rank].level_mean + scaling.shift;
    float gp_stdv = models[kmer_rank].level_stdv * scaling.var; //scaling.var = 1;
    return (level - gp_mean) / gp_stdv;
}


EventalignSummary summarize_alignment(uint32_t strand_idx,
                                      const EventAlignmentParameters& params,
                                      const std::vector<event_alignment_t>& alignments)
{
    EventalignSummary summary;
 
    summary.num_events = 0;
    summary.num_steps = 0;
    summary.num_stays = 0;
    summary.num_skips = 0;
    summary.sum_z_score = 0;
    summary.sum_duration = 0;
    summary.alignment_edit_distance = 0;
    summary.reference_span = 0;

    //assert(params.alphabet == "");
    //const PoreModel* pore_model = params.get_model();
    //uint32_t k = pore_model->k;
    uint32_t k = KMER_SIZE;

    size_t prev_ref_pos = std::string::npos;

    // the number of unique reference positions seen in the alignment
    //size_t num_unique_ref_pos = 0;

    for(size_t i = 0; i < alignments.size(); ++i) {
        const event_alignment_t& ea = alignments[i];

        summary.num_events += 1;

        // movement information
        size_t ref_move = ea.ref_position - prev_ref_pos;
        if(ref_move == 0) {
            summary.num_stays += 1;
        } else if(i != 0 && ref_move > 1) {
            summary.num_skips += 1;
        } else if(i != 0 && ref_move == 1) {
            summary.num_steps += 1;
        }

        //todo
        // event information
        summary.sum_duration += get_duration(params.et, ea.event_idx);

        //todo
        if(ea.hmm_state == 'M') {
            //fprintf(stderr,"%s\n",ea.model_kmer);
            uint32_t rank = get_kmer_rank(ea.model_kmer, k);
            double z = z_score(params.et, params.model, params.scalings, rank, ea.event_idx, 0);
            //double z = 0;
            summary.sum_z_score += z;
        }

        prev_ref_pos = ea.ref_position;
    }

    int nm = bam_aux2i(bam_aux_get(params.record, "NM"));
    summary.alignment_edit_distance = nm;
    if(!alignments.empty()) {
        summary.reference_span = alignments.back().ref_position - alignments.front().ref_position + 1;
    }
    return summary;
}


void emit_event_alignment_tsv_header(FILE* fp, int8_t print_read_names, int8_t write_samples)
{
    fprintf(fp, "%s\t%s\t%s\t%s\t%s\t", "contig", "position", "reference_kmer",
            (print_read_names? "read_name" : "read_index"), "strand");
    fprintf(fp, "%s\t%s\t%s\t%s\t", "event_index", "event_level_mean", "event_stdv", "event_length");
    fprintf(fp, "%s\t%s\t%s\t%s", "model_kmer", "model_mean", "model_stdv", "standardized_level");

    if(write_samples) {
        fprintf(fp, "\t%s", "samples");
    }
    fprintf(fp, "\n");
}

//event_mean = get_fully_scaled_level(ea.event_idx, ea.strand_idx);
// Return the observed current level after correcting for drift, shift and scale
inline float get_fully_scaled_level(const event_table* events,scalings_t scalings, uint32_t event_idx){
    float level = (events->event)[event_idx].mean;
    //float time = get_duration(events, event_idx)/ sample_rate;

    //float get_drift_scaled_level =  level - time * scalings.drift;
    //todo : is drift 0?
    float get_drift_scaled_level =  level ;
    return (get_drift_scaled_level - scalings.shift) / scalings.scale;   
}

inline model_t get_scaled_gaussian_from_pore_model_state(const event_table* events, model_t* models, scalings_t scaling,
                     uint32_t kmer_rank,
                     uint32_t event_idx)
{
    float gp_mean =
    scaling.scale * models[kmer_rank].level_mean + scaling.shift;
    float gp_stdv = models[kmer_rank].level_stdv * scaling.var; 

    model_t scaled_model;
    scaled_model.level_mean = gp_mean;
    scaled_model.level_stdv = gp_stdv;

    return scaled_model;
}

//
// std::vector<float> get_scaled_samples_for_event((const event_table* events,scalings_t scaling, uint32_t event_idx, float sample_rate) const
// {
//     double event_start_time = (events->event)[event_idx].start_time;
//     double event_duration = (events->event)[event_idx].duration;

//     size_t start_idx = this->get_sample_index_at_time(event_start_time * this->sample_rate);
//     size_t end_idx = this->get_sample_index_at_time((event_start_time + event_duration) * this->sample_rate);

//     std::vector<float> out;
//     for(size_t i = start_idx; i < end_idx; ++i) {
//         double curr_sample_time = (this->sample_start_time + i) / this->sample_rate;
//         //fprintf(stderr, "event_start: %.5lf sample start: %.5lf curr: %.5lf rate: %.2lf\n", event_start_time, this->sample_start_time / this->sample_rate, curr_sample_time, this->sample_rate);
//         double s = this->samples[i];
//         // apply scaling corrections
//         double scaled_s = s - this->scalings[strand_idx].shift;
//         assert(curr_sample_time >= (this->sample_start_time / this->sample_rate));
//         scaled_s -= (curr_sample_time - (this->sample_start_time / this->sample_rate)) * this->scalings[strand_idx].drift;
//         scaled_s /= this->scalings[strand_idx].scale;
//         out.push_back(scaled_s);
//     }
//     return out;
// }

void emit_event_alignment_tsv(FILE* fp,
                              uint32_t strand_idx,
                              const event_table* et, model_t* model, scalings_t scalings,
                              const std::vector<event_alignment_t>& alignments, 
                              int8_t print_read_names, int8_t scale_events, int8_t write_samples,
                              char* read_name, char *ref_name)
{
    //assert(params.alphabet == "");
    //const PoreModel* pore_model = params.get_model();
    //uint32_t k = pore_model->k;
    for(size_t i = 0; i < alignments.size(); ++i) {

        const event_alignment_t& ea = alignments[i];

        // basic information
        if (!print_read_names)
        {
            fprintf(fp, "%s\t%d\t%s\t%d\t%c\t",
                    ref_name, //ea.ref_name.c_str(),
                    ea.ref_position,
                    ea.ref_kmer,
                    ea.read_idx,
                    't'); //"tc"[ea.strand_idx]); 
        }
        else
        {
            fprintf(fp, "%s\t%d\t%s\t%s\t%c\t",
                    ref_name, //ea.ref_name.c_str(),
                    ea.ref_position,
                    ea.ref_kmer,
                    read_name, //sr.read_name.c_str(),
                    't'); //"tc"[ea.strand_idx]);
        }

        // event information
        float event_mean = (et->event)[ea.event_idx].mean;
        float event_stdv = (et->event)[ea.event_idx].stdv;
        float event_duration = get_duration(et, ea.event_idx);
        uint32_t rank = get_kmer_rank(ea.model_kmer, KMER_SIZE);
        float model_mean = 0.0;
        float model_stdv = 0.0;

        if(scale_events) {

            // scale reads to the model
            event_mean = get_fully_scaled_level(et, scalings,ea.event_idx);

            // unscaled model parameters
            if(ea.hmm_state != 'B') {
                model_t model1 = model[rank];
                model_mean = model1.level_mean;
                model_stdv = model1.level_stdv;
            }
        } else {

            // scale model to the reads
            if(ea.hmm_state != 'B') {
                
                model_t model1 = get_scaled_gaussian_from_pore_model_state(et, model, scalings, rank, ea.event_idx );
                model_mean = model1.level_mean;
                model_stdv = model1.level_stdv;
            }
        }

        float standard_level = (event_mean - model_mean) / (sqrt(scalings.var) * model_stdv);
        fprintf(fp, "%d\t%.2lf\t%.3lf\t%.5lf\t", ea.event_idx, event_mean, event_stdv, event_duration);
        fprintf(fp, "%s\t%.2lf\t%.2lf\t%.2lf", ea.model_kmer,
                                               model_mean,
                                               model_stdv,
                                               standard_level);

        if(write_samples) {
            assert(0);
            //todo : implement this
            // std::vector<float> samples = get_scaled_samples_for_event(ea.strand_idx, ea.event_idx);
            // std::stringstream sample_ss;
            // std::copy(samples.begin(), samples.end(), std::ostream_iterator<float>(sample_ss, ","));

            // // remove training comma
            // std::string sample_str = sample_ss.str();
            // sample_str.resize(sample_str.size() - 1);
            // fprintf(fp, "\t%s", sample_str.c_str());
        }
        fprintf(fp, "\n");
    }
}


// Realign the read in event space
void realign_read(std::vector<event_alignment_t>* event_alignment_result,EventalignSummary *summary, FILE *summary_fp, char* ref,
                  const bam_hdr_t* hdr,
                  const bam1_t* record,int32_t read_length,
                  size_t read_idx,
                  //int region_start,
                  //int region_end, 
                  event_table* events, model_t* model, index_pair_t* base_to_event_map, scalings_t scalings,
                  double events_per_base)
{
    // Load a squiggle read for the mapped read
    std::string read_name = bam_get_qname(record);

    // load read -- hasindu : we have to get rid of this sr
    //SquiggleRead sr(read_name, read_db, opt::write_samples ? SRF_LOAD_RAW_SAMPLES : 0);

    //hiruna 
    //commented this
    // if(opt::verbose > 1) {
    //     fprintf(stderr, "Realigning %s [%zu]\n",
    //             read_name.c_str(), events->n);
    // }

    //for(int strand_idx = 0; strand_idx < 2; ++strand_idx) { -- hasindu : only 1 stand in our case

        // Do not align this strand if it was not sequenced
        // if(!sr.has_events_for_strand(strand_idx)) {
        //     continue;
        // }

        EventAlignmentParameters params;
        //params.sr = &sr; -- hasindu : we have to get rid of this sr
        params.et = events; // hasindu : what is required inside the sr is to be added like this
        params.model = model; // hasindu : what is required inside the sr is to be added like this
        //params.fai = fai;
        //params.hdr = hdr;
        params.record = record;
       // params.strand_idx = strand_idx; -- hasindu : only 1 stand in our case
        
        params.read_idx = read_idx;
        params.read_length = read_length;
        //params.region_start = region_start;
        //params.region_end = region_end;

        params.region_start = -1;
        params.region_end = -1;

        params.base_to_event_map = base_to_event_map;
        params.scalings = scalings;


        params.events_per_base = events_per_base; // this is in the struct db_t. in nanopolish_arm this is the value they have calculate.


        std::vector<event_alignment_t> alignment = align_read_to_ref(params,ref);
        *event_alignment_result = alignment;


        //-- hasindu : output writing will be done outside of this function
        //EventalignSummary summary;
        //FILE* summary_fp = fopen("test/ecoli_2kb_region/f5c_event_align.summary","a");
        //F_CHK(summary_fp,"test/ecoli_2kb_region/f5c_event_align.summary");
        //todo : fix this 
        //FILE *summary_fp = stderr;
        if(summary_fp != NULL) {
            *summary = summarize_alignment(0, params, alignment);
        }

        //fprintf(summary_fp,"sfsafsaf\n");
        // if(opt::output_sam) {
        //     emit_event_alignment_sam(sam_fp, sr, hdr, record, alignment);
        // } else {
        //     emit_event_alignment_tsv(tsv_fp, sr, strand_idx, params, alignment);
        // }
        //emit_event_alignment_tsv(stdout,0,events,model,scalings,alignment, 1, 1, 0,
        //                      bam_get_qname(record), hdr->target_name[record->core.tid]);

        // if(summary_fp != NULL)
        //     fprintf(stderr, "debug summary_fp != NULL alignment.size() = %d \n",alignment.size());
        
        // if(summary.num_events > 0)
        //     fprintf(stderr, "debug summary.num_events > 0 read_idx = %d \n",read_idx);
        
        // if(summary_fp != NULL && summary.num_events > 0) {
        //     size_t strand_idx = 0;
        //     fprintf(summary_fp, "%zu\t%s\t", read_idx, read_name.c_str());
        //     fprintf(summary_fp, "%s\t%s\t%s\t",".", "dna", strand_idx == 0 ? "template" : "complement");
        //     fprintf(summary_fp, "%d\t%d\t%d\t%d\t", summary.num_events, summary.num_steps, summary.num_skips, summary.num_stays);
        //     fprintf(summary_fp, "%.2lf\t%.3lf\t%.3lf\t%.3lf\t%.3lf\n", summary.sum_duration/(4000.0), scalings.shift, scalings.scale, 0.0, scalings.var);

        // }
        // fclose(summary_fp);
        
    //}
}

///mnt/f/hasindu2008.git/nanopolish/nanopolish eventalign  -b test/ecoli_2kb_region/reads.sorted.bam -g test/ecoli_2kb_region/draft.fa -r test/ecoli_2kb_region/reads.fasta -t 1 --summary=test/ecoli_2kb_region/eventalign.summary.exp > test/ecoli_2kb_region/eventalign.exp
```
