## Supplementary material for "GPU Accelerated Adaptive Banded Event Alignment for Rapid Comparative Nanopore Signal Analysis": f5c-v0.2-beta-release: events.c

```
#include <assert.h>
#include <errno.h>
#include <float.h>
#include <inttypes.h>
#include <math.h>
#include <stdbool.h>
#include <stdint.h>
#include <stdio.h>
#include <stdlib.h>
#include <string.h>
#include <unistd.h>

#include <htslib/faidx.h>
#include <htslib/hts.h>
#include <htslib/sam.h>

#include "f5c.h"
#include "f5cmisc.h"
#include "fast5lite.h"
#include "nanopolish_read_db.h"

#ifndef DISABLE_KSORT
#include "ksort.h"

KSORT_INIT_GENERIC(float)
#endif

/**
  The following code taken from scrappie at https://github.com/nanoporetech/scrappie (c) 2016 Oxford Nanopore Technologies Ltd.
  scrappie is licensed under the Mozilla Public License 2.0
  https://github.com/nanoporetech/scrappie/blob/master/LICENSE.md
*/

typedef struct {
    size_t window_length1;
    size_t window_length2;
    float threshold1;
    float threshold2;
    float peak_height;
} detector_param;

static detector_param const event_detection_defaults = {.window_length1 = 3,
                                                        .window_length2 = 6,
                                                        .threshold1 = 1.4f,
                                                        .threshold2 = 9.0f,
                                                        .peak_height = 0.2f};

static detector_param const event_detection_rna = {.window_length1 = 7,
                                                   .window_length2 = 14,
                                                   .threshold1 = 2.5f,
                                                   .threshold2 = 9.0f,
                                                   .peak_height = 1.0f};

// From scrappie
typedef struct {
    //	Information for scaling raw data from ADC values to pA
    float digitisation;
    float offset;
    float range;
    float sample_rate;
} fast5_raw_scaling;

typedef struct {
    size_t n;
    size_t start;
    size_t end;
    float* raw;
} raw_table;

#ifdef DISABLE_KSORT
int floatcmp(const void* x, const void* y) {
    float d = *(float*)x - *(float*)y;
    return d > 0 ? 1 : -1;
}
#endif

/** Quantiles from n array
 *
 *	Using a relatively inefficent qsort resulting in O(n log n)
 *	performance but better performance is possible for small np.
 *	The array p is modified inplace, containing which quantiles to
 *	calculation on input and the quantiles on output; on error, p
 *	is filled with the value NAN.
 *
 *	@param x  An array to calculate quantiles from
 *	@param nx Length of array x
 *	@param p  An array containing quantiles to calculate [in/out]
 *	@param np Length of array p
 *
 *	@return void
 **/
void quantilef(const float* x, size_t nx, float* p, size_t np) {
    if (NULL == p) {
        return;
    }
    for (unsigned int i = 0; i < np; i++) {
        assert(p[i] >= 0.0f && p[i] <= 1.0f);
    }
    if (NULL == x) {
        for (unsigned i = 0; i < np; i++) {
            p[i] = NAN;
        }
        return;
    }
    // Sort array
    float* space = (float*)malloc(nx * sizeof(float));
    if (NULL == space) {
        for (unsigned int i = 0; i < np; i++) {
            p[i] = NAN;
        }
        return;
    }
    memcpy(space, x, nx * sizeof(float));
#ifdef DISABLE_KSORT
    qsort(space, nx, sizeof(float), floatcmp);
#else
    ks_mergesort(float, nx, space, 0);
#endif

    // Extract quantiles
    for (unsigned int i = 0; i < np; i++) {
        const size_t idx = p[i] * (nx - 1);
        const float remf = p[i] * (nx - 1) - idx;
        if (idx < nx - 1) {
            p[i] = (1.0 - remf) * space[idx] + remf * space[idx + 1];
        } else {
            // Should only occur when p is exactly 1.0
            p[i] = space[idx];
        }
    }

    free(space);
    return;
}

/** Median of an array
 *
 *	Using a relatively inefficent qsort resulting in O(n log n)
 *	performance but O(n) is possible.
 *
 *	@param x An array to calculate median of
 *	@param n Length of array
 *
 *	@return Median of array on success, NAN otherwise.
 **/
float medianf(const float* x, size_t n) {
#ifdef DISABLE_KSORT
    float p = 0.5;
    quantilef(x, n, &p, 1);
    return p;
#else
    float *copy = (float *)malloc(n * sizeof(float));
    memcpy(copy, x, n * sizeof(float));
    float m = ks_ksmall_float(n, copy, n / 2);
    free(copy);
    return m;
#endif
}

/** Median Absolute Deviation of an array
 *
 *	@param x   An array to calculate the MAD of
 *	@param n   Length of array
 *	@param med Median of the array.	 If NAN then median is calculated.
 *
 *	@return MAD of array on success, NAN otherwise.
 **/
float madf(const float* x, size_t n, const float* med) {
    const float mad_scaling_factor = 1.4826;
    if (NULL == x) {
        return NAN;
    }
    if (1 == n) {
        return 0.0f;
    }

    float* absdiff = (float*)malloc(n * sizeof(float));
    if (NULL == absdiff) {
        return NAN;
    }

    const float _med = (NULL == med) ? medianf(x, n) : *med;

    for (size_t i = 0; i < n; i++) {
        absdiff[i] = fabsf(x[i] - _med);
    }

    const float mad = medianf(absdiff, n);
    free(absdiff);
    return mad * mad_scaling_factor;
}

/** Simple segmentation of a raw read by thresholding the MAD
 *
 *	The MAD of the raw signal is calculated for non-overlapping chunks and then
 *	thresholded to find regions at the beginning and end of the signal that have
 *	unusually low variation (generally a stall or open pore).  The threshhold is
 *	derived from the distribution of the calaculated MADs.
 *
 *	The threshold is chosen to be high since a single chunk above it will trigger
 *	the end of the trimming: the threshhold is chosen so it is unlikely to be
 *	exceeded in the leader but commonly exceeded in the main read.
 *
 *	@param rt         Structure containing raw signal
 *	@param chunk_size Size of non-overlapping chunks
 *	@param perc	  The quantile to be calculated to use for threshholding
 *
 *	@return A range structure containing new start and end for read
 **/
raw_table trim_raw_by_mad(raw_table rt, int chunk_size, float perc) {
    assert(chunk_size > 1);
    assert(perc >= 0.0 && perc <= 1.0);

    const size_t nsample = rt.end - rt.start;
    const size_t nchunk = nsample / chunk_size;
    // Truncation of end to be consistent with Sloika
    rt.end = nchunk * chunk_size;

    float* madarr = (float*)malloc(nchunk * sizeof(float));
    NULL_CHK(madarr);
    for (size_t i = 0; i < nchunk; i++) {
        madarr[i] = madf(rt.raw + rt.start + i * chunk_size, chunk_size, NULL);
    }
    quantilef(madarr, nchunk, &perc, 1);

    const float thresh = perc;
    for (size_t i = 0; i < nchunk; i++) {
        if (madarr[i] > thresh) {
            break;
        }
        rt.start += chunk_size;
    }
    for (size_t i = nchunk; i > 0; i--) {
        if (madarr[i - 1] > thresh) {
            break;
        }
        rt.end -= chunk_size;
    }
    assert(rt.end > rt.start);

    free(madarr);

    return rt;
}

raw_table trim_and_segment_raw(raw_table rt, int trim_start, int trim_end,
                               int varseg_chunk, float varseg_thresh) {
    NULL_CHK(rt.raw);

    rt = trim_raw_by_mad(rt, varseg_chunk, varseg_thresh);
    NULL_CHK(rt.raw);

    rt.start += trim_start;
    rt.end -= trim_end;

    if (rt.start >= rt.end) {
        free(rt.raw);
        return (raw_table){0};
    }

    return rt;
}

////////////////////////////////////////////////////////////////////////

typedef struct {
    int DEF_PEAK_POS;
    float DEF_PEAK_VAL;
    float* signal;
    size_t signal_length;
    float threshold;
    size_t window_length;
    size_t masked_to;
    int peak_pos;
    float peak_value;
    bool valid_peak;
} Detector;
typedef Detector* DetectorPtr;

/**
 * Compute cumulative sum and sum of squares for a vector of data
 *
 *	 Element i sum (sumsq) is the sum (sum of squares) up to but excluding element i of the inputy data.
 *
 *	 @param data     double[d_length]     Data to be summed over (in)
 *	 @param sum      double[d_length + 1] Vector to store sum (out)
 *	 @param sumsq    double[d_length + 1] Vector to store sum of squares (out)
 *       @param d_length Length of data vector
 **/
void compute_sum_sumsq(const float* data, double* sum, double* sumsq,
                       size_t d_length) {
    assert(d_length > 0);

    sum[0] = 0.0f;
    sumsq[0] = 0.0f;
    for (size_t i = 0; i < d_length; ++i) {
        sum[i + 1] = sum[i] + data[i];
        sumsq[i + 1] = sumsq[i] + data[i] * data[i];
    }
}

/**
 *  Compute windowed t-statistic from summary information
 *
 *       @param sum      double[d_length] Cumulative sums of data (in)
 *	 @param sumsq    double[d_length] Cumulative sum of squares of data (in)
 *	 @param d_length Length of data vector
 *	 @param w_length Window length to calculate t-statistic over
 *
 *	 @returns float array containing tstats.  Returns NULL on error
 **/
float* compute_tstat(const double* sum, const double* sumsq, size_t d_length,
                     size_t w_length) {
    assert(d_length > 0);
    assert(w_length > 0);

    float* tstat = (float*)calloc(d_length, sizeof(float));

    const float eta = FLT_MIN;
    const float w_lengthf = (float)w_length;

    // Quick return:
    //	 t-test not defined for number of points less than 2
    //	 need at least as many points as twice the window length
    if (d_length < 2 * w_length || w_length < 2) {
        return tstat;
    }
    // fudge boundaries
    for (size_t i = 0; i < w_length; ++i) {
        tstat[i] = 0;
        tstat[d_length - i - 1] = 0;
    }

    // get to work on the rest
    for (size_t i = w_length; i <= d_length - w_length; ++i) {
        double sum1 = sum[i];
        double sumsq1 = sumsq[i];
        if (i > w_length) {
            sum1 -= sum[i - w_length];
            sumsq1 -= sumsq[i - w_length];
        }
        float sum2 = (float)(sum[i + w_length] - sum[i]);
        float sumsq2 = (float)(sumsq[i + w_length] - sumsq[i]);
        float mean1 = sum1 / w_lengthf;
        float mean2 = sum2 / w_lengthf;
        float combined_var = sumsq1 / w_lengthf - mean1 * mean1 +
                             sumsq2 / w_lengthf - mean2 * mean2;

        // Prevent problem due to very small variances
        combined_var = fmaxf(combined_var, eta);

        // t-stat
        //	Formula is a simplified version of Student's t-statistic for the
        //	special case where there are two samples of equal size with
        //	differing variance
        const float delta_mean = mean2 - mean1;
        tstat[i] = fabs(delta_mean) / sqrt(combined_var / w_lengthf);
    }

    return tstat;
}

/**
 *
 *	 @returns array of length nsample whose elements contain peak positions
 *	 Remaining elements are padded by zeros.
 **/
size_t* short_long_peak_detector(DetectorPtr short_detector,
                                 DetectorPtr long_detector,
                                 const float peak_height) {
    assert(short_detector->signal_length == long_detector->signal_length);

    const size_t ndetector = 2;
    DetectorPtr detectors[] = {short_detector, long_detector};

    size_t* peaks =
        (size_t*)calloc(short_detector->signal_length, sizeof(size_t));

    size_t peak_count = 0;
    for (size_t i = 0; i < short_detector->signal_length; i++) {
        for (unsigned int k = 0; k < ndetector; k++) {
            DetectorPtr detector = detectors[k];
            // Carry on if we've been masked out
            if (detector->masked_to >= i) {
                continue;
            }

            float current_value = detector->signal[i];

            if (detector->peak_pos == detector->DEF_PEAK_POS) {
                // CASE 1: We've not yet recorded a maximum
                if (current_value < detector->peak_value) {
                    // Either record a deeper minimum...
                    detector->peak_value = current_value;
                } else if (current_value - detector->peak_value > peak_height) {
                    // ...or we've seen a qualifying maximum
                    detector->peak_value = current_value;
                    detector->peak_pos = i;
                    // otherwise, wait to rise high enough to be considered a
                    // peak
                }
            } else {
                // CASE 2: In an existing peak, waiting to see if it is good
                if (current_value > detector->peak_value) {
                    // Update the peak
                    detector->peak_value = current_value;
                    detector->peak_pos = i;
                }
                // Dominate other tstat signals if we're going to fire at some
                // point
                if (detector == short_detector) {
                    if (detector->peak_value > detector->threshold) {
                        long_detector->masked_to =
                            detector->peak_pos + detector->window_length;
                        long_detector->peak_pos = long_detector->DEF_PEAK_POS;
                        long_detector->peak_value = long_detector->DEF_PEAK_VAL;
                        long_detector->valid_peak = false;
                    }
                }
                // Have we convinced ourselves we've seen a peak
                if (detector->peak_value - current_value > peak_height &&
                    detector->peak_value > detector->threshold) {
                    detector->valid_peak = true;
                }
                // Finally, check the distance if this is a good peak
                if (detector->valid_peak &&
                    (i - detector->peak_pos) > detector->window_length / 2) {
                    // Emit the boundary and reset
                    peaks[peak_count] = detector->peak_pos;
                    peak_count++;
                    detector->peak_pos = detector->DEF_PEAK_POS;
                    detector->peak_value = current_value;
                    detector->valid_peak = false;
                }
            }
        }
    }

    return peaks;
}

/** Create an event given boundaries
 *
 *	Note: Bounds are CADLAG (i.e. lower bound is contained in the interval but the upper bound is not).
 *
 *	@param start   Index of lower bound
 *	@param end     Index of upper bound
 *	@param sums
 *	@param sumsqs
 *	@param nsample Total number of samples in read
 *
 *	@returns An initialised event. A 'null' event is returned on error.
 **/
event_t create_event(size_t start, size_t end, double const* sums,
                     double const* sumsqs, size_t nsample) {
    assert(start < nsample);
    assert(end <= nsample);

    event_t event = {0};
    //event.pos = -1;
    //event.state = -1;

    event.start = (uint64_t)start;
    event.length = (float)(end - start);
    event.mean = (float)(sums[end] - sums[start]) / event.length;
    const float deltasqr = (sumsqs[end] - sumsqs[start]);
    const float var = deltasqr / event.length - event.mean * event.mean;
    event.stdv = sqrtf(fmaxf(var, 0.0f));
    return event;
}

event_table create_events(size_t const* peaks, double const* sums,
                          double const* sumsqs, size_t nsample) {
    event_table et = {0};

    // Count number of events found
    size_t n = 1;
    for (size_t i = 0; i < nsample; ++i) {
        if (peaks[i] > 0 && peaks[i] < nsample) {
            n++;
        }
    }

    et.event = (event_t*)calloc(n, sizeof(event_t));

    et.n = n;
    et.end = et.n;

    // First event -- starts at zero
    et.event[0] = create_event(0, peaks[0], sums, sumsqs, nsample);
    // Other events -- peak[i-1] -> peak[i]
    for (size_t ev = 1; ev < n - 1; ev++) {
        et.event[ev] =
            create_event(peaks[ev - 1], peaks[ev], sums, sumsqs, nsample);
    }
    // Last event -- ends at nsample
    et.event[n - 1] =
        create_event(peaks[n - 2], nsample, sums, sumsqs, nsample);

    return et;
}

event_table detect_events(raw_table const rt, detector_param const edparam) {
    event_table et = {0};

    double* sums = (double*)calloc(rt.n + 1, sizeof(double));
    double* sumsqs = (double*)calloc(rt.n + 1, sizeof(double));

    compute_sum_sumsq(rt.raw, sums, sumsqs, rt.n);
    float* tstat1 = compute_tstat(sums, sumsqs, rt.n, edparam.window_length1);
    float* tstat2 = compute_tstat(sums, sumsqs, rt.n, edparam.window_length2);

    Detector short_detector = {.DEF_PEAK_POS = -1,
                               .DEF_PEAK_VAL = FLT_MAX,
                               .signal = tstat1,
                               .signal_length = rt.n,
                               .threshold = edparam.threshold1,
                               .window_length = edparam.window_length1,
                               .masked_to = 0,
                               .peak_pos = -1,
                               .peak_value = FLT_MAX,
                               .valid_peak = false};

    Detector long_detector = {.DEF_PEAK_POS = -1,
                              .DEF_PEAK_VAL = FLT_MAX,
                              .signal = tstat2,
                              .signal_length = rt.n,
                              .threshold = edparam.threshold2,
                              .window_length = edparam.window_length2,
                              .masked_to = 0,
                              .peak_pos = -1,
                              .peak_value = FLT_MAX,
                              .valid_peak = false};

    size_t* peaks = short_long_peak_detector(&short_detector, &long_detector,
                                             edparam.peak_height);

    et = create_events(peaks, sums, sumsqs, rt.n);

    free(peaks);
    free(tstat2);
    free(tstat1);
    free(sumsqs);
    free(sums);

    return et;
}

// interface to scrappie functions
event_table getevents(size_t nsample, float* rawptr) {
    event_table et;
    raw_table rt = (raw_table){nsample, 0, nsample, rawptr};

    // trim using scrappie's internal method
    // parameters taken directly from scrappie defaults
    int trim_start = 200;
    int trim_end = 10;
    int varseg_chunk = 100;
    float varseg_thresh = 0.0;
    trim_and_segment_raw(rt, trim_start, trim_end, varseg_chunk, varseg_thresh);

    const detector_param* ed_params = &event_detection_defaults; // for dna only
    et = detect_events(rt, *ed_params);

    return et;
}
```
