## Supplementary material for "GPU Accelerated Adaptive Banded Event Alignment for Rapid Comparative Nanopore Signal Analysis": f5c-v0.2-beta-release: f5c.c

```
/* @f5c
**
** f5c interface 
** @author: Hasindu Gamaarachchi
** @@
******************************************************************************/

#include <assert.h>
#include <math.h>
#include <pthread.h>
#include <stdio.h>
#include <stdlib.h>
#include <string.h>

#include "f5c.h"
#include "f5cmisc.h"


/*
todo :
Error counter for consecutive failures in the skip unreadable mode
not all the memory allocations are needed for eventalign mode
*/

core_t* init_core(const char* bamfilename, const char* fastafile,
                  const char* fastqfile, const char* tmpfile, opt_t opt,double realtime0, int8_t mode) {
    core_t* core = (core_t*)malloc(sizeof(core_t));
    MALLOC_CHK(core);

    // load bam file
    core->m_bam_fh = sam_open(bamfilename, "r");
    NULL_CHK(core->m_bam_fh);

    // load bam index file
    core->m_bam_idx = sam_index_load(core->m_bam_fh, bamfilename);
    if(core->m_bam_idx==NULL){
        ERROR("could not load the .bai index file for %s", bamfilename);
        fprintf(stderr, "Please run 'samtools index %s'\n", bamfilename);
        exit(EXIT_FAILURE);
    }

    // read the bam header
    core->m_hdr = sam_hdr_read(core->m_bam_fh);
    NULL_CHK(core->m_hdr);

    core->itr = sam_itr_queryi(core->m_bam_idx, HTS_IDX_START, 0, 0);
    NULL_CHK(core->itr);

    //open the bam file for writing skipped ultra long reads
    core->ultra_long_tmp=NULL; //todo :  at the moment this is used to detect if the load balance mode is enabled. A better method in the opt flags.
    if(tmpfile!=NULL){
        core->ultra_long_tmp = sam_open(tmpfile, "wb");
        NULL_CHK(core->ultra_long_tmp);

        //write the header to the temporary file
        int ret_sw=sam_hdr_write(core->ultra_long_tmp,core->m_hdr);
        NEG_CHK(ret_sw);
    }

    if(opt.flag & F5C_WR_RAW_DUMP){
        core->raw_dump = fopen("f5c.tmp.bin","wb");
        F_CHK(core->raw_dump,"f5c.tmp.bin");
    }
    if(opt.flag & F5C_RD_RAW_DUMP){
        core->raw_dump = fopen("f5c.tmp.bin","rb");
        F_CHK(core->raw_dump,"f5c.tmp.bin");
    }    

    // reference file
    core->fai = fai_load(fastafile);
    NULL_CHK(core->fai);

    // readbb
    core->readbb = new ReadDB;
    core->readbb->load(fastqfile);

    //model
    core->model = (model_t*)malloc(sizeof(model_t) * NUM_KMER); //4096 is 4^6 which is hardcoded now
    MALLOC_CHK(core->model);
    core->cpgmodel = (model_t*)malloc(sizeof(model_t) * NUM_KMER_METH); //15625 is 4^6 which os hardcoded now
    MALLOC_CHK(core->cpgmodel);

    //load the model from files
    if (opt.model_file) {
        read_model(core->model, opt.model_file);
    } else {
        set_model(core->model);
    }

    //todo (low priority) : load the cpg model from file
    set_cpgmodel(core->cpgmodel);

    core->opt = opt;

    //realtime0
    core->realtime0=realtime0;

    core->load_db_time=0;
    core->process_db_time=0;

    core->db_bam_time=0;
    core->db_fasta_time=0;
    core->db_fast5_time=0;
    core->db_fast5_open_time=0;
    core->db_fast5_read_time=0;

    core->event_time=0;
    core->align_time=0;
    core->est_scale_time=0;
    core->meth_time=0;   

    //cuda stuff
#ifdef HAVE_CUDA
    if (!(core->opt.flag & F5C_DISABLE_CUDA)) {
        init_cuda(core);
    }
#endif

    core->sum_bases=0;
    core->total_reads=0; //total number mapped entries in the bam file (after filtering based on flags, mapq etc)
    core->bad_fast5_file=0; //empty fast5 path returned by readdb, could not open fast5
    core->ultra_long_skipped=0;
    core->qc_fail_reads=0;
    core->failed_calibration_reads=0;
    core->failed_alignment_reads=0;

    //eventalign related
    core->mode = mode;
    if(mode==1){
        core->event_summary_fp = fopen("f5c_event_align.summary.txt","w");
        F_CHK(core->event_summary_fp,"f5c_event_align.summary.txt");
    }


    return core;
}

void free_core(core_t* core) {
    free(core->model);
    free(core->cpgmodel);
    delete core->readbb;
    fai_destroy(core->fai);
    sam_itr_destroy(core->itr);
    bam_hdr_destroy(core->m_hdr);
    hts_idx_destroy(core->m_bam_idx);
    sam_close(core->m_bam_fh);
    if(core->ultra_long_tmp!=NULL){
        sam_close(core->ultra_long_tmp);
    }
    if(core->opt.flag&F5C_WR_RAW_DUMP || core->opt.flag&F5C_RD_RAW_DUMP){
        fclose(core->raw_dump);
    }    
#ifdef HAVE_CUDA
    if (!(core->opt.flag & F5C_DISABLE_CUDA)) {
        free_cuda(core);
    }
#endif
    //eventalign related
    if(core->mode==1){
        fclose(core->event_summary_fp);
    }    
    free(core);
}

db_t* init_db(core_t* core) {
    db_t* db = (db_t*)(malloc(sizeof(db_t)));
    MALLOC_CHK(db);

    db->capacity_bam_rec = core->opt.batch_size;
    db->n_bam_rec = 0;

    db->bam_rec = (bam1_t**)(malloc(sizeof(bam1_t*) * db->capacity_bam_rec));
    MALLOC_CHK(db->bam_rec);

    int32_t i = 0;
    for (i = 0; i < db->capacity_bam_rec; ++i) {
        db->bam_rec[i] = bam_init1();
        NULL_CHK(db->bam_rec[i]);
    }

    db->fasta_cache = (char**)(malloc(sizeof(char*) * db->capacity_bam_rec));
    MALLOC_CHK(db->fasta_cache);
    db->read = (char**)(malloc(sizeof(char*) * db->capacity_bam_rec));
    MALLOC_CHK(db->read);
    db->read_len = (int32_t*)(malloc(sizeof(int32_t) * db->capacity_bam_rec));
    MALLOC_CHK(db->read_len);

    db->f5 = (fast5_t**)malloc(sizeof(fast5_t*) * db->capacity_bam_rec);
    MALLOC_CHK(db->f5);

    db->et = (event_table*)malloc(sizeof(event_table) * db->capacity_bam_rec);
    MALLOC_CHK(db->et);

    db->scalings =
        (scalings_t*)malloc(sizeof(scalings_t) * db->capacity_bam_rec);
    MALLOC_CHK(db->scalings);

    db->event_align_pairs =
        (AlignedPair**)malloc(sizeof(AlignedPair*) * db->capacity_bam_rec);
    MALLOC_CHK(db->event_align_pairs);
    db->n_event_align_pairs =
        (int32_t*)malloc(sizeof(int32_t) * db->capacity_bam_rec);
    MALLOC_CHK(db->n_event_align_pairs);

    db->event_alignment = (event_alignment_t**)malloc(
        sizeof(event_alignment_t*) * db->capacity_bam_rec);
    MALLOC_CHK(db->event_alignment);
    db->n_event_alignment =
        (int32_t*)malloc(sizeof(int32_t*) * db->capacity_bam_rec);
    MALLOC_CHK(db->n_event_alignment);

    db->events_per_base =
        (double*)malloc(sizeof(double*) * db->capacity_bam_rec);
    MALLOC_CHK(db->events_per_base);

    db->base_to_event_map =
        (index_pair_t**)malloc(sizeof(index_pair_t*) * db->capacity_bam_rec);
    MALLOC_CHK(db->base_to_event_map);

    db->read_stat_flag = (int32_t *)malloc(sizeof(int32_t) * db->capacity_bam_rec);
    MALLOC_CHK(db->read_stat_flag);

    db->site_score_map = (std::map<int, ScoredSite> **)malloc(sizeof(std::map<int, ScoredSite> *) * db->capacity_bam_rec);
    MALLOC_CHK(db->site_score_map);

    for (i = 0; i < db->capacity_bam_rec; ++i) {
        db->site_score_map[i] = new std::map<int, ScoredSite>;
        NULL_CHK(db->site_score_map[i]);
    }

    db->total_reads=0;
    db->bad_fast5_file=0;
    db->ultra_long_skipped=0;

    //eventalign related
    if(core->mode==1){
        db->eventalign_summary = (EventalignSummary *)malloc(sizeof(EventalignSummary) * db->capacity_bam_rec);
        db->event_alignment_result = (std::vector<event_alignment_t> **)malloc(sizeof(std::vector<event_alignment_t> *) * db->capacity_bam_rec);
        MALLOC_CHK(db->event_alignment_result);
        for (i = 0; i < db->capacity_bam_rec; ++i) {
            db->event_alignment_result[i] = new std::vector<event_alignment_t> ;
            NULL_CHK(db->event_alignment_result[i]);
        }
    }
    else{
        db->eventalign_summary = NULL;
        db->event_alignment_result = NULL;
    }

    return db;
}

static inline void handle_bad_fast5(core_t* core, db_t* db,std::string fast5_path_str, std::string qname){
    db->bad_fast5_file++;
    if (core->opt.flag & F5C_SKIP_UNREADABLE) {
        WARNING("Fast5 file [%s] for read [%s] is unreadable and will be skipped",
                fast5_path_str.c_str(),qname.c_str());
    } else {
        ERROR("Fast5 file [%s] could not be opened for read [%s]", fast5_path_str.c_str(), qname.c_str());
        exit(EXIT_FAILURE);
    }
    return;
}


//make this inline for performance reasons
void f5write(FILE* fp, void *buf, size_t element_size, size_t num_elements){
	size_t ret=fwrite(buf,element_size,num_elements,fp);
	if(ret!=num_elements){
		fprintf(stderr,"Writing error has occurred :%s\n",strerror(errno));
		exit(EXIT_FAILURE);
	}
}

static inline void f5read(FILE* fp, void *buf, size_t element_size, size_t num_elements){
	size_t ret=fread(buf,element_size,num_elements,fp);
	if(ret!=num_elements){
		fprintf(stderr,"Reading error has occurred :%s\n",strerror(errno));
		exit(EXIT_FAILURE);
	}
}


static inline int read_from_fast5_dump(core_t *core, db_t *db , int32_t i){

    //return 1 if success, 0 if failed
    db->f5[i] = (fast5_t*)calloc(1, sizeof(fast5_t));
    MALLOC_CHK(db->f5[i]);

    f5read(core->raw_dump,&(db->f5[i]->nsample), sizeof(hsize_t), 1);

    if(db->f5[i]->nsample>0){
        db->f5[i]->rawptr = (float*)calloc(db->f5[i]->nsample, sizeof(float));
        MALLOC_CHK( db->f5[i]->rawptr);
        f5read(core->raw_dump,db->f5[i]->rawptr, sizeof(float), db->f5[i]->nsample);
        f5read(core->raw_dump,&(db->f5[i]->digitisation), sizeof(float), 1);
        f5read(core->raw_dump,&(db->f5[i]->offset), sizeof(float), 1);
        f5read(core->raw_dump,&(db->f5[i]->range), sizeof(float), 1);
        f5read(core->raw_dump,&(db->f5[i]->sample_rate), sizeof(float), 1);
        return 1;
    }
    else{
        return 0;
    }


}

static inline int read_from_fast5_files(core_t *core, db_t *db, std::string qname, std::string fast5_path_str, int32_t i){
    char* fast5_path =
        (char*)malloc(fast5_path_str.size() + 10); // is +10 needed? do errorcheck
    strcpy(fast5_path, fast5_path_str.c_str());

    //fprintf(stderr,"readname : %s\n",qname.c_str());
    int8_t success=0;

    double t = realtime();
    hid_t hdf5_file = fast5_open(fast5_path);
    double ot = realtime() - t;
    core->db_fast5_open_time += ot;
    core->db_fast5_time += ot;
    if (hdf5_file >= 0) {
        db->f5[i] = (fast5_t*)calloc(1, sizeof(fast5_t));
        MALLOC_CHK(db->f5[i]);
        t = realtime();
        int32_t ret=fast5_read(hdf5_file, db->f5[i]);
        double rt = realtime() - t;
        core->db_fast5_read_time += rt;
        core->db_fast5_time += rt;
        if(ret<0){
            handle_bad_fast5(core, db,fast5_path,qname);
            if(core->opt.flag & F5C_WR_RAW_DUMP){
                hsize_t tmp_nsample = 0;
                f5write(core->raw_dump,&tmp_nsample, sizeof(hsize_t), 1);
            }
            free(fast5_path);
            return 0;
        }
        t = realtime();
        fast5_close(hdf5_file);
        core->db_fast5_time += realtime() - t;

        if (core->opt.flag & F5C_PRINT_RAW) {
            printf(">%s\tPATH:%s\tLN:%llu\n", qname.c_str(), fast5_path,
                db->f5[i]->nsample);
            uint32_t j = 0;
            for (j = 0; j < db->f5[i]->nsample; j++) {
                printf("%d\t", (int)db->f5[i]->rawptr[j]);
            }
            printf("\n");
        }
        if(core->opt.flag & F5C_WR_RAW_DUMP){
            //write the fast5 dump to the binary file pointer core->raw_dump
            f5write(core->raw_dump,&(db->f5[i]->nsample), sizeof(hsize_t), 1);
            f5write(core->raw_dump,db->f5[i]->rawptr, sizeof(float), db->f5[i]->nsample);
            f5write(core->raw_dump,&(db->f5[i]->digitisation), sizeof(float), 1);
            f5write(core->raw_dump,&(db->f5[i]->offset), sizeof(float), 1);
            f5write(core->raw_dump,&(db->f5[i]->range), sizeof(float), 1);
            f5write(core->raw_dump,&(db->f5[i]->sample_rate), sizeof(float), 1);
        }

        //db->n_bam_rec++;
        //t = realtime();
        //status.num_bases += read_length;
        //core->db_fasta_time += realtime() - t;
        success=1;
    } else {
        handle_bad_fast5(core, db,fast5_path,qname);
        if(core->opt.flag & F5C_WR_RAW_DUMP){
            hsize_t tmp_nsample = 0;
            f5write(core->raw_dump,&tmp_nsample, sizeof(hsize_t), 1);
        }
        return 0;
    }
    free(fast5_path);
    assert(success==1);
    return 1;
}

ret_status_t load_db(core_t* core, db_t* db) {

    double load_start = realtime();

    // get bams
    bam1_t* record;
    int32_t result = 0;
    db->n_bam_rec = 0;
    db->sum_bases = 0;
    db->total_reads = 0;
    db->bad_fast5_file = 0;
    db->ultra_long_skipped =0;

    ret_status_t status={0,0};
    int32_t i = 0;
    double t = 0;
    while (db->n_bam_rec < db->capacity_bam_rec && status.num_bases<core->opt.batch_size_bases) {
        i=db->n_bam_rec;
        record = db->bam_rec[i];
        t = realtime();
        result = sam_itr_next(core->m_bam_fh, core->itr, record);
        core->db_bam_time += realtime() - t;

        if (result < 0) {
            break;
        } else {
            if ((record->core.flag & BAM_FUNMAP) == 0 &&
                record->core.qual >= core->opt.min_mapq) {
                // printf("%s\t%d\n",bam_get_qname(db->bam_rec[db->n_bam_rec]),result);

                if(!(core->opt.flag & F5C_SECONDARY_YES)){
                    if((record->core.flag & BAM_FSECONDARY)){
                        continue;
                    }
                }

                db->total_reads++; // candidate read

                std::string qname = bam_get_qname(record);
                t = realtime();
                //todo : make efficient (redudantly accessed below, can be combined with it?)
                int64_t read_length=core->readbb->get_read_sequence(qname).size();
                std::string fast5_path_str = core->readbb->get_signal_path(qname);
                core->db_fasta_time += realtime() - t;

                //skipping ultra-long-reads
                if(core->ultra_long_tmp!=NULL && read_length > core->opt.ultra_thresh){
                    db->ultra_long_skipped++;
                    int ret_wr=sam_write1(core->ultra_long_tmp,core->m_hdr,record);
                    NEG_CHK(ret_wr);  
                    continue;
                }

                if(fast5_path_str==""){
                    handle_bad_fast5(core, db,fast5_path_str,qname);
                    continue;
                }

                int8_t read_status = 0;    
                if (core->opt.flag & F5C_RD_RAW_DUMP){
                    t = realtime();                  
                    read_status=read_from_fast5_dump(core, db,i);     
                    double rt = realtime() - t;
                    core->db_fast5_read_time += rt;
                    core->db_fast5_time += rt;                      
                }
                else{    
                   read_status=read_from_fast5_files(core, db, qname,fast5_path_str,i);
                }
                if(read_status==1){
                    db->n_bam_rec++;
                    status.num_bases += read_length;
                }

            }
        }
    }
    // fprintf(stderr,"%s:: %d queries read\n",__func__,db->n_bam_rec);

    // get ref sequences (todo can make efficient by taking the the start and end of the sorted bam)
    for (i = 0; i < db->n_bam_rec; i++) {
        bam1_t* record = db->bam_rec[i];
        char* ref_name = core->m_hdr->target_name[record->core.tid];
        // printf("refname : %s\n",ref_name);
        int32_t ref_start_pos = record->core.pos;
        int32_t ref_end_pos = bam_endpos(record);
        assert(ref_end_pos >= ref_start_pos);

        // Extract the reference sequence for this region
        int32_t fetched_len = 0;
        t = realtime();
        char* refseq = faidx_fetch_seq(core->fai, ref_name, ref_start_pos, ref_end_pos, &fetched_len); // todo : error handle?
        core->db_fasta_time += realtime() - t;
        db->fasta_cache[i] = refseq;
        // printf("seq : %s\n",db->fasta_cache[i]);

        // get the fast5

        std::string qname = bam_get_qname(db->bam_rec[i]);
        t = realtime();
        std::string read_seq = core->readbb->get_read_sequence(qname);
        core->db_fasta_time += realtime() - t;

        //get the read in ascci
        db->read[i] =
            (char*)malloc(read_seq.size() + 1); // todo : is +1 needed? do errorcheck
        strcpy(db->read[i], read_seq.c_str());
        db->read_len[i] = strlen(db->read[i]);
        db->sum_bases += db->read_len[i];

        db->read_stat_flag[i] = 0; //reset the flag
    }
    // fprintf(stderr,"%s:: %d fast5 read\n",__func__,db->n_bam_rec);
    if(core->opt.verbosity>1){
        STDERR("Average read len %.0f",db->sum_bases/(float)db->n_bam_rec);
    }
    status.num_reads=db->n_bam_rec;
    assert(status.num_bases==db->sum_bases);

    double load_end = realtime();
    core->load_db_time += (load_end-load_start);

    return status;
}

#ifdef WORK_STEAL
static inline int32_t steal_work(pthread_arg_t* all_args, int32_t n_threads)
{

	int32_t i, c_i = -1;
	int32_t k;
	for (i = 0; i < n_threads; ++i){
        pthread_arg_t args = all_args[i];
        //fprintf(stderr,"endi : %d, starti : %d\n",args.endi,args.starti);
		if (args.endi-args.starti > STEAL_THRESH) {
            //fprintf(stderr,"gap : %d\n",args.endi-args.starti);
            c_i = i;
            break;
        }
    }
    if(c_i<0){
        return -1;
    }
	k = __sync_fetch_and_add(&(all_args[c_i].starti), 1);
    //fprintf(stderr,"k : %d, end %d, start %d\n",k,all_args[c_i].endi,all_args[c_i].starti);
	return k >= all_args[c_i].endi ? -1 : k;
}
#endif

void* pthread_single(void* voidargs) {
    int32_t i;
    pthread_arg_t* args = (pthread_arg_t*)voidargs;
    db_t* db = args->db;
    core_t* core = args->core;

#ifndef WORK_STEAL
    for (i = args->starti; i < args->endi; i++) {
        args->func(core,db,i);
    }
#else
    pthread_arg_t* all_args = (pthread_arg_t*)(args->all_pthread_args);
    //adapted from kthread.c in minimap2
	for (;;) {
		i = __sync_fetch_and_add(&args->starti, 1);
		if (i >= args->endi) {
            break;
        }
		args->func(core,db,i);
	}
	while ((i = steal_work(all_args,core->opt.num_thread)) >= 0){
		args->func(core,db,i);
    }
#endif

    //fprintf(stderr,"Thread %d done\n",(myargs->position)/THREADS);
    pthread_exit(0);
}


void pthread_db(core_t* core, db_t* db, void (*func)(core_t*,db_t*,int)){
    //create threads
    pthread_t tids[core->opt.num_thread];
    pthread_arg_t pt_args[core->opt.num_thread];
    int32_t t, ret;
    int32_t i = 0;
    int32_t num_thread = core->opt.num_thread;
    int32_t step = (db->n_bam_rec + num_thread - 1) / num_thread;
    //todo : check for higher num of threads than the data
    //current works but many threads are created despite

    //set the data structures
    for (t = 0; t < num_thread; t++) {
        pt_args[t].core = core;
        pt_args[t].db = db;
        pt_args[t].starti = i;
        i += step;
        if (i > db->n_bam_rec) {
            pt_args[t].endi = db->n_bam_rec;
        } else {
            pt_args[t].endi = i;
        }
        pt_args[t].func=func;
    #ifdef WORK_STEAL
        pt_args[t].all_pthread_args =  (void *)pt_args;
    #endif
        //fprintf(stderr,"t%d : %d-%d\n",t,pt_args[t].starti,pt_args[t].endi);

    }

    //create threads
    for(t = 0; t < core->opt.num_thread; t++){
        ret = pthread_create(&tids[t], NULL, pthread_single,
                                (void*)(&pt_args[t]));
        NEG_CHK(ret);
    }

    //pthread joining
    for (t = 0; t < core->opt.num_thread; t++) {
        int ret = pthread_join(tids[t], NULL);
        NEG_CHK(ret);
    }
}


void event_single(core_t* core,db_t* db, int32_t i) {

    float* rawptr = db->f5[i]->rawptr;
    float range = db->f5[i]->range;
    float digitisation = db->f5[i]->digitisation;
    float offset = db->f5[i]->offset;
    int32_t nsample = db->f5[i]->nsample;

    // convert to pA
    float raw_unit = range / digitisation;
    for (int32_t j = 0; j < nsample; j++) {
        rawptr[j] = (rawptr[j] + offset) * raw_unit;
    }
    db->et[i] = getevents(db->f5[i]->nsample, rawptr);

    // if(db->et[i].n/(float)db->read_len[i] > 20){
    //     fprintf(stderr,"%s\tevents_per_base\t%f\tread_len\t%d\n",bam_get_qname(db->bam_rec[i]), db->et[i].n/(float)db->read_len[i],db->read_len[i]);
    // }

    //get the scalings
    db->scalings[i] = estimate_scalings_using_mom(
        db->read[i], db->read_len[i], core->model, db->et[i]);

}

void event_db(core_t* core, db_t* db){

    if (core->opt.num_thread == 1) {
        int32_t i=0;
        for (i = 0; i < db->n_bam_rec; i++) {
            event_single(core,db,i);
        }

    }

    else {
        pthread_db(core,db,event_single);
    }

}


void scaling_single(core_t* core, db_t* db, int32_t i){

    db->event_alignment[i] = NULL;
    db->n_event_alignment[i] = 0;
    db->events_per_base[i] = 0; //todo : is double needed? not just int8?

    int32_t n_kmers = db->read_len[i] - KMER_SIZE + 1;
    db->base_to_event_map[i]=(index_pair_t*)(malloc(sizeof(index_pair_t) * n_kmers));
    MALLOC_CHK(db->base_to_event_map[i]);

    if (db->n_event_align_pairs[i] > 0) {
        // prepare data structures for the final calibration

        db->event_alignment[i] = (event_alignment_t*)malloc(
            sizeof(event_alignment_t) * db->n_event_align_pairs[i]);
        MALLOC_CHK(db->event_alignment[i]);

        // for (int j = 0; j < n_event_align_pairs; ++j) {
        //     fprintf(stderr, "%d-%d\n",event_align_pairs[j].ref_pos,event_align_pairs[j].read_pos);
        // }


        //todo : verify if this n is needed is needed
        db->n_event_alignment[i] = postalign(
            db->event_alignment[i],db->base_to_event_map[i], &db->events_per_base[i], db->read[i],
            n_kmers, db->event_align_pairs[i], db->n_event_align_pairs[i]);

        //fprintf(stderr,"n_event_alignment %d\n",n_events);

        // run recalibration to get the best set of scaling parameters and the residual
        // between the (scaled) event levels and the model.

        // internally this function will set shift/scale/etc of the pore model
        bool calibrated = recalibrate_model(
            core->model, db->et[i], &db->scalings[i],
            db->event_alignment[i], db->n_event_alignment[i], 1);

        // QC calibration
        if (!calibrated || db->scalings[i].var > MIN_CALIBRATION_VAR) {
            //     events[strand_idx].clear();
            free(db->event_alignment[i]);
            //free(db->event_align_pairs[i]);
            db->read_stat_flag[i] |= FAILED_CALIBRATION;
            return;
        }

        free(db->event_alignment[i]);

    } else {
        // Could not align, fail this read
        // this->events[strand_idx].clear();
        // this->events_per_base[strand_idx] = 0.0f;
        //free(db->event_align_pairs[i]);
        db->read_stat_flag[i] |= FAILED_ALIGNMENT;
        return;
    }

    // Filter poor quality reads that have too many "stays"

    if (db->events_per_base[i] > 5.0) {
        //     events[0].clear();
        //     events[1].clear();
        //free(db->event_align_pairs[i]);
        db->read_stat_flag[i] |= FAILED_QUALITY_CHK;
        return;
    }


}

void scaling_db(core_t* core, db_t* db){
    if (core->opt.num_thread == 1) {
        int32_t i=0;
        for (i = 0; i < db->n_bam_rec; i++) {
            scaling_single(core,db,i);
        }

    }
    else {
        pthread_db(core,db,scaling_single);
    }
}

void align_single(core_t* core, db_t* db, int32_t i) {
    db->n_event_align_pairs[i] = align(
            db->event_align_pairs[i], db->read[i], db->read_len[i], db->et[i],
            core->model, db->scalings[i], db->f5[i]->sample_rate);
        //fprintf(stderr,"readlen %d,n_events %d\n",db->read_len[i],n_event_align_pairs);
}


void align_db(core_t* core, db_t* db) {
#ifdef HAVE_CUDA
    if (!(core->opt.flag & F5C_DISABLE_CUDA)) {
        //STDERR("%s","Performing on cuda");
        align_cuda(core, db);
    }
#endif

    if (core->opt.flag & F5C_DISABLE_CUDA) {
        //fprintf(stderr, "cpu\n");
        if (core->opt.num_thread == 1) {
            int i;
            for (i = 0; i < db->n_bam_rec; i++) {
                align_single(core, db, i);
            }
        } else {
            pthread_db(core, db, align_single);
        }
    }
}


void meth_single(core_t* core, db_t* db, int32_t i){
    if(!db->read_stat_flag[i]){
        if(core->mode==0){
            calculate_methylation_for_read(db->site_score_map[i], db->fasta_cache[i], db->bam_rec[i], db->read_len[i], db->et[i].event, db->base_to_event_map[i],
            db->scalings[i], core->cpgmodel,db->events_per_base[i]);
        }
        else if (core->mode==1){
            realign_read(db->event_alignment_result[i], &(db->eventalign_summary[i]),core->event_summary_fp, db->fasta_cache[i],core->m_hdr,
                  db->bam_rec[i],db->read_len[i],
                  i,
                  &(db->et[i]), core->model,db->base_to_event_map[i],db->scalings[i],db->events_per_base[i]);
        }
    }
}

void meth_db(core_t* core, db_t* db) {
    if (core->opt.num_thread == 1) {
        int i;
        for (i = 0; i < db->n_bam_rec; i++) {
            meth_single(core, db, i);
        }
    }
    else {
        pthread_db(core, db, meth_single);
    }
}


void process_single(core_t* core, db_t* db,int32_t i) {

    event_single(core,db,i);

    db->event_align_pairs[i] = (AlignedPair*)malloc(
        sizeof(AlignedPair) * db->et[i].n * 2); //todo : find a good heuristic to save memory //todo : save memory by freeing here itself
    MALLOC_CHK(db->event_align_pairs[i]);

    align_single(core, db,i);

    db->event_alignment[i] = NULL;
    db->n_event_alignment[i] = 0;
    db->events_per_base[i] = 0; //todo : is double needed? not just float?

    int32_t n_kmers = db->read_len[i] - KMER_SIZE + 1;
    db->base_to_event_map[i]=(index_pair_t*)(malloc(sizeof(index_pair_t) * n_kmers));
    MALLOC_CHK(db->base_to_event_map[i]);

    if (db->n_event_align_pairs[i] > 0) {
        // prepare data structures for the final calibration

        db->event_alignment[i] = (event_alignment_t*)malloc(
            sizeof(event_alignment_t) * db->n_event_align_pairs[i]);
        MALLOC_CHK(db->event_alignment[i]);

        // for (int j = 0; j < n_event_align_pairs; ++j) {
        //     fprintf(stderr, "%d-%d\n",event_align_pairs[j].ref_pos,event_align_pairs[j].read_pos);
        // }


        //todo : verify if this n is needed is needed
        db->n_event_alignment[i] = postalign(
            db->event_alignment[i],db->base_to_event_map[i], &db->events_per_base[i], db->read[i],
            n_kmers, db->event_align_pairs[i], db->n_event_align_pairs[i]);

        //fprintf(stderr,"n_event_alignment %d\n",n_events);

        // run recalibration to get the best set of scaling parameters and the residual
        // between the (scaled) event levels and the model.

        // internally this function will set shift/scale/etc of the pore model
        bool calibrated = recalibrate_model(
            core->model, db->et[i], &db->scalings[i],
            db->event_alignment[i], db->n_event_alignment[i], 1);

        // QC calibration
        if (!calibrated || db->scalings[i].var > MIN_CALIBRATION_VAR) {
            //     events[strand_idx].clear();
            free(db->event_alignment[i]);
            //free(db->event_align_pairs[i]);
            db->read_stat_flag[i] |= FAILED_CALIBRATION;
            return;
        }

        free(db->event_alignment[i]);

    } else {
        // Could not align, fail this read
        // this->events[strand_idx].clear();
        // this->events_per_base[strand_idx] = 0.0f;
        //free(db->event_align_pairs[i]);
        db->read_stat_flag[i] |= FAILED_ALIGNMENT;
        return;
    }

    // Filter poor quality reads that have too many "stays"

    if (db->events_per_base[i] > 5.0) {
        //     events[0].clear();
        //     events[1].clear();
        //free(db->event_align_pairs[i]);
        db->read_stat_flag[i] |= FAILED_QUALITY_CHK;
        return;
    }

    if(core->mode==0){
        calculate_methylation_for_read(db->site_score_map[i], db->fasta_cache[i], db->bam_rec[i], db->read_len[i], db->et[i].event, db->base_to_event_map[i],
            db->scalings[i], core->cpgmodel,db->events_per_base[i]);
    }

    else if(core->mode==1){
        //hack
        realign_read(db->event_alignment_result[i], &(db->eventalign_summary[i]),core->event_summary_fp,db->fasta_cache[i],core->m_hdr,
                  db->bam_rec[i],db->read_len[i],
                  i,
                  &(db->et[i]), core->model,db->base_to_event_map[i],db->scalings[i],db->events_per_base[i]);    
    }
}

void process_db(core_t* core, db_t* db) {

    double process_start = realtime();

    if((core->opt.flag&F5C_SEC_PROF) || (!(core->opt.flag & F5C_DISABLE_CUDA))){

        double realtime0=core->realtime0;
        int32_t i;

        double event_start = realtime();
        event_db(core,db);
        double event_end = realtime();
        core->event_time += (event_end-event_start);

        fprintf(stderr, "[%s::%.3f*%.2f] Events computed\n", __func__,
                realtime() - realtime0, cputime() / (realtime() - realtime0));

        for (i = 0; i < db->n_bam_rec; i++) {
            db->event_align_pairs[i] = (AlignedPair*)malloc(
                sizeof(AlignedPair) * db->et[i].n * 2); //todo : find a good heuristic to save memory
            MALLOC_CHK(db->event_align_pairs[i]);
        }

        double align_start = realtime();
        align_db(core, db);
        double align_end = realtime();
        core->align_time += (align_end-align_start);

        fprintf(stderr, "[%s::%.3f*%.2f] Banded alignment done\n", __func__,
                realtime() - realtime0, cputime() / (realtime() - realtime0));

        double est_scale_start = realtime();
        scaling_db(core,db);
        double est_scale_end = realtime();
        core->est_scale_time += (est_scale_end-est_scale_start);

        fprintf(stderr, "[%s::%.3f*%.2f] Scaling calibration done\n", __func__,
                realtime() - realtime0, cputime() / (realtime() - realtime0));

        double meth_start = realtime();
        meth_db(core,db);
        double meth_end = realtime();
        core->meth_time += (meth_end-meth_start);

        fprintf(stderr, "[%s::%.3f*%.2f] HMM done\n", __func__,
                realtime() - realtime0, cputime() / (realtime() - realtime0));


    }
    else{
        if (core->opt.num_thread == 1) {
            int32_t i=0;
            for (i = 0; i < db->n_bam_rec; i++) {
                process_single(core,db,i);
            }

        }
        else {
            pthread_db(core,db,process_single);
        }

    }

    double process_end= realtime();
    core->process_db_time += (process_end-process_start);

    return;
}

void output_db(core_t* core, db_t* db) {
    if (core->opt.flag & F5C_PRINT_EVENTS) {
        int32_t i = 0;
        for (i = 0; i < db->n_bam_rec; i++) {
            printf(">%s\tLN:%d\tEVENTSTART:%d\tEVENTEND:%d\n",
                   bam_get_qname(db->bam_rec[i]), (int)db->et[i].n,
                   (int)db->et[i].start, (int)db->et[i].end);
            uint32_t j = 0;
            for (j = 0; j < db->et[i].n; j++) {
                printf("{%d,%f,%f,%f}\t", (int)db->et[i].event[j].start,
                       db->et[i].event[j].length, db->et[i].event[j].mean,
                       db->et[i].event[j].stdv);
            }
            printf("\n");
        }
    }
    if (core->opt.flag & F5C_PRINT_BANDED_ALN) {
        int32_t i = 0;
        for (i = 0; i < db->n_bam_rec; i++) {
            if((db->read_stat_flag[i]) & FAILED_ALIGNMENT){
                continue;
            }
            printf(">%s\tN_ALGN_PAIR:%d\t{ref_os,read_pos}\n",
                   bam_get_qname(db->bam_rec[i]),
                   (int)db->n_event_align_pairs[i]);
            AlignedPair* event_align_pairs = db->event_align_pairs[i];
            int32_t j = 0;
            for (j = 0; j < db->n_event_align_pairs[i]; j++) {
                printf("{%d,%d}\t", event_align_pairs[j].ref_pos,
                       event_align_pairs[j].read_pos);
            }
            printf("\n");
        }
    }

    if (core->opt.flag & F5C_PRINT_SCALING) {
        int32_t i = 0;
        printf("read\tshift\tscale\tvar\n");

        for (i = 0; i < db->n_bam_rec; i++) {
            if((db->read_stat_flag[i])&(FAILED_ALIGNMENT|FAILED_CALIBRATION)){
                continue;
            }
            printf("%s\t%.2lf\t%.2lf\t%.2lf\n", bam_get_qname(db->bam_rec[i]),
                   db->scalings[i].shift, db->scalings[i].scale,
                   db->scalings[i].var);
        }
    }

    core->sum_bases += db->sum_bases;
    core->total_reads += db->total_reads;
    core->bad_fast5_file += db->bad_fast5_file;
    core->ultra_long_skipped += db->ultra_long_skipped;

    int32_t i = 0;
    for (i = 0; i < db->n_bam_rec; i++){
        if(!db->read_stat_flag[i]){
            char* qname = bam_get_qname(db->bam_rec[i]);
            char* contig = core->m_hdr->target_name[db->bam_rec[i]->core.tid];
            
            if(core->mode==0) {
                std::map<int, ScoredSite> *site_score_map = db->site_score_map[i];
                // write all sites for this read
                for(auto iter = site_score_map->begin(); iter != site_score_map->end(); ++iter) {

                    const ScoredSite& ss = iter->second;
                    double sum_ll_m = ss.ll_methylated[0]; //+ ss.ll_methylated[1];
                    double sum_ll_u = ss.ll_unmethylated[0]; //+ ss.ll_unmethylated[1];
                    double diff = sum_ll_m - sum_ll_u;

                    // fprintf(stderr, "%s\t%d\t%d\t", ss.chromosome.c_str(), ss.start_position, ss.end_position);
                    // fprintf(stderr, "%s\t%.2lf\t", qname, diff);
                    // fprintf(stderr, "%.2lf\t%.2lf\t", sum_ll_m, sum_ll_u);
                    // fprintf(stderr, "%d\t%d\t%s\n", ss.strands_scored, ss.n_cpg, ss.sequence.c_str());

                    printf("%s\t%d\t%d\t", contig, ss.start_position, ss.end_position);
                    printf("%s\t%.2lf\t", qname, diff);
                    printf("%.2lf\t%.2lf\t", sum_ll_m, sum_ll_u);
                    printf("%d\t%d\t%s\n", ss.strands_scored, ss.n_cpg, ss.sequence.c_str());

                }
            }

            else if(core->mode==1){
                FILE* summary_fp = core->event_summary_fp;
                EventalignSummary summary = db->eventalign_summary[i];
                scalings_t scalings = db->scalings[i];
                if(summary_fp != NULL && summary.num_events > 0) {
                    size_t strand_idx = 0;
                    fprintf(summary_fp, "%d\t%s\t", i, qname);
                    fprintf(summary_fp, "%s\t%s\t%s\t",".", "dna", strand_idx == 0 ? "template" : "complement");
                    fprintf(summary_fp, "%d\t%d\t%d\t%d\t", summary.num_events, summary.num_steps, summary.num_skips, summary.num_stays);
                    fprintf(summary_fp, "%.2lf\t%.3lf\t%.3lf\t%.3lf\t%.3lf\n", summary.sum_duration/(db->f5[i]->sample_rate), scalings.shift, scalings.scale, 0.0, scalings.var);
                }
                std::vector<event_alignment_t> *event_alignment_result = db->event_alignment_result[i];
                 emit_event_alignment_tsv(stdout,0,&(db->et[i]),core->model,db->scalings[i],*event_alignment_result, 1, 0, 0,
                              qname, contig);
            }
        }
        else{
            if((db->read_stat_flag[i])&FAILED_CALIBRATION){
                core->failed_calibration_reads++;
            }
            else if ((db->read_stat_flag[i])&FAILED_ALIGNMENT){
                core->failed_alignment_reads++;
            }
            else if ((db->read_stat_flag[i])&FAILED_QUALITY_CHK){
                core->qc_fail_reads++;
            }
            else{
                assert(0);
            }
        }
    }

}

void free_db_tmp(db_t* db) {
    int32_t i = 0;
    for (i = 0; i < db->n_bam_rec; ++i) {
        bam_destroy1(db->bam_rec[i]);
        db->bam_rec[i] = bam_init1();
        free(db->fasta_cache[i]);
        free(db->read[i]);
        free(db->f5[i]->rawptr);
        free(db->f5[i]);
        free(db->et[i].event);
        free(db->event_align_pairs[i]);
        free(db->base_to_event_map[i]);
        delete db->site_score_map[i];
        db->site_score_map[i] = new std::map<int, ScoredSite>;
        
        if(db->event_alignment_result){ //eventalign related
            delete db->event_alignment_result[i];
            db->event_alignment_result[i] = new std::vector<event_alignment_t>;
        }
    }
}

void free_db(db_t* db) {
    int32_t i = 0;
    for (i = 0; i < db->capacity_bam_rec; ++i) {
        bam_destroy1(db->bam_rec[i]);
    }
    free(db->bam_rec);
    free(db->fasta_cache);
    free(db->read);
    free(db->read_len);
    free(db->et);
    free(db->f5);
    free(db->scalings);
    free(db->event_align_pairs);
    free(db->n_event_align_pairs);
    free(db->event_alignment);
    free(db->n_event_alignment);
    free(db->events_per_base);
    free(db->base_to_event_map);
    free(db->read_stat_flag);
    for (i = 0; i < db->capacity_bam_rec; ++i) {
        delete db->site_score_map[i];
    }
    free(db->site_score_map);
    //eventalign related
    if(db->eventalign_summary){
        free(db->eventalign_summary);
    }
    if(db->event_alignment_result){
        for (i = 0; i < db->capacity_bam_rec; ++i) {
            delete db->event_alignment_result[i];
        }
        free(db->event_alignment_result);        
    }

    free(db);
}

void init_opt(opt_t* opt) {
    memset(opt, 0, sizeof(opt_t));
    opt->min_mapq = 30;
    opt->batch_size = 512;
    opt->batch_size_bases = 2*1000*1000;
    opt->num_thread = 8;
#ifndef HAVE_CUDA
    opt->flag |= F5C_DISABLE_CUDA;
    opt->batch_size_bases = 5*1000*1000;
#endif

    opt->flag |= F5C_SKIP_UNREADABLE;
    opt->debug_break=-1;
    opt->ultra_thresh=100000;

    opt->cuda_block_size=64;
    opt->cuda_dev_id=0;
    opt->cuda_mem_frac=1.0f; //later set by cuda_init()

    //effective only if  CPU_GPU_PROC  is set
    opt->cuda_max_readlen=3.0f;
    opt->cuda_avg_events_per_kmer=2.0f; //only if CUDA_DYNAMIC_MALLOC is unset
    opt->cuda_max_avg_events_per_kmer=5.0f;
}
```
