## Supplementary material for "GPU Accelerated Adaptive Banded Event Alignment for Rapid Comparative Nanopore Signal Analysis": f5c-v0.2-beta-release: f5c.cu

```
/* @f5c
**
** f5c interface 
** @author: Hasindu Gamaarachchi
** @@
******************************************************************************/

#include <errno.h>
#include <stdint.h>
#include <stdio.h>
#include <stdlib.h>
#include <string.h>
#include <assert.h>

#include "error.h"
#include "f5c.h"
#include "f5cmisc.cuh"
#include "f5cmisc.h"


void init_cuda(core_t* core){


    cuda_exists();
    int32_t cuda_device_num = core->opt.cuda_dev_id;

    cudaDeviceProp prop;
    cudaGetDeviceProperties(&prop, cuda_device_num);
    CUDA_CHK();
    STDERR("Running on %s (device id %d)",prop.name, cuda_device_num);

    //fprintf(stderr,"AVG_EVENTS_PER_KMER %f\n",AVG_EVENTS_PER_KMER);
    //fprintf(stderr,"AVG_EVENTS_PER_KMER %f\n",AVG_EVENTS_PER_KMER_GPU_THRESH);
    //fprintf(stderr,"readfac %f\n",core->opt.cuda_max_readlen);
    assert(AVG_EVENTS_PER_KMER>0 && AVG_EVENTS_PER_KMER>0);

    core->cuda = (cuda_data_t*)malloc(sizeof(cuda_data_t));
    MALLOC_CHK(core->cuda);

    core->align_kernel_time=0;
    core->align_pre_kernel_time=0;
    core->align_core_kernel_time=0;
    core->align_post_kernel_time=0;
    core->align_cuda_malloc=0;
    core->extra_load_cpu=0;
    core->align_cuda_memcpy=0;
    core->align_cuda_postprocess=0;
    core->align_cuda_preprocess=0;

    core->previous_mem = -1;
    core->previous_count_mem = 0;
    core->previous_load = -1;
    core->previous_count_load = 0;

#ifdef CUDA_PRE_MALLOC

    int32_t n_bam_rec = core->opt.batch_size;
    //cpu arrays
    core->cuda->read_ptr_host = (ptr_t*)malloc(sizeof(ptr_t) * n_bam_rec);
    MALLOC_CHK(core->cuda->read_ptr_host);
    core->cuda->n_events_host = (int32_t*)malloc(sizeof(int32_t) * n_bam_rec);
    MALLOC_CHK(core->cuda->n_events_host);
    core->cuda->event_ptr_host = (ptr_t*)malloc(sizeof(ptr_t) * n_bam_rec);
    MALLOC_CHK(core->cuda->event_ptr_host);

    core->cuda->read_len_host = (int32_t*)malloc(sizeof(int32_t) * n_bam_rec);
    MALLOC_CHK(core->cuda->read_len_host);
    core->cuda->scalings_host = (scalings_t*)malloc(sizeof(scalings_t) * n_bam_rec);
    MALLOC_CHK(core->cuda->scalings_host);
    core->cuda->n_event_align_pairs_host = (int32_t*)malloc(sizeof(int32_t) * n_bam_rec);
    MALLOC_CHK(core->cuda->n_event_align_pairs_host);

    //cuda arrays
    if(core->opt.verbosity>1) print_size("read_ptr array",n_bam_rec * sizeof(ptr_t));
    cudaMalloc((void**)&(core->cuda->read_ptr), n_bam_rec * sizeof(ptr_t));
    CUDA_CHK();
    if(core->opt.verbosity>1) print_size("read_lens",n_bam_rec * sizeof(int32_t));
    cudaMalloc((void**)&(core->cuda->read_len), n_bam_rec * sizeof(int32_t));
    CUDA_CHK();
    //n_events
    if(core->opt.verbosity>1) print_size("n_events",n_bam_rec * sizeof(int32_t));
    cudaMalloc((void**)&(core->cuda->n_events), n_bam_rec * sizeof(int32_t));
    CUDA_CHK();
    //event ptr
    if(core->opt.verbosity>1) print_size("event ptr",n_bam_rec * sizeof(ptr_t));
    cudaMalloc((void**)&(core->cuda->event_ptr), n_bam_rec * sizeof(ptr_t));
    CUDA_CHK();
    //scalings : already linear
    if(core->opt.verbosity>1) print_size("Scalings",n_bam_rec * sizeof(scalings_t));
    cudaMalloc((void**)&(core->cuda->scalings), n_bam_rec * sizeof(scalings_t));
    CUDA_CHK();
    cudaMalloc((void**)&(core->cuda->model),
            NUM_KMER * sizeof(model_t));
    CUDA_CHK();

    if(core->opt.verbosity>1) print_size("n_event_align_pairs",n_bam_rec * sizeof(int32_t));
    cudaMalloc((void**)&(core->cuda->n_event_align_pairs), n_bam_rec * sizeof(int32_t));
    CUDA_CHK();

    //model : already linear //move to cuda_init
    cudaMemcpy(core->cuda->model, core->model, NUM_KMER * sizeof(model_t),
    cudaMemcpyHostToDevice);
    CUDA_CHK();

#ifndef CUDA_DYNAMIC_MALLOC
    // //dynamic arrays
    //compute the maximum
    uint64_t free_mem = 0;
    if(prop.integrated==1){ //in tegra free mem should be sought differently
        free_mem=tegra_freemem(cuda_device_num);
    }
    else{
        free_mem=cuda_freemem(cuda_device_num);
    }

    double factor =  1 * sizeof(char) + //read_capacity
                    AVG_EVENTS_PER_KMER * sizeof(event_t) + //event_table_capacity
                    1 * sizeof(model_t) + //model_kmer_cache_capacity
                    (AVG_EVENTS_PER_KMER * 2) * sizeof(AlignedPair) +  //event_align_pairs_capacity
                    (AVG_EVENTS_PER_KMER + 1) * ALN_BANDWIDTH * sizeof(float) + //bands_capacity
                    (AVG_EVENTS_PER_KMER + 1) * ALN_BANDWIDTH * sizeof(uint8_t)  + //trace_capacity
                    (AVG_EVENTS_PER_KMER + 1) * sizeof(EventKmerPair) ; //band_lower_left_capacity

    uint64_t sum_read_len = 0;

    //if unset by user (or set to weird values by user)
    if(core->opt.cuda_mem_frac>=1.0f || core->opt.cuda_mem_frac<=0.0f){
        if(prop.integrated==1){ //for tegra we have to reserve some space for RAM
            sum_read_len= floor(free_mem*TEGRA_MEM_FACTOR/factor);
        }
        else{
            sum_read_len= floor(free_mem*MEM_FACTOR/factor);
        }
    }
    else{
        sum_read_len= floor(free_mem*(core->opt.cuda_mem_frac)/factor);
    }

    core->cuda->max_sum_read_len = sum_read_len;
    uint64_t sum_n_events = floor(sum_read_len * AVG_EVENTS_PER_KMER);
    core->cuda->max_sum_n_events = sum_n_events;

    uint64_t read_capacity = sum_read_len * sizeof(char);
    uint64_t event_table_capacity = sum_n_events * sizeof(event_t);
    uint64_t model_kmer_cache_capacity= sum_read_len * sizeof(model_t);
    uint64_t event_align_pairs_capacity= sum_n_events * 2 * sizeof(AlignedPair);
    uint64_t bands_capacity = (sum_n_events + sum_read_len) * ALN_BANDWIDTH * sizeof(float) ;
    uint64_t trace_capacity = (sum_n_events + sum_read_len) * ALN_BANDWIDTH * sizeof(uint8_t) ;
    uint64_t band_lower_left_capacity = (sum_n_events + sum_read_len) * sizeof(EventKmerPair);

    assert(read_capacity + event_table_capacity + model_kmer_cache_capacity + event_align_pairs_capacity
    + bands_capacity + trace_capacity + band_lower_left_capacity <= free_mem);

    if(core->opt.verbosity>1) print_size("read_capacity",read_capacity);
    if(core->opt.verbosity>1) print_size("event_table_capacity",event_table_capacity);
    if(core->opt.verbosity>1) print_size("model_kmer_cache_capacity",model_kmer_cache_capacity);
    if(core->opt.verbosity>1) print_size("event_align_pairs_capacity",event_align_pairs_capacity);
    if(core->opt.verbosity>1) print_size("bands_capacity",bands_capacity);
    if(core->opt.verbosity>1) print_size("trace_capacity",trace_capacity);
    if(core->opt.verbosity>1) print_size("band_lower_left_capacity",band_lower_left_capacity);


    //input arrays
    cudaMalloc((void**)&(core->cuda->read), read_capacity); //with null char
    CUDA_CHK();
    cudaMalloc((void**)&(core->cuda->event_table), event_table_capacity);
    CUDA_CHK();
    cudaMalloc((void**)&(core->cuda->model_kmer_cache), model_kmer_cache_capacity);
    CUDA_CHK();

    /**allocate output arrays for cuda**/
    cudaMalloc((void**)&(core->cuda->event_align_pairs),event_align_pairs_capacity); //todo : need better huristic
    CUDA_CHK();

    //scratch arrays
    cudaMalloc((void**)&(core->cuda->bands), bands_capacity);
    CUDA_CHK();
    cudaMalloc((void**)&(core->cuda->trace), trace_capacity);
    CUDA_CHK();
    cudaMalloc((void**)&(core->cuda->band_lower_left), band_lower_left_capacity);
    CUDA_CHK();

    STDERR("Max GPU capacity %.1fM bases",core->cuda->max_sum_read_len/(1000.0*1000.0));
    int64_t  num_bases_gap = core->cuda->max_sum_read_len - core->opt.batch_size_bases;
    if(num_bases_gap>500000){
        INFO("Your GPU can accommodate upto %.1fM bases. You may increase -B option (currently %.1fM) for better performance!",
        core->cuda->max_sum_read_len/(1000.0*1000.0), core->opt.batch_size_bases/((1000.0*1000.0)));
    }
    else if(num_bases_gap<-500000){
        INFO("Your GPU can accommodate only %.1fM bases. You may decrease -B option (currently %.1fM) for better performance!",
        core->cuda->max_sum_read_len/(1000.0*1000.0), core->opt.batch_size_bases/((1000.0*1000.0)));
    }


#endif

#endif

    return;
}

void free_cuda(core_t* core){

#ifdef CUDA_PRE_MALLOC
    free(core->cuda->event_ptr_host);
    free(core->cuda->n_events_host);
    free(core->cuda->read_ptr_host);
    free(core->cuda->read_len_host);
    free(core->cuda->scalings_host);
    free(core->cuda->n_event_align_pairs_host);

    cudaFree(core->cuda->read_ptr);
    cudaFree(core->cuda->read_len);
    cudaFree(core->cuda->n_events);
    cudaFree(core->cuda->event_ptr);
    cudaFree(core->cuda->model); //constant memory
    cudaFree(core->cuda->scalings);
    cudaFree(core->cuda->n_event_align_pairs);

#ifndef CUDA_DYNAMIC_MALLOC
    cudaFree(core->cuda->read);
    cudaFree(core->cuda->event_table);
    cudaFree(core->cuda->model_kmer_cache);
    cudaFree(core->cuda->event_align_pairs);
    cudaFree(core->cuda->bands);
    cudaFree(core->cuda->trace);
    cudaFree(core->cuda->band_lower_left);
#endif
#endif

    free(core->cuda);
    return;
}


#ifndef CPU_GPU_PROC

void align_cuda(core_t* core, db_t* db) {
    int32_t i;
    int32_t n_bam_rec = db->n_bam_rec;
    double realtime1;

    /**cuda pointers*/
    char* read;        //flattened reads sequences
    ptr_t* read_ptr; //index pointer for flattedned "reads"
    int32_t* read_len;
    int64_t sum_read_len;
    int32_t* n_events;
    event_t* event_table;
    ptr_t* event_ptr;
    int64_t sum_n_events;
    scalings_t* scalings;
    AlignedPair* event_align_pairs;
    int32_t* n_event_align_pairs;
    float *bands;
    uint8_t *trace;
    EventKmerPair* band_lower_left;

realtime1 = realtime();

#ifdef CUDA_PRE_MALLOC
    ptr_t* read_ptr_host = core->cuda->read_ptr_host;
#else
    //get the total size and create the pointers
    ptr_t* read_ptr_host = (ptr_t*)malloc(sizeof(ptr_t) * n_bam_rec);
    MALLOC_CHK(read_ptr_host);
#endif
    sum_read_len = 0;

    //read sequences : needflattening
    for (i = 0; i < n_bam_rec; i++) {
        read_ptr_host[i] = sum_read_len;
        sum_read_len += (db->read_len[i] + 1); //with null term
    }
    //form the temporary flattened array on host
    char* read_host = (char*)malloc(sizeof(char) * sum_read_len);
    MALLOC_CHK(read_host);
    for (i = 0; i < n_bam_rec; i++) {
        ptr_t idx = read_ptr_host[i];
        strcpy(&read_host[idx], db->read[i]);
    }

    //now the events : need flattening
    //num events : need flattening
    //get the total size and create the pointers
#ifdef CUDA_PRE_MALLOC
    int32_t* n_events_host = core->cuda->n_events_host;
    ptr_t* event_ptr_host = core->cuda->event_ptr_host;
#else
    int32_t* n_events_host = (int32_t*)malloc(sizeof(int32_t) * n_bam_rec);
    MALLOC_CHK(n_events_host);
    ptr_t* event_ptr_host = (ptr_t*)malloc(sizeof(ptr_t) * n_bam_rec);
    MALLOC_CHK(event_ptr_host);
#endif

    sum_n_events = 0;
    for (i = 0; i < n_bam_rec; i++) {
        n_events_host[i] = db->et[i].n;
        event_ptr_host[i] = sum_n_events;
        sum_n_events += db->et[i].n;
    }

    //event table flatten
    //form the temporary flattened array on host
    event_t* event_table_host =
        (event_t*)malloc(sizeof(event_t) * sum_n_events);
    MALLOC_CHK(event_table_host);
    for (i = 0; i < n_bam_rec; i++) {
        ptr_t idx = event_ptr_host[i];
        memcpy(&event_table_host[idx], db->et[i].event,
               sizeof(event_t) * db->et[i].n);
    }

    AlignedPair* event_align_pairs_host =
        (AlignedPair*)malloc(2 * sum_n_events * sizeof(AlignedPair));
    MALLOC_CHK(event_align_pairs_host);

core->align_cuda_preprocess += (realtime() - realtime1);

    /** Start GPU mallocs**/
realtime1 = realtime();

#ifdef CUDA_PRE_MALLOC
    read_ptr =core->cuda->read_ptr;
    read_len=core->cuda->read_len;
    n_events=core->cuda->n_events;
    event_ptr=core->cuda->event_ptr;
    scalings=core->cuda->scalings;
    model_t* model = core->cuda->model;
#else

    if(core->opt.verbosity>1) print_size("read_ptr array",n_bam_rec * sizeof(ptr_t));
    cudaMalloc((void**)&read_ptr, n_bam_rec * sizeof(ptr_t));
    CUDA_CHK();

    if(core->opt.verbosity>1) print_size("read_lens",n_bam_rec * sizeof(int32_t));
    cudaMalloc((void**)&read_len, n_bam_rec * sizeof(int32_t));
    CUDA_CHK();
    //n_events
    if(core->opt.verbosity>1) print_size("n_events",n_bam_rec * sizeof(int32_t));
    cudaMalloc((void**)&n_events, n_bam_rec * sizeof(int32_t));
    CUDA_CHK();
    //event ptr
    if(core->opt.verbosity>1) print_size("event ptr",n_bam_rec * sizeof(ptr_t));
    cudaMalloc((void**)&event_ptr, n_bam_rec * sizeof(ptr_t));
    CUDA_CHK();
    //scalings : already linear
    if(core->opt.verbosity>1) print_size("Scalings",n_bam_rec * sizeof(scalings_t));
    cudaMalloc((void**)&scalings, n_bam_rec * sizeof(scalings_t));
    CUDA_CHK();
    //model : already linear
    model_t* model;
    cudaMalloc((void**)&model,
            NUM_KMER * sizeof(model_t));
    CUDA_CHK();
#endif


    if(core->opt.verbosity>1) print_size("read array",sum_read_len * sizeof(char));
    cudaMalloc((void**)&read, sum_read_len * sizeof(char)); //with null char
    CUDA_CHK();
    if(core->opt.verbosity>1) print_size("event table",sum_n_events * sizeof(event_t));
    cudaMalloc((void**)&event_table, sum_n_events * sizeof(event_t));
    CUDA_CHK();
    model_t* model_kmer_cache;
    cudaMalloc((void**)&model_kmer_cache, sum_read_len * sizeof(model_t));
    CUDA_CHK();

    /**allocate output arrays for cuda**/
    if(core->opt.verbosity>1) print_size("event align pairs",2 * sum_n_events *sizeof(AlignedPair));
    cudaMalloc((void**)&event_align_pairs,
            2 * sum_n_events *
                sizeof(AlignedPair)); //todo : need better huristic
    CUDA_CHK();
#ifdef CUDA_PRE_MALLOC
    n_event_align_pairs=core->cuda->n_event_align_pairs;
#else
    if(core->opt.verbosity>1) print_size("n_event_align_pairs",n_bam_rec * sizeof(int32_t));
    cudaMalloc((void**)&n_event_align_pairs, n_bam_rec * sizeof(int32_t));
    CUDA_CHK();
#endif
    //scratch arrays
    size_t sum_n_bands = sum_n_events + sum_read_len; //todo : can be optimised
    if(core->opt.verbosity>1) print_size("bands",sizeof(float) * sum_n_bands * ALN_BANDWIDTH);
    cudaMalloc((void**)&bands,sizeof(float) * sum_n_bands * ALN_BANDWIDTH);
    CUDA_CHK();
    if(core->opt.verbosity>1) print_size("trace",sizeof(uint8_t) * sum_n_bands * ALN_BANDWIDTH);
    cudaMalloc((void**)&trace, sizeof(uint8_t) * sum_n_bands * ALN_BANDWIDTH);
    CUDA_CHK();
    cudaMemset(trace,0,sizeof(uint8_t) * sum_n_bands * ALN_BANDWIDTH); //initialise the trace array to 0
    if(core->opt.verbosity>1) print_size("band_lower_left",sizeof(EventKmerPair)* sum_n_bands);
    cudaMalloc((void**)&band_lower_left, sizeof(EventKmerPair)* sum_n_bands);
    CUDA_CHK();
core->align_cuda_malloc += (realtime() - realtime1);

    /* cuda mem copys*/
realtime1 =realtime();
    cudaMemcpy(read_ptr, read_ptr_host, n_bam_rec * sizeof(ptr_t),
               cudaMemcpyHostToDevice);
    CUDA_CHK();
    cudaMemcpy(read, read_host, sum_read_len * sizeof(char),
               cudaMemcpyHostToDevice);
    CUDA_CHK();
    //read length : already linear hence direct copy
    cudaMemcpy(read_len, db->read_len, n_bam_rec * sizeof(int32_t),
               cudaMemcpyHostToDevice);
    CUDA_CHK();
    cudaMemcpy(n_events, n_events_host, n_bam_rec * sizeof(int32_t),
               cudaMemcpyHostToDevice);
    CUDA_CHK();
    cudaMemcpy(event_ptr, event_ptr_host, n_bam_rec * sizeof(ptr_t),
               cudaMemcpyHostToDevice);
    CUDA_CHK();
    cudaMemcpy(event_table, event_table_host, sizeof(event_t) * sum_n_events,
               cudaMemcpyHostToDevice);
    CUDA_CHK();

#ifndef CUDA_PRE_MALLOC
//model : already linear //move to cuda_init
    cudaMemcpy(model, core->model, NUM_KMER * sizeof(model_t),
            cudaMemcpyHostToDevice);
    CUDA_CHK();
#endif
    //can be interleaved
    cudaMemcpy(scalings, db->scalings, sizeof(scalings_t) * n_bam_rec,
               cudaMemcpyHostToDevice);
    CUDA_CHK();
core->align_cuda_memcpy += (realtime() - realtime1);


realtime1 = realtime();

    /*pre kernel*/
    assert(BLOCK_LEN_BANDWIDTH>=ALN_BANDWIDTH);
    dim3 gridpre(1,(db->n_bam_rec + BLOCK_LEN_READS - 1) / BLOCK_LEN_READS);
    dim3 blockpre(BLOCK_LEN_BANDWIDTH,BLOCK_LEN_READS);
	if(core->opt.verbosity>1) fprintf(stderr,"grid %d,%d, block %d,%d\n",gridpre.x,gridpre.y, blockpre.x,blockpre.y);

    align_kernel_pre_2d<<<gridpre, blockpre>>>( read,
        read_len, read_ptr, n_events,
        event_ptr, model, n_bam_rec, model_kmer_cache,bands,trace,band_lower_left);

    cudaDeviceSynchronize();CUDA_CHK();
    if(core->opt.verbosity>1) fprintf(stderr, "[%s::%.3f*%.2f] align-pre kernel done\n", __func__,
            realtime() - realtime1, cputime() / (realtime() - realtime1));
core->align_kernel_time += (realtime() - realtime1);
core->align_pre_kernel_time += (realtime() - realtime1);

realtime1 = realtime();

    /* core kernel*/
    assert(BLOCK_LEN_BANDWIDTH>=ALN_BANDWIDTH);
    dim3 grid1(1,(db->n_bam_rec + BLOCK_LEN_READS - 1) / BLOCK_LEN_READS);
    dim3 block1(BLOCK_LEN_BANDWIDTH,BLOCK_LEN_READS);
    align_kernel_core_2d_shm<<<grid1, block1>>>(read_len, read_ptr, event_table, n_events,
            event_ptr, scalings, n_bam_rec, model_kmer_cache,bands,trace,band_lower_left );

    cudaDeviceSynchronize();CUDA_CHK();
    if(core->opt.verbosity>1) fprintf(stderr, "[%s::%.3f*%.2f] align-core kernel done\n", __func__,
    realtime() - realtime1, cputime() / (realtime() - realtime1));
    core->align_kernel_time += (realtime() - realtime1);
core->align_core_kernel_time += (realtime() - realtime1);

realtime1 = realtime();

    /*post kernel*/
    int32_t BLOCK_LEN = core->opt.cuda_block_size;
    dim3 gridpost((db->n_bam_rec + BLOCK_LEN - 1) / BLOCK_LEN);
    dim3 blockpost(BLOCK_LEN);
    #ifndef WARP_HACK
        align_kernel_post<<<gridpost, blockpost>>>(event_align_pairs, n_event_align_pairs,
            read_len, read_ptr, event_table, n_events,
            event_ptr,scalings, n_bam_rec, model_kmer_cache,bands,trace,band_lower_left );

    #else
        assert(BLOCK_LEN>=32);
        dim3 grid1post((db->n_bam_rec + (BLOCK_LEN/32) - 1) / (BLOCK_LEN/32));
        if(core->opt.verbosity>1) fprintf(stderr,"grid new %d\n",grid1post.x);
        align_kernel_post<<<grid1post, blockpost>>>(event_align_pairs, n_event_align_pairs,
            read_len, read_ptr, event_table, n_events,
            event_ptr, scalings, n_bam_rec, model_kmer_cache,bands,trace,band_lower_left );
    #endif
    cudaDeviceSynchronize();CUDA_CHK();
    if(core->opt.verbosity>1) fprintf(stderr, "[%s::%.3f*%.2f] align-post kernel done\n", __func__,
            realtime() - realtime1, cputime() / (realtime() - realtime1));
    core->align_kernel_time += (realtime() - realtime1);
core->align_post_kernel_time += (realtime() - realtime1);


    //fprintf(stderr,"readlen %d,n_events %d\n",db->read_len[i],n_event_align_pairs);

#ifdef CUDA_DEBUG

    cudaDeviceSynchronize();
    CUDA_CHK();

#endif

    /** copyback ans**/
realtime1 =  realtime();
    cudaMemcpy(db->n_event_align_pairs, n_event_align_pairs,
               n_bam_rec * sizeof(int32_t), cudaMemcpyDeviceToHost);
    CUDA_CHK();

    cudaMemcpy(event_align_pairs_host, event_align_pairs,
               2 * sum_n_events * sizeof(AlignedPair), cudaMemcpyDeviceToHost);
    CUDA_CHK();
core->align_cuda_memcpy += (realtime() - realtime1);

realtime1 =  realtime();
#ifndef CUDA_PRE_MALLOC
    cudaFree(read_ptr);
    cudaFree(read_len);
    cudaFree(n_events);
    cudaFree(event_ptr);
    cudaFree(model); //constant memory
    cudaFree(scalings);
    cudaFree(n_event_align_pairs);
#endif
    cudaFree(read); //with null char
    cudaFree(event_table);
    cudaFree(event_align_pairs);
    cudaFree(bands);
    cudaFree(trace);
    cudaFree(band_lower_left);
    cudaFree(model_kmer_cache);

core->align_cuda_malloc += (realtime() - realtime1);

    /** post work**/
realtime1 =  realtime();
    //copy back
    for (i = 0; i < n_bam_rec; i++) {
        ptr_t idx = event_ptr_host[i];
        memcpy(db->event_align_pairs[i], &event_align_pairs_host[idx * 2],
               sizeof(AlignedPair) * db->n_event_align_pairs[i]);
    }

    //free the temp arrays on host
#ifndef CUDA_PRE_MALLOC
    free(read_ptr_host);
    free(n_events_host);
    free(event_ptr_host);
#endif
    free(read_host);
    free(event_table_host);
    free(event_align_pairs_host);


core->align_cuda_postprocess += (realtime() - realtime1);

}


#else


#ifdef WORK_STEAL
static inline int32_t steal_work(pthread_arg_t* all_args, int32_t n_threads)
{

	int32_t i, c_i = -1;
	int32_t k;
	for (i = 0; i < n_threads; ++i){
        pthread_arg_t args = all_args[i];
        //fprintf(stderr,"endi : %d, starti : %d\n",args.endi,args.starti);
		if (args.endi-args.starti > STEAL_THRESH_CUDA) {
            //fprintf(stderr,"gap : %d\n",args.endi-args.starti);
            c_i = i;
            break;
        }
    }
    if(c_i<0){
        return -1;
    }
	k = __sync_fetch_and_add(&(all_args[c_i].starti), 1);
    //fprintf(stderr,"k : %d, end %d, start %d\n",k,all_args[c_i].endi,all_args[c_i].starti);
	return k >= all_args[c_i].endi ? -1 : k;
}
#endif

void* pthread_cusingle(void* voidargs) {

    double realtime1 = realtime();

    int32_t i,j;
    pthread_arg_t* args = (pthread_arg_t*)voidargs;
    db_t* db = args->db;
    core_t* core = args->core;

#ifndef WORK_STEAL
    for (i = args->starti; i < args->endi; i++) {
        j=args->ultra_long_reads[i];
        args->func(core,db,j);
    }
#else
    pthread_arg_t* all_args = (pthread_arg_t*)(args->all_pthread_args);
    //adapted from kthread
	for (;;) {
		i = __sync_fetch_and_add(&args->starti, 1);
		if (i >= args->endi) {
            break;
        }
        j=args->ultra_long_reads[i];
        if(core->opt.verbosity>2) fprintf(stderr, "[%s::%.3fsec] Thread (%d-%d) : read %d events %ld assigned\n", __func__,
            realtime() - realtime1, args->starti,args->endi, db->read_len[j], db->et[j].n );
		args->func(core,db,j);
        if(core->opt.verbosity>2) fprintf(stderr, "[%s::%.3fsec] Thread (%d-%d) : read %d events %ld done\n", __func__,
            realtime() - realtime1, args->starti,args->endi, db->read_len[j], db->et[j].n );
	}
	while ((i = steal_work(all_args,core->opt.num_thread)) >= 0){
        j=args->ultra_long_reads[i];
        if(core->opt.verbosity>2) fprintf(stderr, "[%s::%.3fsec] Thread (%d-%d) : stolen read %d events %ld assigned\n", __func__,
            realtime() - realtime1, args->starti,args->endi, db->read_len[j], db->et[j].n );
		args->func(core,db,j);
        if(core->opt.verbosity>2) fprintf(stderr, "[%s::%.3fsec] Thread (%d-%d) : stolen read %d events %ld done\n", __func__,
            realtime() - realtime1, args->starti,args->endi, db->read_len[j], db->et[j].n );
    }
#endif
    if(core->opt.verbosity>2) fprintf(stderr, "[%s::%.3fsec] Thread (%d-%d) done\n", __func__,
    realtime() - realtime1, args->starti,args->endi );
    //fprintf(stderr,"Thread %d done\n",(myargs->position)/THREADS);
    pthread_exit(0);
}


void pthread_cudb(core_t* core, db_t* db, int32_t* ultra_long_reads, int32_t  n_ultra_long_reads,void (*func)(core_t*,db_t*,int)){
    //create threads
    pthread_t tids[core->opt.num_thread];
    pthread_arg_t pt_args[core->opt.num_thread];
    int32_t t, ret;
    int32_t i = 0;
    int32_t num_thread = core->opt.num_thread;
    int32_t step = (n_ultra_long_reads + num_thread - 1) / num_thread;
    //todo : check for higher num of threads than the data
    //current works but many threads are created despite

    //set the data structures
    for (t = 0; t < num_thread; t++) {
        pt_args[t].core = core;
        pt_args[t].db = db;
        pt_args[t].starti = i;
        i += step;
        if (i > n_ultra_long_reads) {
            pt_args[t].endi = n_ultra_long_reads;
        } else {
            pt_args[t].endi = i;
        }
        pt_args[t].func=func;
        pt_args[t].ultra_long_reads=ultra_long_reads;
    #ifdef WORK_STEAL
        pt_args[t].all_pthread_args =  (void *)pt_args;
    #endif
        //fprintf(stderr,"t%d : %d-%d\n",t,pt_args[t].starti,pt_args[t].endi);

    }

    //create threads
    for(t = 0; t < core->opt.num_thread; t++){
        ret = pthread_create(&tids[t], NULL, pthread_cusingle,
                                (void*)(&pt_args[t]));
        NEG_CHK(ret);
    }

    //pthread joining
    for (t = 0; t < core->opt.num_thread; t++) {
        int ret = pthread_join(tids[t], NULL);
        NEG_CHK(ret);
    }
}


void* align_cudb(void* voidargs){

    double realtime1 = realtime();
    pthread_arg_t* args = (pthread_arg_t*)voidargs;
    db_t* db = args->db;
    core_t* core = args->core;
    int32_t* ultra_long_reads = args->ultra_long_reads;
    int32_t n_ultra_long_reads = args->endi;
    //fprintf(stderr,"ultra long guys : %d\n",n_ultra_long_reads);
    //fprintf(stderr, "cpu\n");
    if (core->opt.num_thread == 1) {
        int j;
        for(j=0;j<n_ultra_long_reads;j++) {
            int32_t i = ultra_long_reads[j];
            align_single(core, db, i);
            // db->n_event_align_pairs[i] =
            //     align(db->event_align_pairs[i], db->read[i],
            //           db->read_len[i], db->et[i], core->model,
            //           db->scalings[i], db->f5[i]->sample_rate);
            //fprintf(stderr,"readlen %d,n_events %d\n",db->read_len[i],n_event_align_pairs);
        }
    } else {
        pthread_cudb(core, db, ultra_long_reads,n_ultra_long_reads,align_single);

    }


    args->ret1 = realtime() - realtime1;
    if(core->opt.verbosity>1) fprintf(stderr, "[%s::%.3fsec] %d reads processed on cpu\n", __func__,
    realtime() - realtime1, n_ultra_long_reads);


    return NULL;
}

pthread_t align_cudb_async(pthread_arg_t **pt_args_ptr,core_t* core, db_t* db, int32_t* ultra_long_reads, int32_t  n_ultra_long_reads) {

    assert(*pt_args_ptr==NULL);
    *pt_args_ptr = (pthread_arg_t *)malloc(sizeof(pthread_arg_t));
    pthread_arg_t *pt_args=*pt_args_ptr;
    MALLOC_CHK(pt_args);
    pt_args->core = core;
    pt_args->db = db;
    pt_args->starti = 0;
    pt_args->endi = n_ultra_long_reads;
    pt_args->ultra_long_reads=ultra_long_reads;

    pthread_t tid;
    int ret = pthread_create(&tid, NULL, align_cudb,(void*)(pt_args));
    NEG_CHK(ret);

    return tid;
}

double align_cudb_async_join(pthread_arg_t *pt_args, pthread_t tid) {
    int ret = pthread_join(tid, NULL);
    NEG_CHK(ret);
    assert(pt_args);
    double time_cpu = pt_args->ret1;
    free(pt_args);
    return time_cpu;

}

//check if we have run out of space in the pre-allocated gpu arrays
static inline int8_t if_gpu_mem_free(core_t* core, db_t* db, int32_t i,int64_t sum_read_len,int64_t sum_n_events){
#ifdef CUDA_DYNAMIC_MALLOC
    return 1;
#else
    if((sum_read_len+(db->read_len[i] + 1) <= (int64_t)core->cuda->max_sum_read_len) &&
       (sum_n_events+db->et[i].n <= (core->cuda->max_sum_n_events)) ){
        return 1;
    }
    else{
        return 0;
    }
#endif
}

//if a suitable candidate to be run on GPU
//ultra-long reads as well as the reads with too many average events per base
//are done of CPU
static inline int8_t if_on_gpu(core_t* core, db_t* db, int32_t i){
    if(db->read_len[i]<(core->opt.cuda_max_readlen * db->sum_bases/(float)db->n_bam_rec) && (db->et[i].n)/(float)(db->read_len[i]) < AVG_EVENTS_PER_KMER_GPU_THRESH ){
        return 1;
    }
    else{
        return 0;
    }
}


#define LB_T1_DEC_K 0
#define LB_T2_INC_MAX_LF 1
#define LB_T3_INC_MAX_EPK 2
#define LB_T4_DEC_ULTRA_INC_T_CPU 3
#define LB_T5_DEC_MAX_LF_EPK 4


static inline void load_balance_advisor(core_t* core, int32_t state){
    if(core->previous_load==state){
        core->previous_count_load++;
        if(core->previous_count_load>3){
            switch (core->previous_load) {
                case LB_T1_DEC_K                    : INFO("%s","CPU got too much work. Try decreasing -K. See http://bit.ly/f5cperf");   break;
                case LB_T2_INC_MAX_LF               : INFO("%s","CPU got too much work. Try increasing --cuda-max-lf. See http://bit.ly/f5cperf");   break;
                case LB_T3_INC_MAX_EPK              : INFO("%s", "CPU got too much work. Try increasing --cuda-max-epk. See http://bit.ly/f5cperf");   break;
                case LB_T4_DEC_ULTRA_INC_T_CPU      : INFO("%s", "CPU got too much work. Try --skip-ultra, decreasing --ultra-thresh or increasing -t. Else, CPU is too weaker than GPUa and just ignore. See http://bit.ly/f5cperf");   break;
                case LB_T5_DEC_MAX_LF_EPK           : INFO("%s", "GPU got too much work. Try increasing --ultra-thresh, decreasing --cuda-max-lf, decreasing --cuda-max-epk. Else, CPU is too powerful than GPU and just ignore. See http://bit.ly/f5cperf");   break;
                default :
                    break;

            }

        }
    }
    else{
        core->previous_load=state;
        core->previous_count_load=0;
    }
}


void load_balance(core_t *core, db_t *db, double cpu_process_time,double gpu_process_time,
    int32_t stat_n_gpu_mem_out, int32_t stat_n_too_many_events, int32_t stat_n_ultra_long_reads,
    float read_array_usage, float event_array_usage){

    fprintf(stderr,"[%s] Processing time : CPU %.1f sec, GPU %.1f sec\n",__func__,cpu_process_time,gpu_process_time);
    double factor = (cpu_process_time-gpu_process_time)/(cpu_process_time+gpu_process_time);
    if (core->opt.verbosity>1) fprintf(stderr,"[%s] factor %f\n",__func__,factor);
    float thresh_factor=0.3;
    float thresh_reads=0.1;
    //float thresh=0.3;

    //cpu-gpu load balance
    if(factor>thresh_factor){ //cpu too much time
        if (core->opt.verbosity>1) fprintf(stderr,"[%s] CPU too much time\n",__func__);

        if(stat_n_gpu_mem_out > db->n_bam_rec * thresh_reads || 
            stat_n_ultra_long_reads> db->n_bam_rec * thresh_reads || 
            stat_n_too_many_events > db->n_bam_rec * thresh_reads){

            if(stat_n_gpu_mem_out > db->n_bam_rec * thresh_reads){ //gpu run out of memory
                load_balance_advisor(core,LB_T1_DEC_K);
                if (core->opt.verbosity>1) INFO("%s", "CPU did most work. If this message repeats, consider decreasing -K or -B");
            }
            else{
                if(stat_n_ultra_long_reads> db->n_bam_rec * thresh_reads){ //ultra long reads
                    load_balance_advisor(core,LB_T2_INC_MAX_LF);
                    if (core->opt.verbosity>1) INFO("%s","CPU got too many very long reads to process. If this message repeats, consider increasing --cuda-max-lf");
                }
                else{
                    if(stat_n_too_many_events > db->n_bam_rec * thresh_reads){//reads with too many events
                                load_balance_advisor(core,LB_T3_INC_MAX_EPK);
                                if (core->opt.verbosity>1) INFO("%s","CPU got too many over segmented reads to process. If this message repeats, consider  increasing --cuda-max-epk");
                    }
                    else{
                        if (core->opt.verbosity>1) INFO("%s", "Impossible exception\n");
                    }
                }

            }
        }
        else{
            load_balance_advisor(core,LB_T4_DEC_ULTRA_INC_T_CPU);
            if (core->opt.verbosity>1) INFO("%s", "CPU took too much time. If this message repeats, consider using --skip-ultra or decreasing --ultra-thresh or increasing number of CPU threads. If you tried all that means your CPU is not powerful enough to match the GPU and just ignore.");
        }

    }
    
    else if(factor<-thresh_factor){ //gpu too much time
        load_balance_advisor(core,LB_T5_DEC_MAX_LF_EPK);
        if (core->opt.verbosity>1) INFO("%s", "GPU got too much work. If this message repeats, consider increasing --ultra-thresh or decreasing --cuda-max-lf or decreasing --cuda-max-epk. If you tried all that means your GPU is not powerful enough to match the CPU and just ignore.");
    }
    else{
        if (core->opt.verbosity>1) fprintf(stderr,"[%s] No load balancing required\n",__func__);
    }
}


#define MEM_S1_EPK_INC_MAX_DEC_AVG 0
#define MEM_S2_EPK_DEC_MAX_INC_AVG 1
#define MEM_S3_INC_B 2
#define MEM_S4_INC_K 3

static inline void memory_balance_advisor(core_t* core, int32_t state){
    if(core->previous_mem==state){
        core->previous_count_mem++;
        if(core->previous_count_mem>3){
            switch (core->previous_mem) {
                case MEM_S1_EPK_INC_MAX_DEC_AVG     : INFO("%s","GPU event arrays under-utilised. Try increasing --max-epk or (decreasing --avg-epk). See http://bit.ly/f5cperf");   break;
                case MEM_S2_EPK_DEC_MAX_INC_AVG     : INFO("%s", "GPU read arrays under-utilised. Try decreasing --max-epk or (increasing --avg-epk). See http://bit.ly/f5cperf");   break;
                case MEM_S3_INC_B                   : INFO("%s","GPU arrays under-utilised. Try increasing -B. See http://bit.ly/f5cperf");   break;
                case MEM_S4_INC_K                   : INFO("%s","GPU arrays under-utilised. Try increasing -K. See http://bit.ly/f5cperf");   break;
                default :
                    break;

            }

        }
    }
    else{
        core->previous_mem=state;
        core->previous_count_mem=0;
    }
}


void memory_balance(core_t *core, db_t *db, double cpu_process_time,double gpu_process_time,
    int32_t stat_n_gpu_mem_out, int32_t stat_n_too_many_events, int32_t stat_n_ultra_long_reads,
    float read_array_usage, float event_array_usage){

    //float thresh_factor=0.3;
    //float thresh_reads=0.1;
    float thresh=0.3;

    //memory usage isssues

    //read arrays > 70%
    if(read_array_usage>100-thresh*100){
        
        //event arrays > 70%
        if(event_array_usage>100-thresh*100){
            if (core->opt.verbosity>1) fprintf(stderr,"[%s] GPU array usage good\n",__func__);
        }
        else{
            //read arrays-event arrays > 30%
            if(read_array_usage-event_array_usage>thresh*100){
                memory_balance_advisor(core,MEM_S1_EPK_INC_MAX_DEC_AVG);
                if (core->opt.verbosity>1) INFO("%s", "GPU event arrays under-utilised. If this message repeats, consider increasing --max-epk or (decreasing --avg-epk)");

            }
            else{
                if (core->opt.verbosity>1) fprintf(stderr,"[%s] GPU array usage alright\n",__func__);
            }

        }
    }
    else{
        //event arrays > 70%
        if(event_array_usage>100-thresh*100){
            //event arrays-read arrays > 30%
            if(event_array_usage-read_array_usage>thresh*100){
                memory_balance_advisor(core,MEM_S2_EPK_DEC_MAX_INC_AVG);
                if (core->opt.verbosity>1) INFO("%s", "GPU read arrays under-utilised. If this message repeats, consider decreasing --max-epk or (increasing --avg-epk)");

            }
            else{
                if (core->opt.verbosity>1) fprintf(stderr,"[%s] GPU array usage alright\n",__func__);
            }
        }
        else{
            //db->n_bam_rec is n, core->opt.batch_size is K
            //db->sum_bases is b, core->opt.batch_size_bases B
            // n<K
            if(db->n_bam_rec < core->opt.batch_size){
                //b<B
                if(db->sum_bases < core->opt.batch_size_bases){
                    if (core->opt.verbosity>1) fprintf(stderr,"[%s] Probably the last batch\n",__func__);
                }
                else{
                        memory_balance_advisor(core,MEM_S3_INC_B);
                        if (core->opt.verbosity>1) INFO("%s", "GPU arrays are not fully utilised. If this message repeats, consider increasing the --max-bases (-B option)");
                }
            }            
            else{
                //b<B
                if(db->sum_bases < core->opt.batch_size_bases){       
                    memory_balance_advisor(core,MEM_S4_INC_K);
                    if (core->opt.verbosity>1) INFO("%s", "GPU arrays are not fully utilised. If this message repeats, consider increasing the --batchsize (-K option)");
                }
                else{
                    if (core->opt.verbosity>1) INFO("%s", "Unhandled exception\n");
                }
            }
        }
    }
 
}


void align_cuda(core_t* core, db_t* db) {
    int32_t i,j;
    int32_t n_bam_rec = db->n_bam_rec;
    int32_t n_bam_rec_cuda;
    double realtime1;
    int32_t n_ultra_long_reads=0;

    int32_t stat_n_ultra_long_reads=0; //number of ultralong reads processed on CPU
    int32_t stat_n_too_many_events=0;  //number of reads with high avg events per base that are processed on CPU
    int32_t stat_n_gpu_mem_out=0;      //number of reads run on CPU due to the GPU memory running out
    int32_t sum_bases_cpu=0;           //The total sum of bases run on GPU

    int32_t ultra_long_reads[n_bam_rec]; //not only ultra-long reads, but also ones with large number of average events per base

    //cpu temp pointers
    ptr_t* read_ptr_host;
    int32_t* n_events_host;
    ptr_t* event_ptr_host;
    event_t* event_table_host;
    AlignedPair* event_align_pairs_host;
    int32_t* read_len_host;
    scalings_t* scalings_host;
    int32_t* n_event_align_pairs_host;
    char* read_host;

    /**cuda pointers*/
    char* read;        //flattened reads sequences
    ptr_t* read_ptr; //index pointer for flattedned "reads"
    int32_t* read_len;
    int64_t sum_read_len;
    int32_t* n_events;
    event_t* event_table;
    ptr_t* event_ptr;
    int64_t sum_n_events;
    scalings_t* scalings;
    AlignedPair* event_align_pairs;
    int32_t* n_event_align_pairs;
    float *bands;
    uint8_t *trace;
    EventKmerPair* band_lower_left;
    model_t* model_kmer_cache;
    model_t* model;

realtime1 = realtime();

    read_ptr_host = core->cuda->read_ptr_host;

    sum_read_len = 0;
    sum_n_events = 0;
    //read sequences : needflattening
    for (i = 0,j=0; i < n_bam_rec; i++) {
        if(if_on_gpu(core, db, i) && if_gpu_mem_free(core, db, i,sum_read_len,sum_n_events)){
            read_ptr_host[j] = sum_read_len;
            sum_read_len += (db->read_len[i] + 1); //with null term
            sum_n_events += db->et[i].n;
            j++;
        }
        else{
            if ((db->et[i].n)/(float)(db->read_len[i]) < AVG_EVENTS_PER_KMER_MAX){
                ultra_long_reads[n_ultra_long_reads]=i;
                n_ultra_long_reads++;
                sum_bases_cpu += db->read_len[i];
                if(db->read_len[i]>=(core->opt.cuda_max_readlen * db->sum_bases/(float)db->n_bam_rec)){
                    stat_n_ultra_long_reads++;
                    if(core->opt.verbosity>2)STDERR("readlen>=%.0fkbases\t%d",(core->opt.cuda_max_readlen * db->sum_bases/(float)db->n_bam_rec)/1000,db->read_len[i]);
                }
                else if ((db->et[i].n)/(float)(db->read_len[i]) >= AVG_EVENTS_PER_KMER_GPU_THRESH){
                    stat_n_too_many_events++;
                }
                else{
                    stat_n_gpu_mem_out++;
                }
            }
            else{//todo : too many avg events per base, even for the CPU
                db->n_event_align_pairs[i]=0;
            }

        }
    }
    n_bam_rec_cuda = j;


    //can start processing on the ultra long reads on the CPU
    pthread_arg_t *tmparg=NULL;
    pthread_t tid =  align_cudb_async(&tmparg,core, db, ultra_long_reads, n_ultra_long_reads);

    double realtime_process_start=realtime();

    read_len_host = core->cuda->read_len_host;
    scalings_host = core->cuda->scalings_host;
    n_event_align_pairs_host = core->cuda->n_event_align_pairs_host;

    //form the temporary flattened array on host
    read_host = (char*)malloc(sizeof(char) * sum_read_len);
    MALLOC_CHK(read_host);
    sum_read_len = 0;
    sum_n_events = 0;
    for (i = 0,j=0; i < n_bam_rec; i++) {
        if(if_on_gpu(core, db, i) && if_gpu_mem_free(core, db, i,sum_read_len,sum_n_events)){
            ptr_t idx = read_ptr_host[j];
            strcpy(&read_host[idx], db->read[i]);
            read_len_host[j]=db->read_len[i];
            scalings_host[j]=db->scalings[i];
            j++;
            sum_read_len += (db->read_len[i] + 1); //with null term
            sum_n_events += db->et[i].n;
        }

    }


    //now the events : need flattening
    //num events : need flattening
    //get the total size and create the pointers
    n_events_host = core->cuda->n_events_host;
    event_ptr_host = core->cuda->event_ptr_host;

    sum_read_len = 0;
    sum_n_events = 0;
    for (i = 0,j=0; i < n_bam_rec; i++) {
        if(if_on_gpu(core, db, i) && if_gpu_mem_free(core, db, i,sum_read_len,sum_n_events)){
            n_events_host[j] = db->et[i].n;
            event_ptr_host[j] = sum_n_events;
            sum_n_events += db->et[i].n;
            j++;
            sum_read_len += (db->read_len[i] + 1); //with null term
        }
    }

    //event table flatten
    //form the temporary flattened array on host
    event_table_host =
        (event_t*)malloc(sizeof(event_t) * sum_n_events);
    MALLOC_CHK(event_table_host);
    sum_read_len = 0;
    sum_n_events = 0;
    for (i = 0,j=0; i < n_bam_rec; i++) {
        if(if_on_gpu(core, db, i) && if_gpu_mem_free(core, db, i,sum_read_len,sum_n_events)){
            ptr_t idx = event_ptr_host[j];
            memcpy(&event_table_host[idx], db->et[i].event,
                sizeof(event_t) * db->et[i].n);
                j++;
            sum_read_len += (db->read_len[i] + 1); //with null term
            sum_n_events += db->et[i].n;
            }
    }

    event_align_pairs_host =
        (AlignedPair*)malloc(2 * sum_n_events * sizeof(AlignedPair));
    MALLOC_CHK(event_align_pairs_host);

core->align_cuda_preprocess += (realtime() - realtime1);

    /** Start GPU mallocs**/
realtime1 = realtime();


    read_ptr =core->cuda->read_ptr;
    read_len=core->cuda->read_len;
    n_events=core->cuda->n_events;
    event_ptr=core->cuda->event_ptr;
    scalings=core->cuda->scalings;
    model = core->cuda->model;
    n_event_align_pairs=core->cuda->n_event_align_pairs;

#ifndef CUDA_DYNAMIC_MALLOC

    assert(sum_read_len <= (int64_t)core->cuda->max_sum_read_len);
    assert(sum_n_events <= (int64_t)(core->cuda->max_sum_n_events));
    //fprintf(stderr,"%d %d\n", sum_read_len,sum_n_events);
    if(core->opt.verbosity>1) STDERR("%.2f %% of GPU read arrays and %.2f %% of GPU event arrays were utilised",
        sum_read_len/(float)(core->cuda->max_sum_read_len)*100 ,
        sum_n_events/(float)(core->cuda->max_sum_n_events)*100);

    read=(core->cuda->read);
    event_table=(core->cuda->event_table);
    model_kmer_cache=(core->cuda->model_kmer_cache);
    event_align_pairs=(core->cuda->event_align_pairs);
    bands=(core->cuda->bands);
    trace=(core->cuda->trace);
    band_lower_left=(core->cuda->band_lower_left);

    cudaMemset(trace,0,sizeof(uint8_t) * (sum_n_events + sum_read_len) * ALN_BANDWIDTH); //initialise the trace array to 0
    CUDA_CHK();

#else
    if(core->opt.verbosity>1) print_size("read array",sum_read_len * sizeof(char));
    cudaMalloc((void**)&read, sum_read_len * sizeof(char)); //with null char
    CUDA_CHK();
    if(core->opt.verbosity>1) print_size("event table",sum_n_events * sizeof(event_t));
    cudaMalloc((void**)&event_table, sum_n_events * sizeof(event_t));
    CUDA_CHK();
    if(core->opt.verbosity>1) print_size("model kmer cache",sum_read_len * sizeof(model_t));
    cudaMalloc((void**)&model_kmer_cache, sum_read_len * sizeof(model_t));
    CUDA_CHK();

    /**allocate output arrays for cuda**/
    if(core->opt.verbosity>1) print_size("event align pairs",2 * sum_n_events *sizeof(AlignedPair));
    cudaMalloc((void**)&event_align_pairs,
            2 * sum_n_events *
                sizeof(AlignedPair)); //todo : need better huristic
    CUDA_CHK();


    //scratch arrays
    size_t sum_n_bands = sum_n_events + sum_read_len; //todo : can be optimised
    if(core->opt.verbosity>1) print_size("bands",sizeof(float) * sum_n_bands * ALN_BANDWIDTH);
    cudaMalloc((void**)&bands,sizeof(float) * sum_n_bands * ALN_BANDWIDTH);
    CUDA_CHK();
    if(core->opt.verbosity>1) print_size("trace",sizeof(uint8_t) * sum_n_bands * ALN_BANDWIDTH);
    cudaMalloc((void**)&trace, sizeof(uint8_t) * sum_n_bands * ALN_BANDWIDTH);
    CUDA_CHK();
    cudaMemset(trace,0,sizeof(uint8_t) * sum_n_bands * ALN_BANDWIDTH); //initialise the trace array to 0
    CUDA_CHK();
    if(core->opt.verbosity>1) print_size("band_lower_left",sizeof(EventKmerPair)* sum_n_bands);
    cudaMalloc((void**)&band_lower_left, sizeof(EventKmerPair)* sum_n_bands);
    CUDA_CHK();

#endif


core->align_cuda_malloc += (realtime() - realtime1);

    /* cuda mem copys*/
realtime1 =realtime();
    cudaMemcpy(read_ptr, read_ptr_host, n_bam_rec_cuda * sizeof(ptr_t),
               cudaMemcpyHostToDevice);
    CUDA_CHK();
    cudaMemcpy(read, read_host, sum_read_len * sizeof(char),
               cudaMemcpyHostToDevice);
    CUDA_CHK();
    //read length : already linear hence direct copy
    cudaMemcpy(read_len, read_len_host, n_bam_rec_cuda * sizeof(int32_t),
               cudaMemcpyHostToDevice);
    CUDA_CHK();
    cudaMemcpy(n_events, n_events_host, n_bam_rec_cuda * sizeof(int32_t),
               cudaMemcpyHostToDevice);
    CUDA_CHK();
    cudaMemcpy(event_ptr, event_ptr_host, n_bam_rec_cuda * sizeof(ptr_t),
               cudaMemcpyHostToDevice);
    CUDA_CHK();
    cudaMemcpy(event_table, event_table_host, sizeof(event_t) * sum_n_events,
               cudaMemcpyHostToDevice);
    CUDA_CHK();


    //can be interleaved
    cudaMemcpy(scalings, scalings_host, sizeof(scalings_t) * n_bam_rec_cuda,
               cudaMemcpyHostToDevice);
    CUDA_CHK();
core->align_cuda_memcpy += (realtime() - realtime1);


realtime1 = realtime();

    if(n_bam_rec_cuda>0){ 
    /*pre kernel*/
    assert(BLOCK_LEN_BANDWIDTH>=ALN_BANDWIDTH);
        dim3 gridpre(1,(n_bam_rec_cuda + BLOCK_LEN_READS - 1) / BLOCK_LEN_READS);
        dim3 blockpre(BLOCK_LEN_BANDWIDTH,BLOCK_LEN_READS);
        if(core->opt.verbosity>1) STDERR("grid %d,%d, block %d,%d",gridpre.x,gridpre.y, blockpre.x,blockpre.y);
        align_kernel_pre_2d<<<gridpre, blockpre>>>( read,
            read_len, read_ptr, n_events,
            event_ptr, model, n_bam_rec_cuda, model_kmer_cache,bands,trace,band_lower_left);
        cudaDeviceSynchronize();CUDA_CHK();
        if(core->opt.verbosity>1) fprintf(stderr, "[%s::%.3fsec] align-pre kernel done\n", __func__,
            realtime() - realtime1);
    }
core->align_kernel_time += (realtime() - realtime1);
core->align_pre_kernel_time += (realtime() - realtime1);

realtime1 = realtime();

    /* core kernel*/
    if(n_bam_rec_cuda>0){ 
        assert(BLOCK_LEN_BANDWIDTH>=ALN_BANDWIDTH);
        dim3 grid1(1,(n_bam_rec_cuda + BLOCK_LEN_READS - 1) / BLOCK_LEN_READS);
        dim3 block1(BLOCK_LEN_BANDWIDTH,BLOCK_LEN_READS);
        align_kernel_core_2d_shm<<<grid1, block1>>>(read_len, read_ptr, event_table, n_events,
                event_ptr, scalings, n_bam_rec_cuda, model_kmer_cache,bands,trace,band_lower_left );

        cudaDeviceSynchronize();CUDA_CHK();
        if(core->opt.verbosity>1) fprintf(stderr, "[%s::%.3fsec] align-core kernel done\n", __func__,
        realtime() - realtime1);
    }
core->align_kernel_time += (realtime() - realtime1);
core->align_core_kernel_time += (realtime() - realtime1);

realtime1 = realtime();

    /*post kernel*/
    if(n_bam_rec_cuda>0){ 
        int32_t BLOCK_LEN = core->opt.cuda_block_size;
        dim3 gridpost((n_bam_rec_cuda + BLOCK_LEN - 1) / BLOCK_LEN);
        dim3 blockpost(BLOCK_LEN);
        #ifndef WARP_HACK
            align_kernel_post<<<gridpost, blockpost>>>(event_align_pairs, n_event_align_pairs,
                read_len, read_ptr, event_table, n_events,
                event_ptr,scalings, n_bam_rec_cuda, model_kmer_cache,bands,trace,band_lower_left );

        #else
            assert(BLOCK_LEN>=32);
            dim3 grid1post((n_bam_rec_cuda + (BLOCK_LEN/32) - 1) / (BLOCK_LEN/32));
            if(core->opt.verbosity>1) STDERR("grid new %d",grid1post.x);
            align_kernel_post<<<grid1post, blockpost>>>(event_align_pairs, n_event_align_pairs,
                read_len, read_ptr, event_table, n_events,
                event_ptr, scalings, n_bam_rec_cuda, model_kmer_cache,bands,trace,band_lower_left );
        #endif
        cudaDeviceSynchronize();CUDA_CHK();
        if(core->opt.verbosity>1) fprintf(stderr, "[%s::%.3fsec] align-post kernel done\n", __func__,
                realtime() - realtime1);
    }
    core->align_kernel_time += (realtime() - realtime1);
core->align_post_kernel_time += (realtime() - realtime1);


    //fprintf(stderr,"readlen %d,n_events %d\n",db->read_len[i],n_event_align_pairs);

#ifdef CUDA_DEBUG

    cudaDeviceSynchronize();
    CUDA_CHK();

#endif

    /** copyback ans**/
realtime1 =  realtime();
    cudaMemcpy(n_event_align_pairs_host, n_event_align_pairs,
               n_bam_rec_cuda * sizeof(int32_t), cudaMemcpyDeviceToHost);
    CUDA_CHK();

    cudaMemcpy(event_align_pairs_host, event_align_pairs,
               2 * sum_n_events * sizeof(AlignedPair), cudaMemcpyDeviceToHost);
    CUDA_CHK();
core->align_cuda_memcpy += (realtime() - realtime1);

realtime1 =  realtime();

#ifdef CUDA_DYNAMIC_MALLOC
    cudaFree(read); //with null char
    cudaFree(event_table);
    cudaFree(event_align_pairs);
    cudaFree(bands);
    cudaFree(trace);
    cudaFree(band_lower_left);
    cudaFree(model_kmer_cache);
#endif

core->align_cuda_malloc += (realtime() - realtime1);

    /** post work**/
realtime1 =  realtime();
    //copy back
    sum_read_len = 0;
    sum_n_events = 0;
    for (i = 0,j=0; i < n_bam_rec; i++) {
        if(if_on_gpu(core, db, i) && if_gpu_mem_free(core, db, i,sum_read_len,sum_n_events)){
            ptr_t idx = event_ptr_host[j];
            db->n_event_align_pairs[i]=n_event_align_pairs_host[j];
    #ifdef REVERSAL_ON_CPU
            int c;
            int end = db->n_event_align_pairs[i] - 1;
            AlignedPair* out_2= db->event_align_pairs[i];
            AlignedPair* in_2= &event_align_pairs_host[idx * 2];
            for (c = 0; c < db->n_event_align_pairs[i] ; c++) {
                out_2[c].ref_pos = in_2[end].ref_pos;
                out_2[c].read_pos = in_2[end].read_pos;
                end--;
            }
    #else
            memcpy(db->event_align_pairs[i], &event_align_pairs_host[idx * 2],
                sizeof(AlignedPair) * db->n_event_align_pairs[i]);
    #endif
            j++;
            sum_read_len += (db->read_len[i] + 1); //with null term
            sum_n_events += db->et[i].n;

        }
    }

    //free the temp arrays on host
    free(read_host);
    free(event_table_host);
    free(event_align_pairs_host);

core->align_cuda_postprocess += (realtime() - realtime1);

    double gpu_process_time = realtime()-realtime_process_start;

realtime1 =  realtime();
    double cpu_process_time = align_cudb_async_join(tmparg,tid);
core->extra_load_cpu += (realtime() - realtime1);

    if(core->opt.verbosity>1) {
        fprintf(stderr, "[%s::%.3fsec] CPU extra processing done (>=%.0fkbases:%d|>=%.1fevents:%d|gpu_mem_out:%d)\n",
        __func__,realtime() - realtime1,((core->opt.cuda_max_readlen * db->sum_bases/(float)db->n_bam_rec))/1000,
        stat_n_ultra_long_reads, AVG_EVENTS_PER_KMER_GPU_THRESH,stat_n_too_many_events, stat_n_gpu_mem_out);
    }


    STDERR("Load : CPU %d entries (%.1fM bases), GPU %d entries (%.1fM bases)",
    n_bam_rec-n_bam_rec_cuda, (float)sum_bases_cpu/(1000*1000),n_bam_rec_cuda, (float)sum_read_len/(1000*1000));

    load_balance(core,db,cpu_process_time,gpu_process_time,stat_n_gpu_mem_out,stat_n_too_many_events, stat_n_ultra_long_reads,
        sum_read_len/(float)(core->cuda->max_sum_read_len)*100 ,
        sum_n_events/(float)(core->cuda->max_sum_n_events)*100);
    memory_balance(core,db,cpu_process_time,gpu_process_time,stat_n_gpu_mem_out,stat_n_too_many_events, stat_n_ultra_long_reads,
        sum_read_len/(float)(core->cuda->max_sum_read_len)*100 ,
        sum_n_events/(float)(core->cuda->max_sum_n_events)*100);
}


#endif
```
