## Supplementary material for "GPU Accelerated Adaptive Banded Event Alignment for Rapid Comparative Nanopore Signal Analysis": f5c-v0.2-beta-release: f5c.h

```
/* @f5c
**
** f5c interface 
** @author: Hasindu Gamaarachchi
** @@
******************************************************************************/

#ifndef F5C_H
#define F5C_H

#include <htslib/faidx.h>
#include <htslib/hts.h>
#include <htslib/sam.h>

#include "fast5lite.h"
#include "nanopolish_read_db.h"

//required for eventalign
#include <vector>

#define F5C_VERSION "0.1-beta"

/* hard coded numbers*/
#define KMER_SIZE 6 //hard coded for now; todo : change to dynamic?
#define NUM_KMER 4096   //num k-mers for 6-mers DNA
#define NUM_KMER_METH 15625 //number k-mers for 6-mers with methylated C
//#define HAVE_CUDA 1 //if compiled for CUDA or not
#define ALN_BANDWIDTH 100 // the band size in adaptive_banded_dynamic_alignment

/*flags related to the user specified options (opt_t)*/
#define F5C_PRINT_RAW 0x001     //print the raw signal to stdio
#define F5C_SECONDARY_YES 0x002 //consider secondary reads
#define F5C_SKIP_UNREADABLE 0x004 //Skip unreadable fast5 and continue rather than exiting
#define F5C_PRINT_EVENTS 0x008 //print the event table
#define F5C_PRINT_BANDED_ALN 0x010 //print the event alignment
#define F5C_PRINT_SCALING 0x020 //print the estimated scalings
#define F5C_DISABLE_CUDA 0x040 //disable cuda (only when compile for cuda)
//#define F5C_DEBUG_BRK 0x080 //break after the first batch //removed can be reused
#define F5C_SEC_PROF 0x100 //profile section by section (only effective on the CPU mode)
#define F5C_WR_RAW_DUMP 0x200 //to say if we should write the raw dump of the fast5
#define F5C_RD_RAW_DUMP 0x400 //to say if we should read the raw dump fof the fast5


/*flags for a read status (related to db_t->read_stat_flag)*/
#define FAILED_CALIBRATION 0x001 //if the calibration failed
#define FAILED_ALIGNMENT 0x002 //if the alignment failed
#define FAILED_QUALITY_CHK  0x004 //if the quality check failed


/* other hard coded options */

/*CPU thread scheduling options for multithreading framework for processing*/
#define WORK_STEAL 1 //simple work stealing enabled or not (no work stealing mean no load balancing)
#define STEAL_THRESH 1 //stealing threshold for the CPU only sections
#define STEAL_THRESH_CUDA 0 //stealing threshold for the CPU part in a GPU accelerated workload

//set if input, processing and output are not to be interleaved (serial mode) - useful for debugging
//#define IO_PROC_NO_INTERLEAVE 1

//#define ALIGN_2D_ARRAY 1 //for CPU whether to use a 1D array or a 2D array
// note : (2D arrays are very slow due to mallocs when the number of threads is high)

#define CACHED_LOG 1 //if the log values of scalings and the model k-mers are cached
//#define LOAD_SD_MEANSSTDV 1 //if the sd_mean and the sd_stdv is to be loaded (they are unused anyway)

#define ESL_LOG_SUM 1 // enable the fast log sum for HMM


typedef int64_t ptr_t;

/* data structures */

//user options
typedef struct {
    int32_t min_mapq;           //minimum mapq
    const char* model_file;     //name of the model file
    uint32_t flag;              //flags
    int32_t batch_size;         //max reads loaded at once
    int64_t batch_size_bases;   //max bases loaded at once

    int32_t num_thread;
    int8_t verbosity;
    int32_t debug_break;
    int64_t ultra_thresh;

    //todo : these are required only for HAVE_CUDA (but need to chnage the meth_main accordingly)
    int32_t cuda_block_size;
    float cuda_max_readlen;
    float cuda_avg_events_per_kmer;
    float cuda_max_avg_events_per_kmer;
    int32_t cuda_dev_id;
    float cuda_mem_frac;
} opt_t;

// a single event : adapted from taken from scrappie
typedef struct {
    uint64_t start;
    float length; //todo : cant be made int?
    float mean;
    float stdv;
    //int32_t pos;   //todo : always -1 can be removed
    //int32_t state; //todo : always -1 can be removed
} event_t;

// event table : adapted from scrappie
typedef struct {
    size_t n;     //todo : int32_t not enough?
    size_t start; //todo : always 0?
    size_t end;   //todo : always equal to n?
    event_t* event;
} event_table;

//k-mer model
typedef struct {
    float level_mean;
    float level_stdv;

#ifdef CACHED_LOG
    //calculated for efficiency
    float level_log_stdv;
#endif

#ifdef LOAD_SD_MEANSSTDV
    //float sd_mean;
    //float sd_stdv;
    //float weight;
#endif
} model_t;

//scaling parameters for the signal : taken from nanopolish
typedef struct {
    // direct parameters that must be set
    float scale;
    float shift;
    //float drift; = 0 always?
    float var; // set later when calibrating
    //float scale_sd;
    //float var_sd;

    // derived parameters that are cached for efficiency
#ifdef CACHED_LOG
    float log_var;
#endif
    //float scaled_var;
    //float log_scaled_var;
} scalings_t;

//from nanopolish
typedef struct {
        int event_idx;
        int kmer_idx;
} EventKmerPair;

//from nanopolish
typedef struct {
    int ref_pos;
    int read_pos;
} AlignedPair;

//from nanopolish
typedef struct {
    int32_t start;
    int32_t stop; // inclusive
} index_pair_t;

//from nanopolish
typedef struct {
    // ref data
    //char* ref_name;
    char ref_kmer[KMER_SIZE + 1];
    int32_t ref_position;

    // event data
    int32_t read_idx;
    //int32_t strand_idx;
    int32_t event_idx;
    bool rc;

    // hmm data
    char model_kmer[KMER_SIZE + 1];
    char hmm_state;
} event_alignment_t;

//from nanopolish
struct ScoredSite
{
    //toto : clean up unused
    ScoredSite()
    {
        ll_unmethylated[0] = 0;
        ll_unmethylated[1] = 0;
        ll_methylated[0] = 0;
        ll_methylated[1] = 0;
        strands_scored = 0;
    }

    std::string chromosome;
    int start_position;
    int end_position;
    int n_cpg;
    std::string sequence;

    // scores per strand
    double ll_unmethylated[2];
    double ll_methylated[2];
    int strands_scored;

    //
    static bool sort_by_position(const ScoredSite& a, const ScoredSite& b) { return a.start_position < b.start_position; }
};

//eventalign related
// Summarize the event alignment for a read strand
typedef struct 
{
    // //cleanup this part    
    // EventalignSummary() {
    //     num_events = 0;
    //     num_steps = 0;
    //     num_stays = 0;
    //     num_skips = 0;
    //     sum_z_score = 0;
    //     sum_duration = 0;
    //     alignment_edit_distance = 0;
    //     reference_span = 0;
    // }

    int num_events;
    int num_steps;
    int num_stays;
    int num_skips;

    double sum_duration;
    double sum_z_score;
    int alignment_edit_distance;
    int reference_span;
}EventalignSummary;


// a data batch (dynamic data based on the reads)
typedef struct {
    // region string
    //char* region;

    // bam records
    bam1_t** bam_rec;
    int32_t capacity_bam_rec; // will these overflow?
    int32_t n_bam_rec;

    // fasta cache //can optimise later by caching a common string for all
    // records in the batch
    char** fasta_cache;

    //read sequence //todo : optimise by grabbing it from bam seq. is it possible due to clipping?
    char** read;
    int32_t* read_len;

    // fast5 file //should flatten this to reduce mallocs
    fast5_t** f5;

    //event table
    event_table* et;

    //scalings
    scalings_t* scalings;

    //aligned pairs
    AlignedPair** event_align_pairs;
    int32_t* n_event_align_pairs;

    //event alignments
    event_alignment_t** event_alignment;
    int32_t* n_event_alignment;
    double* events_per_base; //todo : do we need double?

    index_pair_t** base_to_event_map;

    int32_t* read_stat_flag;

    //extreme ugly hack till converted to C
    // An output map from reference positions to scored CpG sites
    std::map<int, ScoredSite> **site_score_map;

    //stats //set by the load_db
    int64_t sum_bases;
    int64_t total_reads; //total number mapped entries in the bam file (after filtering based on flags, mapq etc)
    int64_t bad_fast5_file; //empty fast5 path returned by readdb, could not open fast5
    int64_t ultra_long_skipped; //ultra long reads that are skipped

    //eventalign related
    EventalignSummary *eventalign_summary;
    //another extremely ugly hack till converted to C
    //TODO : convert this to a C array and get rid of include <vector>
    std::vector<event_alignment_t> **event_alignment_result;


} db_t;

//cuda core data structure : allocated array pointers
#ifdef HAVE_CUDA
    typedef struct{
    ptr_t* event_ptr_host;
    int32_t* n_events_host;
    ptr_t* read_ptr_host;
    int32_t* read_len_host;
    scalings_t* scalings_host;
    int32_t* n_event_align_pairs_host;

    char* read;        //flattened reads sequences
    ptr_t* read_ptr; //index pointer for flattedned "reads"
    int32_t* read_len;
    int64_t sum_read_len;
    int32_t* n_events;
    event_t* event_table;
    ptr_t* event_ptr;
    int64_t sum_n_events;
    scalings_t* scalings;
    AlignedPair* event_align_pairs;
    int32_t* n_event_align_pairs;
    float *bands;
    uint8_t *trace;
    EventKmerPair* band_lower_left;
    model_t* model_kmer_cache;
    model_t* model;

    //dynamic arrays
    uint64_t max_sum_read_len;
    uint64_t max_sum_n_events;


    } cuda_data_t;
#endif

//core data structure (mostly static data throughout the program lifetime)
typedef struct {
    // bam file related
    htsFile* m_bam_fh;
    hts_idx_t* m_bam_idx;
    bam_hdr_t* m_hdr;
    hts_itr_t* itr;

    //bam file for writing the skipped ultra long reads to be later processed
    htsFile* ultra_long_tmp;

    //temporary file for dumping
    FILE *raw_dump;

    // fa related
    faidx_t* fai;

    // readbb
    ReadDB* readbb;

    // models
    model_t* model; //dna model
    model_t* cpgmodel;

    // options
    opt_t opt;

    //realtime0
    double realtime0;

    double load_db_time;
    double process_db_time;

    //loading time breakdown
    double db_bam_time;
    double db_fasta_time;
    double db_fast5_time;
    double db_fast5_open_time;
    double db_fast5_read_time;

    //processing time break down
    double event_time;
    double align_time;
    double est_scale_time;
    double meth_time;


#ifdef HAVE_CUDA

    //cuda arrays
    cuda_data_t* cuda;

    double align_kernel_time;
    double align_pre_kernel_time;
    double align_core_kernel_time;
    double align_post_kernel_time;
    double extra_load_cpu;
    double align_cuda_malloc;
    double align_cuda_memcpy;
    double align_cuda_postprocess;
    double align_cuda_preprocess;
    double align_cuda_total_kernel;

    //perf stats (can reduce to 16 bit integers)
    int32_t previous_mem;
    int32_t previous_count_mem;
    int32_t previous_load;
    int32_t previous_count_load;

#endif

    //stats //set by output_db
    int64_t sum_bases;
    int64_t total_reads; //total number mapped entries in the bam file (after filtering based on flags, mapq etc)
    int64_t bad_fast5_file; //empty fast5 path returned by readdb, could not open fast5
    int64_t ultra_long_skipped; //ultra long reads that are skipped
    int64_t qc_fail_reads;
    int64_t failed_calibration_reads;
    int64_t failed_alignment_reads;

    //eventalign related
    int8_t mode;
    FILE *event_summary_fp;


} core_t;

//argument wrapper for the multithreading framework for processing
typedef struct {
    core_t* core;
    db_t* db;
    int32_t starti;
    int32_t endi;
    void (*func)(core_t*,db_t*,int);
#ifdef WORK_STEAL
    void *all_pthread_args;
#endif
#ifdef HAVE_CUDA
    int32_t *ultra_long_reads; //reads that are assigned to the CPU due to the unsuitability to process on the GPU
    double ret1;    //return value
#endif
} pthread_arg_t;

//argument wrapper for Input/processing/output interleave thread framework
typedef struct {
    core_t* core;
    db_t* db;
    //conditional variable for notifying the processing to the output threads
    pthread_cond_t cond;
    pthread_mutex_t mutex;
    int8_t finished;
} pthread_arg2_t;

//return status by the load_db - used for termination when all the data is processed
typedef struct {
    int32_t num_reads;
    int64_t num_bases;
} ret_status_t;


/* Function prototype for major functions */

db_t* init_db(core_t* core);
ret_status_t load_db(core_t* dg, db_t* db);
core_t* init_core(const char* bamfilename, const char* fastafile,
                  const char* fastqfile, const char* tmpfile, opt_t opt,double realtime0, int8_t mode);
void process_db(core_t* dg, db_t* db);
void pthread_db(core_t* core, db_t* db, void (*func)(core_t*,db_t*,int));
void align_db(core_t* core, db_t* db);
void align_single(core_t* core, db_t* db, int32_t i);
void output_db(core_t* core, db_t* db);
void free_core(core_t* core);
void free_db_tmp(db_t* db);
void free_db(db_t* db);
void init_opt(opt_t* opt);

#ifdef HAVE_CUDA
    void init_cuda(core_t* core);
    void free_cuda(core_t* core);

#endif

/* Function prototypes for other non-major functions are in f5cmisc.h (and f5cmisc.cuh for CUDA)*/

#endif
```
