## Supplementary material for "GPU Accelerated Adaptive Banded Event Alignment for Rapid Comparative Nanopore Signal Analysis": f5c-v0.2-beta-release: f5cmisc.h

```
#ifndef F5CMISC_H
#define F5CMISC_H

#include "error.h"
#include "f5c.h"
#include <sys/resource.h>
#include <sys/time.h>

#define MIN_CALIBRATION_VAR 2.5
#define MAX_EVENT_TO_BP_RATIO 20

// Flags to modify the behaviour of the HMM
enum HMMAlignmentFlags
{
    HAF_ALLOW_PRE_CLIP = 1, // allow events to go unmatched before the aligning region
    HAF_ALLOW_POST_CLIP = 2 // allow events to go unmatched after the aligning region
};

event_table getevents(size_t nsample, float* rawptr);
void read_model(model_t* model, const char* file);
void set_model(model_t* model);
void set_cpgmodel(model_t* model);
scalings_t estimate_scalings_using_mom(char* sequence, int32_t sequence_len,
                                       model_t* pore_model, event_table et);
int32_t align(AlignedPair* out_2, char* sequence, int32_t sequence_len,
              event_table events, model_t* models, scalings_t scaling,
              float sample_rate);
int32_t postalign(event_alignment_t* alignment, index_pair_t* base_to_event_map, double* events_per_base,
                  char* sequence, int32_t n_kmers, AlignedPair* event_alignment,
                  int32_t n_events);
bool recalibrate_model(model_t* pore_model, event_table et,
                       scalings_t* scallings,
                       const event_alignment_t* alignment_output,
                       int32_t num_alignments, bool scale_var);

float profile_hmm_score(const char *m_seq,const char *m_rc_seq, event_t* event, scalings_t scaling,  model_t* cpgmodel, uint32_t event_start_idx,
    uint32_t event_stop_idx,
    uint8_t strand,
    int8_t event_stride,
    uint8_t rc,double events_per_base,uint32_t hmm_flags
);
void calculate_methylation_for_read(std::map<int, ScoredSite>* site_score_map, char* ref, bam1_t* record, int32_t read_length, event_t* event, index_pair_t* base_to_event_map,
scalings_t scaling, model_t* cpgmodel,double events_per_base);

void emit_event_alignment_tsv(FILE* fp,
                              uint32_t strand_idx,
                              const event_table* et, model_t* model, scalings_t scalings,
                              const std::vector<event_alignment_t>& alignments, 
                              int8_t print_read_names, int8_t scale_events, int8_t write_samples,
                              char* read_name, char *ref_name);
void emit_event_alignment_tsv_header(FILE* fp, int8_t print_read_names, int8_t write_samples);

void realign_read(std::vector<event_alignment_t>* event_alignment_result, EventalignSummary *summary, FILE *summary_fp,char* ref,
                  const bam_hdr_t* hdr,
                  const bam1_t* record, int32_t read_length,
                  size_t read_idx,
                  //int region_start,
                  //int region_end, 
                  event_table* events, model_t* model,index_pair_t* base_to_event_map,scalings_t scaling,double events_per_base );

//basically the functions in nanopolish_profile_hmm_r9.*
float profile_hmm_score_r9(const char *m_seq,
                                const char *m_rc_seq,
                                event_t* event,
                                scalings_t scaling,
                                model_t* cpgmodel,
                                uint32_t event_start_idx,
                                uint32_t event_stop_idx,
                                uint8_t strand,
                                int8_t event_stride,
                                uint8_t rc,
                                double events_per_base,
                                uint32_t hmm_flags);


#ifdef HAVE_CUDA
void align_cuda(core_t* core, db_t* db);
#endif

// taken from minimap2/misc
static inline double realtime(void) {
    struct timeval tp;
    struct timezone tzp;
    gettimeofday(&tp, &tzp);
    return tp.tv_sec + tp.tv_usec * 1e-6;
}

// taken from minimap2/misc
static inline double cputime(void) {
    struct rusage r;
    getrusage(RUSAGE_SELF, &r);
    return r.ru_utime.tv_sec + r.ru_stime.tv_sec +
           1e-6 * (r.ru_utime.tv_usec + r.ru_stime.tv_usec);
}

//taken from minimap2
static inline long peakrss(void)
{
	struct rusage r;
	getrusage(RUSAGE_SELF, &r);
#ifdef __linux__
	return r.ru_maxrss * 1024;
#else
	return r.ru_maxrss;
#endif
}

// Prints to the provided buffer a nice number of bytes (KB, MB, GB, etc)
//from https://www.mbeckler.org/blog/?p=114
static inline void print_size(const char* name, uint64_t bytes)
{
    const char* suffixes[7];
    suffixes[0] = "B";
    suffixes[1] = "KB";
    suffixes[2] = "MB";
    suffixes[3] = "GB";
    suffixes[4] = "TB";
    suffixes[5] = "PB";
    suffixes[6] = "EB";
    uint64_t s = 0; // which suffix to use
    double count = bytes;
    while (count >= 1024 && s < 7)
    {
        s++;
        count /= 1024;
    }
    if (count - floor(count) == 0.0)
        fprintf(stderr, "[%s] %s : %d %s\n", __func__ , name, (int)count, suffixes[s]);
    else
        fprintf(stderr, "[%s] %s : %.1f %s\n", __func__, name, count, suffixes[s]);
}


#endif
```
