## Supplementary material for "GPU Accelerated Adaptive Banded Event Alignment for Rapid Comparative Nanopore Signal Analysis": f5c-v0.2-beta-release: fast5lite.h

```
#ifndef FAST5LITE_H
#define FAST5LITE_H

#ifndef HAVE_CONFIG_H
#    define HAVE_CONFIG_H
#    include "config.h"
#endif

#ifdef HAVE_HDF5_SERIAL_HDF5_H
#    include <hdf5/serial/hdf5.h>
#endif

#ifdef HAVE_HDF5_H
#    include <hdf5.h>
#endif

#ifdef HAVE_HDF5_HDF5_H
#    include <hdf5/hdf5.h>
#endif

#ifdef HAVE___HDF5_INCLUDE_HDF5_H
#    include <hdf5.h>
#endif

#include <errno.h>
#include <math.h>
#include <stdint.h>
#include <stdlib.h>
#include <string.h>

#include "error.h"

typedef struct {
    float* rawptr;   // raw signal (float is not the best datatype type though)
    hsize_t nsample; // number of samples

    //	Information for scaling raw data from ADC values to pA (are these duplicates?)
    float digitisation;
    float offset;
    float range;
    float sample_rate;

    // computed scaling paramtersd
    float scale;
    float shift;
    float drift;
    float var;
    float scale_sd;
    float var_sd;

    // derived parameters that are cached for efficiency. do we need these?
    float log_var;
    float scaled_var;
    float log_scaled_var;

} fast5_t;

// The following was adapted from Nanopolish : nanopolish_fast5_io.cpp

#define RAW_ROOT "/Raw/Reads/"

static inline hid_t fast5_open(char* filename) {
    hid_t hdf5file = H5Fopen(filename, H5F_ACC_RDONLY, H5P_DEFAULT);
    return hdf5file;
}

static inline void fast5_close(hid_t hdf5_file) { H5Fclose(hdf5_file); }

static inline float fast5_read_float_attribute(hid_t group,
                                               const char* attribute) {
    float val = NAN;

    hid_t attr = H5Aopen(group, attribute, H5P_DEFAULT);
    if (attr < 0) {
        WARNING("Failed to open attribute '%s'.", attribute);
        return NAN;
    }

    herr_t ret = H5Aread(attr, H5T_NATIVE_FLOAT, &val);
    if (ret < 0) {
        WARNING("Failed to read attribute '%s'.", attribute);
        return NAN;
    }

    H5Aclose(attr);

    return val;
}

static inline int32_t fast5_read(hid_t hdf5_file, fast5_t* f5) {
    f5->rawptr = NULL;
    hid_t space;
    hsize_t nsample;
    herr_t status;

    // retrieve the size of the read name
    ssize_t size = H5Lget_name_by_idx(hdf5_file, RAW_ROOT, H5_INDEX_NAME,
                                      H5_ITER_INC, 0, NULL, 0, H5P_DEFAULT);

    if (size < 0) {
        WARNING("Failed to retrieve the size of the read name.%s", "");
        return -1;
    }

    // copy the read name out of the fast5
    char* read_name = (char*)calloc(1 + size, sizeof(char));
    ssize_t ret =
        H5Lget_name_by_idx(hdf5_file, RAW_ROOT, H5_INDEX_NAME, H5_ITER_INC, 0,
                           read_name, 1 + size, H5P_DEFAULT);
    if (ret < 0) {
        WARNING("Failed to retrieve the read name.%s", "");
        return -1;
    }

    // not the most efficient and safest, but being a bit lazy for the moment
    char* signal_path = (char*)malloc(
        (size + 1 + strlen(RAW_ROOT) + strlen("/Signal")) * sizeof(char));
    MALLOC_CHK(signal_path);

    sprintf(signal_path, "%s%s%s", RAW_ROOT, read_name, "/Signal");

#ifdef DEBUG_SIGNAL_PATH
    printf("Signal path : %s\n", signal_path);
#endif
    free(read_name);

    hid_t dset = H5Dopen(hdf5_file, signal_path, H5P_DEFAULT);
    if (dset < 0) {
        WARNING("Failed to open dataset '%s' to read raw signal.", signal_path);
        free(signal_path);
        return -1;
        // goto cleanup2;
    }

    space = H5Dget_space(dset);
    if (space < 0) {
        WARNING("Failed to create copy of dataspace for raw signal %s.",
                signal_path);
        H5Dclose(dset);
        free(signal_path);
        return -1;
        // goto cleanup3;
    }

    int32_t ret1 = H5Sget_simple_extent_dims(space, &nsample, NULL);
    if (ret1 < 0) {
        WARNING("Failed to get the dataspace dimension for raw signal %s.",
                signal_path);
        H5Sclose(space);
        H5Dclose(dset);
        free(signal_path);
        return -1;
    }

    f5->nsample = nsample;
    f5->rawptr = (float*)calloc(nsample, sizeof(float));
    status = H5Dread(dset, H5T_NATIVE_FLOAT, H5S_ALL, H5S_ALL, H5P_DEFAULT,
                     f5->rawptr);

    if (status < 0) {
        free(f5->rawptr);
        WARNING("Failed to read raw data from dataset %s.", signal_path);
        H5Sclose(space);
        H5Dclose(dset);
        free(signal_path);
        return -1;
    }

    H5Sclose(space);
    H5Dclose(dset);

    // get channel parameters
    const char* scaling_path = "/UniqueGlobalKey/channel_id";

    hid_t scaling_group = H5Gopen(hdf5_file, scaling_path, H5P_DEFAULT);
    if (scaling_group < 0) {
        WARNING("Failed to open group %s.", scaling_path);
        free(signal_path);
        return -1;
    }

    f5->digitisation =
        fast5_read_float_attribute(scaling_group, "digitisation");
    f5->offset = fast5_read_float_attribute(scaling_group, "offset");
    f5->range = fast5_read_float_attribute(scaling_group, "range");
    f5->sample_rate =
        fast5_read_float_attribute(scaling_group, "sampling_rate");

    if (f5->digitisation == NAN || f5->offset == NAN || f5->range == NAN ||
        f5->sample_rate == NAN) {
        WARNING("Read a NAN value for scaling parameters for '%s'.",
                signal_path);
        H5Gclose(scaling_group);
        free(signal_path);
        return -1;
    }

    H5Gclose(scaling_group);
    free(signal_path);

    return 0;
}


//todo : make to C (the following are used by the indexer)
#include <string>
static inline std::string fast5_get_fixed_string_attribute(hid_t hdf5_file, const std::string& group_name, const std::string& attribute_name)
{
    size_t storage_size;
    char* buffer;
    hid_t group, attribute, attribute_type;
    int ret;
    std::string out;

    // according to http://hdf-forum.184993.n3.nabble.com/check-if-dataset-exists-td194725.html
    // we should use H5Lexists to check for the existence of a group/dataset using an arbitrary path
    ret = H5Lexists(hdf5_file, group_name.c_str(), H5P_DEFAULT);
    if(ret <= 0) {
        return "";
    }

    // Open the group /Raw/Reads/Read_nnn
    group = H5Gopen(hdf5_file, group_name.c_str(), H5P_DEFAULT);
    if(group < 0) {
#ifdef DEBUG_FAST5_IO
        fprintf(stderr, "could not open group %s\n", group_name.c_str());
#endif
        goto close_group;
    }

    // Ensure attribute exists
    ret = H5Aexists(group, attribute_name.c_str());
    if(ret <= 0) {
        goto close_group;
    }

    // Open the attribute
    attribute = H5Aopen(group, attribute_name.c_str(), H5P_DEFAULT);
    if(attribute < 0) {
#ifdef DEBUG_FAST5_IO
        fprintf(stderr, "could not open attribute: %s\n", attribute_name.c_str());
#endif
        goto close_attr;
    }

    // Get data type and check it is a fixed-length string
    attribute_type = H5Aget_type(attribute);
    if(H5Tis_variable_str(attribute_type)) {
#ifdef DEBUG_FAST5_IO
        fprintf(stderr, "variable length string detected -- ignoring attribute\n");
#endif
        goto close_type;
    }

    // Get the storage size and allocate
    storage_size = H5Aget_storage_size(attribute);
    buffer = (char*)calloc(storage_size + 1, sizeof(char));

    // finally read the attribute
    ret = H5Aread(attribute, attribute_type, buffer);
    if(ret >= 0) {
        out = buffer;
    }

    // clean up
    free(buffer);
close_type:
    H5Tclose(attribute_type);
close_attr:
    H5Aclose(attribute);
close_group:
    H5Gclose(group);

    return out;
}


//from nanopolish_fast5_io.cpp
static inline std::string fast5_get_read_id(hid_t hdf5_file)
{

    std::string out = "";

    // Get the path to the raw read group
    // retrieve the size of the read name
    ssize_t size = H5Lget_name_by_idx(hdf5_file, RAW_ROOT, H5_INDEX_NAME,
                                      H5_ITER_INC, 0, NULL, 0, H5P_DEFAULT);

    if (size < 0) {
        WARNING("Failed to retrieve the size of the read name.%s", "");
        return out;
    }

    // copy the read name out of the fast5
    char* read_name = (char*)calloc(1 + size, sizeof(char));
    ssize_t ret =
        H5Lget_name_by_idx(hdf5_file, RAW_ROOT, H5_INDEX_NAME, H5_ITER_INC, 0,
                           read_name, 1 + size, H5P_DEFAULT);
    if (ret < 0) {
        WARNING("Failed to retrieve the read name.%s", "");
        return out;
    }

    std::string raw_read_group= std::string(RAW_ROOT) + read_name;
    free(read_name);


    return fast5_get_fixed_string_attribute(hdf5_file, raw_read_group, "read_id");
}

#endif
```
