## Supplementary material for "GPU Accelerated Adaptive Banded Event Alignment for Rapid Comparative Nanopore Signal Analysis": f5c-v0.2-beta-release: freq.c

```
/* @f5c
**
** calculate methylation frequency
** @author: Thomas Lam
** @@
******************************************************************************/

#include <stdlib.h>
#include <stdio.h>
#include <unistd.h>
#include <stdbool.h>
#include <math.h>
#include <string.h>
#include <getopt.h>
#include "khash.h"

#define KEY_SIZE 3
#define TSV_HEADER_LENGTH 160

#define ALLOC(type, size) (type *)safe_malloc((size) * sizeof(type))

// TODO : a number of inefficient mallocs are done in this code, which can be removed
// for example getline can use a onetime buffer
// need to check for empty newline chars
// todo : can improve peak memory by disk based sorting - futuristic next level

static const char usage[] = "Usage: %s [options...]\n"
                            "\n"
                            "  -c [float]        Call threshold. Default is 2.5.\n"
                            "  -i [file]         Input file. Read from stdin if not specified.\n"
                            "  -s                Split groups\n";

struct site_stats {
    int num_reads;
    int posterior_methylated;
    int called_sites;
    int called_sites_methylated;
    int group_size;
    char* sequence;
};

struct tsv_record {
    char* chromosome;
    int start;
    int end;
    double log_lik_ratio;
    int num_cpgs;
    char *sequence;
};

KHASH_MAP_INIT_STR(str, struct site_stats);

void* safe_malloc(size_t size) {
    void* p = malloc(size);
    if (!p) {
        perror("Cannot allocated memory");
        exit(EXIT_FAILURE);
    }
    return p;
}

/* Format chromosome, start and end to colon-delimited string. */
char* make_key(char* chromosome, int start, int end) {
    int start_strlen = snprintf(NULL, 0, "%d", start);
    int end_strlen = snprintf(NULL, 0, "%d", end);
    int key_strlen = strlen(chromosome) + start_strlen + end_strlen + 3;
    char* key = ALLOC(char, key_strlen);
    snprintf(key, key_strlen, "%s:%d:%d", chromosome, start, end);
    return key;
}

/* Split colon-delimited keys */
char** split_key(char* key, int size) {
    char** tok = ALLOC(char*, size);
    char* cpy = strdup(key);
    char* cpy_start = cpy;

    char* t = strtok(cpy, ":");
    if (t) {
        tok[0] = strdup(t);
    }

    for (int i = 1; i < size; i++) {
        char* t = strtok(NULL, ":");
        if (t) {
            tok[i] = strdup(t);
        }
    }

    free(cpy_start);

    return tok;
}

int cmp_key(const void* a, const void* b) {
    char* key_a = *(char**)a;
    char* key_b = *(char**)b;

    char** toks_a = split_key(key_a, KEY_SIZE);
    char** toks_b = split_key(key_b, KEY_SIZE);

    int chromosome_neq = strcmp(toks_a[0], toks_b[0]);

    if (chromosome_neq) {
        for (int i = 0; i < KEY_SIZE; i++) {
            free(toks_a[i]);
            free(toks_b[i]);
        }
        free(toks_a);
        free(toks_b);
        return chromosome_neq;
    }

    int start_a = atoi(toks_a[1]);
    int start_b = atoi(toks_b[1]);
    int end_a = atoi(toks_a[2]);
    int end_b = atoi(toks_b[2]);

    for (int i = 0; i < KEY_SIZE; i++) {
        free(toks_a[i]);
        free(toks_b[i]);
    }
    free(toks_a);
    free(toks_b);

    if (start_a == start_b) {
        return end_a - end_b;
    }

    return start_a - start_b;
}

void update_call_stats(khash_t(str)* sites, char* key, int num_called_cpg_sites,
        bool is_methylated, char* sequence) {
    struct site_stats ss = {
        .num_reads = 0,
        .posterior_methylated = 0,
        .called_sites = 0,
        .called_sites_methylated = 0,
        .group_size = num_called_cpg_sites,
        .sequence = strdup(sequence)
    };

    int absent;
    khint_t k = kh_put(str, sites, key, &absent);

    if (absent == -1) {
        fprintf(stderr, "Failed to insert key: %s\n", key);
        exit(EXIT_FAILURE);
    } else if (absent > 0) {
        kh_key(sites,k)=key;
        kh_value(sites, k) = ss;
    }
    else{
        free(ss.sequence);
        free(key);
    }

    kh_value(sites, k).num_reads++;
    kh_value(sites, k).called_sites += num_called_cpg_sites;

    if (is_methylated) {
        kh_value(sites, k).called_sites_methylated += num_called_cpg_sites;
    }
}

struct tsv_record* get_tsv_line(FILE* fp) {
    struct tsv_record* record = ALLOC(struct tsv_record, 1);
    char* buf = NULL;
    size_t buf_size = 0;

    if (getline(&buf, &buf_size, fp) == -1) {
        free(record);
        if(buf_size>0){
            free(buf);
        }
        return NULL;

    }

    record->chromosome = strdup(strtok(buf, "\t"));
    record->start = atoi(strtok(NULL, "\t"));
    record->end = atoi(strtok(NULL, "\t"));
    strtok(NULL, "\t");
    record->log_lik_ratio = strtod(strtok(NULL, "\t"), NULL);
    strtok(NULL, "\t");
    strtok(NULL, "\t");
    strtok(NULL, "\t");
    record->num_cpgs = atoi(strtok(NULL, "\t"));
    record->sequence = strdup(strtok(NULL, "\t\n"));

    free(buf);

    return record;
}

int freq_main(int argc, char **argv) {
    FILE* input = stdin;
    double call_threshold = 2.5;
    bool split_groups = false;

    int c;

    //extern char* optarg;
    //extern int optind, optopt;

    while ((c = getopt(argc, argv, "c:i:s")) != -1) {
        switch(c) {
            case 'c':
                /* TODO handle overflow when calling strtod */
                call_threshold = strtod(optarg, NULL);
                break;
            case 'i': {
                          FILE* in = fopen(optarg, "r");
                          if (in == NULL) {
                              perror("Failed to open file.");
                              exit(EXIT_FAILURE);
                          }
                          input = in;
                          break;
                      }
            case 's':
                      split_groups = true;
                      break;
            case ':':
                      fprintf(stderr, "Option -%c requires an operand\n", optopt);
                      fprintf(stderr, usage, argv[0]);
                      exit(EXIT_FAILURE);
            case '?':
                      fprintf(stderr, "Unrecognized option: -%c\n", optopt);
                      fprintf(stderr, usage, argv[0]);
                      exit(EXIT_FAILURE);
            default:
                      break;
        }
    }

    khash_t(str)* sites = kh_init(str);
    struct tsv_record* record;
    /* ignore header */
    char tmp[TSV_HEADER_LENGTH];
    char *ret=fgets(tmp, TSV_HEADER_LENGTH, input);
    if(ret==NULL){
        fprintf(stderr,"Bad file format with no header?\n");
        exit(1);
    }

    while ((record = get_tsv_line(input)) != NULL) {
        int num_sites = record->num_cpgs;
        double llr = record->log_lik_ratio;

        if (fabs(llr) < call_threshold) {
            free(record->sequence);
            free(record->chromosome);
            free(record);
            continue;
        }

        char* sequence = record->sequence;
        bool is_methylated = llr > 0;

        if (split_groups && num_sites > 1) {
            char* c = record->chromosome;
            int s = record->start;
            /* TODO following variable is unused in original Python script */
            /* int e = records[i].end; */
            char* substring = strstr(sequence, "CG");
            /* str.find() is equivalent to the offset of the needle pointer location */
            /* and the haystack pointer location */
            long cg_pos = substring == NULL ? -1 : substring - sequence;
            long first_cg_pos = cg_pos;

            while (cg_pos != -1) {
                char* key = make_key(c, s + cg_pos - first_cg_pos, s + cg_pos - first_cg_pos);
                const char* sg = "split-group";
                update_call_stats(sites, key, 1, is_methylated, (char*)sg);
                char* substring = strstr(sequence + cg_pos + 1, "CG");
                cg_pos = substring == NULL ? -1 : substring - sequence;
            }
        } else {
            char* key = make_key(record->chromosome, record->start, record->end);
            update_call_stats(sites, key, num_sites, is_methylated, sequence);

        }
        free(record->sequence);
        free(record->chromosome);
        free(record);
    }

    printf("chromosome\tstart\tend\tnum_cpgs_in_group\tcalled_sites\tcalled_sites_methylated\tmethylated_frequency\tgroup_sequence\n");

    char** sorted_keys = ALLOC(char*, kh_size(sites));
    int size = 0;

    for (khint_t k = kh_begin(sites); k != kh_end(sites); k++) {
        if (kh_exist(sites, k)) {
            sorted_keys[size++] = (char*)kh_key(sites, k);
        }
    }

    qsort(sorted_keys, size, sizeof(char*), cmp_key);

    for (int i = 0; i < size; i++) {
        khint_t k = kh_get(str, sites, sorted_keys[i]);
        if (kh_value(sites, k).called_sites > 0) {
            char** toks = split_key((char*)kh_key(sites, k), KEY_SIZE);
            char* c = toks[0];
            char* s = toks[1];
            char* e = toks[2];
            struct site_stats site = kh_value(sites, k);
            double f = (double)site.called_sites_methylated / site.called_sites;
            printf("%s\t%s\t%s\t%d\t%d\t%d\t%.3lf\t%s\n", c, s, e, site.group_size, site.called_sites, site.called_sites_methylated, f, site.sequence);
            free(site.sequence);
            free(c);
            free(s);
            free(e);
            free(toks);
            //free((char*)kh_key(sites, k));
        }
    }


    for (khint_t k = kh_begin(sites); k != kh_end(sites); k++) {
        if (kh_exist(sites, k)) {
            free((char*)kh_key(sites, k));
        }
    }

    fclose(input);
    kh_destroy(str, sites);
    free(sorted_keys);
    return 0;
}
```
