## Supplementary material for "GPU Accelerated Adaptive Banded Event Alignment for Rapid Comparative Nanopore Signal Analysis": f5c-v0.2-beta-release: hmm.c

```
#include "f5c.h"
#include "f5cmisc.h"
#include "matrix.h"
#include <math.h>
#include <assert.h>
#include <vector>
#include "logsum.h"


//#define INPUT_DEBUG 1
#define TRANS_START_TO_CLIP 0.5
#define TRANS_CLIP_SELF 0.9

#define TRUE 1
#define FALSE 0

//contains extracted code from nanopolish hmm and matrix


//todo : can make more efficient using bit encoding
static inline uint32_t get_rank(char base) {
    if (base == 'A') { //todo: do we neeed simple alpha?
        return 0;
    } else if (base == 'C') {
        return 1;
    } else if (base == 'G') {
        return 2;
    } else if (base == 'M') {
        return 3;
    } else if (base == 'T') {
        return 4;
    } else {
        WARNING("A None ACGMT base found : %c", base);
        return 0;
    }
}

// return the lexicographic rank of the kmer amongst all strings of
// length k for this alphabet
static inline uint32_t get_kmer_rank(const char* str, uint32_t k) {
    uint32_t p = 1;
    uint32_t r = 0;

    // from last base to first
    for (uint32_t i = 0; i < k; ++i) {
        //r += rank(str[k - i - 1]) * p;
        //p *= size();
        r += get_rank(str[k - i - 1]) * p;
        p *= 5;
    }
    return r;
}


static inline float log_normal_pdf(float x, float gp_mean, float gp_stdv,
                                   float gp_log_stdv) {
    /*INCOMPLETE*/
    float log_inv_sqrt_2pi = -0.918938f; // Natural logarithm
    float a = (x - gp_mean) / gp_stdv;
    return log_inv_sqrt_2pi - gp_log_stdv + (-0.5f * a * a);
    // return 1;
}

static inline float log_probability_match_r9(scalings_t scaling,
                                             model_t* models,
                                             event_t* event, int event_idx,
                                             uint32_t kmer_rank, uint8_t strand,
                                             float sample_rate) {
    // event level mean, scaled with the drift value
    strand = 0;
    assert(kmer_rank < 15625);
    //float level = read.get_drift_scaled_level(event_idx, strand);

    //float time =
    //    (events.event[event_idx].start - events.event[0].start) / sample_rate;
    float unscaledLevel = event[event_idx].mean;
    float scaledLevel = unscaledLevel;
    //float scaledLevel = unscaledLevel - time * scaling.shift;

    //fprintf(stderr, "level %f\n",scaledLevel);
    //GaussianParameters gp = read.get_scaled_gaussian_from_pore_model_state(pore_model, strand, kmer_rank);
    float gp_mean =
        scaling.scale * models[kmer_rank].level_mean + scaling.shift;
    float gp_stdv = models[kmer_rank].level_stdv * scaling.var;
    // float gp_stdv = 0;
    // float gp_log_stdv = models[kmer_rank].level_log_stdv + scaling.log_var;
    // if(models[kmer_rank].level_stdv <0.01 ){
    //  fprintf(stderr,"very small std dev %f\n",models[kmer_rank].level_stdv);
    // }
#ifdef CACHED_LOG
    float gp_log_stdv =
        models[kmer_rank].level_log_stdv + scaling.log_var;
#else
    float gp_log_stdv =
        log(models[kmer_rank].level_stdv) + log(scaling.var);
#endif

    float lp = log_normal_pdf(scaledLevel, gp_mean, gp_stdv, gp_log_stdv);
    return lp;
}


//following Code is from Nanopolish HMM

enum ProfileStateR9
{
    PSR9_KMER_SKIP = 0,
    PSR9_BAD_EVENT,
    PSR9_MATCH,
    PSR9_NUM_STATES = 3,
    PSR9_PRE_SOFT // intentionally after PS_NUM_STATES
};

enum HMMMovementType
{
    HMT_FROM_SAME_M = 0,
    HMT_FROM_PREV_M,
    HMT_FROM_SAME_B,
    HMT_FROM_PREV_B,
    HMT_FROM_PREV_K,
    HMT_FROM_SOFT,
    HMT_NUM_MOVEMENT_TYPES
};
typedef struct { float x[HMT_NUM_MOVEMENT_TYPES]; } HMMUpdateScores;


//all the blocks(functions, structures) are added in reverse order. at bottom is the first block called and on top is the latest block called.

// Allocate a vector with the model probabilities of skipping the remaining
// events after the alignment of event i
inline std::vector<float> make_post_flanking(const uint32_t e_start,
                                             const uint32_t num_events,
                                             int8_t event_stride,
                                             uint32_t event_stop_idx)
{
    // post_flank[i] means that the i-th event was the last one
    // aligned and the remainder should be emitted from the background model
    std::vector<float> post_flank(num_events, 0.0f);

    // base case, all events aligned
    post_flank[num_events - 1] = log(1 - TRANS_START_TO_CLIP);

    if(num_events > 1) {
        // base case, all events aligned but 1
        {
            uint32_t event_idx = e_start + (num_events - 1) * event_stride; // last event
            assert(event_idx == event_stop_idx);
            // post_flank[num_events - 2] = log(TRANS_START_TO_CLIP) + // transition from pre to background state
            //                              log_probability_background(*data.read, event_idx, data.strand) + // emit from background
            //                              log(1 - TRANS_CLIP_SELF); // transition to silent pre state
            post_flank[num_events - 2] = log(TRANS_START_TO_CLIP) + // transition from pre to background state
                                         -3.0f + // emit from background
                                         log(1 - TRANS_CLIP_SELF); // transition to silent pre state
        }

        for(int i = num_events - 3; i >= 0; --i) {
            //uint32_t event_idx = e_start + (i + 1) * event_stride;
            // post_flank[i] = log(TRANS_CLIP_SELF) +
            //                 log_probability_background(*data.read, event_idx, data.strand) + // emit from background
            //                 post_flank[i + 1]; // this accounts for the transition from start, and to silent pre
            post_flank[i] = log(TRANS_CLIP_SELF) +
                            -3.0f + // emit from background
                            post_flank[i + 1]; // this accounts for the transition from start, and to silent pre
        }
    }
    return post_flank;
}


// Allocate a vector with the model probabilities of skipping the first i events
inline std::vector<float> make_pre_flanking(const uint32_t e_start,
                                            const uint32_t num_events,
                                            int8_t event_stride)
{
    std::vector<float> pre_flank(num_events + 1, 0.0f);

    // base cases

    // no skipping
    pre_flank[0] = log(1 - TRANS_START_TO_CLIP);

    // skipping the first event
    // this includes the transition probability into and out of the skip state
    // pre_flank[1] = log(TRANS_START_TO_CLIP) + // transition from start to the background state
    //                log_probability_background(*data.read, e_start, data.strand) + // emit from background
    //                log(1 - TRANS_CLIP_SELF); // transition to silent pre state
    pre_flank[1] = log(TRANS_START_TO_CLIP) + // transition from start to the background state
                   -3.0f + // emit from background
                   log(1 - TRANS_CLIP_SELF); // transition to silent pre state

    // skip the remaining events
    for(size_t i = 2; i < pre_flank.size(); ++i) {
        //uint32_t event_idx = e_start + (i - 1) * event_stride;
        // pre_flank[i] = log(TRANS_CLIP_SELF) +
        //                log_probability_background(*data.read, event_idx, data.strand) + // emit from background
        //                pre_flank[i - 1]; // this accounts for the transition from the start & to the silent pre
        pre_flank[i] = log(TRANS_CLIP_SELF) +
                       -3.0f + // emit from background
                       pre_flank[i - 1]; // this accounts for the transition from the start & to the silent pre

    }

    return pre_flank;
}


// Pre-computed transitions from the previous block
// into the current block of states. Log-scaled.
struct BlockTransitions
{
    // Transition from m state (match event to k-mer)
    float lp_mm_self;
    float lp_mb;
    float lp_mk;
    float lp_mm_next;

    // Transitions from b state (bad event that should be ignored)
    float lp_bb;
    float lp_bk;
    float lp_bm_next; // movement to next k-mer
    float lp_bm_self; // movement to k-mer that we came from

    // Transitions from k state (no observation from k-mer)
    float lp_kk;
    float lp_km;
};

inline std::vector<BlockTransitions> calculate_transitions(uint32_t num_kmers, const char *m_seq,
                                                                                const char *m_rc_seq,
                                                                                event_t* event,
                                                                                scalings_t scaling,
                                                                                model_t* cpgmodel,
                                                                                uint32_t event_start_idx,
                                                                                uint32_t event_stop_idx,
                                                                                uint8_t strand,
                                                                                int8_t event_stride,
                                                                                uint8_t rc,
                                                                                double events_per_base)
{
    std::vector<BlockTransitions> transitions(num_kmers);

    //double read_events_per_base = data.read->events_per_base[data.strand];
    double read_events_per_base = events_per_base;
    for(uint32_t ki = 0; ki < num_kmers; ++ki) {

        // probability of skipping k_i from k_(i - 1)
        //float p_stay = 0.4;
        float p_stay = 1 - (1 / read_events_per_base);
#ifndef USE_EXTERNAL_PARAMS
        float p_skip = 0.0025;
        float p_bad = 0.001;
        float p_bad_self = p_bad;
        float p_skip_self = 0.3;
#else
        extern float g_p_skip, g_p_skip_self, g_p_bad, g_p_bad_self;
        float p_skip = g_p_skip;
        float p_skip_self = g_p_skip_self;
        float p_bad = g_p_bad;
        float p_bad_self = g_p_bad_self;
#endif
        // transitions from match state in previous block
        float p_mk = p_skip; // probability of not observing an event at all
        float p_mb = p_bad; // probabilty of observing a bad event
        float p_mm_self = p_stay; // probability of observing additional events from this k-mer
        float p_mm_next = 1.0f - p_mm_self - p_mk - p_mb; // normal movement from state to state

        // transitions from event split state in previous block
        float p_bb = p_bad_self;
        float p_bk, p_bm_next, p_bm_self;
        p_bk = p_bm_next = p_bm_self = (1.0f - p_bb) / 3;

        // transitions from kmer skip state in previous block
        float p_kk = p_skip_self;
        float p_km = 1.0f - p_kk;
        // p_kb not needed, equivalent to B->K

        // log-transform and store
        BlockTransitions& bt = transitions[ki];

        bt.lp_mk = log(p_mk);
        bt.lp_mb = log(p_mb);
        bt.lp_mm_self = log(p_mm_self);
        bt.lp_mm_next = log(p_mm_next);

        bt.lp_bb = log(p_bb);
        bt.lp_bk = log(p_bk);
        bt.lp_bm_next = log(p_bm_next);
        bt.lp_bm_self = log(p_bm_self);

        bt.lp_kk = log(p_kk);
        bt.lp_km = log(p_km);
    }

    return transitions;
}


// This function fills in a matrix with the result of running the HMM.
// The templated ProfileHMMOutput class allows one to run either Viterbi
// or the Forward algorithm.
template<class ProfileHMMOutput>
inline float profile_hmm_fill_generic_r9(const char *m_seq,
                                         const char *m_rc_seq,
                                        event_t* event,
                                        scalings_t scaling,
                                        model_t* cpgmodel,
                                        uint32_t event_start_idx,
                                        uint32_t event_stop_idx,
                                        uint8_t strand,
                                        int8_t event_stride,
                                        uint8_t rc,
                                        const uint32_t e_start,
                                        double events_per_base,
                                        uint32_t hmm_flags,
                                        ProfileHMMOutput& output){
    // PROFILE_FUNC("profile_hmm_fill_generic")
    // HMMInputSequence sequence = _sequence;
    // HMMInputData data = _data;
    assert( (rc && event_stride == -1) || (!rc && event_stride == 1));

#if HMM_REVERSE_FIX
    if(event_stride == -1) {
        // sequence.swap();
        char *temp = m_seq;
        m_seq = m_rc_seq;
        m_rc_seq = temp;
        uint32_t tmp = event_stop_idx;
        event_stop_idx = event_start_idx;
        event_start_idx = tmp;
        event_stride = 1;
        rc = false;
    }
#endif

    //e_start = event_start_idx;

    // Calculate number of blocks
    // A block of the HMM is a set of states for one kmer
    uint32_t num_blocks = output.get_num_columns() / PSR9_NUM_STATES;
    //fprintf(stderr,"%d %d\n",output.get_num_columns(),PSR9_NUM_STATES);
    uint32_t last_event_row_idx = output.get_num_rows() - 1;

    // Precompute the transition probabilites for each kmer block
    uint32_t num_kmers = num_blocks - 2; // two terminal blocks
    uint32_t last_kmer_idx = num_kmers - 1;


    std::vector<BlockTransitions> transitions = calculate_transitions(num_kmers,
                                                                        m_seq,
                                                                        m_rc_seq,
                                                                        event,
                                                                        scaling,
                                                                        cpgmodel,
                                                                        event_start_idx,
                                                                        event_stop_idx,
                                                                        strand,
                                                                        event_stride,
                                                                        rc,
                                                                        events_per_base);

    // Precompute kmer ranks
    // const uint32_t k = data.pore_model->k;
    //const uint32_t k = KMER_SIZE;
    // Make sure the HMMInputSequence's alphabet matches the state space of the read


    ////change start

    // assert( data.pore_model->states.size() == sequence.get_num_kmer_ranks(k) );

    std::vector<uint32_t> kmer_ranks(num_kmers);

    //todo : pre-calculate
    int32_t seq_len = strlen(m_seq);

    //check : this might be reverse cmplement kmer rnak
    for(size_t ki = 0; ki < num_kmers; ++ki){
        const char* substring = 0;
        if(rc==0){
            substring=m_seq+ki;
        }
        else{
            substring=m_rc_seq+seq_len-ki-KMER_SIZE;
        }

        // kmer_ranks[ki] = sequence.get_kmer_rank(ki, k, data.rc);
        kmer_ranks[ki] = get_kmer_rank(substring,KMER_SIZE);
    }


    ///change over

    size_t num_events = output.get_num_rows() - 1;

    std::vector<float> pre_flank = make_pre_flanking(e_start, num_events,event_stride);
    std::vector<float> post_flank = make_post_flanking(e_start, num_events,event_stride,event_stop_idx);

    // The model is currently constrainted to always transition
    // from the terminal/clipped state to the first kmer (and from the
    // last kmer to the terminal/clipping state so these are log(1.0).
    // They are kept as variables as it might be relaxed later.
    float lp_sm, lp_ms;
    lp_sm = lp_ms = 0.0f;

    // the penalty is controlled by the transition probability
    float BAD_EVENT_PENALTY = 0.0f;

    // Fill in matrix
    for(uint32_t row = 1; row < output.get_num_rows(); row++) {

        // Skip the first block which is the start state, it was initialized above
        // Similarily skip the last block, which is calculated in the terminate() function
        for(uint32_t block = 1; block < num_blocks - 1; block++) {

            // retrieve transitions
            uint32_t kmer_idx = block - 1;
            BlockTransitions& bt = transitions[kmer_idx];

            uint32_t prev_block = block - 1;
            uint32_t prev_block_offset = PSR9_NUM_STATES * prev_block;
            uint32_t curr_block_offset = PSR9_NUM_STATES * block;

            // Emission probabilities
            uint32_t event_idx = e_start + (row - 1) * event_stride;
            uint32_t rank = kmer_ranks[kmer_idx];
            // float lp_emission_m = log_probability_match_r9(*data.read, *data.pore_model, rank, event_idx, data.strand);
            float lp_emission_m =
                log_probability_match_r9(scaling, cpgmodel, event, event_idx,rank, strand, 0);
            //fprintf(stderr,"m_seq %s, event_idx %d, kmer_rank %d, log prob : %f\n",m_seq,event_idx,rank,lp_emission_m);
            //fprintf(stderr,"e_start %d, row %d, event_stride %d, block %d, num_block %d\n",e_start,row,event_stride,block,num_blocks);
            float lp_emission_b = BAD_EVENT_PENALTY;

            HMMUpdateScores scores;

            // state PSR9_MATCH
            scores.x[HMT_FROM_SAME_M] = bt.lp_mm_self + output.get(row - 1, curr_block_offset + PSR9_MATCH);
            scores.x[HMT_FROM_PREV_M] = bt.lp_mm_next + output.get(row - 1, prev_block_offset + PSR9_MATCH);
            scores.x[HMT_FROM_SAME_B] = bt.lp_bm_self + output.get(row - 1, curr_block_offset + PSR9_BAD_EVENT);
            scores.x[HMT_FROM_PREV_B] = bt.lp_bm_next + output.get(row - 1, prev_block_offset + PSR9_BAD_EVENT);
            scores.x[HMT_FROM_PREV_K] = bt.lp_km + output.get(row - 1, prev_block_offset + PSR9_KMER_SKIP);

            // m_s is the probability of going from the start state
            // to this kmer. The start state is (currently) only
            // allowed to go to the first kmer. If ALLOW_PRE_CLIP
            // is defined, we allow all events before this one to be skipped,
            // with a penalty;
            scores.x[HMT_FROM_SOFT] = (kmer_idx == 0 &&
                                        (event_idx == e_start ||
                                             (hmm_flags & HAF_ALLOW_PRE_CLIP))) ? lp_sm + pre_flank[row - 1] : -INFINITY;

            output.update_cell(row, curr_block_offset + PSR9_MATCH, scores, lp_emission_m);

            // state PSR9_BAD_EVENT
            scores.x[HMT_FROM_SAME_M] = bt.lp_mb + output.get(row - 1, curr_block_offset + PSR9_MATCH);
            scores.x[HMT_FROM_PREV_M] = -INFINITY; // not allowed
            scores.x[HMT_FROM_SAME_B] = bt.lp_bb + output.get(row - 1, curr_block_offset + PSR9_BAD_EVENT);
            scores.x[HMT_FROM_PREV_B] = -INFINITY;
            scores.x[HMT_FROM_PREV_K] = -INFINITY;
            scores.x[HMT_FROM_SOFT] = -INFINITY;
            output.update_cell(row, curr_block_offset + PSR9_BAD_EVENT, scores, lp_emission_b);

            // state PSR9_KMER_SKIP
            scores.x[HMT_FROM_SAME_M] = -INFINITY;
            scores.x[HMT_FROM_PREV_M] = bt.lp_mk + output.get(row, prev_block_offset + PSR9_MATCH);
            scores.x[HMT_FROM_SAME_B] = -INFINITY;
            scores.x[HMT_FROM_PREV_B] = bt.lp_bk + output.get(row, prev_block_offset + PSR9_BAD_EVENT);
            scores.x[HMT_FROM_PREV_K] = bt.lp_kk + output.get(row, prev_block_offset + PSR9_KMER_SKIP);
            scores.x[HMT_FROM_SOFT] = -INFINITY;
            output.update_cell(row, curr_block_offset + PSR9_KMER_SKIP, scores, 0.0f); // no emission

            // If POST_CLIP is enabled we allow the last kmer to transition directly
            // to the end after any event. Otherwise we only allow it from the
            // last kmer/event match.
            if(kmer_idx == last_kmer_idx && ( (hmm_flags & HAF_ALLOW_POST_CLIP) || row == last_event_row_idx)) {
                float lp1 = lp_ms + output.get(row, curr_block_offset + PSR9_MATCH) + post_flank[row - 1];
                float lp2 = lp_ms + output.get(row, curr_block_offset + PSR9_BAD_EVENT) + post_flank[row - 1];
                float lp3 = lp_ms + output.get(row, curr_block_offset + PSR9_KMER_SKIP) + post_flank[row - 1];

                output.update_end(lp1, row, curr_block_offset + PSR9_MATCH);
                output.update_end(lp2, row, curr_block_offset + PSR9_BAD_EVENT);
                output.update_end(lp3, row, curr_block_offset + PSR9_KMER_SKIP);
            }

#ifdef DEBUG_LOCAL_ALIGNMENT
            printf("[%d %d] start: %.2lf  pre: %.2lf fm: %.2lf\n", event_idx, kmer_idx, m_s + lp_emission_m, pre_flank[row - 1], output.get(row, curr_block_offset + PSR9_MATCH));
            printf("[%d %d]   end: %.2lf post: %.2lf\n", event_idx, kmer_idx, lp_end, post_flank[row - 1]);
#endif

#ifdef DEBUG_FILL
            printf("Row %u block %u\n", row, block);

            printf("\tPSR9_MATCH -- Transitions: [%.3lf %.3lf %.3lf %.3lf %.3lf] Prev: [%.2lf %.2lf %.2lf %.2lf %.2lf] out: %.2lf\n",
                    bt.lp_mm_self, bt.lp_mm_next, bt.lp_bm_self, bt.lp_bm_next, bt.lp_km,
                    output.get(row - 1, prev_block_offset + PSR9_MATCH),
                    output.get(row - 1, curr_block_offset + PSR9_MATCH),
                    output.get(row - 1, prev_block_offset + PSR9_BAD_EVENT),
                    output.get(row - 1, curr_block_offset + PSR9_BAD_EVENT),
                    output.get(row - 1, prev_block_offset + PSR9_KMER_SKIP),
                    output.get(row, curr_block_offset + PSR9_MATCH));
            printf("\tPSR9_BAD_EVENT -- Transitions: [%.3lf %.3lf] Prev: [%.2lf %.2lf] out: %.2lf\n",
                    bt.lp_mb, bt.lp_bb,
                    output.get(row - 1, curr_block_offset + PSR9_MATCH),
                    output.get(row - 1, curr_block_offset + PSR9_BAD_EVENT),
                    output.get(row, curr_block_offset + PSR9_BAD_EVENT));

            printf("\tPSR9_KMER_SKIP -- Transitions: [%.3lf %.3lf %.3lf] Prev: [%.2lf %.2lf %.2lf] sum: %.2lf\n",
                    bt.lp_mk, bt.lp_bk, bt.lp_kk,
                    output.get(row, prev_block_offset + PSR9_MATCH),
                    output.get(row, prev_block_offset + PSR9_BAD_EVENT),
                    output.get(row, prev_block_offset + PSR9_KMER_SKIP),
                    output.get(row, curr_block_offset + PSR9_KMER_SKIP));

            printf("\tEMISSION: %.2lf %.2lf\n", lp_emission_m, lp_emission_b);
#endif
        }
    }

    return output.get_end();
}


// Add the log-scaled values a and b using a transform to avoid precision errors
inline double add_logs(const double a, const double b)
{
#if ESL_LOG_SUM
    return p7_FLogsum(a, b);
#else
    if(a == -INFINITY && b == -INFINITY)
        return -INFINITY;

    if(a > b) {
        double diff = b - a;
        return a + log(1.0 + exp(diff));
    } else {
        double diff = a - b;
        return b + log(1.0 + exp(diff));
    }
#endif
}


class ProfileHMMForwardOutputR9
{
    public:
        ProfileHMMForwardOutputR9(FloatMatrix* p) : p_fm(p), lp_end(-INFINITY) {
		//p7_FLogsumInit();
        //commented by hasindu
	}

        //
        inline void update_cell(uint32_t row, uint32_t col, const HMMUpdateScores& scores, float lp_emission)
        {
            float sum = scores.x[0];
            for(auto i = 1; i < HMT_NUM_MOVEMENT_TYPES; ++i) {
                sum = add_logs(sum, scores.x[i]);
            }
            sum += lp_emission;
            set(*p_fm, row, col, sum);
        }

        // add in the probability of ending the alignment at row,col
        inline void update_end(float v, uint32_t, uint32_t)
        {
            lp_end = add_logs(lp_end, v);
        }

        // get the log probability stored at a particular row/column
        inline float get(uint32_t row, uint32_t col) const
        {
            return ::get(*p_fm, row, col);
        }

        // get the log probability for the end state
        inline float get_end() const
        {
            return lp_end;
        }

        inline size_t get_num_columns() const
        {
            return p_fm->n_cols;
        }

        inline size_t get_num_rows() const
        {
            return p_fm->n_rows;
        }

    private:
        ProfileHMMForwardOutputR9(); // not allowed
        FloatMatrix* p_fm;
        float lp_end;
};


void profile_hmm_forward_initialize_r9(FloatMatrix& fm)
{
    // initialize forward calculation
    for(uint32_t si = 0; si < fm.n_cols; si++) {
        set(fm, 0, si, -INFINITY);
    }

    for(uint32_t ri = 0; ri < fm.n_rows; ri++) {
        set(fm, ri, PSR9_KMER_SKIP, -INFINITY);
        set(fm, ri, PSR9_BAD_EVENT, -INFINITY);
        set(fm, ri, PSR9_MATCH, -INFINITY);
    }
}


float profile_hmm_score_r9(const char *m_seq,
                                const char *m_rc_seq,
                                event_t* event,
                                scalings_t scaling,
                                model_t* cpgmodel,
                                uint32_t event_start_idx,
                                uint32_t event_stop_idx,
                                uint8_t strand,
                                int8_t event_stride,
                                uint8_t rc,
                                double events_per_base,
                                uint32_t hmm_flags)
{
    const uint32_t k = KMER_SIZE; //hardcoded had this const uint32_t k = data.pore_model->k;
    uint32_t n_kmers = strlen(m_seq) - k + 1;

    uint32_t n_states = PSR9_NUM_STATES * (n_kmers + 2); // + 2 for explicit terminal states

    uint32_t e_start = event_start_idx;
    uint32_t e_end = event_stop_idx;
    uint32_t n_events = 0;
    if(e_end > e_start)
        n_events = e_end - e_start + 1;
    else
        n_events = e_start - e_end + 1;

    uint32_t n_rows = n_events + 1;

    // Allocate a matrix to hold the HMM result
    FloatMatrix fm;
    allocate_matrix(fm, n_rows, n_states);

    profile_hmm_forward_initialize_r9(fm);

    ProfileHMMForwardOutputR9 output(&fm);

    float score = profile_hmm_fill_generic_r9(m_seq,
                                                m_rc_seq,
                                                event,
                                                scaling,
                                                cpgmodel,
                                                event_start_idx,
                                                event_stop_idx,
                                                strand,
                                                event_stride,
                                                rc,
                                                e_start,
                                                events_per_base,
                                                hmm_flags,
                                                output);

    // cleanup
    free_matrix(fm);
    return score;
}


float profile_hmm_score(
	const char *m_seq,
	const char *m_rc_seq,
	event_t* event,
	scalings_t scaling,
	model_t* cpgmodel,
	uint32_t event_start_idx,
    uint32_t event_stop_idx,
    uint8_t strand,
    int8_t event_stride,
    uint8_t rc,
    double events_per_base,
    uint32_t hmm_flags
){
    #ifdef INPUT_DEBUG

        fprintf(stderr,"m_seq : %s\n",m_seq);
        fprintf(stderr,"m_rc_seq : %s\n",m_rc_seq);
        fprintf(stderr,"event_start_idx %d, event_stop_idx %d, event_stride %d, rc %d\n",event_start_idx,event_stop_idx,event_stride,rc);

    #endif


	return profile_hmm_score_r9(m_seq,
								m_rc_seq,
								event,
								scaling,
								cpgmodel,
								event_start_idx,
							    event_stop_idx,
							    strand,
							    event_stride,
							    rc,
							    events_per_base,
							    hmm_flags);


}
```
