## Supplementary material for "GPU Accelerated Adaptive Banded Event Alignment for Rapid Comparative Nanopore Signal Analysis": f5c-v0.2-beta-release: khash.h

```
/* The MIT License

   Copyright (c) 2008, 2009, 2011 by Attractive Chaos <>

   Permission is hereby granted, free of charge, to any person obtaining
   a copy of this software and associated documentation files (the
   "Software"), to deal in the Software without restriction, including
   without limitation the rights to use, copy, modify, merge, publish,
   distribute, sublicense, and/or sell copies of the Software, and to
   permit persons to whom the Software is furnished to do so, subject to
   the following conditions:

   The above copyright notice and this permission notice shall be
   included in all copies or substantial portions of the Software.

   THE SOFTWARE IS PROVIDED "AS IS", WITHOUT WARRANTY OF ANY KIND,
   EXPRESS OR IMPLIED, INCLUDING BUT NOT LIMITED TO THE WARRANTIES OF
   MERCHANTABILITY, FITNESS FOR A PARTICULAR PURPOSE AND
   NONINFRINGEMENT. IN NO EVENT SHALL THE AUTHORS OR COPYRIGHT HOLDERS
   BE LIABLE FOR ANY CLAIM, DAMAGES OR OTHER LIABILITY, WHETHER IN AN
   ACTION OF CONTRACT, TORT OR OTHERWISE, ARISING FROM, OUT OF OR IN
   CONNECTION WITH THE SOFTWARE OR THE USE OR OTHER DEALINGS IN THE
   SOFTWARE.
*/

/*
  An example:

#include "khash.h"
KHASH_MAP_INIT_INT(32, char)
int main() {
	int ret, is_missing;
	khiter_t k;
	khash_t(32) *h = kh_init(32);
	k = kh_put(32, h, 5, &ret);
	kh_value(h, k) = 10;
	k = kh_get(32, h, 10);
	is_missing = (k == kh_end(h));
	k = kh_get(32, h, 5);
	kh_del(32, h, k);
	for (k = kh_begin(h); k != kh_end(h); ++k)
		if (kh_exist(h, k)) kh_value(h, k) = 1;
	kh_destroy(32, h);
	return 0;
}
*/

/*
  2013-05-02 (0.2.8):

	* Use quadratic probing. When the capacity is power of 2, stepping function
	  i*(i+1)/2 guarantees to traverse each bucket. It is better than double
	  hashing on cache performance and is more robust than linear probing.

	  In theory, double hashing should be more robust than quadratic probing.
	  However, my implementation is probably not for large hash tables, because
	  the second hash function is closely tied to the first hash function,
	  which reduce the effectiveness of double hashing.

	Reference: http://research.cs.vt.edu/AVresearch/hashing/quadratic.php

  2011-12-29 (0.2.7):

    * Minor code clean up; no actual effect.

  2011-09-16 (0.2.6):

	* The capacity is a power of 2. This seems to dramatically improve the
	  speed for simple keys. Thank Zilong Tan for the suggestion. Reference:

	   - http://code.google.com/p/ulib/
	   - http://nothings.org/computer/judy/

	* Allow to optionally use linear probing which usually has better
	  performance for random input. Double hashing is still the default as it
	  is more robust to certain non-random input.

	* Added Wang's integer hash function (not used by default). This hash
	  function is more robust to certain non-random input.

  2011-02-14 (0.2.5):

    * Allow to declare global functions.

  2009-09-26 (0.2.4):

    * Improve portability

  2008-09-19 (0.2.3):

	* Corrected the example
	* Improved interfaces

  2008-09-11 (0.2.2):

	* Improved speed a little in kh_put()

  2008-09-10 (0.2.1):

	* Added kh_clear()
	* Fixed a compiling error

  2008-09-02 (0.2.0):

	* Changed to token concatenation which increases flexibility.

  2008-08-31 (0.1.2):

	* Fixed a bug in kh_get(), which has not been tested previously.

  2008-08-31 (0.1.1):

	* Added destructor
*/

#ifndef __AC_KHASH_H
#define __AC_KHASH_H

/*!
  @header

  Generic hash table library.
 */

#define AC_VERSION_KHASH_H "0.2.8"

#include <stdlib.h>
#include <string.h>
#include <limits.h>

/* compiler specific configuration */

#if UINT_MAX == 0xffffffffu
typedef unsigned int khint32_t;
#elif ULONG_MAX == 0xffffffffu
typedef unsigned long khint32_t;
#endif

#if ULONG_MAX == ULLONG_MAX
typedef unsigned long khint64_t;
#else
typedef unsigned long long khint64_t;
#endif

#ifndef kh_inline
#ifdef _MSC_VER
#define kh_inline __inline
#else
#define kh_inline inline
#endif
#endif /* kh_inline */

#ifndef klib_unused
#if (defined __clang__ && __clang_major__ >= 3) ||                             \
	(defined __GNUC__ && __GNUC__ >= 3)
#define klib_unused __attribute__((__unused__))
#else
#define klib_unused
#endif
#endif /* klib_unused */

typedef khint32_t khint_t;
typedef khint_t khiter_t;

#define __ac_isempty(flag, i) ((flag[i >> 4] >> ((i & 0xfU) << 1)) & 2)
#define __ac_isdel(flag, i) ((flag[i >> 4] >> ((i & 0xfU) << 1)) & 1)
#define __ac_iseither(flag, i) ((flag[i >> 4] >> ((i & 0xfU) << 1)) & 3)
#define __ac_set_isdel_false(flag, i)                                          \
	(flag[i >> 4] &= ~(1ul << ((i & 0xfU) << 1)))
#define __ac_set_isempty_false(flag, i)                                        \
	(flag[i >> 4] &= ~(2ul << ((i & 0xfU) << 1)))
#define __ac_set_isboth_false(flag, i)                                         \
	(flag[i >> 4] &= ~(3ul << ((i & 0xfU) << 1)))
#define __ac_set_isdel_true(flag, i) (flag[i >> 4] |= 1ul << ((i & 0xfU) << 1))

#define __ac_fsize(m) ((m) < 16 ? 1 : (m) >> 4)

#ifndef kroundup32
#define kroundup32(x)                                                          \
	(--(x), (x) |= (x) >> 1, (x) |= (x) >> 2, (x) |= (x) >> 4,             \
	 (x) |= (x) >> 8, (x) |= (x) >> 16, ++(x))
#endif

#ifndef kcalloc
#define kcalloc(N, Z) calloc(N, Z)
#endif
#ifndef kmalloc
#define kmalloc(Z) malloc(Z)
#endif
#ifndef krealloc
#define krealloc(P, Z) realloc(P, Z)
#endif
#ifndef kfree
#define kfree(P) free(P)
#endif

static const double __ac_HASH_UPPER = 0.77;

#define __KHASH_TYPE(name, khkey_t, khval_t)                                   \
	typedef struct kh_##name##_s {                                         \
		khint_t n_buckets, size, n_occupied, upper_bound;              \
		khint32_t *flags;                                              \
		khkey_t *keys;                                                 \
		khval_t *vals;                                                 \
	} kh_##name##_t;

#define __KHASH_PROTOTYPES(name, khkey_t, khval_t)                             \
	extern kh_##name##_t *kh_init_##name(void);                            \
	extern void kh_destroy_##name(kh_##name##_t *h);                       \
	extern void kh_clear_##name(kh_##name##_t *h);                         \
	extern khint_t kh_get_##name(const kh_##name##_t *h, khkey_t key);     \
	extern int kh_resize_##name(kh_##name##_t *h, khint_t new_n_buckets);  \
	extern khint_t kh_put_##name(kh_##name##_t *h, khkey_t key, int *ret); \
	extern void kh_del_##name(kh_##name##_t *h, khint_t x);

#define __KHASH_IMPL(name, SCOPE, khkey_t, khval_t, kh_is_map, __hash_func,                                              \
		     __hash_equal)                                                                                       \
	SCOPE kh_##name##_t *kh_init_##name(void)                                                                        \
	{                                                                                                                \
		return (kh_##name##_t *)kcalloc(1, sizeof(kh_##name##_t));                                               \
	}                                                                                                                \
	SCOPE void kh_destroy_##name(kh_##name##_t *h)                                                                   \
	{                                                                                                                \
		if (h) {                                                                                                 \
			kfree((void *)h->keys);                                                                          \
			kfree(h->flags);                                                                                 \
			kfree((void *)h->vals);                                                                          \
			kfree(h);                                                                                        \
		}                                                                                                        \
	}                                                                                                                \
	SCOPE void kh_clear_##name(kh_##name##_t *h)                                                                     \
	{                                                                                                                \
		if (h && h->flags) {                                                                                     \
			memset(h->flags, 0xaa,                                                                           \
			       __ac_fsize(h->n_buckets) * sizeof(khint32_t));                                            \
			h->size = h->n_occupied = 0;                                                                     \
		}                                                                                                        \
	}                                                                                                                \
	SCOPE khint_t kh_get_##name(const kh_##name##_t *h, khkey_t key)                                                 \
	{                                                                                                                \
		if (h->n_buckets) {                                                                                      \
			khint_t k, i, last, mask, step = 0;                                                              \
			mask = h->n_buckets - 1;                                                                         \
			k = __hash_func(key);                                                                            \
			i = k & mask;                                                                                    \
			last = i;                                                                                        \
			while (!__ac_isempty(h->flags, i) &&                                                             \
			       (__ac_isdel(h->flags, i) ||                                                               \
				!__hash_equal(h->keys[i], key))) {                                                       \
				i = (i + (++step)) & mask;                                                               \
				if (i == last)                                                                           \
					return h->n_buckets;                                                             \
			}                                                                                                \
			return __ac_iseither(h->flags, i) ? h->n_buckets : i;                                            \
		} else                                                                                                   \
			return 0;                                                                                        \
	}                                                                                                                \
	SCOPE int kh_resize_##name(kh_##name##_t *h, khint_t new_n_buckets)                                              \
	{ /* This function uses 0.25*n_buckets bytes of working space instead of [sizeof(key_t+val_t)+.25]*n_buckets. */ \
		khint32_t *new_flags = 0;                                                                                \
		khint_t j = 1;                                                                                           \
		{                                                                                                        \
			kroundup32(new_n_buckets);                                                                       \
			if (new_n_buckets < 4)                                                                           \
				new_n_buckets = 4;                                                                       \
			if (h->size >=                                                                                   \
			    (khint_t)(new_n_buckets * __ac_HASH_UPPER + 0.5))                                            \
				j = 0; /* requested size is too small */                                                 \
			else { /* hash table size to be changed (shrink or expand); rehash */                            \
				new_flags = (khint32_t *)kmalloc(                                                        \
					__ac_fsize(new_n_buckets) *                                                      \
					sizeof(khint32_t));                                                              \
				if (!new_flags)                                                                          \
					return -1;                                                                       \
				memset(new_flags, 0xaa,                                                                  \
				       __ac_fsize(new_n_buckets) *                                                       \
					       sizeof(khint32_t));                                                       \
				if (h->n_buckets <                                                                       \
				    new_n_buckets) { /* expand */                                                        \
					khkey_t *new_keys = (khkey_t *)krealloc(                                         \
						(void *)h->keys,                                                         \
						new_n_buckets *                                                          \
							sizeof(khkey_t));                                                \
					if (!new_keys) {                                                                 \
						kfree(new_flags);                                                        \
						return -1;                                                               \
					}                                                                                \
					h->keys = new_keys;                                                              \
					if (kh_is_map) {                                                                 \
						khval_t *new_vals = (khval_t *)krealloc(                                 \
							(void *)h->vals,                                                 \
							new_n_buckets *                                                  \
								sizeof(khval_t));                                        \
						if (!new_vals) {                                                         \
							kfree(new_flags);                                                \
							return -1;                                                       \
						}                                                                        \
						h->vals = new_vals;                                                      \
					}                                                                                \
				} /* otherwise shrink */                                                                 \
			}                                                                                                \
		}                                                                                                        \
		if (j) { /* rehashing is needed */                                                                       \
			for (j = 0; j != h->n_buckets; ++j) {                                                            \
				if (__ac_iseither(h->flags, j) == 0) {                                                   \
					khkey_t key = h->keys[j];                                                        \
					khval_t val;                                                                     \
					khint_t new_mask;                                                                \
					new_mask = new_n_buckets - 1;                                                    \
					if (kh_is_map)                                                                   \
						val = h->vals[j];                                                        \
					__ac_set_isdel_true(h->flags, j);                                                \
					while (1) { /* kick-out process; sort of like in Cuckoo hashing */               \
						khint_t k, i, step = 0;                                                  \
						k = __hash_func(key);                                                    \
						i = k & new_mask;                                                        \
						while (!__ac_isempty(                                                    \
							new_flags, i))                                                   \
							i = (i + (++step)) &                                             \
							    new_mask;                                                    \
						__ac_set_isempty_false(                                                  \
							new_flags, i);                                                   \
						if (i < h->n_buckets &&                                                  \
						    __ac_iseither(h->flags,                                              \
								  i) ==                                                  \
							    0) { /* kick out the existing element */                     \
							{                                                                \
								khkey_t tmp =                                            \
									h->keys[i];                                      \
								h->keys[i] =                                             \
									key;                                             \
								key = tmp;                                               \
							}                                                                \
							if (kh_is_map) {                                                 \
								khval_t tmp =                                            \
									h->vals[i];                                      \
								h->vals[i] =                                             \
									val;                                             \
								val = tmp;                                               \
							}                                                                \
							__ac_set_isdel_true(                                             \
								h->flags,                                                \
								i); /* mark it as deleted in the old hash table */       \
						} else { /* write the element and jump out of the loop */                \
							h->keys[i] = key;                                                \
							if (kh_is_map)                                                   \
								h->vals[i] =                                             \
									val;                                             \
							break;                                                           \
						}                                                                        \
					}                                                                                \
				}                                                                                        \
			}                                                                                                \
			if (h->n_buckets >                                                                               \
			    new_n_buckets) { /* shrink the hash table */                                                 \
				h->keys = (khkey_t *)krealloc(                                                           \
					(void *)h->keys,                                                                 \
					new_n_buckets * sizeof(khkey_t));                                                \
				if (kh_is_map)                                                                           \
					h->vals = (khval_t *)krealloc(                                                   \
						(void *)h->vals,                                                         \
						new_n_buckets *                                                          \
							sizeof(khval_t));                                                \
			}                                                                                                \
			kfree(h->flags); /* free the working space */                                                    \
			h->flags = new_flags;                                                                            \
			h->n_buckets = new_n_buckets;                                                                    \
			h->n_occupied = h->size;                                                                         \
			h->upper_bound = (khint_t)(                                                                      \
				h->n_buckets * __ac_HASH_UPPER + 0.5);                                                   \
		}                                                                                                        \
		return 0;                                                                                                \
	}                                                                                                                \
	SCOPE khint_t kh_put_##name(kh_##name##_t *h, khkey_t key, int *ret)                                             \
	{                                                                                                                \
		khint_t x;                                                                                               \
		if (h->n_occupied >=                                                                                     \
		    h->upper_bound) { /* update the hash table */                                                        \
			if (h->n_buckets > (h->size << 1)) {                                                             \
				if (kh_resize_##name(h, h->n_buckets - 1) <                                              \
				    0) { /* clear "deleted" elements */                                                  \
					*ret = -1;                                                                       \
					return h->n_buckets;                                                             \
				}                                                                                        \
			} else if (kh_resize_##name(h, h->n_buckets + 1) <                                               \
				   0) { /* expand the hash table */                                                      \
				*ret = -1;                                                                               \
				return h->n_buckets;                                                                     \
			}                                                                                                \
		} /* TODO: to implement automatically shrinking; resize() already support shrinking */                   \
		{                                                                                                        \
			khint_t k, i, site, last, mask = h->n_buckets - 1,                                               \
						  step = 0;                                                              \
			x = site = h->n_buckets;                                                                         \
			k = __hash_func(key);                                                                            \
			i = k & mask;                                                                                    \
			if (__ac_isempty(h->flags, i))                                                                   \
				x = i; /* for speed up */                                                                \
			else {                                                                                           \
				last = i;                                                                                \
				while (!__ac_isempty(h->flags, i) &&                                                     \
				       (__ac_isdel(h->flags, i) ||                                                       \
					!__hash_equal(h->keys[i], key))) {                                               \
					if (__ac_isdel(h->flags, i))                                                     \
						site = i;                                                                \
					i = (i + (++step)) & mask;                                                       \
					if (i == last) {                                                                 \
						x = site;                                                                \
						break;                                                                   \
					}                                                                                \
				}                                                                                        \
				if (x == h->n_buckets) {                                                                 \
					if (__ac_isempty(h->flags, i) &&                                                 \
					    site != h->n_buckets)                                                        \
						x = site;                                                                \
					else                                                                             \
						x = i;                                                                   \
				}                                                                                        \
			}                                                                                                \
		}                                                                                                        \
		if (__ac_isempty(h->flags, x)) { /* not present at all */                                                \
			h->keys[x] = key;                                                                                \
			__ac_set_isboth_false(h->flags, x);                                                              \
			++h->size;                                                                                       \
			++h->n_occupied;                                                                                 \
			*ret = 1;                                                                                        \
		} else if (__ac_isdel(h->flags, x)) { /* deleted */                                                      \
			h->keys[x] = key;                                                                                \
			__ac_set_isboth_false(h->flags, x);                                                              \
			++h->size;                                                                                       \
			*ret = 2;                                                                                        \
		} else                                                                                                   \
			*ret = 0; /* Don't touch h->keys[x] if present and not deleted */                                \
		return x;                                                                                                \
	}                                                                                                                \
	SCOPE void kh_del_##name(kh_##name##_t *h, khint_t x)                                                            \
	{                                                                                                                \
		if (x != h->n_buckets && !__ac_iseither(h->flags, x)) {                                                  \
			__ac_set_isdel_true(h->flags, x);                                                                \
			--h->size;                                                                                       \
		}                                                                                                        \
	}

#define KHASH_DECLARE(name, khkey_t, khval_t)                                  \
	__KHASH_TYPE(name, khkey_t, khval_t)                                   \
	__KHASH_PROTOTYPES(name, khkey_t, khval_t)

#define KHASH_INIT2(name, SCOPE, khkey_t, khval_t, kh_is_map, __hash_func,     \
		    __hash_equal)                                              \
	__KHASH_TYPE(name, khkey_t, khval_t)                                   \
	__KHASH_IMPL(name, SCOPE, khkey_t, khval_t, kh_is_map, __hash_func,    \
		     __hash_equal)

#define KHASH_INIT(name, khkey_t, khval_t, kh_is_map, __hash_func,             \
		   __hash_equal)                                               \
	KHASH_INIT2(name, static kh_inline klib_unused, khkey_t, khval_t,      \
		    kh_is_map, __hash_func, __hash_equal)

/* --- BEGIN OF HASH FUNCTIONS --- */

/*! @function
  @abstract     Integer hash function
  @param  key   The integer [khint32_t]
  @return       The hash value [khint_t]
 */
#define kh_int_hash_func(key) (khint32_t)(key)
/*! @function
  @abstract     Integer comparison function
 */
#define kh_int_hash_equal(a, b) ((a) == (b))
/*! @function
  @abstract     64-bit integer hash function
  @param  key   The integer [khint64_t]
  @return       The hash value [khint_t]
 */
#define kh_int64_hash_func(key) (khint32_t)((key) >> 33 ^ (key) ^ (key) << 11)
/*! @function
  @abstract     64-bit integer comparison function
 */
#define kh_int64_hash_equal(a, b) ((a) == (b))
/*! @function
  @abstract     const char* hash function
  @param  s     Pointer to a null terminated string
  @return       The hash value
 */
static kh_inline khint_t __ac_X31_hash_string(const char *s)
{
	khint_t h = (khint_t)*s;
	if (h)
		for (++s; *s; ++s)
			h = (h << 5) - h + (khint_t)*s;
	return h;
}
/*! @function
  @abstract     Another interface to const char* hash function
  @param  key   Pointer to a null terminated string [const char*]
  @return       The hash value [khint_t]
 */
#define kh_str_hash_func(key) __ac_X31_hash_string(key)
/*! @function
  @abstract     Const char* comparison function
 */
#define kh_str_hash_equal(a, b) (strcmp(a, b) == 0)

static kh_inline khint_t __ac_Wang_hash(khint_t key)
{
	key += ~(key << 15);
	key ^= (key >> 10);
	key += (key << 3);
	key ^= (key >> 6);
	key += ~(key << 11);
	key ^= (key >> 16);
	return key;
}
#define kh_int_hash_func2(key) __ac_Wang_hash((khint_t)key)

/* --- END OF HASH FUNCTIONS --- */

/* Other convenient macros... */

/*!
  @abstract Type of the hash table.
  @param  name  Name of the hash table [symbol]
 */
#define khash_t(name) kh_##name##_t

/*! @function
  @abstract     Initiate a hash table.
  @param  name  Name of the hash table [symbol]
  @return       Pointer to the hash table [khash_t(name)*]
 */
#define kh_init(name) kh_init_##name()

/*! @function
  @abstract     Destroy a hash table.
  @param  name  Name of the hash table [symbol]
  @param  h     Pointer to the hash table [khash_t(name)*]
 */
#define kh_destroy(name, h) kh_destroy_##name(h)

/*! @function
  @abstract     Reset a hash table without deallocating memory.
  @param  name  Name of the hash table [symbol]
  @param  h     Pointer to the hash table [khash_t(name)*]
 */
#define kh_clear(name, h) kh_clear_##name(h)

/*! @function
  @abstract     Resize a hash table.
  @param  name  Name of the hash table [symbol]
  @param  h     Pointer to the hash table [khash_t(name)*]
  @param  s     New size [khint_t]
 */
#define kh_resize(name, h, s) kh_resize_##name(h, s)

/*! @function
  @abstract     Insert a key to the hash table.
  @param  name  Name of the hash table [symbol]
  @param  h     Pointer to the hash table [khash_t(name)*]
  @param  k     Key [type of keys]
  @param  r     Extra return code: -1 if the operation failed;
                0 if the key is present in the hash table;
                1 if the bucket is empty (never used); 2 if the element in
				the bucket has been deleted [int*]
  @return       Iterator to the inserted element [khint_t]
 */
#define kh_put(name, h, k, r) kh_put_##name(h, k, r)

/*! @function
  @abstract     Retrieve a key from the hash table.
  @param  name  Name of the hash table [symbol]
  @param  h     Pointer to the hash table [khash_t(name)*]
  @param  k     Key [type of keys]
  @return       Iterator to the found element, or kh_end(h) if the element is absent [khint_t]
 */
#define kh_get(name, h, k) kh_get_##name(h, k)

/*! @function
  @abstract     Remove a key from the hash table.
  @param  name  Name of the hash table [symbol]
  @param  h     Pointer to the hash table [khash_t(name)*]
  @param  k     Iterator to the element to be deleted [khint_t]
 */
#define kh_del(name, h, k) kh_del_##name(h, k)

/*! @function
  @abstract     Test whether a bucket contains data.
  @param  h     Pointer to the hash table [khash_t(name)*]
  @param  x     Iterator to the bucket [khint_t]
  @return       1 if containing data; 0 otherwise [int]
 */
#define kh_exist(h, x) (!__ac_iseither((h)->flags, (x)))

/*! @function
  @abstract     Get key given an iterator
  @param  h     Pointer to the hash table [khash_t(name)*]
  @param  x     Iterator to the bucket [khint_t]
  @return       Key [type of keys]
 */
#define kh_key(h, x) ((h)->keys[x])

/*! @function
  @abstract     Get value given an iterator
  @param  h     Pointer to the hash table [khash_t(name)*]
  @param  x     Iterator to the bucket [khint_t]
  @return       Value [type of values]
  @discussion   For hash sets, calling this results in segfault.
 */
#define kh_val(h, x) ((h)->vals[x])

/*! @function
  @abstract     Alias of kh_val()
 */
#define kh_value(h, x) ((h)->vals[x])

/*! @function
  @abstract     Get the start iterator
  @param  h     Pointer to the hash table [khash_t(name)*]
  @return       The start iterator [khint_t]
 */
#define kh_begin(h) (khint_t)(0)

/*! @function
  @abstract     Get the end iterator
  @param  h     Pointer to the hash table [khash_t(name)*]
  @return       The end iterator [khint_t]
 */
#define kh_end(h) ((h)->n_buckets)

/*! @function
  @abstract     Get the number of elements in the hash table
  @param  h     Pointer to the hash table [khash_t(name)*]
  @return       Number of elements in the hash table [khint_t]
 */
#define kh_size(h) ((h)->size)

/*! @function
  @abstract     Get the number of buckets in the hash table
  @param  h     Pointer to the hash table [khash_t(name)*]
  @return       Number of buckets in the hash table [khint_t]
 */
#define kh_n_buckets(h) ((h)->n_buckets)

/*! @function
  @abstract     Iterate over the entries in the hash table
  @param  h     Pointer to the hash table [khash_t(name)*]
  @param  kvar  Variable to which key will be assigned
  @param  vvar  Variable to which value will be assigned
  @param  code  Block of code to execute
 */
#define kh_foreach(h, kvar, vvar, code)                                        \
	{                                                                      \
		khint_t __i;                                                   \
		for (__i = kh_begin(h); __i != kh_end(h); ++__i) {             \
			if (!kh_exist(h, __i))                                 \
				continue;                                      \
			(kvar) = kh_key(h, __i);                               \
			(vvar) = kh_val(h, __i);                               \
			code;                                                  \
		}                                                              \
	}

/*! @function
  @abstract     Iterate over the values in the hash table
  @param  h     Pointer to the hash table [khash_t(name)*]
  @param  vvar  Variable to which value will be assigned
  @param  code  Block of code to execute
 */
#define kh_foreach_value(h, vvar, code)                                        \
	{                                                                      \
		khint_t __i;                                                   \
		for (__i = kh_begin(h); __i != kh_end(h); ++__i) {             \
			if (!kh_exist(h, __i))                                 \
				continue;                                      \
			(vvar) = kh_val(h, __i);                               \
			code;                                                  \
		}                                                              \
	}

/* More convenient interfaces */

/*! @function
  @abstract     Instantiate a hash set containing integer keys
  @param  name  Name of the hash table [symbol]
 */
#define KHASH_SET_INIT_INT(name)                                               \
	KHASH_INIT(name, khint32_t, char, 0, kh_int_hash_func,                 \
		   kh_int_hash_equal)

/*! @function
  @abstract     Instantiate a hash map containing integer keys
  @param  name  Name of the hash table [symbol]
  @param  khval_t  Type of values [type]
 */
#define KHASH_MAP_INIT_INT(name, khval_t)                                      \
	KHASH_INIT(name, khint32_t, khval_t, 1, kh_int_hash_func,              \
		   kh_int_hash_equal)

/*! @function
  @abstract     Instantiate a hash set containing 64-bit integer keys
  @param  name  Name of the hash table [symbol]
 */
#define KHASH_SET_INIT_INT64(name)                                             \
	KHASH_INIT(name, khint64_t, char, 0, kh_int64_hash_func,               \
		   kh_int64_hash_equal)

/*! @function
  @abstract     Instantiate a hash map containing 64-bit integer keys
  @param  name  Name of the hash table [symbol]
  @param  khval_t  Type of values [type]
 */
#define KHASH_MAP_INIT_INT64(name, khval_t)                                    \
	KHASH_INIT(name, khint64_t, khval_t, 1, kh_int64_hash_func,            \
		   kh_int64_hash_equal)

typedef const char *kh_cstr_t;
/*! @function
  @abstract     Instantiate a hash map containing const char* keys
  @param  name  Name of the hash table [symbol]
 */
#define KHASH_SET_INIT_STR(name)                                               \
	KHASH_INIT(name, kh_cstr_t, char, 0, kh_str_hash_func,                 \
		   kh_str_hash_equal)

/*! @function
  @abstract     Instantiate a hash map containing const char* keys
  @param  name  Name of the hash table [symbol]
  @param  khval_t  Type of values [type]
 */
#define KHASH_MAP_INIT_STR(name, khval_t)                                      \
	KHASH_INIT(name, kh_cstr_t, khval_t, 1, kh_str_hash_func,              \
		   kh_str_hash_equal)

#endif /* __AC_KHASH_H */
```
