## Supplementary material for "GPU Accelerated Adaptive Banded Event Alignment for Rapid Comparative Nanopore Signal Analysis": f5c-v0.2-beta-release: ksort.h

```
/* The MIT License

   Copyright (c) 2008, 2011 Attractive Chaos <>

   Permission is hereby granted, free of charge, to any person obtaining
   a copy of this software and associated documentation files (the
   "Software"), to deal in the Software without restriction, including
   without limitation the rights to use, copy, modify, merge, publish,
   distribute, sublicense, and/or sell copies of the Software, and to
   permit persons to whom the Software is furnished to do so, subject to
   the following conditions:

   The above copyright notice and this permission notice shall be
   included in all copies or substantial portions of the Software.

   THE SOFTWARE IS PROVIDED "AS IS", WITHOUT WARRANTY OF ANY KIND,
   EXPRESS OR IMPLIED, INCLUDING BUT NOT LIMITED TO THE WARRANTIES OF
   MERCHANTABILITY, FITNESS FOR A PARTICULAR PURPOSE AND
   NONINFRINGEMENT. IN NO EVENT SHALL THE AUTHORS OR COPYRIGHT HOLDERS
   BE LIABLE FOR ANY CLAIM, DAMAGES OR OTHER LIABILITY, WHETHER IN AN
   ACTION OF CONTRACT, TORT OR OTHERWISE, ARISING FROM, OUT OF OR IN
   CONNECTION WITH THE SOFTWARE OR THE USE OR OTHER DEALINGS IN THE
   SOFTWARE.
*/

/*
  2011-04-10 (0.1.6):

  	* Added sample

  2011-03 (0.1.5):

	* Added shuffle/permutation

  2008-11-16 (0.1.4):

    * Fixed a bug in introsort() that happens in rare cases.

  2008-11-05 (0.1.3):

    * Fixed a bug in introsort() for complex comparisons.

	* Fixed a bug in mergesort(). The previous version is not stable.

  2008-09-15 (0.1.2):

	* Accelerated introsort. On my Mac (not on another Linux machine),
	  my implementation is as fast as std::sort on random input.

	* Added combsort and in introsort, switch to combsort if the
	  recursion is too deep.

  2008-09-13 (0.1.1):

	* Added k-small algorithm

  2008-09-05 (0.1.0):

	* Initial version

*/

#ifndef AC_KSORT_H
#define AC_KSORT_H

#include <stdlib.h>
#include <string.h>

typedef struct {
	void *left, *right;
	int depth;
} ks_isort_stack_t;

#define KSORT_SWAP(type_t, a, b) { register type_t t=(a); (a)=(b); (b)=t; }

#define KSORT_INIT(name, type_t, __sort_lt)								\
	void ks_mergesort_##name(size_t n, type_t array[], type_t temp[])	\
	{																	\
		type_t *a2[2], *a, *b;											\
		int curr, shift;												\
																		\
		a2[0] = array;													\
		a2[1] = temp? temp : (type_t*)malloc(sizeof(type_t) * n);		\
		for (curr = 0, shift = 0; (1ul<<shift) < n; ++shift) {			\
			a = a2[curr]; b = a2[1-curr];								\
			if (shift == 0) {											\
				type_t *p = b, *i, *eb = a + n;							\
				for (i = a; i < eb; i += 2) {							\
					if (i == eb - 1) *p++ = *i;							\
					else {												\
						if (__sort_lt(*(i+1), *i)) {					\
							*p++ = *(i+1); *p++ = *i;					\
						} else {										\
							*p++ = *i; *p++ = *(i+1);					\
						}												\
					}													\
				}														\
			} else {													\
				size_t i, step = 1ul<<shift;							\
				for (i = 0; i < n; i += step<<1) {						\
					type_t *p, *j, *k, *ea, *eb;						\
					if (n < i + step) {									\
						ea = a + n; eb = a;								\
					} else {											\
						ea = a + i + step;								\
						eb = a + (n < i + (step<<1)? n : i + (step<<1)); \
					}													\
					j = a + i; k = a + i + step; p = b + i;				\
					while (j < ea && k < eb) {							\
						if (__sort_lt(*k, *j)) *p++ = *k++;				\
						else *p++ = *j++;								\
					}													\
					while (j < ea) *p++ = *j++;							\
					while (k < eb) *p++ = *k++;							\
				}														\
			}															\
			curr = 1 - curr;											\
		}																\
		if (curr == 1) {												\
			type_t *p = a2[0], *i = a2[1], *eb = array + n;				\
			for (; p < eb; ++i) *p++ = *i;								\
		}																\
		if (temp == 0) free(a2[1]);										\
	}																	\
	void ks_heapadjust_##name(size_t i, size_t n, type_t l[])			\
	{																	\
		size_t k = i;													\
		type_t tmp = l[i];												\
		while ((k = (k << 1) + 1) < n) {								\
			if (k != n - 1 && __sort_lt(l[k], l[k+1])) ++k;				\
			if (__sort_lt(l[k], tmp)) break;							\
			l[i] = l[k]; i = k;											\
		}																\
		l[i] = tmp;														\
	}																	\
	void ks_heapmake_##name(size_t lsize, type_t l[])					\
	{																	\
		size_t i;														\
		for (i = (lsize >> 1) - 1; i != (size_t)(-1); --i)				\
			ks_heapadjust_##name(i, lsize, l);							\
	}																	\
	void ks_heapsort_##name(size_t lsize, type_t l[])					\
	{																	\
		size_t i;														\
		for (i = lsize - 1; i > 0; --i) {								\
			type_t tmp;													\
			tmp = *l; *l = l[i]; l[i] = tmp; ks_heapadjust_##name(0, i, l); \
		}																\
	}																	\
	static inline void __ks_insertsort_##name(type_t *s, type_t *t)			\
	{																	\
		type_t *i, *j, swap_tmp;										\
		for (i = s + 1; i < t; ++i)										\
			for (j = i; j > s && __sort_lt(*j, *(j-1)); --j) {			\
				swap_tmp = *j; *j = *(j-1); *(j-1) = swap_tmp;			\
			}															\
	}																	\
	void ks_combsort_##name(size_t n, type_t a[])						\
	{																	\
		const double shrink_factor = 1.2473309501039786540366528676643; \
		int do_swap;													\
		size_t gap = n;													\
		type_t tmp, *i, *j;												\
		do {															\
			if (gap > 2) {												\
				gap = (size_t)(gap / shrink_factor);					\
				if (gap == 9 || gap == 10) gap = 11;					\
			}															\
			do_swap = 0;												\
			for (i = a; i < a + n - gap; ++i) {							\
				j = i + gap;											\
				if (__sort_lt(*j, *i)) {								\
					tmp = *i; *i = *j; *j = tmp;						\
					do_swap = 1;										\
				}														\
			}															\
		} while (do_swap || gap > 2);									\
		if (gap != 1) __ks_insertsort_##name(a, a + n);					\
	}																	\
	void ks_introsort_##name(size_t n, type_t a[])						\
	{																	\
		int d;															\
		ks_isort_stack_t *top, *stack;									\
		type_t rp, swap_tmp;											\
		type_t *s, *t, *i, *j, *k;										\
																		\
		if (n < 1) return;												\
		else if (n == 2) {												\
			if (__sort_lt(a[1], a[0])) { swap_tmp = a[0]; a[0] = a[1]; a[1] = swap_tmp; } \
			return;														\
		}																\
		for (d = 2; 1ul<<d < n; ++d);									\
		stack = (ks_isort_stack_t*)malloc(sizeof(ks_isort_stack_t) * ((sizeof(size_t)*d)+2)); \
		top = stack; s = a; t = a + (n-1); d <<= 1;						\
		while (1) {														\
			if (s < t) {												\
				if (--d == 0) {											\
					ks_combsort_##name(t - s + 1, s);					\
					t = s;												\
					continue;											\
				}														\
				i = s; j = t; k = i + ((j-i)>>1) + 1;					\
				if (__sort_lt(*k, *i)) {								\
					if (__sort_lt(*k, *j)) k = j;						\
				} else k = __sort_lt(*j, *i)? i : j;					\
				rp = *k;												\
				if (k != t) { swap_tmp = *k; *k = *t; *t = swap_tmp; }	\
				for (;;) {												\
					do ++i; while (__sort_lt(*i, rp));					\
					do --j; while (i <= j && __sort_lt(rp, *j));		\
					if (j <= i) break;									\
					swap_tmp = *i; *i = *j; *j = swap_tmp;				\
				}														\
				swap_tmp = *i; *i = *t; *t = swap_tmp;					\
				if (i-s > t-i) {										\
					if (i-s > 16) { top->left = s; top->right = i-1; top->depth = d; ++top; } \
					s = t-i > 16? i+1 : t;								\
				} else {												\
					if (t-i > 16) { top->left = i+1; top->right = t; top->depth = d; ++top; } \
					t = i-s > 16? i-1 : s;								\
				}														\
			} else {													\
				if (top == stack) {										\
					free(stack);										\
					__ks_insertsort_##name(a, a+n);						\
					return;												\
				} else { --top; s = (type_t*)top->left; t = (type_t*)top->right; d = top->depth; } \
			}															\
		}																\
	}																	\
	/* This function is adapted from: http://ndevilla.free.fr/median/ */ \
	/* 0 <= kk < n */													\
	type_t ks_ksmall_##name(size_t n, type_t arr[], size_t kk)			\
	{																	\
		type_t *low, *high, *k, *ll, *hh, *mid;							\
		low = arr; high = arr + n - 1; k = arr + kk;					\
		for (;;) {														\
			if (high <= low) return *k;									\
			if (high == low + 1) {										\
				if (__sort_lt(*high, *low)) KSORT_SWAP(type_t, *low, *high); \
				return *k;												\
			}															\
			mid = low + (high - low) / 2;								\
			if (__sort_lt(*high, *mid)) KSORT_SWAP(type_t, *mid, *high); \
			if (__sort_lt(*high, *low)) KSORT_SWAP(type_t, *low, *high); \
			if (__sort_lt(*low, *mid)) KSORT_SWAP(type_t, *mid, *low);	\
			KSORT_SWAP(type_t, *mid, *(low+1));							\
			ll = low + 1; hh = high;									\
			for (;;) {													\
				do ++ll; while (__sort_lt(*ll, *low));					\
				do --hh; while (__sort_lt(*low, *hh));					\
				if (hh < ll) break;										\
				KSORT_SWAP(type_t, *ll, *hh);							\
			}															\
			KSORT_SWAP(type_t, *low, *hh);								\
			if (hh <= k) low = ll;										\
			if (hh >= k) high = hh - 1;									\
		}																\
	}																	\
	void ks_shuffle_##name(size_t n, type_t a[])						\
	{																	\
		int i, j;														\
		for (i = n; i > 1; --i) {										\
			type_t tmp;													\
			j = (int)(drand48() * i);									\
			tmp = a[j]; a[j] = a[i-1]; a[i-1] = tmp;					\
		}																\
	}

#define ks_mergesort(name, n, a, t) ks_mergesort_##name(n, a, t)
#define ks_introsort(name, n, a) ks_introsort_##name(n, a)
#define ks_combsort(name, n, a) ks_combsort_##name(n, a)
#define ks_heapsort(name, n, a) ks_heapsort_##name(n, a)
#define ks_heapmake(name, n, a) ks_heapmake_##name(n, a)
#define ks_heapadjust(name, i, n, a) ks_heapadjust_##name(i, n, a)
#define ks_ksmall(name, n, a, k) ks_ksmall_##name(n, a, k)
#define ks_shuffle(name, n, a) ks_shuffle_##name(n, a)

#define ks_lt_generic(a, b) ((a) < (b))
#define ks_lt_str(a, b) (strcmp((a), (b)) < 0)

typedef const char *ksstr_t;

#define KSORT_INIT_GENERIC(type_t) KSORT_INIT(type_t, type_t, ks_lt_generic)
#define KSORT_INIT_STR KSORT_INIT(str, ksstr_t, ks_lt_str)

#endif
```
