## Supplementary material for "GPU Accelerated Adaptive Banded Event Alignment for Rapid Comparative Nanopore Signal Analysis": f5c-v0.2-beta-release: logsum.h

```
//
// logsum -- a port of Sean Eddy's fast table-driven log sum
// This code was originally part of HMMER. This version is used with
// Sean Eddy's permission as public domain code.
//

#ifndef LOGSUM_H
#define LOGSUM_H

/* p7_LOGSUM_SCALE defines the precision of the calculation; the
 * default of 1000.0 means rounding differences to the nearest 0.001
 * nat. p7_LOGSUM_TBL defines the size of the lookup table; the
 * default of 16000 means entries are calculated for differences of 0
 * to 16.000 nats (when p7_LOGSUM_SCALE is 1000.0).  e^{-p7_LOGSUM_TBL /
 * p7_LOGSUM_SCALE} should be on the order of the machine FLT_EPSILON,
 * typically 1.2e-7.
 */
#define p7_LOGSUM_TBL   16000
#define p7_LOGSUM_SCALE 1000.f
#define ESL_MAX(a,b)    (((a)>(b))?(a):(b))
#define ESL_MIN(a,b)    (((a)<(b))?(a):(b))
#define eslINFINITY     INFINITY

 /* Synopsis:  Initialize the p7_Logsum() function.
 *
 * Purpose:   Initialize the lookup table for <p7_FLogsum()>.
 *            This function must be called once before any
 *            call to <p7_FLogsum()>.
 *
 *            The precision of the lookup table is determined
 *            by the compile-time <p7_LOGSUM_TBL> constant.
 *
 * Returns:   <eslOK> on success.
 */
static inline int p7_FLogsumInit(void)
{

	//   static int firsttime = TRUE;
	//   if (!firsttime) return 1;
	//   firsttime = FALSE;

    extern float flogsum_lookup[p7_LOGSUM_TBL];
	int i;
	for (i = 0; i < p7_LOGSUM_TBL; i++) {
	  flogsum_lookup[i] = log(1. + exp((double) -i / p7_LOGSUM_SCALE));
	}
	return 1;
}

/* Function:  p7_FLogsum()
 * Synopsis:  Approximate $\log(e^a + e^b)$.
 *
 * Purpose:   Returns a fast table-driven approximation to
 *            $\log(e^a + e^b)$.
 *
 *            Either <a> or <b> (or both) may be $-\infty$,
 *            but neither may be $+\infty$ or <NaN>.
 *
 * Note:      This function is a critical optimization target, because
 *            it's in the inner loop of generic Forward() algorithms.*/
static inline float p7_FLogsum(float a, float b){

  extern float flogsum_lookup[p7_LOGSUM_TBL]; /* p7_LOGSUM_TBL=16000: (A-B) = 0..16 nats, steps of 0.001 */

  const float max = ESL_MAX(a, b);
  const float min = ESL_MIN(a, b);

  //return (min == -eslINFINITY || (max-min) >= 15.7f) ? max : max + log(1.0 + exp(min-max));  /* SRE: While debugging SSE impl. Remember to remove! */

  return (min == -eslINFINITY || (max-min) >= 15.7f) ? max : max + flogsum_lookup[(int)((max-min)*p7_LOGSUM_SCALE)];
}

#endif
```
