## Supplementary material for "GPU Accelerated Adaptive Banded Event Alignment for Rapid Comparative Nanopore Signal Analysis": f5c-v0.2-beta-release: main.c

```
/* @f5c
**
** main 
** @author: Hasindu Gamaarachchi
** @@
******************************************************************************/

#include <stdlib.h>
#include <unistd.h>
#include <stdio.h>
#include <string.h>
#include <signal.h>
#include "f5cmisc.h"
#include "error.h"

#ifdef HAVE_EXECINFO_H
    #include <execinfo.h>
#endif

//make the segmentation faults a bit cool
void sig_handler(int sig) {
#ifdef HAVE_EXECINFO_H
    void* array[100];
    size_t size = backtrace(array, 100);
    ERROR("I regret to inform that a segmentation fault occurred. But at least "
          "it is better than a wrong answer%s",
          ".");
    fprintf(stderr,
            "[%s::DEBUG]\033[1;35m Here is the backtrace in case it is of any "
            "use:\n",
            __func__);
    backtrace_symbols_fd(&array[2], size - 1, STDERR_FILENO);
    fprintf(stderr, "\033[0m\n");
#else
    ERROR("I regret to inform that a segmentation fault occurred. But at least "
          "it is better than a wrong answer%s",
          ".");
#endif
    exit(EXIT_FAILURE);
}

int meth_main(int argc, char* argv[], int8_t mode);
int index_main(int argc, char** argv);
int freq_main(int argc, char **argv);

int print_usage(){

    fprintf(stderr,"Usage: f5c <command> [options]\n\n");
    fprintf(stderr,"command:\n");
    fprintf(stderr,"         index               Build an index mapping from basecalled reads to the signals measured by the sequencer (same as nanopolish index)\n");
    fprintf(stderr,"         call-methylation    Classify nucleotides as methylated or not (optimised nanopolish call-methylation)\n");
    fprintf(stderr,"         meth-freq           Calculate methylation frequency at genomic CpG sites (optimised nanopolish calculate_methylation_frequency.py)\n");
    fprintf(stderr,"         eventalign          Align nanopore events to reference k-mers (optimised nanopolish eventalign)\n\n");


    exit(EXIT_FAILURE);
}


int main(int argc, char* argv[]){

    double realtime0 = realtime();
    signal(SIGSEGV, sig_handler);

    int ret=1;

    if(argc<2){
        return print_usage();
    }
    if(strcmp(argv[1],"index")==0){
        ret=index_main(argc-1, argv+1);
    }
    else if(strcmp(argv[1],"call-methylation")==0){
        ret=meth_main(argc-1, argv+1,0);
    }
    else if(strcmp(argv[1],"eventalign")==0){
        ret=meth_main(argc-1, argv+1,1);
    }    
    else if(strcmp(argv[1],"meth-freq")==0){
        ret=freq_main(argc-1, argv+1);
    }
    else{
        fprintf(stderr,"[f5c] Unrecognised command %s\n",argv[1]);
        print_usage();
    }

    fprintf(stderr, "\n[%s] CMD:", __func__);
    for (int i = 0; i < argc; ++i) {
        fprintf(stderr, " %s", argv[i]);
    }

    fprintf(stderr, "\n[%s] Real time: %.3f sec; CPU time: %.3f sec; Peak RAM: %.3f GB\n\n",
            __func__, realtime() - realtime0, cputime(),peakrss() / 1024.0 / 1024.0 / 1024.0);

    return ret;
}
```
