## Supplementary material for "GPU Accelerated Adaptive Banded Event Alignment for Rapid Comparative Nanopore Signal Analysis": f5c-v0.2-beta-release: matrix.h

```
//---------------------------------------------------------
// Copyright 2015 Ontario Institute for Cancer Research
// Written by Jared Simpson
//---------------------------------------------------------
//
// nanopolish_matrix -- matrix manipulation functions
//
#ifndef MATRIX_H
#define MATRIX_H

#include <math.h>
#include <stdlib.h>
#include <stdio.h>
#include <assert.h>
#include <string.h>

//
// Template Matrix for POD types
//
template<typename T>
struct Matrix
{
    T* cells;
    uint32_t n_rows;
    uint32_t n_cols;
};

typedef Matrix<double> DoubleMatrix;
typedef Matrix<float> FloatMatrix;
typedef Matrix<uint32_t> UInt32Matrix;
typedef Matrix<uint8_t> UInt8Matrix;

//
template<typename T>
void allocate_matrix(Matrix<T>& matrix, uint32_t n_rows, uint32_t n_cols)
{
    matrix.n_rows = n_rows;
    matrix.n_cols = n_cols;

    uint32_t N = matrix.n_rows * matrix.n_cols;
    matrix.cells = (T*)malloc(N * sizeof(T));
    memset(matrix.cells, 0, N * sizeof(T));
}

//
template<typename T>
void free_matrix(Matrix<T>& matrix)
{
    assert(matrix.cells != NULL);
    free(matrix.cells);
    matrix.cells = NULL;
}

// Copy a matrix and its contents
template<typename T>
void copy_matrix(Matrix<T>& new_matrix, const Matrix<T>& old_matrix)
{
    allocate_matrix(new_matrix, old_matrix.n_rows, old_matrix.n_cols);
    uint32_t bytes = sizeof(T) * new_matrix.n_rows * new_matrix.n_cols;
    memcpy(new_matrix.cells, old_matrix.cells, bytes);
}

template<typename T>
inline uint32_t cell(const Matrix<T>& matrix, uint32_t row, uint32_t col)
{
    return row * matrix.n_cols + col;
}

template<typename T, typename U>
inline void set(Matrix<T>& matrix, uint32_t row, uint32_t col, U v)
{
    uint32_t c = cell(matrix, row, col);
    matrix.cells[c] = v;
}

template<typename T>
inline T get(const Matrix<T>& matrix, uint32_t row, uint32_t col)
{
    uint32_t c = cell(matrix, row, col);
    return matrix.cells[c];
}

//
inline void print_matrix(const DoubleMatrix& matrix, bool do_exp = false)
{
    for(uint32_t i = 0; i < matrix.n_rows; ++i) {
        for(uint32_t j = 0; j < matrix.n_cols; ++j) {
            uint32_t c = cell(matrix, i, j);
            double v = matrix.cells[c];
            if(do_exp)
                v = exp(v);
            printf("%.3lf\t", v);
        }
        printf("\n");
    }
}

#endif
```
