## Supplementary material for "GPU Accelerated Adaptive Banded Event Alignment for Rapid Comparative Nanopore Signal Analysis": f5c-v0.2-beta-release: meth.c

```
#include <assert.h>
#include <stdlib.h>
#include <stdio.h>
#include <algorithm>

#include "f5c.h"
#include "f5cmisc.h"

#define METHYLATED_SYMBOL 'M'

#define METH_DEBUG 1

//following Code is adapted from Nanopolish

int32_t get_aligned_segments(const bam1_t* record, int read_stride, AlignedPair *segments, int32_t read_len)
{
    // Initialize first segment
    int32_t segment_size = 0;

    // This code is derived from bam_fillmd1_core
    //uint8_t *ref = NULL;
    //uint8_t *seq = bam_get_seq(record);
    uint32_t *cigar = bam_get_cigar(record);
    const bam1_core_t *c = &record->core;

    // read pos is an index into the original sequence that is present in the FASTQ
    // on the strand matching the reference
    int read_pos = 0;

    // query pos is an index in the query string that is recorded in the bam
    // we record this as a sanity check
    //int query_pos = 0;

    int ref_pos = c->pos;

    for (uint32_t ci = 0; ci < c->n_cigar; ++ci) {

        int cigar_len = cigar[ci] >> 4;
        int cigar_op = cigar[ci] & 0xf;

        // Set the amount that the ref/read positions should be incremented
        // based on the cigar operation
        int read_inc = 0;
        int ref_inc = 0;

        // Process match between the read and the reference
        bool is_aligned = false;
        if(cigar_op == BAM_CMATCH || cigar_op == BAM_CEQUAL || cigar_op == BAM_CDIFF) {
            is_aligned = true;
            read_inc = read_stride;
            ref_inc = 1;
        } else if(cigar_op == BAM_CDEL) {
            ref_inc = 1;
        } else if(cigar_op == BAM_CREF_SKIP) {
            // end the current segment and start a new one
            fprintf(stderr, "Error: spliced alignments detected when loading read %s\n", bam_get_qname(record));
            fprintf(stderr, "Please align the reads to the genome using a non-spliced aligner\n");
            exit(EXIT_FAILURE);
        } else if(cigar_op == BAM_CINS) {
            read_inc = read_stride;
        } else if(cigar_op == BAM_CSOFT_CLIP) {
            read_inc = 1; // special case, do not use read_stride
        } else if(cigar_op == BAM_CHARD_CLIP) {
            read_inc = 0;
        } else {
            printf("Cigar: %d\n", cigar_op);
            assert(false && "Unhandled cigar operation");
        }

        // Iterate over the pairs of aligned bases
        for(int j = 0; j < cigar_len; ++j) {
            if(is_aligned) {
                AlignedPair p = {
                    .ref_pos = ref_pos,
                    .read_pos = read_pos
                };
                segments[segment_size++] = p;
                assert(segment_size <= segment_size);
            }

            // increment
            read_pos += read_inc;
            ref_pos += ref_inc;
        }
    }
    return segment_size;
}


// helper for get_closest_event_to
int get_next_event(int start, int stop, int stride,index_pair_t* base_to_event_map)
{
    while(start != stop) {

        int ei = base_to_event_map[start].start;
        if(ei != -1)
            return ei;
        start += stride;
    }
    return -1;
}


int get_closest_event_to(int k_idx, index_pair_t* base_to_event_map, int base_to_event_map_size)
{
    int stop_before = std::max(0, k_idx - 1000);
    int stop_after = std::min(k_idx + 1000, base_to_event_map_size - 1);

    int event_before = get_next_event(k_idx, stop_before, -1, base_to_event_map);
    int event_after = get_next_event(k_idx, stop_after, 1, base_to_event_map);

    // TODO: better selection of "best" event to return
    if(event_before == -1)
        return event_after;
    return event_before;
}

inline int32_t flip_k_strand(int32_t read_length,int32_t k_idx, uint32_t k)
{
    return read_length - k_idx - k;
}

int32_t get_event_alignment_record(const bam1_t* record,int32_t read_length, index_pair_t* base_to_event_map, AlignedPair **aligned_events_ptr)
{

    uint8_t rc = bam_is_rev(record);
    //int8_t stride;

    // copy read base-to-reference alignment
    AlignedPair *seq_record = (AlignedPair *)malloc(read_length*sizeof(AlignedPair));
    MALLOC_CHK(seq_record);

    int32_t segments_size = get_aligned_segments(record, 1, seq_record,read_length);
    assert(segments_size <= read_length && read_length >= 0);

    *aligned_events_ptr = (AlignedPair *)malloc(segments_size* sizeof(AlignedPair));
    AlignedPair *aligned_events = *aligned_events_ptr;
    MALLOC_CHK(aligned_events);

    int32_t aligned_events_size = 0;
    //this->sr = sr;
    int32_t k = KMER_SIZE;
    //size_t read_length = this->sr->read_sequence.length();

    for(int32_t i = 0; i < segments_size; ++i) {
        // skip positions at the boundary
        if(seq_record[i].read_pos < k) {
            continue;
        }

        if(seq_record[i].read_pos + k >= read_length) {
            continue;
        }

        int32_t kmer_pos_ref_strand = seq_record[i].read_pos;
        int32_t kmer_pos_read_strand = rc ? flip_k_strand(read_length,kmer_pos_ref_strand, k) : kmer_pos_ref_strand;
        int32_t event_idx = get_closest_event_to(kmer_pos_read_strand, base_to_event_map, read_length-KMER_SIZE + 1);
        AlignedPair p = {
            .ref_pos = seq_record[i].ref_pos,
	    .read_pos = (int)event_idx
	    };
	    aligned_events[aligned_events_size++] = p;
	    assert(aligned_events_size <= segments_size);
    }
    //this->rc = strand_idx == 0 ? seq_record.rc : !seq_record.rc;
    //this->strand = strand_idx;

    if(aligned_events_size) {
        //stride = aligned_events.front().read_pos < aligned_events.back().read_pos ? 1 : -1;

        // check for a degenerate alignment and discard the events if so
        if(aligned_events[0].read_pos == aligned_events[aligned_events_size - 1].read_pos) {
            free(aligned_events);
	    *aligned_events_ptr = NULL;
	    aligned_events_size = 0;
        }
    }
    // else {
    //     stride = 1;
    // }
    free(seq_record);

    return aligned_events_size;
}


// IUPAC alphabet
bool isUnambiguous(char c) {
    switch (c) {
        case 'A':
        case 'C':
        case 'G':
        case 'T':
            return true;
        default:
            return false;
    }
}

// Returns true if c is a valid ambiguity code
bool isAmbiguous(char c) {
    switch (c) {
        case 'M':
        case 'R':
        case 'W':
        case 'S':
        case 'Y':
        case 'K':
        case 'V':
        case 'H':
        case 'D':
        case 'B':
        case 'N':
            return true;
        default:
            return false;
    }
}

std::string getPossibleSymbols(char c) {
    switch (c) {
        case 'A':
            return "A";
        case 'C':
            return "C";
        case 'G':
            return "G";
        case 'T':
            return "T";
        case 'M':
            return "AC";
        case 'R':
            return "AG";
        case 'W':
            return "AT";
        case 'S':
            return "CG";
        case 'Y':
            return "CT";
        case 'K':
            return "GT";
        case 'V':
            return "ACG";
        case 'H':
            return "ACT";
        case 'D':
            return "AGT";
        case 'B':
            return "CGT";
        case 'N':
            return "ACGT";
        default:
            return "";
    }
}

// Returns true if c is a valid symbol in this alphabet
bool isValid(char c) { return isUnambiguous(c) || isAmbiguous(c); }

const char* complement_dna = "TGCA";
const uint8_t rank_dna[256] = {
    0, 0, 0, 0, 0, 0, 0, 0, 0, 0, 0, 0, 0, 0, 0, 0, 0, 0, 0, 0, 0, 0, 0, 0,
    0, 0, 0, 0, 0, 0, 0, 0, 0, 0, 0, 0, 0, 0, 0, 0, 0, 0, 0, 0, 0, 0, 0, 0,
    0, 0, 0, 0, 0, 0, 0, 0, 0, 0, 0, 0, 0, 0, 0, 0, 0, 0, 0, 1, 0, 0, 0, 2,
    0, 0, 0, 0, 0, 0, 0, 0, 0, 0, 0, 0, 3, 0, 0, 0, 0, 0, 0, 0, 0, 0, 0, 0,
    0, 0, 0, 0, 0, 0, 0, 0, 0, 0, 0, 0, 0, 0, 0, 0, 0, 0, 0, 0, 0, 0, 0, 0,
    0, 0, 0, 0, 0, 0, 0, 0, 0, 0, 0, 0, 0, 0, 0, 0, 0, 0, 0, 0, 0, 0, 0, 0,
    0, 0, 0, 0, 0, 0, 0, 0, 0, 0, 0, 0, 0, 0, 0, 0, 0, 0, 0, 0, 0, 0, 0, 0,
    0, 0, 0, 0, 0, 0, 0, 0, 0, 0, 0, 0, 0, 0, 0, 0, 0, 0, 0, 0, 0, 0, 0, 0,
    0, 0, 0, 0, 0, 0, 0, 0, 0, 0, 0, 0, 0, 0, 0, 0, 0, 0, 0, 0, 0, 0, 0, 0,
    0, 0, 0, 0, 0, 0, 0, 0, 0, 0, 0, 0, 0, 0, 0, 0, 0, 0, 0, 0, 0, 0, 0, 0,
    0, 0, 0, 0, 0, 0, 0, 0, 0, 0, 0, 0, 0, 0, 0, 0};

// reverse-complement a DNA string
std::string reverse_complement(const std::string& str) {
    std::string out(str.length(), 'A');
    size_t i = 0;             // input
    int j = str.length() - 1; // output
    while (i < str.length()) {
        // complement a single base
        assert(str[i] != METHYLATED_SYMBOL);
        out[j--] = complement_dna[rank_dna[(int)str[i++]]];
    }
    return out;
}

std::string disambiguate(const std::string& str) {
    // create output and convert lower case to upper case
    std::string out(str);
    std::transform(out.begin(), out.end(), out.begin(), ::toupper);

    size_t i = 0;
    while (i < out.length()) {
        size_t stride = 1;
        //bool is_recognition_site = false;

        assert(isValid(out[i]));
        out[i] = getPossibleSymbols(out[i])[0];
        stride = 1;


        i += stride;
    }
    return out;
}


struct RecognitionMatch
{
    unsigned offset; // the matched position in the recognition site
    unsigned length; // the length of the match, 0 indicates no match
    bool covers_methylated_site; // does the match cover an M base?
};

const uint32_t num_recognition_sites = 1;
const uint32_t recognition_length = 2;
const char* recognition_sites[] = { "CG" };
const char* recognition_sites_methylated[] = { "MG" };
const char* recognition_sites_methylated_complement[] = { "GM" };


// Check whether a recognition site starts at position i of str
inline RecognitionMatch match_to_site(const std::string& str, size_t i, const char* recognition, size_t rl)
{
    RecognitionMatch match;
    match.length = 0;
    match.offset = 0;
    match.covers_methylated_site = false;

    // Case 1: str is a substring of recognition
    const char* p = strstr(recognition, str.c_str());
    if(i == 0 && p != NULL) {
        match.offset = p - recognition;
        match.length = str.length();
    } else {
        // Case 2: the suffix str[i..n] is a prefix of recognition
        size_t cl = std::min(rl, str.length() - i);
        if(str.compare(i, cl, recognition, cl) == 0) {
            match.offset = 0;
            match.length = cl;
        }
    }

    //printf("Match site: %s %s %s %d %d\n", str.c_str(), str.substr(i).c_str(), recognition, match.offset, match.length);
    if(match.length > 0) {
        match.covers_methylated_site =
            str.substr(i, match.length).find_first_of(METHYLATED_SYMBOL) != std::string::npos;
    }

    return match;
}


// If the alphabet supports methylated bases, convert str
// to a methylated string using the recognition sites
std::string methylate(const std::string& str)
{
    std::string out(str);
    size_t i = 0;
    while(i < out.length()) {
        size_t stride = 1;

        // Does this location match a recognition site?
        for(size_t j = 0; j < num_recognition_sites; ++j) {

            RecognitionMatch match = match_to_site(str, i, recognition_sites[j], recognition_length);
            // Require the recognition site to be completely matched
            if(match.length == recognition_length) {
                // Replace by the methylated version
                out.replace(i, recognition_length, recognition_sites_methylated[j]);
                stride = match.length; // skip to end of match
                break;
            }
        }

        i += stride;
    }
    return out;
}

// reverse-complement a string meth aware
// when the string contains methylated bases, the methylation
// symbol transfered to the output strand in the appropriate position
std::string reverse_complement_meth(const std::string& str)
{
    std::string out(str.length(), 'A');
    size_t i = 0; // input
    int j = str.length() - 1; // output
    while(i < str.length()) {
        int recognition_index = -1;
        RecognitionMatch match;

        // Does this location (partially) match a methylated recognition site?
        for(size_t j = 0; j < num_recognition_sites; ++j) {
            match = match_to_site(str, i, recognition_sites_methylated[j], recognition_length);
            if(match.length > 0 && match.covers_methylated_site) {
                recognition_index = j;
                break;
            }
        }

        // If this subsequence matched a methylated recognition site,
        // copy the complement of the site to the output
        if(recognition_index != -1) {
            for(size_t k = match.offset; k < match.offset + match.length; ++k) {
                out[j--] = recognition_sites_methylated_complement[recognition_index][k];
                i += 1;
            }
        } else {
            // complement a single base
            assert(str[i] != METHYLATED_SYMBOL);
            //out[j--] = complement(str[i++]);
            out[j--] = complement_dna[rank_dna[(int)str[i++]]];
        }
    }
    return out;
}

int aligned_pair_lower_bound(const AlignedPair *a, int low, int high, int v) {
    while (low < high) {
        int mid = low + (high - low) / 2;
        if (a[mid].ref_pos < v)
                low = mid + 1;
        else
                high = mid;
        }
    return low;
}

bool find_iter_by_ref_bounds(const AlignedPair *pairs, size_t pairs_size,
                             int ref_start, int ref_stop,
                             int *start_iter,
                             int *stop_iter) {
    *start_iter = aligned_pair_lower_bound(pairs, 0, pairs_size, ref_start);
    *stop_iter = aligned_pair_lower_bound(pairs, 0, pairs_size, ref_stop);

    if ((size_t)*start_iter == pairs_size || (size_t)*stop_iter == pairs_size)
        return false;

    // require at least one aligned reference base at or outside the boundary
    bool left_bounded =
        pairs[*start_iter].ref_pos <= ref_start ||
        (*start_iter != 0 && pairs[*start_iter - 1].ref_pos <= ref_start);

    bool right_bounded =
        pairs[*stop_iter].ref_pos >= ref_stop ||
        ((size_t)*stop_iter != pairs_size && pairs[*stop_iter + 1].ref_pos >= ref_start);

    return left_bounded && right_bounded;
}

bool find_by_ref_bounds(const AlignedPair *pairs, size_t pairs_size, int ref_start,
                        int ref_stop, int& read_start, int& read_stop) {
    int start_i;
    int stop_i;
    bool bounded = find_iter_by_ref_bounds(pairs, pairs_size, ref_start, ref_stop,
                                           &start_i, &stop_i);
    if (bounded) {
        read_start = pairs[start_i].read_pos;
        read_stop = pairs[stop_i].read_pos;
        return true;
    }
    return false;
}


// struct ScoredSite
// {
//     ScoredSite()
//     {
//         ll_unmethylated[0] = 0;
//         ll_unmethylated[1] = 0;
//         ll_methylated[0] = 0;
//         ll_methylated[1] = 0;
//         strands_scored = 0;
//     }

//     std::string chromosome;
//     int start_position;
//     int end_position;
//     int n_cpg;
//     std::string sequence;

//     // scores per strand
//     double ll_unmethylated[2];
//     double ll_methylated[2];
//     int strands_scored;

//     //
//     static bool sort_by_position(const ScoredSite& a, const ScoredSite& b) { return a.start_position < b.start_position; }

// };


// Test CpG sites in this read for methylation
void calculate_methylation_for_read(std::map<int, ScoredSite>* site_score_map, char* ref, bam1_t* record, int32_t read_length, event_t* event, index_pair_t* base_to_event_map,
scalings_t scaling, model_t* cpgmodel,double events_per_base) {


    //todo : have to check this, if required have to implement
    // if(!sr.has_events_for_strand(strand_idx)) {
    //     continue;
    // }

    //size_t k = sr.get_model_k(strand_idx);
    uint32_t k = KMER_SIZE;

    //todo : this is required in a generic version of the tool
    // // check if there is a cpg model for this strand
    // if(!PoreModelSet::has_model(sr.get_model_kit_name(strand_idx),
    //                             "cpg",
    //                             sr.get_model_strand_name(strand_idx),
    //                             k))
    // {
    //     continue;
    // }

    int ref_start_pos = record->core.pos;
    //int ref_end_pos =  bam_endpos(record);

    // Remove non-ACGT bases from this reference segment
    std::string ref_seq = ref; //todo : this is a hack due to pure laziness convert to C
    ref_seq = disambiguate(ref_seq); //todo :  convert to C

    // Scan the sequence for CpGs
    assert(ref_seq.size() != 0);
    int *cpg_sites = (int *)calloc(ref_seq.size() - 1, sizeof(int));
    MALLOC_CHK(cpg_sites);
    size_t cpg_sites_size = 0;
    for (size_t i = 0; i < ref_seq.size() - 1; ++i) {
        if (ref_seq[i] == 'C' && ref_seq[i + 1] == 'G') {
            cpg_sites[cpg_sites_size++] = i;
        }
    }

    // Batch the CpGs together into groups that are separated by some minimum distance
    int min_separation = 10;
    std::pair<int, int> *groups = (std::pair<int, int> *)calloc(cpg_sites_size, sizeof(std::pair<int, int>));
    MALLOC_CHK(groups);
    size_t group_size = 0;

    size_t curr_idx = 0;
    while (curr_idx < cpg_sites_size) {
        // Find the endpoint of this group of sites
        size_t end_idx = curr_idx + 1;
        while (end_idx < cpg_sites_size) {
            if (cpg_sites[end_idx] - cpg_sites[end_idx - 1] > min_separation)
                break;
            end_idx += 1;
        }
	groups[group_size++] = std::make_pair(curr_idx, end_idx);
        curr_idx = end_idx;
    }

    for (size_t group_idx = 0; group_idx < group_size; ++group_idx) {
        size_t start_idx = groups[group_idx].first;
        size_t end_idx = groups[group_idx].second;

        // the coordinates on the reference substring for this group of sites
        int sub_start_pos = cpg_sites[start_idx] - min_separation;
        int sub_end_pos = cpg_sites[end_idx - 1] + min_separation;
        int span = cpg_sites[end_idx - 1] - cpg_sites[start_idx];

        // skip if too close to the start of the read alignment or
        // if the reference range is too large to efficiently call
        if (sub_start_pos <= min_separation || span > 200) {
            continue;
        }

        //todo : convert to C
        std::string subseq =
            ref_seq.substr(sub_start_pos, sub_end_pos - sub_start_pos + 1);
        std::string rc_subseq = reverse_complement(subseq); //todo : fix this to meth aware generic complement function

        int calling_start = sub_start_pos + ref_start_pos;
        int calling_end = sub_end_pos + ref_start_pos;

        // Build the event-to-reference map for this read from the bam record
        AlignedPair *event_align_record = NULL;
        int32_t event_align_record_size = get_event_alignment_record(record, read_length, base_to_event_map, &event_align_record);

        // using the reference-to-event map, look up the event indices for this segment
        int e1, e2;
        bool bounded = find_by_ref_bounds(
             event_align_record, event_align_record_size, calling_start, calling_end, e1,
            e2);

        if (event_align_record_size) {
            free(event_align_record);
        }

        double ratio = fabs(e2 - e1) / (calling_start - calling_end);

        // Only process this region if the the read is aligned within the boundaries
        // and the span between the start/end is not unusually short
        if (!bounded || abs(e2 - e1) <= 10 || ratio > MAX_EVENT_TO_BP_RATIO) {
            continue;
        }

        uint32_t hmm_flags = HAF_ALLOW_PRE_CLIP | HAF_ALLOW_POST_CLIP;

        // Set up event data
        uint8_t strand = 0;
        uint32_t event_start_idx=e1;
        uint32_t event_stop_idx=e2;
        int8_t event_stride = event_start_idx <= event_stop_idx ? 1 : -1;
        uint8_t rc = bam_is_rev(record); //only for strand 0

        const char* m_seq = subseq.c_str();
        const char* m_rc_seq = rc_subseq.c_str();

        double unmethylated_score=profile_hmm_score(m_seq,m_rc_seq, event, scaling, cpgmodel, event_start_idx, event_stop_idx,
        strand,event_stride,rc,events_per_base,hmm_flags);

        // Methylate all CpGs in the sequence and score again
        std::string mcpg_subseq = methylate(subseq);
        std::string rc_mcpg_subseq = reverse_complement_meth(mcpg_subseq);

        double methylated_score=profile_hmm_score(mcpg_subseq.c_str(),rc_mcpg_subseq.c_str(), event, scaling, cpgmodel, event_start_idx, event_stop_idx,
        strand,event_stride,rc,events_per_base,hmm_flags);
        //fprintf(stderr,"meth score %f\n",methylated_score);

        // Aggregate score
        int start_position = cpg_sites[start_idx] + ref_start_pos;
        auto iter = site_score_map->find(start_position);
        if (iter == site_score_map->end()) {
            // insert new score into the map
            ScoredSite ss;
            //ss.chromosome = contig;
            ss.start_position = start_position;
            ss.end_position = cpg_sites[end_idx - 1] + ref_start_pos;
            ss.n_cpg = end_idx - start_idx;

            // extract the CpG site(s) with a k-mers worth of surrounding context
            size_t site_output_start = cpg_sites[start_idx] - k + 1;
            size_t site_output_end = cpg_sites[end_idx - 1] + k;
            ss.sequence = ref_seq.substr(site_output_start,
                                         site_output_end - site_output_start);

            // insert into the map
            iter =
                site_score_map->insert(std::make_pair(start_position, ss)).first;
        }
        // set strand-specific score
        // upon output below the strand scores will be summed
        int strand_idx=0;
        iter->second.ll_unmethylated[strand_idx] = unmethylated_score;
        iter->second.ll_methylated[strand_idx] = methylated_score;
        iter->second.strands_scored += 1;
    } // for group
    free(groups);
    free(cpg_sites);
}
```
