## Supplementary material for "GPU Accelerated Adaptive Banded Event Alignment for Rapid Comparative Nanopore Signal Analysis": f5c-v0.2-beta-release: meth_main.c

```
#include "f5c.h"
#include "f5cmisc.h"
#include <assert.h>
#include <getopt.h>
#include <pthread.h>
#include <stdint.h>
#include <stdio.h>
#include <stdlib.h>
#include <unistd.h>
#include "logsum.h"

/* Input/processing/output interleave framework :
unless IO_PROC_NO_INTERLEAVE is set input, processing and output are interleaved
main thread
1. allocates and loads a databatch
2. create `pthread_processor` thread which will perform the processing
(note that `pthread_processor` is the process-controller that will spawn user specified number of processing threads - see below)
3. create the `pthread_post_processor` thread that will print the output and free the databatch one the `pthread_processor` is done
4. allocates and load another databatch
5. wait till the previous `pthread_processor` is done and perform 2
6. wait till the previous `pthread_post_processor` is done and perform 3
7. goto 4 untill all the input is processed
*/

/* Multithreaded framework for processing


*/


/* CUDA acceleration


*/

/*
TODO :
--print-* options should take a filename
a step controller if one requires to perform only upto a certain step such as alignment, ignoring the reset
--the names of functions and variablrs starting with meth_ are confusing as it stands for both meth and event now. should be fixed later
*/

/*
LIMITATIONS :
Does not support multi strand reads (2D and 1D^2 reads) at the moment
Only works for DNA at the moment
*/

//fast logsum data structure
float flogsum_lookup[p7_LOGSUM_TBL]; //todo : get rid of global vars

static struct option long_options[] = {
    {"reads", required_argument, 0, 'r'},          //0 fastq/fasta read file
    {"bam", required_argument, 0, 'b'},            //1 sorted bam file
    {"genome", required_argument, 0, 'g'},         //2 reference genome
    {"threads", required_argument, 0, 't'},        //3 number of threads [8]
    {"batchsize", required_argument, 0, 'K'},      //4 batchsize - number of reads loaded at once [512]
    {"max-bases", required_argument, 0, 'B'},      //5 batchsize - number of bases loaded at once
    {"verbose", required_argument, 0, 'v'},        //6 verbosity level [1]
    {"help", no_argument, 0, 'h'},                 //7
    {"version", no_argument, 0, 'V'},              //8
    {"min-mapq", required_argument, 0, 0},         //9 consider only reads with MAPQ>=min-mapq [30]
    {"secondary", required_argument, 0, 0},        //10 consider secondary alignments or not [yes]
    {"kmer-model", required_argument, 0, 0},       //11 custom k-mer model file (used for debugging)
    {"skip-unreadable", required_argument, 0, 0},  //12 skip any unreadable fast5 or terminate program [yes]
    {"print-events", required_argument, 0, 0},     //13 prints the event table (used for debugging)
    {"print-banded-aln", required_argument, 0, 0}, //14 prints the event alignment (used for debugging)
    {"print-scaling", required_argument, 0, 0},    //15 prints the estimated scalings (used for debugging)
    {"print-raw", required_argument, 0, 0},        //16 prints the raw signal (used for debugging)
    {"disable-cuda", required_argument, 0, 0},     //17 disable running on CUDA [no] (only if compiled for CUDA)
    {"cuda-block-size",required_argument, 0, 0},   //18
    {"debug-break",required_argument, 0, 0},       //19 break after processing the first batch (used for debugging)
    {"profile-cpu",required_argument, 0, 0},       //20 perform section by section (used for profiling - for CPU only)
    {"cuda-max-lf",required_argument, 0, 0},       //21 reads <= cuda-max-lf*avg_readlen on GPU, rest on CPU (only if compiled for CUDA)
    {"cuda-avg-epk",required_argument, 0, 0},      //22 average number of events per kmer - for allocating GPU arrays (only if compiled for CUDA)
    {"cuda-max-epk",required_argument, 0, 0},      //23 reads <= cuda_max_epk on GPU, rest on CPU (only if compiled for CUDA)
    {"cuda-dev-id",required_argument, 0, 0},       //24 cuda device ID to run on (only if compiled for CUDA)
    {"cuda-mem-frac",required_argument, 0, 0},     //25 fraction of the free GPU memory to use (only if compiled for CUDA)
    {"skip-ultra",required_argument, 0, 0},        //26 skip the ultra long reads for better load balancing
    {"ultra-thresh",required_argument, 0, 0},      //27 the threadshold for skipping ultra long reads
    {"write-dump",required_argument, 0, 0},        //28 write the raw data as a dump
    {"read-dump",required_argument, 0, 0},         //29 read the raw data as a dump
    {0, 0, 0, 0}};


static inline int64_t mm_parse_num(const char* str) //taken from minimap2
{
    double x;
    char* p;
    x = strtod(str, &p);
    if (*p == 'G' || *p == 'g')
        x *= 1e9;
    else if (*p == 'M' || *p == 'm')
        x *= 1e6;
    else if (*p == 'K' || *p == 'k')
        x *= 1e3;
    return (int64_t)(x + .499);
}

//parse yes or no arguments : taken from minimap2
static inline void yes_or_no(opt_t* opt, uint64_t flag, int long_idx,
                             const char* arg,
                             int yes_to_set)
{
    if (yes_to_set) {
        if (strcmp(arg, "yes") == 0 || strcmp(arg, "y") == 0) {
            opt->flag |= flag;
        } else if (strcmp(arg, "no") == 0 || strcmp(arg, "n") == 0) {
            opt->flag &= ~flag;
        } else {
            WARNING("option '--%s' only accepts 'yes' or 'no'.",
                    long_options[long_idx].name);
        }
    } else {
        if (strcmp(arg, "yes") == 0 || strcmp(arg, "y") == 0) {
            opt->flag &= ~flag;
        } else if (strcmp(arg, "no") == 0 || strcmp(arg, "n") == 0) {
            opt->flag |= flag;
        } else {
            WARNING("option '--%s' only accepts 'yes' or 'no'.",
                    long_options[long_idx].name);
        }
    }
}


//function that processes a databatch - for pthreads when I/O and processing are interleaved
void* pthread_processor(void* voidargs) {
    pthread_arg2_t* args = (pthread_arg2_t*)voidargs;
    db_t* db = args->db;
    core_t* core = args->core;
    double realtime0=core->realtime0;

    //process
    process_db(core, db);

    fprintf(stderr, "[%s::%.3f*%.2f] %d Entries (%.1fM bases) processed\n", __func__,
                realtime() - realtime0, cputime() / (realtime() - realtime0),
                db->n_bam_rec,db->sum_bases/(1000.0*1000.0));

    //need to inform the output thread that we completed the processing
    pthread_mutex_lock(&args->mutex);
    pthread_cond_signal(&args->cond);
    args->finished=1;
    pthread_mutex_unlock(&args->mutex);

    if(core->opt.verbosity > 1){
        fprintf(stderr, "[%s::%.3f*%.2f] Signal sent by processor thread!\n", __func__,
                realtime() - realtime0, cputime() / (realtime() - realtime0));
    }

    pthread_exit(0);
}


//function that prints the output and free - for pthreads when I/O and processing are interleaved
void* pthread_post_processor(void* voidargs){
    pthread_arg2_t* args = (pthread_arg2_t*)voidargs;
    db_t* db = args->db;
    core_t* core = args->core;
    double realtime0=core->realtime0;

    //wait until the processing thread has informed us
    pthread_mutex_lock(&args->mutex);
    while(args->finished==0){
        pthread_cond_wait(&args->cond, &args->mutex);
    }
    pthread_mutex_unlock(&args->mutex);

    if(core->opt.verbosity > 1){
        fprintf(stderr, "[%s::%.3f*%.2f] Signal got by post-processor thread!\n", __func__,
                realtime() - realtime0, cputime() / (realtime() - realtime0));
    }

    //output and free
    output_db(core, db);
    free_db_tmp(db);
    free_db(db);
    free(args);
    pthread_exit(0);
}

//todo : need to print error message and arg check with respect to eventalign
int meth_main(int argc, char* argv[], int8_t mode) {

    double realtime0 = realtime();

    //signal(SIGSEGV, sig_handler);

    const char* optstring = "r:b:g:t:B:K:v:hV";
    int longindex = 0;
    int32_t c = -1;

    char* bamfilename = NULL;
    char* fastafile = NULL;
    char* fastqfile = NULL;
    char *tmpfile = NULL;

    FILE *fp_help = stderr;

    opt_t opt;
    init_opt(&opt); //initialise options to defaults

    //parse the user args
    while ((c = getopt_long(argc, argv, optstring, long_options, &longindex)) >= 0) {
        if (c == 'r') {
            fastqfile = optarg;
        } else if (c == 'b') {
            bamfilename = optarg;
        } else if (c == 'g') {
            fastafile = optarg;
        } else if (c == 'B') {
            opt.batch_size_bases = mm_parse_num(optarg);
            if(opt.batch_size_bases<=0){
                ERROR("%s","Maximum number of bases should be larger than 0.");
                exit(EXIT_FAILURE);
            }
        } else if (c == 'K') {
            opt.batch_size = atoi(optarg);
            if (opt.batch_size < 1) {
                ERROR("Batch size should larger than 0. You entered %d",opt.batch_size);
                exit(EXIT_FAILURE);
            }
        } else if (c == 't') {
            opt.num_thread = atoi(optarg);
            if (opt.num_thread < 1) {
                ERROR("Number of threads should larger than 0. You entered %d", opt.num_thread);
                exit(EXIT_FAILURE);
            }
        }
        else if (c=='v'){
            opt.verbosity = atoi(optarg);
        }
        else if (c=='V'){
            fprintf(stderr,"F5C %s\n",F5C_VERSION);
            exit(EXIT_SUCCESS);
        }
        else if (c=='h'){
            fp_help = stdout;
        }
        else if (c == 0 && longindex == 9) {
            opt.min_mapq = atoi(optarg); //todo : check whether this is between 0 and 60
        } else if (c == 0 && longindex == 10) { //consider secondary mappings or not
            yes_or_no(&opt, F5C_SECONDARY_YES, longindex, optarg, 1);
        } else if (c == 0 && longindex == 11) { //custom model file
            opt.model_file = optarg;
        } else if (c == 0 && longindex == 12) {
            yes_or_no(&opt, F5C_SKIP_UNREADABLE, longindex, optarg, 1);
        } else if (c == 0 && longindex == 13) {
            yes_or_no(&opt, F5C_PRINT_EVENTS, longindex, optarg, 1);
        } else if (c == 0 && longindex == 14) {
            yes_or_no(&opt, F5C_PRINT_BANDED_ALN, longindex, optarg, 1);
        } else if (c == 0 && longindex == 15) {
            yes_or_no(&opt, F5C_PRINT_SCALING, longindex, optarg, 1);
        } else if (c == 0 && longindex == 16) {
            yes_or_no(&opt, F5C_PRINT_RAW, longindex, optarg, 1);
        } else if (c == 0 && longindex == 17) {
#ifdef HAVE_CUDA
            yes_or_no(&opt, F5C_DISABLE_CUDA, longindex, optarg, 1);
#else
            WARNING("%s", "disable-cuda has no effect when compiled for the CPU");
#endif
        } else if(c == 0 && longindex == 18){
            opt.cuda_block_size = atoi(optarg); //todo : warnining for cpu only mode, check limits
        }else if(c == 0 && longindex == 19){ //debug break
            //yes_or_no(&opt, F5C_DEBUG_BRK, longindex, optarg, 1);
            opt.debug_break = atoi(optarg);
        }else if(c == 0 && longindex == 20){ //sectional benchmark todo : warning for gpu mode
            yes_or_no(&opt, F5C_SEC_PROF, longindex, optarg, 1);
        }else if(c == 0 && longindex == 21){ //cuda todo : warning for cpu mode, error check
            opt.cuda_max_readlen = atof(optarg);
        }else if(c == 0 && longindex == 22){ //cuda todo : warning for cpu mode, warning for dynamic cuda malloc mode, error check
            opt.cuda_avg_events_per_kmer = atof(optarg);
        }else if(c == 0 && longindex == 23){ //cuda todo : warning for cpu mode, error check
            opt.cuda_max_avg_events_per_kmer = atof(optarg);
        }else if(c == 0 && longindex == 24){
            opt.cuda_dev_id = atoi(optarg);
        } else if(c == 0 && longindex == 25){ //todo : warning for CPU mode, warning for dynamic malloc mode
            opt.cuda_mem_frac = atof(optarg);
        } else if(c == 0 && longindex == 26){ //check for empty strings
            tmpfile = optarg;
        } else if(c == 0 && longindex == 27){ 
            if(tmpfile==NULL){
                WARNING("%s", "ultra-thresh has no effect without skip-ultra");
            }
            opt.ultra_thresh = atoi(optarg);
        } else if(c == 0 && longindex == 28){ //write the raw dump of the fast5 files
            yes_or_no(&opt, F5C_WR_RAW_DUMP, longindex, optarg, 1);
        } else if(c == 0 && longindex == 29){ //read the raw dump of the fast5 files
            yes_or_no(&opt, F5C_RD_RAW_DUMP, longindex, optarg, 1);
        }        
    }

    if (fastqfile == NULL || bamfilename == NULL || fastafile == NULL || fp_help == stdout) {
        fprintf(
            fp_help,
            "Usage: f5c %s [OPTIONS] -r reads.fa -b alignments.bam -g genome.fa\n",mode==1 ? "eventalign" : "call-methylation");
        fprintf(fp_help,"   -r FILE                    fastq/fasta read file\n");
        fprintf(fp_help,"   -b FILE                    sorted bam file\n");
        fprintf(fp_help,"   -g FILE                    reference genome\n");
        fprintf(fp_help,"   -t INT                     number of threads [%d]\n",opt.num_thread);
        fprintf(fp_help,"   -K INT                     batch size (max number of reads loaded at once) [%d]\n",opt.batch_size);
        fprintf(fp_help,"   -B FLOAT[K/M/G]            max number of bases loaded at once [%.1fM]\n",opt.batch_size_bases/(float)(1000*1000));
        fprintf(fp_help,"   -h                         help\n");
        fprintf(fp_help,"   --min-mapq INT             minimum mapping quality [%d]\n",opt.min_mapq);
        fprintf(fp_help,"   --secondary=yes|no         consider secondary mappings or not [%s]\n",(opt.flag&F5C_SECONDARY_YES)?"yes":"no");
        fprintf(fp_help,"   --skip-unreadable=yes|no   skip any unreadable fast5 or terminate program [%s]\n",(opt.flag&F5C_SKIP_UNREADABLE?"yes":"no"));
        fprintf(fp_help,"   --verbose INT              verbosity level [%d]\n",opt.verbosity);
        fprintf(fp_help,"   --version                  print version\n");
#ifdef HAVE_CUDA
        fprintf(fp_help,"   --disable-cuda=yes|no      disable running on CUDA [%s]\n",(opt.flag&F5C_DISABLE_CUDA?"yes":"no"));
        fprintf(fp_help,"   --cuda-dev-id INT          CUDA device ID to run kernels on [%d]\n",opt.cuda_dev_id);
        fprintf(fp_help,"   --cuda-max-lf FLOAT        reads with length <= cuda-max-lf*avg_readlen on GPU, rest on CPU [%.1f]\n",opt.cuda_max_readlen);
        fprintf(fp_help,"   --cuda-avg-epk FLOAT       average number of events per kmer - for allocating GPU arrays [%.1f]\n",opt.cuda_avg_events_per_kmer);
        fprintf(fp_help,"   --cuda-max-epk FLOAT       reads with events per kmer <= cuda_max_epk on GPU, rest on CPU [%.1f]\n",opt.cuda_max_avg_events_per_kmer);
#endif


        fprintf(fp_help,"advanced options:\n");
        fprintf(fp_help,"   --kmer-model FILE          custom k-mer model file\n");
        fprintf(fp_help,"   --print-events=yes|no      prints the event table\n");
        fprintf(fp_help,"   --print-banded-aln=yes|no  prints the event alignment\n");
        fprintf(fp_help,"   --print-scaling=yes|no     prints the estimated scalings\n");
        fprintf(fp_help,"   --print-raw=yes|no         prints the raw signal\n");
        fprintf(fp_help,"   --debug-break [INT]        break after processing the specified batch\n");
        fprintf(fp_help,"   --profile-cpu=yes|no       process section by section (used for profiling on CPU)\n");
        fprintf(fp_help,"   --skip-ultra FILE          skip ultra long reads and write those entries to the bam file provided as the argument\n");
        fprintf(fp_help,"   --ultra-thresh [INT]       threshold to skip ultra long reads [%ld]\n",opt.ultra_thresh);
        fprintf(fp_help,"   --write-dump=yes|no        write the fast5 dump to a file or not\n");
        fprintf(fp_help,"   --read-dump=yes|no         read from a fast5 dump file or not\n");
#ifdef HAVE_CUDA
        fprintf(fp_help,"   - cuda-mem-frac FLOAT      Fraction of free GPU memory to allocate [0.9 (0.7 for tegra)]\n");
        fprintf(fp_help,"   --cuda-block-size\n");
#endif
        if(fp_help == stdout){
            exit(EXIT_SUCCESS);
        }
        exit(EXIT_FAILURE);
    }

    //initialise the core data structure
    core_t* core = init_core(bamfilename, fastafile, fastqfile, tmpfile, opt,realtime0,mode);

    #ifdef ESL_LOG_SUM
        p7_FLogsumInit();
    #endif

    //print the header
    if(mode==0){
        fprintf(stdout, "chromosome\tstart\tend\tread_name\t"
                                 "log_lik_ratio\tlog_lik_methylated\tlog_lik_unmethylated\t"
                                 "num_calling_strands\tnum_cpgs\tsequence\n");
    }
    else if(mode==1){
        if(core->event_summary_fp!=NULL){
            fprintf(core->event_summary_fp,"read_index\tread_name\tfast5_path\tmodel_name\tstrand\tnum_events\t");
            fprintf(core->event_summary_fp,"num_steps\tnum_skips\tnum_stays\ttotal_duration\tshift\tscale\tdrift\tvar\n");
        }
        emit_event_alignment_tsv_header(stdout, 1, 0);
    }
    int32_t counter=0;

 #ifdef IO_PROC_NO_INTERLEAVE   //If input, processing and output are not interleaved (serial mode)

    //initialise a databatch
    db_t* db = init_db(core);

    ret_status_t status = {core->opt.batch_size,core->opt.batch_size_bases};
    while (status.num_reads >= core->opt.batch_size || status.num_bases>=core->opt.batch_size_bases) {

        //load a databatch
        status = load_db(core, db);

        fprintf(stderr, "[%s::%.3f*%.2f] %d Entries (%.1fM bases) loaded\n", __func__,
                realtime() - realtime0, cputime() / (realtime() - realtime0),
                status.num_reads,status.num_bases/(1000.0*1000.0));

        //process a databatch
        process_db(core, db);

        fprintf(stderr, "[%s::%.3f*%.2f] %d Entries (%.1fM bases) processed\n", __func__,
                realtime() - realtime0, cputime() / (realtime() - realtime0),
                status.num_reads,status.num_bases/(1000.0*1000.0));

        //output print
        output_db(core, db);

        //free temporary
        free_db_tmp(db);

        if(opt.debug_break==counter){
            break;
        }
        counter++;
    }

    //free the databatch
    free_db(db);

#else   //input, processing and output are interleaved (default)

    ret_status_t status = {core->opt.batch_size,core->opt.batch_size_bases};
    int8_t first_flag_p=0;
    int8_t first_flag_pp=0;
    pthread_t tid_p; //process thread
    pthread_t tid_pp; //post-process thread


    while (status.num_reads >= core->opt.batch_size || status.num_bases>=core->opt.batch_size_bases) {

        //init and load a databatch
        db_t* db = init_db(core);
        status = load_db(core, db);

        fprintf(stderr, "[%s::%.3f*%.2f] %d Entries (%.1fM bases) loaded\n", __func__,
                realtime() - realtime0, cputime() / (realtime() - realtime0),
                status.num_reads,status.num_bases/(1000.0*1000.0));

        if(first_flag_p){ //if not the first time of the "process" wait for the previous "process"
            int ret = pthread_join(tid_p, NULL);
            NEG_CHK(ret);
            if(opt.verbosity>1){
                fprintf(stderr, "[%s::%.3f*%.2f] Joined to processor thread %ld\n", __func__,
                realtime() - realtime0, cputime() / (realtime() - realtime0),
                tid_p);
            }
        }
        first_flag_p=1;

        //set up args
        pthread_arg2_t *pt_arg = (pthread_arg2_t*)malloc(sizeof(pthread_arg2_t));
        pt_arg->core=core;
        pt_arg->db=db;
        pt_arg->cond = PTHREAD_COND_INITIALIZER;
        pt_arg->mutex = PTHREAD_MUTEX_INITIALIZER;
        pt_arg->finished = 0;

        //process thread launch
        int ret = pthread_create(&tid_p, NULL, pthread_processor,
                                (void*)(pt_arg));
        NEG_CHK(ret);
        if(opt.verbosity>1){
            fprintf(stderr, "[%s::%.3f*%.2f] Spawned processor thread %ld\n", __func__,
                realtime() - realtime0, cputime() / (realtime() - realtime0),
                tid_p);
        }

        if(first_flag_pp){ //if not the first time of the post-process wait for the previous post-process
            int ret = pthread_join(tid_pp, NULL);
            NEG_CHK(ret);
            if(opt.verbosity>1){
                fprintf(stderr, "[%s::%.3f*%.2f] Joined to post-processor thread %ld\n", __func__,
                realtime() - realtime0, cputime() / (realtime() - realtime0),
                tid_pp);
            }
        }
        first_flag_pp=1;

        //post-process thread launch (output and freeing thread)
        ret = pthread_create(&tid_pp, NULL, pthread_post_processor,
                                (void*)(pt_arg));
        NEG_CHK(ret);
        if(opt.verbosity>1){
            fprintf(stderr, "[%s::%.3f*%.2f] Spawned post-processor thread %ld\n", __func__,
                realtime() - realtime0, cputime() / (realtime() - realtime0),
                tid_pp);
        }

        if(opt.debug_break==counter){
            break;
        }
        counter++;
    }

    //final round
    int ret = pthread_join(tid_p, NULL);
    NEG_CHK(ret);
    if(opt.verbosity>1){
        fprintf(stderr, "[%s::%.3f*%.2f] Joined to last processor thread %ld\n", __func__,
                realtime() - realtime0, cputime() / (realtime() - realtime0),
                tid_p);
    }
    ret = pthread_join(tid_pp, NULL);
    NEG_CHK(ret);
    if(opt.verbosity>1){
    fprintf(stderr, "[%s::%.3f*%.2f] Joined to last post-processor thread %ld\n", __func__,
                realtime() - realtime0, cputime() / (realtime() - realtime0),
                tid_pp);
    }


#endif


    fprintf(stderr, "\n[%s] total entries: %ld, qc fail: %ld, could not calibrate: %ld, no alignment: %ld, bad fast5: %ld",
             __func__,core->total_reads, core->qc_fail_reads, core->failed_calibration_reads, core->failed_alignment_reads, core->bad_fast5_file);
    fprintf(stderr,"\n[%s] total bases: %.1f Mbases",__func__,core->sum_bases/(float)(1000*1000));

    fprintf(stderr, "\n[%s] Data loading time: %.3f sec", __func__,core->load_db_time);
    fprintf(stderr, "\n[%s]     - bam load time: %.3f sec", __func__, core->db_bam_time);
    fprintf(stderr, "\n[%s]     - fasta load time: %.3f sec", __func__, core->db_fasta_time);
    fprintf(stderr, "\n[%s]     - fast5 load time: %.3f sec", __func__, core->db_fast5_time);
    fprintf(stderr, "\n[%s]         - fast5 open time: %.3f sec", __func__, core->db_fast5_open_time);
    fprintf(stderr, "\n[%s]         - fast5 read time: %.3f sec", __func__, core->db_fast5_read_time);

    fprintf(stderr, "\n[%s] Data processing time: %.3f sec", __func__,core->process_db_time);

    if((core->opt.flag&F5C_SEC_PROF) || (!(core->opt.flag & F5C_DISABLE_CUDA))){
        fprintf(stderr, "\n[%s]     - Events time: %.3f sec",
                __func__, core->event_time);
        fprintf(stderr, "\n[%s]     - Alignment time: %.3f sec",
                __func__, core->align_time);
        #ifdef HAVE_CUDA
            if (!(core->opt.flag & F5C_DISABLE_CUDA)) {
                fprintf(stderr, "\n[%s]           -cpu preprocess time: %.3f sec",
                    __func__, core->align_cuda_preprocess);
            #ifdef CUDA_DYNAMIC_MALLOC
                fprintf(stderr, "\n[%s]           -cuda malloc time: %.3f sec",
                    __func__, core->align_cuda_malloc);
            #endif
                fprintf(stderr, "\n[%s]           -cuda data transfer time: %.3f sec",
                    __func__, core->align_cuda_memcpy);
                fprintf(stderr, "\n[%s]           -cuda kernel time: %.3f sec",
                    __func__, core->align_kernel_time);
                fprintf(stderr, "\n[%s]                -align-pre kernel only time: %.3f sec",
                    __func__, core->align_pre_kernel_time);
                fprintf(stderr, "\n[%s]                -align-core kernel only time: %.3f sec",
                    __func__, core->align_core_kernel_time);
                fprintf(stderr, "\n[%s]                -align-post kernel only time: %.3f sec",
                    __func__, core->align_post_kernel_time);

                fprintf(stderr, "\n[%s]           -cpu postprocess time: %.3f sec",
                    __func__, core->align_cuda_postprocess);
                fprintf(stderr, "\n[%s]           -additional cpu processing time (load imbalance): %.3f sec",
                    __func__, core->extra_load_cpu);
            }
        #endif
        fprintf(stderr, "\n[%s]     - Estimate scaling time: %.3f sec",
                __func__, core->est_scale_time);
        fprintf(stderr, "\n[%s]     - HMM time: %.3f sec",
                __func__, core->meth_time);

    }

    fprintf(stderr,"\n");

    if(core->ultra_long_skipped>0){
        assert(tmpfile!=NULL);
        WARNING("%ld ultra long reads (>%.1f kbases) were skipped.",core->ultra_long_skipped,core->opt.ultra_thresh/1000.0);
        fprintf(stderr," Please run samtools index on '%s' followed by f5c with a larger -B on the CPU.\n",tmpfile);
    }


#ifndef IO_PROC_NO_INTERLEAVE
    if((core->load_db_time - core->process_db_time) > (core->process_db_time*0.2) ){
        INFO("Performance bounded by file I/O. File I/O took %.3f sec than processing",core->load_db_time - core->process_db_time);
    }
#endif

    //free the core data structure
    free_core(core);


    return 0;
}
```
