## Supplementary material for "GPU Accelerated Adaptive Banded Event Alignment for Rapid Comparative Nanopore Signal Analysis": f5c-v0.2-beta-release: model.c

```
#include "model.h"
#include "f5c.h"
#include "f5cmisc.h"
#include <stdint.h>
#include <stdio.h>
#include <stdlib.h>

//#define DEBUG_MODEL_PRINT 1

void read_model(model_t* model, const char* file) {
    FILE* fp = fopen(file, "r");
    F_CHK(fp, file);

    //these two are discarded from the model. hollow vars
    char kmer[10];
    float weight;

    //buffers for geline
    char* buffer =
        (char*)malloc(sizeof(char) * (100)); //READ+newline+nullcharacter
    MALLOC_CHK(buffer);
    size_t bufferSize = 100;
    ssize_t readlinebytes = 0;

    uint32_t num_k = 0;
    uint32_t i = 0;

    while ((readlinebytes = getline(&buffer, &bufferSize, fp)) != -1) {
        if (buffer[0] == '#' ||
            strcmp(
                buffer,
                "kmer\tlevel_mean\tlevel_stdv\tsd_mean\tsd_stdv\tweight\n") ==
                0 ||
            buffer[0] == '\n' || buffer[0] == '\r') { //comments and header
            //todo : (make generic)
            //fprintf(stderr, "%s\n", buffer);
            continue;
        } else {
            //as sd_mean and sd_stdv seems not to be used just read to the summy weight
            #ifdef LOAD_SD_MEANSSTDV
                int32_t ret =
                    sscanf(buffer, "%s\t%f\t%f\t%f\t%f\t%f", kmer,
                        &model[num_k].level_mean, &model[num_k].level_stdv,
                        &model[num_k].sd_mean, &model[num_k].sd_stdv, &weight);
            #else
                int32_t ret =
                    sscanf(buffer, "%s\t%f\t%f\t%f\t%f\t%f", kmer,
                        &model[num_k].level_mean, &model[num_k].level_stdv,
                        &weight, &weight, &weight);
            #endif
            #ifdef CACHED_LOG
                model[num_k].level_log_stdv=log(model[num_k].level_stdv);
            #endif
            num_k++;
            if (ret != 6) {
                ERROR("File %s is corrupted at line %d", file, i);
            }
            if (num_k > NUM_KMER) {
                ERROR("File %s has too many entries. Expected %d kmers in the "
                      "model, but file had more than that",
                      file, NUM_KMER);
                exit(EXIT_FAILURE);
            }
        }
        i++;
    }

    if (num_k != NUM_KMER) {
        ERROR("File %s prematurely ended. Expected %d kmers in the model, but "
              "file had only%d",
              file, NUM_KMER, num_k);
        exit(EXIT_FAILURE);
    }

#ifdef DEBUG_MODEL_PRINT
    i = 0;
    fprintf(stderr, "level_mean\tlevel_stdv\tsd_mean\tsd_stdv\n");
    for (i = 0; i < NUM_KMER; i++) {
        fprintf(stderr, "%f\t%f\t%f\t%f\n", model[i].level_mean,
                model[i].level_stdv, model[i].sd_mean, model[i].sd_stdv);
    }
#endif

    free(buffer);
    fclose(fp);
}

//this function can be made more efficient by setting the address to the global variable
void set_model(model_t* model) {
    uint32_t i = 0;
    for (i = 0; i < NUM_KMER; i++) {
        model[i].level_mean =
            r9_4_450bps_nucleotide_6mer_template_model_builtin_data[i * 4 + 0];
        model[i].level_stdv =
            r9_4_450bps_nucleotide_6mer_template_model_builtin_data[i * 4 + 1];
    #ifdef LOAD_SD_MEANSSTDV
        model[i].sd_mean =
            r9_4_450bps_nucleotide_6mer_template_model_builtin_data[i * 4 + 2];
        model[i].sd_stdv =
            r9_4_450bps_nucleotide_6mer_template_model_builtin_data[i * 4 + 3];
    #endif
    #ifdef CACHED_LOG
        model[i].level_log_stdv=log(model[i].level_stdv);
    #endif
    }
#ifdef DEBUG_MODEL_PRINT
    i = 0;
    fprintf(stderr, "level_mean\tlevel_stdv\tsd_mean\tsd_stdv\n");
    for (i = 0; i < NUM_KMER; i++) {
        fprintf(stderr, "%f\t%f\t%f\t%f\n", model[i].level_mean,
                model[i].level_stdv, model[i].sd_mean, model[i].sd_stdv);
    }
#endif
}

//todo : this function can be made more efficient by setting the address to the global variable
//todo : duplicate function can be removed
void set_cpgmodel(model_t* model) {
    uint32_t i = 0;
    for (i = 0; i < NUM_KMER_METH; i++) {
        model[i].level_mean =
            r9_4_450bps_cpg_6mer_template_model_builtin_data[i * 4 + 0];
        model[i].level_stdv =
            r9_4_450bps_cpg_6mer_template_model_builtin_data[i * 4 + 1];
    #ifdef LOAD_SD_MEANSSTDV
        model[i].sd_mean =
            r9_4_450bps_cpg_6mer_template_model_builtin_data[i * 4 + 2];
        model[i].sd_stdv =
            r9_4_450bps_cpg_6mer_template_model_builtin_data[i * 4 + 3];
    #endif
    #ifdef CACHED_LOG
        model[i].level_log_stdv=log(model[i].level_stdv);
    #endif
    }
#ifdef DEBUG_MODEL_PRINT
    i = 0;
    fprintf(stderr, "level_mean\tlevel_stdv\tsd_mean\tsd_stdv\n");
    for (i = 0; i < NUM_KMER; i++) {
        fprintf(stderr, "%f\t%f\t%f\t%f\n", model[i].level_mean,
                model[i].level_stdv, model[i].sd_mean, model[i].sd_stdv);
    }
#endif
}
```
