## Supplementary material for "GPU Accelerated Adaptive Banded Event Alignment for Rapid Comparative Nanopore Signal Analysis": f5c-v0.2-beta-release: model.h

```
#ifndef MODEL_H
#define MODEL_H

//extracted from nanopolish

static float r9_4_450bps_nucleotide_6mer_template_model_builtin_data[] = {
    86.48634,  1.51785, 0.94148, 0.60936, // AAAAAA
    83.94884,  1.51785, 1.07705, 0.74561, // AAAAAC
    85.47537,  1.51785, 0.95343, 0.62100, // AAAAAG
    84.42391,  1.51785, 1.10608, 0.77595, // AAAAAT
    77.09728,  1.70501, 1.29686, 0.93621, // AAAACA
    75.70630,  1.70501, 1.31939, 0.96072, // AAAACC
    76.92824,  1.70501, 1.30207, 0.94186, // AAAACG
    76.81141,  1.70501, 1.17932, 0.81186, // AAAACT
    84.56326,  1.53757, 0.93295, 0.60480, // AAAAGA
    82.76070,  1.53757, 0.96369, 0.63495, // AAAAGC
    83.76557,  1.53757, 0.93570, 0.60748, // AAAAGG
    82.86687,  1.53757, 1.07858, 0.75182, // AAAAGT
    76.49238,  1.54253, 1.14524, 0.84930, // AAAATA
    74.62771,  1.54253, 1.11274, 0.81340, // AAAATC
    75.94387,  1.54253, 1.09532, 0.79438, // AAAATG
    75.80068,  1.54253, 1.28978, 1.01505, // AAAATT
    88.25543,  1.96203, 1.48065, 1.34391, // AAACAA
    86.68990,  1.96203, 1.27700, 1.07641, // AAACAC
    87.32014,  1.96203, 1.40026, 1.23597, // AAACAG
    87.45209,  1.96203, 1.31446, 1.12412, // AAACAT
    84.58290,  1.83613, 1.49328, 1.42524, // AAACCA
    83.47830,  1.83613, 1.40063, 1.29469, // AAACCC
    84.50585,  1.83613, 1.45041, 1.36432, // AAACCG
    84.41226,  1.83613, 1.38692, 1.27573, // AAACCT
    87.18447,  1.55453, 1.30654, 1.16200, // AAACGA
    85.63648,  1.55453, 1.21094, 1.03683, // AAACGC
    86.33688,  1.55453, 1.29411, 1.14546, // AAACGG
    86.06472,  1.55453, 1.24344, 1.07884, // AAACGT
    84.13415,  1.58806, 1.28615, 1.13271, // AAACTA
    82.01609,  1.58806, 1.12216, 0.92313, // AAACTC
    83.27923,  1.58806, 1.22077, 1.04745, // AAACTG
    82.58670,  1.58806, 0.95274, 0.72217, // AAACTT
    84.35344,  1.46160, 0.97542, 0.66709, // AAAGAA
    80.39255,  1.46160, 1.17615, 0.88326, // AAAGAC
    83.61615,  1.46160, 1.00518, 0.69785, // AAAGAG
    81.85876,  1.46160, 1.16532, 0.87110, // AAAGAT
    79.14905,  1.42710, 1.04709, 0.74755, // AAAGCA
    77.56133,  1.42710, 1.04196, 0.74206, // AAAGCC
    78.87292,  1.42710, 0.98261, 0.67957, // AAAGCG
    78.67537,  1.42710, 0.93431, 0.63008, // AAAGCT
    83.20780,  1.40989, 0.98410, 0.67678, // AAAGGA
    80.68861,  1.40989, 1.02292, 0.71723, // AAAGGC
    82.25816,  1.40989, 0.93946, 0.63127, // AAAGGG
    81.70047,  1.40989, 1.07170, 0.76914, // AAAGGT
    77.65304,  1.48703, 1.00723, 0.72430, // AAAGTA
    75.18822,  1.48703, 1.20345, 0.94594, // AAAGTC
    76.72314,  1.48703, 1.07134, 0.79454, // AAAGTG
    76.25087,  1.48703, 1.30532, 1.06856, // AAAGTT
    82.70040,  1.98024, 1.64571, 1.60071, // AAATAA
    80.33955,  1.98024, 1.57619, 1.50036, // AAATAC
    81.96701,  1.98024, 1.53106, 1.43639, // AAATAG
    81.67422,  1.98024, 1.71069, 1.69644, // AAATAT
    76.03344,  2.22770, 2.11887, 2.65176, // AAATCA
    74.91406,  2.22770, 2.03424, 2.49449, // AAATCC
    76.19823,  2.22770, 2.15565, 2.72110, // AAATCG
    76.11483,  2.22770, 2.14848, 2.70753, // AAATCT
    80.08087,  1.67617, 1.56632, 1.53101, // AAATGA
    77.61264,  1.67617, 1.61261, 1.59937, // AAATGC
    79.18708,  1.67617, 1.64897, 1.65377, // AAATGG
    78.44148,  1.67617, 1.66765, 1.68195, // AAATGT
    70.76241,  2.00867, 1.67811, 1.58287, // AAATTA
    67.99951,  2.00867, 1.83641, 1.81206, // AAATTC
    70.12356,  2.00867, 1.71772, 1.63924, // AAATTG
    69.30979,  2.00867, 1.73599, 1.66547, // AAATTT
    107.92456, 2.19132, 1.79213, 1.53404, // AACAAA
    104.86114, 2.19132, 1.80497, 1.55056, // AACAAC
    106.83318, 2.19132, 1.87422, 1.64064, // AACAAG
    106.15898, 2.19132, 1.96452, 1.76063, // AACAAT
    99.65940,  1.74166, 1.36504, 1.00221, // AACACA
    98.13202,  1.74166, 1.33763, 0.97218, // AACACC
    99.53501,  1.74166, 1.32214, 0.95534, // AACACG
    99.48971,  1.74166, 1.43867, 1.08438, // AACACT
    105.26973, 2.28987, 1.92656, 1.69338, // AACAGA
    102.31212, 2.28987, 1.77407, 1.49637, // AACAGC
    104.13857, 2.28987, 1.74932, 1.46516, // AACAGG
    103.70156, 2.28987, 1.92458, 1.69077, // AACAGT
    99.05354,  2.17760, 1.55825, 1.29235, // AACATA
    97.47739,  2.17760, 1.61265, 1.36061, // AACATC
    98.40492,  2.17760, 1.58327, 1.32361, // AACATG
    98.80829,  2.17760, 1.84242, 1.66153, // AACATT
    107.94456, 1.88264, 1.53183, 1.35379, // AACCAA
    105.07232, 1.88264, 1.51285, 1.32871, // AACCAC
    107.00329, 1.88264, 1.48473, 1.29184, // AACCAG
    106.26763, 1.88264, 1.65588, 1.52152, // AACCAT
    103.93410, 1.75332, 1.29734, 1.06459, // AACCCA
    102.22953, 1.75332, 1.24222, 0.99747, // AACCCC
    103.83467, 1.75332, 1.34040, 1.11804, // AACCCG
    103.20345, 1.75332, 1.47108, 1.28546, // AACCCT
    106.59327, 1.90443, 1.60306, 1.39462, // AACCGA
    103.84816, 1.90443, 1.47383, 1.22942, // AACCGC
    105.63889, 1.90443, 1.54235, 1.31615, // AACCGG
    104.87454, 1.90443, 1.68020, 1.49648, // AACCGT
    103.12480, 1.83098, 1.49866, 1.19909, // AACCTA
    100.84405, 1.83098, 1.52449, 1.23022, // AACCTC
    102.21601, 1.83098, 1.48098, 1.17793, // AACCTG
    101.83993, 1.83098, 1.69160, 1.43795, // AACCTT
    103.12972, 2.24722, 1.81869, 1.59373, // AACGAA
    98.44624,  2.24722, 1.72440, 1.47142, // AACGAC
    102.04100, 2.24722, 1.78587, 1.55078, // AACGAG
    100.50147, 2.24722, 1.95330, 1.77391, // AACGAT
    98.63875,  2.02317, 1.43135, 1.10889, // AACGCA
    96.96738,  2.02317, 1.50183, 1.19180, // AACGCC
    98.42319,  2.02317, 1.53638, 1.23315, // AACGCG
    98.63125,  2.02317, 1.58425, 1.29123, // AACGCT
    102.87807, 2.22668, 1.82532, 1.59411, // AACGGA
    99.63194,  2.22668, 1.69529, 1.42684, // AACGGC
    101.89877, 2.22668, 1.74356, 1.48821, // AACGGG
    101.25285, 2.22668, 1.88906, 1.67833, // AACGGT
    97.84676,  2.64556, 1.75347, 1.45050, // AACGTA
    96.07217,  2.64556, 1.79988, 1.50847, // AACGTC
    97.18586,  2.64556, 1.75924, 1.45768, // AACGTG
    97.66835,  2.64556, 2.09559, 1.89511, // AACGTT
    101.57266, 1.69847, 1.59479, 1.18964, // AACTAA
    98.06116,  1.69847, 1.40164, 0.98019, // AACTAC
    100.42033, 1.69847, 1.58045, 1.17362, // AACTAG
    99.67084,  1.69847, 1.51928, 1.10615, // AACTAT
    93.66733,  1.86136, 1.57047, 1.41151, // AACTCA
    91.54694,  1.86136, 1.59310, 1.44213, // AACTCC
    93.52490,  1.86136, 1.54560, 1.37810, // AACTCG
    93.05050,  1.86136, 1.57869, 1.42260, // AACTCT
    98.74313,  1.64486, 1.41332, 1.09402, // AACTGA
    95.15271,  1.64486, 1.32356, 0.99148, // AACTGC
    97.38793,  1.64486, 1.38744, 1.06412, // AACTGG
    96.27599,  1.64486, 1.35980, 1.03248, // AACTGT
    88.93059,  1.46961, 1.29167, 0.96950, // AACTTA
    85.82739,  1.46961, 1.18574, 0.85272, // AACTTC
    88.07340,  1.46961, 1.18719, 0.85428, // AACTTG
    87.03033,  1.46961, 1.07811, 0.73929, // AACTTT
    79.83489,  1.82916, 1.12430, 0.71403, // AAGAAA
    76.76366,  1.82916, 1.35326, 0.94291, // AAGAAC
    79.01269,  1.82916, 1.13240, 0.72177, // AAGAAG
    77.18936,  1.82916, 1.34279, 0.93199, // AAGAAT
    69.06004,  2.14481, 1.64737, 1.23807, // AAGACA
    67.42395,  2.14481, 1.63794, 1.22745, // AAGACC
    68.61974,  2.14481, 1.66378, 1.25661, // AAGACG
    68.60189,  2.14481, 1.59024, 1.17423, // AAGACT
    78.23783,  1.82854, 1.09014, 0.73888, // AAGAGA
    76.26280,  1.82854, 1.15687, 0.80774, // AAGAGC
    77.26173,  1.82854, 1.16133, 0.81242, // AAGAGG
    76.41294,  1.82854, 1.18627, 0.83874, // AAGAGT
    68.66484,  1.85459, 1.47689, 1.21422, // AAGATA
    67.31463,  1.85459, 1.34951, 1.06057, // AAGATC
    68.11579,  1.85459, 1.38770, 1.10590, // AAGATG
    67.98461,  1.85459, 1.36842, 1.08293, // AAGATT
    87.07535,  1.87706, 1.47547, 1.22356, // AAGCAA
    84.83646,  1.87706, 1.20873, 0.90724, // AAGCAC
    86.01732,  1.87706, 1.40134, 1.13252, // AAGCAG
    85.60647,  1.87706, 1.26641, 0.97295, // AAGCAT
    82.42552,  1.70618, 1.43823, 1.25952, // AAGCCA
    81.30370,  1.70618, 1.39259, 1.20004, // AAGCCC
    82.45521,  1.70618, 1.44790, 1.27224, // AAGCCG
    82.33154,  1.70618, 1.41867, 1.23391, // AAGCCT
    84.88051,  1.56014, 1.28937, 1.05556, // AAGCGA
    81.78064,  1.56014, 1.19997, 0.94770, // AAGCGC
    83.72481,  1.56014, 1.31411, 1.08609, // AAGCGG
    82.51428,  1.56014, 1.27584, 1.03900, // AAGCGT
    81.54698,  1.42173, 1.28659, 1.02607, // AAGCTA
    79.38379,  1.42173, 1.14310, 0.85930, // AAGCTC
    80.91313,  1.42173, 1.24512, 0.97686, // AAGCTG
    80.06805,  1.42173, 0.98504, 0.68738, // AAGCTT
    78.78397,  1.83346, 1.17273, 0.78158, // AAGGAA
    73.52237,  1.83346, 1.57040, 1.21112, // AAGGAC
    77.44999,  1.83346, 1.24215, 0.85198, // AAGGAG
    75.14774,  1.83346, 1.48421, 1.11279, // AAGGAT
    73.06399,  1.67690, 1.40742, 1.12077, // AAGGCA
    71.63335,  1.67690, 1.38351, 1.09233, // AAGGCC
    72.82221,  1.67690, 1.31019, 1.00666, // AAGGCG
    72.48132,  1.67690, 1.19811, 0.88029, // AAGGCT
    79.08838,  1.68720, 1.06266, 0.73225, // AAGGGA
    76.17658,  1.68720, 1.23284, 0.91501, // AAGGGC
    77.93815,  1.68720, 1.01128, 0.67978, // AAGGGG
    77.11319,  1.68720, 1.26818, 0.95463, // AAGGGT
    71.73488,  1.76089, 1.19129, 0.85060, // AAGGTA
    69.86706,  1.76089, 1.41274, 1.09848, // AAGGTC
    71.17182,  1.76089, 1.27893, 0.94617, // AAGGTG
    70.98170,  1.76089, 1.45329, 1.14611, // AAGGTT
    77.23804,  2.04805, 1.62302, 1.46888, // AAGTAA
    73.87071,  2.04805, 1.75202, 1.64743, // AAGTAC
    76.10582,  2.04805, 1.62763, 1.47515, // AAGTAG
    75.33943,  2.04805, 1.65682, 1.51500, // AAGTAT
    69.35298,  2.42133, 2.02437, 2.11329, // AAGTCA
    68.05680,  2.42133, 2.20431, 2.40122, // AAGTCC
    69.41673,  2.42133, 2.08053, 2.20184, // AAGTCG
    69.38230,  2.42133, 2.25893, 2.49103, // AAGTCT
    73.35763,  1.99693, 1.64171, 1.59332, // AAGTGA
    69.44204,  1.99693, 1.99644, 2.13670, // AAGTGC
    72.06195,  1.99693, 1.71446, 1.70039, // AAGTGG
    70.51905,  1.99693, 1.97265, 2.09862, // AAGTGT
    64.50249,  2.35636, 2.22607, 2.19760, // AAGTTA
    61.27854,  2.35636, 2.19616, 2.15345, // AAGTTC
    63.79433,  2.35636, 2.18102, 2.13122, // AAGTTG
    62.61819,  2.35636, 2.07986, 1.98468, // AAGTTT
    110.62635, 3.16286, 2.50002, 2.09543, // AATAAA
    108.31980, 3.16286, 2.51648, 2.11615, // AATAAC
    109.93493, 3.16286, 2.40830, 1.98118, // AATAAG
    110.61756, 3.16286, 2.74380, 2.40927, // AATAAT
    105.95822, 3.09766, 2.25473, 1.78717, // AATACA
    105.46224, 3.09766, 2.35632, 1.90929, // AATACC
    105.35066, 3.09766, 2.18793, 1.70833, // AATACG
    107.32040, 3.09766, 2.50950, 2.09848, // AATACT
    108.50323, 3.02439, 2.51831, 2.28269, // AATAGA
    106.94645, 3.02439, 2.38518, 2.10410, // AATAGC
    108.47818, 3.02439, 2.23292, 1.90588, // AATAGG
    109.47666, 3.02439, 2.63960, 2.44959, // AATAGT
    106.66735, 3.75132, 2.60030, 2.14247, // AATATA
    108.00626, 3.75132, 2.80050, 2.39460, // AATATC
    106.31226, 3.75132, 2.64585, 2.19902, // AATATG
    110.24256, 3.75132, 2.98101, 2.62982, // AATATT
    119.70278, 2.97261, 2.56604, 2.71939, // AATCAA
    117.13152, 2.97261, 2.51237, 2.63451, // AATCAC
    118.75132, 2.97261, 2.65432, 2.86093, // AATCAG
    118.52388, 2.97261, 2.63883, 2.83591, // AATCAT
    118.30807, 3.04768, 2.50700, 2.50391, // AATCCA
    116.44033, 3.04768, 2.45317, 2.42369, // AATCCC
    118.47767, 3.04768, 2.60978, 2.65945, // AATCCG
    117.90892, 3.04768, 2.62540, 2.68337, // AATCCT
    119.68484, 3.00027, 2.68172, 2.78683, // AATCGA
    116.67011, 3.00027, 2.53482, 2.56100, // AATCGC
    118.87008, 3.00027, 2.52142, 2.54073, // AATCGG
    118.74556, 3.00027, 2.72499, 2.85455, // AATCGT
    118.85739, 3.10410, 2.68266, 2.69276, // AATCTA
    117.34390, 3.10410, 2.67939, 2.68784, // AATCTC
    118.18109, 3.10410, 2.81019, 2.88705, // AATCTG
    118.77124, 3.10410, 2.81799, 2.89908, // AATCTT
    107.65314, 2.99484, 2.38696, 2.13157, // AATGAA
    103.87279, 2.99484, 2.42541, 2.18329, // AATGAC
    106.80360, 2.99484, 2.17702, 1.85664, // AATGAG
    106.47893, 2.99484, 2.60505, 2.43029, // AATGAT
    106.69589, 3.13415, 2.35365, 2.00915, // AATGCA
    106.34136, 3.13415, 2.38223, 2.04586, // AATGCC
    106.62565, 3.13415, 2.32879, 1.97740, // AATGCG
    108.29426, 3.13415, 2.52659, 2.23460, // AATGCT
    108.49845, 2.93123, 2.42027, 2.12453, // AATGGA
    105.97477, 2.93123, 2.45913, 2.17591, // AATGGC
    107.72020, 2.93123, 2.39505, 2.09142, // AATGGG
    108.60154, 2.93123, 2.57384, 2.32992, // AATGGT
    108.80152, 3.56800, 2.75687, 2.41235, // AATGTA
    110.07764, 3.56800, 2.89791, 2.59982, // AATGTC
    108.79268, 3.56800, 2.87882, 2.57419, // AATGTG
    112.47230, 3.56800, 3.09831, 2.87411, // AATGTT
    107.88063, 2.18027, 1.86753, 1.84138, // AATTAA
    103.82113, 2.18027, 1.70083, 1.60042, // AATTAC
    106.86058, 2.18027, 1.80516, 1.74991, // AATTAG
    105.33516, 2.18027, 1.78368, 1.71878, // AATTAT
    101.47680, 2.09022, 2.07889, 2.03614, // AATTCA
    99.49711,  2.09022, 1.95655, 1.85908, // AATTCC
    101.49873, 2.09022, 1.96630, 1.87299, // AATTCG
    101.55540, 2.09022, 1.83787, 1.69252, // AATTCT
    105.36436, 2.20088, 1.82854, 1.74520, // AATTGA
    101.44947, 2.20088, 1.69701, 1.56033, // AATTGC
    104.50041, 2.20088, 1.71423, 1.58413, // AATTGG
    103.18267, 2.20088, 1.72035, 1.59263, // AATTGT
    97.31223,  1.92511, 1.77757, 1.61419, // AATTTA
    95.12487,  1.92511, 1.63780, 1.42759, // AATTTC
    96.87934,  1.92511, 1.72709, 1.54592, // AATTTG
    96.66201,  1.92511, 1.65003, 1.44361, // AATTTT
    92.71184,  2.70987, 1.59825, 1.19732, // ACAAAA
    89.28397,  2.70987, 1.81392, 1.44767, // ACAAAC
    91.93551,  2.70987, 1.69696, 1.30994, // ACAAAG
    90.32442,  2.70987, 1.85900, 1.50197, // ACAAAT
    83.13679,  2.16112, 1.51177, 1.20984, // ACAACA
    82.21645,  2.16112, 1.54321, 1.24778, // ACAACC
    83.18673,  2.16112, 1.54582, 1.25095, // ACAACG
    82.76916,  2.16112, 1.49251, 1.18680, // ACAACT
    91.01502,  2.81796, 1.60187, 1.24600, // ACAAGA
    88.61560,  2.81796, 1.71362, 1.37863, // ACAAGC
    90.15783,  2.81796, 1.61613, 1.26266, // ACAAGG
    88.88455,  2.81796, 1.87238, 1.57459, // ACAAGT
    82.18981,  2.24890, 1.64608, 1.39736, // ACAATA
    80.65211,  2.24890, 1.97752, 1.83998, // ACAATC
    81.65630,  2.24890, 1.76664, 1.55366, // ACAATG
    81.09841,  2.24890, 2.00648, 1.88054, // ACAATT
    94.96909,  1.22481, 1.05715, 0.80493, // ACACAA
    92.34914,  1.22481, 1.06102, 0.80936, // ACACAC
    94.13071,  1.22481, 1.04348, 0.78937, // ACACAG
    93.39945,  1.22481, 0.97070, 0.70824, // ACACAT
    91.76716,  1.29044, 1.19393, 0.96929, // ACACCA
    90.32106,  1.29044, 1.15189, 0.91854, // ACACCC
    91.49946,  1.29044, 1.10979, 0.86865, // ACACCG
    91.20304,  1.29044, 1.12044, 0.88119, // ACACCT
    94.08571,  1.39798, 1.00994, 0.73926, // ACACGA
    91.54810,  1.39798, 1.08903, 0.82779, // ACACGC
    93.27921,  1.39798, 1.05984, 0.79473, // ACACGG
    92.34441,  1.39798, 1.07297, 0.80954, // ACACGT
    90.18549,  1.56257, 1.12931, 0.82924, // ACACTA
    87.93344,  1.56257, 1.12747, 0.82720, // ACACTC
    89.42717,  1.56257, 1.12618, 0.82579, // ACACTG
    89.02177,  1.56257, 1.09133, 0.78776, // ACACTT
    88.19817,  2.60118, 1.64646, 1.26460, // ACAGAA
    82.73936,  2.60118, 1.92949, 1.60431, // ACAGAC
    87.04281,  2.60118, 1.73953, 1.37333, // ACAGAG
    84.61652,  2.60118, 1.92427, 1.59781, // ACAGAT
    83.01817,  2.08236, 1.42329, 1.19082, // ACAGCA
    81.45033,  2.08236, 1.43105, 1.20057, // ACAGCC
    82.88876,  2.08236, 1.40829, 1.17205, // ACAGCG
    82.80060,  2.08236, 1.52849, 1.32526, // ACAGCT
    87.98536,  2.50147, 1.58798, 1.26495, // ACAGGA
    83.96549,  2.50147, 1.73486, 1.44444, // ACAGGC
    86.51230,  2.50147, 1.55245, 1.22273, // ACAGGG
    85.46983,  2.50147, 1.82887, 1.56343, // ACAGGT
    80.35500,  2.40625, 1.91397, 1.72963, // ACAGTA
    76.81655,  2.40625, 2.09972, 1.98743, // ACAGTC
    78.88416,  2.40625, 1.89939, 1.70991, // ACAGTG
    78.04571,  2.40625, 2.06168, 1.93366, // ACAGTT
    89.15223,  1.95200, 1.62646, 1.31493, // ACATAA
    85.54059,  1.95200, 1.69418, 1.39791, // ACATAC
    88.49882,  1.95200, 1.54416, 1.21641, // ACATAG
    87.28997,  1.95200, 1.68679, 1.38878, // ACATAT
    82.32278,  2.22932, 2.14178, 2.16515, // ACATCA
    80.65778,  2.22932, 2.23229, 2.30384, // ACATCC
    82.23759,  2.22932, 2.00829, 1.96592, // ACATCG
    81.80179,  2.22932, 2.09412, 2.09328, // ACATCT
    86.59766,  2.18506, 1.56195, 1.23545, // ACATGA
    82.80009,  2.18506, 1.79534, 1.52245, // ACATGC
    85.50663,  2.18506, 1.74257, 1.45582, // ACATGG
    83.96140,  2.18506, 1.83102, 1.56805, // ACATGT
    76.51898,  2.43089, 2.12527, 1.99846, // ACATTA
    73.42809,  2.43089, 2.23765, 2.15905, // ACATTC
    75.79693,  2.43089, 2.19356, 2.09555, // ACATTG
    74.69007,  2.43089, 2.04004, 1.87946, // ACATTT
    103.97738, 1.63085, 1.23603, 1.02003, // ACCAAA
    101.47398, 1.63085, 1.45536, 1.30325, // ACCAAC
    103.08320, 1.63085, 1.29590, 1.09503, // ACCAAG
    102.37968, 1.63085, 1.49323, 1.35444, // ACCAAT
    96.35808,  1.46322, 1.05360, 0.74833, // ACCACA
    95.03571,  1.46322, 1.10929, 0.80845, // ACCACC
    96.08636,  1.46322, 1.09260, 0.79027, // ACCACG
    95.95961,  1.46322, 1.15982, 0.86431, // ACCACT
    101.89768, 1.84203, 1.39859, 1.15660, // ACCAGA
    99.59446,  1.84203, 1.44253, 1.21154, // ACCAGC
    100.71046, 1.84203, 1.39727, 1.15497, // ACCAGG
    100.26964, 1.84203, 1.59888, 1.41375, // ACCAGT
    95.04159,  1.79927, 1.30899, 0.96780, // ACCATA
    93.32734,  1.79927, 1.50756, 1.19616, // ACCATC
    94.58491,  1.79927, 1.39952, 1.06991, // ACCATG
    94.37825,  1.79927, 1.64925, 1.36870, // ACCATT
    103.40436, 1.44981, 0.87226, 0.61194, // ACCCAA
    100.50332, 1.44981, 0.96606, 0.71325, // ACCCAC
    102.49018, 1.44981, 0.89341, 0.63433, // ACCCAG
    101.52654, 1.44981, 1.06095, 0.82088, // ACCCAT
    99.35502,  1.40182, 0.89591, 0.64955, // ACCCCA
    97.85843,  1.40182, 0.89646, 0.65015, // ACCCCC
    99.21716,  1.40182, 0.90089, 0.65497, // ACCCCG
    99.09357,  1.40182, 0.87739, 0.62951, // ACCCCT
    101.93673, 1.43192, 0.92684, 0.68212, // ACCCGA
    99.34464,  1.43192, 0.94800, 0.70561, // ACCCGC
    100.73264, 1.43192, 0.90440, 0.65750, // ACCCGG
    100.16651, 1.43192, 1.03949, 0.81019, // ACCCGT
    98.28366,  1.44710, 0.97684, 0.73578, // ACCCTA
    96.48633,  1.44710, 1.13600, 0.92274, // ACCCTC
    97.66747,  1.44710, 0.99337, 0.75453, // ACCCTG
    96.88325,  1.44710, 1.19237, 0.99226, // ACCCTT
    99.07035,  1.82025, 1.36048, 1.11248, // ACCGAA
    94.17784,  1.82025, 1.59801, 1.41621, // ACCGAC
    97.84924,  1.82025, 1.41891, 1.18491, // ACCGAG
    95.99364,  1.82025, 1.67392, 1.51830, // ACCGAT
    94.48171,  1.58093, 1.12144, 0.78696, // ACCGCA
    92.97802,  1.58093, 1.21249, 0.88471, // ACCGCC
    94.16967,  1.58093, 1.18195, 0.85150, // ACCGCG
    94.08148,  1.58093, 1.21946, 0.89235, // ACCGCT
    98.80269,  1.74812, 1.35912, 1.15166, // ACCGGA
    95.47873,  1.74812, 1.45115, 1.27059, // ACCGGC
    97.57446,  1.74812, 1.36074, 1.15372, // ACCGGG
    96.85988,  1.74812, 1.51550, 1.35604, // ACCGGT
    92.62404,  2.13467, 1.44073, 1.19456, // ACCGTA
    90.17103,  2.13467, 1.67103, 1.49214, // ACCGTC
    91.68521,  2.13467, 1.59610, 1.39291, // ACCGTG
    91.56364,  2.13467, 1.79201, 1.65708, // ACCGTT
    98.01512,  1.77408, 1.30411, 1.02284, // ACCTAA
    95.25879,  1.77408, 1.37864, 1.11177, // ACCTAC
    97.16902,  1.77408, 1.05253, 0.74164, // ACCTAG
    96.13042,  1.77408, 1.36413, 1.09426, // ACCTAT
    91.24900,  2.16210, 1.71292, 1.55628, // ACCTCA
    90.14287,  2.16210, 1.86109, 1.76252, // ACCTCC
    91.39747,  2.16210, 1.62527, 1.43836, // ACCTCG
    91.26808,  2.16210, 1.78964, 1.66200, // ACCTCT
    95.41651,  1.63830, 1.26908, 0.98862, // ACCTGA
    92.07513,  1.63830, 1.41720, 1.16665, // ACCTGC
    94.01394,  1.63830, 1.29676, 1.02113, // ACCTGG
    93.17519,  1.63830, 1.44582, 1.20217, // ACCTGT
    86.81065,  1.91311, 1.71890, 1.48956, // ACCTTA
    83.39007,  1.91311, 1.70994, 1.47793, // ACCTTC
    85.81538,  1.91311, 1.56463, 1.29360, // ACCTTG
    84.69539,  1.91311, 1.58306, 1.31652, // ACCTTT
    85.68487,  2.36795, 1.56701, 1.16930, // ACGAAA
    82.10622,  2.36795, 1.88115, 1.53799, // ACGAAC
    84.58063,  2.36795, 1.70811, 1.33074, // ACGAAG
    83.06389,  2.36795, 1.92702, 1.59459, // ACGAAT
    75.36940,  2.00332, 1.81079, 1.55061, // ACGACA
    74.19597,  2.00332, 1.65631, 1.35647, // ACGACC
    75.24330,  2.00332, 1.71221, 1.42572, // ACGACG
    75.11467,  2.00332, 1.66362, 1.36547, // ACGACT
    84.40649,  2.41291, 1.51583, 1.11615, // ACGAGA
    82.02665,  2.41291, 1.78219, 1.42291, // ACGAGC
    83.11420,  2.41291, 1.51980, 1.12053, // ACGAGG
    82.40086,  2.41291, 1.83001, 1.48055, // ACGAGT
    74.29910,  1.85855, 1.56104, 1.41982, // ACGATA
    72.41119,  1.85855, 1.80103, 1.75951, // ACGATC
    73.39194,  1.85855, 1.56900, 1.43069, // ACGATG
    73.68440,  1.85855, 1.93844, 1.96467, // ACGATT
    90.72405,  1.44291, 1.25549, 0.98457, // ACGCAA
    87.92699,  1.44291, 1.18231, 0.89975, // ACGCAC
    89.69155,  1.44291, 1.23807, 0.96416, // ACGCAG
    88.90817,  1.44291, 1.16388, 0.87880, // ACGCAT
    86.80542,  1.52741, 1.41921, 1.19162, // ACGCCA
    85.42382,  1.52741, 1.44306, 1.22178, // ACGCCC
    86.59406,  1.52741, 1.41830, 1.19047, // ACGCCG
    86.47021,  1.52741, 1.36153, 1.11971, // ACGCCT
    88.86568,  1.58181, 1.28360, 0.98233, // ACGCGA
    85.75870,  1.58181, 1.35971, 1.07099, // ACGCGC
    87.67576,  1.58181, 1.34631, 1.05519, // ACGCGG
    86.47437,  1.58181, 1.34700, 1.05600, // ACGCGT
    85.13681,  1.56040, 1.41851, 1.12330, // ACGCTA
    82.69009,  1.56040, 1.39000, 1.08960, // ACGCTC
    84.23896,  1.56040, 1.39386, 1.09415, // ACGCTG
    83.58706,  1.56040, 1.18494, 0.85761, // ACGCTT
    84.93959,  2.44172, 1.76816, 1.37549, // ACGGAA
    79.42779,  2.44172, 2.03723, 1.70112, // ACGGAC
    83.71984,  2.44172, 1.82726, 1.44503, // ACGGAG
    81.12399,  2.44172, 2.01232, 1.67002, // ACGGAT
    79.99905,  2.03186, 1.49247, 1.20792, // ACGGCA
    78.44174,  2.03186, 1.50469, 1.22280, // ACGGCC
    79.78446,  2.03186, 1.48313, 1.19660, // ACGGCG
    79.70291,  2.03186, 1.50224, 1.21981, // ACGGCT
    85.58088,  2.35270, 1.63675, 1.21654, // ACGGGA
    82.61027,  2.35270, 1.81525, 1.42087, // ACGGGC
    84.24855,  2.35270, 1.59567, 1.17102, // ACGGGG
    83.63375,  2.35270, 1.85320, 1.46566, // ACGGGT
    78.32032,  2.46096, 1.84312, 1.56160, // ACGGTA
    75.53449,  2.46096, 2.00903, 1.77714, // ACGGTC
    77.16704,  2.46096, 1.92296, 1.66416, // ACGGTG
    76.54799,  2.46096, 1.99035, 1.75241, // ACGGTT
    82.12717,  2.14468, 1.80972, 1.57415, // ACGTAA
    78.53428,  2.14468, 1.93097, 1.73496, // ACGTAC
    81.44443,  2.14468, 1.89084, 1.68117, // ACGTAG
    80.16145,  2.14468, 2.00056, 1.82960, // ACGTAT
    75.28892,  2.49565, 2.48078, 2.86185, // ACGTCA
    73.78885,  2.49565, 2.54924, 2.98112, // ACGTCC
    75.21510,  2.49565, 2.45874, 2.82380, // ACGTCG
    75.03445,  2.49565, 2.47828, 2.85752, // ACGTCT
    78.72390,  2.22005, 1.90469, 1.77665, // ACGTGA
    74.67260,  2.22005, 2.14974, 2.13031, // ACGTGC
    77.41127,  2.22005, 2.11114, 2.07319, // ACGTGG
    75.76270,  2.22005, 2.24488, 2.27328, // ACGTGT
    69.69304,  2.47313, 2.39457, 2.49536, // ACGTTA
    66.19745,  2.47313, 2.51962, 2.69335, // ACGTTC
    68.73888,  2.47313, 2.44079, 2.56795, // ACGTTG
    67.60651,  2.47313, 2.40054, 2.50469, // ACGTTT
    109.60510, 2.64098, 1.78499, 1.34274, // ACTAAA
    107.06679, 2.64098, 1.77711, 1.33385, // ACTAAC
    108.71678, 2.64098, 1.82866, 1.39231, // ACTAAG
    108.65006, 2.64098, 1.98981, 1.58036, // ACTAAT
    104.91342, 2.58253, 1.53278, 1.06551, // ACTACA
    104.37825, 2.58253, 1.69406, 1.23803, // ACTACC
    104.60198, 2.58253, 1.50915, 1.04096, // ACTACG
    105.78065, 2.58253, 1.81195, 1.36948, // ACTACT
    107.84126, 2.51606, 1.83695, 1.55379, // ACTAGA
    105.68891, 2.51606, 1.61234, 1.27770, // ACTAGC
    106.36396, 2.51606, 1.53484, 1.18670, // ACTAGG
    107.10914, 2.51606, 1.60849, 1.27313, // ACTAGT
    104.96065, 3.03892, 1.91641, 1.40773, // ACTATA
    105.35614, 3.03892, 2.02830, 1.53279, // ACTATC
    104.56493, 3.03892, 1.87534, 1.36271, // ACTATG
    106.93514, 3.03892, 2.25275, 1.79414, // ACTATT
    115.09190, 2.42793, 1.81878, 1.59109, // ACTCAA
    112.57760, 2.42793, 1.69623, 1.43303, // ACTCAC
    114.34963, 2.42793, 1.82823, 1.60352, // ACTCAG
    113.79365, 2.42793, 1.94818, 1.76388, // ACTCAT
    113.71476, 2.47041, 1.82491, 1.51937, // ACTCCA
    111.72680, 2.47041, 1.74480, 1.42044, // ACTCCC
    113.64929, 2.47041, 1.84678, 1.54677, // ACTCCG
    112.87568, 2.47041, 1.95376, 1.68309, // ACTCCT
    113.91984, 2.49891, 1.79737, 1.48675, // ACTCGA
    111.63731, 2.49891, 1.85362, 1.55710, // ACTCGC
    114.07047, 2.49891, 1.98026, 1.71936, // ACTCGG
    113.36890, 2.49891, 2.00748, 1.75494, // ACTCGT
    113.72638, 2.72204, 1.82568, 1.50849, // ACTCTA
    111.73290, 2.72204, 1.99910, 1.72846, // ACTCTC
    113.20368, 2.72204, 1.94200, 1.65494, // ACTCTG
    112.99610, 2.72204, 2.31939, 2.16006, // ACTCTT
    105.11124, 2.50593, 1.75335, 1.35976, // ACTGAA
    101.17513, 2.50593, 1.64828, 1.23939, // ACTGAC
    104.20205, 2.50593, 1.72515, 1.32709, // ACTGAG
    103.16431, 2.50593, 1.82084, 1.43903, // ACTGAT
    102.97577, 2.56584, 1.57276, 1.10065, // ACTGCA
    102.18670, 2.56584, 1.62044, 1.15109, // ACTGCC
    102.89065, 2.56584, 1.64559, 1.17798, // ACTGCG
    103.88761, 2.56584, 1.76473, 1.30821, // ACTGCT
    105.30268, 2.47346, 1.75997, 1.38612, // ACTGGA
    102.53315, 2.47346, 1.64528, 1.25286, // ACTGGC
    104.40768, 2.47346, 1.73224, 1.35349, // ACTGGG
    104.49876, 2.47346, 1.86940, 1.51738, // ACTGGT
    103.07858, 3.11250, 1.91169, 1.39461, // ACTGTA
    103.11917, 3.11250, 2.04214, 1.53976, // ACTGTC
    103.12730, 3.11250, 1.91520, 1.39845, // ACTGTG
    104.95890, 3.11250, 2.23889, 1.76757, // ACTGTT
    104.24170, 1.86007, 1.35553, 0.99397, // ACTTAA
    100.05889, 1.86007, 1.25649, 0.88704, // ACTTAC
    103.66617, 1.86007, 1.31032, 0.94465, // ACTTAG
    101.76082, 1.86007, 1.34247, 0.97963, // ACTTAT
    98.45005,  1.79286, 1.45211, 1.11851, // ACTTCA
    96.66331,  1.79286, 1.44106, 1.10576, // ACTTCC
    98.39871,  1.79286, 1.42524, 1.08761, // ACTTCG
    98.02968,  1.79286, 1.42580, 1.08825, // ACTTCT
    102.26170, 1.88803, 1.27431, 0.94949, // ACTTGA
    98.11485,  1.88803, 1.25882, 0.93224, // ACTTGC
    101.32567, 1.88803, 1.36817, 1.05630, // ACTTGG
    99.81328,  1.88803, 1.30757, 0.98691, // ACTTGT
    94.60815,  1.89728, 1.23750, 0.84797, // ACTTTA
    92.09168,  1.89728, 1.20209, 0.81183, // ACTTTC
    93.95156,  1.89728, 1.22881, 0.83906, // ACTTTG
    94.31763,  1.89728, 1.23664, 0.84708, // ACTTTT
    79.73762,  1.67748, 1.08810, 0.72343, // AGAAAA
    77.58039,  1.67748, 1.04629, 0.68214, // AGAAAC
    78.65132,  1.67748, 1.07330, 0.70873, // AGAAAG
    77.97715,  1.67748, 1.05928, 0.69488, // AGAAAT
    69.45981,  2.18144, 1.55604, 1.21560, // AGAACA
    68.24815,  2.18144, 1.55865, 1.21866, // AGAACC
    69.54785,  2.18144, 1.56447, 1.22549, // AGAACG
    69.47773,  2.18144, 1.42127, 1.06115, // AGAACT
    77.36004,  1.59541, 1.10042, 0.74694, // AGAAGA
    76.82254,  1.59541, 0.97460, 0.62257, // AGAAGC
    76.63338,  1.59541, 1.09712, 0.74359, // AGAAGG
    76.60311,  1.59541, 1.00034, 0.64740, // AGAAGT
    69.75056,  1.95147, 1.32986, 0.97283, // AGAATA
    68.00686,  1.95147, 1.23796, 0.87374, // AGAATC
    69.03248,  1.95147, 1.25434, 0.89114, // AGAATG
    69.03891,  1.95147, 1.13737, 0.76944, // AGAATT
    84.95638,  2.48599, 1.78971, 1.65509, // AGACAA
    83.40053,  2.48599, 1.60586, 1.40672, // AGACAC
    84.01266,  2.48599, 1.63755, 1.44857, // AGACAG
    84.30642,  2.48599, 1.56983, 1.35965, // AGACAT
    80.71250,  2.19720, 1.83649, 1.83058, // AGACCA
    79.65685,  2.19720, 1.62768, 1.52742, // AGACCC
    80.75777,  2.19720, 1.78318, 1.75145, // AGACCG
    80.84230,  2.19720, 1.72728, 1.66974, // AGACCT
    83.75337,  1.98191, 1.62836, 1.52196, // AGACGA
    82.32478,  1.98191, 1.50701, 1.35505, // AGACGC
    82.85512,  1.98191, 1.60290, 1.48640, // AGACGG
    82.61912,  1.98191, 1.57058, 1.44168, // AGACGT
    80.39252,  1.92945, 1.50382, 1.39992, // AGACTA
    79.03189,  1.92945, 1.41325, 1.27537, // AGACTC
    79.92163,  1.92945, 1.57927, 1.50658, // AGACTG
    79.45908,  1.92945, 1.19775, 0.99508, // AGACTT
    79.70502,  1.41776, 1.00120, 0.70704, // AGAGAA
    75.89352,  1.41776, 1.16624, 0.88889, // AGAGAC
    78.34065,  1.41776, 1.03765, 0.74601, // AGAGAG
    77.11158,  1.41776, 1.16682, 0.88955, // AGAGAT
    72.78265,  1.59099, 1.21805, 0.86020, // AGAGCA
    71.45409,  1.59099, 1.18467, 0.82508, // AGAGCC
    72.79337,  1.59099, 1.12391, 0.76242, // AGAGCG
    72.84450,  1.59099, 1.10945, 0.74776, // AGAGCT
    78.91211,  1.35821, 0.98338, 0.67166, // AGAGGA
    76.74545,  1.35821, 1.08303, 0.77629, // AGAGGC
    78.20516,  1.35821, 0.97898, 0.66715, // AGAGGG
    77.52585,  1.35821, 1.08809, 0.78174, // AGAGGT
    72.64988,  1.45706, 0.94952, 0.63402, // AGAGTA
    71.00046,  1.45706, 1.12795, 0.82087, // AGAGTC
    72.14156,  1.45706, 1.04330, 0.73022, // AGAGTG
    72.12275,  1.45706, 1.21518, 0.91792, // AGAGTT
    78.16891,  2.58682, 2.01072, 2.00564, // AGATAA
    75.89173,  2.58682, 1.89705, 1.83800, // AGATAC
    77.37135,  2.58682, 1.87561, 1.80693, // AGATAG
    77.43397,  2.58682, 2.05402, 2.07077, // AGATAT
    71.31328,  2.70241, 2.29287, 2.70016, // AGATCA
    70.60058,  2.70241, 2.46108, 3.00267, // AGATCC
    71.48606,  2.70241, 2.29132, 2.69741, // AGATCG
    72.29044,  2.70241, 2.62501, 3.30762, // AGATCT
    75.57488,  2.05805, 1.82321, 1.91469, // AGATGA
    73.09137,  2.05805, 1.87844, 2.00234, // AGATGC
    74.50635,  2.05805, 1.80686, 1.88899, // AGATGG
    74.12339,  2.05805, 2.01780, 2.22926, // AGATGT
    67.46160,  2.35300, 1.96410, 2.00682, // AGATTA
    64.66484,  2.35300, 1.82858, 1.80274, // AGATTC
    66.47004,  2.35300, 1.79933, 1.75965, // AGATTG
    66.16008,  2.35300, 1.73700, 1.66902, // AGATTT
    105.59051, 2.21768, 1.79959, 1.48473, // AGCAAA
    102.45988, 2.21768, 1.92923, 1.64802, // AGCAAC
    104.51222, 2.21768, 1.84686, 1.54362, // AGCAAG
    103.80151, 2.21768, 1.99074, 1.72747, // AGCAAT
    96.59164,  1.72978, 1.22791, 0.85677, // AGCACA
    95.09218,  1.72978, 1.27845, 0.91020, // AGCACC
    96.42328,  1.72978, 1.25632, 0.88667, // AGCACG
    96.68609,  1.72978, 1.45681, 1.10718, // AGCACT
    102.73184, 2.29394, 1.78239, 1.43898, // AGCAGA
    99.73497,  2.29394, 1.74636, 1.39557, // AGCAGC
    101.49318, 2.29394, 1.78162, 1.43805, // AGCAGG
    101.01826, 2.29394, 1.99085, 1.69867, // AGCAGT
    96.08068,  2.05386, 1.45939, 1.12231, // AGCATA
    94.57679,  2.05386, 1.62236, 1.31545, // AGCATC
    95.46815,  2.05386, 1.46005, 1.12307, // AGCATG
    96.14382,  2.05386, 1.75748, 1.48317, // AGCATT
    106.23421, 1.96521, 1.47199, 1.26140, // AGCCAA
    103.37831, 1.96521, 1.52091, 1.32481, // AGCCAC
    105.35533, 1.96521, 1.49291, 1.28839, // AGCCAG
    104.70667, 1.96521, 1.69935, 1.56467, // AGCCAT
    102.61347, 1.76337, 1.35848, 1.13096, // AGCCCA
    100.69997, 1.76337, 1.26908, 1.02118, // AGCCCC
    102.45967, 1.76337, 1.33123, 1.09709, // AGCCCG
    101.79677, 1.76337, 1.45288, 1.25086, // AGCCCT
    104.92154, 1.93464, 1.56010, 1.39917, // AGCCGA
    102.30698, 1.93464, 1.55199, 1.38828, // AGCCGC
    103.88504, 1.93464, 1.55128, 1.38734, // AGCCGG
    103.44232, 1.93464, 1.70782, 1.60254, // AGCCGT
    101.54913, 1.85738, 1.38877, 1.05146, // AGCCTA
    99.30654,  1.85738, 1.52241, 1.20682, // AGCCTC
    100.90244, 1.85738, 1.46404, 1.13809, // AGCCTG
    100.44952, 1.85738, 1.67194, 1.38892, // AGCCTT
    101.13829, 2.24827, 1.89422, 1.65705, // AGCGAA
    96.90534,  2.24827, 1.75699, 1.48028, // AGCGAC
    100.51203, 2.24827, 1.70520, 1.41531, // AGCGAG
    98.86572,  2.24827, 1.95617, 1.73899, // AGCGAT
    95.93565,  1.86985, 1.45399, 1.10557, // AGCGCA
    94.43324,  1.86985, 1.44846, 1.09927, // AGCGCC
    95.92198,  1.86985, 1.45659, 1.10854, // AGCGCG
    96.01057,  1.86985, 1.61437, 1.29345, // AGCGCT
    100.98811, 2.16392, 1.91151, 1.70788, // AGCGGA
    98.00531,  2.16392, 1.72836, 1.46840, // AGCGGC
    100.15058, 2.16392, 1.76206, 1.51155, // AGCGGG
    99.63023,  2.16392, 1.90136, 1.69429, // AGCGGT
    95.73931,  2.46720, 1.74294, 1.44600, // AGCGTA
    94.22061,  2.46720, 1.81332, 1.53445, // AGCGTC
    95.20073,  2.46720, 1.81042, 1.53078, // AGCGTG
    96.12240,  2.46720, 2.09402, 1.90421, // AGCGTT
    99.63621,  1.80194, 1.67024, 1.36156, // AGCTAA
    96.45278,  1.80194, 1.47981, 1.13548, // AGCTAC
    98.46221,  1.80194, 1.58090, 1.25379, // AGCTAG
    97.76014,  1.80194, 1.53013, 1.19388, // AGCTAT
    91.71822,  2.01289, 1.67351, 1.42475, // AGCTCA
    89.79097,  2.01289, 1.74461, 1.51651, // AGCTCC
    91.84239,  2.01289, 1.60820, 1.34218, // AGCTCG
    91.37889,  2.01289, 1.71803, 1.48199, // AGCTCT
    96.88074,  1.67840, 1.51234, 1.20029, // AGCTGA
    93.46389,  1.67840, 1.37694, 1.04276, // AGCTGC
    95.56791,  1.67840, 1.47689, 1.15834, // AGCTGG
    94.55177,  1.67840, 1.45287, 1.13019, // AGCTGT
    87.36690,  1.51841, 1.41299, 1.10879, // AGCTTA
    84.42880,  1.51841, 1.24603, 0.91820, // AGCTTC
    86.85296,  1.51841, 1.29414, 0.97188, // AGCTTG
    85.66451,  1.51841, 1.10217, 0.76386, // AGCTTT
    72.74415,  1.83378, 1.22764, 0.79505, // AGGAAA
    69.89161,  1.83378, 1.35461, 0.92154, // AGGAAC
    71.22479,  1.83378, 1.29147, 0.85786, // AGGAAG
    70.58931,  1.83378, 1.32620, 0.89270, // AGGAAT
    60.91353,  2.32589, 1.84078, 1.43995, // AGGACA
    60.08858,  2.32589, 1.92056, 1.53457, // AGGACC
    60.98795,  2.32589, 1.93153, 1.54774, // AGGACG
    61.03662,  2.32589, 1.71638, 1.29648, // AGGACT
    70.55430,  1.86110, 1.29856, 0.93297, // AGGAGA
    69.34039,  1.86110, 1.23683, 0.86724, // AGGAGC
    69.38303,  1.86110, 1.36777, 1.00855, // AGGAGG
    69.31639,  1.86110, 1.21569, 0.84511, // AGGAGT
    61.69028,  1.92277, 1.73037, 1.42446, // AGGATA
    60.33079,  1.92277, 1.55428, 1.21266, // AGGATC
    61.01395,  1.92277, 1.60928, 1.27759, // AGGATG
    61.42190,  1.92277, 1.28752, 0.91427, // AGGATT
    83.65624,  1.89614, 1.60776, 1.39868, // AGGCAA
    81.50612,  1.89614, 1.32000, 1.04051, // AGGCAC
    82.56152,  1.89614, 1.56674, 1.34549, // AGGCAG
    82.51457,  1.89614, 1.36378, 1.09271, // AGGCAT
    79.25195,  1.71082, 1.60340, 1.43590, // AGGCCA
    78.17076,  1.71082, 1.55874, 1.37633, // AGGCCC
    79.11551,  1.71082, 1.59689, 1.42717, // AGGCCG
    79.05207,  1.71082, 1.57318, 1.39549, // AGGCCT
    82.02143,  1.57550, 1.41475, 1.17987, // AGGCGA
    79.62117,  1.57550, 1.31215, 1.05387, // AGGCGC
    80.90574,  1.57550, 1.38856, 1.14727, // AGGCGG
    80.29601,  1.57550, 1.33347, 1.07967, // AGGCGT
    78.40385,  1.47431, 1.42184, 1.16218, // AGGCTA
    76.46661,  1.47431, 1.29132, 1.00588, // AGGCTC
    77.72343,  1.47431, 1.36160, 1.08912, // AGGCTG
    77.21187,  1.47431, 1.09321, 0.78353, // AGGCTT
    75.08225,  1.70210, 1.10878, 0.72685, // AGGGAA
    70.38135,  1.70210, 1.37412, 1.00281, // AGGGAC
    74.05506,  1.70210, 1.08724, 0.70578, // AGGGAG
    71.98229,  1.70210, 1.29842, 0.92109, // AGGGAT
    68.22513,  1.76808, 1.44825, 1.06143, // AGGGCA
    66.74489,  1.76808, 1.46362, 1.07838, // AGGGCC
    68.28452,  1.76808, 1.41889, 1.02932, // AGGGCG
    68.07767,  1.76808, 1.37433, 0.98121, // AGGGCT
    75.37677,  1.51256, 1.02860, 0.69371, // AGGGGA
    73.13694,  1.51256, 1.10436, 0.77175, // AGGGGC
    74.57971,  1.51256, 0.95862, 0.62413, // AGGGGG
    73.97048,  1.51256, 1.10899, 0.77661, // AGGGGT
    68.05667,  1.63484, 1.13416, 0.81137, // AGGGTA
    65.93227,  1.63484, 1.24686, 0.93526, // AGGGTC
    67.13477,  1.63484, 1.12686, 0.80355, // AGGGTG
    67.61058,  1.63484, 1.35120, 1.05508, // AGGGTT
    73.90667,  2.38667, 1.94833, 1.80553, // AGGTAA
    70.68024,  2.38667, 1.70329, 1.47585, // AGGTAC
    72.57155,  2.38667, 1.90831, 1.75019, // AGGTAG
    72.32862,  2.38667, 1.90007, 1.73887, // AGGTAT
    66.21875,  2.69199, 2.00760, 2.12192, // AGGTCA
    65.06782,  2.69199, 2.16087, 2.36949, // AGGTCC
    66.18776,  2.69199, 1.91538, 1.97741, // AGGTCG
    66.28898,  2.69199, 2.21170, 2.45359, // AGGTCT
    69.90868,  2.13251, 1.71347, 1.59926, // AGGTGA
    66.77344,  2.13251, 1.77829, 1.69087, // AGGTGC
    68.68470,  2.13251, 1.62552, 1.47772, // AGGTGG
    67.84169,  2.13251, 1.79891, 1.72035, // AGGTGT
    62.14313,  2.62114, 2.07388, 1.97327, // AGGTTA
    59.36188,  2.62114, 2.30480, 2.31187, // AGGTTC
    61.55308,  2.62114, 2.01714, 1.89285, // AGGTTG
    60.58549,  2.62114, 2.00778, 1.87969, // AGGTTT
    107.31818, 2.96792, 2.47623, 2.00239, // AGTAAA
    104.71904, 2.96792, 2.48563, 2.01380, // AGTAAC
    106.24774, 2.96792, 2.45718, 1.97934, // AGTAAG
    106.89103, 2.96792, 2.62386, 2.18411, // AGTAAT
    100.41577, 2.66382, 2.08182, 1.59872, // AGTACA
    100.23167, 2.66382, 2.22721, 1.76909, // AGTACC
    100.47641, 2.66382, 2.06477, 1.57912, // AGTACG
    102.00699, 2.66382, 2.15706, 1.68617, // AGTACT
    105.19299, 2.77966, 2.33354, 1.93877, // AGTAGA
    102.80583, 2.77966, 2.30493, 1.90322, // AGTAGC
    104.33295, 2.77966, 2.17747, 1.74755, // AGTAGG
    105.04105, 2.77966, 2.62265, 2.31000, // AGTAGT
    101.99600, 3.77006, 2.51775, 2.00755, // AGTATA
    103.02473, 3.77006, 2.62019, 2.13132, // AGTATC
    101.71340, 3.77006, 2.63292, 2.14686, // AGTATG
    105.25232, 3.77006, 2.94594, 2.54087, // AGTATT
    115.74456, 2.78871, 2.71665, 2.89390, // AGTCAA
    113.45636, 2.78871, 2.60991, 2.72502, // AGTCAC
    115.02895, 2.78871, 2.59796, 2.70633, // AGTCAG
    114.52908, 2.78871, 2.71690, 2.89428, // AGTCAT
    114.16661, 2.92671, 2.50562, 2.55218, // AGTCCA
    112.41097, 2.92671, 2.44614, 2.46185, // AGTCCC
    114.07268, 2.92671, 2.68237, 2.82694, // AGTCCG
    113.52250, 2.92671, 2.51036, 2.55943, // AGTCCT
    115.50966, 2.90558, 2.77885, 2.95622, // AGTCGA
    112.77381, 2.90558, 2.53831, 2.58082, // AGTCGC
    114.52685, 2.90558, 2.71568, 2.85600, // AGTCGG
    114.77168, 2.90558, 2.74253, 2.89845, // AGTCGT
    113.82369, 3.02381, 3.10125, 3.31683, // AGTCTA
    113.10517, 3.02381, 2.68468, 2.67151, // AGTCTC
    114.17053, 3.02381, 2.58891, 2.52984, // AGTCTG
    114.23695, 3.02381, 2.94227, 3.06507, // AGTCTT
    104.60878, 2.86549, 2.47043, 2.23788, // AGTGAA
    101.14965, 2.86549, 2.55404, 2.35245, // AGTGAC
    103.62359, 2.86549, 2.39201, 2.13218, // AGTGAG
    103.68409, 2.86549, 2.70593, 2.56540, // AGTGAT
    102.74835, 2.92533, 2.24864, 1.91265, // AGTGCA
    102.19906, 2.92533, 2.26848, 1.93801, // AGTGCC
    102.89103, 2.92533, 2.37726, 2.07907, // AGTGCG
    104.54464, 2.92533, 2.53149, 2.28464, // AGTGCT
    105.47302, 2.85435, 2.45545, 2.24024, // AGTGGA
    103.23315, 2.85435, 2.44236, 2.22234, // AGTGGC
    104.68215, 2.85435, 2.31369, 2.04906, // AGTGGG
    105.36966, 2.85435, 2.67063, 2.54107, // AGTGGT
    105.40292, 3.38667, 2.78922, 2.54947, // AGTGTA
    105.82330, 3.38667, 2.88082, 2.67608, // AGTGTC
    105.42987, 3.38667, 2.73224, 2.47174, // AGTGTG
    108.30825, 3.38667, 3.12153, 3.01841, // AGTGTT
    104.15935, 2.40487, 2.07483, 2.08587, // AGTTAA
    100.28468, 2.40487, 1.81180, 1.70207, // AGTTAC
    103.49304, 2.40487, 1.86412, 1.77632, // AGTTAG
    102.11459, 2.40487, 1.83710, 1.73785, // AGTTAT
    97.29695,  2.21733, 2.19881, 2.08101, // AGTTCA
    95.36013,  2.21733, 2.26075, 2.16956, // AGTTCC
    97.34270,  2.21733, 2.18418, 2.06027, // AGTTCG
    97.15650,  2.21733, 2.11452, 1.96250, // AGTTCT
    101.92706, 2.36448, 1.90687, 1.75635, // AGTTGA
    97.83344,  2.36448, 1.86502, 1.69886, // AGTTGC
    100.90786, 2.36448, 1.90940, 1.75985, // AGTTGG
    99.57770,  2.36448, 1.88914, 1.73192, // AGTTGT
    93.64647,  1.91331, 1.91003, 1.76480, // AGTTTA
    91.49898,  1.91331, 1.73059, 1.52205, // AGTTTC
    93.03756,  1.91331, 1.88580, 1.73132, // AGTTTG
    93.01751,  1.91331, 1.53996, 1.27762, // AGTTTT
    93.79473,  2.42434, 1.57687, 1.22726, // ATAAAA
    91.15586,  2.42434, 1.89717, 1.61958, // ATAAAC
    92.95292,  2.42434, 1.62758, 1.28693, // ATAAAG
    91.89059,  2.42434, 1.88984, 1.61020, // ATAAAT
    84.68293,  2.30963, 1.69423, 1.46061, // ATAACA
    83.48879,  2.30963, 1.61840, 1.36365, // ATAACC
    84.38443,  2.30963, 1.64764, 1.40078, // ATAACG
    84.23409,  2.30963, 1.57393, 1.30784, // ATAACT
    92.29336,  2.47862, 1.68310, 1.33125, // ATAAGA
    90.33165,  2.47862, 1.83482, 1.51526, // ATAAGC
    91.08240,  2.47862, 1.67035, 1.31616, // ATAAGG
    90.97774,  2.47862, 1.90871, 1.60770, // ATAAGT
    84.02277,  2.47960, 1.78173, 1.57934, // ATAATA
    82.55840,  2.47960, 2.15061, 2.09438, // ATAATC
    83.59893,  2.47960, 1.93051, 1.78124, // ATAATG
    83.25645,  2.47960, 2.17920, 2.13629, // ATAATT
    95.13486,  1.32011, 1.26931, 1.02022, // ATACAA
    93.12879,  1.32011, 1.17206, 0.90524, // ATACAC
    94.32318,  1.32011, 1.17395, 0.90743, // ATACAG
    94.09217,  1.32011, 1.13887, 0.86706, // ATACAT
    92.38954,  1.41774, 1.40558, 1.12366, // ATACCA
    91.18998,  1.41774, 1.37829, 1.09109, // ATACCC
    92.20567,  1.41774, 1.38253, 1.09614, // ATACCG
    91.98577,  1.41774, 1.30744, 1.00806, // ATACCT
    94.14073,  1.34132, 1.21208, 0.90475, // ATACGA
    92.09594,  1.34132, 1.21240, 0.90510, // ATACGC
    93.18780,  1.34132, 1.23697, 0.93276, // ATACGG
    92.64420,  1.34132, 1.20091, 0.89227, // ATACGT
    91.12710,  1.52174, 1.41339, 1.17198, // ATACTA
    89.44629,  1.52174, 1.26653, 0.99414, // ATACTC
    90.62898,  1.52174, 1.35891, 1.10486, // ATACTG
    90.24396,  1.52174, 1.37557, 1.12525, // ATACTT
    91.42696,  2.32328, 1.60769, 1.32628, // ATAGAA
    86.69851,  2.32328, 1.83709, 1.62004, // ATAGAC
    90.37252,  2.32328, 1.64517, 1.37293, // ATAGAG
    88.32654,  2.32328, 2.05073, 1.91070, // ATAGAT
    86.51265,  2.28919, 1.56045, 1.37884, // ATAGCA
    85.46653,  2.28919, 1.78523, 1.68725, // ATAGCC
    86.45716,  2.28919, 1.60563, 1.43915, // ATAGCG
    85.95473,  2.28919, 1.71304, 1.58595, // ATAGCT
    91.11246,  2.34031, 1.55603, 1.25638, // ATAGGA
    88.36086,  2.34031, 1.79962, 1.56266, // ATAGGC
    90.05070,  2.34031, 1.60771, 1.31949, // ATAGGG
    89.44830,  2.34031, 1.86582, 1.64968, // ATAGGT
    84.71810,  2.72334, 1.96004, 1.87494, // ATAGTA
    82.42144,  2.72334, 2.03688, 1.98627, // ATAGTC
    83.76555,  2.72334, 1.94091, 1.84756, // ATAGTG
    83.64063,  2.72334, 2.28158, 2.35475, // ATAGTT
    90.55660,  1.88263, 1.80275, 1.66447, // ATATAA
    87.66190,  1.88263, 1.83923, 1.71525, // ATATAC
    89.81416,  1.88263, 1.71064, 1.53855, // ATATAG
    88.87365,  1.88263, 1.86705, 1.75431, // ATATAT
    85.68595,  2.46291, 2.35953, 2.49103, // ATATCA
    84.45935,  2.46291, 2.38703, 2.53470, // ATATCC
    85.37966,  2.46291, 2.24608, 2.31355, // ATATCG
    85.28440,  2.46291, 2.31993, 2.42857, // ATATCT
    88.40309,  2.11919, 1.78046, 1.56483, // ATATGA
    85.29203,  2.11919, 2.06531, 1.95500, // ATATGC
    87.34669,  2.11919, 1.90620, 1.73349, // ATATGG
    86.32354,  2.11919, 2.04659, 1.92847, // ATATGT
    80.72200,  2.69506, 2.52054, 2.58799, // ATATTA
    77.56828,  2.69506, 2.48029, 2.52626, // ATATTC
    79.80331,  2.69506, 2.41837, 2.43225, // ATATTG
    79.07776,  2.69506, 2.33927, 2.31390, // ATATTT
    104.69457, 2.08052, 1.65247, 1.41304, // ATCAAA
    102.35254, 2.08052, 1.83989, 1.66013, // ATCAAC
    103.83233, 2.08052, 1.68580, 1.45600, // ATCAAG
    103.25146, 2.08052, 1.95166, 1.81367, // ATCAAT
    98.08033,  2.10126, 1.67386, 1.39585, // ATCACA
    96.85752,  2.10126, 1.67189, 1.39338, // ATCACC
    97.77408,  2.10126, 1.61007, 1.31682, // ATCACG
    97.54661,  2.10126, 1.74883, 1.49067, // ATCACT
    103.14687, 2.40061, 1.84262, 1.64081, // ATCAGA
    101.17145, 2.40061, 1.94509, 1.77956, // ATCAGC
    102.15399, 2.40061, 1.87611, 1.68574, // ATCAGG
    101.83061, 2.40061, 2.05478, 1.93220, // ATCAGT
    96.41120,  2.47361, 1.82769, 1.59362, // ATCATA
    94.93353,  2.47361, 1.97392, 1.78865, // ATCATC
    95.74034,  2.47361, 1.85877, 1.63444, // ATCATG
    95.84220,  2.47361, 2.16147, 2.04953, // ATCATT
    103.89771, 1.63253, 1.39898, 1.21060, // ATCCAA
    101.14796, 1.63253, 1.41598, 1.23273, // ATCCAC
    103.06665, 1.63253, 1.27224, 1.04987, // ATCCAG
    102.18616, 1.63253, 1.52237, 1.37424, // ATCCAT
    100.29815, 1.60868, 1.21153, 0.98116, // ATCCCA
    99.31157,  1.60868, 1.36765, 1.17679, // ATCCCC
    100.56298, 1.60868, 1.29170, 1.08014, // ATCCCG
    100.02892, 1.60868, 1.37158, 1.18186, // ATCCCT
    102.71989, 1.82797, 1.45262, 1.21134, // ATCCGA
    100.44501, 1.82797, 1.50033, 1.27151, // ATCCGC
    101.86598, 1.82797, 1.43010, 1.18329, // ATCCGG
    101.15044, 1.82797, 1.57180, 1.36343, // ATCCGT
    99.78581,  1.91501, 1.42880, 1.21337, // ATCCTA
    97.38258,  1.91501, 1.73183, 1.61918, // ATCCTC
    98.70789,  1.91501, 1.57036, 1.39809, // ATCCTG
    98.40446,  1.91501, 1.80432, 1.72190, // ATCCTT
    101.68077, 2.59137, 1.86527, 1.68529, // ATCGAA
    97.17925,  2.59137, 2.09177, 2.00140, // ATCGAC
    100.62107, 2.59137, 1.94115, 1.78917, // ATCGAG
    98.73655,  2.59137, 2.21640, 2.18291, // ATCGAT
    97.05357,  2.48599, 1.76431, 1.56306, // ATCGCA
    95.50119,  2.48599, 1.81133, 1.62596, // ATCGCC
    96.57899,  2.48599, 1.80250, 1.61408, // ATCGCG
    96.38580,  2.48599, 1.84527, 1.67187, // ATCGCT
    101.51893, 2.51279, 1.87047, 1.66318, // ATCGGA
    98.27156,  2.51279, 2.00178, 1.84135, // ATCGGC
    100.40557, 2.51279, 1.81423, 1.58873, // ATCGGG
    99.59241,  2.51279, 2.05045, 1.90890, // ATCGGT
    94.49653,  5.74004, 1.91023, 1.75033, // ATCGTA
    91.78190,  5.74004, 1.79642, 1.59626, // ATCGTC
    91.98362,  5.74004, 1.77468, 1.56738, // ATCGTG
    93.43773,  5.74004, 1.92510, 1.77080, // ATCGTT
    99.33426,  1.86097, 1.66516, 1.33296, // ATCTAA
    96.19987,  1.86097, 1.75171, 1.43822, // ATCTAC
    97.96393,  1.86097, 1.39076, 1.01745, // ATCTAG
    97.50568,  1.86097, 1.82747, 1.53253, // ATCTAT
    94.03527,  2.32915, 2.10394, 2.08340, // ATCTCA
    92.34256,  2.32915, 2.15828, 2.16463, // ATCTCC
    93.57149,  2.32915, 2.17261, 2.18622, // ATCTCG
    93.39215,  2.32915, 2.20544, 2.23597, // ATCTCT
    97.20684,  2.03287, 1.75945, 1.53289, // ATCTGA
    93.97878,  2.03287, 1.87688, 1.68888, // ATCTGC
    95.94911,  2.03287, 1.73864, 1.50577, // ATCTGG
    94.73559,  2.03287, 1.77508, 1.55335, // ATCTGT
    89.23135,  2.37489, 2.19609, 2.14834, // ATCTTA
    85.81331,  2.37489, 2.19508, 2.14685, // ATCTTC
    88.27375,  2.37489, 2.06379, 1.95715, // ATCTTG
    87.03908,  2.37489, 2.01551, 1.88888, // ATCTTT
    87.19755,  3.07534, 1.94009, 1.52771, // ATGAAA
    84.06041,  3.07534, 2.23581, 1.88999, // ATGAAC
    85.80763,  3.07534, 1.94070, 1.52842, // ATGAAG
    84.93302,  3.07534, 2.29793, 1.96929, // ATGAAT
    76.96090,  2.53545, 1.92617, 1.69681, // ATGACA
    75.72704,  2.53545, 1.79974, 1.53251, // ATGACC
    76.73404,  2.53545, 1.83494, 1.57769, // ATGACG
    76.55043,  2.53545, 1.72531, 1.43843, // ATGACT
    86.42476,  3.17429, 1.93846, 1.61228, // ATGAGA
    84.44112,  3.17429, 2.08360, 1.79671, // ATGAGC
    85.33302,  3.17429, 1.91537, 1.58356, // ATGAGG
    84.97895,  3.17429, 2.10524, 1.82477, // ATGAGT
    76.29342,  2.69379, 1.86362, 1.63677, // ATGATA
    75.54548,  2.69379, 2.36282, 2.33668, // ATGATC
    75.91030,  2.69379, 2.07416, 1.92183, // ATGATG
    75.93039,  2.69379, 2.34114, 2.30459, // ATGATT
    93.31503,  1.48208, 1.34877, 1.09880, // ATGCAA
    90.83253,  1.48208, 1.25738, 0.98904, // ATGCAC
    92.34616,  1.48208, 1.34120, 1.08956, // ATGCAG
    91.74390,  1.48208, 1.16966, 0.88737, // ATGCAT
    90.04483,  1.66578, 1.55970, 1.30892, // ATGCCA
    88.88185,  1.66578, 1.52228, 1.26210, // ATGCCC
    89.89598,  1.66578, 1.52564, 1.26628, // ATGCCG
    89.61947,  1.66578, 1.45460, 1.17887, // ATGCCT
    92.06637,  1.67467, 1.34368, 0.99526, // ATGCGA
    89.53870,  1.67467, 1.42637, 1.08853, // ATGCGC
    90.86742,  1.67467, 1.40515, 1.06433, // ATGCGG
    90.16696,  1.67467, 1.43259, 1.09566, // ATGCGT
    88.72254,  1.83904, 1.56710, 1.36488, // ATGCTA
    86.64499,  1.83904, 1.51788, 1.30108, // ATGCTC
    87.88986,  1.83904, 1.48527, 1.25938, // ATGCTG
    87.77665,  1.83904, 1.62826, 1.44555, // ATGCTT
    89.69845,  3.22959, 2.06265, 1.86345, // ATGGAA
    84.81982,  3.22959, 2.32679, 2.23263, // ATGGAC
    88.46465,  3.22959, 2.13094, 1.95675, // ATGGAG
    86.60223,  3.22959, 2.41193, 2.35627, // ATGGAT
    85.04505,  2.83028, 1.83263, 1.59952, // ATGGCA
    83.76943,  2.83028, 1.90176, 1.69087, // ATGGCC
    84.98438,  2.83028, 1.89036, 1.67570, // ATGGCG
    84.45677,  2.83028, 1.97253, 1.78613, // ATGGCT
    90.16198,  2.98826, 1.92663, 1.61767, // ATGGGA
    87.62250,  2.98826, 2.06233, 1.79156, // ATGGGC
    88.72899,  2.98826, 1.85858, 1.53273, // ATGGGG
    88.33417,  2.98826, 2.09716, 1.83714, // ATGGGT
    84.15925,  3.42265, 2.22849, 1.94542, // ATGGTA
    81.66863,  3.42265, 2.38819, 2.15824, // ATGGTC
    82.74259,  3.42265, 2.28373, 2.01820, // ATGGTG
    83.09398,  3.42265, 2.54805, 2.37852, // ATGGTT
    87.56436,  2.22980, 1.90357, 1.62371, // ATGTAA
    84.03006,  2.22980, 2.03149, 1.79010, // ATGTAC
    86.47526,  2.22980, 2.03863, 1.79954, // ATGTAG
    85.48093,  2.22980, 2.03236, 1.79125, // ATGTAT
    82.69946,  2.83133, 2.63435, 2.66436, // ATGTCA
    81.50008,  2.83133, 2.70387, 2.77052, // ATGTCC
    82.42726,  2.83133, 2.50788, 2.47481, // ATGTCG
    82.13821,  2.83133, 2.48871, 2.44649, // ATGTCT
    85.00967,  2.65515, 2.18261, 1.99618, // ATGTGA
    81.34280,  2.65515, 2.38566, 2.28111, // ATGTGC
    83.40364,  2.65515, 2.26082, 2.10442, // ATGTGG
    82.34214,  2.65515, 2.28172, 2.13367, // ATGTGT
    78.09946,  3.04048, 2.69835, 2.77889, // ATGTTA
    74.37517,  3.04048, 2.72964, 2.82737, // ATGTTC
    77.09206,  3.04048, 2.69614, 2.77547, // ATGTTG
    75.85129,  3.04048, 2.55241, 2.55652, // ATGTTT
    107.16362, 2.16371, 1.49912, 1.18194, // ATTAAA
    104.95414, 2.16371, 1.69367, 1.41934, // ATTAAC
    106.41662, 2.16371, 1.48251, 1.16235, // ATTAAG
    106.35831, 2.16371, 1.73534, 1.47203, // ATTAAT
    101.74633, 2.01953, 1.29689, 0.91929, // ATTACA
    101.08249, 2.01953, 1.36380, 0.99134, // ATTACC
    101.72532, 2.01953, 1.30086, 0.92351, // ATTACG
    102.42604, 2.01953, 1.41064, 1.04285, // ATTACT
    105.35916, 2.04194, 1.45085, 1.13485, // ATTAGA
    103.46301, 2.04194, 1.63101, 1.35266, // ATTAGC
    104.65520, 2.04194, 1.53412, 1.23394, // ATTAGG
    104.99768, 2.04194, 1.66538, 1.39565, // ATTAGT
    101.57655, 2.50557, 1.63863, 1.31333, // ATTATA
    101.44099, 2.50557, 1.81357, 1.52916, // ATTATC
    100.92647, 2.50557, 1.68931, 1.37473, // ATTATG
    103.23744, 2.50557, 1.87435, 1.60667, // ATTATT
    110.52115, 2.19857, 1.36739, 1.04675, // ATTCAA
    108.65768, 2.19857, 1.36330, 1.04206, // ATTCAC
    109.93881, 2.19857, 1.36114, 1.03959, // ATTCAG
    109.55669, 2.19857, 1.47687, 1.17494, // ATTCAT
    108.67945, 2.17160, 1.32279, 0.97959, // ATTCCA
    107.19467, 2.17160, 1.31956, 0.97601, // ATTCCC
    108.53621, 2.17160, 1.40284, 1.06985, // ATTCCG
    108.09868, 2.17160, 1.42231, 1.09219, // ATTCCT
    109.85938, 2.22607, 1.48296, 1.18243, // ATTCGA
    107.72661, 2.22607, 1.38529, 1.06757, // ATTCGC
    109.51163, 2.22607, 1.41513, 1.10224, // ATTCGG
    109.12734, 2.22607, 1.53435, 1.24442, // ATTCGT
    108.88093, 2.26458, 1.40010, 1.08926, // ATTCTA
    107.89257, 2.26458, 1.49782, 1.20527, // ATTCTC
    108.54936, 2.26458, 1.48496, 1.18978, // ATTCTG
    108.75394, 2.26458, 1.62198, 1.35820, // ATTCTT
    103.68880, 2.11369, 1.50205, 1.29065, // ATTGAA
    99.90611,  2.11369, 1.71114, 1.56931, // ATTGAC
    102.70965, 2.11369, 1.54398, 1.34507, // ATTGAG
    101.54006, 2.11369, 1.85854, 1.77640, // ATTGAT
    100.77890, 2.06630, 1.27285, 0.93220, // ATTGCA
    100.02740, 2.06630, 1.33577, 1.00216, // ATTGCC
    100.77610, 2.06630, 1.32718, 0.99251, // ATTGCG
    101.46161, 2.06630, 1.37337, 1.04477, // ATTGCT
    103.64245, 1.98654, 1.42783, 1.19349, // ATTGGA
    101.27455, 1.98654, 1.51760, 1.30780, // ATTGGC
    102.88093, 1.98654, 1.41716, 1.18014, // ATTGGG
    103.09235, 1.98654, 1.55136, 1.35167, // ATTGGT
    101.11055, 2.51049, 1.54362, 1.21925, // ATTGTA
    100.88935, 2.51049, 1.72791, 1.44398, // ATTGTC
    100.84393, 2.51049, 1.65559, 1.35428, // ATTGTG
    102.78337, 2.51049, 1.79033, 1.52293, // ATTGTT
    101.71343, 1.70840, 1.08220, 0.75389, // ATTTAA
    98.38569,  1.70840, 1.02777, 0.69774, // ATTTAC
    100.81662, 1.70840, 1.05562, 0.72630, // ATTTAG
    99.67506,  1.70840, 1.05025, 0.72076, // ATTTAT
    96.27519,  1.56977, 1.19292, 0.90778, // ATTTCA
    94.48898,  1.56977, 1.15448, 0.86426, // ATTTCC
    96.04806,  1.56977, 1.16328, 0.87417, // ATTTCG
    95.89435,  1.56977, 1.17229, 0.88434, // ATTTCT
    100.16951, 1.62351, 1.00007, 0.67717, // ATTTGA
    96.52232,  1.62351, 1.03624, 0.71424, // ATTTGC
    98.89950,  1.62351, 1.01909, 0.69658, // ATTTGG
    97.46554,  1.62351, 1.04119, 0.71937, // ATTTGT
    92.64328,  1.51391, 1.13431, 0.81682, // ATTTTA
    90.43820,  1.51391, 1.10265, 0.78287, // ATTTTC
    92.17087,  1.51391, 1.03481, 0.71174, // ATTTTG
    92.21591,  1.51391, 1.03607, 0.71304, // ATTTTT
    86.81562,  1.51785, 1.05997, 0.72794, // CAAAAA
    83.85032,  1.51785, 1.28891, 0.97609, // CAAAAC
    86.04444,  1.51785, 1.11668, 0.78713, // CAAAAG
    84.48643,  1.51785, 1.31591, 1.00692, // CAAAAT
    76.43798,  1.70501, 1.51867, 1.18640, // CAAACA
    75.27268,  1.70501, 1.45700, 1.11487, // CAAACC
    76.67243,  1.70501, 1.49200, 1.15528, // CAAACG
    76.34599,  1.70501, 1.34246, 0.98603, // CAAACT
    84.69193,  1.53757, 1.07529, 0.74838, // CAAAGA
    82.74224,  1.53757, 1.18789, 0.86895, // CAAAGC
    83.80544,  1.53757, 1.10766, 0.78242, // CAAAGG
    82.82370,  1.53757, 1.27292, 0.96390, // CAAAGT
    76.19247,  1.54253, 1.32188, 1.05318, // CAAATA
    73.94960,  1.54253, 1.20799, 0.92005, // CAAATC
    75.36262,  1.54253, 1.24307, 0.96041, // CAAATG
    75.25496,  1.54253, 1.48202, 1.25025, // CAAATT
    89.08822,  1.96203, 1.41203, 1.25157, // CAACAA
    87.22138,  1.96203, 1.21474, 0.99866, // CAACAC
    88.18537,  1.96203, 1.30512, 1.11216, // CAACAG
    88.05052,  1.96203, 1.20035, 0.98096, // CAACAT
    85.41713,  1.83613, 1.49102, 1.42202, // CAACCA
    84.15069,  1.83613, 1.36144, 1.24073, // CAACCC
    85.35102,  1.83613, 1.41150, 1.30978, // CAACCG
    85.06306,  1.83613, 1.34393, 1.21687, // CAACCT
    87.97785,  1.55453, 1.17444, 0.99030, // CAACGA
    86.21400,  1.55453, 1.13424, 0.93990, // CAACGC
    87.10736,  1.55453, 1.21104, 1.03696, // CAACGG
    86.72755,  1.55453, 1.14390, 0.95193, // CAACGT
    84.42248,  1.58806, 1.21695, 1.04253, // CAACTA
    82.48828,  1.58806, 1.13613, 0.94042, // CAACTC
    83.96413,  1.58806, 1.21996, 1.04640, // CAACTG
    83.09859,  1.58806, 0.92727, 0.69341, // CAACTT
    85.08038,  1.46160, 1.09305, 0.79133, // CAAGAA
    80.37536,  1.46160, 1.41330, 1.16345, // CAAGAC
    84.00512,  1.46160, 1.15152, 0.85567, // CAAGAG
    81.73993,  1.46160, 1.35624, 1.09371, // CAAGAT
    78.59103,  1.42710, 1.19950, 0.91657, // CAAGCA
    77.31484,  1.42710, 1.15734, 0.86867, // CAAGCC
    78.57010,  1.42710, 1.12321, 0.83052, // CAAGCG
    78.12357,  1.42710, 1.05215, 0.75298, // CAAGCT
    83.76934,  1.40989, 1.08415, 0.78258, // CAAGGA
    80.73310,  1.40989, 1.20464, 0.91660, // CAAGGC
    82.70274,  1.40989, 1.04511, 0.74070, // CAAGGG
    81.86256,  1.40989, 1.26150, 0.98226, // CAAGGT
    77.01656,  1.48703, 1.19640, 0.93765, // CAAGTA
    74.72120,  1.48703, 1.49101, 1.30450, // CAAGTC
    76.12311,  1.48703, 1.27961, 1.03715, // CAAGTG
    75.59901,  1.48703, 1.46633, 1.27225, // CAAGTT
    82.59722,  1.98024, 1.58228, 1.50907, // CAATAA
    80.23488,  1.98024, 1.43132, 1.29833, // CAATAC
    81.90842,  1.98024, 1.49654, 1.38808, // CAATAG
    81.37406,  1.98024, 1.56482, 1.48416, // CAATAT
    75.75470,  2.22770, 1.78694, 2.05373, // CAATCA
    74.65851,  2.22770, 1.87371, 2.20511, // CAATCC
    75.96251,  2.22770, 1.80635, 2.08727, // CAATCG
    75.74439,  2.22770, 1.87779, 2.21231, // CAATCT
    79.91138,  1.67617, 1.36550, 1.24621, // CAATGA
    77.48927,  1.67617, 1.49675, 1.43014, // CAATGC
    79.04619,  1.67617, 1.43102, 1.33697, // CAATGG
    78.29229,  1.67617, 1.54090, 1.49388, // CAATGT
    70.65836,  2.00867, 1.93329, 1.95732, // CAATTA
    67.95390,  2.00867, 2.04579, 2.13063, // CAATTC
    70.11130,  2.00867, 1.95478, 1.99004, // CAATTG
    69.20659,  2.00867, 1.83616, 1.81167, // CAATTT
    106.55732, 2.19132, 1.62999, 1.33064, // CACAAA
    103.61414, 2.19132, 1.56215, 1.24844, // CACAAC
    105.30123, 2.19132, 1.59723, 1.29073, // CACAAG
    104.86095, 2.19132, 1.81583, 1.56457, // CACAAT
    98.03615,  1.74166, 1.07277, 0.69823, // CACACA
    96.60375,  1.74166, 1.09403, 0.71909, // CACACC
    97.96045,  1.74166, 1.08854, 0.71368, // CACACG
    97.86225,  1.74166, 1.20908, 0.83545, // CACACT
    103.50060, 2.28987, 1.63204, 1.32031, // CACAGA
    100.77056, 2.28987, 1.50958, 1.17454, // CACAGC
    102.44903, 2.28987, 1.50902, 1.17388, // CACAGG
    101.98982, 2.28987, 1.64728, 1.33886, // CACAGT
    97.52787,  2.17760, 1.21455, 0.88930, // CACATA
    95.87153,  2.17760, 1.32818, 1.01698, // CACATC
    96.93895,  2.17760, 1.26461, 0.94485, // CACATG
    96.99620,  2.17760, 1.56600, 1.30200, // CACATT
    106.62476, 1.88264, 1.27184, 1.02419, // CACCAA
    103.98404, 1.88264, 1.31919, 1.08192, // CACCAC
    105.85818, 1.88264, 1.28414, 1.03909, // CACCAG
    105.24145, 1.88264, 1.51882, 1.33658, // CACCAT
    102.70828, 1.75332, 1.11442, 0.84757, // CACCCA
    100.91353, 1.75332, 1.08234, 0.81124, // CACCCC
    102.63448, 1.75332, 1.10970, 0.84219, // CACCCG
    101.95535, 1.75332, 1.20743, 0.95586, // CACCCT
    105.21069, 1.90443, 1.40549, 1.14491, // CACCGA
    102.64821, 1.90443, 1.30003, 1.01849, // CACCGC
    104.20774, 1.90443, 1.29735, 1.01535, // CACCGG
    103.62328, 1.90443, 1.40655, 1.14621, // CACCGT
    101.25187, 1.83098, 1.30625, 0.97574, // CACCTA
    99.67359,  1.83098, 1.25861, 0.92285, // CACCTC
    100.97987, 1.83098, 1.28755, 0.95487, // CACCTG
    100.43581, 1.83098, 1.45893, 1.15173, // CACCTT
    101.11489, 2.24722, 1.49862, 1.19211, // CACGAA
    96.29776,  2.24722, 1.40991, 1.08784, // CACGAC
    99.93821,  2.24722, 1.46548, 1.15279, // CACGAG
    98.37729,  2.24722, 1.62384, 1.34460, // CACGAT
    95.92705,  2.02317, 1.13880, 0.78694, // CACGCA
    94.73979,  2.02317, 1.18990, 0.84049, // CACGCC
    96.11709,  2.02317, 1.19548, 0.84641, // CACGCG
    96.04508,  2.02317, 1.30168, 0.96167, // CACGCT
    100.66809, 2.22668, 1.61661, 1.32867, // CACGGA
    97.29486,  2.22668, 1.41218, 1.08478, // CACGGC
    99.66144,  2.22668, 1.49791, 1.18505, // CACGGG
    98.94610,  2.22668, 1.62796, 1.34268, // CACGGT
    95.22340,  2.64556, 1.33682, 0.96556, // CACGTA
    93.11048,  2.64556, 1.50426, 1.15255, // CACGTC
    94.24708,  2.64556, 1.47420, 1.11817, // CACGTG
    94.85048,  2.64556, 1.68166, 1.36232, // CACGTT
    100.32149, 1.69847, 1.40820, 0.98708, // CACTAA
    96.98708,  1.69847, 1.28716, 0.86259, // CACTAC
    99.58239,  1.69847, 1.34421, 0.92057, // CACTAG
    98.35078,  1.69847, 1.38804, 0.96596, // CACTAT
    92.08676,  1.86136, 1.49377, 1.30937, // CACTCA
    90.53936,  1.86136, 1.61647, 1.47398, // CACTCC
    92.25001,  1.86136, 1.59089, 1.43913, // CACTCG
    92.04478,  1.86136, 1.59545, 1.44532, // CACTCT
    97.10306,  1.64486, 1.33084, 0.99967, // CACTGA
    93.49241,  1.64486, 1.24836, 0.90819, // CACTGC
    95.85222,  1.64486, 1.27417, 0.93650, // CACTGG
    94.70450,  1.64486, 1.29863, 0.96360, // CACTGT
    87.39047,  1.46961, 1.20250, 0.87086, // CACTTA
    84.56300,  1.46961, 1.24603, 0.91857, // CACTTC
    86.62217,  1.46961, 1.13987, 0.80372, // CACTTG
    85.83269,  1.46961, 1.07642, 0.73755, // CACTTT
    78.43748,  1.82916, 1.26788, 0.85510, // CAGAAA
    75.85913,  1.82916, 1.56883, 1.17696, // CAGAAC
    77.08668,  1.82916, 1.33389, 0.92273, // CAGAAG
    76.30950,  1.82916, 1.54722, 1.15273, // CAGAAT
    67.34887,  2.14481, 1.86316, 1.48913, // CAGACA
    66.32630,  2.14481, 1.87829, 1.50730, // CAGACC
    67.34776,  2.14481, 1.84872, 1.47185, // CAGACG
    67.48376,  2.14481, 1.81787, 1.43517, // CAGACT
    76.42675,  1.82854, 1.31162, 0.97512, // CAGAGA
    74.86750,  1.82854, 1.40216, 1.07781, // CAGAGC
    75.18738,  1.82854, 1.37510, 1.04677, // CAGAGG
    74.96157,  1.82854, 1.42244, 1.10128, // CAGAGT
    67.41734,  1.85459, 1.66387, 1.45195, // CAGATA
    66.16665,  1.85459, 1.44213, 1.17160, // CAGATC
    66.84723,  1.85459, 1.55884, 1.31666, // CAGATG
    66.55666,  1.85459, 1.54524, 1.29948, // CAGATT
    86.69550,  1.87706, 1.50906, 1.26558, // CAGCAA
    84.42123,  1.87706, 1.23853, 0.94100, // CAGCAC
    85.65235,  1.87706, 1.37079, 1.09569, // CAGCAG
    85.15599,  1.87706, 1.25884, 0.96424, // CAGCAT
    82.36197,  1.70618, 1.52389, 1.37369, // CAGCCA
    80.84537,  1.70618, 1.38859, 1.19487, // CAGCCC
    82.26861,  1.70618, 1.49104, 1.32951, // CAGCCG
    81.92858,  1.70618, 1.39173, 1.19893, // CAGCCT
    84.75762,  1.56014, 1.31804, 1.09096, // CAGCGA
    82.01971,  1.56014, 1.18841, 0.93404, // CAGCGC
    83.55608,  1.56014, 1.33545, 1.11265, // CAGCGG
    82.52782,  1.56014, 1.24375, 1.00004, // CAGCGT
    80.88727,  1.42173, 1.26239, 0.99727, // CAGCTA
    79.24282,  1.42173, 1.11691, 0.82993, // CAGCTC
    80.43945,  1.42173, 1.17465, 0.89511, // CAGCTG
    79.90986,  1.42173, 0.99619, 0.69909, // CAGCTT
    77.78104,  1.83346, 1.30527, 0.91775, // CAGGAA
    72.96374,  1.83346, 1.71768, 1.38543, // CAGGAC
    76.56782,  1.83346, 1.40180, 1.02141, // CAGGAG
    74.35727,  1.83346, 1.65844, 1.31439, // CAGGAT
    72.08294,  1.67690, 1.50261, 1.23637, // CAGGCA
    70.63449,  1.67690, 1.48910, 1.21974, // CAGGCC
    71.92395,  1.67690, 1.44127, 1.16143, // CAGGCG
    71.68837,  1.67690, 1.31434, 1.01143, // CAGGCT
    78.00536,  1.68720, 1.18922, 0.86688, // CAGGGA
    75.37337,  1.68720, 1.37318, 1.07562, // CAGGGC
    76.96801,  1.68720, 1.16083, 0.83602, // CAGGGG
    76.11175,  1.68720, 1.40579, 1.11416, // CAGGGT
    70.96085,  1.76089, 1.26601, 0.93187, // CAGGTA
    69.21876,  1.76089, 1.56124, 1.27615, // CAGGTC
    70.35166,  1.76089, 1.32927, 1.00258, // CAGGTG
    70.18719,  1.76089, 1.54529, 1.25665, // CAGGTT
    76.53448,  2.04805, 1.59630, 1.43276, // CAGTAA
    73.40594,  2.04805, 1.68936, 1.55986, // CAGTAC
    75.87202,  2.04805, 1.60659, 1.44663, // CAGTAG
    74.76321,  2.04805, 1.75277, 1.64850, // CAGTAT
    69.06535,  2.42133, 2.22866, 2.44113, // CAGTCA
    67.81113,  2.42133, 2.23318, 2.44855, // CAGTCC
    69.27582,  2.42133, 2.08051, 2.20180, // CAGTCG
    68.96241,  2.42133, 2.22287, 2.43162, // CAGTCT
    73.20490,  1.99693, 1.67817, 1.64669, // CAGTGA
    69.65252,  1.99693, 2.00998, 2.15846, // CAGTGC
    72.15619,  1.99693, 1.72222, 1.71194, // CAGTGG
    70.66767,  1.99693, 1.92874, 2.02894, // CAGTGT
    64.15501,  2.35636, 2.28022, 2.27826, // CAGTTA
    61.21348,  2.35636, 2.24417, 2.22444, // CAGTTC
    63.58264,  2.35636, 2.17729, 2.12576, // CAGTTG
    62.57969,  2.35636, 2.17873, 2.12787, // CAGTTT
    108.22131, 3.16286, 2.14775, 1.66853, // CATAAA
    105.45731, 3.16286, 2.25084, 1.79009, // CATAAC
    107.12266, 3.16286, 2.15662, 1.67887, // CATAAG
    107.65532, 3.16286, 2.42710, 2.00442, // CATAAT
    102.51687, 3.09766, 1.83692, 1.31419, // CATACA
    102.72060, 3.09766, 1.94011, 1.42647, // CATACC
    102.72067, 3.09766, 1.89805, 1.38034, // CATACG
    104.14664, 3.09766, 2.17161, 1.68925, // CATACT
    105.39625, 3.02439, 2.13016, 1.77584, // CATAGA
    103.97977, 3.02439, 2.06723, 1.69772, // CATAGC
    104.79926, 3.02439, 2.09827, 1.73611, // CATAGG
    106.28338, 3.02439, 2.13666, 1.78397, // CATAGT
    103.60892, 3.75132, 2.25257, 1.72743, // CATATA
    104.56725, 3.75132, 2.48025, 1.99583, // CATATC
    102.74219, 3.75132, 2.25924, 1.73510, // CATATG
    106.70454, 3.75132, 2.69236, 2.25726, // CATATT
    117.50043, 2.97261, 2.40814, 2.47228, // CATCAA
    115.17048, 2.97261, 2.42047, 2.49130, // CATCAC
    116.82723, 2.97261, 2.42200, 2.49366, // CATCAG
    116.39495, 2.97261, 2.51480, 2.63835, // CATCAT
    115.89070, 3.04768, 2.43822, 2.40157, // CATCCA
    114.18528, 3.04768, 2.37485, 2.30857, // CATCCC
    116.28932, 3.04768, 2.41774, 2.37137, // CATCCG
    115.52491, 3.04768, 2.49477, 2.48561, // CATCCT
    116.92871, 3.00027, 2.52931, 2.55266, // CATCGA
    114.21307, 3.00027, 2.34075, 2.27260, // CATCGC
    116.30778, 3.00027, 2.43944, 2.41782, // CATCGG
    116.18255, 3.00027, 2.48651, 2.48815, // CATCGT
    116.29742, 3.10410, 2.51726, 2.44762, // CATCTA
    115.05333, 3.10410, 2.56161, 2.51257, // CATCTC
    116.22001, 3.10410, 2.43656, 2.33086, // CATCTG
    116.22684, 3.10410, 2.77596, 2.83446, // CATCTT
    104.67927, 2.99484, 2.09340, 1.75070, // CATGAA
    100.85987, 2.99484, 2.04153, 1.68603, // CATGAC
    103.63814, 2.99484, 2.04858, 1.69478, // CATGAG
    103.23510, 2.99484, 2.30184, 2.01858, // CATGAT
    103.26977, 3.13415, 1.93940, 1.50279, // CATGCA
    102.87227, 3.13415, 2.03240, 1.61217, // CATGCC
    103.20089, 3.13415, 2.02635, 1.60499, // CATGCG
    104.62381, 3.13415, 2.20328, 1.81972, // CATGCT
    105.09490, 2.93123, 2.21219, 1.85653, // CATGGA
    102.79268, 2.93123, 2.04718, 1.65274, // CATGGC
    104.33709, 2.93123, 2.09882, 1.71566, // CATGGG
    104.99463, 2.93123, 2.25863, 1.91529, // CATGGT
    105.52403, 3.56800, 2.46314, 2.03729, // CATGTA
    106.16709, 3.56800, 2.56727, 2.16783, // CATGTC
    105.33589, 3.56800, 2.42090, 1.98510, // CATGTG
    108.29126, 3.56800, 2.71499, 2.35760, // CATGTT
    104.49619, 2.18027, 1.93972, 1.94918, // CATTAA
    100.91951, 2.18027, 1.64005, 1.51541, // CATTAC
    103.72965, 2.18027, 1.83686, 1.79622, // CATTAG
    102.67349, 2.18027, 1.71403, 1.61909, // CATTAT
    98.48690,  2.09022, 1.95374, 1.85508, // CATTCA
    96.59479,  2.09022, 1.77414, 1.60525, // CATTCC
    98.37010,  2.09022, 1.86800, 1.73432, // CATTCG
    98.26777,  2.09022, 1.96180, 1.86657, // CATTCT
    102.40200, 2.20088, 1.76540, 1.65559, // CATTGA
    98.27582,  2.20088, 1.65904, 1.50824, // CATTGC
    101.31999, 2.20088, 1.77785, 1.67313, // CATTGG
    99.92969,  2.20088, 1.77428, 1.66810, // CATTGT
    94.48623,  1.92511, 1.65966, 1.45626, // CATTTA
    92.21787,  1.92511, 1.51390, 1.26870, // CATTTC
    93.98173,  1.92511, 1.61291, 1.39517, // CATTTG
    94.05437,  1.92511, 1.44413, 1.18201, // CATTTT
    92.48449,  2.70987, 1.47522, 1.06177, // CCAAAA
    89.46815,  2.70987, 1.71315, 1.32873, // CCAAAC
    91.43517,  2.70987, 1.53251, 1.12421, // CCAAAG
    90.15980,  2.70987, 1.78962, 1.41867, // CCAAAT
    83.15695,  2.16112, 1.61209, 1.33224, // CCAACA
    81.84383,  2.16112, 1.45955, 1.14770, // CCAACC
    82.99686,  2.16112, 1.50544, 1.20225, // CCAACG
    82.75683,  2.16112, 1.43315, 1.11670, // CCAACT
    91.20168,  2.81796, 1.54275, 1.17766, // CCAAGA
    88.22276,  2.81796, 1.57504, 1.21482, // CCAAGC
    90.14237,  2.81796, 1.45401, 1.07752, // CCAAGG
    88.52196,  2.81796, 1.76295, 1.43858, // CCAAGT
    81.97053,  2.24890, 1.45837, 1.16529, // CCAATA
    80.36940,  2.24890, 1.86483, 1.68497, // CCAATC
    81.44488,  2.24890, 1.61926, 1.36335, // CCAATG
    80.85945,  2.24890, 1.81600, 1.61922, // CCAATT
    95.50360,  1.22481, 1.00045, 0.74105, // CCACAA
    92.84240,  1.22481, 0.91946, 0.65291, // CCACAC
    94.62827,  1.22481, 0.97627, 0.71435, // CCACAG
    94.19127,  1.22481, 0.90524, 0.63782, // CCACAT
    92.22045,  1.29044, 1.07628, 0.82961, // CCACCA
    90.85456,  1.29044, 1.07630, 0.82963, // CCACCC
    92.04114,  1.29044, 1.03503, 0.78238, // CCACCG
    91.65253,  1.29044, 0.96936, 0.70911, // CCACCT
    94.31410,  1.39798, 0.91034, 0.63265, // CCACGA
    91.81315,  1.39798, 0.99786, 0.72604, // CCACGC
    93.36288,  1.39798, 0.93863, 0.66237, // CCACGG
    93.18299,  1.39798, 0.96166, 0.68690, // CCACGT
    90.53136,  1.56257, 1.06684, 0.76138, // CCACTA
    88.26866,  1.56257, 1.02163, 0.71351, // CCACTC
    89.92248,  1.56257, 1.05351, 0.74716, // CCACTG
    89.26021,  1.56257, 1.05064, 0.74411, // CCACTT
    88.31136,  2.60118, 1.61217, 1.22530, // CCAGAA
    82.79872,  2.60118, 1.90210, 1.57028, // CCAGAC
    87.06283,  2.60118, 1.69234, 1.31782, // CCAGAG
    84.50436,  2.60118, 1.87194, 1.53307, // CCAGAT
    82.94839,  2.08236, 1.37063, 1.12535, // CCAGCA
    81.39129,  2.08236, 1.37632, 1.13236, // CCAGCC
    82.89686,  2.08236, 1.35986, 1.11211, // CCAGCG
    82.73071,  2.08236, 1.46891, 1.24854, // CCAGCT
    87.78580,  2.50147, 1.57422, 1.24853, // CCAGGA
    83.97988,  2.50147, 1.67091, 1.36531, // CCAGGC
    86.47442,  2.50147, 1.56698, 1.23993, // CCAGGG
    85.40992,  2.50147, 1.78419, 1.50648, // CCAGGT
    80.23698,  2.40625, 1.69188, 1.43749, // CCAGTA
    76.87663,  2.40625, 1.89487, 1.70380, // CCAGTC
    78.95811,  2.40625, 1.69845, 1.44587, // CCAGTG
    78.25879,  2.40625, 1.99376, 1.83890, // CCAGTT
    89.03524,  1.95200, 1.44379, 1.09976, // CCATAA
    85.64921,  1.95200, 1.60414, 1.28796, // CCATAC
    88.22545,  1.95200, 1.41055, 1.06200, // CCATAG
    87.23583,  1.95200, 1.56555, 1.24177, // CCATAT
    82.08498,  2.22932, 2.06212, 2.04548, // CCATCA
    80.58589,  2.22932, 2.11889, 2.13052, // CCATCC
    82.13112,  2.22932, 2.00535, 1.96160, // CCATCG
    81.80091,  2.22932, 2.01893, 1.98156, // CCATCT
    86.32370,  2.18506, 1.53708, 1.20605, // CCATGA
    82.62480,  2.18506, 1.72595, 1.43504, // CCATGC
    85.39210,  2.18506, 1.62637, 1.31266, // CCATGG
    84.09425,  2.18506, 1.83548, 1.57378, // CCATGT
    76.61840,  2.43089, 2.15111, 2.03502, // CCATTA
    73.31518,  2.43089, 2.03530, 1.87291, // CCATTC
    75.69504,  2.43089, 2.05684, 1.90272, // CCATTG
    74.92189,  2.43089, 1.99302, 1.81486, // CCATTT
    105.19016, 1.63085, 1.15916, 0.92637, // CCCAAA
    102.70161, 1.63085, 1.28581, 1.08226, // CCCAAC
    103.85233, 1.63085, 1.15443, 0.92070, // CCCAAG
    103.60858, 1.63085, 1.41371, 1.24770, // CCCAAT
    97.12289,  1.46322, 0.93570, 0.62631, // CCCACA
    95.73720,  1.46322, 0.98169, 0.67305, // CCCACC
    96.70835,  1.46322, 0.90501, 0.59575, // CCCACG
    96.72498,  1.46322, 1.03604, 0.72971, // CCCACT
    102.43429, 1.84203, 1.31807, 1.05818, // CCCAGA
    99.98067,  1.84203, 1.35566, 1.10377, // CCCAGC
    101.24734, 1.84203, 1.23456, 0.95921, // CCCAGG
    100.67631, 1.84203, 1.54810, 1.34694, // CCCAGT
    95.92987,  1.79927, 1.16042, 0.80779, // CCCATA
    94.08101,  1.79927, 1.32364, 0.98409, // CCCATC
    95.29163,  1.79927, 1.23702, 0.88909, // CCCATG
    95.03883,  1.79927, 1.51880, 1.20957, // CCCATT
    104.58109, 1.44981, 0.92138, 0.66435, // CCCCAA
    101.84125, 1.44981, 0.90989, 0.65196, // CCCCAC
    103.76841, 1.44981, 0.90080, 0.64222, // CCCCAG
    103.06699, 1.44981, 0.99552, 0.74613, // CCCCAT
    100.32566, 1.40182, 0.85028, 0.60057, // CCCCCA
    98.89078,  1.40182, 0.80402, 0.55223, // CCCCCC
    100.48122, 1.40182, 0.84418, 0.59411, // CCCCCG
    99.96987,  1.40182, 0.81810, 0.56679, // CCCCCT
    102.83618, 1.43192, 0.99724, 0.76130, // CCCCGA
    100.50779, 1.43192, 0.93336, 0.68934, // CCCCGC
    101.99010, 1.43192, 0.91479, 0.66886, // CCCCGG
    101.55851, 1.43192, 1.00540, 0.77066, // CCCCGT
    99.42695,  1.44710, 0.90556, 0.65673, // CCCCTA
    97.44506,  1.44710, 1.02178, 0.78713, // CCCCTC
    98.51893,  1.44710, 0.90849, 0.65992, // CCCCTG
    98.07747,  1.44710, 1.06242, 0.83456, // CCCCTT
    99.51858,  1.82025, 1.22151, 0.94645, // CCCGAA
    94.99282,  1.82025, 1.47034, 1.24992, // CCCGAC
    98.38138,  1.82025, 1.23197, 0.95864, // CCCGAG
    96.70874,  1.82025, 1.48032, 1.26267, // CCCGAT
    94.80287,  1.58093, 0.95819, 0.62153, // CCCGCA
    93.50752,  1.58093, 1.06394, 0.72722, // CCCGCC
    94.69036,  1.58093, 1.03922, 0.70202, // CCCGCG
    94.69532,  1.58093, 1.12764, 0.79349, // CCCGCT
    99.10076,  1.74812, 1.30342, 1.08160, // CCCGGA
    95.90636,  1.74812, 1.34606, 1.13510, // CCCGGC
    98.03599,  1.74812, 1.23281, 0.99491, // CCCGGG
    97.38603,  1.74812, 1.45623, 1.27728, // CCCGGT
    93.23158,  2.13467, 1.28975, 1.01180, // CCCGTA
    90.78889,  2.13467, 1.59479, 1.39120, // CCCGTC
    92.54264,  2.13467, 1.40844, 1.15462, // CCCGTG
    92.40885,  2.13467, 1.65942, 1.47662, // CCCGTT
    99.46478,  1.77408, 1.19425, 0.89636, // CCCTAA
    96.54081,  1.77408, 1.25172, 0.96183, // CCCTAC
    98.26053,  1.77408, 1.11690, 0.81070, // CCCTAG
    97.58854,  1.77408, 1.23985, 0.94818, // CCCTAT
    92.63162,  2.16210, 1.61561, 1.42556, // CCCTCA
    90.87105,  2.16210, 1.58668, 1.38745, // CCCTCC
    92.29789,  2.16210, 1.50531, 1.28210, // CCCTCG
    92.35864,  2.16210, 1.66382, 1.48985, // CCCTCT
    96.12946,  1.63830, 1.11899, 0.81853, // CCCTGA
    93.04349,  1.63830, 1.25832, 0.97607, // CCCTGC
    94.99957,  1.63830, 1.17616, 0.88206, // CCCTGG
    94.10107,  1.63830, 1.24420, 0.95969, // CCCTGT
    87.24337,  1.91311, 1.44426, 1.14723, // CCCTTA
    84.41210,  1.91311, 1.61398, 1.35528, // CCCTTC
    86.83067,  1.91311, 1.48188, 1.19234, // CCCTTG
    85.66470,  1.91311, 1.46142, 1.16774, // CCCTTT
    85.05826,  2.36795, 1.49522, 1.08987, // CCGAAA
    81.97938,  2.36795, 1.82372, 1.46809, // CCGAAC
    83.85231,  2.36795, 1.56292, 1.16472, // CCGAAG
    82.80197,  2.36795, 1.79978, 1.43929, // CCGAAT
    74.97206,  2.00332, 1.65424, 1.35394, // CCGACA
    73.88856,  2.00332, 1.72083, 1.43650, // CCGACC
    74.97089,  2.00332, 1.63414, 1.32933, // CCGACG
    74.85252,  2.00332, 1.53845, 1.21429, // CCGACT
    83.74456,  2.41291, 1.48333, 1.08044, // CCGAGA
    81.14395,  2.41291, 1.55549, 1.16023, // CCGAGC
    82.47967,  2.41291, 1.43404, 1.02703, // CCGAGG
    81.46509,  2.41291, 1.68807, 1.31168, // CCGAGT
    73.94447,  1.85855, 1.45498, 1.27760, // CCGATA
    71.90463,  1.85855, 1.67125, 1.57280, // CCGATC
    73.21882,  1.85855, 1.51441, 1.35667, // CCGATG
    73.04808,  1.85855, 1.81180, 1.77532, // CCGATT
    91.41069,  1.44291, 1.11817, 0.82754, // CCGCAA
    88.73922,  1.44291, 1.07078, 0.77549, // CCGCAC
    90.40692,  1.44291, 1.13480, 0.84608, // CCGCAG
    89.64388,  1.44291, 1.05717, 0.76076, // CCGCAT
    87.52690,  1.52741, 1.33680, 1.08935, // CCGCCA
    86.14175,  1.52741, 1.28265, 1.02383, // CCGCCC
    87.36814,  1.52741, 1.29479, 1.03840, // CCGCCG
    86.97927,  1.52741, 1.22652, 0.95737, // CCGCCT
    89.55020,  1.58181, 1.14925, 0.83221, // CCGCGA
    86.46680,  1.58181, 1.24933, 0.94325, // CCGCGC
    88.24212,  1.58181, 1.22603, 0.91700, // CCGCGG
    87.17906,  1.58181, 1.19613, 0.88365, // CCGCGT
    85.73738,  1.56040, 1.33556, 1.02622, // CCGCTA
    83.37869,  1.56040, 1.31790, 1.00593, // CCGCTC
    84.94247,  1.56040, 1.29590, 0.98085, // CCGCTG
    84.26868,  1.56040, 1.10545, 0.77277, // CCGCTT
    84.65142,  2.44172, 1.65995, 1.25117, // CCGGAA
    79.25381,  2.44172, 2.00944, 1.66643, // CCGGAC
    83.32889,  2.44172, 1.73850, 1.34104, // CCGGAG
    80.91036,  2.44172, 2.02037, 1.68005, // CCGGAT
    79.52553,  2.03186, 1.47211, 1.18329, // CCGGCA
    78.08105,  2.03186, 1.46493, 1.17464, // CCGGCC
    79.29594,  2.03186, 1.43488, 1.13869, // CCGGCG
    79.37801,  2.03186, 1.42772, 1.13018, // CCGGCT
    85.14868,  2.35270, 1.56743, 1.14007, // CCGGGA
    82.15414,  2.35270, 1.70967, 1.29874, // CCGGGC
    84.03296,  2.35270, 1.49843, 1.06563, // CCGGGG
    83.07996,  2.35270, 1.72738, 1.31897, // CCGGGT
    77.73256,  2.46096, 1.65744, 1.33167, // CCGGTA
    74.95130,  2.46096, 1.90555, 1.64162, // CCGGTC
    76.55398,  2.46096, 1.77825, 1.47989, // CCGGTG
    76.08887,  2.46096, 1.98292, 1.74261, // CCGGTT
    82.22761,  2.14468, 1.74967, 1.49645, // CCGTAA
    78.41309,  2.14468, 1.82580, 1.59518, // CCGTAC
    81.08679,  2.14468, 1.78410, 1.54084, // CCGTAG
    80.25475,  2.14468, 1.90168, 1.69565, // CCGTAT
    74.78274,  2.49565, 2.27147, 2.50741, // CCGTCA
    73.48590,  2.49565, 2.25938, 2.48741, // CCGTCC
    75.07600,  2.49565, 2.35403, 2.64534, // CCGTCG
    74.64642,  2.49565, 2.31295, 2.57640, // CCGTCT
    78.50660,  2.22005, 1.87389, 1.73372, // CCGTGA
    74.74633,  2.22005, 2.13354, 2.10628, // CCGTGC
    77.32992,  2.22005, 1.97566, 1.87686, // CCGTGG
    76.05888,  2.22005, 2.19043, 2.19108, // CCGTGT
    69.34909,  2.47313, 2.27971, 2.31799, // CCGTTA
    66.14638,  2.47313, 2.33794, 2.40736, // CCGTTC
    68.79461,  2.47313, 2.32479, 2.38708, // CCGTTG
    67.57072,  2.47313, 2.23380, 2.24831, // CCGTTT
    111.56102, 2.64098, 2.00895, 1.60322, // CCTAAA
    109.24124, 2.64098, 2.04016, 1.64072, // CCTAAC
    110.80713, 2.64098, 2.09390, 1.70598, // CCTAAG
    111.17403, 2.64098, 2.26104, 1.91425, // CCTAAT
    106.84759, 2.58253, 1.72467, 1.27173, // CCTACA
    106.57158, 2.58253, 1.86359, 1.42844, // CCTACC
    106.59891, 2.58253, 1.77169, 1.32409, // CCTACG
    108.17754, 2.58253, 1.85240, 1.41560, // CCTACT
    109.99890, 2.51606, 1.99770, 1.76214, // CCTAGA
    107.27774, 2.51606, 1.78137, 1.48381, // CCTAGC
    109.55804, 2.51606, 1.67283, 1.35028, // CCTAGG
    108.81934, 2.51606, 2.36546, 2.27049, // CCTAGT
    106.69030, 3.03892, 2.08422, 1.59661, // CCTATA
    107.28340, 3.03892, 2.08957, 1.60277, // CCTATC
    106.38890, 3.03892, 2.14633, 1.66851, // CCTATG
    108.93144, 3.03892, 2.36658, 1.93182, // CCTATT
    117.34217, 2.42793, 2.04315, 1.89443, // CCTCAA
    114.91764, 2.42793, 1.88952, 1.68482, // CCTCAC
    116.88408, 2.42793, 2.02384, 1.86763, // CCTCAG
    116.11351, 2.42793, 2.21849, 2.14345, // CCTCAT
    116.09311, 2.47041, 1.94381, 1.67026, // CCTCCA
    114.19616, 2.47041, 1.96818, 1.70176, // CCTCCC
    116.19286, 2.47041, 1.89371, 1.60610, // CCTCCG
    115.63091, 2.47041, 2.14908, 1.94169, // CCTCCT
    117.13667, 2.49891, 2.03913, 1.79660, // CCTCGA
    114.05355, 2.49891, 1.99874, 1.74348, // CCTCGC
    116.45515, 2.49891, 2.02864, 1.78275, // CCTCGG
    115.99207, 2.49891, 2.26022, 2.09657, // CCTCGT
    115.79001, 2.72204, 2.12746, 1.89757, // CCTCTA
    114.38041, 2.72204, 2.18690, 1.97765, // CCTCTC
    115.32904, 2.72204, 2.23761, 2.04684, // CCTCTG
    115.61773, 2.72204, 2.29600, 2.12748, // CCTCTT
    107.36810, 2.50593, 2.00353, 1.66094, // CCTGAA
    103.39825, 2.50593, 1.93335, 1.57444, // CCTGAC
    106.44910, 2.50593, 1.95978, 1.60684, // CCTGAG
    105.47241, 2.50593, 2.17999, 1.88514, // CCTGAT
    105.22691, 2.56584, 1.77168, 1.31593, // CCTGCA
    104.57382, 2.56584, 1.83748, 1.38993, // CCTGCC
    105.13309, 2.56584, 1.82080, 1.37104, // CCTGCG
    105.99859, 2.56584, 1.99585, 1.57344, // CCTGCT
    107.86286, 2.47346, 1.93532, 1.59835, // CCTGGA
    104.79034, 2.47346, 1.92086, 1.58047, // CCTGGC
    106.73592, 2.47346, 1.95588, 1.62388, // CCTGGG
    106.97619, 2.47346, 2.07576, 1.77545, // CCTGGT
    105.63187, 3.11250, 2.12047, 1.62919, // CCTGTA
    105.42110, 3.11250, 2.27390, 1.80918, // CCTGTC
    105.29760, 3.11250, 2.22370, 1.74961, // CCTGTG
    107.42442, 3.11250, 2.46420, 2.04098, // CCTGTT
    106.48452, 1.86007, 1.45840, 1.10923, // CCTTAA
    102.31224, 1.86007, 1.28920, 0.92191, // CCTTAC
    105.46701, 1.86007, 1.45809, 1.10888, // CCTTAG
    104.29298, 1.86007, 1.41457, 1.05961, // CCTTAT
    100.10352, 1.79286, 1.47500, 1.14506, // CCTTCA
    98.18619,  1.79286, 1.51484, 1.19177, // CCTTCC
    100.13250, 1.79286, 1.48822, 1.16049, // CCTTCG
    99.85296,  1.79286, 1.51504, 1.19200, // CCTTCT
    104.17926, 1.88803, 1.41369, 1.10946, // CCTTGA
    100.40203, 1.88803, 1.28826, 0.96513, // CCTTGC
    103.28348, 1.88803, 1.32410, 1.00568, // CCTTGG
    102.06027, 1.88803, 1.37744, 1.06706, // CCTTGT
    96.08204,  1.89728, 1.41445, 1.03620, // CCTTTA
    94.18767,  1.89728, 1.26949, 0.88106, // CCTTTC
    95.79528,  1.89728, 1.38068, 0.99932, // CCTTTG
    95.74019,  1.89728, 1.34993, 0.96611, // CCTTTT
    80.61632,  1.67748, 1.10686, 0.74223, // CGAAAA
    78.22116,  1.67748, 1.24776, 0.88837, // CGAAAC
    79.49026,  1.67748, 1.12708, 0.76266, // CGAAAG
    78.70362,  1.67748, 1.26078, 0.90230, // CGAAAT
    69.69091,  2.18144, 1.82435, 1.54320, // CGAACA
    68.70090,  2.18144, 1.78468, 1.49315, // CGAACC
    69.78103,  2.18144, 1.73312, 1.42891, // CGAACG
    69.70701,  2.18144, 1.64673, 1.32341, // CGAACT
    78.08350,  1.59541, 1.18919, 0.83913, // CGAAGA
    76.83452,  1.59541, 1.12885, 0.77608, // CGAAGC
    77.26550,  1.59541, 1.16480, 0.81345, // CGAAGG
    76.83505,  1.59541, 1.13682, 0.78432, // CGAAGT
    69.86426,  1.95147, 1.72005, 1.43099, // CGAATA
    68.24103,  1.95147, 1.48557, 1.14859, // CGAATC
    69.08545,  1.95147, 1.58568, 1.26662, // CGAATG
    68.63222,  1.95147, 1.37190, 1.01932, // CGAATT
    86.02693,  2.48599, 1.75206, 1.60313, // CGACAA
    84.71163,  2.48599, 1.44087, 1.19559, // CGACAC
    85.20410,  2.48599, 1.64876, 1.46347, // CGACAG
    85.35769,  2.48599, 1.48821, 1.25500, // CGACAT
    82.08572,  2.19720, 1.67253, 1.59099, // CGACCA
    81.03963,  2.19720, 1.52456, 1.38460, // CGACCC
    82.02268,  2.19720, 1.66799, 1.58451, // CGACCG
    82.01235,  2.19720, 1.56258, 1.43672, // CGACCT
    84.74454,  1.98191, 1.47726, 1.31512, // CGACGA
    83.32297,  1.98191, 1.36705, 1.17073, // CGACGC
    83.94138,  1.98191, 1.44387, 1.27078, // CGACGG
    83.69828,  1.98191, 1.45192, 1.28143, // CGACGT
    81.65850,  1.92945, 1.43374, 1.30321, // CGACTA
    80.01686,  1.92945, 1.31443, 1.14397, // CGACTC
    81.13747,  1.92945, 1.43620, 1.30657, // CGACTG
    80.47222,  1.92945, 1.11490, 0.89364, // CGACTT
    80.39930,  1.41776, 1.04069, 0.74929, // CGAGAA
    76.31226,  1.41776, 1.35772, 1.11655, // CGAGAC
    79.45215,  1.41776, 1.11487, 0.83081, // CGAGAG
    77.56085,  1.41776, 1.33854, 1.09299, // CGAGAT
    73.39241,  1.59099, 1.36246, 1.01762, // CGAGCA
    71.99532,  1.59099, 1.37790, 1.03497, // CGAGCC
    73.49537,  1.59099, 1.31894, 0.96926, // CGAGCG
    73.32665,  1.59099, 1.26982, 0.91562, // CGAGCT
    79.39735,  1.35821, 1.01957, 0.70907, // CGAGGA
    77.65581,  1.35821, 1.13177, 0.82929, // CGAGGC
    78.61253,  1.35821, 1.03553, 0.72579, // CGAGGG
    78.11218,  1.35821, 1.13827, 0.83644, // CGAGGT
    72.99261,  1.45706, 1.14076, 0.83490, // CGAGTA
    71.08400,  1.45706, 1.42932, 1.17094, // CGAGTC
    72.58897,  1.45706, 1.28072, 0.99317, // CGAGTG
    72.18960,  1.45706, 1.48256, 1.23696, // CGAGTT
    78.97860,  2.58682, 2.02382, 2.02527, // CGATAA
    76.65542,  2.58682, 1.95614, 1.92454, // CGATAC
    78.00293,  2.58682, 1.86972, 1.79842, // CGATAG
    78.23180,  2.58682, 2.12814, 2.18387, // CGATAT
    71.71263,  2.70241, 2.20719, 2.55022, // CGATCA
    70.48569,  2.70241, 2.11490, 2.39197, // CGATCC
    71.91043,  2.70241, 2.15705, 2.46382, // CGATCG
    71.98006,  2.70241, 2.30373, 2.71936, // CGATCT
    76.02798,  2.05805, 1.81971, 1.90918, // CGATGA
    73.27182,  2.05805, 1.86174, 1.97571, // CGATGC
    74.96712,  2.05805, 1.90529, 2.04544, // CGATGG
    74.30764,  2.05805, 1.99801, 2.19655, // CGATGT
    67.45881,  2.35300, 1.96833, 2.01330, // CGATTA
    64.88022,  2.35300, 2.12497, 2.25835, // CGATTC
    66.91346,  2.35300, 1.94627, 1.97955, // CGATTG
    65.84366,  2.35300, 1.83378, 1.81042, // CGATTT
    106.29852, 2.21768, 1.80415, 1.49037, // CGCAAA
    103.04753, 2.21768, 1.86283, 1.56368, // CGCAAC
    105.28698, 2.21768, 1.85430, 1.55295, // CGCAAG
    104.54742, 2.21768, 1.96340, 1.69200, // CGCAAT
    96.61939,  1.72978, 1.20353, 0.83138, // CGCACA
    95.28675,  1.72978, 1.20961, 0.83769, // CGCACC
    96.66242,  1.72978, 1.23832, 0.86769, // CGCACG
    96.46912,  1.72978, 1.32526, 0.96065, // CGCACT
    103.20676, 2.29394, 1.85311, 1.52547, // CGCAGA
    100.29749, 2.29394, 1.72462, 1.36959, // CGCAGC
    102.17607, 2.29394, 1.75823, 1.40982, // CGCAGG
    101.69526, 2.29394, 1.89408, 1.57634, // CGCAGT
    96.36338,  2.05386, 1.45579, 1.11816, // CGCATA
    94.75573,  2.05386, 1.50372, 1.17384, // CGCATC
    95.68007,  2.05386, 1.45356, 1.11559, // CGCATG
    96.10603,  2.05386, 1.73657, 1.45678, // CGCATT
    107.06578, 1.96521, 1.43974, 1.22018, // CGCCAA
    104.28495, 1.96521, 1.46962, 1.25836, // CGCCAC
    106.39096, 1.96521, 1.49064, 1.28545, // CGCCAG
    105.59818, 1.96521, 1.64296, 1.48744, // CGCCAT
    103.23351, 1.76337, 1.27698, 1.03072, // CGCCCA
    101.29667, 1.76337, 1.24337, 0.99030, // CGCCCC
    102.97258, 1.76337, 1.31795, 1.08072, // CGCCCG
    102.37858, 1.76337, 1.40933, 1.19505, // CGCCCT
    105.60820, 1.93464, 1.63408, 1.49987, // CGCCGA
    103.04052, 1.93464, 1.45061, 1.25450, // CGCCGC
    104.65536, 1.93464, 1.44976, 1.25340, // CGCCGG
    104.13710, 1.93464, 1.62200, 1.48327, // CGCCGT
    101.98089, 1.85738, 1.39477, 1.05828, // CGCCTA
    99.93215,  1.85738, 1.48134, 1.15831, // CGCCTC
    101.40303, 1.85738, 1.42092, 1.08818, // CGCCTG
    100.85404, 1.85738, 1.60245, 1.30323, // CGCCTT
    101.54665, 2.24827, 1.78713, 1.51853, // CGCGAA
    96.95696,  2.24827, 1.68414, 1.38918, // CGCGAC
    100.61981, 2.24827, 1.68677, 1.39243, // CGCGAG
    98.99945,  2.24827, 1.83885, 1.58492, // CGCGAT
    95.75826,  1.86985, 1.34712, 0.98595, // CGCGCA
    94.41418,  1.86985, 1.38165, 1.02410, // CGCGCC
    95.92411,  1.86985, 1.39508, 1.03907, // CGCGCG
    95.88654,  1.86985, 1.54422, 1.21007, // CGCGCT
    101.06557, 2.16392, 1.77581, 1.52929, // CGCGGA
    97.96149,  2.16392, 1.65079, 1.37067, // CGCGGC
    100.10790, 2.16392, 1.68434, 1.41267, // CGCGGG
    99.68512,  2.16392, 1.86464, 1.64546, // CGCGGT
    95.59369,  2.46720, 1.63121, 1.30921, // CGCGTA
    93.79967,  2.46720, 1.72582, 1.42474, // CGCGTC
    95.02257,  2.46720, 1.75540, 1.46152, // CGCGTG
    95.49458,  2.46720, 1.95280, 1.71487, // CGCGTT
    100.48064, 1.80194, 1.56079, 1.22995, // CGCTAA
    97.03599,  1.80194, 1.45010, 1.10145, // CGCTAC
    99.64590,  1.80194, 1.38356, 1.02652, // CGCTAG
    98.50710,  1.80194, 1.48504, 1.14150, // CGCTAT
    91.83596,  2.01289, 1.67968, 1.43265, // CGCTCA
    89.78576,  2.01289, 1.67993, 1.43296, // CGCTCC
    91.80073,  2.01289, 1.67621, 1.42821, // CGCTCG
    91.29607,  2.01289, 1.65113, 1.39627, // CGCTCT
    97.44864,  1.67840, 1.44872, 1.12536, // CGCTGA
    93.77013,  1.67840, 1.37062, 1.03559, // CGCTGC
    95.97398,  1.67840, 1.41488, 1.08615, // CGCTGG
    94.90259,  1.67840, 1.46179, 1.14061, // CGCTGT
    87.63206,  1.51841, 1.36703, 1.05514, // CGCTTA
    84.58854,  1.51841, 1.26005, 0.93373, // CGCTTC
    87.03947,  1.51841, 1.34753, 1.03265, // CGCTTG
    85.83871,  1.51841, 1.13541, 0.79868, // CGCTTT
    73.18831,  1.83378, 1.39908, 0.96728, // CGGAAA
    70.68729,  1.83378, 1.66928, 1.26062, // CGGAAC
    71.89471,  1.83378, 1.44724, 1.01766, // CGGAAG
    71.16753,  1.83378, 1.59950, 1.18241, // CGGAAT
    61.48468,  2.32589, 2.09738, 1.75130, // CGGACA
    60.56091,  2.32589, 2.03665, 1.67579, // CGGACC
    61.59076,  2.32589, 2.05180, 1.69453, // CGGACG
    61.66219,  2.32589, 2.00949, 1.64238, // CGGACT
    70.74671,  1.86110, 1.50747, 1.16694, // CGGAGA
    69.61236,  1.86110, 1.44178, 1.09150, // CGGAGC
    69.95651,  1.86110, 1.48307, 1.13873, // CGGAGG
    69.72196,  1.86110, 1.43658, 1.08560, // CGGAGT
    61.95182,  1.92277, 1.86559, 1.59465, // CGGATA
    60.53403,  1.92277, 1.61557, 1.28508, // CGGATC
    61.39601,  1.92277, 1.73567, 1.43101, // CGGATG
    61.26469,  1.92277, 1.41105, 1.04896, // CGGATT
    85.50017,  1.89614, 1.45737, 1.20710, // CGGCAA
    83.28443,  1.89614, 1.25411, 0.96359, // CGGCAC
    84.45537,  1.89614, 1.40096, 1.13769, // CGGCAG
    84.10385,  1.89614, 1.28960, 1.00478, // CGGCAT
    81.12591,  1.71082, 1.59401, 1.42331, // CGGCCA
    79.79892,  1.71082, 1.48907, 1.28509, // CGGCCC
    80.93330,  1.71082, 1.57437, 1.39708, // CGGCCG
    80.96980,  1.71082, 1.48611, 1.28126, // CGGCCT
    83.81948,  1.57550, 1.35655, 1.10783, // CGGCGA
    81.30843,  1.57550, 1.24370, 0.97250, // CGGCGC
    82.85162,  1.57550, 1.36906, 1.12318, // CGGCGG
    81.93362,  1.57550, 1.27071, 1.00435, // CGGCGT
    79.81521,  1.47431, 1.37075, 1.10011, // CGGCTA
    77.88229,  1.47431, 1.17586, 0.87404, // CGGCTC
    79.20148,  1.47431, 1.27911, 0.99165, // CGGCTG
    78.66933,  1.47431, 0.99993, 0.68542, // CGGCTT
    76.29136,  1.70210, 1.32467, 0.94917, // CGGGAA
    71.53490,  1.70210, 1.61041, 1.27229, // CGGGAC
    75.25186,  1.70210, 1.41396, 1.04673, // CGGGAG
    73.10256,  1.70210, 1.60728, 1.26858, // CGGGAT
    69.54305,  1.76808, 1.69237, 1.34082, // CGGGCA
    67.96891,  1.76808, 1.66763, 1.31152, // CGGGCC
    69.54033,  1.76808, 1.58993, 1.22093, // CGGGCG
    69.28697,  1.76808, 1.48603, 1.10324, // CGGGCT
    76.74035,  1.51256, 1.18037, 0.85278, // CGGGGA
    74.48854,  1.51256, 1.27863, 0.96145, // CGGGGC
    75.82767,  1.51256, 1.12236, 0.79069, // CGGGGG
    74.95259,  1.51256, 1.29713, 0.98240, // CGGGGT
    69.04707,  1.63484, 1.28933, 0.98346, // CGGGTA
    67.10291,  1.63484, 1.53817, 1.28149, // CGGGTC
    68.24370,  1.63484, 1.36037, 1.06585, // CGGGTG
    68.38537,  1.63484, 1.59963, 1.35905, // CGGGTT
    74.95188,  2.38667, 1.66414, 1.42527, // CGGTAA
    72.20048,  2.38667, 1.84154, 1.65914, // CGGTAC
    73.96011,  2.38667, 1.63382, 1.38649, // CGGTAG
    73.40125,  2.38667, 1.78345, 1.58126, // CGGTAT
    67.25835,  2.69199, 2.19819, 2.43115, // CGGTCA
    66.37642,  2.69199, 2.29665, 2.59630, // CGGTCC
    67.59165,  2.69199, 2.22516, 2.47603, // CGGTCG
    67.44332,  2.69199, 2.28429, 2.57537, // CGGTCT
    71.33638,  2.13251, 1.70473, 1.58704, // CGGTGA
    67.70429,  2.13251, 1.93681, 1.92192, // CGGTGC
    70.28187,  2.13251, 1.83037, 1.76569, // CGGTGG
    68.80485,  2.13251, 2.03696, 2.07290, // CGGTGT
    63.06148,  2.62114, 2.29401, 2.29565, // CGGTTA
    60.30055,  2.62114, 2.38254, 2.42981, // CGGTTC
    62.52588,  2.62114, 2.18369, 2.13206, // CGGTTG
    61.46048,  2.62114, 2.13911, 2.06711, // CGGTTT
    107.28313, 2.96792, 2.39383, 1.90328, // CGTAAA
    104.67476, 2.96792, 2.46809, 1.99253, // CGTAAC
    106.26889, 2.96792, 2.40998, 1.92257, // CGTAAG
    106.83595, 2.96792, 2.64964, 2.21637, // CGTAAT
    100.49188, 2.66382, 2.06139, 1.57525, // CGTACA
    100.26306, 2.66382, 2.13489, 1.66024, // CGTACC
    100.36310, 2.66382, 1.98274, 1.48596, // CGTACG
    101.76846, 2.66382, 2.28929, 1.84357, // CGTACT
    104.43415, 2.77966, 2.30021, 1.89738, // CGTAGA
    102.67902, 2.77966, 2.26717, 1.85664, // CGTAGC
    104.21127, 2.77966, 2.21900, 1.79778, // CGTAGG
    104.76315, 2.77966, 2.57992, 2.25379, // CGTAGT
    101.68157, 3.77006, 2.41063, 1.88081, // CGTATA
    102.59785, 3.77006, 2.68063, 2.20547, // CGTATC
    101.35161, 3.77006, 2.40451, 1.87365, // CGTATG
    104.75438, 3.77006, 2.92640, 2.51564, // CGTATT
    116.61618, 2.78871, 2.69276, 2.85580, // CGTCAA
    114.14309, 2.78871, 2.60269, 2.71373, // CGTCAC
    115.60012, 2.78871, 2.26977, 2.21006, // CGTCAG
    115.05073, 2.78871, 2.69324, 2.85656, // CGTCAT
    114.71154, 2.92671, 2.47884, 2.51137, // CGTCCA
    112.62600, 2.92671, 2.50574, 2.55237, // CGTCCC
    114.49881, 2.92671, 2.61106, 2.71496, // CGTCCG
    114.17426, 2.92671, 2.52779, 2.58613, // CGTCCT
    115.67037, 2.90558, 2.74962, 2.90971, // CGTCGA
    113.35969, 2.90558, 2.52887, 2.56643, // CGTCGC
    115.00071, 2.90558, 2.68967, 2.81506, // CGTCGG
    114.89961, 2.90558, 2.73943, 2.89355, // CGTCGT
    114.47505, 3.02381, 2.78705, 2.82575, // CGTCTA
    113.45484, 3.02381, 2.81043, 2.86139, // CGTCTC
    114.45543, 3.02381, 2.71821, 2.72170, // CGTCTG
    114.55539, 3.02381, 3.00914, 3.17017, // CGTCTT
    104.35754, 2.86549, 2.42399, 2.17508, // CGTGAA
    100.46515, 2.86549, 2.42368, 2.17467, // CGTGAC
    103.46031, 2.86549, 2.38396, 2.12142, // CGTGAG
    103.03005, 2.86549, 2.56425, 2.36658, // CGTGAT
    102.53775, 2.92533, 2.16682, 1.80921, // CGTGCA
    101.80738, 2.92533, 2.26172, 1.92936, // CGTGCC
    102.62811, 2.92533, 2.23697, 1.89778, // CGTGCG
    103.79042, 2.92533, 2.40748, 2.11884, // CGTGCT
    104.85398, 2.85435, 2.39764, 2.16159, // CGTGGA
    102.76602, 2.85435, 2.42744, 2.20201, // CGTGGC
    104.54431, 2.85435, 2.36628, 2.11932, // CGTGGG
    105.16465, 2.85435, 2.52981, 2.34276, // CGTGGT
    105.10491, 3.38667, 2.67057, 2.38854, // CGTGTA
    105.44857, 3.38667, 2.78980, 2.55026, // CGTGTC
    104.50054, 3.38667, 2.67322, 2.39209, // CGTGTG
    107.64272, 3.38667, 3.06524, 2.93712, // CGTGTT
    104.18338, 2.40487, 1.96442, 1.92160, // CGTTAA
    100.10445, 2.40487, 1.83609, 1.73641, // CGTTAC
    102.85776, 2.40487, 2.03415, 2.02482, // CGTTAG
    101.48648, 2.40487, 1.92609, 1.86563, // CGTTAT
    96.81851,  2.21733, 2.19092, 2.06982, // CGTTCA
    95.03131,  2.21733, 2.15752, 2.02267, // CGTTCC
    96.98690,  2.21733, 2.22985, 2.12522, // CGTTCG
    96.65888,  2.21733, 2.26738, 2.17911, // CGTTCT
    101.31512, 2.36448, 1.93676, 1.79781, // CGTTGA
    96.96511,  2.36448, 1.92659, 1.78366, // CGTTGC
    100.35532, 2.36448, 1.92492, 1.78135, // CGTTGG
    98.62143,  2.36448, 2.05611, 1.96652, // CGTTGT
    92.90026,  1.91331, 1.97194, 1.85129, // CGTTTA
    90.57875,  1.91331, 1.81369, 1.63297, // CGTTTC
    92.47031,  1.91331, 1.85573, 1.69009, // CGTTTG
    92.34155,  1.91331, 1.67573, 1.45024, // CGTTTT
    93.83520,  2.42434, 1.47917, 1.11499, // CTAAAA
    91.32745,  2.42434, 1.86001, 1.57222, // CTAAAC
    92.99120,  2.42434, 1.56945, 1.21861, // CTAAAG
    92.09466,  2.42434, 1.82407, 1.52688, // CTAAAT
    84.90479,  2.30963, 1.74080, 1.52124, // CTAACA
    84.08237,  2.30963, 1.67422, 1.43481, // CTAACC
    84.73976,  2.30963, 1.73537, 1.51413, // CTAACG
    84.92922,  2.30963, 1.68383, 1.44718, // CTAACT
    92.04232,  2.47862, 1.51978, 1.14226, // CTAAGA
    90.55991,  2.47862, 1.69212, 1.34197, // CTAAGC
    91.31530,  2.47862, 1.61818, 1.25498, // CTAAGG
    90.88775,  2.47862, 1.88894, 1.58279, // CTAAGT
    84.47675,  2.47960, 1.72841, 1.50898, // CTAATA
    82.97981,  2.47960, 2.23227, 2.21480, // CTAATC
    83.52836,  2.47960, 1.93015, 1.78074, // CTAATG
    83.40727,  2.47960, 2.05533, 1.95675, // CTAATT
    95.87570,  1.32011, 1.20590, 0.94473, // CTACAA
    93.95905,  1.32011, 1.07195, 0.79178, // CTACAC
    94.95349,  1.32011, 1.16528, 0.89739, // CTACAG
    94.79761,  1.32011, 1.08282, 0.80385, // CTACAT
    92.87069,  1.41774, 1.32561, 1.02915, // CTACCA
    92.11476,  1.41774, 1.35993, 1.06936, // CTACCC
    92.90431,  1.41774, 1.33390, 1.03881, // CTACCG
    92.48348,  1.41774, 1.19754, 0.88366, // CTACCT
    94.72702,  1.34132, 1.09195, 0.77363, // CTACGA
    92.75737,  1.34132, 1.17049, 0.85858, // CTACGC
    93.93024,  1.34132, 1.21049, 0.90297, // CTACGG
    93.39007,  1.34132, 1.16337, 0.85076, // CTACGT
    91.89184,  1.52174, 1.26907, 0.99713, // CTACTA
    90.11145,  1.52174, 1.18873, 0.90396, // CTACTC
    91.20982,  1.52174, 1.28920, 1.02095, // CTACTG
    91.04497,  1.52174, 1.38162, 1.13267, // CTACTT
    92.03515,  2.32328, 1.48052, 1.17206, // CTAGAA
    86.58915,  2.32328, 1.88099, 1.67846, // CTAGAC
    90.81340,  2.32328, 1.50341, 1.19935, // CTAGAG
    88.66793,  2.32328, 1.57341, 1.28408, // CTAGAT
    86.58766,  2.28919, 1.52284, 1.32929, // CTAGCA
    85.84305,  2.28919, 1.52679, 1.33447, // CTAGCC
    86.41926,  2.28919, 1.59582, 1.42599, // CTAGCG
    86.19618,  2.28919, 1.58859, 1.41631, // CTAGCT
    90.87900,  2.34031, 1.53875, 1.23551, // CTAGGA
    88.13920,  2.34031, 1.56188, 1.26347, // CTAGGC
    90.37954,  2.34031, 1.34794, 1.01297, // CTAGGG
    89.75688,  2.34031, 1.53973, 1.23668, // CTAGGT
    84.70667,  2.72334, 1.77447, 1.61507, // CTAGTA
    81.72098,  2.72334, 2.43672, 2.59896, // CTAGTC
    83.74847,  2.72334, 1.94613, 1.85501, // CTAGTG
    83.10821,  2.72334, 2.11420, 2.10044, // CTAGTT
    90.99058,  1.88263, 1.67039, 1.48457, // CTATAA
    87.95557,  1.88263, 1.85946, 1.74363, // CTATAC
    90.17969,  1.88263, 1.70979, 1.53740, // CTATAG
    89.61549,  1.88263, 1.81465, 1.68097, // CTATAT
    86.42782,  2.46291, 2.37511, 2.51574, // CTATCA
    85.03745,  2.46291, 2.34207, 2.46342, // CTATCC
    86.22179,  2.46291, 2.29368, 2.38747, // CTATCG
    86.11740,  2.46291, 2.38860, 2.53721, // CTATCT
    88.95411,  2.11919, 1.81522, 1.61088, // CTATGA
    86.11184,  2.11919, 2.01192, 1.87968, // CTATGC
    87.61123,  2.11919, 1.82595, 1.62517, // CTATGG
    86.83031,  2.11919, 1.91978, 1.75205, // CTATGT
    81.74064,  2.69506, 2.27582, 2.22039, // CTATTA
    78.84256,  2.69506, 2.51210, 2.57501, // CTATTC
    80.51645,  2.69506, 2.27787, 2.22340, // CTATTG
    79.95026,  2.69506, 2.33309, 2.30473, // CTATTT
    103.75284, 2.08052, 1.46574, 1.18042, // CTCAAA
    101.65030, 2.08052, 1.68072, 1.44943, // CTCAAC
    102.78811, 2.08052, 1.51512, 1.24057, // CTCAAG
    102.47257, 2.08052, 1.75574, 1.54755, // CTCAAT
    97.42459,  2.10126, 1.42485, 1.09627, // CTCACA
    96.40523,  2.10126, 1.46219, 1.13963, // CTCACC
    97.14872,  2.10126, 1.34811, 1.00890, // CTCACG
    97.18569,  2.10126, 1.57068, 1.26879, // CTCACT
    102.00417, 2.40061, 1.72813, 1.49029, // CTCAGA
    100.18904, 2.40061, 1.73447, 1.49849, // CTCAGC
    100.88282, 2.40061, 1.68337, 1.43276, // CTCAGG
    100.61515, 2.40061, 1.87408, 1.68301, // CTCAGT
    96.46494,  2.47361, 1.62795, 1.33966, // CTCATA
    94.82559,  2.47361, 1.91621, 1.71079, // CTCATC
    95.56206,  2.47361, 1.67176, 1.39410, // CTCATG
    95.66259,  2.47361, 1.98540, 1.80429, // CTCATT
    103.09791, 1.63253, 1.15379, 0.90672, // CTCCAA
    100.38223, 1.63253, 1.30646, 1.09251, // CTCCAC
    101.86579, 1.63253, 1.17786, 0.93524, // CTCCAG
    101.43084, 1.63253, 1.39065, 1.19981, // CTCCAT
    99.47862,  1.60868, 1.22897, 1.00241, // CTCCCA
    98.34879,  1.60868, 1.21807, 0.98911, // CTCCCC
    99.88779,  1.60868, 1.14270, 0.89875, // CTCCCG
    98.87650,  1.60868, 1.24571, 1.02297, // CTCCCT
    101.69224, 1.82797, 1.29773, 1.02286, // CTCCGA
    99.46408,  1.82797, 1.34790, 1.08274, // CTCCGC
    100.69082, 1.82797, 1.28809, 1.01148, // CTCCGG
    100.21928, 1.82797, 1.38855, 1.13209, // CTCCGT
    99.00657,  1.91501, 1.47403, 1.27144, // CTCCTA
    97.16174,  1.91501, 1.54782, 1.36810, // CTCCTC
    98.07807,  1.91501, 1.42031, 1.20257, // CTCCTG
    97.68373,  1.91501, 1.60841, 1.44921, // CTCCTT
    100.65687, 2.59137, 1.68887, 1.45197, // CTCGAA
    96.71906,  2.59137, 2.00884, 1.88356, // CTCGAC
    99.57282,  2.59137, 1.82801, 1.63505, // CTCGAG
    98.11287,  2.59137, 2.11099, 2.02905, // CTCGAT
    95.83203,  2.48599, 1.60792, 1.35991, // CTCGCA
    94.55094,  2.48599, 1.68849, 1.46339, // CTCGCC
    95.66071,  2.48599, 1.65291, 1.41737, // CTCGCG
    95.51672,  2.48599, 1.75626, 1.55238, // CTCGCT
    100.20112, 2.51279, 1.95622, 1.77884, // CTCGGA
    97.40229,  2.51279, 1.82780, 1.60658, // CTCGGC
    99.14961,  2.51279, 1.79267, 1.56050, // CTCGGG
    98.85401,  2.51279, 1.97876, 1.80968, // CTCGGT
    94.24153,  5.74004, 1.99271, 1.86492, // CTCGTA
    90.99083,  5.74004, 1.79043, 1.58828, // CTCGTC
    91.68634,  5.74004, 1.73107, 1.50995, // CTCGTG
    93.71592,  5.74004, 2.40862, 2.47825, // CTCGTT
    98.76451,  1.86097, 1.64516, 1.30902, // CTCTAA
    95.73300,  1.86097, 1.66056, 1.32744, // CTCTAC
    97.20427,  1.86097, 1.66889, 1.33744, // CTCTAG
    96.84145,  1.86097, 1.63015, 1.29114, // CTCTAT
    93.60275,  2.32915, 2.08763, 2.05922, // CTCTCA
    92.08123,  2.32915, 2.04538, 1.99702, // CTCTCC
    93.54484,  2.32915, 2.03866, 1.98720, // CTCTCG
    93.14379,  2.32915, 2.05564, 2.01207, // CTCTCT
    96.46740,  2.03287, 1.59698, 1.32554, // CTCTGA
    93.61345,  2.03287, 1.77429, 1.55232, // CTCTGC
    95.23281,  2.03287, 1.71975, 1.48130, // CTCTGG
    94.39212,  2.03287, 1.67318, 1.42153, // CTCTGT
    89.35339,  2.37489, 2.09401, 2.00030, // CTCTTA
    86.10896,  2.37489, 2.05041, 1.93815, // CTCTTC
    88.48011,  2.37489, 1.95206, 1.80038, // CTCTTG
    87.30630,  2.37489, 1.86753, 1.68471, // CTCTTT
    86.26018,  3.07534, 1.84792, 1.42014, // CTGAAA
    83.26909,  3.07534, 2.19778, 1.84197, // CTGAAC
    85.06820,  3.07534, 1.86613, 1.44118, // CTGAAG
    84.19621,  3.07534, 2.28153, 1.94825, // CTGAAT
    76.53880,  2.53545, 1.86892, 1.62172, // CTGACA
    75.60886,  2.53545, 1.85017, 1.59738, // CTGACC
    76.41002,  2.53545, 1.88116, 1.63768, // CTGACG
    76.42019,  2.53545, 1.71029, 1.41970, // CTGACT
    85.48499,  3.17429, 1.82911, 1.47780, // CTGAGA
    83.63721,  3.17429, 2.00794, 1.69974, // CTGAGC
    84.47508,  3.17429, 1.90830, 1.57480, // CTGAGG
    83.80269,  3.17429, 2.13983, 1.86993, // CTGAGT
    75.98277,  2.69379, 1.83061, 1.59348, // CTGATA
    74.84260,  2.69379, 2.31263, 2.26262, // CTGATC
    75.43744,  2.69379, 2.04514, 1.88164, // CTGATG
    75.40919,  2.69379, 2.23904, 2.15548, // CTGATT
    92.53569,  1.48208, 1.30412, 1.04470, // CTGCAA
    90.25391,  1.48208, 1.20387, 0.92658, // CTGCAC
    91.43292,  1.48208, 1.27280, 1.00729, // CTGCAG
    91.15977,  1.48208, 1.20811, 0.93148, // CTGCAT
    89.34187,  1.66578, 1.51101, 1.24810, // CTGCCA
    88.05671,  1.66578, 1.53058, 1.27243, // CTGCCC
    89.08256,  1.66578, 1.46013, 1.18560, // CTGCCG
    88.82595,  1.66578, 1.42271, 1.14031, // CTGCCT
    91.31416,  1.67467, 1.27230, 0.91702, // CTGCGA
    89.02297,  1.67467, 1.32841, 0.97835, // CTGCGC
    90.21052,  1.67467, 1.34358, 0.99515, // CTGCGG
    89.48765,  1.67467, 1.34656, 0.99846, // CTGCGT
    87.98689,  1.83904, 1.45373, 1.21947, // CTGCTA
    86.36876,  1.83904, 1.39455, 1.14577, // CTGCTC
    87.55145,  1.83904, 1.42748, 1.18660, // CTGCTG
    87.28739,  1.83904, 1.44225, 1.20505, // CTGCTT
    88.12282,  3.22959, 1.87855, 1.61963, // CTGGAA
    83.88883,  3.22959, 2.37693, 2.30517, // CTGGAC
    87.21368,  3.22959, 1.96124, 1.72773, // CTGGAG
    85.26362,  3.22959, 2.27902, 2.16423, // CTGGAT
    83.21175,  2.83028, 1.64206, 1.35663, // CTGGCA
    82.19038,  2.83028, 1.84797, 1.61964, // CTGGCC
    83.14869,  2.83028, 1.69879, 1.42753, // CTGGCG
    82.97039,  2.83028, 1.80806, 1.56746, // CTGGCT
    88.50582,  2.98826, 1.78113, 1.43793, // CTGGGA
    86.35001,  2.98826, 1.95858, 1.65808, // CTGGGC
    87.48003,  2.98826, 1.85944, 1.53380, // CTGGGG
    87.00809,  2.98826, 2.01326, 1.72801, // CTGGGT
    82.45380,  3.42265, 1.96257, 1.60781, // CTGGTA
    80.33838,  3.42265, 2.25711, 1.98301, // CTGGTC
    81.23353,  3.42265, 2.08555, 1.76128, // CTGGTG
    81.43126,  3.42265, 2.36074, 2.12114, // CTGGTT
    86.58095,  2.22980, 1.88084, 1.59472, // CTGTAA
    83.23369,  2.22980, 1.95649, 1.69189, // CTGTAC
    85.35781,  2.22980, 1.89075, 1.60734, // CTGTAG
    84.48607,  2.22980, 1.90788, 1.62923, // CTGTAT
    81.53175,  2.83133, 2.44295, 2.37932, // CTGTCA
    81.01804,  2.83133, 2.64736, 2.68411, // CTGTCC
    81.46674,  2.83133, 2.42760, 2.35693, // CTGTCG
    81.68073,  2.83133, 2.53205, 2.51068, // CTGTCT
    84.16144,  2.65515, 2.07424, 1.84937, // CTGTGA
    80.70274,  2.65515, 2.22481, 2.05435, // CTGTGC
    82.40705,  2.65515, 1.98866, 1.73611, // CTGTGG
    81.65345,  2.65515, 2.22830, 2.05918, // CTGTGT
    77.58057,  3.04048, 2.44390, 2.39524, // CTGTTA
    74.22467,  3.04048, 2.66047, 2.72057, // CTGTTC
    76.48339,  3.04048, 2.54599, 2.54687, // CTGTTG
    75.56915,  3.04048, 2.47889, 2.44686, // CTGTTT
    105.16833, 2.16371, 1.37828, 1.04195, // CTTAAA
    102.95003, 2.16371, 1.53473, 1.22431, // CTTAAC
    104.24486, 2.16371, 1.34146, 1.00047, // CTTAAG
    104.42122, 2.16371, 1.63037, 1.34051, // CTTAAT
    100.52270, 2.01953, 1.15417, 0.77180, // CTTACA
    99.54297,  2.01953, 1.22020, 0.83896, // CTTACC
    99.94436,  2.01953, 1.17932, 0.79716, // CTTACG
    101.14171, 2.01953, 1.31589, 0.93957, // CTTACT
    102.70838, 2.04194, 1.45109, 1.13513, // CTTAGA
    101.54715, 2.04194, 1.36894, 1.04011, // CTTAGC
    102.15628, 2.04194, 1.31907, 0.98380, // CTTAGG
    103.16533, 2.04194, 1.50803, 1.20260, // CTTAGT
    100.19883, 2.50557, 1.45480, 1.09864, // CTTATA
    100.84588, 2.50557, 1.62906, 1.30184, // CTTATC
    100.11145, 2.50557, 1.50116, 1.15158, // CTTATG
    102.19914, 2.50557, 1.73776, 1.43429, // CTTATT
    108.82288, 2.19857, 1.44137, 1.13284, // CTTCAA
    107.10557, 2.19857, 1.45677, 1.15104, // CTTCAC
    107.93431, 2.19857, 1.47598, 1.17389, // CTTCAG
    107.86059, 2.19857, 1.54439, 1.25644, // CTTCAT
    106.76766, 2.17160, 1.39519, 1.06110, // CTTCCA
    105.28866, 2.17160, 1.37963, 1.04340, // CTTCCC
    106.65699, 2.17160, 1.46087, 1.13691, // CTTCCG
    106.52162, 2.17160, 1.45374, 1.12859, // CTTCCT
    108.09875, 2.22607, 1.54907, 1.26237, // CTTCGA
    105.74055, 2.22607, 1.39507, 1.07889, // CTTCGC
    107.31774, 2.22607, 1.50934, 1.21412, // CTTCGG
    107.24615, 2.22607, 1.61332, 1.34172, // CTTCGT
    107.74388, 2.26458, 1.48754, 1.19288, // CTTCTA
    106.45938, 2.26458, 1.59105, 1.31953, // CTTCTC
    107.20884, 2.26458, 1.51672, 1.22816, // CTTCTG
    107.21221, 2.26458, 1.75799, 1.53256, // CTTCTT
    101.21652, 2.11369, 1.41775, 1.18353, // CTTGAA
    98.27224,  2.11369, 1.61164, 1.43445, // CTTGAC
    100.34789, 2.11369, 1.38457, 1.14224, // CTTGAG
    99.74285,  2.11369, 1.66867, 1.51126, // CTTGAT
    98.83545,  2.06630, 1.22297, 0.87794, // CTTGCA
    97.96221,  2.06630, 1.24786, 0.90488, // CTTGCC
    98.54362,  2.06630, 1.16354, 0.81473, // CTTGCG
    99.45121,  2.06630, 1.27748, 0.93729, // CTTGCT
    100.67903, 1.98654, 1.40924, 1.17026, // CTTGGA
    99.21224,  1.98654, 1.42700, 1.19245, // CTTGGC
    99.91388,  1.98654, 1.34201, 1.08752, // CTTGGG
    100.76840, 1.98654, 1.61229, 1.43209, // CTTGGT
    99.33635,  2.51049, 1.52199, 1.19371, // CTTGTA
    99.18424,  2.51049, 1.64659, 1.34325, // CTTGTC
    98.92414,  2.51049, 1.57069, 1.25146, // CTTGTG
    101.34917, 2.51049, 1.75455, 1.47750, // CTTGTT
    99.38382,  1.70840, 1.20443, 0.88517, // CTTTAA
    96.12679,  1.70840, 1.09604, 0.76841, // CTTTAC
    98.20597,  1.70840, 1.18937, 0.86861, // CTTTAG
    97.61660,  1.70840, 1.15108, 0.82701, // CTTTAT
    94.47535,  1.56977, 1.30421, 1.03774, // CTTTCA
    92.83063,  1.56977, 1.31825, 1.05454, // CTTTCC
    94.37139,  1.56977, 1.26016, 0.98561, // CTTTCG
    94.30088,  1.56977, 1.28813, 1.01861, // CTTTCT
    97.71116,  1.62351, 1.11118, 0.79311, // CTTTGA
    94.25998,  1.62351, 1.08676, 0.76710, // CTTTGC
    96.71993,  1.62351, 1.11237, 0.79438, // CTTTGG
    95.46304,  1.62351, 1.13491, 0.81864, // CTTTGT
    91.42952,  1.51391, 1.09707, 0.77694, // CTTTTA
    89.63750,  1.51391, 1.08868, 0.76803, // CTTTTC
    91.05377,  1.51391, 1.06903, 0.74734, // CTTTTG
    91.46416,  1.51391, 0.98342, 0.65938, // CTTTTT
    84.83351,  1.51785, 1.01351, 0.68061, // GAAAAA
    82.24157,  1.51785, 1.03761, 0.70503, // GAAAAC
    83.67791,  1.51785, 1.01968, 0.68683, // GAAAAG
    82.79356,  1.51785, 1.10196, 0.77162, // GAAAAT
    74.70693,  1.70501, 1.26783, 0.90495, // GAAACA
    73.43240,  1.70501, 1.22891, 0.86361, // GAAACC
    74.82775,  1.70501, 1.21877, 0.85294, // GAAACG
    74.58650,  1.70501, 1.14717, 0.77889, // GAAACT
    82.40766,  1.53757, 0.99032, 0.66144, // GAAAGA
    80.79589,  1.53757, 0.96005, 0.63135, // GAAAGC
    81.59670,  1.53757, 0.96151, 0.63279, // GAAAGG
    80.75521,  1.53757, 1.04220, 0.71410, // GAAAGT
    74.86594,  1.54253, 1.08847, 0.78694, // GAAATA
    73.11850,  1.54253, 0.96506, 0.65697, // GAAATC
    74.13222,  1.54253, 1.02942, 0.72378, // GAAATG
    74.05160,  1.54253, 1.10352, 0.80332, // GAAATT
    87.90414,  1.96203, 1.73391, 1.70306, // GAACAA
    86.21222,  1.96203, 1.59118, 1.49717, // GAACAC
    87.00499,  1.96203, 1.68617, 1.63322, // GAACAG
    87.07323,  1.96203, 1.56815, 1.46479, // GAACAT
    83.94132,  1.83613, 1.71004, 1.74657, // GAACCA
    82.83934,  1.83613, 1.58180, 1.55384, // GAACCC
    83.82087,  1.83613, 1.62743, 1.62155, // GAACCG
    83.62945,  1.83613, 1.63622, 1.63472, // GAACCT
    86.52070,  1.55453, 1.52300, 1.46242, // GAACGA
    84.90051,  1.55453, 1.34564, 1.21455, // GAACGC
    85.82282,  1.55453, 1.48361, 1.40605, // GAACGG
    85.41703,  1.55453, 1.43757, 1.34112, // GAACGT
    83.64428,  1.58806, 1.44419, 1.34776, // GAACTA
    81.70897,  1.58806, 1.30134, 1.15284, // GAACTC
    82.87008,  1.58806, 1.42060, 1.31489, // GAACTG
    82.31319,  1.58806, 1.15357, 0.96216, // GAACTT
    83.00746,  1.46160, 1.09674, 0.79534, // GAAGAA
    79.25123,  1.46160, 1.12650, 0.82793, // GAAGAC
    82.08597,  1.46160, 1.02257, 0.71603, // GAAGAG
    80.32113,  1.46160, 1.15901, 0.86402, // GAAGAT
    76.94267,  1.42710, 1.08455, 0.78802, // GAAGCA
    75.50147,  1.42710, 1.02165, 0.72047, // GAAGCC
    76.87143,  1.42710, 1.02705, 0.72619, // GAAGCG
    76.76763,  1.42710, 0.95294, 0.64902, // GAAGCT
    82.14809,  1.40989, 1.09922, 0.79895, // GAAGGA
    79.57441,  1.40989, 1.03804, 0.73319, // GAAGGC
    81.22374,  1.40989, 0.99847, 0.69167, // GAAGGG
    80.38671,  1.40989, 1.10642, 0.80681, // GAAGGT
    75.45234,  1.48703, 0.95657, 0.67035, // GAAGTA
    73.89876,  1.48703, 1.07690, 0.80074, // GAAGTC
    75.04359,  1.48703, 1.01930, 0.73736, // GAAGTG
    74.78915,  1.48703, 1.15248, 0.88649, // GAAGTT
    81.13429,  1.98024, 1.80273, 1.83518, // GAATAA
    78.84102,  1.98024, 1.72914, 1.72397, // GAATAC
    80.36050,  1.98024, 1.67996, 1.65094, // GAATAG
    80.15035,  1.98024, 1.90614, 1.99534, // GAATAT
    73.97533,  2.22770, 2.12293, 2.65938, // GAATCA
    73.16045,  2.22770, 2.25524, 2.91183, // GAATCC
    74.12232,  2.22770, 2.08062, 2.58028, // GAATCG
    74.15972,  2.22770, 2.25431, 2.91003, // GAATCT
    78.42674,  1.67617, 1.71565, 1.75509, // GAATGA
    75.82070,  1.67617, 1.74494, 1.80022, // GAATGC
    77.39373,  1.67617, 1.69885, 1.72938, // GAATGG
    76.73931,  1.67617, 1.85741, 1.97705, // GAATGT
    69.60846,  2.00867, 1.87329, 1.86690, // GAATTA
    66.95419,  2.00867, 1.77602, 1.72340, // GAATTC
    69.07015,  2.00867, 1.79342, 1.74879, // GAATTG
    68.51136,  2.00867, 1.65812, 1.55468, // GAATTT
    106.82549, 2.19132, 1.88070, 1.64916, // GACAAA
    103.52554, 2.19132, 1.94872, 1.73943, // GACAAC
    105.66968, 2.19132, 1.89339, 1.66588, // GACAAG
    104.87078, 2.19132, 2.05847, 1.88842, // GACAAT
    98.00244,  1.74166, 1.30842, 0.94051, // GACACA
    96.73920,  1.74166, 1.33002, 0.96389, // GACACC
    97.96068,  1.74166, 1.35904, 0.99561, // GACACG
    97.94517,  1.74166, 1.47185, 1.12210, // GACACT
    103.62596, 2.28987, 1.83777, 1.57768, // GACAGA
    100.80088, 2.28987, 1.79721, 1.52574, // GACAGC
    102.59385, 2.28987, 1.77165, 1.49331, // GACAGG
    102.05144, 2.28987, 1.99390, 1.78294, // GACAGT
    97.41274,  2.17760, 1.49582, 1.21547, // GACATA
    95.91196,  2.17760, 1.66658, 1.42944, // GACATC
    96.73982,  2.17760, 1.56814, 1.30467, // GACATG
    97.26743,  2.17760, 1.84903, 1.67048, // GACATT
    106.81534, 1.88264, 1.57972, 1.41777, // GACCAA
    103.74683, 1.88264, 1.55773, 1.38827, // GACCAC
    106.05382, 1.88264, 1.56461, 1.39747, // GACCAG
    105.21138, 1.88264, 1.69401, 1.57438, // GACCAT
    102.66255, 1.75332, 1.42284, 1.22275, // GACCCA
    101.31167, 1.75332, 1.26974, 1.03080, // GACCCC
    102.68555, 1.75332, 1.39512, 1.18719, // GACCCG
    102.07374, 1.75332, 1.51096, 1.33808, // GACCCT
    105.35864, 1.90443, 1.67532, 1.48996, // GACCGA
    102.50432, 1.90443, 1.58862, 1.37582, // GACCGC
    104.32265, 1.90443, 1.58945, 1.37690, // GACCGG
    103.63503, 1.90443, 1.69765, 1.51985, // GACCGT
    101.84760, 1.83098, 1.42937, 1.11690, // GACCTA
    99.64319,  1.83098, 1.65401, 1.39028, // GACCTC
    101.16424, 1.83098, 1.48513, 1.18289, // GACCTG
    100.34227, 1.83098, 1.76028, 1.52640, // GACCTT
    102.12872, 2.24722, 1.86305, 1.65239, // GACGAA
    97.34564,  2.24722, 1.86906, 1.66040, // GACGAC
    100.96513, 2.24722, 1.89046, 1.68900, // GACGAG
    99.56364,  2.24722, 1.96942, 1.79592, // GACGAT
    97.11296,  2.02317, 1.49778, 1.18698, // GACGCA
    95.66850,  2.02317, 1.55308, 1.25332, // GACGCC
    97.23326,  2.02317, 1.54381, 1.24212, // GACGCG
    97.07303,  2.02317, 1.68464, 1.41590, // GACGCT
    101.63598, 2.22668, 1.92494, 1.72637, // GACGGA
    98.53540,  2.22668, 1.76730, 1.51870, // GACGGC
    100.73732, 2.22668, 1.83402, 1.60551, // GACGGG
    100.23248, 2.22668, 1.96107, 1.77521, // GACGGT
    96.60034,  2.64556, 1.83804, 1.55670, // GACGTA
    94.53590,  2.64556, 1.96479, 1.72046, // GACGTC
    95.83574,  2.64556, 1.87368, 1.60220, // GACGTG
    96.42843,  2.64556, 2.11707, 1.92432, // GACGTT
    100.67500, 1.69847, 1.70739, 1.31782, // GACTAA
    96.95079,  1.69847, 1.50070, 1.08593, // GACTAC
    99.38701,  1.69847, 1.43735, 1.01789, // GACTAG
    98.40117,  1.69847, 1.60193, 1.19763, // GACTAT
    92.33290,  1.86136, 1.58318, 1.42867, // GACTCA
    90.54727,  1.86136, 1.78560, 1.71125, // GACTCC
    92.38296,  1.86136, 1.59909, 1.45027, // GACTCG
    92.00403,  1.86136, 1.59803, 1.44882, // GACTCT
    97.53059,  1.64486, 1.47226, 1.16317, // GACTGA
    94.00903,  1.64486, 1.45054, 1.13753, // GACTGC
    96.28244,  1.64486, 1.42514, 1.10779, // GACTGG
    95.20364,  1.64486, 1.49573, 1.19110, // GACTGT
    87.53082,  1.46961, 1.29250, 0.97044, // GACTTA
    84.64753,  1.46961, 1.24025, 0.91219, // GACTTC
    87.05997,  1.46961, 1.27069, 0.94597, // GACTTG
    86.00065,  1.46961, 1.11380, 0.77631, // GACTTT
    77.16147,  1.82916, 1.09930, 0.69035, // GAGAAA
    74.38516,  1.82916, 1.26717, 0.85438, // GAGAAC
    75.87596,  1.82916, 1.12952, 0.71901, // GAGAAG
    74.75730,  1.82916, 1.26713, 0.85434, // GAGAAT
    65.53258,  2.14481, 1.67459, 1.26888, // GAGACA
    64.87668,  2.14481, 1.72655, 1.32839, // GAGACC
    65.36853,  2.14481, 1.64119, 1.23110, // GAGACG
    65.45330,  2.14481, 1.65865, 1.25080, // GAGACT
    75.01429,  1.82854, 1.13452, 0.78445, // GAGAGA
    73.70988,  1.82854, 1.06286, 0.71131, // GAGAGC
    73.60582,  1.82854, 1.15430, 0.80506, // GAGAGG
    73.67346,  1.82854, 1.12243, 0.77195, // GAGAGT
    65.88961,  1.85459, 1.58796, 1.35373, // GAGATA
    64.38126,  1.85459, 1.37495, 1.09069, // GAGATC
    65.19205,  1.85459, 1.46775, 1.20296, // GAGATG
    65.17221,  1.85459, 1.19741, 0.88641, // GAGATT
    85.93084,  1.87706, 1.60969, 1.39426, // GAGCAA
    83.54329,  1.87706, 1.35855, 1.08104, // GAGCAC
    84.88293,  1.87706, 1.50350, 1.25858, // GAGCAG
    84.36567,  1.87706, 1.41629, 1.15068, // GAGCAT
    81.36761,  1.70618, 1.63321, 1.52414, // GAGCCA
    80.07623,  1.70618, 1.67238, 1.57930, // GAGCCC
    81.29425,  1.70618, 1.54369, 1.40056, // GAGCCG
    81.04504,  1.70618, 1.47601, 1.30946, // GAGCCT
    83.61566,  1.56014, 1.46044, 1.27246, // GAGCGA
    80.80411,  1.56014, 1.35693, 1.13960, // GAGCGC
    82.52107,  1.56014, 1.46077, 1.27289, // GAGCGG
    81.46428,  1.56014, 1.46260, 1.27527, // GAGCGT
    80.23185,  1.42173, 1.42918, 1.20129, // GAGCTA
    77.92246,  1.42173, 1.30564, 1.04895, // GAGCTC
    79.57163,  1.42173, 1.34078, 1.09158, // GAGCTG
    78.95648,  1.42173, 1.13530, 0.85052, // GAGCTT
    77.04816,  1.83346, 1.12313, 0.73252, // GAGGAA
    72.34175,  1.83346, 1.40406, 1.02388, // GAGGAC
    75.95770,  1.83346, 1.15661, 0.76552, // GAGGAG
    73.72581,  1.83346, 1.41847, 1.03968, // GAGGAT
    70.90958,  1.67690, 1.34162, 1.04310, // GAGGCA
    69.69790,  1.67690, 1.45154, 1.17388, // GAGGCC
    70.86570,  1.67690, 1.33481, 1.03515, // GAGGCG
    70.46071,  1.67690, 1.26093, 0.95042, // GAGGCT
    77.15888,  1.68720, 1.04282, 0.71184, // GAGGGA
    74.79310,  1.68720, 1.15803, 0.83300, // GAGGGC
    76.19856,  1.68720, 0.96247, 0.63117, // GAGGGG
    75.55031,  1.68720, 1.12521, 0.79784, // GAGGGT
    70.21683,  1.76089, 1.08710, 0.74148, // GAGGTA
    68.49592,  1.76089, 1.35193, 1.02832, // GAGGTC
    69.45042,  1.76089, 1.19910, 0.85897, // GAGGTG
    69.59032,  1.76089, 1.30530, 0.97559, // GAGGTT
    75.55785,  2.04805, 1.91474, 1.88220, // GAGTAA
    72.41631,  2.04805, 1.72680, 1.61199, // GAGTAC
    74.13450,  2.04805, 1.68651, 1.55590, // GAGTAG
    73.56683,  2.04805, 1.85321, 1.79220, // GAGTAT
    67.44377,  2.42133, 1.90328, 1.92654, // GAGTCA
    66.16784,  2.42133, 2.03034, 2.12264, // GAGTCC
    67.40195,  2.42133, 1.88764, 1.90284, // GAGTCG
    67.60882,  2.42133, 2.02940, 2.12117, // GAGTCT
    71.35480,  1.99693, 1.63389, 1.58195, // GAGTGA
    67.93321,  1.99693, 1.86340, 1.92671, // GAGTGC
    70.29641,  1.99693, 1.68937, 1.66321, // GAGTGG
    69.14875,  1.99693, 1.81087, 1.84582, // GAGTGT
    62.44920,  2.35636, 1.98804, 1.85471, // GAGTTA
    59.83591,  2.35636, 2.12447, 2.04887, // GAGTTC
    62.06153,  2.35636, 1.98230, 1.84668, // GAGTTG
    61.09120,  2.35636, 1.97088, 1.83075, // GAGTTT
    110.29144, 3.16286, 2.51512, 2.11444, // GATAAA
    107.75428, 3.16286, 2.62327, 2.25228, // GATAAC
    109.12917, 3.16286, 2.49680, 2.09137, // GATAAG
    109.95282, 3.16286, 2.75051, 2.41811, // GATAAT
    104.94406, 3.09766, 2.24738, 1.77844, // GATACA
    104.56754, 3.09766, 2.26325, 1.79730, // GATACC
    104.59646, 3.09766, 2.20320, 1.72625, // GATACG
    106.10046, 3.09766, 2.49050, 2.07469, // GATACT
    107.82439, 3.02439, 2.43924, 2.17603, // GATAGA
    106.23379, 3.02439, 2.44904, 2.18917, // GATAGC
    107.11508, 3.02439, 2.47949, 2.23013, // GATAGG
    108.28670, 3.02439, 2.64292, 2.45421, // GATAGT
    106.04032, 3.75132, 2.60275, 2.14551, // GATATA
    106.56336, 3.75132, 2.90007, 2.52345, // GATATC
    105.81466, 3.75132, 2.57638, 2.11298, // GATATG
    109.20086, 3.75132, 2.99616, 2.64990, // GATATT
    117.94606, 2.97261, 2.59314, 2.76259, // GATCAA
    115.45112, 2.97261, 2.50046, 2.61581, // GATCAC
    117.32716, 2.97261, 2.45902, 2.55105, // GATCAG
    116.50731, 2.97261, 2.69600, 2.92858, // GATCAT
    116.84959, 3.04768, 2.61672, 2.67007, // GATCCA
    114.68282, 3.04768, 2.68803, 2.77996, // GATCCC
    116.78833, 3.04768, 2.63678, 2.70083, // GATCCG
    116.26587, 3.04768, 2.75139, 2.87883, // GATCCT
    117.70063, 3.00027, 2.68980, 2.79944, // GATCGA
    114.83337, 3.00027, 2.59841, 2.65797, // GATCGC
    116.87186, 3.00027, 2.69259, 2.80379, // GATCGG
    116.84079, 3.00027, 2.84283, 3.04169, // GATCGT
    116.28159, 3.10410, 2.75925, 2.80890, // GATCTA
    115.39571, 3.10410, 2.84031, 2.93359, // GATCTC
    116.37083, 3.10410, 2.70420, 2.72526, // GATCTG
    116.63258, 3.10410, 3.00893, 3.19866, // GATCTT
    106.92771, 2.99484, 2.47760, 2.25414, // GATGAA
    103.30190, 2.99484, 2.50704, 2.29443, // GATGAC
    105.98437, 2.99484, 2.40545, 2.15640, // GATGAG
    105.86841, 2.99484, 2.73938, 2.62067, // GATGAT
    105.66971, 3.13415, 2.27132, 1.90465, // GATGCA
    105.21145, 3.13415, 2.41012, 2.08188, // GATGCC
    105.78298, 3.13415, 2.36101, 2.01857, // GATGCG
    107.04909, 3.13415, 2.61428, 2.35194, // GATGCT
    107.70086, 2.93123, 2.48554, 2.21104, // GATGGA
    105.35589, 2.93123, 2.44836, 2.16163, // GATGGC
    106.86672, 2.93123, 2.39242, 2.08797, // GATGGG
    107.70268, 2.93123, 2.52413, 2.26275, // GATGGT
    107.62158, 3.56800, 2.71104, 2.35245, // GATGTA
    108.86927, 3.56800, 2.79783, 2.46632, // GATGTC
    107.70553, 3.56800, 2.82056, 2.49644, // GATGTG
    110.89342, 3.56800, 3.12099, 2.90573, // GATGTT
    106.24377, 2.18027, 1.94588, 1.95846, // GATTAA
    102.27310, 2.18027, 1.80130, 1.74431, // GATTAC
    104.92267, 2.18027, 2.06077, 2.13446, // GATTAG
    103.78559, 2.18027, 1.87847, 1.85758, // GATTAT
    100.14968, 2.09022, 2.02363, 1.95550, // GATTCA
    98.29873,  2.09022, 2.08883, 2.05076, // GATTCC
    100.13099, 2.09022, 2.02862, 1.96274, // GATTCG
    100.02409, 2.09022, 1.97428, 1.88441, // GATTCT
    103.90612, 2.20088, 1.91878, 1.87597, // GATTGA
    99.85786,  2.20088, 1.82681, 1.74272, // GATTGC
    102.86287, 2.20088, 1.91880, 1.87599, // GATTGG
    101.63967, 2.20088, 1.93684, 1.90252, // GATTGT
    96.30701,  1.92511, 1.71371, 1.52798, // GATTTA
    93.91281,  1.92511, 1.67682, 1.47891, // GATTTC
    95.62872,  1.92511, 1.67328, 1.47423, // GATTTG
    95.92146,  1.92511, 1.71976, 1.53608, // GATTTT
    91.52287,  2.70987, 1.50203, 1.09083, // GCAAAA
    88.27341,  2.70987, 1.73014, 1.34854, // GCAAAC
    90.82009,  2.70987, 1.56481, 1.15994, // GCAAAG
    89.14020,  2.70987, 1.81654, 1.45081, // GCAAAT
    82.19198,  2.16112, 1.57555, 1.28720, // GCAACA
    80.91370,  2.16112, 1.50214, 1.19830, // GCAACC
    82.05805,  2.16112, 1.51513, 1.21388, // GCAACG
    81.55514,  2.16112, 1.41497, 1.09552, // GCAACT
    89.87186,  2.81796, 1.54015, 1.17468, // GCAAGA
    87.14250,  2.81796, 1.67362, 1.33064, // GCAAGC
    88.82630,  2.81796, 1.49253, 1.12062, // GCAAGG
    87.63187,  2.81796, 1.77269, 1.45052, // GCAAGT
    81.00738,  2.24890, 1.51884, 1.23851, // GCAATA
    79.51646,  2.24890, 1.86386, 1.68365, // GCAATC
    80.41199,  2.24890, 1.57476, 1.30753, // GCAATG
    80.08215,  2.24890, 1.83912, 1.65024, // GCAATT
    94.01706,  1.22481, 1.04078, 0.78631, // GCACAA
    91.48792,  1.22481, 0.99328, 0.73310, // GCACAC
    93.14011,  1.22481, 1.00906, 0.75063, // GCACAG
    92.49433,  1.22481, 0.93071, 0.66493, // GCACAT
    90.57220,  1.29044, 1.10446, 0.86240, // GCACCA
    89.33187,  1.29044, 1.08633, 0.84125, // GCACCC
    90.47731,  1.29044, 1.09162, 0.84741, // GCACCG
    90.21233,  1.29044, 1.04377, 0.79231, // GCACCT
    92.87086,  1.39798, 0.97792, 0.70439, // GCACGA
    90.39981,  1.39798, 1.04648, 0.77975, // GCACGC
    91.93219,  1.39798, 1.02330, 0.75398, // GCACGG
    91.29054,  1.39798, 1.00428, 0.73306, // GCACGT
    88.96387,  1.56257, 1.08366, 0.77946, // GCACTA
    87.01270,  1.56257, 1.09485, 0.79156, // GCACTC
    88.35424,  1.56257, 1.09495, 0.79168, // GCACTG
    87.97573,  1.56257, 1.03844, 0.73118, // GCACTT
    87.12311,  2.60118, 1.60460, 1.21668, // GCAGAA
    81.71243,  2.60118, 1.85964, 1.51799, // GCAGAC
    86.03571,  2.60118, 1.65622, 1.27586, // GCAGAG
    83.57467,  2.60118, 1.91465, 1.58585, // GCAGAT
    82.00798,  2.08236, 1.33620, 1.08321, // GCAGCA
    80.51316,  2.08236, 1.38837, 1.14727, // GCAGCC
    81.78686,  2.08236, 1.34373, 1.09239, // GCAGCG
    81.78746,  2.08236, 1.45456, 1.23028, // GCAGCT
    86.65682,  2.50147, 1.53041, 1.19678, // GCAGGA
    82.99176,  2.50147, 1.67807, 1.37410, // GCAGGC
    85.52396,  2.50147, 1.54565, 1.21470, // GCAGGG
    84.42905,  2.50147, 1.74494, 1.45705, // GCAGGT
    79.31435,  2.40625, 1.67038, 1.41017, // GCAGTA
    76.09673,  2.40625, 1.95056, 1.77946, // GCAGTC
    78.15703,  2.40625, 1.70866, 1.45893, // GCAGTG
    77.41588,  2.40625, 1.93599, 1.75956, // GCAGTT
    87.93751,  1.95200, 1.50474, 1.17012, // GCATAA
    84.69618,  1.95200, 1.59070, 1.27181, // GCATAC
    87.15618,  1.95200, 1.55442, 1.22855, // GCATAG
    86.16746,  1.95200, 1.62991, 1.31912, // GCATAT
    81.17729,  2.22932, 1.98717, 1.93499, // GCATCA
    79.83222,  2.22932, 2.13952, 2.16173, // GCATCC
    81.17351,  2.22932, 1.98700, 1.93474, // GCATCG
    80.94912,  2.22932, 2.04634, 2.02205, // GCATCT
    85.32210,  2.18506, 1.52590, 1.19292, // GCATGA
    81.86078,  2.18506, 1.82753, 1.56357, // GCATGC
    84.19991,  2.18506, 1.64245, 1.33217, // GCATGG
    82.99030,  2.18506, 1.75378, 1.46989, // GCATGT
    75.59882,  2.43089, 2.10834, 1.97464, // GCATTA
    72.57817,  2.43089, 2.10695, 1.97267, // GCATTC
    74.83872,  2.43089, 2.09655, 1.95808, // GCATTG
    74.01700,  2.43089, 1.95712, 1.76605, // GCATTT
    103.17460, 1.63085, 1.22025, 1.00056, // GCCAAA
    100.61496, 1.63085, 1.41065, 1.24365, // GCCAAC
    102.15368, 1.63085, 1.21637, 0.99579, // GCCAAG
    101.40038, 1.63085, 1.51217, 1.38029, // GCCAAT
    95.31901,  1.46322, 1.03724, 0.73098, // GCCACA
    94.26997,  1.46322, 1.11264, 0.81211, // GCCACC
    95.31003,  1.46322, 1.08941, 0.78681, // GCCACG
    95.05689,  1.46322, 1.17856, 0.88534, // GCCACT
    100.88631, 1.84203, 1.35768, 1.10623, // GCCAGA
    98.62416,  1.84203, 1.44632, 1.21632, // GCCAGC
    99.75961,  1.84203, 1.37206, 1.12386, // GCCAGG
    99.35961,  1.84203, 1.58323, 1.39305, // GCCAGT
    94.27914,  1.79927, 1.26504, 0.91947, // GCCATA
    92.44036,  1.79927, 1.47115, 1.15310, // GCCATC
    93.62866,  1.79927, 1.39470, 1.06439, // GCCATG
    93.50286,  1.79927, 1.60995, 1.32007, // GCCATT
    102.43057, 1.44981, 0.90302, 0.64459, // GCCCAA
    100.10357, 1.44981, 0.92107, 0.66401, // GCCCAC
    101.70197, 1.44981, 0.87369, 0.61345, // GCCCAG
    101.10914, 1.44981, 1.01185, 0.76456, // GCCCAT
    98.90899,  1.40182, 0.90851, 0.66330, // GCCCCA
    97.18743,  1.40182, 0.88455, 0.63723, // GCCCCC
    98.56439,  1.40182, 0.90251, 0.65674, // GCCCCG
    98.18067,  1.40182, 0.84318, 0.59306, // GCCCCT
    101.15453, 1.43192, 0.93938, 0.69602, // GCCCGA
    98.80123,  1.43192, 0.94427, 0.70145, // GCCCGC
    100.08150, 1.43192, 0.90036, 0.65310, // GCCCGG
    99.60123,  1.43192, 1.05182, 0.82464, // GCCCGT
    97.49046,  1.44710, 0.93653, 0.69070, // GCCCTA
    95.74057,  1.44710, 1.10276, 0.88253, // GCCCTC
    97.04746,  1.44710, 0.98271, 0.74242, // GCCCTG
    96.36536,  1.44710, 1.17309, 0.96829, // GCCCTT
    98.25696,  1.82025, 1.32775, 1.07258, // GCCGAA
    93.56539,  1.82025, 1.56859, 1.37728, // GCCGAC
    96.94579,  1.82025, 1.34545, 1.09411, // GCCGAG
    95.28702,  1.82025, 1.56774, 1.37616, // GCCGAT
    93.63631,  1.58093, 1.10678, 0.77157, // GCCGCA
    92.28131,  1.58093, 1.18911, 0.85925, // GCCGCC
    93.56640,  1.58093, 1.15398, 0.82146, // GCCGCG
    93.32514,  1.58093, 1.23237, 0.90656, // GCCGCT
    97.86780,  1.74812, 1.36209, 1.15545, // GCCGGA
    94.73272,  1.74812, 1.46687, 1.29130, // GCCGGC
    96.72327,  1.74812, 1.29556, 1.07182, // GCCGGG
    96.11043,  1.74812, 1.49743, 1.33186, // GCCGGT
    91.95446,  2.13467, 1.47325, 1.23524, // GCCGTA
    89.41518,  2.13467, 1.61343, 1.41566, // GCCGTC
    91.09525,  2.13467, 1.52748, 1.30406, // GCCGTG
    91.02063,  2.13467, 1.80790, 1.67917, // GCCGTT
    97.36578,  1.77408, 1.28904, 1.00516, // GCCTAA
    94.54892,  1.77408, 1.30564, 1.02464, // GCCTAC
    96.47539,  1.77408, 1.20395, 0.90729, // GCCTAG
    95.78051,  1.77408, 1.38814, 1.12328, // GCCTAT
    90.87483,  2.16210, 1.69166, 1.52739, // GCCTCA
    89.23277,  2.16210, 1.78332, 1.65320, // GCCTCC
    90.71387,  2.16210, 1.69480, 1.53165, // GCCTCG
    90.65554,  2.16210, 1.80431, 1.68247, // GCCTCT
    94.62085,  1.63830, 1.20718, 0.91718, // GCCTGA
    91.45570,  1.63830, 1.39103, 1.13449, // GCCTGC
    93.48036,  1.63830, 1.30817, 1.03465, // GCCTGG
    92.59635,  1.63830, 1.40314, 1.14934, // GCCTGT
    86.41753,  1.91311, 1.71619, 1.48603, // GCCTTA
    83.07268,  1.91311, 1.68484, 1.44551, // GCCTTC
    85.38181,  1.91311, 1.64286, 1.39182, // GCCTTG
    84.35484,  1.91311, 1.52964, 1.25045, // GCCTTT
    84.86228,  2.36795, 1.55448, 1.15531, // GCGAAA
    81.48370,  2.36795, 1.83172, 1.47777, // GCGAAC
    83.60064,  2.36795, 1.63074, 1.24135, // GCGAAG
    82.24277,  2.36795, 1.74621, 1.37550, // GCGAAT
    74.43381,  2.00332, 1.77849, 1.50930, // GCGACA
    73.35608,  2.00332, 1.78081, 1.51226, // GCGACC
    74.25036,  2.00332, 1.73720, 1.45705, // GCGACG
    74.11810,  2.00332, 1.63085, 1.32532, // GCGACT
    83.21643,  2.41291, 1.49998, 1.09868, // GCGAGA
    80.98331,  2.41291, 1.66231, 1.28178, // GCGAGC
    82.50385,  2.41291, 1.56264, 1.16824, // GCGAGG
    81.48023,  2.41291, 1.71857, 1.34739, // GCGAGT
    73.59305,  1.85855, 1.52418, 1.36982, // GCGATA
    71.48757,  1.85855, 1.67717, 1.58116, // GCGATC
    72.64844,  1.85855, 1.50873, 1.34905, // GCGATG
    72.66030,  1.85855, 1.74946, 1.68448, // GCGATT
    90.21027,  1.44291, 1.20926, 0.93069, // GCGCAA
    87.64048,  1.44291, 1.14117, 0.85321, // GCGCAC
    89.21907,  1.44291, 1.19076, 0.90941, // GCGCAG
    88.58031,  1.44291, 1.11367, 0.82255, // GCGCAT
    86.51197,  1.52741, 1.35900, 1.11659, // GCGCCA
    85.20144,  1.52741, 1.34207, 1.09580, // GCGCCC
    86.39913,  1.52741, 1.35478, 1.11140, // GCGCCG
    86.17426,  1.52741, 1.31852, 1.06707, // GCGCCT
    88.46920,  1.58181, 1.23903, 0.93162, // GCGCGA
    85.37314,  1.58181, 1.28978, 0.98943, // GCGCGC
    87.29837,  1.58181, 1.30678, 1.00906, // GCGCGG
    86.00411,  1.58181, 1.31241, 1.01558, // GCGCGT
    84.66758,  1.56040, 1.39153, 1.09141, // GCGCTA
    82.26661,  1.56040, 1.33029, 1.02016, // GCGCTC
    83.71771,  1.56040, 1.35287, 1.04624, // GCGCTG
    83.18393,  1.56040, 1.16362, 0.83457, // GCGCTT
    84.29619,  2.44172, 1.69482, 1.29081, // GCGGAA
    78.85926,  2.44172, 1.98637, 1.63783, // GCGGAC
    82.95075,  2.44172, 1.72548, 1.32599, // GCGGAG
    80.52359,  2.44172, 1.98819, 1.64008, // GCGGAT
    79.25942,  2.03186, 1.49949, 1.21646, // GCGGCA
    77.73205,  2.03186, 1.43681, 1.14099, // GCGGCC
    79.05709,  2.03186, 1.46984, 1.18056, // GCGGCG
    78.83721,  2.03186, 1.46133, 1.17032, // GCGGCT
    84.89113,  2.35270, 1.57863, 1.15231, // GCGGGA
    81.98453,  2.35270, 1.75998, 1.35647, // GCGGGC
    83.71165,  2.35270, 1.59297, 1.16805, // GCGGGG
    82.92384,  2.35270, 1.79350, 1.39542, // GCGGGT
    77.65904,  2.46096, 1.78977, 1.49430, // GCGGTA
    75.15196,  2.46096, 1.95062, 1.70020, // GCGGTC
    76.52673,  2.46096, 1.79363, 1.49914, // GCGGTG
    76.02077,  2.46096, 2.04645, 1.82702, // GCGGTT
    81.47946,  2.14468, 1.78829, 1.54627, // GCGTAA
    77.98011,  2.14468, 1.94354, 1.75194, // GCGTAC
    80.64812,  2.14468, 1.78445, 1.54130, // GCGTAG
    79.70191,  2.14468, 1.98478, 1.80799, // GCGTAT
    74.49931,  2.49565, 2.44921, 2.80740, // GCGTCA
    73.10376,  2.49565, 2.41990, 2.75715, // GCGTCC
    74.66631,  2.49565, 2.44280, 2.79637, // GCGTCG
    74.19457,  2.49565, 2.38156, 2.69189, // GCGTCT
    78.04438,  2.22005, 1.99424, 1.90340, // GCGTGA
    74.04066,  2.22005, 2.18015, 2.17567, // GCGTGC
    76.67219,  2.22005, 2.03059, 1.95569, // GCGTGG
    75.31801,  2.22005, 2.16674, 2.15564, // GCGTGT
    69.09482,  2.47313, 2.42943, 2.55005, // GCGTTA
    65.72543,  2.47313, 2.50984, 2.67770, // GCGTTC
    68.27786,  2.47313, 2.40349, 2.50932, // GCGTTG
    67.21544,  2.47313, 2.34327, 2.41560, // GCGTTT
    108.41118, 2.64098, 1.77505, 1.33154, // GCTAAA
    105.67410, 2.64098, 1.75093, 1.30449, // GCTAAC
    107.17450, 2.64098, 1.72228, 1.27260, // GCTAAG
    107.59218, 2.64098, 1.97352, 1.56099, // GCTAAT
    103.55463, 2.58253, 1.59582, 1.13191, // GCTACA
    103.15189, 2.58253, 1.64624, 1.18598, // GCTACC
    103.48069, 2.58253, 1.55712, 1.09099, // GCTACG
    104.48680, 2.58253, 1.70671, 1.25191, // GCTACT
    105.61262, 2.51606, 1.64556, 1.31739, // GCTAGA
    104.37572, 2.51606, 1.53851, 1.19095, // GCTAGC
    105.02213, 2.51606, 1.50959, 1.15754, // GCTAGG
    105.66307, 2.51606, 1.75774, 1.45438, // GCTAGT
    103.47713, 3.03892, 1.85978, 1.34579, // GCTATA
    103.98588, 3.03892, 1.94134, 1.43529, // GCTATC
    103.04804, 3.03892, 1.80779, 1.28975, // GCTATG
    105.75739, 3.03892, 2.12403, 1.64257, // GCTATT
    114.05536, 2.42793, 1.78090, 1.54165, // GCTCAA
    111.53572, 2.42793, 1.81041, 1.58012, // GCTCAC
    113.27859, 2.42793, 1.82836, 1.60369, // GCTCAG
    112.74832, 2.42793, 1.88986, 1.68528, // GCTCAT
    112.55961, 2.47041, 1.79289, 1.47957, // GCTCCA
    110.63764, 2.47041, 1.84785, 1.54811, // GCTCCC
    112.33961, 2.47041, 1.84265, 1.54159, // GCTCCG
    111.76638, 2.47041, 1.96072, 1.69210, // GCTCCT
    113.54418, 2.49891, 1.96382, 1.69800, // GCTCGA
    110.64809, 2.49891, 1.81937, 1.51414, // GCTCGC
    112.74133, 2.49891, 1.90632, 1.62397, // GCTCGG
    112.38180, 2.49891, 1.99322, 1.73627, // GCTCGT
    112.19057, 2.72204, 1.91925, 1.62593, // GCTCTA
    111.00245, 2.72204, 2.04338, 1.78620, // GCTCTC
    111.87465, 2.72204, 1.85419, 1.54396, // GCTCTG
    111.97963, 2.72204, 2.16250, 1.94464, // GCTCTT
    103.98489, 2.50593, 1.74517, 1.35026, // GCTGAA
    100.06145, 2.50593, 1.62889, 1.21759, // GCTGAC
    103.20416, 2.50593, 1.70809, 1.30745, // GCTGAG
    102.17115, 2.50593, 1.88510, 1.51588, // GCTGAT
    101.79282, 2.56584, 1.60232, 1.13183, // GCTGCA
    101.02847, 2.56584, 1.61141, 1.14147, // GCTGCC
    101.75007, 2.56584, 1.60997, 1.13994, // GCTGCG
    102.47593, 2.56584, 1.79039, 1.33684, // GCTGCT
    104.25535, 2.47346, 1.70688, 1.32387, // GCTGGA
    101.53446, 2.47346, 1.69230, 1.30695, // GCTGGC
    103.35922, 2.47346, 1.73455, 1.35619, // GCTGGG
    103.48027, 2.47346, 1.84359, 1.48606, // GCTGGT
    102.34954, 3.11250, 1.86787, 1.34694, // GCTGTA
    102.17880, 3.11250, 1.96329, 1.45145, // GCTGTC
    101.94829, 3.11250, 1.99102, 1.48231, // GCTGTG
    104.10260, 3.11250, 2.21114, 1.73481, // GCTGTT
    103.19134, 1.86007, 1.41971, 1.06539, // GCTTAA
    99.14475,  1.86007, 1.27678, 0.90861, // GCTTAC
    101.93374, 1.86007, 1.39278, 1.03521, // GCTTAG
    100.84643, 1.86007, 1.35400, 0.99229, // GCTTAT
    97.39882,  1.79286, 1.44074, 1.10539, // GCTTCA
    95.54023,  1.79286, 1.41621, 1.07729, // GCTTCC
    97.41529,  1.79286, 1.39702, 1.05546, // GCTTCG
    97.06837,  1.79286, 1.41249, 1.07304, // GCTTCT
    101.29570, 1.88803, 1.29015, 0.96725, // GCTTGA
    97.09176,  1.88803, 1.23972, 0.91109, // GCTTGC
    100.36874, 1.88803, 1.29791, 0.97599, // GCTTGG
    98.74198,  1.88803, 1.36547, 1.05318, // GCTTGT
    93.77220,  1.89728, 1.31059, 0.92420, // GCTTTA
    91.31969,  1.89728, 1.17292, 0.78247, // GCTTTC
    93.07373,  1.89728, 1.25573, 0.86678, // GCTTTG
    93.28001,  1.89728, 1.23974, 0.85027, // GCTTTT
    76.42876,  1.67748, 1.20914, 0.84745, // GGAAAA
    74.24755,  1.67748, 1.02846, 0.66478, // GGAAAC
    75.10789,  1.67748, 1.15758, 0.79382, // GGAAAG
    74.41582,  1.67748, 1.04002, 0.67602, // GGAAAT
    65.60024,  2.18144, 1.53850, 1.19511, // GGAACA
    64.52218,  2.18144, 1.55502, 1.21441, // GGAACC
    65.69705,  2.18144, 1.57070, 1.23282, // GGAACG
    65.83860,  2.18144, 1.47206, 1.11854, // GGAACT
    73.90522,  1.59541, 1.22062, 0.87261, // GGAAGA
    73.20713,  1.59541, 1.02403, 0.67054, // GGAAGC
    73.06817,  1.59541, 1.17155, 0.82052, // GGAAGG
    72.71406,  1.59541, 1.03267, 0.67904, // GGAAGT
    66.34885,  1.95147, 1.37464, 1.02238, // GGAATA
    64.78872,  1.95147, 1.28777, 0.92700, // GGAATC
    65.82346,  1.95147, 1.34147, 0.98559, // GGAATG
    65.79904,  1.95147, 1.04726, 0.67984, // GGAATT
    81.57317,  2.48599, 2.00479, 1.96223, // GGACAA
    80.22381,  2.48599, 1.69098, 1.52003, // GGACAC
    80.62650,  2.48599, 1.98723, 1.93651, // GGACAG
    80.92524,  2.48599, 1.80655, 1.67850, // GGACAT
    77.34988,  2.19720, 1.88480, 1.90329, // GGACCA
    76.57879,  2.19720, 1.99876, 2.07848, // GGACCC
    77.47107,  2.19720, 1.94229, 1.99103, // GGACCG
    77.49146,  2.19720, 1.81767, 1.80252, // GGACCT
    80.24197,  1.98191, 1.78277, 1.74350, // GGACGA
    79.16421,  1.98191, 1.66132, 1.56841, // GGACGC
    79.48362,  1.98191, 1.78382, 1.74504, // GGACGG
    79.47032,  1.98191, 1.70483, 1.63043, // GGACGT
    77.27275,  1.92945, 1.58859, 1.51995, // GGACTA
    75.91172,  1.92945, 1.44300, 1.31586, // GGACTC
    76.84676,  1.92945, 1.60177, 1.53890, // GGACTG
    76.48946,  1.92945, 1.42750, 1.29472, // GGACTT
    76.36283,  1.41776, 1.06438, 0.77502, // GGAGAA
    73.05837,  1.41776, 1.17851, 0.90295, // GGAGAC
    75.37160,  1.41776, 1.03067, 0.73849, // GGAGAG
    73.90196,  1.41776, 1.12547, 0.84269, // GGAGAT
    69.36461,  1.59099, 1.32171, 0.97231, // GGAGCA
    68.15218,  1.59099, 1.30430, 0.95316, // GGAGCC
    69.42309,  1.59099, 1.26520, 0.91063, // GGAGCG
    69.00127,  1.59099, 1.21061, 0.85233, // GGAGCT
    75.62603,  1.35821, 1.04998, 0.74103, // GGAGGA
    74.15381,  1.35821, 1.04183, 0.73242, // GGAGGC
    74.89660,  1.35821, 1.00117, 0.68996, // GGAGGG
    74.39183,  1.35821, 1.07561, 0.76833, // GGAGGT
    69.78615,  1.45706, 1.01432, 0.70001, // GGAGTA
    67.64786,  1.45706, 0.97397, 0.65865, // GGAGTC
    68.71390,  1.45706, 0.97191, 0.65657, // GGAGTG
    69.35717,  1.45706, 1.18548, 0.88446, // GGAGTT
    75.03130,  2.58682, 2.15807, 2.23010, // GGATAA
    72.71990,  2.58682, 2.04672, 2.05975, // GGATAC
    74.28249,  2.58682, 1.89827, 1.83976, // GGATAG
    74.34094,  2.58682, 2.20999, 2.31106, // GGATAT
    68.39130,  2.70241, 2.34118, 2.78594, // GGATCA
    67.47663,  2.70241, 2.49718, 3.06898, // GGATCC
    68.47040,  2.70241, 2.28211, 2.68118, // GGATCG
    68.67501,  2.70241, 2.50861, 3.09009, // GGATCT
    72.36504,  2.05805, 1.90139, 2.03915, // GGATGA
    69.99222,  2.05805, 2.07033, 2.31688, // GGATGC
    71.41088,  2.05805, 1.92488, 2.07706, // GGATGG
    70.93967,  2.05805, 2.04656, 2.27709, // GGATGT
    64.91248,  2.35300, 2.14540, 2.29098, // GGATTA
    62.64008,  2.35300, 1.96828, 2.01322, // GGATTC
    64.67501,  2.35300, 2.03664, 2.11900, // GGATTG
    63.83970,  2.35300, 1.95683, 1.99568, // GGATTT
    102.94936, 2.21768, 1.75155, 1.42568, // GGCAAA
    99.91965,  2.21768, 1.77093, 1.44941, // GGCAAC
    101.93896, 2.21768, 1.74636, 1.41935, // GGCAAG
    101.15088, 2.21768, 1.93663, 1.65752, // GGCAAT
    93.70521,  1.72978, 1.18593, 0.81322, // GGCACA
    92.42839,  1.72978, 1.18877, 0.81613, // GGCACC
    93.53596,  1.72978, 1.19555, 0.82312, // GGCACG
    93.66882,  1.72978, 1.29596, 0.92897, // GGCACT
    99.90589,  2.29394, 1.72290, 1.36755, // GGCAGA
    97.26656,  2.29394, 1.69668, 1.33645, // GGCAGC
    98.72476,  2.29394, 1.71685, 1.36035, // GGCAGG
    98.43349,  2.29394, 1.88239, 1.56177, // GGCAGT
    93.60106,  2.05386, 1.34312, 0.99090, // GGCATA
    91.98458,  2.05386, 1.52712, 1.20134, // GGCATC
    92.79190,  2.05386, 1.39197, 1.04545, // GGCATG
    93.29199,  2.05386, 1.69656, 1.40673, // GGCATT
    103.42067, 1.96521, 1.46725, 1.25531, // GGCCAA
    100.87798, 1.96521, 1.48383, 1.27665, // GGCCAC
    102.86031, 1.96521, 1.47623, 1.26686, // GGCCAG
    101.99640, 1.96521, 1.60107, 1.43091, // GGCCAT
    99.37486,  1.76337, 1.31758, 1.08027, // GGCCCA
    97.21331,  1.76337, 1.24706, 0.99470, // GGCCCC
    99.49274,  1.76337, 1.33168, 1.09765, // GGCCCG
    98.97484,  1.76337, 1.33098, 1.09679, // GGCCCT
    101.93524, 1.93464, 1.52782, 1.35598, // GGCCGA
    99.44598,  1.93464, 1.45611, 1.26164, // GGCCGC
    101.00467, 1.93464, 1.43210, 1.23057, // GGCCGG
    100.49345, 1.93464, 1.57410, 1.41806, // GGCCGT
    99.37073,  1.85738, 1.37760, 1.03880, // GGCCTA
    96.79442,  1.85738, 1.43023, 1.09889, // GGCCTC
    97.95267,  1.85738, 1.42942, 1.09796, // GGCCTG
    97.43053,  1.85738, 1.57334, 1.26788, // GGCCTT
    98.54724,  2.24827, 1.69666, 1.40468, // GGCGAA
    94.40113,  2.24827, 1.62432, 1.31582, // GGCGAC
    97.43606,  2.24827, 1.68328, 1.38810, // GGCGAG
    96.15804,  2.24827, 1.84360, 1.59106, // GGCGAT
    93.23451,  1.86985, 1.32243, 0.95897, // GGCGCA
    91.77401,  1.86985, 1.48274, 1.13852, // GGCGCC
    93.16639,  1.86985, 1.37276, 1.01423, // GGCGCG
    93.30621,  1.86985, 1.45602, 1.10789, // GGCGCT
    98.12397,  2.16392, 1.74452, 1.48904, // GGCGGA
    95.32863,  2.16392, 1.63934, 1.35643, // GGCGGC
    97.39498,  2.16392, 1.70506, 1.43881, // GGCGGG
    97.03192,  2.16392, 1.80463, 1.56666, // GGCGGT
    93.09077,  2.46720, 1.60905, 1.28262, // GGCGTA
    91.51494,  2.46720, 1.72397, 1.42245, // GGCGTC
    92.69795,  2.46720, 1.65468, 1.33756, // GGCGTG
    93.36643,  2.46720, 1.92246, 1.67505, // GGCGTT
    97.22256,  1.80194, 1.58639, 1.26033, // GGCTAA
    93.68629,  1.80194, 1.42238, 1.07002, // GGCTAC
    96.28740,  1.80194, 1.56277, 1.23228, // GGCTAG
    95.09904,  1.80194, 1.53694, 1.20186, // GGCTAT
    88.87416,  2.01289, 1.66435, 1.41308, // GGCTCA
    87.28216,  2.01289, 1.66105, 1.40888, // GGCTCC
    89.10844,  2.01289, 1.75864, 1.53484, // GGCTCG
    88.80643,  2.01289, 1.83354, 1.63394, // GGCTCT
    94.19141,  1.67840, 1.47745, 1.15900, // GGCTGA
    90.89795,  1.67840, 1.40575, 1.07567, // GGCTGC
    92.76305,  1.67840, 1.45282, 1.13014, // GGCTGG
    91.94792,  1.67840, 1.46294, 1.14196, // GGCTGT
    84.86449,  1.51841, 1.36745, 1.05561, // GGCTTA
    82.16587,  1.51841, 1.26247, 0.93642, // GGCTTC
    84.34678,  1.51841, 1.29724, 0.97537, // GGCTTG
    83.40787,  1.51841, 1.16746, 0.83273, // GGCTTT
    71.85641,  1.83378, 1.24145, 0.80851, // GGGAAA
    68.86553,  1.83378, 1.38459, 0.95230, // GGGAAC
    70.29393,  1.83378, 1.32015, 0.88659, // GGGAAG
    69.39585,  1.83378, 1.36001, 0.92706, // GGGAAT
    59.57767,  2.32589, 1.85393, 1.45541, // GGGACA
    59.01548,  2.32589, 1.95737, 1.57890, // GGGACC
    59.68467,  2.32589, 1.84439, 1.44419, // GGGACG
    60.00105,  2.32589, 1.80805, 1.40173, // GGGACT
    69.49930,  1.86110, 1.33045, 0.96755, // GGGAGA
    67.88051,  1.86110, 1.21124, 0.84047, // GGGAGC
    68.27814,  1.86110, 1.38142, 1.02368, // GGGAGG
    68.31421,  1.86110, 1.23623, 0.86661, // GGGAGT
    60.44173,  1.92277, 1.66063, 1.33922, // GGGATA
    59.15800,  1.92277, 1.60670, 1.27451, // GGGATC
    59.71201,  1.92277, 1.68821, 1.37272, // GGGATG
    60.16389,  1.92277, 1.35897, 0.99142, // GGGATT
    82.64892,  1.89614, 1.63929, 1.44002, // GGGCAA
    80.45375,  1.89614, 1.38372, 1.11675, // GGGCAC
    81.63085,  1.89614, 1.59955, 1.38799, // GGGCAG
    81.37913,  1.89614, 1.45424, 1.20321, // GGGCAT
    78.40047,  1.71082, 1.65610, 1.50727, // GGGCCA
    77.73981,  1.71082, 1.99051, 1.98613, // GGGCCC
    78.39283,  1.71082, 1.62160, 1.46042, // GGGCCG
    78.20453,  1.71082, 1.56999, 1.39125, // GGGCCT
    81.19677,  1.57550, 1.46616, 1.24476, // GGGCGA
    78.81711,  1.57550, 1.40735, 1.17063, // GGGCGC
    80.10901,  1.57550, 1.45462, 1.23010, // GGGCGG
    79.32203,  1.57550, 1.40932, 1.17309, // GGGCGT
    77.55115,  1.47431, 1.47323, 1.22575, // GGGCTA
    75.59463,  1.47431, 1.39624, 1.13093, // GGGCTC
    76.91374,  1.47431, 1.39896, 1.13425, // GGGCTG
    76.62732,  1.47431, 1.19940, 0.90042, // GGGCTT
    74.38149,  1.70210, 1.09106, 0.70949, // GGGGAA
    70.10105,  1.70210, 1.38654, 1.01643, // GGGGAC
    73.50237,  1.70210, 1.12873, 0.74656, // GGGGAG
    71.41461,  1.70210, 1.31015, 0.93360, // GGGGAT
    67.60135,  1.76808, 1.43190, 1.04351, // GGGGCA
    66.32658,  1.76808, 1.52315, 1.14482, // GGGGCC
    67.62506,  1.76808, 1.40970, 1.01933, // GGGGCG
    67.43186,  1.76808, 1.36054, 0.96648, // GGGGCT
    74.24110,  1.51256, 0.95770, 0.62324, // GGGGGA
    72.76470,  1.51256, 1.07652, 0.74274, // GGGGGC
    73.67002,  1.51256, 0.95007, 0.61580, // GGGGGG
    73.25758,  1.51256, 1.05196, 0.71747, // GGGGGT
    67.75823,  1.63484, 1.12094, 0.79723, // GGGGTA
    65.27034,  1.63484, 1.18913, 0.87107, // GGGGTC
    66.67078,  1.63484, 1.13546, 0.81276, // GGGGTG
    67.10002,  1.63484, 1.40054, 1.11340, // GGGGTT
    72.79406,  2.38667, 1.94312, 1.79830, // GGGTAA
    69.76211,  2.38667, 1.71198, 1.48717, // GGGTAC
    71.42324,  2.38667, 1.82880, 1.64195, // GGGTAG
    71.47527,  2.38667, 2.00748, 1.88838, // GGGTAT
    65.17704,  2.69199, 1.97020, 2.06291, // GGGTCA
    63.75708,  2.69199, 1.87986, 1.92266, // GGGTCC
    65.35388,  2.69199, 2.04698, 2.18466, // GGGTCG
    65.12871,  2.69199, 2.06290, 2.21020, // GGGTCT
    69.18746,  2.13251, 1.73578, 1.63060, // GGGTGA
    65.95898,  2.13251, 1.76295, 1.66903, // GGGTGC
    67.82625,  2.13251, 1.73673, 1.63193, // GGGTGG
    66.88860,  2.13251, 1.79689, 1.71747, // GGGTGT
    61.20158,  2.62114, 2.02204, 1.89976, // GGGTTA
    58.50559,  2.62114, 2.06800, 1.96490, // GGGTTC
    60.40935,  2.62114, 1.92575, 1.76568, // GGGTTG
    59.71892,  2.62114, 1.87208, 1.69239, // GGGTTT
    104.16621, 2.96792, 2.31935, 1.81516, // GGTAAA
    101.53993, 2.96792, 2.40969, 1.92223, // GGTAAC
    103.16483, 2.96792, 2.26666, 1.75365, // GGTAAG
    103.49939, 2.96792, 2.53955, 2.07969, // GGTAAT
    97.18379,  2.66382, 1.93077, 1.42792, // GGTACA
    96.75782,  2.66382, 1.96027, 1.46077, // GGTACC
    97.12915,  2.66382, 1.88245, 1.37465, // GGTACG
    98.30350,  2.66382, 2.13196, 1.65682, // GGTACT
    101.42769, 2.77966, 2.25449, 1.84109, // GGTAGA
    99.56871,  2.77966, 2.27660, 1.86824, // GGTAGC
    100.78021, 2.77966, 2.11831, 1.67682, // GGTAGG
    101.86021, 2.77966, 2.44823, 2.08344, // GGTAGT
    98.64816,  3.77006, 2.27722, 1.72685, // GGTATA
    99.21754,  3.77006, 2.54459, 2.03974, // GGTATC
    97.62968,  3.77006, 2.09966, 1.52888, // GGTATG
    101.10003, 3.77006, 2.73411, 2.27181, // GGTATT
    112.06960, 2.78871, 2.53260, 2.60484, // GGTCAA
    109.89436, 2.78871, 2.40851, 2.41576, // GGTCAC
    111.44981, 2.78871, 2.45508, 2.48616, // GGTCAG
    110.80568, 2.78871, 2.51432, 2.57670, // GGTCAT
    110.66923, 2.92671, 2.43558, 2.44592, // GGTCCA
    108.71304, 2.92671, 2.23555, 2.15087, // GGTCCC
    110.15598, 2.92671, 2.51785, 2.57089, // GGTCCG
    109.78380, 2.92671, 2.30155, 2.24683, // GGTCCT
    111.52377, 2.90558, 2.61395, 2.69703, // GGTCGA
    108.99776, 2.90558, 2.34047, 2.28505, // GGTCGC
    110.87136, 2.90558, 2.45603, 2.45635, // GGTCGG
    110.76239, 2.90558, 2.46623, 2.47167, // GGTCGT
    110.35278, 3.02381, 2.53239, 2.44745, // GGTCTA
    109.88322, 3.02381, 2.48373, 2.37724, // GGTCTC
    110.07636, 3.02381, 2.45910, 2.34197, // GGTCTG
    110.41875, 3.02381, 2.75201, 2.77264, // GGTCTT
    101.51945, 2.86549, 2.28107, 1.98558, // GGTGAA
    98.20638,  2.86549, 2.30375, 2.01526, // GGTGAC
    100.84754, 2.86549, 2.33457, 2.05585, // GGTGAG
    100.33789, 2.86549, 2.47979, 2.25062, // GGTGAT
    99.46721,  2.92533, 2.12857, 1.76152, // GGTGCA
    98.92722,  2.92533, 2.13172, 1.76543, // GGTGCC
    99.44743,  2.92533, 2.22279, 1.87976, // GGTGCG
    100.47287, 2.92533, 2.41781, 2.13249, // GGTGCT
    102.11686, 2.85435, 2.35847, 2.10883, // GGTGGA
    99.98094,  2.85435, 2.29813, 2.02842, // GGTGGC
    101.02743, 2.85435, 2.21498, 1.91934, // GGTGGG
    102.08704, 2.85435, 2.37320, 2.12862, // GGTGGT
    101.78662, 3.38667, 2.42086, 2.06149, // GGTGTA
    102.24146, 3.38667, 2.59309, 2.28534, // GGTGTC
    101.58790, 3.38667, 2.53749, 2.21223, // GGTGTG
    104.47687, 3.38667, 2.88132, 2.67678, // GGTGTT
    100.33186, 2.40487, 2.12998, 2.16959, // GGTTAA
    96.62729,  2.40487, 1.93945, 1.88508, // GGTTAC
    99.15139,  2.40487, 2.06799, 2.07557, // GGTTAG
    98.21548,  2.40487, 1.91512, 1.84973, // GGTTAT
    94.09996,  2.21733, 2.20585, 2.09102, // GGTTCA
    92.42303,  2.21733, 1.90801, 1.68214, // GGTTCC
    94.18104,  2.21733, 2.05318, 1.87773, // GGTTCG
    94.05363,  2.21733, 2.14615, 2.00670, // GGTTCT
    97.87643,  2.36448, 2.01894, 1.91345, // GGTTGA
    94.24000,  2.36448, 1.79985, 1.61058, // GGTTGC
    97.13676,  2.36448, 1.90082, 1.74800, // GGTTGG
    95.62856,  2.36448, 1.97561, 1.85218, // GGTTGT
    90.55120,  1.91331, 1.86036, 1.69642, // GGTTTA
    88.32043,  1.91331, 1.68147, 1.45771, // GGTTTC
    90.01876,  1.91331, 1.77743, 1.58425, // GGTTTG
    89.51721,  1.91331, 1.58087, 1.32886, // GGTTTT
    91.47827,  2.42434, 1.47445, 1.10965, // GTAAAA
    88.65387,  2.42434, 1.75671, 1.44309, // GTAAAC
    90.50221,  2.42434, 1.55983, 1.20742, // GTAAAG
    89.29395,  2.42434, 1.79989, 1.49662, // GTAAAT
    81.82573,  2.30963, 1.73209, 1.50985, // GTAACA
    80.81677,  2.30963, 1.62785, 1.37562, // GTAACC
    81.75360,  2.30963, 1.67439, 1.43503, // GTAACG
    81.76245,  2.30963, 1.54680, 1.27417, // GTAACT
    89.74247,  2.47862, 1.48770, 1.10629, // GTAAGA
    87.72171,  2.47862, 1.61753, 1.25422, // GTAAGC
    88.76883,  2.47862, 1.54824, 1.17450, // GTAAGG
    87.81236,  2.47862, 1.72551, 1.38189, // GTAAGT
    81.25912,  2.47960, 1.60901, 1.35535, // GTAATA
    79.96964,  2.47960, 1.94188, 1.79700, // GTAATC
    80.42898,  2.47960, 1.68809, 1.45649, // GTAATG
    80.40883,  2.47960, 1.98573, 1.85821, // GTAATT
    92.93383,  1.32011, 1.16550, 0.89765, // GTACAA
    91.11572,  1.32011, 1.08543, 0.80676, // GTACAC
    92.12358,  1.32011, 1.14228, 0.87096, // GTACAG
    91.79051,  1.32011, 1.05461, 0.77264, // GTACAT
    90.08694,  1.41774, 1.32246, 1.02547, // GTACCA
    88.89257,  1.41774, 1.28303, 0.97996, // GTACCC
    89.98041,  1.41774, 1.29485, 0.99353, // GTACCG
    89.81302,  1.41774, 1.21308, 0.90092, // GTACCT
    91.84640,  1.34132, 1.15751, 0.84433, // GTACGA
    89.88847,  1.34132, 1.14746, 0.83337, // GTACGC
    90.99482,  1.34132, 1.15337, 0.83982, // GTACGG
    90.33903,  1.34132, 1.09965, 0.78183, // GTACGT
    89.14305,  1.52174, 1.34571, 1.08881, // GTACTA
    87.69767,  1.52174, 1.07971, 0.78250, // GTACTC
    88.25791,  1.52174, 1.25152, 0.97652, // GTACTG
    88.10663,  1.52174, 1.21415, 0.93311, // GTACTT
    89.36855,  2.32328, 1.46296, 1.15128, // GTAGAA
    84.53736,  2.32328, 1.83673, 1.61957, // GTAGAC
    88.15302,  2.32328, 1.60557, 1.32365, // GTAGAG
    86.17426,  2.32328, 1.85569, 1.64470, // GTAGAT
    83.89273,  2.28919, 1.38505, 1.15302, // GTAGCA
    82.43136,  2.28919, 1.42142, 1.19874, // GTAGCC
    83.84035,  2.28919, 1.45874, 1.24626, // GTAGCG
    83.75964,  2.28919, 1.55741, 1.37482, // GTAGCT
    88.56635,  2.34031, 1.57789, 1.28295, // GTAGGA
    86.11981,  2.34031, 1.68167, 1.41158, // GTAGGC
    87.69378,  2.34031, 1.46845, 1.15181, // GTAGGG
    87.02742,  2.34031, 1.72222, 1.46294, // GTAGGT
    82.53431,  2.72334, 1.72354, 1.54604, // GTAGTA
    80.28733,  2.72334, 1.90415, 1.79533, // GTAGTC
    81.61561,  2.72334, 1.72703, 1.55075, // GTAGTG
    81.43765,  2.72334, 1.98864, 1.91614, // GTAGTT
    88.06446,  1.88263, 1.71881, 1.54958, // GTATAA
    85.26141,  1.88263, 1.78830, 1.64449, // GTATAC
    87.02028,  1.88263, 1.61101, 1.40611, // GTATAG
    86.56721,  1.88263, 1.75708, 1.60162, // GTATAT
    82.90651,  2.46291, 2.29072, 2.38286, // GTATCA
    81.79616,  2.46291, 2.32343, 2.43407, // GTATCC
    82.89747,  2.46291, 2.30951, 2.41223, // GTATCG
    82.89172,  2.46291, 2.29959, 2.39671, // GTATCT
    85.91856,  2.11919, 1.64959, 1.39550, // GTATGA
    82.84562,  2.11919, 1.91672, 1.74785, // GTATGC
    84.77322,  2.11919, 1.81800, 1.61457, // GTATGG
    83.72264,  2.11919, 1.88372, 1.70290, // GTATGT
    78.30410,  2.69506, 2.31913, 2.28408, // GTATTA
    75.13481,  2.69506, 2.33506, 2.30765, // GTATTC
    77.26291,  2.69506, 2.29886, 2.25420, // GTATTG
    76.52266,  2.69506, 2.17784, 2.07856, // GTATTT
    102.58827, 2.08052, 1.62019, 1.37183, // GTCAAA
    100.24962, 2.08052, 1.80267, 1.61001, // GTCAAC
    101.57558, 2.08052, 1.73346, 1.51818, // GTCAAG
    101.10493, 2.08052, 1.82250, 1.63664, // GTCAAT
    95.74167,  2.10126, 1.42242, 1.09346, // GTCACA
    94.94477,  2.10126, 1.60290, 1.30804, // GTCACC
    95.69148,  2.10126, 1.53966, 1.23140, // GTCACG
    95.18746,  2.10126, 1.50201, 1.18650, // GTCACT
    100.65451, 2.40061, 1.67755, 1.42533, // GTCAGA
    98.61494,  2.40061, 1.82767, 1.62088, // GTCAGC
    99.71725,  2.40061, 1.74717, 1.51497, // GTCAGG
    99.43729,  2.40061, 1.92506, 1.75215, // GTCAGT
    94.21125,  2.47361, 1.80561, 1.56484, // GTCATA
    92.62737,  2.47361, 1.87321, 1.65352, // GTCATC
    93.60753,  2.47361, 1.82770, 1.59364, // GTCATG
    93.59040,  2.47361, 2.01227, 1.84104, // GTCATT
    101.73698, 1.63253, 1.22876, 0.99652, // GTCCAA
    99.43852,  1.63253, 1.35619, 1.15548, // GTCCAC
    101.22380, 1.63253, 1.24909, 1.02134, // GTCCAG
    100.28255, 1.63253, 1.43772, 1.26123, // GTCCAT
    98.63390,  1.60868, 1.20702, 0.97568, // GTCCCA
    96.75032,  1.60868, 1.18789, 0.95258, // GTCCCC
    98.20252,  1.60868, 1.16358, 0.92349, // GTCCCG
    98.03053,  1.60868, 1.28546, 1.07232, // GTCCCT
    100.68426, 1.82797, 1.29791, 1.02308, // GTCCGA
    98.47872,  1.82797, 1.40698, 1.15470, // GTCCGC
    99.81807,  1.82797, 1.27567, 0.99689, // GTCCGG
    99.08231,  1.82797, 1.46479, 1.22660, // GTCCGT
    97.26094,  1.91501, 1.14967, 0.87578, // GTCCTA
    95.30014,  1.91501, 1.44143, 1.22949, // GTCCTC
    96.81827,  1.91501, 1.45871, 1.25167, // GTCCTG
    96.09497,  1.91501, 1.62577, 1.47273, // GTCCTT
    99.46575,  2.59137, 1.74631, 1.52667, // GTCGAA
    95.11484,  2.59137, 2.09233, 2.00220, // GTCGAC
    98.27732,  2.59137, 1.83182, 1.64017, // GTCGAG
    96.68127,  2.59137, 2.02560, 1.90718, // GTCGAT
    94.84823,  2.48599, 1.54808, 1.28470, // GTCGCA
    93.40353,  2.48599, 1.68402, 1.45759, // GTCGCC
    94.53344,  2.48599, 1.62588, 1.38275, // GTCGCG
    94.43036,  2.48599, 1.77432, 1.57637, // GTCGCT
    99.17000,  2.51279, 1.71040, 1.45432, // GTCGGA
    96.10623,  2.51279, 1.88136, 1.67771, // GTCGGC
    98.21251,  2.51279, 1.82460, 1.60237, // GTCGGG
    97.38291,  2.51279, 2.02844, 1.87826, // GTCGGT
    93.49116,  5.74004, 1.89813, 1.73373, // GTCGTA
    91.27454,  5.74004, 2.10904, 2.03058, // GTCGTC
    92.96333,  5.74004, 1.93755, 1.78802, // GTCGTG
    92.50608,  5.74004, 1.91348, 1.75480, // GTCGTT
    97.20339,  1.86097, 1.62183, 1.28127, // GTCTAA
    94.14489,  1.86097, 1.63312, 1.29467, // GTCTAC
    96.40072,  1.86097, 1.71766, 1.39650, // GTCTAG
    95.41914,  1.86097, 1.74664, 1.43199, // GTCTAT
    91.90459,  2.32915, 2.15105, 2.15376, // GTCTCA
    90.37184,  2.32915, 2.11079, 2.09359, // GTCTCC
    91.59987,  2.32915, 2.07812, 2.04517, // GTCTCG
    91.20402,  2.32915, 2.08590, 2.05666, // GTCTCT
    95.08401,  2.03287, 1.64094, 1.38064, // GTCTGA
    91.85663,  2.03287, 1.81000, 1.59941, // GTCTGC
    93.92966,  2.03287, 1.72403, 1.48683, // GTCTGG
    92.81320,  2.03287, 1.82282, 1.61644, // GTCTGT
    86.91404,  2.37489, 2.08710, 1.99040, // GTCTTA
    84.05151,  2.37489, 2.08628, 1.98923, // GTCTTC
    85.95131,  2.37489, 1.93905, 1.78242, // GTCTTG
    85.30013,  2.37489, 2.05228, 1.94081, // GTCTTT
    85.34118,  3.07534, 1.82938, 1.39882, // GTGAAA
    81.97262,  3.07534, 2.03028, 1.63546, // GTGAAC
    84.07770,  3.07534, 1.90765, 1.48954, // GTGAAG
    83.01400,  3.07534, 2.14695, 1.77844, // GTGAAT
    74.83836,  2.53545, 1.89597, 1.65706, // GTGACA
    73.91491,  2.53545, 1.86887, 1.62166, // GTGACC
    74.78076,  2.53545, 1.85556, 1.60436, // GTGACG
    74.77096,  2.53545, 1.69993, 1.40682, // GTGACT
    84.44492,  3.17429, 1.79970, 1.44230, // GTGAGA
    82.22070,  3.17429, 1.88356, 1.54428, // GTGAGC
    83.39819,  3.17429, 1.86563, 1.52228, // GTGAGG
    82.44459,  3.17429, 2.03734, 1.73720, // GTGAGT
    74.44972,  2.69379, 1.74880, 1.48787, // GTGATA
    73.32149,  2.69379, 2.15983, 2.04212, // GTGATC
    73.66173,  2.69379, 1.85521, 1.62570, // GTGATG
    73.92065,  2.69379, 2.14246, 2.01754, // GTGATT
    91.43005,  1.48208, 1.29466, 1.03335, // GTGCAA
    88.97947,  1.48208, 1.22138, 0.94687, // GTGCAC
    90.46120,  1.48208, 1.27689, 1.01214, // GTGCAG
    89.92898,  1.48208, 1.21305, 0.93720, // GTGCAT
    88.12535,  1.66578, 1.51400, 1.25181, // GTGCCA
    86.99090,  1.66578, 1.47590, 1.20485, // GTGCCC
    87.94424,  1.66578, 1.46995, 1.19758, // GTGCCG
    87.76856,  1.66578, 1.45268, 1.17654, // GTGCCT
    90.18016,  1.67467, 1.32817, 0.97807, // GTGCGA
    87.78096,  1.67467, 1.35531, 1.00822, // GTGCGC
    89.09269,  1.67467, 1.36016, 1.01363, // GTGCGG
    88.17006,  1.67467, 1.33712, 0.98798, // GTGCGT
    86.88256,  1.83904, 1.52358, 1.30842, // GTGCTA
    84.93876,  1.83904, 1.41585, 1.17213, // GTGCTC
    86.08563,  1.83904, 1.47565, 1.24716, // GTGCTG
    85.98037,  1.83904, 1.48283, 1.25628, // GTGCTT
    87.53023,  3.22959, 1.93193, 1.68915, // GTGGAA
    82.87095,  3.22959, 2.31030, 2.20893, // GTGGAC
    86.43826,  3.22959, 1.94327, 1.70404, // GTGGAG
    84.27046,  3.22959, 2.23788, 2.10588, // GTGGAT
    82.31577,  2.83028, 1.62622, 1.33704, // GTGGCA
    81.57911,  2.83028, 1.82331, 1.58734, // GTGGCC
    82.26793,  2.83028, 1.66899, 1.39014, // GTGGCG
    82.35203,  2.83028, 1.80201, 1.55961, // GTGGCT
    87.99480,  2.98826, 1.81644, 1.48090, // GTGGGA
    85.47941,  2.98826, 1.98039, 1.68585, // GTGGGC
    86.94466,  2.98826, 1.71797, 1.36213, // GTGGGG
    86.33459,  2.98826, 2.02295, 1.74049, // GTGGGT
    81.90012,  3.42265, 2.05934, 1.72818, // GTGGTA
    79.59412,  3.42265, 2.20673, 1.91699, // GTGGTC
    80.58593,  3.42265, 2.12910, 1.81673, // GTGGTG
    80.83785,  3.42265, 2.34397, 2.09858, // GTGGTT
    85.15663,  2.22980, 1.93499, 1.66408, // GTGTAA
    81.64370,  2.22980, 2.07233, 1.84435, // GTGTAC
    84.26899,  2.22980, 1.94352, 1.67509, // GTGTAG
    82.96525,  2.22980, 2.00567, 1.75608, // GTGTAT
    79.81156,  2.83133, 2.49998, 2.46313, // GTGTCA
    78.17194,  2.83133, 2.42315, 2.35046, // GTGTCC
    79.87530,  2.83133, 2.50247, 2.46680, // GTGTCG
    79.51664,  2.83133, 2.51313, 2.48258, // GTGTCT
    82.48641,  2.65515, 2.14261, 1.94155, // GTGTGA
    78.73788,  2.65515, 2.28857, 2.14329, // GTGTGC
    81.15679,  2.65515, 2.24286, 2.07939, // GTGTGG
    79.95704,  2.65515, 2.29864, 2.15745, // GTGTGT
    75.01523,  3.04048, 2.70018, 2.78171, // GTGTTA
    71.79837,  3.04048, 2.63516, 2.68185, // GTGTTC
    73.99449,  3.04048, 2.55317, 2.55766, // GTGTTG
    73.04893,  3.04048, 2.47416, 2.43986, // GTGTTT
    104.98309, 2.16371, 1.43973, 1.11240, // GTTAAA
    102.79086, 2.16371, 1.58255, 1.28197, // GTTAAC
    104.17725, 2.16371, 1.45860, 1.13434, // GTTAAG
    104.15990, 2.16371, 1.65787, 1.37457, // GTTAAT
    99.67702,  2.01953, 1.21174, 0.83025, // GTTACA
    99.15126,  2.01953, 1.27946, 0.90081, // GTTACC
    99.58219,  2.01953, 1.22536, 0.84429, // GTTACG
    100.35581, 2.01953, 1.34855, 0.97476, // GTTACT
    103.10860, 2.04194, 1.39157, 1.06601, // GTTAGA
    101.33784, 2.04194, 1.52240, 1.21982, // GTTAGC
    101.97894, 2.04194, 1.48526, 1.17546, // GTTAGG
    102.59455, 2.04194, 1.63479, 1.35737, // GTTAGT
    99.40517,  2.50557, 1.44195, 1.08412, // GTTATA
    99.46733,  2.50557, 1.70288, 1.39133, // GTTATC
    98.86304,  2.50557, 1.62983, 1.30276, // GTTATG
    101.03991, 2.50557, 1.78232, 1.48981, // GTTATT
    108.44203, 2.19857, 1.36265, 1.04131, // GTTCAA
    106.52641, 2.19857, 1.41891, 1.10646, // GTTCAC
    108.01739, 2.19857, 1.38370, 1.06554, // GTTCAG
    107.51442, 2.19857, 1.54093, 1.25221, // GTTCAT
    106.88314, 2.17160, 1.37027, 1.03280, // GTTCCA
    105.27742, 2.17160, 1.37628, 1.03960, // GTTCCC
    106.71414, 2.17160, 1.36695, 1.02904, // GTTCCG
    106.34612, 2.17160, 1.43333, 1.10491, // GTTCCT
    108.13160, 2.22607, 1.45104, 1.14446, // GTTCGA
    105.87703, 2.22607, 1.39271, 1.07615, // GTTCGC
    107.58331, 2.22607, 1.42238, 1.11072, // GTTCGG
    107.45630, 2.22607, 1.55811, 1.27344, // GTTCGT
    106.80446, 2.26458, 1.56791, 1.29084, // GTTCTA
    105.69961, 2.26458, 1.55459, 1.27443, // GTTCTC
    106.51769, 2.26458, 1.49548, 1.20244, // GTTCTG
    106.68303, 2.26458, 1.69596, 1.45217, // GTTCTT
    101.47866, 2.11369, 1.46447, 1.24252, // GTTGAA
    98.00233,  2.11369, 1.65688, 1.49526, // GTTGAC
    100.65991, 2.11369, 1.45244, 1.22724, // GTTGAG
    99.73427,  2.11369, 1.75609, 1.63155, // GTTGAT
    98.84041,  2.06630, 1.23715, 0.89325, // GTTGCA
    98.01209,  2.06630, 1.33071, 0.99648, // GTTGCC
    98.91706,  2.06630, 1.27064, 0.92977, // GTTGCG
    99.55696,  2.06630, 1.38579, 1.05898, // GTTGCT
    101.60254, 1.98654, 1.49026, 1.27261, // GTTGGA
    99.29000,  1.98654, 1.44736, 1.21806, // GTTGGC
    100.66989, 1.98654, 1.39821, 1.15654, // GTTGGG
    100.96343, 1.98654, 1.48975, 1.27196, // GTTGGT
    99.01444,  2.51049, 1.57589, 1.25767, // GTTGTA
    99.03220,  2.51049, 1.67098, 1.37321, // GTTGTC
    98.91975,  2.51049, 1.59641, 1.28231, // GTTGTG
    100.86573, 2.51049, 1.76040, 1.48490, // GTTGTT
    99.78830,  1.70840, 1.12987, 0.80426, // GTTTAA
    96.21788,  1.70840, 1.04056, 0.71081, // GTTTAC
    98.85820,  1.70840, 1.14939, 0.82519, // GTTTAG
    97.65538,  1.70840, 1.08895, 0.76096, // GTTTAT
    94.49598,  1.56977, 1.18595, 0.89984, // GTTTCA
    92.74278,  1.56977, 1.18711, 0.90117, // GTTTCC
    94.45128,  1.56977, 1.20071, 0.91669, // GTTTCG
    94.12721,  1.56977, 1.18075, 0.89392, // GTTTCT
    97.77427,  1.62351, 1.04164, 0.71983, // GTTTGA
    94.73253,  1.62351, 1.01339, 0.69075, // GTTTGC
    96.98571,  1.62351, 1.05309, 0.73174, // GTTTGG
    95.70799,  1.62351, 1.07646, 0.75622, // GTTTGT
    91.21152,  1.51391, 1.09706, 0.77692, // GTTTTA
    88.91613,  1.51391, 1.09798, 0.77790, // GTTTTC
    90.72744,  1.51391, 1.07626, 0.75493, // GTTTTG
    90.26131,  1.51391, 0.98819, 0.66419, // GTTTTT
    88.53078,  1.51785, 1.04825, 0.71590, // TAAAAA
    85.76570,  1.51785, 1.29167, 0.97923, // TAAAAC
    87.79532,  1.51785, 1.09756, 0.76701, // TAAAAG
    86.35140,  1.51785, 1.33038, 1.02357, // TAAAAT
    78.56690,  1.70501, 1.41457, 1.06653, // TAAACA
    77.45211,  1.70501, 1.41067, 1.06212, // TAAACC
    78.70234,  1.70501, 1.40818, 1.05931, // TAAACG
    78.41393,  1.70501, 1.24338, 0.87890, // TAAACT
    86.60806,  1.53757, 1.03914, 0.71096, // TAAAGA
    84.61097,  1.53757, 1.16995, 0.84933, // TAAAGC
    85.62810,  1.53757, 1.08461, 0.75812, // TAAAGG
    84.76401,  1.53757, 1.25015, 0.93815, // TAAAGT
    78.23306,  1.54253, 1.18119, 0.88959, // TAAATA
    76.29628,  1.54253, 1.35270, 1.09022, // TAAATC
    77.36571,  1.54253, 1.18517, 0.89410, // TAAATG
    77.38321,  1.54253, 1.52256, 1.30190, // TAAATT
    90.92847,  1.96203, 1.38066, 1.21010, // TAACAA
    88.91254,  1.96203, 1.14774, 0.91719, // TAACAC
    90.10884,  1.96203, 1.25407, 1.04756, // TAACAG
    89.84615,  1.96203, 1.14323, 0.91178, // TAACAT
    87.10763,  1.83613, 1.35635, 1.23378, // TAACCA
    85.95794,  1.83613, 1.31104, 1.17247, // TAACCC
    87.12905,  1.83613, 1.35384, 1.23036, // TAACCG
    86.95059,  1.83613, 1.24371, 1.08332, // TAACCT
    89.69204,  1.55453, 1.14701, 0.95581, // TAACGA
    87.67810,  1.55453, 1.09222, 0.88815, // TAACGC
    88.81745,  1.55453, 1.19051, 1.01070, // TAACGG
    88.27069,  1.55453, 1.09095, 0.88661, // TAACGT
    86.43258,  1.58806, 1.28098, 1.12589, // TAACTA
    84.36992,  1.58806, 1.10521, 0.90229, // TAACTC
    85.65073,  1.58806, 1.12051, 0.92109, // TAACTG
    84.87685,  1.58806, 0.93079, 0.69736, // TAACTT
    86.82051,  1.46160, 1.12707, 0.82856, // TAAGAA
    82.50102,  1.46160, 1.42750, 1.18103, // TAAGAC
    85.93323,  1.46160, 1.16373, 0.86932, // TAAGAG
    83.60207,  1.46160, 1.44931, 1.20820, // TAAGAT
    80.63391,  1.42710, 1.16076, 0.87252, // TAAGCA
    78.85534,  1.42710, 1.10092, 0.80592, // TAAGCC
    80.56624,  1.42710, 1.07986, 0.78292, // TAAGCG
    80.02239,  1.42710, 1.09146, 0.79557, // TAAGCT
    85.75052,  1.40989, 1.09476, 0.79409, // TAAGGA
    82.79740,  1.40989, 1.19238, 0.90265, // TAAGGC
    84.68214,  1.40989, 1.04527, 0.74086, // TAAGGG
    83.91311,  1.40989, 1.30124, 1.02904, // TAAGGT
    79.12672,  1.48703, 1.21288, 0.95709, // TAAGTA
    76.62803,  1.48703, 1.44204, 1.24077, // TAAGTC
    78.33126,  1.48703, 1.26271, 1.01667, // TAAGTG
    77.96321,  1.48703, 1.59025, 1.43688, // TAAGTT
    84.20895,  1.98024, 1.50588, 1.40110, // TAATAA
    82.00571,  1.98024, 1.51851, 1.41876, // TAATAC
    83.38998,  1.98024, 1.40432, 1.26177, // TAATAG
    83.24214,  1.98024, 1.52794, 1.43200, // TAATAT
    77.27125,  2.22770, 1.76574, 2.01729, // TAATCA
    76.40816,  2.22770, 1.92251, 2.29182, // TAATCC
    77.44228,  2.22770, 1.76527, 2.01648, // TAATCG
    77.24688,  2.22770, 1.74794, 1.98685, // TAATCT
    81.50683,  1.67617, 1.41977, 1.32125, // TAATGA
    79.03523,  1.67617, 1.51592, 1.45771, // TAATGC
    80.68196,  1.67617, 1.40260, 1.29735, // TAATGG
    79.80955,  1.67617, 1.51219, 1.45233, // TAATGT
    72.01753,  2.00867, 1.90015, 1.90721, // TAATTA
    69.30088,  2.00867, 1.94130, 1.96950, // TAATTC
    71.59567,  2.00867, 1.87523, 1.86980, // TAATTG
    70.76287,  2.00867, 1.83364, 1.80795, // TAATTT
    106.45496, 2.19132, 1.57068, 1.25868, // TACAAA
    103.46311, 2.19132, 1.56254, 1.24890, // TACAAC
    105.46405, 2.19132, 1.51941, 1.19756, // TACAAG
    104.77095, 2.19132, 1.64143, 1.34467, // TACAAT
    97.87495,  1.74166, 1.03375, 0.66048, // TACACA
    96.50944,  1.74166, 1.07960, 0.70491, // TACACC
    97.85826,  1.74166, 1.07618, 0.70156, // TACACG
    97.59567,  1.74166, 1.14367, 0.76858, // TACACT
    103.50685, 2.28987, 1.57936, 1.25691, // TACAGA
    100.72278, 2.28987, 1.53685, 1.20650, // TACAGC
    102.34865, 2.28987, 1.53831, 1.20823, // TACAGG
    101.97361, 2.28987, 1.62676, 1.31391, // TACAGT
    96.96352,  2.17760, 1.21873, 0.89389, // TACATA
    95.66174,  2.17760, 1.29315, 0.97701, // TACATC
    96.23039,  2.17760, 1.25129, 0.92996, // TACATG
    96.82573,  2.17760, 1.45466, 1.16565, // TACATT
    106.36283, 1.88264, 1.21948, 0.96161, // TACCAA
    103.52762, 1.88264, 1.26103, 1.01116, // TACCAC
    105.47664, 1.88264, 1.23387, 0.97867, // TACCAG
    104.74809, 1.88264, 1.35542, 1.12679, // TACCAT
    102.13969, 1.75332, 1.09768, 0.82855, // TACCCA
    100.42075, 1.75332, 1.06461, 0.79139, // TACCCC
    102.27253, 1.75332, 1.09803, 0.82894, // TACCCG
    101.56056, 1.75332, 1.13717, 0.87365, // TACCCT
    104.89360, 1.90443, 1.36780, 1.09917, // TACCGA
    102.22541, 1.90443, 1.24941, 0.95960, // TACCGC
    103.84748, 1.90443, 1.27284, 0.98672, // TACCGG
    103.05126, 1.90443, 1.39336, 1.13012, // TACCGT
    101.04011, 1.83098, 1.25729, 0.92140, // TACCTA
    98.94952,  1.83098, 1.28097, 0.94755, // TACCTC
    100.50265, 1.83098, 1.20605, 0.86566, // TACCTG
    99.88463,  1.83098, 1.44897, 1.13995, // TACCTT
    101.47914, 2.24722, 1.49205, 1.18427, // TACGAA
    97.03130,  2.24722, 1.51610, 1.21302, // TACGAC
    100.58111, 2.24722, 1.53171, 1.23180, // TACGAG
    98.85062,  2.24722, 1.72358, 1.47036, // TACGAT
    96.29177,  2.02317, 1.18780, 0.83827, // TACGCA
    95.04988,  2.02317, 1.17086, 0.82041, // TACGCC
    96.49731,  2.02317, 1.22257, 0.87535, // TACGCG
    96.37102,  2.02317, 1.33527, 0.99914, // TACGCT
    101.07238, 2.22668, 1.52491, 1.21724, // TACGGA
    97.68104,  2.22668, 1.44678, 1.12490, // TACGGC
    100.24026, 2.22668, 1.50612, 1.19481, // TACGGG
    99.38421,  2.22668, 1.64494, 1.36375, // TACGGT
    95.54601,  2.64556, 1.34510, 0.97455, // TACGTA
    93.63168,  2.64556, 1.47465, 1.11868, // TACGTC
    94.65124,  2.64556, 1.45408, 1.09536, // TACGTG
    95.14513,  2.64556, 1.67447, 1.35359, // TACGTT
    100.18495, 1.69847, 1.30999, 0.88565, // TACTAA
    96.95703,  1.69847, 1.28624, 0.86167, // TACTAC
    99.05523,  1.69847, 1.32066, 0.89648, // TACTAG
    98.35818,  1.69847, 1.25276, 0.82825, // TACTAT
    92.35163,  1.86136, 1.51088, 1.33193, // TACTCA
    90.71414,  1.86136, 1.48240, 1.29446, // TACTCC
    92.34909,  1.86136, 1.45622, 1.26031, // TACTCG
    92.11751,  1.86136, 1.59732, 1.44786, // TACTCT
    97.26936,  1.64486, 1.23651, 0.89529, // TACTGA
    93.77413,  1.64486, 1.19443, 0.84999, // TACTGC
    95.92278,  1.64486, 1.20872, 0.86528, // TACTGG
    94.81972,  1.64486, 1.25162, 0.91176, // TACTGT
    87.25996,  1.46961, 1.14176, 0.80572, // TACTTA
    84.62581,  1.46961, 1.25815, 0.93200, // TACTTC
    86.76065,  1.46961, 1.12569, 0.78877, // TACTTG
    86.00177,  1.46961, 1.05356, 0.71419, // TACTTT
    80.55679,  1.82916, 1.34686, 0.93623, // TAGAAA
    77.48297,  1.82916, 1.59610, 1.20779, // TAGAAC
    79.44515,  1.82916, 1.39628, 0.98823, // TAGAAG
    78.03953,  1.82916, 1.57940, 1.18887, // TAGAAT
    69.19734,  2.14481, 1.93839, 1.58023, // TAGACA
    68.10677,  2.14481, 1.86980, 1.49710, // TAGACC
    68.97301,  2.14481, 1.72200, 1.32315, // TAGACG
    68.90267,  2.14481, 1.70331, 1.30166, // TAGACT
    78.64136,  1.82854, 1.27735, 0.93715, // TAGAGA
    76.66943,  1.82854, 1.45954, 1.14465, // TAGAGC
    77.27561,  1.82854, 1.24382, 0.90050, // TAGAGG
    77.19863,  1.82854, 1.45405, 1.13819, // TAGAGT
    69.06633,  1.85459, 1.61310, 1.38601, // TAGATA
    67.73844,  1.85459, 1.35435, 1.06628, // TAGATC
    68.56502,  1.85459, 1.51024, 1.25557, // TAGATG
    68.51548,  1.85459, 1.62437, 1.40056, // TAGATT
    86.97801,  1.87706, 1.36709, 1.09125, // TAGCAA
    84.57760,  1.87706, 1.08398, 0.77048, // TAGCAC
    85.92300,  1.87706, 1.28925, 0.99939, // TAGCAG
    85.36749,  1.87706, 1.10522, 0.79324, // TAGCAT
    82.63835,  1.70618, 1.33413, 1.12527, // TAGCCA
    81.32429,  1.70618, 1.27170, 1.04722, // TAGCCC
    82.45350,  1.70618, 1.24378, 1.01292, // TAGCCG
    82.11752,  1.70618, 1.24639, 1.01611, // TAGCCT
    84.36410,  1.56014, 1.24450, 1.00094, // TAGCGA
    81.35600,  1.56014, 1.21873, 0.97002, // TAGCGC
    83.15054,  1.56014, 1.17242, 0.91525, // TAGCGG
    82.07118,  1.56014, 1.23290, 0.98698, // TAGCGT
    81.53271,  1.42173, 1.26420, 0.99940, // TAGCTA
    79.26478,  1.42173, 1.10998, 0.82222, // TAGCTC
    80.94764,  1.42173, 1.19038, 0.91316, // TAGCTG
    80.24891,  1.42173, 1.00337, 0.70666, // TAGCTT
    80.01890,  1.83346, 1.46665, 1.09311, // TAGGAA
    74.62770,  1.83346, 1.89373, 1.60380, // TAGGAC
    78.45331,  1.83346, 1.55655, 1.19513, // TAGGAG
    76.29148,  1.83346, 1.69548, 1.35867, // TAGGAT
    73.80171,  1.67690, 1.46621, 1.19172, // TAGGCA
    72.25192,  1.67690, 1.44969, 1.17163, // TAGGCC
    73.39346,  1.67690, 1.50625, 1.24087, // TAGGCG
    73.43874,  1.67690, 1.32299, 1.02144, // TAGGCT
    80.32358,  1.68720, 1.23642, 0.91900, // TAGGGA
    77.41749,  1.68720, 1.42840, 1.14114, // TAGGGC
    79.19140,  1.68720, 1.19575, 0.87403, // TAGGGG
    78.20710,  1.68720, 1.43927, 1.15419, // TAGGGT
    72.73568,  1.76089, 1.40538, 1.08990, // TAGGTA
    70.66788,  1.76089, 1.66241, 1.40219, // TAGGTC
    72.07976,  1.76089, 1.52375, 1.23047, // TAGGTG
    71.77100,  1.76089, 1.73351, 1.49310, // TAGGTT
    77.96780,  2.04805, 1.60956, 1.45064, // TAGTAA
    74.34658,  2.04805, 1.68446, 1.55306, // TAGTAC
    77.15253,  2.04805, 1.59692, 1.43360, // TAGTAG
    75.80110,  2.04805, 1.90187, 1.86325, // TAGTAT
    70.34646,  2.42133, 2.32320, 2.59808, // TAGTCA
    68.86867,  2.42133, 2.45032, 2.81423, // TAGTCC
    70.01596,  2.42133, 2.11218, 2.25227, // TAGTCG
    70.05103,  2.42133, 2.37287, 2.68184, // TAGTCT
    74.16668,  1.99693, 1.68145, 1.65152, // TAGTGA
    70.40746,  1.99693, 2.03056, 2.19170, // TAGTGC
    72.97021,  1.99693, 1.87816, 1.94965, // TAGTGG
    71.22354,  1.99693, 1.98209, 2.11370, // TAGTGT
    64.98179,  2.35636, 2.13700, 2.06703, // TAGTTA
    61.86187,  2.35636, 2.41609, 2.48489, // TAGTTC
    64.22523,  2.35636, 2.24405, 2.22427, // TAGTTG
    63.27101,  2.35636, 2.18727, 2.14039, // TAGTTT
    110.64584, 3.16286, 2.21893, 1.75215, // TATAAA
    108.35993, 3.16286, 2.33482, 1.89121, // TATAAC
    109.70926, 3.16286, 2.17434, 1.69961, // TATAAG
    110.01330, 3.16286, 2.61621, 2.24319, // TATAAT
    104.54216, 3.09766, 1.79706, 1.27165, // TATACA
    105.20455, 3.09766, 2.09643, 1.60230, // TATACC
    105.31443, 3.09766, 1.92222, 1.40678, // TATACG
    106.55406, 3.09766, 2.28143, 1.81900, // TATACT
    108.37237, 3.02439, 2.10034, 1.73867, // TATAGA
    106.37793, 3.02439, 2.16133, 1.81496, // TATAGC
    107.46821, 3.02439, 2.14942, 1.79997, // TATAGG
    108.80078, 3.02439, 2.37806, 2.09469, // TATAGT
    106.32974, 3.75132, 2.32298, 1.80905, // TATATA
    107.52383, 3.75132, 2.52285, 2.04747, // TATATC
    105.85022, 3.75132, 2.44323, 1.95132, // TATATG
    109.53560, 3.75132, 2.72107, 2.29346, // TATATT
    119.08138, 2.97261, 2.38936, 2.44342, // TATCAA
    116.64436, 2.97261, 2.34762, 2.37968, // TATCAC
    118.40572, 2.97261, 2.34141, 2.37025, // TATCAG
    117.80427, 2.97261, 2.43901, 2.51998, // TATCAT
    117.98549, 3.04768, 2.24938, 2.12805, // TATCCA
    115.79225, 3.04768, 2.20052, 2.05909, // TATCCC
    117.78732, 3.04768, 2.22441, 2.09271, // TATCCG
    117.18889, 3.04768, 2.50667, 2.50341, // TATCCT
    118.61262, 3.00027, 2.49509, 2.50103, // TATCGA
    115.74231, 3.00027, 2.38344, 2.33505, // TATCGC
    118.08500, 3.00027, 2.36293, 2.30498, // TATCGG
    117.62897, 3.00027, 2.46633, 2.45792, // TATCGT
    117.88430, 3.10410, 2.36295, 2.22603, // TATCTA
    116.66588, 3.10410, 2.49604, 2.41673, // TATCTC
    117.77047, 3.10410, 2.45321, 2.35480, // TATCTG
    117.90044, 3.10410, 2.66563, 2.66716, // TATCTT
    106.87456, 2.99484, 2.19874, 1.88450, // TATGAA
    103.46319, 2.99484, 2.13518, 1.80337, // TATGAC
    106.13889, 2.99484, 1.99640, 1.63043, // TATGAG
    105.72421, 2.99484, 2.29151, 2.00501, // TATGAT
    105.49231, 3.13415, 1.98592, 1.55719, // TATGCA
    104.95270, 3.13415, 2.14093, 1.74302, // TATGCC
    105.49572, 3.13415, 2.04695, 1.62952, // TATGCG
    106.82756, 3.13415, 2.29070, 1.92908, // TATGCT
    107.50048, 2.93123, 2.18803, 1.82619, // TATGGA
    105.18488, 2.93123, 2.09599, 1.71219, // TATGGC
    106.69801, 2.93123, 2.07913, 1.69157, // TATGGG
    107.36226, 2.93123, 2.25258, 1.90760, // TATGGT
    107.31879, 3.56800, 2.47436, 2.05121, // TATGTA
    107.85663, 3.56800, 2.58961, 2.19618, // TATGTC
    107.13375, 3.56800, 2.44950, 2.02039, // TATGTG
    110.48252, 3.56800, 2.92715, 2.63928, // TATGTT
    106.93805, 2.18027, 1.64195, 1.51804, // TATTAA
    103.17009, 2.18027, 1.47674, 1.29478, // TATTAC
    106.02449, 2.18027, 1.51161, 1.34093, // TATTAG
    104.49588, 2.18027, 1.54459, 1.38505, // TATTAT
    100.72848, 2.09022, 1.71995, 1.53227, // TATTCA
    98.70827,  2.09022, 1.73088, 1.54689, // TATTCC
    100.65656, 2.09022, 1.69850, 1.50370, // TATTCG
    100.35470, 2.09022, 1.69430, 1.49811, // TATTCT
    104.45090, 2.20088, 1.57124, 1.39011, // TATTGA
    100.62956, 2.20088, 1.49380, 1.28862, // TATTGC
    103.69423, 2.20088, 1.51226, 1.31258, // TATTGG
    102.37981, 2.20088, 1.58313, 1.40592, // TATTGT
    96.74776,  1.92511, 1.47291, 1.21752, // TATTTA
    94.51683,  1.92511, 1.36919, 1.09122, // TATTTC
    96.22989,  1.92511, 1.40760, 1.13745, // TATTTG
    96.29356,  1.92511, 1.46191, 1.20391, // TATTTT
    91.70968,  2.70987, 1.43588, 1.01957, // TCAAAA
    88.58592,  2.70987, 1.70326, 1.31723, // TCAAAC
    90.91896,  2.70987, 1.58622, 1.18383, // TCAAAG
    89.22599,  2.70987, 1.70043, 1.31395, // TCAAAT
    82.67843,  2.16112, 1.59090, 1.30606, // TCAACA
    81.43304,  2.16112, 1.49398, 1.18854, // TCAACC
    82.71295,  2.16112, 1.53763, 1.24101, // TCAACG
    81.97429,  2.16112, 1.38415, 1.05992, // TCAACT
    90.18497,  2.81796, 1.46312, 1.08766, // TCAAGA
    87.54521,  2.81796, 1.63134, 1.28054, // TCAAGC
    88.91764,  2.81796, 1.59201, 1.23451, // TCAAGG
    88.03776,  2.81796, 1.76757, 1.44425, // TCAAGT
    81.44014,  2.24890, 1.53942, 1.26377, // TCAATA
    79.99393,  2.24890, 1.90423, 1.73865, // TCAATC
    81.07820,  2.24890, 1.65661, 1.41079, // TCAATG
    80.36772,  2.24890, 1.86789, 1.68912, // TCAATT
    93.98041,  1.22481, 1.02920, 0.77323, // TCACAA
    91.62725,  1.22481, 0.99209, 0.73178, // TCACAC
    93.32495,  1.22481, 0.98633, 0.72542, // TCACAG
    92.65333,  1.22481, 0.93521, 0.66976, // TCACAT
    90.90121,  1.29044, 1.13735, 0.90122, // TCACCA
    89.55020,  1.29044, 1.14805, 0.91396, // TCACCC
    90.72044,  1.29044, 1.08032, 0.83429, // TCACCG
    90.36226,  1.29044, 1.07550, 0.82870, // TCACCT
    93.17081,  1.39798, 0.96943, 0.69524, // TCACGA
    90.48583,  1.39798, 1.03690, 0.76906, // TCACGC
    92.14853,  1.39798, 0.98158, 0.70835, // TCACGG
    91.40069,  1.39798, 0.98509, 0.71215, // TCACGT
    89.22172,  1.56257, 1.18511, 0.89144, // TCACTA
    87.18163,  1.56257, 1.11422, 0.81266, // TCACTC
    88.60986,  1.56257, 1.13884, 0.83975, // TCACTG
    88.14816,  1.56257, 1.08670, 0.78275, // TCACTT
    87.60585,  2.60118, 1.66860, 1.29019, // TCAGAA
    82.21065,  2.60118, 1.96494, 1.64873, // TCAGAC
    86.58014,  2.60118, 1.67937, 1.30271, // TCAGAG
    84.01977,  2.60118, 1.88126, 1.54453, // TCAGAT
    82.38516,  2.08236, 1.37661, 1.13272, // TCAGCA
    80.72782,  2.08236, 1.41674, 1.18261, // TCAGCC
    82.04578,  2.08236, 1.35170, 1.10212, // TCAGCG
    82.14005,  2.08236, 1.57461, 1.38569, // TCAGCT
    86.99697,  2.50147, 1.58055, 1.25608, // TCAGGA
    83.22526,  2.50147, 1.71773, 1.42310, // TCAGGC
    85.81632,  2.50147, 1.56340, 1.23569, // TCAGGG
    84.53612,  2.50147, 1.84652, 1.58611, // TCAGGT
    79.60826,  2.40625, 1.73029, 1.48672, // TCAGTA
    76.59953,  2.40625, 1.93601, 1.75959, // TCAGTC
    78.46030,  2.40625, 1.81795, 1.60112, // TCAGTG
    77.81209,  2.40625, 2.01495, 1.86830, // TCAGTT
    87.86733,  1.95200, 1.51373, 1.18062, // TCATAA
    84.60316,  1.95200, 1.65213, 1.34619, // TCATAC
    87.19821,  1.95200, 1.49017, 1.15317, // TCATAG
    86.16081,  1.95200, 1.61150, 1.29684, // TCATAT
    81.20019,  2.22932, 2.02189, 1.98592, // TCATCA
    79.80195,  2.22932, 2.06742, 2.05337, // TCATCC
    81.17764,  2.22932, 2.02140, 1.98520, // TCATCG
    80.75664,  2.22932, 2.05581, 2.03610, // TCATCT
    85.53164,  2.18506, 1.58623, 1.26436, // TCATGA
    82.02848,  2.18506, 1.84311, 1.58361, // TCATGC
    84.57131,  2.18506, 1.69250, 1.39352, // TCATGG
    83.03519,  2.18506, 1.80777, 1.53829, // TCATGT
    75.80826,  2.43089, 2.13063, 2.00603, // TCATTA
    72.81026,  2.43089, 2.16581, 2.05592, // TCATTC
    74.96353,  2.43089, 2.10415, 1.96875, // TCATTG
    74.16296,  2.43089, 1.99140, 1.81264, // TCATTT
    102.62817, 1.63085, 1.14508, 0.90955, // TCCAAA
    100.44174, 1.63085, 1.22738, 1.00934, // TCCAAC
    101.80829, 1.63085, 1.18129, 0.95302, // TCCAAG
    101.03114, 1.63085, 1.36851, 1.18834, // TCCAAT
    95.11154,  1.46322, 0.92952, 0.62011, // TCCACA
    94.13069,  1.46322, 0.99983, 0.69179, // TCCACC
    95.24184,  1.46322, 0.93561, 0.62621, // TCCACG
    94.98765,  1.46322, 1.05312, 0.74782, // TCCACT
    100.28665, 1.84203, 1.23013, 0.95406, // TCCAGA
    98.09435,  1.84203, 1.29051, 1.02517, // TCCAGC
    99.23889,  1.84203, 1.18560, 0.90272, // TCCAGG
    98.71456,  1.84203, 1.39697, 1.15459, // TCCAGT
    94.08304,  1.79927, 1.13987, 0.78643, // TCCATA
    92.27877,  1.79927, 1.38479, 1.05307, // TCCATC
    93.68181,  1.79927, 1.25905, 0.91294, // TCCATG
    93.26788,  1.79927, 1.49766, 1.18440, // TCCATT
    101.82450, 1.44981, 0.94407, 0.68904, // TCCCAA
    99.50680,  1.44981, 0.89127, 0.63205, // TCCCAC
    101.45114, 1.44981, 0.88766, 0.62822, // TCCCAG
    100.64840, 1.44981, 0.95060, 0.69620, // TCCCAT
    98.30203,  1.40182, 0.91854, 0.67432, // TCCCCA
    96.96535,  1.40182, 0.84469, 0.59465, // TCCCCC
    98.11396,  1.40182, 0.88836, 0.64135, // TCCCCG
    97.74620,  1.40182, 0.84009, 0.58980, // TCCCCT
    100.83899, 1.43192, 0.94518, 0.70247, // TCCCGA
    98.31509,  1.43192, 0.91553, 0.66967, // TCCCGC
    99.50875,  1.43192, 0.89965, 0.65232, // TCCCGG
    99.14404,  1.43192, 0.99482, 0.75853, // TCCCGT
    96.69544,  1.44710, 0.85599, 0.60355, // TCCCTA
    95.44964,  1.44710, 1.01579, 0.78022, // TCCCTC
    96.41916,  1.44710, 0.93000, 0.68349, // TCCCTG
    95.99334,  1.44710, 1.12930, 0.91459, // TCCCTT
    97.51656,  1.82025, 1.21415, 0.93792, // TCCGAA
    93.25948,  1.82025, 1.43472, 1.20477, // TCCGAC
    96.37002,  1.82025, 1.22872, 0.95485, // TCCGAG
    94.97409,  1.82025, 1.56238, 1.36910, // TCCGAT
    93.35043,  1.58093, 1.01241, 0.67503, // TCCGCA
    91.96959,  1.58093, 1.09859, 0.76303, // TCCGCC
    93.21119,  1.58093, 1.05052, 0.71350, // TCCGCG
    93.01557,  1.58093, 1.12798, 0.79385, // TCCGCT
    97.46855,  1.74812, 1.23405, 0.99641, // TCCGGA
    94.37234,  1.74812, 1.32324, 1.10636, // TCCGGC
    96.33818,  1.74812, 1.19036, 0.94396, // TCCGGG
    95.72074,  1.74812, 1.44435, 1.26168, // TCCGGT
    91.55007,  2.13467, 1.30797, 1.03330, // TCCGTA
    89.30436,  2.13467, 1.56665, 1.35454, // TCCGTC
    90.85081,  2.13467, 1.41791, 1.16630, // TCCGTG
    90.75122,  2.13467, 1.64165, 1.45297, // TCCGTT
    97.10615,  1.77408, 1.31017, 1.02998, // TCCTAA
    94.12696,  1.77408, 1.18901, 0.89046, // TCCTAC
    96.29827,  1.77408, 1.01092, 0.69809, // TCCTAG
    95.61737,  1.77408, 1.22704, 0.93353, // TCCTAT
    90.37477,  2.16210, 1.63649, 1.45329, // TCCTCA
    89.29270,  2.16210, 1.64251, 1.46131, // TCCTCC
    90.30381,  2.16210, 1.52233, 1.30390, // TCCTCG
    90.34597,  2.16210, 1.70452, 1.54485, // TCCTCT
    94.34019,  1.63830, 1.18747, 0.89481, // TCCTGA
    91.09617,  1.63830, 1.34437, 1.07789, // TCCTGC
    93.07600,  1.63830, 1.25918, 0.97708, // TCCTGG
    92.30254,  1.63830, 1.31664, 1.04472, // TCCTGT
    85.84151,  1.91311, 1.65267, 1.40430, // TCCTTA
    82.88176,  1.91311, 1.65349, 1.40534, // TCCTTC
    85.12990,  1.91311, 1.56571, 1.29493, // TCCTTG
    84.11910,  1.91311, 1.51485, 1.23236, // TCCTTT
    84.52575,  2.36795, 1.53673, 1.13557, // TCGAAA
    81.42666,  2.36795, 1.83106, 1.47697, // TCGAAC
    83.50028,  2.36795, 1.60841, 1.21594, // TCGAAG
    81.96874,  2.36795, 1.67753, 1.29516, // TCGAAT
    74.69163,  2.00332, 1.79145, 1.52583, // TCGACA
    73.60515,  2.00332, 1.70497, 1.41669, // TCGACC
    74.51726,  2.00332, 1.67714, 1.38214, // TCGACG
    74.56388,  2.00332, 1.56856, 1.25012, // TCGACT
    82.97938,  2.41291, 1.48687, 1.08431, // TCGAGA
    80.81811,  2.41291, 1.66655, 1.28669, // TCGAGC
    81.94141,  2.41291, 1.55053, 1.15469, // TCGAGG
    81.26791,  2.41291, 1.80749, 1.45330, // TCGAGT
    73.80137,  1.85855, 1.54114, 1.39275, // TCGATA
    71.59896,  1.85855, 1.68551, 1.59298, // TCGATC
    72.65103,  1.85855, 1.52823, 1.37528, // TCGATG
    72.81994,  1.85855, 1.85325, 1.83658, // TCGATT
    89.46263,  1.44291, 1.24637, 0.97386, // TCGCAA
    86.82018,  1.44291, 1.18769, 0.90590, // TCGCAC
    88.39232,  1.44291, 1.21353, 0.93562, // TCGCAG
    87.83583,  1.44291, 1.13777, 0.84940, // TCGCAT
    85.40644,  1.52741, 1.43737, 1.21456, // TCGCCA
    84.12684,  1.52741, 1.43097, 1.20645, // TCGCCC
    85.29427,  1.52741, 1.44072, 1.21880, // TCGCCG
    85.07948,  1.52741, 1.40622, 1.17529, // TCGCCT
    87.38775,  1.58181, 1.26945, 0.96613, // TCGCGA
    84.28562,  1.58181, 1.36821, 1.08105, // TCGCGC
    86.12401,  1.58181, 1.37605, 1.09035, // TCGCGG
    84.90946,  1.58181, 1.36077, 1.07224, // TCGCGT
    83.67479,  1.56040, 1.45030, 1.16127, // TCGCTA
    81.59317,  1.56040, 1.45632, 1.16850, // TCGCTC
    82.99140,  1.56040, 1.40957, 1.11270, // TCGCTG
    82.35617,  1.56040, 1.17232, 0.84395, // TCGCTT
    84.06893,  2.44172, 1.73572, 1.33781, // TCGGAA
    78.74478,  2.44172, 2.09350, 1.77209, // TCGGAC
    82.70516,  2.44172, 1.73467, 1.33660, // TCGGAG
    80.53028,  2.44172, 1.98971, 1.64196, // TCGGAT
    79.09616,  2.03186, 1.51282, 1.23271, // TCGGCA
    77.93863,  2.03186, 1.50341, 1.22124, // TCGGCC
    78.93172,  2.03186, 1.45101, 1.15794, // TCGGCG
    78.52940,  2.03186, 1.41477, 1.11484, // TCGGCT
    84.47601,  2.35270, 1.68298, 1.26843, // TCGGGA
    81.63329,  2.35270, 1.78368, 1.38397, // TCGGGC
    83.41569,  2.35270, 1.56947, 1.14230, // TCGGGG
    82.49225,  2.35270, 1.86532, 1.48006, // TCGGGT
    77.59504,  2.46096, 1.81453, 1.52542, // TCGGTA
    74.95520,  2.46096, 2.02466, 1.79792, // TCGGTC
    76.36677,  2.46096, 1.82725, 1.54148, // TCGGTG
    75.87329,  2.46096, 1.98587, 1.74650, // TCGGTT
    81.22164,  2.14468, 1.78346, 1.54000, // TCGTAA
    77.85720,  2.14468, 1.94772, 1.75760, // TCGTAC
    80.16911,  2.14468, 1.69986, 1.43301, // TCGTAG
    79.44127,  2.14468, 1.95215, 1.76359, // TCGTAT
    74.16061,  2.49565, 2.37511, 2.68095, // TCGTCA
    72.91151,  2.49565, 2.47830, 2.85755, // TCGTCC
    74.23101,  2.49565, 2.34246, 2.62586, // TCGTCG
    73.97536,  2.49565, 2.42103, 2.75907, // TCGTCT
    77.71739,  2.22005, 1.84336, 1.69153, // TCGTGA
    73.81925,  2.22005, 2.08535, 2.03532, // TCGTGC
    76.57344,  2.22005, 1.94735, 1.83666, // TCGTGG
    75.09615,  2.22005, 2.15204, 2.13374, // TCGTGT
    68.89922,  2.47313, 2.41023, 2.51988, // TCGTTA
    65.62652,  2.47313, 2.41447, 2.52653, // TCGTTC
    68.08385,  2.47313, 2.25506, 2.28049, // TCGTTG
    67.06562,  2.47313, 2.20843, 2.21012, // TCGTTT
    109.32029, 2.64098, 1.90571, 1.48123, // TCTAAA
    106.60539, 2.64098, 1.88766, 1.46023, // TCTAAC
    108.75147, 2.64098, 1.85287, 1.42006, // TCTAAG
    108.48686, 2.64098, 2.16352, 1.79176, // TCTAAT
    104.77799, 2.58253, 1.59639, 1.13251, // TCTACA
    104.20514, 2.58253, 1.73020, 1.27785, // TCTACC
    104.39760, 2.58253, 1.66446, 1.20572, // TCTACG
    105.45799, 2.58253, 1.73492, 1.28309, // TCTACT
    106.82774, 2.51606, 1.69776, 1.38057, // TCTAGA
    104.75727, 2.51606, 1.76755, 1.46657, // TCTAGC
    106.47783, 2.51606, 1.85961, 1.58263, // TCTAGG
    106.36920, 2.51606, 1.93203, 1.67597, // TCTAGT
    104.99840, 3.03892, 1.92449, 1.41664, // TCTATA
    104.80833, 3.03892, 2.07253, 1.58320, // TCTATC
    104.25745, 3.03892, 1.99449, 1.49462, // TCTATG
    106.72082, 3.03892, 2.32739, 1.88403, // TCTATT
    115.10708, 2.42793, 1.92898, 1.73787, // TCTCAA
    112.48690, 2.42793, 1.76759, 1.52440, // TCTCAC
    114.36494, 2.42793, 2.00453, 1.84096, // TCTCAG
    113.64446, 2.42793, 2.05150, 1.90605, // TCTCAT
    113.59854, 2.47041, 1.87198, 1.57854, // TCTCCA
    111.98342, 2.47041, 1.80394, 1.49325, // TCTCCC
    113.65360, 2.47041, 1.82312, 1.51713, // TCTCCG
    113.06756, 2.47041, 1.90487, 1.62032, // TCTCCT
    114.68038, 2.49891, 1.87670, 1.58627, // TCTCGA
    111.67784, 2.49891, 1.85072, 1.55344, // TCTCGC
    113.78234, 2.49891, 1.93550, 1.66140, // TCTCGG
    113.60303, 2.49891, 1.90566, 1.62312, // TCTCGT
    113.11863, 2.72204, 2.04049, 1.78241, // TCTCTA
    111.81451, 2.72204, 2.10076, 1.86196, // TCTCTC
    112.89933, 2.72204, 1.96921, 1.68984, // TCTCTG
    113.03027, 2.72204, 2.16615, 1.94957, // TCTCTT
    105.08194, 2.50593, 1.86261, 1.48883, // TCTGAA
    101.22679, 2.50593, 1.87597, 1.50487, // TCTGAC
    104.20780, 2.50593, 1.70527, 1.30422, // TCTGAG
    103.46960, 2.50593, 1.98152, 1.63365, // TCTGAT
    103.06920, 2.56584, 1.68340, 1.21881, // TCTGCA
    102.21203, 2.56584, 1.75431, 1.29664, // TCTGCC
    102.86303, 2.56584, 1.70330, 1.24049, // TCTGCG
    103.81452, 2.56584, 1.89121, 1.45133, // TCTGCT
    105.25017, 2.47346, 1.90217, 1.55746, // TCTGGA
    102.57546, 2.47346, 1.78469, 1.41543, // TCTGGC
    104.31067, 2.47346, 1.81145, 1.44738, // TCTGGG
    104.43064, 2.47346, 1.98929, 1.66567, // TCTGGT
    103.35835, 3.11250, 2.03625, 1.53311, // TCTGTA
    103.11007, 3.11250, 2.05164, 1.55053, // TCTGTC
    103.08028, 3.11250, 2.01599, 1.51029, // TCTGTG
    105.41488, 3.11250, 2.31698, 1.86084, // TCTGTT
    104.06920, 1.86007, 1.40660, 1.05066, // TCTTAA
    100.07019, 1.86007, 1.30296, 0.93671, // TCTTAC
    103.34857, 1.86007, 1.54052, 1.20423, // TCTTAG
    101.92415, 1.86007, 1.37565, 1.01618, // TCTTAT
    98.36303,  1.79286, 1.49806, 1.17202, // TCTTCA
    96.34539,  1.79286, 1.46923, 1.13834, // TCTTCC
    98.13657,  1.79286, 1.50774, 1.18340, // TCTTCG
    97.92099,  1.79286, 1.46292, 1.13102, // TCTTCT
    102.01414, 1.88803, 1.38506, 1.07592, // TCTTGA
    98.05287,  1.88803, 1.29226, 0.96962, // TCTTGC
    100.87473, 1.88803, 1.38715, 1.07836, // TCTTGG
    99.77419,  1.88803, 1.39549, 1.08811, // TCTTGT
    94.54996,  1.89728, 1.32606, 0.94060, // TCTTTA
    92.29034,  1.89728, 1.25227, 0.86319, // TCTTTC
    93.92674,  1.89728, 1.28265, 0.89479, // TCTTTG
    94.35495,  1.89728, 1.34124, 0.95681, // TCTTTT
    79.16058,  1.67748, 1.18047, 0.81748, // TGAAAA
    76.73379,  1.67748, 1.27795, 0.92080, // TGAAAC
    77.87040,  1.67748, 1.19847, 0.83625, // TGAAAG
    77.42006,  1.67748, 1.24510, 0.88552, // TGAAAT
    68.18430,  2.18144, 1.89381, 1.63217, // TGAACA
    67.45056,  2.18144, 1.89881, 1.63864, // TGAACC
    68.41971,  2.18144, 1.80396, 1.51740, // TGAACG
    68.26325,  2.18144, 1.71431, 1.40571, // TGAACT
    76.42236,  1.59541, 1.25480, 0.90953, // TGAAGA
    75.18910,  1.59541, 1.17744, 0.82673, // TGAAGC
    75.59541,  1.59541, 1.27716, 0.93394, // TGAAGG
    75.22149,  1.59541, 1.18127, 0.83076, // TGAAGT
    67.70199,  1.95147, 1.77525, 1.50043, // TGAATA
    65.97852,  1.95147, 1.63087, 1.32115, // TGAATC
    67.09391,  1.95147, 1.70060, 1.40678, // TGAATG
    66.67355,  1.95147, 1.42776, 1.08220, // TGAATT
    84.45728,  2.48599, 1.58686, 1.38183, // TGACAA
    82.89401,  2.48599, 1.40247, 1.14812, // TGACAC
    83.52969,  2.48599, 1.55040, 1.33448, // TGACAG
    83.70919,  2.48599, 1.45090, 1.20810, // TGACAT
    80.63030,  2.19720, 1.63281, 1.53465, // TGACCA
    79.61343,  2.19720, 1.52625, 1.38690, // TGACCC
    80.61523,  2.19720, 1.64086, 1.54602, // TGACCG
    80.51824,  2.19720, 1.51042, 1.36537, // TGACCT
    83.24167,  1.98191, 1.43135, 1.25429, // TGACGA
    81.70218,  1.98191, 1.33075, 1.12440, // TGACGC
    82.24404,  1.98191, 1.39807, 1.21080, // TGACGG
    82.12770,  1.98191, 1.33680, 1.13208, // TGACGT
    80.00070,  1.92945, 1.47058, 1.35376, // TGACTA
    78.44945,  1.92945, 1.24365, 1.05283, // TGACTC
    79.51767,  1.92945, 1.42755, 1.29479, // TGACTG
    78.86855,  1.92945, 1.11819, 0.89760, // TGACTT
    79.42930,  1.41776, 1.10036, 0.81465, // TGAGAA
    75.34649,  1.41776, 1.41715, 1.19066, // TGAGAC
    78.55391,  1.41776, 1.12934, 0.84704, // TGAGAG
    76.42423,  1.41776, 1.34358, 1.09916, // TGAGAT
    72.50338,  1.59099, 1.42543, 1.08898, // TGAGCA
    71.18738,  1.59099, 1.53487, 1.21677, // TGAGCC
    72.56386,  1.59099, 1.43324, 1.09794, // TGAGCG
    72.27828,  1.59099, 1.34354, 0.99650, // TGAGCT
    78.56913,  1.35821, 1.03248, 0.72259, // TGAGGA
    76.43449,  1.35821, 1.15865, 0.85900, // TGAGGC
    77.75268,  1.35821, 1.02673, 0.71655, // TGAGGG
    77.12340,  1.35821, 1.13360, 0.83129, // TGAGGT
    71.93143,  1.45706, 1.19061, 0.89021, // TGAGTA
    70.03049,  1.45706, 1.33381, 1.05556, // TGAGTC
    71.31526,  1.45706, 1.16663, 0.86346, // TGAGTG
    71.18597,  1.45706, 1.43684, 1.18019, // TGAGTT
    77.03777,  2.58682, 1.87593, 1.80739, // TGATAA
    74.78199,  2.58682, 1.78715, 1.68061, // TGATAC
    76.17129,  2.58682, 1.87250, 1.80244, // TGATAG
    75.86323,  2.58682, 1.82603, 1.73576, // TGATAT
    70.39360,  2.70241, 2.03025, 2.24980, // TGATCA
    69.21458,  2.70241, 2.09047, 2.35063, // TGATCC
    70.47746,  2.70241, 1.95333, 2.12317, // TGATCG
    70.28094,  2.70241, 2.09227, 2.35367, // TGATCT
    74.11821,  2.05805, 1.66699, 1.67395, // TGATGA
    71.66881,  2.05805, 1.68484, 1.70092, // TGATGC
    73.20023,  2.05805, 1.65160, 1.65082, // TGATGG
    72.48205,  2.05805, 1.69763, 1.72032, // TGATGT
    65.87358,  2.35300, 2.03751, 2.12036, // TGATTA
    63.30257,  2.35300, 2.22321, 2.41674, // TGATTC
    65.28221,  2.35300, 2.05967, 2.15505, // TGATTG
    64.46161,  2.35300, 1.95543, 1.99353, // TGATTT
    103.69726, 2.21768, 1.52460, 1.15776, // TGCAAA
    100.67031, 2.21768, 1.62311, 1.27177, // TGCAAC
    102.64072, 2.21768, 1.60748, 1.25345, // TGCAAG
    101.97483, 2.21768, 1.70050, 1.36380, // TGCAAT
    94.60938,  1.72978, 1.06279, 0.68990, // TGCACA
    93.25694,  1.72978, 1.06479, 0.69184, // TGCACC
    94.50801,  1.72978, 1.09940, 0.72586, // TGCACG
    94.31217,  1.72978, 1.16088, 0.78759, // TGCACT
    100.45631, 2.29394, 1.51303, 1.12544, // TGCAGA
    97.67679,  2.29394, 1.51167, 1.12393, // TGCAGC
    99.35155,  2.29394, 1.54424, 1.16044, // TGCAGG
    98.84938,  2.29394, 1.73508, 1.38208, // TGCAGT
    93.87780,  2.05386, 1.21205, 0.84944, // TGCATA
    92.33253,  2.05386, 1.29948, 0.94300, // TGCATC
    93.20118,  2.05386, 1.26809, 0.90904, // TGCATG
    93.47661,  2.05386, 1.49705, 1.16603, // TGCATT
    104.21886, 1.96521, 1.25242, 0.98997, // TGCCAA
    101.55918, 1.96521, 1.31381, 1.06364, // TGCCAC
    103.53122, 1.96521, 1.26742, 1.00781, // TGCCAG
    102.73140, 1.96521, 1.41788, 1.19249, // TGCCAT
    99.93724,  1.76337, 1.15762, 0.88965, // TGCCCA
    98.17657,  1.76337, 1.08568, 0.80802, // TGCCCC
    100.19984, 1.76337, 1.13432, 0.86291, // TGCCCG
    99.71909,  1.76337, 1.21594, 0.95771, // TGCCCT
    102.91038, 1.93464, 1.35872, 1.13721, // TGCCGA
    100.35901, 1.93464, 1.29023, 1.05231, // TGCCGC
    101.79792, 1.93464, 1.26996, 1.02761, // TGCCGG
    101.26978, 1.93464, 1.46483, 1.27299, // TGCCGT
    99.34717,  1.85738, 1.19395, 0.83816, // TGCCTA
    97.09324,  1.85738, 1.30217, 0.95465, // TGCCTC
    98.72299,  1.85738, 1.27628, 0.92632, // TGCCTG
    97.79432,  1.85738, 1.43884, 1.10883, // TGCCTT
    98.98564,  2.24827, 1.48495, 1.15016, // TGCGAA
    94.71091,  2.24827, 1.49414, 1.16085, // TGCGAC
    97.98421,  2.24827, 1.48191, 1.14662, // TGCGAG
    96.54693,  2.24827, 1.63063, 1.32350, // TGCGAT
    93.85392,  1.86985, 1.15731, 0.78509, // TGCGCA
    92.42639,  1.86985, 1.20773, 0.83695, // TGCGCC
    93.90451,  1.86985, 1.21255, 0.84197, // TGCGCG
    93.84689,  1.86985, 1.31139, 0.94698, // TGCGCT
    98.73246,  2.16392, 1.59790, 1.30532, // TGCGGA
    95.70381,  2.16392, 1.42767, 1.10239, // TGCGGC
    97.74447,  2.16392, 1.50435, 1.19238, // TGCGGG
    97.29791,  2.16392, 1.61957, 1.33196, // TGCGGT
    93.23988,  2.46720, 1.40808, 1.04999, // TGCGTA
    91.44193,  2.46720, 1.51099, 1.16718, // TGCGTC
    92.65072,  2.46720, 1.50249, 1.15733, // TGCGTG
    93.15525,  2.46720, 1.75611, 1.46241, // TGCGTT
    97.86649,  1.80194, 1.44082, 1.09090, // TGCTAA
    94.52138,  1.80194, 1.24378, 0.87495, // TGCTAC
    96.91853,  1.80194, 1.51322, 1.17415, // TGCTAG
    95.85533,  1.80194, 1.39505, 1.03933, // TGCTAT
    89.78006,  2.01289, 1.58073, 1.30794, // TGCTCA
    88.19181,  2.01289, 1.67106, 1.42163, // TGCTCC
    89.88053,  2.01289, 1.58787, 1.31680, // TGCTCG
    89.67403,  2.01289, 1.65469, 1.40080, // TGCTCT
    94.97751,  1.67840, 1.29962, 0.95617, // TGCTGA
    91.47430,  1.67840, 1.22528, 0.87532, // TGCTGC
    93.61357,  1.67840, 1.31875, 0.97736, // TGCTGG
    92.41649,  1.67840, 1.30197, 0.95877, // TGCTGT
    85.30525,  1.51841, 1.25215, 0.92497, // TGCTTA
    82.54217,  1.51841, 1.24014, 0.91169, // TGCTTC
    84.67812,  1.51841, 1.22371, 0.89363, // TGCTTG
    83.90327,  1.51841, 1.09855, 0.76009, // TGCTTT
    73.55141,  1.83378, 1.40877, 0.97735, // TGGAAA
    71.09789,  1.83378, 1.69493, 1.28979, // TGGAAC
    72.13115,  1.83378, 1.44149, 1.01160, // TGGAAG
    71.53808,  1.83378, 1.70758, 1.30426, // TGGAAT
    61.89434,  2.32589, 2.04139, 1.68165, // TGGACA
    61.16762,  2.32589, 2.06003, 1.70473, // TGGACC
    62.05338,  2.32589, 2.06606, 1.71222, // TGGACG
    62.26475,  2.32589, 2.03403, 1.67256, // TGGACT
    71.36653,  1.86110, 1.42224, 1.06939, // TGGAGA
    70.13208,  1.86110, 1.53952, 1.20436, // TGGAGC
    70.37449,  1.86110, 1.50406, 1.16298, // TGGAGG
    70.25263,  1.86110, 1.49500, 1.15248, // TGGAGT
    62.04922,  1.92277, 1.87827, 1.61094, // TGGATA
    60.81068,  1.92277, 1.73381, 1.42871, // TGGATC
    61.54061,  1.92277, 1.78852, 1.49687, // TGGATG
    61.53769,  1.92277, 1.47549, 1.12162, // TGGATT
    84.29265,  1.89614, 1.36631, 1.09574, // TGGCAA
    81.95968,  1.89614, 1.15746, 0.85437, // TGGCAC
    83.09512,  1.89614, 1.28564, 1.00015, // TGGCAG
    82.58472,  1.89614, 1.15997, 0.85715, // TGGCAT
    80.00475,  1.71082, 1.36462, 1.12740, // TGGCCA
    78.87824,  1.71082, 1.33991, 1.09692, // TGGCCC
    79.94080,  1.71082, 1.32317, 1.07643, // TGGCCG
    79.80222,  1.71082, 1.25246, 0.99130, // TGGCCT
    82.44543,  1.57550, 1.22443, 0.94998, // TGGCGA
    80.33366,  1.57550, 1.13348, 0.84612, // TGGCGC
    81.37433,  1.57550, 1.22079, 0.94575, // TGGCGG
    80.74356,  1.57550, 1.14530, 0.85940, // TGGCGT
    78.93726,  1.47431, 1.21961, 0.92328, // TGGCTA
    77.21709,  1.47431, 1.19730, 0.89805, // TGGCTC
    78.45449,  1.47431, 1.16734, 0.86456, // TGGCTG
    78.07039,  1.47431, 1.00925, 0.69502, // TGGCTT
    76.73966,  1.70210, 1.38217, 1.01162, // TGGGAA
    72.19540,  1.70210, 1.59210, 1.25065, // TGGGAC
    75.54713,  1.70210, 1.48650, 1.12831, // TGGGAG
    73.51320,  1.70210, 1.65079, 1.32043, // TGGGAT
    70.30996,  1.76808, 1.69424, 1.34303, // TGGGCA
    69.01608,  1.76808, 1.66755, 1.31143, // TGGGCC
    70.17382,  1.76808, 1.59974, 1.23225, // TGGGCG
    70.03176,  1.76808, 1.49926, 1.11800, // TGGGCT
    76.92651,  1.51256, 1.22887, 0.90588, // TGGGGA
    74.66691,  1.51256, 1.38318, 1.08175, // TGGGGC
    76.00694,  1.51256, 1.26697, 0.94833, // TGGGGG
    75.31955,  1.51256, 1.38211, 1.08049, // TGGGGT
    69.55675,  1.63484, 1.30631, 1.00294, // TGGGTA
    67.89434,  1.63484, 1.59531, 1.35356, // TGGGTC
    68.85489,  1.63484, 1.41802, 1.13431, // TGGGTG
    68.95910,  1.63484, 1.64463, 1.41680, // TGGGTT
    75.25013,  2.38667, 1.71531, 1.49151, // TGGTAA
    71.83601,  2.38667, 1.81465, 1.62294, // TGGTAC
    74.26741,  2.38667, 1.64274, 1.39787, // TGGTAG
    73.42921,  2.38667, 1.76952, 1.56278, // TGGTAT
    67.63649,  2.69199, 2.37313, 2.72707, // TGGTCA
    66.22692,  2.69199, 2.30193, 2.60526, // TGGTCC
    67.88821,  2.69199, 2.28491, 2.57641, // TGGTCG
    67.32619,  2.69199, 2.31013, 2.61919, // TGGTCT
    71.51229,  2.13251, 1.79387, 1.71314, // TGGTGA
    67.99517,  2.13251, 2.03442, 2.06902, // TGGTGC
    70.39136,  2.13251, 1.91996, 1.89690, // TGGTGG
    69.08756,  2.13251, 2.01945, 2.04623, // TGGTGT
    63.21822,  2.62114, 2.45197, 2.53680, // TGGTTA
    60.21297,  2.62114, 2.36296, 2.39992, // TGGTTC
    62.49468,  2.62114, 2.30413, 2.31086, // TGGTTG
    61.49278,  2.62114, 2.22290, 2.18974, // TGGTTT
    105.81796, 2.96792, 2.15611, 1.62694, // TGTAAA
    103.23673, 2.96792, 2.21803, 1.69752, // TGTAAC
    105.04695, 2.96792, 2.10865, 1.57351, // TGTAAG
    105.07939, 2.96792, 2.35712, 1.85967, // TGTAAT
    99.12379,  2.66382, 1.83210, 1.31987, // TGTACA
    98.81876,  2.66382, 1.93741, 1.43529, // TGTACC
    99.07463,  2.66382, 1.82661, 1.31395, // TGTACG
    100.10449, 2.66382, 2.07384, 1.58954, // TGTACT
    103.08804, 2.77966, 1.99794, 1.53595, // TGTAGA
    101.21233, 2.77966, 2.03007, 1.57315, // TGTAGC
    102.70309, 2.77966, 2.01064, 1.55061, // TGTAGG
    103.43531, 2.77966, 2.18645, 1.75838, // TGTAGT
    100.30152, 3.77006, 2.06712, 1.49347, // TGTATA
    100.65660, 3.77006, 2.25349, 1.69993, // TGTATC
    99.49768,  3.77006, 2.19565, 1.63491, // TGTATG
    102.78049, 3.77006, 2.53528, 2.02856, // TGTATT
    114.53235, 2.78871, 2.21441, 2.12970, // TGTCAA
    112.17964, 2.78871, 2.15908, 2.05038, // TGTCAC
    113.62317, 2.78871, 2.14292, 2.02740, // TGTCAG
    113.09150, 2.78871, 2.33650, 2.30824, // TGTCAT
    112.56917, 2.92671, 2.11216, 1.97528, // TGTCCA
    110.56637, 2.92671, 2.19073, 2.08651, // TGTCCC
    112.28557, 2.92671, 2.17305, 2.06130, // TGTCCG
    111.43088, 2.92671, 2.34128, 2.30526, // TGTCCT
    113.67591, 2.90558, 2.34425, 2.29059, // TGTCGA
    110.99639, 2.90558, 2.16551, 2.03366, // TGTCGC
    112.91592, 2.90558, 2.21307, 2.10103, // TGTCGG
    112.86219, 2.90558, 2.29002, 2.21156, // TGTCGT
    112.76392, 3.02381, 2.27526, 2.08431, // TGTCTA
    111.37611, 3.02381, 2.30837, 2.12998, // TGTCTC
    112.32996, 3.02381, 2.32052, 2.14683, // TGTCTG
    112.45399, 3.02381, 2.40951, 2.27150, // TGTCTT
    102.82494, 2.86549, 2.06137, 1.70574, // TGTGAA
    99.45197,  2.86549, 2.10992, 1.76635, // TGTGAC
    102.56018, 2.86549, 1.73220, 1.31394, // TGTGAG
    101.37281, 2.86549, 2.31706, 2.03275, // TGTGAT
    100.72656, 2.92533, 1.95072, 1.54542, // TGTGCA
    100.15558, 2.92533, 1.96292, 1.55994, // TGTGCC
    100.71899, 2.92533, 2.02982, 1.64036, // TGTGCG
    102.26354, 2.92533, 2.17431, 1.81859, // TGTGCT
    103.64007, 2.85435, 2.19656, 1.89544, // TGTGGA
    101.36063, 2.85435, 1.99603, 1.64190, // TGTGGC
    102.56450, 2.85435, 2.08888, 1.75779, // TGTGGG
    103.31843, 2.85435, 2.22132, 1.92759, // TGTGGT
    102.65395, 3.38667, 2.31481, 1.92752, // TGTGTA
    103.24591, 3.38667, 2.33248, 1.94963, // TGTGTC
    102.64075, 3.38667, 2.32845, 1.94459, // TGTGTG
    105.52966, 3.38667, 2.75156, 2.49802, // TGTGTT
    102.18583, 2.40487, 1.78688, 1.66707, // TGTTAA
    98.46040,  2.40487, 1.58949, 1.39862, // TGTTAC
    101.35644, 2.40487, 1.73806, 1.59923, // TGTTAG
    100.01274, 2.40487, 1.68537, 1.52705, // TGTTAT
    95.75321,  2.21733, 1.83155, 1.58205, // TGTTCA
    93.92644,  2.21733, 1.75875, 1.48867, // TGTTCC
    95.90757,  2.21733, 1.82115, 1.56860, // TGTTCG
    95.68536,  2.21733, 1.77454, 1.50877, // TGTTCT
    99.82126,  2.36448, 1.68798, 1.46279, // TGTTGA
    96.01412,  2.36448, 1.58407, 1.32982, // TGTTGC
    98.65133,  2.36448, 1.62601, 1.38298, // TGTTGG
    97.41030,  2.36448, 1.64466, 1.40684, // TGTTGT
    92.11586,  1.91331, 1.54905, 1.28895, // TGTTTA
    89.61881,  1.91331, 1.43565, 1.15003, // TGTTTC
    91.58383,  1.91331, 1.48829, 1.21385, // TGTTTG
    91.23515,  1.91331, 1.38869, 1.09407, // TGTTTT
    92.78675,  2.42434, 1.39438, 1.02050, // TTAAAA
    90.21307,  2.42434, 1.70760, 1.38300, // TTAAAC
    91.77014,  2.42434, 1.43465, 1.06503, // TTAAAG
    90.87389,  2.42434, 1.76461, 1.45284, // TTAAAT
    83.71534,  2.30963, 1.63531, 1.38509, // TTAACA
    82.77787,  2.30963, 1.58021, 1.31568, // TTAACC
    83.65963,  2.30963, 1.59059, 1.32866, // TTAACG
    83.53102,  2.30963, 1.48891, 1.20331, // TTAACT
    91.20453,  2.47862, 1.47278, 1.08968, // TTAAGA
    89.13221,  2.47862, 1.56451, 1.19306, // TTAAGC
    90.04791,  2.47862, 1.49988, 1.11990, // TTAAGG
    89.52987,  2.47862, 1.72416, 1.38027, // TTAAGT
    82.78767,  2.47960, 1.59142, 1.33319, // TTAATA
    81.57276,  2.47960, 1.93782, 1.79136, // TTAATC
    82.20635,  2.47960, 1.66815, 1.43076, // TTAATG
    81.98709,  2.47960, 1.96643, 1.83118, // TTAATT
    94.46538,  1.32011, 1.11494, 0.83988, // TTACAA
    92.52684,  1.32011, 0.99692, 0.71012, // TTACAC
    93.55903,  1.32011, 1.04779, 0.76516, // TTACAG
    93.27573,  1.32011, 0.95727, 0.66818, // TTACAT
    91.44035,  1.41774, 1.21555, 0.90367, // TTACCA
    90.35008,  1.41774, 1.22845, 0.91810, // TTACCC
    91.38978,  1.41774, 1.19805, 0.88423, // TTACCG
    91.16695,  1.41774, 1.16287, 0.84557, // TTACCT
    93.41556,  1.34132, 1.04884, 0.72827, // TTACGA
    91.33059,  1.34132, 1.07550, 0.75621, // TTACGC
    92.39860,  1.34132, 1.08706, 0.76844, // TTACGG
    91.92480,  1.34132, 1.04011, 0.71920, // TTACGT
    90.43376,  1.52174, 1.17962, 0.89359, // TTACTA
    88.54356,  1.52174, 1.15048, 0.86069, // TTACTC
    89.77245,  1.52174, 1.18979, 0.90517, // TTACTG
    89.48426,  1.52174, 1.18924, 0.90455, // TTACTT
    90.45838,  2.32328, 1.49052, 1.18396, // TTAGAA
    86.13132,  2.32328, 1.99225, 1.82956, // TTAGAC
    89.19069,  2.32328, 1.58126, 1.29371, // TTAGAG
    87.55009,  2.32328, 1.86929, 1.66282, // TTAGAT
    85.28627,  2.28919, 1.38014, 1.14689, // TTAGCA
    84.23158,  2.28919, 1.57912, 1.40366, // TTAGCC
    85.23459,  2.28919, 1.47897, 1.27226, // TTAGCG
    85.25903,  2.28919, 1.56307, 1.38232, // TTAGCT
    90.20923,  2.34031, 1.52579, 1.21993, // TTAGGA
    87.13703,  2.34031, 1.67319, 1.40091, // TTAGGC
    88.92265,  2.34031, 1.46876, 1.15218, // TTAGGG
    88.02151,  2.34031, 1.71388, 1.45233, // TTAGGT
    83.78704,  2.72334, 1.70724, 1.52416, // TTAGTA
    81.05575,  2.72334, 2.00973, 1.94670, // TTAGTC
    82.60521,  2.72334, 1.73752, 1.56490, // TTAGTG
    82.46024,  2.72334, 2.05146, 2.00763, // TTAGTT
    89.11374,  1.88263, 1.52282, 1.29225, // TTATAA
    86.59217,  1.88263, 1.64616, 1.45238, // TTATAC
    88.72957,  1.88263, 1.51943, 1.28793, // TTATAG
    87.94378,  1.88263, 1.75459, 1.59822, // TTATAT
    84.08888,  2.46291, 2.16538, 2.18998, // TTATCA
    83.08700,  2.46291, 2.27198, 2.35368, // TTATCC
    84.01811,  2.46291, 2.18184, 2.21500, // TTATCG
    83.82846,  2.46291, 2.12140, 2.12361, // TTATCT
    87.17879,  2.11919, 1.59033, 1.32098, // TTATGA
    84.14010,  2.11919, 1.82441, 1.62312, // TTATGC
    86.11398,  2.11919, 1.73862, 1.51000, // TTATGG
    85.15860,  2.11919, 1.92228, 1.75547, // TTATGT
    79.29614,  2.69506, 2.25545, 2.19065, // TTATTA
    76.44361,  2.69506, 2.30636, 2.26525, // TTATTC
    78.37761,  2.69506, 2.26611, 2.20620, // TTATTG
    77.74862,  2.69506, 2.16530, 2.06063, // TTATTT
    102.34449, 2.08052, 1.28967, 0.97424, // TTCAAA
    100.08637, 2.08052, 1.46179, 1.17565, // TTCAAC
    101.18919, 2.08052, 1.42261, 1.12871, // TTCAAG
    101.06379, 2.08052, 1.55589, 1.29099, // TTCAAT
    95.57641,  2.10126, 1.20834, 0.85614, // TTCACA
    94.59384,  2.10126, 1.29014, 0.94453, // TTCACC
    95.51583,  2.10126, 1.22039, 0.86898, // TTCACG
    95.26049,  2.10126, 1.34364, 1.00389, // TTCACT
    100.53990, 2.40061, 1.44198, 1.13591, // TTCAGA
    98.35225,  2.40061, 1.57194, 1.29287, // TTCAGC
    99.36105,  2.40061, 1.46850, 1.16739, // TTCAGG
    98.93122,  2.40061, 1.67453, 1.42149, // TTCAGT
    94.35220,  2.47361, 1.44551, 1.12090, // TTCATA
    92.86572,  2.47361, 1.65532, 1.37359, // TTCATC
    93.75800,  2.47361, 1.51316, 1.20050, // TTCATG
    93.79265,  2.47361, 1.80943, 1.56980, // TTCATT
    101.38962, 1.63253, 0.97949, 0.70922, // TTCCAA
    98.99866,  1.63253, 1.10481, 0.84960, // TTCCAC
    100.68656, 1.63253, 0.99143, 0.72223, // TTCCAG
    99.94924,  1.63253, 1.15680, 0.91027, // TTCCAT
    97.98849,  1.60868, 1.00082, 0.73667, // TTCCCA
    96.46583,  1.60868, 0.98676, 0.72120, // TTCCCC
    97.87840,  1.60868, 1.00327, 0.73937, // TTCCCG
    97.55122,  1.60868, 1.05504, 0.79733, // TTCCCT
    100.00035, 1.82797, 1.11040, 0.80957, // TTCCGA
    97.99526,  1.82797, 1.17839, 0.88506, // TTCCGC
    99.34538,  1.82797, 1.08556, 0.78257, // TTCCGG
    98.63118,  1.82797, 1.26779, 0.98766, // TTCCGT
    96.73313,  1.91501, 1.09589, 0.81505, // TTCCTA
    95.07999,  1.91501, 1.34949, 1.11376, // TTCCTC
    96.51612,  1.91501, 1.21837, 0.95544, // TTCCTG
    95.77914,  1.91501, 1.42061, 1.20296, // TTCCTT
    99.02390,  2.59137, 1.51887, 1.23835, // TTCGAA
    94.97569,  2.59137, 1.76042, 1.54520, // TTCGAC
    97.67307,  2.59137, 1.59024, 1.32665, // TTCGAG
    96.41629,  2.59137, 1.88355, 1.71013, // TTCGAT
    94.51429,  2.48599, 1.35302, 1.04970, // TTCGCA
    93.07963,  2.48599, 1.45462, 1.17013, // TTCGCC
    94.08696,  2.48599, 1.39556, 1.09960, // TTCGCG
    93.96426,  2.48599, 1.52010, 1.25004, // TTCGCT
    98.44544,  2.51279, 1.64512, 1.37186, // TTCGGA
    95.81626,  2.51279, 1.60363, 1.32029, // TTCGGC
    97.57335,  2.51279, 1.46974, 1.15844, // TTCGGG
    97.04990,  2.51279, 1.72628, 1.47461, // TTCGGT
    91.48451,  5.74004, 1.73885, 1.52015, // TTCGTA
    91.65520,  5.74004, 1.74560, 1.52900, // TTCGTC
    92.95654,  5.74004, 1.81267, 1.61797, // TTCGTG
    92.10766,  5.74004, 2.00165, 1.87747, // TTCGTT
    97.10832,  1.86097, 1.46208, 1.09671, // TTCTAA
    94.00372,  1.86097, 1.52057, 1.16317, // TTCTAC
    96.64642,  1.86097, 1.69333, 1.36692, // TTCTAG
    95.13393,  1.86097, 1.48134, 1.11845, // TTCTAT
    91.23243,  2.32915, 1.87244, 1.74918, // TTCTCA
    89.66048,  2.32915, 1.91186, 1.80470, // TTCTCC
    91.15127,  2.32915, 1.85082, 1.71897, // TTCTCG
    90.90960,  2.32915, 1.93274, 1.83435, // TTCTCT
    94.76474,  2.03287, 1.45444, 1.15210, // TTCTGA
    91.56217,  2.03287, 1.60237, 1.33226, // TTCTGC
    93.37877,  2.03287, 1.49719, 1.20325, // TTCTGG
    92.50426,  2.03287, 1.60365, 1.33385, // TTCTGT
    86.80983,  2.37489, 1.93218, 1.77295, // TTCTTA
    83.75956,  2.37489, 1.92185, 1.75876, // TTCTTC
    86.31522,  2.37489, 1.93333, 1.77453, // TTCTTG
    85.03642,  2.37489, 1.77162, 1.55661, // TTCTTT
    85.41832,  3.07534, 1.70837, 1.26235, // TTGAAA
    82.20749,  3.07534, 1.97946, 1.57445, // TTGAAC
    84.02560,  3.07534, 1.75214, 1.31117, // TTGAAG
    83.23353,  3.07534, 2.07802, 1.69348, // TTGAAT
    75.99666,  2.53545, 1.78191, 1.50979, // TTGACA
    75.02045,  2.53545, 1.78324, 1.51149, // TTGACC
    75.77610,  2.53545, 1.80828, 1.54344, // TTGACG
    75.56733,  2.53545, 1.56589, 1.24375, // TTGACT
    84.33818,  3.17429, 1.76023, 1.39511, // TTGAGA
    82.52952,  3.17429, 1.91681, 1.58535, // TTGAGC
    83.61250,  3.17429, 1.70665, 1.33190, // TTGAGG
    82.82443,  3.17429, 1.92948, 1.60110, // TTGAGT
    74.84466,  2.69379, 1.68500, 1.40719, // TTGATA
    73.85601,  2.69379, 2.10707, 1.96776, // TTGATC
    74.25066,  2.69379, 1.84152, 1.60774, // TTGATG
    74.37427,  2.69379, 2.07510, 1.92315, // TTGATT
    91.18978,  1.48208, 1.19933, 0.92135, // TTGCAA
    89.00731,  1.48208, 1.08786, 0.79592, // TTGCAC
    90.19978,  1.48208, 1.19094, 0.91169, // TTGCAG
    89.62757,  1.48208, 1.09996, 0.80924, // TTGCAT
    87.82278,  1.66578, 1.39991, 1.11301, // TTGCCA
    86.62900,  1.66578, 1.38912, 1.10018, // TTGCCC
    87.69509,  1.66578, 1.39143, 1.10292, // TTGCCG
    87.31815,  1.66578, 1.29220, 0.98706, // TTGCCT
    89.79299,  1.67467, 1.18201, 0.82116, // TTGCGA
    87.41386,  1.67467, 1.22950, 0.87114, // TTGCGC
    88.74949,  1.67467, 1.26572, 0.90991, // TTGCGG
    87.78843,  1.67467, 1.20677, 0.84709, // TTGCGT
    86.58272,  1.83904, 1.37863, 1.12621, // TTGCTA
    84.70658,  1.83904, 1.32516, 1.06133, // TTGCTC
    85.87558,  1.83904, 1.35709, 1.09992, // TTGCTG
    85.47926,  1.83904, 1.34336, 1.08327, // TTGCTT
    87.41145,  3.22959, 1.68285, 1.37325, // TTGGAA
    82.78512,  3.22959, 2.13149, 1.95751, // TTGGAC
    85.74534,  3.22959, 1.93354, 1.69126, // TTGGAG
    84.17072,  3.22959, 2.15595, 1.99130, // TTGGAT
    82.15172,  2.83028, 1.50948, 1.19569, // TTGGCA
    80.93054,  2.83028, 1.64631, 1.36190, // TTGGCC
    82.03986,  2.83028, 1.54109, 1.23344, // TTGGCG
    82.24195,  2.83028, 1.71634, 1.44971, // TTGGCT
    87.30877,  2.98826, 1.73747, 1.38538, // TTGGGA
    84.96066,  2.98826, 1.81507, 1.47923, // TTGGGC
    86.39877,  2.98826, 1.67559, 1.31203, // TTGGGG
    85.96758,  2.98826, 1.91899, 1.60806, // TTGGGT
    81.48422,  3.42265, 1.88027, 1.50774, // TTGGTA
    79.31135,  3.42265, 2.06282, 1.73256, // TTGGTC
    80.33437,  3.42265, 2.01689, 1.67502, // TTGGTG
    80.50541,  3.42265, 2.15495, 1.84992, // TTGGTT
    84.86542,  2.22980, 1.73579, 1.41385, // TTGTAA
    81.69345,  2.22980, 1.92093, 1.64598, // TTGTAC
    83.84760,  2.22980, 1.89495, 1.61269, // TTGTAG
    82.95916,  2.22980, 1.82337, 1.52219, // TTGTAT
    79.60963,  2.83133, 2.40129, 2.31872, // TTGTCA
    78.39531,  2.83133, 2.38869, 2.30050, // TTGTCC
    79.43774,  2.83133, 2.31576, 2.19595, // TTGTCG
    79.25458,  2.83133, 2.42014, 2.34607, // TTGTCT
    82.20461,  2.65515, 1.94299, 1.67664, // TTGTGA
    78.63721,  2.65515, 2.11983, 1.91067, // TTGTGC
    80.81706,  2.65515, 1.98385, 1.72980, // TTGTGG
    79.72688,  2.65515, 2.08637, 1.86561, // TTGTGT
    74.82282,  3.04048, 2.43281, 2.37894, // TTGTTA
    71.58618,  3.04048, 2.49978, 2.47786, // TTGTTC
    73.96948,  3.04048, 2.41907, 2.35882, // TTGTTG
    72.98539,  3.04048, 2.38268, 2.30579, // TTGTTT
    104.92316, 2.16371, 1.36049, 1.02184, // TTTAAA
    102.67468, 2.16371, 1.43588, 1.10795, // TTTAAC
    103.84147, 2.16371, 1.34198, 1.00106, // TTTAAG
    104.15069, 2.16371, 1.55096, 1.24377, // TTTAAT
    99.94973,  2.01953, 1.16728, 0.78498, // TTTACA
    99.44129,  2.01953, 1.22834, 0.84737, // TTTACC
    99.73312,  2.01953, 1.19091, 0.80893, // TTTACG
    100.64758, 2.01953, 1.33058, 0.95534, // TTTACT
    103.09441, 2.04194, 1.31172, 0.97559, // TTTAGA
    101.18926, 2.04194, 1.34980, 1.01838, // TTTAGC
    102.01019, 2.04194, 1.26662, 0.92571, // TTTAGG
    102.31191, 2.04194, 1.48380, 1.17373, // TTTAGT
    99.74653,  2.50557, 1.39572, 1.03240, // TTTATA
    100.16592, 2.50557, 1.52283, 1.17660, // TTTATC
    99.66889,  2.50557, 1.46234, 1.10719, // TTTATG
    101.81814, 2.50557, 1.72667, 1.42058, // TTTATT
    109.11987, 2.19857, 1.45131, 1.14458, // TTTCAA
    107.13528, 2.19857, 1.44167, 1.13319, // TTTCAC
    108.47611, 2.19857, 1.45332, 1.14696, // TTTCAG
    108.09867, 2.19857, 1.57007, 1.28790, // TTTCAT
    107.34698, 2.17160, 1.41230, 1.08069, // TTTCCA
    105.84822, 2.17160, 1.34081, 0.99968, // TTTCCC
    107.20150, 2.17160, 1.42904, 1.09996, // TTTCCG
    106.76742, 2.17160, 1.47439, 1.15273, // TTTCCT
    108.32412, 2.22607, 1.55531, 1.27001, // TTTCGA
    106.18324, 2.22607, 1.45348, 1.14736, // TTTCGC
    107.63787, 2.22607, 1.44900, 1.14204, // TTTCGG
    107.27219, 2.22607, 1.57040, 1.28854, // TTTCGT
    107.27312, 2.26458, 1.58491, 1.31190, // TTTCTA
    106.52256, 2.26458, 1.56115, 1.28251, // TTTCTC
    107.21335, 2.26458, 1.56272, 1.28445, // TTTCTG
    107.29086, 2.26458, 1.80283, 1.59157, // TTTCTT
    101.14136, 2.11369, 1.37872, 1.13499, // TTTGAA
    98.13275,  2.11369, 1.45215, 1.22687, // TTTGAC
    100.38283, 2.11369, 1.40071, 1.16226, // TTTGAG
    99.79899,  2.11369, 1.50903, 1.29966, // TTTGAT
    98.65438,  2.06630, 1.19618, 0.84925, // TTTGCA
    98.00416,  2.06630, 1.26399, 0.92248, // TTTGCC
    98.73183,  2.06630, 1.23240, 0.88812, // TTTGCG
    99.53043,  2.06630, 1.34030, 1.00727, // TTTGCT
    101.43908, 1.98654, 1.34378, 1.08967, // TTTGGA
    99.26147,  1.98654, 1.34328, 1.08906, // TTTGGC
    100.65307, 1.98654, 1.26761, 0.99835, // TTTGGG
    100.94995, 1.98654, 1.42336, 1.18789, // TTTGGT
    99.30954,  2.51049, 1.53365, 1.20745, // TTTGTA
    99.36009,  2.51049, 1.56768, 1.24786, // TTTGTC
    99.33139,  2.51049, 1.53273, 1.20636, // TTTGTG
    100.98521, 2.51049, 1.78791, 1.51984, // TTTGTT
    99.56834,  1.70840, 1.15763, 0.83408, // TTTTAA
    96.25565,  1.70840, 1.02715, 0.69711, // TTTTAC
    98.77880,  1.70840, 1.11740, 0.79098, // TTTTAG
    97.57975,  1.70840, 1.08483, 0.75665, // TTTTAT
    94.53220,  1.56977, 1.21410, 0.93207, // TTTTCA
    92.87624,  1.56977, 1.18668, 0.90067, // TTTTCC
    94.46475,  1.56977, 1.20816, 0.92524, // TTTTCG
    94.19426,  1.56977, 1.23478, 0.95598, // TTTTCT
    97.77195,  1.62351, 1.07009, 0.74952, // TTTTGA
    94.38447,  1.62351, 1.00115, 0.67827, // TTTTGC
    96.81804,  1.62351, 1.08120, 0.76123, // TTTTGG
    95.53426,  1.62351, 1.06767, 0.74697, // TTTTGT
    91.07904,  1.51391, 1.03289, 0.70976, // TTTTTA
    89.17389,  1.51391, 1.00204, 0.67820, // TTTTTC
    90.68991,  1.51391, 1.00091, 0.67705, // TTTTTG
    90.67901,  1.51391, 0.90750, 0.58452  // TTTTTT
};

static float r9_4_450bps_cpg_6mer_template_model_builtin_data[] = {
    86.95960,  1.61061, 0.94148, 0.60936, // AAAAAA
    83.44650,  1.65993, 1.07705, 0.74561, // AAAAAC
    84.20150,  1.44083, 0.95343, 0.62100, // AAAAAG
    84.06040,  1.23283, 1.07705, 0.74561, // AAAAAM
    86.50580,  2.79715, 1.10608, 0.77595, // AAAAAT
    77.31630,  1.82344, 1.29686, 0.93621, // AAAACA
    75.37490,  1.66194, 1.31939, 0.96072, // AAAACC
    76.92820,  1.70501, 1.30207, 0.94186, // AAAACG
    76.35400,  1.66292, 1.31939, 0.96072, // AAAACM
    76.15580,  1.62634, 1.17932, 0.81186, // AAAACT
    83.85860,  1.28235, 0.93295, 0.60480, // AAAAGA
    83.29760,  1.40791, 0.96369, 0.63495, // AAAAGC
    83.92940,  1.10518, 0.93570, 0.60748, // AAAAGG
    82.98230,  0.99890, 0.96369, 0.63495, // AAAAGM
    82.41880,  1.58546, 1.07858, 0.75182, // AAAAGT
    77.09730,  2.70501, 1.29686, 0.93621, // AAAAMA
    75.70630,  2.70501, 1.31939, 0.96072, // AAAAMC
    80.68720,  1.37188, 1.30207, 0.94186, // AAAAMG
    75.70630,  2.70501, 1.31939, 0.96072, // AAAAMM
    76.81140,  2.70501, 1.17932, 0.81186, // AAAAMT
    76.70890,  1.65970, 1.14524, 0.84930, // AAAATA
    74.76450,  1.42923, 1.11274, 0.81340, // AAAATC
    75.05440,  1.57364, 1.09532, 0.79438, // AAAATG
    75.57200,  1.44628, 1.11274, 0.81340, // AAAATM
    76.51530,  1.90023, 1.28978, 1.01505, // AAAATT
    88.04510,  2.03674, 1.48065, 1.34391, // AAACAA
    86.45610,  1.81021, 1.27700, 1.07641, // AAACAC
    86.37200,  2.15651, 1.40026, 1.23597, // AAACAG
    87.11870,  1.97353, 1.27700, 1.07641, // AAACAM
    87.84140,  2.01169, 1.31446, 1.12412, // AAACAT
    83.65830,  2.12520, 1.49328, 1.42524, // AAACCA
    83.38350,  1.75518, 1.40063, 1.29469, // AAACCC
    84.50590,  1.83613, 1.45041, 1.36432, // AAACCG
    84.04540,  1.85250, 1.40063, 1.29469, // AAACCM
    84.26480,  1.83453, 1.38692, 1.27573, // AAACCT
    87.18450,  1.55453, 1.30654, 1.16200, // AAACGA
    85.63650,  1.55453, 1.21094, 1.03683, // AAACGC
    86.33690,  1.55453, 1.29411, 1.14546, // AAACGG
    85.63650,  2.55453, 1.21094, 1.03683, // AAACGM
    86.06470,  1.55453, 1.24344, 1.07884, // AAACGT
    84.58290,  2.83613, 1.49328, 1.42524, // AAACMA
    83.47830,  2.83613, 1.40063, 1.29469, // AAACMC
    84.58050,  1.56178, 1.45041, 1.36432, // AAACMG
    83.47830,  2.83613, 1.40063, 1.29469, // AAACMM
    84.41230,  2.83613, 1.38692, 1.27573, // AAACMT
    83.77380,  1.67705, 1.28615, 1.13271, // AAACTA
    81.99770,  1.43213, 1.12216, 0.92313, // AAACTC
    82.16530,  1.92527, 1.22077, 1.04745, // AAACTG
    82.95350,  1.59993, 1.12216, 0.92313, // AAACTM
    82.46180,  1.32164, 0.95274, 0.72217, // AAACTT
    85.38450,  1.35767, 0.97542, 0.66709, // AAAGAA
    80.42880,  1.43673, 1.17615, 0.88326, // AAAGAC
    83.24860,  1.34999, 1.00518, 0.69785, // AAAGAG
    81.36990,  1.62610, 1.17615, 0.88326, // AAAGAM
    81.33540,  1.44262, 1.16532, 0.87110, // AAAGAT
    78.61990,  1.37587, 1.04709, 0.74755, // AAAGCA
    77.80840,  1.42815, 1.04196, 0.74206, // AAAGCC
    78.87290,  1.42710, 0.98261, 0.67957, // AAAGCG
    78.12000,  1.36880, 1.04196, 0.74206, // AAAGCM
    78.58110,  1.26863, 0.93431, 0.63008, // AAAGCT
    83.19630,  1.32845, 0.98410, 0.67678, // AAAGGA
    80.85620,  1.26686, 1.02292, 0.71723, // AAAGGC
    82.51300,  1.19511, 0.93946, 0.63127, // AAAGGG
    81.01680,  1.35137, 1.02292, 0.71723, // AAAGGM
    82.36290,  1.35463, 1.07170, 0.76914, // AAAGGT
    79.14900,  2.42710, 1.04709, 0.74755, // AAAGMA
    77.56130,  2.42710, 1.04196, 0.74206, // AAAGMC
    80.82360,  1.24671, 0.98261, 0.67957, // AAAGMG
    77.56130,  2.42710, 1.04196, 0.74206, // AAAGMM
    78.67540,  2.42710, 0.93431, 0.63008, // AAAGMT
    77.81850,  1.26415, 1.00723, 0.72430, // AAAGTA
    75.23090,  1.55049, 1.20345, 0.94594, // AAAGTC
    76.21280,  1.44832, 1.07134, 0.79454, // AAAGTG
    76.40810,  1.45186, 1.20345, 0.94594, // AAAGTM
    75.92730,  1.69108, 1.30532, 1.06856, // AAAGTT
    88.25540,  2.96203, 1.48065, 1.34391, // AAAMAA
    86.68990,  2.96203, 1.27700, 1.07641, // AAAMAC
    87.32010,  2.96203, 1.40026, 1.23597, // AAAMAG
    86.68990,  2.96203, 1.27700, 1.07641, // AAAMAM
    87.45210,  2.96203, 1.31446, 1.12412, // AAAMAT
    84.58290,  2.83613, 1.49328, 1.42524, // AAAMCA
    83.47830,  2.83613, 1.40063, 1.29469, // AAAMCC
    84.50590,  2.83613, 1.45041, 1.36432, // AAAMCG
    83.47830,  2.83613, 1.40063, 1.29469, // AAAMCM
    84.41230,  2.83613, 1.38692, 1.27573, // AAAMCT
    85.58820,  1.36722, 1.30654, 1.16200, // AAAMGA
    83.73860,  1.16843, 1.21094, 1.03683, // AAAMGC
    85.00510,  1.36125, 1.29411, 1.14546, // AAAMGG
    84.31820,  1.22025, 1.21094, 1.03683, // AAAMGM
    84.02890,  1.22816, 1.24344, 1.07884, // AAAMGT
    84.58290,  2.83613, 1.49328, 1.42524, // AAAMMA
    83.47830,  2.83613, 1.40063, 1.29469, // AAAMMC
    84.50590,  2.83613, 1.45041, 1.36432, // AAAMMG
    83.47830,  2.83613, 1.40063, 1.29469, // AAAMMM
    84.41230,  2.83613, 1.38692, 1.27573, // AAAMMT
    84.13420,  2.58806, 1.28615, 1.13271, // AAAMTA
    82.01610,  2.58806, 1.12216, 0.92313, // AAAMTC
    83.27920,  2.58806, 1.22077, 1.04745, // AAAMTG
    82.01610,  2.58806, 1.12216, 0.92313, // AAAMTM
    82.58670,  2.58806, 0.95274, 0.72217, // AAAMTT
    83.42330,  1.95981, 1.64571, 1.60071, // AAATAA
    80.18000,  1.65854, 1.57619, 1.50036, // AAATAC
    81.81680,  1.61976, 1.53106, 1.43639, // AAATAG
    80.60410,  1.80502, 1.57619, 1.50036, // AAATAM
    80.86300,  1.88127, 1.71069, 1.69644, // AAATAT
    76.79700,  2.25572, 2.11887, 2.65176, // AAATCA
    74.75060,  2.24317, 2.03424, 2.49449, // AAATCC
    76.19820,  2.22770, 2.15565, 2.72110, // AAATCG
    75.58390,  2.41428, 2.03424, 2.49449, // AAATCM
    75.99540,  2.32932, 2.14848, 2.70753, // AAATCT
    79.50020,  1.69057, 1.56632, 1.53101, // AAATGA
    77.60980,  1.55530, 1.61261, 1.59937, // AAATGC
    78.79890,  1.57656, 1.64897, 1.65377, // AAATGG
    78.49260,  1.75191, 1.61261, 1.59937, // AAATGM
    78.45210,  1.70405, 1.66765, 1.68195, // AAATGT
    76.03340,  3.22770, 2.11887, 2.65176, // AAATMA
    74.91410,  3.22770, 2.03424, 2.49449, // AAATMC
    78.35020,  1.62251, 2.15565, 2.72110, // AAATMG
    74.91410,  3.22770, 2.03424, 2.49449, // AAATMM
    76.11480,  3.22770, 2.14848, 2.70753, // AAATMT
    70.41970,  1.97918, 1.67811, 1.58287, // AAATTA
    68.02980,  2.02744, 1.83641, 1.81206, // AAATTC
    69.80860,  2.01268, 1.71772, 1.63924, // AAATTG
    69.56110,  2.23687, 1.83641, 1.81206, // AAATTM
    68.93400,  1.96019, 1.73599, 1.66547, // AAATTT
    109.07600, 2.27967, 1.79213, 1.53404, // AACAAA
    104.75400, 2.17798, 1.80497, 1.55056, // AACAAC
    106.87400, 2.13321, 1.87422, 1.64064, // AACAAG
    106.06800, 1.99188, 1.80497, 1.55056, // AACAAM
    105.84600, 2.27770, 1.96452, 1.76063, // AACAAT
    99.78910,  1.69654, 1.36504, 1.00221, // AACACA
    98.80330,  1.77798, 1.33763, 0.97218, // AACACC
    99.53500,  1.74166, 1.32214, 0.95534, // AACACG
    99.16990,  1.79660, 1.33763, 0.97218, // AACACM
    99.67290,  1.92055, 1.43867, 1.08438, // AACACT
    105.73500, 2.21774, 1.92656, 1.69338, // AACAGA
    102.08600, 2.34423, 1.77407, 1.49637, // AACAGC
    103.91300, 2.23332, 1.74932, 1.46516, // AACAGG
    103.70800, 2.07030, 1.77407, 1.49637, // AACAGM
    103.54700, 2.45351, 1.92458, 1.69077, // AACAGT
    99.65940,  2.74166, 1.36504, 1.00221, // AACAMA
    98.13200,  2.74166, 1.33763, 0.97218, // AACAMC
    104.39800, 1.83486, 1.32214, 0.95534, // AACAMG
    98.13200,  2.74166, 1.33763, 0.97218, // AACAMM
    99.48970,  2.74166, 1.43867, 1.08438, // AACAMT
    99.38340,  1.91647, 1.55825, 1.29235, // AACATA
    98.24840,  2.30793, 1.61265, 1.36061, // AACATC
    98.32950,  2.00613, 1.58327, 1.32361, // AACATG
    98.84990,  2.08584, 1.61265, 1.36061, // AACATM
    99.52910,  2.44129, 1.84242, 1.66153, // AACATT
    107.76200, 1.97012, 1.53183, 1.35379, // AACCAA
    104.90200, 1.97916, 1.51285, 1.32871, // AACCAC
    105.90100, 2.31087, 1.48473, 1.29184, // AACCAG
    106.17100, 1.79461, 1.51285, 1.32871, // AACCAM
    107.06600, 2.07042, 1.65588, 1.52152, // AACCAT
    104.09900, 1.83181, 1.29734, 1.06459, // AACCCA
    102.13600, 1.64249, 1.24222, 0.99747, // AACCCC
    103.83500, 1.75332, 1.34040, 1.11804, // AACCCG
    103.02000, 1.72960, 1.24222, 0.99747, // AACCCM
    103.26000, 1.85650, 1.47108, 1.28546, // AACCCT
    106.59300, 1.90443, 1.60306, 1.39462, // AACCGA
    103.84800, 1.90443, 1.47383, 1.22942, // AACCGC
    105.63900, 1.90443, 1.54235, 1.31615, // AACCGG
    103.84800, 2.90443, 1.47383, 1.22942, // AACCGM
    104.87500, 1.90443, 1.68020, 1.49648, // AACCGT
    103.93400, 2.75332, 1.29734, 1.06459, // AACCMA
    102.23000, 2.75332, 1.24222, 0.99747, // AACCMC
    105.08000, 1.67520, 1.34040, 1.11804, // AACCMG
    102.23000, 2.75332, 1.24222, 0.99747, // AACCMM
    103.20300, 2.75332, 1.47108, 1.28546, // AACCMT
    103.13500, 1.84952, 1.49866, 1.19909, // AACCTA
    100.92900, 1.86761, 1.52449, 1.23022, // AACCTC
    103.02300, 1.92992, 1.48098, 1.17793, // AACCTG
    101.88200, 1.77188, 1.52449, 1.23022, // AACCTM
    102.04900, 1.98311, 1.69160, 1.43795, // AACCTT
    103.13000, 2.24722, 1.81869, 1.59373, // AACGAA
    98.44620,  2.24722, 1.72440, 1.47142, // AACGAC
    102.04100, 2.24722, 1.78587, 1.55078, // AACGAG
    98.44620,  3.24722, 1.72440, 1.47142, // AACGAM
    100.50100, 2.24722, 1.95330, 1.77391, // AACGAT
    98.63880,  2.02317, 1.43135, 1.10889, // AACGCA
    96.96740,  2.02317, 1.50183, 1.19180, // AACGCC
    98.42320,  2.02317, 1.53638, 1.23315, // AACGCG
    96.96740,  3.02317, 1.50183, 1.19180, // AACGCM
    98.63120,  2.02317, 1.58425, 1.29123, // AACGCT
    102.87800, 2.22668, 1.82532, 1.59411, // AACGGA
    99.63190,  2.22668, 1.69529, 1.42684, // AACGGC
    101.89900, 2.22668, 1.74356, 1.48821, // AACGGG
    99.63190,  3.22668, 1.69529, 1.42684, // AACGGM
    101.25300, 2.22668, 1.88906, 1.67833, // AACGGT
    98.63880,  3.02317, 1.43135, 1.10889, // AACGMA
    96.96740,  3.02317, 1.50183, 1.19180, // AACGMC
    98.42320,  3.02317, 1.53638, 1.23315, // AACGMG
    96.96740,  3.02317, 1.50183, 1.19180, // AACGMM
    98.63120,  3.02317, 1.58425, 1.29123, // AACGMT
    97.84680,  2.64556, 1.75347, 1.45050, // AACGTA
    96.07220,  2.64556, 1.79988, 1.50847, // AACGTC
    97.18590,  2.64556, 1.75924, 1.45768, // AACGTG
    96.07220,  3.64556, 1.79988, 1.50847, // AACGTM
    97.66840,  2.64556, 2.09559, 1.89511, // AACGTT
    107.94500, 2.88264, 1.53183, 1.35379, // AACMAA
    105.07200, 2.88264, 1.51285, 1.32871, // AACMAC
    107.00300, 2.88264, 1.48473, 1.29184, // AACMAG
    105.07200, 2.88264, 1.51285, 1.32871, // AACMAM
    106.26800, 2.88264, 1.65588, 1.52152, // AACMAT
    103.93400, 2.75332, 1.29734, 1.06459, // AACMCA
    102.23000, 2.75332, 1.24222, 0.99747, // AACMCC
    103.83500, 2.75332, 1.34040, 1.11804, // AACMCG
    102.23000, 2.75332, 1.24222, 0.99747, // AACMCM
    103.20300, 2.75332, 1.47108, 1.28546, // AACMCT
    102.77800, 1.89806, 1.60306, 1.39462, // AACMGA
    100.35400, 1.87824, 1.47383, 1.22942, // AACMGC
    101.86800, 1.75615, 1.54235, 1.31615, // AACMGG
    101.38100, 1.91946, 1.47383, 1.22942, // AACMGM
    101.47400, 1.99118, 1.68020, 1.49648, // AACMGT
    103.93400, 2.75332, 1.29734, 1.06459, // AACMMA
    102.23000, 2.75332, 1.24222, 0.99747, // AACMMC
    103.83500, 2.75332, 1.34040, 1.11804, // AACMMG
    102.23000, 2.75332, 1.24222, 0.99747, // AACMMM
    103.20300, 2.75332, 1.47108, 1.28546, // AACMMT
    103.12500, 2.83098, 1.49866, 1.19909, // AACMTA
    100.84400, 2.83098, 1.52449, 1.23022, // AACMTC
    102.21600, 2.83098, 1.48098, 1.17793, // AACMTG
    100.84400, 2.83098, 1.52449, 1.23022, // AACMTM
    101.84000, 2.83098, 1.69160, 1.43795, // AACMTT
    101.66000, 1.81097, 1.59479, 1.18964, // AACTAA
    98.16890,  1.52897, 1.40164, 0.98019, // AACTAC
    100.29800, 1.68146, 1.58045, 1.17362, // AACTAG
    99.25930,  1.70844, 1.40164, 0.98019, // AACTAM
    99.60620,  1.64007, 1.51928, 1.10615, // AACTAT
    93.38270,  1.74243, 1.57047, 1.41151, // AACTCA
    91.70640,  1.69783, 1.59310, 1.44213, // AACTCC
    93.52490,  1.86136, 1.54560, 1.37810, // AACTCG
    92.44330,  1.82188, 1.59310, 1.44213, // AACTCM
    93.09320,  1.74621, 1.57869, 1.42260, // AACTCT
    98.60640,  1.68200, 1.41332, 1.09402, // AACTGA
    95.81290,  1.60976, 1.32356, 0.99148, // AACTGC
    96.42190,  1.93059, 1.38744, 1.06412, // AACTGG
    96.49000,  1.52469, 1.32356, 0.99148, // AACTGM
    96.38030,  1.53204, 1.35980, 1.03248, // AACTGT
    93.66730,  2.86136, 1.57047, 1.41151, // AACTMA
    91.54690,  2.86136, 1.59310, 1.44213, // AACTMC
    96.37470,  1.59153, 1.54560, 1.37810, // AACTMG
    91.54690,  2.86136, 1.59310, 1.44213, // AACTMM
    93.05050,  2.86136, 1.57869, 1.42260, // AACTMT
    88.79880,  1.57141, 1.29167, 0.96950, // AACTTA
    85.98470,  1.33709, 1.18574, 0.85272, // AACTTC
    88.16340,  1.49123, 1.18719, 0.85428, // AACTTG
    87.14590,  1.41192, 1.18574, 0.85272, // AACTTM
    86.97300,  1.33328, 1.07811, 0.73929, // AACTTT
    79.98880,  1.76853, 1.12430, 0.71403, // AAGAAA
    76.80700,  1.80246, 1.35326, 0.94291, // AAGAAC
    79.79600,  1.88606, 1.13240, 0.72177, // AAGAAG
    77.15920,  1.48670, 1.35326, 0.94291, // AAGAAM
    77.22380,  2.05394, 1.34279, 0.93199, // AAGAAT
    68.79750,  2.08019, 1.64737, 1.23807, // AAGACA
    67.57860,  2.02281, 1.63794, 1.22745, // AAGACC
    68.61970,  2.14481, 1.66378, 1.25661, // AAGACG
    68.28560,  2.06766, 1.63794, 1.22745, // AAGACM
    68.46190,  1.97099, 1.59024, 1.17423, // AAGACT
    77.87670,  1.65310, 1.09014, 0.73888, // AAGAGA
    76.28590,  1.56443, 1.15687, 0.80774, // AAGAGC
    77.01280,  1.92961, 1.16133, 0.81242, // AAGAGG
    76.55340,  2.05465, 1.15687, 0.80774, // AAGAGM
    76.31460,  1.73124, 1.18627, 0.83874, // AAGAGT
    69.06000,  3.14481, 1.64737, 1.23807, // AAGAMA
    67.42400,  3.14481, 1.63794, 1.22745, // AAGAMC
    73.73740,  1.83066, 1.66378, 1.25661, // AAGAMG
    67.42400,  3.14481, 1.63794, 1.22745, // AAGAMM
    68.60190,  3.14481, 1.59024, 1.17423, // AAGAMT
    68.69620,  1.87700, 1.47689, 1.21422, // AAGATA
    67.24550,  1.64528, 1.34951, 1.06057, // AAGATC
    68.37180,  1.85826, 1.38770, 1.10590, // AAGATG
    68.09610,  1.80633, 1.34951, 1.06057, // AAGATM
    68.37610,  1.66211, 1.36842, 1.08293, // AAGATT
    86.75110,  1.97799, 1.47547, 1.22356, // AAGCAA
    84.52530,  1.80137, 1.20873, 0.90724, // AAGCAC
    85.42220,  1.87047, 1.40134, 1.13252, // AAGCAG
    85.28460,  1.94650, 1.20873, 0.90724, // AAGCAM
    85.34400,  1.83051, 1.26641, 0.97295, // AAGCAT
    82.16160,  1.75052, 1.43823, 1.25952, // AAGCCA
    81.23210,  1.56130, 1.39259, 1.20004, // AAGCCC
    82.45520,  1.70618, 1.44790, 1.27224, // AAGCCG
    82.13940,  1.73756, 1.39259, 1.20004, // AAGCCM
    82.28450,  1.70853, 1.41867, 1.23391, // AAGCCT
    84.88050,  1.56014, 1.28937, 1.05556, // AAGCGA
    81.78060,  1.56014, 1.19997, 0.94770, // AAGCGC
    83.72480,  1.56014, 1.31411, 1.08609, // AAGCGG
    81.78060,  2.56014, 1.19997, 0.94770, // AAGCGM
    82.51430,  1.56014, 1.27584, 1.03900, // AAGCGT
    82.42550,  2.70618, 1.43823, 1.25952, // AAGCMA
    81.30370,  2.70618, 1.39259, 1.20004, // AAGCMC
    82.90800,  1.50963, 1.44790, 1.27224, // AAGCMG
    81.30370,  2.70618, 1.39259, 1.20004, // AAGCMM
    82.33150,  2.70618, 1.41867, 1.23391, // AAGCMT
    81.36030,  1.49209, 1.28659, 1.02607, // AAGCTA
    79.31180,  1.27881, 1.14310, 0.85930, // AAGCTC
    80.42290,  1.45654, 1.24512, 0.97686, // AAGCTG
    80.68470,  1.46464, 1.14310, 0.85930, // AAGCTM
    80.05730,  1.20442, 0.98504, 0.68738, // AAGCTT
    78.76060,  1.76790, 1.17273, 0.78158, // AAGGAA
    73.68170,  1.84933, 1.57040, 1.21112, // AAGGAC
    77.65740,  1.90103, 1.24215, 0.85198, // AAGGAG
    75.08610,  1.83433, 1.57040, 1.21112, // AAGGAM
    75.23070,  1.94961, 1.48421, 1.11279, // AAGGAT
    73.02760,  1.71064, 1.40742, 1.12077, // AAGGCA
    71.73720,  1.74403, 1.38351, 1.09233, // AAGGCC
    72.82220,  1.67690, 1.31019, 1.00666, // AAGGCG
    72.26630,  1.75133, 1.38351, 1.09233, // AAGGCM
    72.69990,  1.63005, 1.19811, 0.88029, // AAGGCT
    79.26740,  1.39686, 1.06266, 0.73225, // AAGGGA
    76.62640,  1.71059, 1.23284, 0.91501, // AAGGGC
    78.23670,  1.42068, 1.01128, 0.67978, // AAGGGG
    76.99180,  1.40533, 1.23284, 0.91501, // AAGGGM
    77.47700,  1.80130, 1.26818, 0.95463, // AAGGGT
    73.06400,  2.67690, 1.40742, 1.12077, // AAGGMA
    71.63330,  2.67690, 1.38351, 1.09233, // AAGGMC
    76.99490,  1.63441, 1.31019, 1.00666, // AAGGMG
    71.63330,  2.67690, 1.38351, 1.09233, // AAGGMM
    72.48130,  2.67690, 1.19811, 0.88029, // AAGGMT
    71.81950,  1.50790, 1.19129, 0.85060, // AAGGTA
    70.19870,  1.80555, 1.41274, 1.09848, // AAGGTC
    71.63120,  1.62171, 1.27893, 0.94617, // AAGGTG
    71.20440,  1.63815, 1.41274, 1.09848, // AAGGTM
    71.14100,  1.97158, 1.45329, 1.14611, // AAGGTT
    87.07530,  2.87706, 1.47547, 1.22356, // AAGMAA
    84.83650,  2.87706, 1.20873, 0.90724, // AAGMAC
    86.01730,  2.87706, 1.40134, 1.13252, // AAGMAG
    84.83650,  2.87706, 1.20873, 0.90724, // AAGMAM
    85.60650,  2.87706, 1.26641, 0.97295, // AAGMAT
    82.42550,  2.70618, 1.43823, 1.25952, // AAGMCA
    81.30370,  2.70618, 1.39259, 1.20004, // AAGMCC
    82.45520,  2.70618, 1.44790, 1.27224, // AAGMCG
    81.30370,  2.70618, 1.39259, 1.20004, // AAGMCM
    82.33150,  2.70618, 1.41867, 1.23391, // AAGMCT
    81.91960,  1.09523, 1.28937, 1.05556, // AAGMGA
    79.45710,  1.13659, 1.19997, 0.94770, // AAGMGC
    81.69820,  1.51333, 1.31411, 1.08609, // AAGMGG
    80.40630,  1.20246, 1.19997, 0.94770, // AAGMGM
    80.20710,  1.35099, 1.27584, 1.03900, // AAGMGT
    82.42550,  2.70618, 1.43823, 1.25952, // AAGMMA
    81.30370,  2.70618, 1.39259, 1.20004, // AAGMMC
    82.45520,  2.70618, 1.44790, 1.27224, // AAGMMG
    81.30370,  2.70618, 1.39259, 1.20004, // AAGMMM
    82.33150,  2.70618, 1.41867, 1.23391, // AAGMMT
    81.54700,  2.42173, 1.28659, 1.02607, // AAGMTA
    79.38380,  2.42173, 1.14310, 0.85930, // AAGMTC
    80.91310,  2.42173, 1.24512, 0.97686, // AAGMTG
    79.38380,  2.42173, 1.14310, 0.85930, // AAGMTM
    80.06800,  2.42173, 0.98504, 0.68738, // AAGMTT
    77.14530,  1.91049, 1.62302, 1.46888, // AAGTAA
    73.81400,  1.90100, 1.75202, 1.64743, // AAGTAC
    76.02460,  1.71731, 1.62763, 1.47515, // AAGTAG
    75.41390,  2.33253, 1.75202, 1.64743, // AAGTAM
    75.25730,  2.07775, 1.65682, 1.51500, // AAGTAT
    69.67360,  2.20843, 2.02437, 2.11329, // AAGTCA
    68.41350,  2.38632, 2.20431, 2.40122, // AAGTCC
    69.41670,  2.42133, 2.08053, 2.20184, // AAGTCG
    69.69810,  2.61777, 2.20431, 2.40122, // AAGTCM
    69.64260,  2.47535, 2.25893, 2.49103, // AAGTCT
    73.35960,  1.80635, 1.64171, 1.59332, // AAGTGA
    69.85430,  1.90063, 1.99644, 2.13670, // AAGTGC
    72.12990,  1.80354, 1.71446, 1.70039, // AAGTGG
    71.70940,  2.28019, 1.99644, 2.13670, // AAGTGM
    70.96740,  1.99813, 1.97265, 2.09862, // AAGTGT
    69.35300,  3.42133, 2.02437, 2.11329, // AAGTMA
    68.05680,  3.42133, 2.20431, 2.40122, // AAGTMC
    72.46680,  1.81469, 2.08053, 2.20184, // AAGTMG
    68.05680,  3.42133, 2.20431, 2.40122, // AAGTMM
    69.38230,  3.42133, 2.25893, 2.49103, // AAGTMT
    64.66840,  2.34374, 2.22607, 2.19760, // AAGTTA
    61.63570,  2.17524, 2.19616, 2.15345, // AAGTTC
    64.01600,  2.33333, 2.18102, 2.13122, // AAGTTG
    63.85630,  2.62104, 2.19616, 2.15345, // AAGTTM
    62.93030,  2.19583, 2.07986, 1.98468, // AAGTTT
    107.92500, 3.19132, 1.79213, 1.53404, // AAMAAA
    104.86100, 3.19132, 1.80497, 1.55056, // AAMAAC
    106.83300, 3.19132, 1.87422, 1.64064, // AAMAAG
    104.86100, 3.19132, 1.80497, 1.55056, // AAMAAM
    106.15900, 3.19132, 1.96452, 1.76063, // AAMAAT
    99.65940,  2.74166, 1.36504, 1.00221, // AAMACA
    98.13200,  2.74166, 1.33763, 0.97218, // AAMACC
    99.53500,  2.74166, 1.32214, 0.95534, // AAMACG
    98.13200,  2.74166, 1.33763, 0.97218, // AAMACM
    99.48970,  2.74166, 1.43867, 1.08438, // AAMACT
    105.27000, 3.28987, 1.92656, 1.69338, // AAMAGA
    102.31200, 3.28987, 1.77407, 1.49637, // AAMAGC
    104.13900, 3.28987, 1.74932, 1.46516, // AAMAGG
    102.31200, 3.28987, 1.77407, 1.49637, // AAMAGM
    103.70200, 3.28987, 1.92458, 1.69077, // AAMAGT
    99.65940,  2.74166, 1.36504, 1.00221, // AAMAMA
    98.13200,  2.74166, 1.33763, 0.97218, // AAMAMC
    99.53500,  2.74166, 1.32214, 0.95534, // AAMAMG
    98.13200,  2.74166, 1.33763, 0.97218, // AAMAMM
    99.48970,  2.74166, 1.43867, 1.08438, // AAMAMT
    99.05350,  3.17760, 1.55825, 1.29235, // AAMATA
    97.47740,  3.17760, 1.61265, 1.36061, // AAMATC
    98.40490,  3.17760, 1.58327, 1.32361, // AAMATG
    97.47740,  3.17760, 1.61265, 1.36061, // AAMATM
    98.80830,  3.17760, 1.84242, 1.66153, // AAMATT
    107.94500, 2.88264, 1.53183, 1.35379, // AAMCAA
    105.07200, 2.88264, 1.51285, 1.32871, // AAMCAC
    107.00300, 2.88264, 1.48473, 1.29184, // AAMCAG
    105.07200, 2.88264, 1.51285, 1.32871, // AAMCAM
    106.26800, 2.88264, 1.65588, 1.52152, // AAMCAT
    103.93400, 2.75332, 1.29734, 1.06459, // AAMCCA
    102.23000, 2.75332, 1.24222, 0.99747, // AAMCCC
    103.83500, 2.75332, 1.34040, 1.11804, // AAMCCG
    102.23000, 2.75332, 1.24222, 0.99747, // AAMCCM
    103.20300, 2.75332, 1.47108, 1.28546, // AAMCCT
    106.59300, 2.90443, 1.60306, 1.39462, // AAMCGA
    103.84800, 2.90443, 1.47383, 1.22942, // AAMCGC
    105.63900, 2.90443, 1.54235, 1.31615, // AAMCGG
    103.84800, 2.90443, 1.47383, 1.22942, // AAMCGM
    104.87500, 2.90443, 1.68020, 1.49648, // AAMCGT
    103.93400, 2.75332, 1.29734, 1.06459, // AAMCMA
    102.23000, 2.75332, 1.24222, 0.99747, // AAMCMC
    103.83500, 2.75332, 1.34040, 1.11804, // AAMCMG
    102.23000, 2.75332, 1.24222, 0.99747, // AAMCMM
    103.20300, 2.75332, 1.47108, 1.28546, // AAMCMT
    103.12500, 2.83098, 1.49866, 1.19909, // AAMCTA
    100.84400, 2.83098, 1.52449, 1.23022, // AAMCTC
    102.21600, 2.83098, 1.48098, 1.17793, // AAMCTG
    100.84400, 2.83098, 1.52449, 1.23022, // AAMCTM
    101.84000, 2.83098, 1.69160, 1.43795, // AAMCTT
    101.93300, 2.23690, 1.81869, 1.59373, // AAMGAA
    97.33020,  2.25920, 1.72440, 1.47142, // AAMGAC
    100.91800, 2.18148, 1.78587, 1.55078, // AAMGAG
    99.36490,  2.03689, 1.72440, 1.47142, // AAMGAM
    99.55820,  2.31861, 1.95330, 1.77391, // AAMGAT
    96.15050,  2.02043, 1.43135, 1.10889, // AAMGCA
    95.23930,  2.07708, 1.50183, 1.19180, // AAMGCC
    98.42320,  3.02317, 1.53638, 1.23315, // AAMGCG
    95.45500,  2.19763, 1.50183, 1.19180, // AAMGCM
    96.94780,  2.08427, 1.58425, 1.29123, // AAMGCT
    101.84200, 2.28977, 1.82532, 1.59411, // AAMGGA
    98.61130,  2.19459, 1.69529, 1.42684, // AAMGGC
    100.92900, 2.18328, 1.74356, 1.48821, // AAMGGG
    99.94250,  2.01990, 1.69529, 1.42684, // AAMGGM
    100.36800, 2.31861, 1.88906, 1.67833, // AAMGGT
    98.63880,  3.02317, 1.43135, 1.10889, // AAMGMA
    96.96740,  3.02317, 1.50183, 1.19180, // AAMGMC
    100.33000, 1.81767, 1.53638, 1.23315, // AAMGMG
    96.96740,  3.02317, 1.50183, 1.19180, // AAMGMM
    98.63120,  3.02317, 1.58425, 1.29123, // AAMGMT
    95.75970,  2.30855, 1.75347, 1.45050, // AAMGTA
    94.21630,  2.48689, 1.79988, 1.50847, // AAMGTC
    94.98800,  2.38699, 1.75924, 1.45768, // AAMGTG
    94.89230,  2.55780, 1.79988, 1.50847, // AAMGTM
    95.05940,  2.80371, 2.09559, 1.89511, // AAMGTT
    107.94500, 2.88264, 1.53183, 1.35379, // AAMMAA
    105.07200, 2.88264, 1.51285, 1.32871, // AAMMAC
    107.00300, 2.88264, 1.48473, 1.29184, // AAMMAG
    105.07200, 2.88264, 1.51285, 1.32871, // AAMMAM
    106.26800, 2.88264, 1.65588, 1.52152, // AAMMAT
    103.93400, 2.75332, 1.29734, 1.06459, // AAMMCA
    102.23000, 2.75332, 1.24222, 0.99747, // AAMMCC
    103.83500, 2.75332, 1.34040, 1.11804, // AAMMCG
    102.23000, 2.75332, 1.24222, 0.99747, // AAMMCM
    103.20300, 2.75332, 1.47108, 1.28546, // AAMMCT
    106.59300, 2.90443, 1.60306, 1.39462, // AAMMGA
    103.84800, 2.90443, 1.47383, 1.22942, // AAMMGC
    105.63900, 2.90443, 1.54235, 1.31615, // AAMMGG
    103.84800, 2.90443, 1.47383, 1.22942, // AAMMGM
    104.87500, 2.90443, 1.68020, 1.49648, // AAMMGT
    103.93400, 2.75332, 1.29734, 1.06459, // AAMMMA
    102.23000, 2.75332, 1.24222, 0.99747, // AAMMMC
    103.83500, 2.75332, 1.34040, 1.11804, // AAMMMG
    102.23000, 2.75332, 1.24222, 0.99747, // AAMMMM
    103.20300, 2.75332, 1.47108, 1.28546, // AAMMMT
    103.12500, 2.83098, 1.49866, 1.19909, // AAMMTA
    100.84400, 2.83098, 1.52449, 1.23022, // AAMMTC
    102.21600, 2.83098, 1.48098, 1.17793, // AAMMTG
    100.84400, 2.83098, 1.52449, 1.23022, // AAMMTM
    101.84000, 2.83098, 1.69160, 1.43795, // AAMMTT
    101.57300, 2.69847, 1.59479, 1.18964, // AAMTAA
    98.06120,  2.69847, 1.40164, 0.98019, // AAMTAC
    100.42000, 2.69847, 1.58045, 1.17362, // AAMTAG
    98.06120,  2.69847, 1.40164, 0.98019, // AAMTAM
    99.67080,  2.69847, 1.51928, 1.10615, // AAMTAT
    93.66730,  2.86136, 1.57047, 1.41151, // AAMTCA
    91.54690,  2.86136, 1.59310, 1.44213, // AAMTCC
    93.52490,  2.86136, 1.54560, 1.37810, // AAMTCG
    91.54690,  2.86136, 1.59310, 1.44213, // AAMTCM
    93.05050,  2.86136, 1.57869, 1.42260, // AAMTCT
    98.74310,  2.64486, 1.41332, 1.09402, // AAMTGA
    95.15270,  2.64486, 1.32356, 0.99148, // AAMTGC
    97.38790,  2.64486, 1.38744, 1.06412, // AAMTGG
    95.15270,  2.64486, 1.32356, 0.99148, // AAMTGM
    96.27600,  2.64486, 1.35980, 1.03248, // AAMTGT
    93.66730,  2.86136, 1.57047, 1.41151, // AAMTMA
    91.54690,  2.86136, 1.59310, 1.44213, // AAMTMC
    93.52490,  2.86136, 1.54560, 1.37810, // AAMTMG
    91.54690,  2.86136, 1.59310, 1.44213, // AAMTMM
    93.05050,  2.86136, 1.57869, 1.42260, // AAMTMT
    88.93060,  2.46961, 1.29167, 0.96950, // AAMTTA
    85.82740,  2.46961, 1.18574, 0.85272, // AAMTTC
    88.07340,  2.46961, 1.18719, 0.85428, // AAMTTG
    85.82740,  2.46961, 1.18574, 0.85272, // AAMTTM
    87.03030,  2.46961, 1.07811, 0.73929, // AAMTTT
    109.48600, 3.40262, 2.50002, 2.09543, // AATAAA
    108.54900, 3.11666, 2.51648, 2.11615, // AATAAC
    109.57800, 3.08313, 2.40830, 1.98118, // AATAAG
    109.90700, 3.22364, 2.51648, 2.11615, // AATAAM
    110.33600, 3.26519, 2.74380, 2.40927, // AATAAT
    105.88800, 3.08880, 2.25473, 1.78717, // AATACA
    105.84200, 3.23614, 2.35632, 1.90929, // AATACC
    105.35100, 3.09766, 2.18793, 1.70833, // AATACG
    106.82600, 3.20029, 2.35632, 1.90929, // AATACM
    107.33900, 3.30663, 2.50950, 2.09848, // AATACT
    108.70700, 2.98678, 2.51831, 2.28269, // AATAGA
    107.01800, 3.05603, 2.38518, 2.10410, // AATAGC
    108.15700, 2.94730, 2.23292, 1.90588, // AATAGG
    108.26300, 3.01302, 2.38518, 2.10410, // AATAGM
    109.24100, 3.21227, 2.63960, 2.44959, // AATAGT
    105.95800, 4.09766, 2.25473, 1.78717, // AATAMA
    105.46200, 4.09766, 2.35632, 1.90929, // AATAMC
    114.84700, 3.14054, 2.18793, 1.70833, // AATAMG
    105.46200, 4.09766, 2.35632, 1.90929, // AATAMM
    107.32000, 4.09766, 2.50950, 2.09848, // AATAMT
    106.78500, 3.51700, 2.60030, 2.14247, // AATATA
    107.75500, 3.62505, 2.80050, 2.39460, // AATATC
    107.17000, 3.51682, 2.64585, 2.19902, // AATATG
    108.60200, 3.66276, 2.80050, 2.39460, // AATATM
    110.36500, 3.70291, 2.98101, 2.62982, // AATATT
    119.33000, 3.10285, 2.56604, 2.71939, // AATCAA
    117.85900, 2.90157, 2.51237, 2.63451, // AATCAC
    117.73700, 3.38971, 2.65432, 2.86093, // AATCAG
    119.85800, 2.58288, 2.51237, 2.63451, // AATCAM
    118.06100, 3.08411, 2.63883, 2.83591, // AATCAT
    118.37900, 3.19966, 2.50700, 2.50391, // AATCCA
    116.36200, 3.06049, 2.45317, 2.42369, // AATCCC
    118.47800, 3.04768, 2.60978, 2.65945, // AATCCG
    118.05700, 2.79552, 2.45317, 2.42369, // AATCCM
    118.17400, 3.04372, 2.62540, 2.68337, // AATCCT
    119.68500, 3.00027, 2.68172, 2.78683, // AATCGA
    116.67000, 3.00027, 2.53482, 2.56100, // AATCGC
    118.87000, 3.00027, 2.52142, 2.54073, // AATCGG
    116.67000, 4.00027, 2.53482, 2.56100, // AATCGM
    118.74600, 3.00027, 2.72499, 2.85455, // AATCGT
    118.30800, 4.04768, 2.50700, 2.50391, // AATCMA
    116.44000, 4.04768, 2.45317, 2.42369, // AATCMC
    120.22300, 2.76665, 2.60978, 2.65945, // AATCMG
    116.44000, 4.04768, 2.45317, 2.42369, // AATCMM
    117.90900, 4.04768, 2.62540, 2.68337, // AATCMT
    118.82500, 3.01181, 2.68266, 2.69276, // AATCTA
    117.75700, 3.03294, 2.67939, 2.68784, // AATCTC
    117.77800, 3.11426, 2.81019, 2.88705, // AATCTG
    118.16200, 2.92674, 2.67939, 2.68784, // AATCTM
    119.09800, 3.12758, 2.81799, 2.89908, // AATCTT
    106.89800, 3.02839, 2.38696, 2.13157, // AATGAA
    103.85200, 2.86822, 2.42541, 2.18329, // AATGAC
    106.63700, 2.85274, 2.17702, 1.85664, // AATGAG
    105.52800, 2.79324, 2.42541, 2.18329, // AATGAM
    105.93100, 3.04333, 2.60505, 2.43029, // AATGAT
    106.23000, 3.25203, 2.35365, 2.00915, // AATGCA
    105.60700, 3.27607, 2.38223, 2.04586, // AATGCC
    106.62600, 3.13415, 2.32879, 1.97740, // AATGCG
    107.04800, 3.17584, 2.38223, 2.04586, // AATGCM
    108.47900, 3.22412, 2.52659, 2.23460, // AATGCT
    108.00600, 2.99782, 2.42027, 2.12453, // AATGGA
    105.69400, 2.91904, 2.45913, 2.17591, // AATGGC
    107.34200, 2.97062, 2.39505, 2.09142, // AATGGG
    106.96100, 2.90706, 2.45913, 2.17591, // AATGGM
    107.71200, 3.11585, 2.57384, 2.32992, // AATGGT
    106.69600, 4.13415, 2.35365, 2.00915, // AATGMA
    106.34100, 4.13415, 2.38223, 2.04586, // AATGMC
    114.60600, 3.26427, 2.32879, 1.97740, // AATGMG
    106.34100, 4.13415, 2.38223, 2.04586, // AATGMM
    108.29400, 4.13415, 2.52659, 2.23460, // AATGMT
    109.07600, 3.48650, 2.75687, 2.41235, // AATGTA
    109.51200, 3.46688, 2.89791, 2.59982, // AATGTC
    108.82000, 3.44348, 2.87882, 2.57419, // AATGTG
    109.81000, 3.55469, 2.89791, 2.59982, // AATGTM
    112.06600, 3.64099, 3.09831, 2.87411, // AATGTT
    119.70300, 3.97261, 2.56604, 2.71939, // AATMAA
    117.13200, 3.97261, 2.51237, 2.63451, // AATMAC
    118.75100, 3.97261, 2.65432, 2.86093, // AATMAG
    117.13200, 3.97261, 2.51237, 2.63451, // AATMAM
    118.52400, 3.97261, 2.63883, 2.83591, // AATMAT
    118.30800, 4.04768, 2.50700, 2.50391, // AATMCA
    116.44000, 4.04768, 2.45317, 2.42369, // AATMCC
    118.47800, 4.04768, 2.60978, 2.65945, // AATMCG
    116.44000, 4.04768, 2.45317, 2.42369, // AATMCM
    117.90900, 4.04768, 2.62540, 2.68337, // AATMCT
    111.96400, 2.92552, 2.68172, 2.78683, // AATMGA
    109.41400, 2.76421, 2.53482, 2.56100, // AATMGC
    111.45000, 2.86922, 2.52142, 2.54073, // AATMGG
    111.75300, 2.94746, 2.53482, 2.56100, // AATMGM
    111.76000, 2.89843, 2.72499, 2.85455, // AATMGT
    118.30800, 4.04768, 2.50700, 2.50391, // AATMMA
    116.44000, 4.04768, 2.45317, 2.42369, // AATMMC
    118.47800, 4.04768, 2.60978, 2.65945, // AATMMG
    116.44000, 4.04768, 2.45317, 2.42369, // AATMMM
    117.90900, 4.04768, 2.62540, 2.68337, // AATMMT
    118.85700, 4.10410, 2.68266, 2.69276, // AATMTA
    117.34400, 4.10410, 2.67939, 2.68784, // AATMTC
    118.18100, 4.10410, 2.81019, 2.88705, // AATMTG
    117.34400, 4.10410, 2.67939, 2.68784, // AATMTM
    118.77100, 4.10410, 2.81799, 2.89908, // AATMTT
    106.59000, 2.38108, 1.86753, 1.84138, // AATTAA
    103.87000, 2.01371, 1.70083, 1.60042, // AATTAC
    106.79900, 2.14208, 1.80516, 1.74991, // AATTAG
    104.36200, 2.01504, 1.70083, 1.60042, // AATTAM
    104.53700, 1.97054, 1.78368, 1.71878, // AATTAT
    101.57800, 2.27164, 2.07889, 2.03614, // AATTCA
    99.43560,  2.11676, 1.95655, 1.85908, // AATTCC
    101.49900, 2.09022, 1.96630, 1.87299, // AATTCG
    100.63800, 2.05671, 1.95655, 1.85908, // AATTCM
    101.11700, 2.13774, 1.83787, 1.69252, // AATTCT
    105.15800, 2.15174, 1.82854, 1.74520, // AATTGA
    100.97300, 2.24609, 1.69701, 1.56033, // AATTGC
    104.53100, 2.13108, 1.71423, 1.58413, // AATTGG
    102.19200, 1.89830, 1.69701, 1.56033, // AATTGM
    103.54600, 2.06883, 1.72035, 1.59263, // AATTGT
    101.47700, 3.09022, 2.07889, 2.03614, // AATTMA
    99.49710,  3.09022, 1.95655, 1.85908, // AATTMC
    106.07400, 2.58814, 1.96630, 1.87299, // AATTMG
    99.49710,  3.09022, 1.95655, 1.85908, // AATTMM
    101.55500, 3.09022, 1.83787, 1.69252, // AATTMT
    96.89640,  1.91411, 1.77757, 1.61419, // AATTTA
    94.73580,  2.16208, 1.63780, 1.42759, // AATTTC
    96.38220,  2.05012, 1.72709, 1.54592, // AATTTG
    96.05760,  1.74862, 1.63780, 1.42759, // AATTTM
    96.43360,  2.01036, 1.65003, 1.44361, // AATTTT
    92.77410,  2.76979, 1.59825, 1.19732, // ACAAAA
    89.90150,  2.93590, 1.81392, 1.44767, // ACAAAC
    92.02500,  2.99317, 1.69696, 1.30994, // ACAAAG
    89.52710,  2.76735, 1.81392, 1.44767, // ACAAAM
    90.56340,  3.04676, 1.85900, 1.50197, // ACAAAT
    83.77760,  2.26690, 1.51177, 1.20984, // ACAACA
    82.28510,  2.22983, 1.54321, 1.24778, // ACAACC
    83.18670,  2.16112, 1.54582, 1.25095, // ACAACG
    82.50810,  2.19423, 1.54321, 1.24778, // ACAACM
    82.99070,  2.34516, 1.49251, 1.18680, // ACAACT
    91.43430,  2.67828, 1.60187, 1.24600, // ACAAGA
    88.90700,  2.99642, 1.71362, 1.37863, // ACAAGC
    90.38190,  2.84319, 1.61613, 1.26266, // ACAAGG
    88.37800,  2.53839, 1.71362, 1.37863, // ACAAGM
    89.56260,  3.24360, 1.87238, 1.57459, // ACAAGT
    83.13680,  3.16112, 1.51177, 1.20984, // ACAAMA
    82.21650,  3.16112, 1.54321, 1.24778, // ACAAMC
    88.29270,  2.48579, 1.54582, 1.25095, // ACAAMG
    82.21650,  3.16112, 1.54321, 1.24778, // ACAAMM
    82.76920,  3.16112, 1.49251, 1.18680, // ACAAMT
    82.66420,  2.41185, 1.64608, 1.39736, // ACAATA
    80.93500,  2.47166, 1.97752, 1.83998, // ACAATC
    82.13360,  2.11712, 1.76664, 1.55366, // ACAATG
    81.95930,  2.25680, 1.97752, 1.83998, // ACAATM
    81.25860,  2.70630, 2.00648, 1.88054, // ACAATT
    95.29530,  1.23926, 1.05715, 0.80493, // ACACAA
    92.73200,  1.30575, 1.06102, 0.80936, // ACACAC
    94.50730,  1.28040, 1.04348, 0.78937, // ACACAG
    93.38940,  1.38053, 1.06102, 0.80936, // ACACAM
    93.69830,  1.20590, 0.97070, 0.70824, // ACACAT
    91.60880,  1.40037, 1.19393, 0.96929, // ACACCA
    90.90560,  1.43708, 1.15189, 0.91854, // ACACCC
    91.49950,  1.29044, 1.10979, 0.86865, // ACACCG
    91.20560,  1.44950, 1.15189, 0.91854, // ACACCM
    91.59960,  1.30552, 1.12044, 0.88119, // ACACCT
    94.08570,  1.39798, 1.00994, 0.73926, // ACACGA
    91.54810,  1.39798, 1.08903, 0.82779, // ACACGC
    93.27920,  1.39798, 1.05984, 0.79473, // ACACGG
    91.54810,  2.39798, 1.08903, 0.82779, // ACACGM
    92.34440,  1.39798, 1.07297, 0.80954, // ACACGT
    91.76720,  2.29044, 1.19393, 0.96929, // ACACMA
    90.32110,  2.29044, 1.15189, 0.91854, // ACACMC
    92.24700,  1.33431, 1.10979, 0.86865, // ACACMG
    90.32110,  2.29044, 1.15189, 0.91854, // ACACMM
    91.20300,  2.29044, 1.12044, 0.88119, // ACACMT
    90.40680,  1.52132, 1.12931, 0.82924, // ACACTA
    88.23050,  1.62082, 1.12747, 0.82720, // ACACTC
    89.69680,  1.56217, 1.12618, 0.82579, // ACACTG
    88.99160,  1.63247, 1.12747, 0.82720, // ACACTM
    89.34520,  1.61376, 1.09133, 0.78776, // ACACTT
    88.77880,  2.69186, 1.64646, 1.26460, // ACAGAA
    83.08260,  2.53914, 1.92949, 1.60431, // ACAGAC
    87.41120,  2.73791, 1.73953, 1.37333, // ACAGAG
    84.40230,  2.68111, 1.92949, 1.60431, // ACAGAM
    84.91760,  2.63475, 1.92427, 1.59781, // ACAGAT
    83.87540,  2.18696, 1.42329, 1.19082, // ACAGCA
    81.80640,  2.44759, 1.43105, 1.20057, // ACAGCC
    82.88880,  2.08236, 1.40829, 1.17205, // ACAGCG
    82.11530,  2.28750, 1.43105, 1.20057, // ACAGCM
    83.44860,  2.03650, 1.52849, 1.32526, // ACAGCT
    88.38040,  2.53803, 1.58798, 1.26495, // ACAGGA
    84.54670,  2.61326, 1.73486, 1.44444, // ACAGGC
    87.12870,  2.56024, 1.55245, 1.22273, // ACAGGG
    85.13670,  2.59534, 1.73486, 1.44444, // ACAGGM
    86.14410,  2.69764, 1.82887, 1.56343, // ACAGGT
    83.01820,  3.08236, 1.42329, 1.19082, // ACAGMA
    81.45030,  3.08236, 1.43105, 1.20057, // ACAGMC
    86.04400,  2.20477, 1.40829, 1.17205, // ACAGMG
    81.45030,  3.08236, 1.43105, 1.20057, // ACAGMM
    82.80060,  3.08236, 1.52849, 1.32526, // ACAGMT
    80.63540,  2.19230, 1.91397, 1.72963, // ACAGTA
    77.24180,  2.47800, 2.09972, 1.98743, // ACAGTC
    79.22750,  2.28339, 1.89939, 1.70991, // ACAGTG
    78.55130,  2.29733, 2.09972, 1.98743, // ACAGTM
    78.50690,  2.67418, 2.06168, 1.93366, // ACAGTT
    94.96910,  2.22481, 1.05715, 0.80493, // ACAMAA
    92.34910,  2.22481, 1.06102, 0.80936, // ACAMAC
    94.13070,  2.22481, 1.04348, 0.78937, // ACAMAG
    92.34910,  2.22481, 1.06102, 0.80936, // ACAMAM
    93.39940,  2.22481, 0.97070, 0.70824, // ACAMAT
    91.76720,  2.29044, 1.19393, 0.96929, // ACAMCA
    90.32110,  2.29044, 1.15189, 0.91854, // ACAMCC
    91.49950,  2.29044, 1.10979, 0.86865, // ACAMCG
    90.32110,  2.29044, 1.15189, 0.91854, // ACAMCM
    91.20300,  2.29044, 1.12044, 0.88119, // ACAMCT
    94.12890,  2.15591, 1.00994, 0.73926, // ACAMGA
    91.46000,  2.29852, 1.08903, 0.82779, // ACAMGC
    93.06120,  2.35805, 1.05984, 0.79473, // ACAMGG
    92.09180,  2.47364, 1.08903, 0.82779, // ACAMGM
    91.87820,  2.24632, 1.07297, 0.80954, // ACAMGT
    91.76720,  2.29044, 1.19393, 0.96929, // ACAMMA
    90.32110,  2.29044, 1.15189, 0.91854, // ACAMMC
    91.49950,  2.29044, 1.10979, 0.86865, // ACAMMG
    90.32110,  2.29044, 1.15189, 0.91854, // ACAMMM
    91.20300,  2.29044, 1.12044, 0.88119, // ACAMMT
    90.18550,  2.56257, 1.12931, 0.82924, // ACAMTA
    87.93340,  2.56257, 1.12747, 0.82720, // ACAMTC
    89.42720,  2.56257, 1.12618, 0.82579, // ACAMTG
    87.93340,  2.56257, 1.12747, 0.82720, // ACAMTM
    89.02180,  2.56257, 1.09133, 0.78776, // ACAMTT
    89.27800,  2.01912, 1.62646, 1.31493, // ACATAA
    85.88680,  2.11105, 1.69418, 1.39791, // ACATAC
    88.68400,  2.14249, 1.54416, 1.21641, // ACATAG
    86.79450,  2.42211, 1.69418, 1.39791, // ACATAM
    87.39450,  2.09812, 1.68679, 1.38878, // ACATAT
    83.00120,  2.37160, 2.14178, 2.16515, // ACATCA
    81.22500,  2.29624, 2.23229, 2.30384, // ACATCC
    82.23760,  2.22932, 2.00829, 1.96592, // ACATCG
    81.68810,  2.50732, 2.23229, 2.30384, // ACATCM
    82.11580,  2.20937, 2.09412, 2.09328, // ACATCT
    86.78490,  2.17729, 1.56195, 1.23545, // ACATGA
    83.23900,  2.35041, 1.79534, 1.52245, // ACATGC
    85.61080,  2.27982, 1.74257, 1.45582, // ACATGG
    83.97920,  2.31808, 1.79534, 1.52245, // ACATGM
    84.12520,  2.23878, 1.83102, 1.56805, // ACATGT
    82.32280,  3.22932, 2.14178, 2.16515, // ACATMA
    80.65780,  3.22932, 2.23229, 2.30384, // ACATMC
    85.02200,  2.26421, 2.00829, 1.96592, // ACATMG
    80.65780,  3.22932, 2.23229, 2.30384, // ACATMM
    81.80180,  3.22932, 2.09412, 2.09328, // ACATMT
    76.67630,  2.43457, 2.12527, 1.99846, // ACATTA
    73.65410,  2.49842, 2.23765, 2.15905, // ACATTC
    75.80940,  2.40883, 2.19356, 2.09555, // ACATTG
    74.81790,  2.52697, 2.23765, 2.15905, // ACATTM
    74.96360,  2.43515, 2.04004, 1.87946, // ACATTT
    103.95400, 1.51735, 1.23603, 1.02003, // ACCAAA
    102.19800, 1.69209, 1.45536, 1.30325, // ACCAAC
    103.13800, 1.56283, 1.29590, 1.09503, // ACCAAG
    102.42400, 1.55055, 1.45536, 1.30325, // ACCAAM
    102.54100, 1.63227, 1.49323, 1.35444, // ACCAAT
    97.27160,  1.49050, 1.05360, 0.74833, // ACCACA
    95.49960,  1.44168, 1.10929, 0.80845, // ACCACC
    96.08640,  1.46322, 1.09260, 0.79027, // ACCACG
    96.03340,  1.42050, 1.10929, 0.80845, // ACCACM
    96.53180,  1.56292, 1.15982, 0.86431, // ACCACT
    103.40100, 2.03668, 1.39859, 1.15660, // ACCAGA
    101.20900, 2.25747, 1.44253, 1.21154, // ACCAGC
    101.36900, 1.78989, 1.39727, 1.15497, // ACCAGG
    100.90800, 1.73811, 1.44253, 1.21154, // ACCAGM
    100.23600, 1.97881, 1.59888, 1.41375, // ACCAGT
    96.35810,  2.46322, 1.05360, 0.74833, // ACCAMA
    95.03570,  2.46322, 1.10929, 0.80845, // ACCAMC
    100.08100, 1.33840, 1.09260, 0.79027, // ACCAMG
    95.03570,  2.46322, 1.10929, 0.80845, // ACCAMM
    95.95960,  2.46322, 1.15982, 0.86431, // ACCAMT
    95.42800,  1.67812, 1.30899, 0.96780, // ACCATA
    93.37690,  1.96321, 1.50756, 1.19616, // ACCATC
    94.51650,  1.74106, 1.39952, 1.06991, // ACCATG
    94.74870,  1.83497, 1.50756, 1.19616, // ACCATM
    94.32140,  1.99312, 1.64925, 1.36870, // ACCATT
    103.61000, 0.97223, 0.87226, 0.61194, // ACCCAA
    100.73800, 1.23988, 0.96606, 0.71325, // ACCCAC
    102.67200, 1.01624, 0.89341, 0.63433, // ACCCAG
    101.51900, 0.99946, 0.96606, 0.71325, // ACCCAM
    101.74400, 1.29932, 1.06095, 0.82088, // ACCCAT
    99.33880,  1.19289, 0.89591, 0.64955, // ACCCCA
    97.89600,  1.11878, 0.89646, 0.65015, // ACCCCC
    99.21720,  1.40182, 0.90089, 0.65497, // ACCCCG
    98.50940,  1.05557, 0.89646, 0.65015, // ACCCCM
    99.23030,  1.06822, 0.87739, 0.62951, // ACCCCT
    101.93700, 1.43192, 0.92684, 0.68212, // ACCCGA
    99.34460,  1.43192, 0.94800, 0.70561, // ACCCGC
    100.73300, 1.43192, 0.90440, 0.65750, // ACCCGG
    99.34460,  2.43192, 0.94800, 0.70561, // ACCCGM
    100.16700, 1.43192, 1.03949, 0.81019, // ACCCGT
    99.35500,  2.40182, 0.89591, 0.64955, // ACCCMA
    97.85840,  2.40182, 0.89646, 0.65015, // ACCCMC
    99.65530,  1.03394, 0.90089, 0.65497, // ACCCMG
    97.85840,  2.40182, 0.89646, 0.65015, // ACCCMM
    99.09360,  2.40182, 0.87739, 0.62951, // ACCCMT
    98.47610,  1.22402, 0.97684, 0.73578, // ACCCTA
    96.67910,  1.42527, 1.13600, 0.92274, // ACCCTC
    98.45050,  1.39223, 0.99337, 0.75453, // ACCCTG
    97.64320,  1.35888, 1.13600, 0.92274, // ACCCTM
    97.27350,  1.52359, 1.19237, 0.99226, // ACCCTT
    99.07040,  1.82025, 1.36048, 1.11248, // ACCGAA
    94.17780,  1.82025, 1.59801, 1.41621, // ACCGAC
    97.84920,  1.82025, 1.41891, 1.18491, // ACCGAG
    94.17780,  2.82025, 1.59801, 1.41621, // ACCGAM
    95.99360,  1.82025, 1.67392, 1.51830, // ACCGAT
    94.48170,  1.58093, 1.12144, 0.78696, // ACCGCA
    92.97800,  1.58093, 1.21249, 0.88471, // ACCGCC
    94.16970,  1.58093, 1.18195, 0.85150, // ACCGCG
    92.97800,  2.58093, 1.21249, 0.88471, // ACCGCM
    94.08150,  1.58093, 1.21946, 0.89235, // ACCGCT
    98.80270,  1.74812, 1.35912, 1.15166, // ACCGGA
    95.47870,  1.74812, 1.45115, 1.27059, // ACCGGC
    97.57450,  1.74812, 1.36074, 1.15372, // ACCGGG
    95.47870,  2.74812, 1.45115, 1.27059, // ACCGGM
    96.85990,  1.74812, 1.51550, 1.35604, // ACCGGT
    94.48170,  2.58093, 1.12144, 0.78696, // ACCGMA
    92.97800,  2.58093, 1.21249, 0.88471, // ACCGMC
    94.16970,  2.58093, 1.18195, 0.85150, // ACCGMG
    92.97800,  2.58093, 1.21249, 0.88471, // ACCGMM
    94.08150,  2.58093, 1.21946, 0.89235, // ACCGMT
    92.62400,  2.13467, 1.44073, 1.19456, // ACCGTA
    90.17100,  2.13467, 1.67103, 1.49214, // ACCGTC
    91.68520,  2.13467, 1.59610, 1.39291, // ACCGTG
    90.17100,  3.13467, 1.67103, 1.49214, // ACCGTM
    91.56360,  2.13467, 1.79201, 1.65708, // ACCGTT
    103.40400, 2.44981, 0.87226, 0.61194, // ACCMAA
    100.50300, 2.44981, 0.96606, 0.71325, // ACCMAC
    102.49000, 2.44981, 0.89341, 0.63433, // ACCMAG
    100.50300, 2.44981, 0.96606, 0.71325, // ACCMAM
    101.52700, 2.44981, 1.06095, 0.82088, // ACCMAT
    99.35500,  2.40182, 0.89591, 0.64955, // ACCMCA
    97.85840,  2.40182, 0.89646, 0.65015, // ACCMCC
    99.21720,  2.40182, 0.90089, 0.65497, // ACCMCG
    97.85840,  2.40182, 0.89646, 0.65015, // ACCMCM
    99.09360,  2.40182, 0.87739, 0.62951, // ACCMCT
    100.64000, 1.29101, 0.92684, 0.68212, // ACCMGA
    97.97330,  1.50625, 0.94800, 0.70561, // ACCMGC
    100.00900, 1.44570, 0.90440, 0.65750, // ACCMGG
    98.96460,  1.28820, 0.94800, 0.70561, // ACCMGM
    99.01530,  1.72116, 1.03949, 0.81019, // ACCMGT
    99.35500,  2.40182, 0.89591, 0.64955, // ACCMMA
    97.85840,  2.40182, 0.89646, 0.65015, // ACCMMC
    99.21720,  2.40182, 0.90089, 0.65497, // ACCMMG
    97.85840,  2.40182, 0.89646, 0.65015, // ACCMMM
    99.09360,  2.40182, 0.87739, 0.62951, // ACCMMT
    98.28370,  2.44710, 0.97684, 0.73578, // ACCMTA
    96.48630,  2.44710, 1.13600, 0.92274, // ACCMTC
    97.66750,  2.44710, 0.99337, 0.75453, // ACCMTG
    96.48630,  2.44710, 1.13600, 0.92274, // ACCMTM
    96.88320,  2.44710, 1.19237, 0.99226, // ACCMTT
    98.36010,  1.60744, 1.30411, 1.02284, // ACCTAA
    95.37600,  1.71475, 1.37864, 1.11177, // ACCTAC
    97.35410,  1.50312, 1.05253, 0.74164, // ACCTAG
    96.55870,  1.88172, 1.37864, 1.11177, // ACCTAM
    96.41820,  1.74373, 1.36413, 1.09426, // ACCTAT
    91.69520,  1.93512, 1.71292, 1.55628, // ACCTCA
    90.36560,  2.04311, 1.86109, 1.76252, // ACCTCC
    91.39750,  2.16210, 1.62527, 1.43836, // ACCTCG
    91.08180,  2.34319, 1.86109, 1.76252, // ACCTCM
    91.48020,  2.10542, 1.78964, 1.66200, // ACCTCT
    95.19070,  1.58620, 1.26908, 0.98862, // ACCTGA
    92.93460,  1.76258, 1.41720, 1.16665, // ACCTGC
    93.72280,  1.66932, 1.29676, 1.02113, // ACCTGG
    93.80990,  1.85177, 1.41720, 1.16665, // ACCTGM
    93.64140,  1.74152, 1.44582, 1.20217, // ACCTGT
    91.24900,  3.16210, 1.71292, 1.55628, // ACCTMA
    90.14290,  3.16210, 1.86109, 1.76252, // ACCTMC
    93.38540,  1.72972, 1.62527, 1.43836, // ACCTMG
    90.14290,  3.16210, 1.86109, 1.76252, // ACCTMM
    91.26810,  3.16210, 1.78964, 1.66200, // ACCTMT
    86.82400,  1.84507, 1.71890, 1.48956, // ACCTTA
    83.93740,  1.83551, 1.70994, 1.47793, // ACCTTC
    86.11700,  1.81074, 1.56463, 1.29360, // ACCTTG
    85.35680,  1.94061, 1.70994, 1.47793, // ACCTTM
    84.82430,  1.73091, 1.58306, 1.31652, // ACCTTT
    85.68490,  2.36795, 1.56701, 1.16930, // ACGAAA
    82.10620,  2.36795, 1.88115, 1.53799, // ACGAAC
    84.58060,  2.36795, 1.70811, 1.33074, // ACGAAG
    82.10620,  3.36795, 1.88115, 1.53799, // ACGAAM
    83.06390,  2.36795, 1.92702, 1.59459, // ACGAAT
    75.36940,  2.00332, 1.81079, 1.55061, // ACGACA
    74.19600,  2.00332, 1.65631, 1.35647, // ACGACC
    75.24330,  2.00332, 1.71221, 1.42572, // ACGACG
    74.19600,  3.00332, 1.65631, 1.35647, // ACGACM
    75.11470,  2.00332, 1.66362, 1.36547, // ACGACT
    84.40650,  2.41291, 1.51583, 1.11615, // ACGAGA
    82.02660,  2.41291, 1.78219, 1.42291, // ACGAGC
    83.11420,  2.41291, 1.51980, 1.12053, // ACGAGG
    82.02660,  3.41291, 1.78219, 1.42291, // ACGAGM
    82.40090,  2.41291, 1.83001, 1.48055, // ACGAGT
    75.36940,  3.00332, 1.81079, 1.55061, // ACGAMA
    74.19600,  3.00332, 1.65631, 1.35647, // ACGAMC
    75.24330,  3.00332, 1.71221, 1.42572, // ACGAMG
    74.19600,  3.00332, 1.65631, 1.35647, // ACGAMM
    75.11470,  3.00332, 1.66362, 1.36547, // ACGAMT
    74.29910,  1.85855, 1.56104, 1.41982, // ACGATA
    72.41120,  1.85855, 1.80103, 1.75951, // ACGATC
    73.39190,  1.85855, 1.56900, 1.43069, // ACGATG
    72.41120,  2.85855, 1.80103, 1.75951, // ACGATM
    73.68440,  1.85855, 1.93844, 1.96467, // ACGATT
    90.72400,  1.44291, 1.25549, 0.98457, // ACGCAA
    87.92700,  1.44291, 1.18231, 0.89975, // ACGCAC
    89.69160,  1.44291, 1.23807, 0.96416, // ACGCAG
    87.92700,  2.44291, 1.18231, 0.89975, // ACGCAM
    88.90820,  1.44291, 1.16388, 0.87880, // ACGCAT
    86.80540,  1.52741, 1.41921, 1.19162, // ACGCCA
    85.42380,  1.52741, 1.44306, 1.22178, // ACGCCC
    86.59410,  1.52741, 1.41830, 1.19047, // ACGCCG
    85.42380,  2.52741, 1.44306, 1.22178, // ACGCCM
    86.47020,  1.52741, 1.36153, 1.11971, // ACGCCT
    88.86570,  1.58181, 1.28360, 0.98233, // ACGCGA
    85.75870,  1.58181, 1.35971, 1.07099, // ACGCGC
    87.67580,  1.58181, 1.34631, 1.05519, // ACGCGG
    85.75870,  2.58181, 1.35971, 1.07099, // ACGCGM
    86.47440,  1.58181, 1.34700, 1.05600, // ACGCGT
    86.80540,  2.52741, 1.41921, 1.19162, // ACGCMA
    85.42380,  2.52741, 1.44306, 1.22178, // ACGCMC
    86.59410,  2.52741, 1.41830, 1.19047, // ACGCMG
    85.42380,  2.52741, 1.44306, 1.22178, // ACGCMM
    86.47020,  2.52741, 1.36153, 1.11971, // ACGCMT
    85.13680,  1.56040, 1.41851, 1.12330, // ACGCTA
    82.69010,  1.56040, 1.39000, 1.08960, // ACGCTC
    84.23900,  1.56040, 1.39386, 1.09415, // ACGCTG
    82.69010,  2.56040, 1.39000, 1.08960, // ACGCTM
    83.58710,  1.56040, 1.18494, 0.85761, // ACGCTT
    84.93960,  2.44172, 1.76816, 1.37549, // ACGGAA
    79.42780,  2.44172, 2.03723, 1.70112, // ACGGAC
    83.71980,  2.44172, 1.82726, 1.44503, // ACGGAG
    79.42780,  3.44172, 2.03723, 1.70112, // ACGGAM
    81.12400,  2.44172, 2.01232, 1.67002, // ACGGAT
    79.99910,  2.03186, 1.49247, 1.20792, // ACGGCA
    78.44170,  2.03186, 1.50469, 1.22280, // ACGGCC
    79.78450,  2.03186, 1.48313, 1.19660, // ACGGCG
    78.44170,  3.03186, 1.50469, 1.22280, // ACGGCM
    79.70290,  2.03186, 1.50224, 1.21981, // ACGGCT
    85.58090,  2.35270, 1.63675, 1.21654, // ACGGGA
    82.61030,  2.35270, 1.81525, 1.42087, // ACGGGC
    84.24850,  2.35270, 1.59567, 1.17102, // ACGGGG
    82.61030,  3.35270, 1.81525, 1.42087, // ACGGGM
    83.63380,  2.35270, 1.85320, 1.46566, // ACGGGT
    79.99910,  3.03186, 1.49247, 1.20792, // ACGGMA
    78.44170,  3.03186, 1.50469, 1.22280, // ACGGMC
    79.78450,  3.03186, 1.48313, 1.19660, // ACGGMG
    78.44170,  3.03186, 1.50469, 1.22280, // ACGGMM
    79.70290,  3.03186, 1.50224, 1.21981, // ACGGMT
    78.32030,  2.46096, 1.84312, 1.56160, // ACGGTA
    75.53450,  2.46096, 2.00903, 1.77714, // ACGGTC
    77.16700,  2.46096, 1.92296, 1.66416, // ACGGTG
    75.53450,  3.46096, 2.00903, 1.77714, // ACGGTM
    76.54800,  2.46096, 1.99035, 1.75241, // ACGGTT
    90.72400,  2.44291, 1.25549, 0.98457, // ACGMAA
    87.92700,  2.44291, 1.18231, 0.89975, // ACGMAC
    89.69160,  2.44291, 1.23807, 0.96416, // ACGMAG
    87.92700,  2.44291, 1.18231, 0.89975, // ACGMAM
    88.90820,  2.44291, 1.16388, 0.87880, // ACGMAT
    86.80540,  2.52741, 1.41921, 1.19162, // ACGMCA
    85.42380,  2.52741, 1.44306, 1.22178, // ACGMCC
    86.59410,  2.52741, 1.41830, 1.19047, // ACGMCG
    85.42380,  2.52741, 1.44306, 1.22178, // ACGMCM
    86.47020,  2.52741, 1.36153, 1.11971, // ACGMCT
    88.86570,  2.58181, 1.28360, 0.98233, // ACGMGA
    85.75870,  2.58181, 1.35971, 1.07099, // ACGMGC
    87.67580,  2.58181, 1.34631, 1.05519, // ACGMGG
    85.75870,  2.58181, 1.35971, 1.07099, // ACGMGM
    86.47440,  2.58181, 1.34700, 1.05600, // ACGMGT
    86.80540,  2.52741, 1.41921, 1.19162, // ACGMMA
    85.42380,  2.52741, 1.44306, 1.22178, // ACGMMC
    86.59410,  2.52741, 1.41830, 1.19047, // ACGMMG
    85.42380,  2.52741, 1.44306, 1.22178, // ACGMMM
    86.47020,  2.52741, 1.36153, 1.11971, // ACGMMT
    85.13680,  2.56040, 1.41851, 1.12330, // ACGMTA
    82.69010,  2.56040, 1.39000, 1.08960, // ACGMTC
    84.23900,  2.56040, 1.39386, 1.09415, // ACGMTG
    82.69010,  2.56040, 1.39000, 1.08960, // ACGMTM
    83.58710,  2.56040, 1.18494, 0.85761, // ACGMTT
    82.12720,  2.14468, 1.80972, 1.57415, // ACGTAA
    78.53430,  2.14468, 1.93097, 1.73496, // ACGTAC
    81.44440,  2.14468, 1.89084, 1.68117, // ACGTAG
    78.53430,  3.14468, 1.93097, 1.73496, // ACGTAM
    80.16140,  2.14468, 2.00056, 1.82960, // ACGTAT
    75.28890,  2.49565, 2.48078, 2.86185, // ACGTCA
    73.78890,  2.49565, 2.54924, 2.98112, // ACGTCC
    75.21510,  2.49565, 2.45874, 2.82380, // ACGTCG
    73.78890,  3.49565, 2.54924, 2.98112, // ACGTCM
    75.03440,  2.49565, 2.47828, 2.85752, // ACGTCT
    78.72390,  2.22005, 1.90469, 1.77665, // ACGTGA
    74.67260,  2.22005, 2.14974, 2.13031, // ACGTGC
    77.41130,  2.22005, 2.11114, 2.07319, // ACGTGG
    74.67260,  3.22005, 2.14974, 2.13031, // ACGTGM
    75.76270,  2.22005, 2.24488, 2.27328, // ACGTGT
    75.28890,  3.49565, 2.48078, 2.86185, // ACGTMA
    73.78890,  3.49565, 2.54924, 2.98112, // ACGTMC
    75.21510,  3.49565, 2.45874, 2.82380, // ACGTMG
    73.78890,  3.49565, 2.54924, 2.98112, // ACGTMM
    75.03440,  3.49565, 2.47828, 2.85752, // ACGTMT
    69.69300,  2.47313, 2.39457, 2.49536, // ACGTTA
    66.19740,  2.47313, 2.51962, 2.69335, // ACGTTC
    68.73890,  2.47313, 2.44079, 2.56795, // ACGTTG
    66.19740,  3.47313, 2.51962, 2.69335, // ACGTTM
    67.60650,  2.47313, 2.40054, 2.50469, // ACGTTT
    103.97700, 2.63085, 1.23603, 1.02003, // ACMAAA
    101.47400, 2.63085, 1.45536, 1.30325, // ACMAAC
    103.08300, 2.63085, 1.29590, 1.09503, // ACMAAG
    101.47400, 2.63085, 1.45536, 1.30325, // ACMAAM
    102.38000, 2.63085, 1.49323, 1.35444, // ACMAAT
    96.35810,  2.46322, 1.05360, 0.74833, // ACMACA
    95.03570,  2.46322, 1.10929, 0.80845, // ACMACC
    96.08640,  2.46322, 1.09260, 0.79027, // ACMACG
    95.03570,  2.46322, 1.10929, 0.80845, // ACMACM
    95.95960,  2.46322, 1.15982, 0.86431, // ACMACT
    101.89800, 2.84203, 1.39859, 1.15660, // ACMAGA
    99.59450,  2.84203, 1.44253, 1.21154, // ACMAGC
    100.71000, 2.84203, 1.39727, 1.15497, // ACMAGG
    99.59450,  2.84203, 1.44253, 1.21154, // ACMAGM
    100.27000, 2.84203, 1.59888, 1.41375, // ACMAGT
    96.35810,  2.46322, 1.05360, 0.74833, // ACMAMA
    95.03570,  2.46322, 1.10929, 0.80845, // ACMAMC
    96.08640,  2.46322, 1.09260, 0.79027, // ACMAMG
    95.03570,  2.46322, 1.10929, 0.80845, // ACMAMM
    95.95960,  2.46322, 1.15982, 0.86431, // ACMAMT
    95.04160,  2.79927, 1.30899, 0.96780, // ACMATA
    93.32730,  2.79927, 1.50756, 1.19616, // ACMATC
    94.58490,  2.79927, 1.39952, 1.06991, // ACMATG
    93.32730,  2.79927, 1.50756, 1.19616, // ACMATM
    94.37830,  2.79927, 1.64925, 1.36870, // ACMATT
    103.40400, 2.44981, 0.87226, 0.61194, // ACMCAA
    100.50300, 2.44981, 0.96606, 0.71325, // ACMCAC
    102.49000, 2.44981, 0.89341, 0.63433, // ACMCAG
    100.50300, 2.44981, 0.96606, 0.71325, // ACMCAM
    101.52700, 2.44981, 1.06095, 0.82088, // ACMCAT
    99.35500,  2.40182, 0.89591, 0.64955, // ACMCCA
    97.85840,  2.40182, 0.89646, 0.65015, // ACMCCC
    99.21720,  2.40182, 0.90089, 0.65497, // ACMCCG
    97.85840,  2.40182, 0.89646, 0.65015, // ACMCCM
    99.09360,  2.40182, 0.87739, 0.62951, // ACMCCT
    101.93700, 2.43192, 0.92684, 0.68212, // ACMCGA
    99.34460,  2.43192, 0.94800, 0.70561, // ACMCGC
    100.73300, 2.43192, 0.90440, 0.65750, // ACMCGG
    99.34460,  2.43192, 0.94800, 0.70561, // ACMCGM
    100.16700, 2.43192, 1.03949, 0.81019, // ACMCGT
    99.35500,  2.40182, 0.89591, 0.64955, // ACMCMA
    97.85840,  2.40182, 0.89646, 0.65015, // ACMCMC
    99.21720,  2.40182, 0.90089, 0.65497, // ACMCMG
    97.85840,  2.40182, 0.89646, 0.65015, // ACMCMM
    99.09360,  2.40182, 0.87739, 0.62951, // ACMCMT
    98.28370,  2.44710, 0.97684, 0.73578, // ACMCTA
    96.48630,  2.44710, 1.13600, 0.92274, // ACMCTC
    97.66750,  2.44710, 0.99337, 0.75453, // ACMCTG
    96.48630,  2.44710, 1.13600, 0.92274, // ACMCTM
    96.88320,  2.44710, 1.19237, 0.99226, // ACMCTT
    98.48420,  1.46232, 1.36048, 1.11248, // ACMGAA
    93.89500,  1.69845, 1.59801, 1.41621, // ACMGAC
    97.16800,  1.55424, 1.41891, 1.18491, // ACMGAG
    95.77590,  1.64479, 1.59801, 1.41621, // ACMGAM
    95.52590,  1.65076, 1.67392, 1.51830, // ACMGAT
    92.82610,  1.35481, 1.12144, 0.78696, // ACMGCA
    91.19420,  1.55617, 1.21249, 0.88471, // ACMGCC
    94.16970,  2.58093, 1.18195, 0.85150, // ACMGCG
    92.44780,  1.78695, 1.21249, 0.88471, // ACMGCM
    93.69470,  1.94812, 1.21946, 0.89235, // ACMGCT
    98.04480,  1.48615, 1.35912, 1.15166, // ACMGGA
    94.93850,  1.64709, 1.45115, 1.27059, // ACMGGC
    96.91280,  1.44860, 1.36074, 1.15372, // ACMGGG
    96.47030,  1.53281, 1.45115, 1.27059, // ACMGGM
    96.62110,  1.64010, 1.51550, 1.35604, // ACMGGT
    94.48170,  2.58093, 1.12144, 0.78696, // ACMGMA
    92.97800,  2.58093, 1.21249, 0.88471, // ACMGMC
    95.52330,  1.47546, 1.18195, 0.85150, // ACMGMG
    92.97800,  2.58093, 1.21249, 0.88471, // ACMGMM
    94.08150,  2.58093, 1.21946, 0.89235, // ACMGMT
    92.41170,  1.91520, 1.44073, 1.19456, // ACMGTA
    89.65200,  2.19307, 1.67103, 1.49214, // ACMGTC
    91.71640,  2.11772, 1.59610, 1.39291, // ACMGTG
    91.07010,  2.04370, 1.67103, 1.49214, // ACMGTM
    90.90410,  2.12405, 1.79201, 1.65708, // ACMGTT
    103.40400, 2.44981, 0.87226, 0.61194, // ACMMAA
    100.50300, 2.44981, 0.96606, 0.71325, // ACMMAC
    102.49000, 2.44981, 0.89341, 0.63433, // ACMMAG
    100.50300, 2.44981, 0.96606, 0.71325, // ACMMAM
    101.52700, 2.44981, 1.06095, 0.82088, // ACMMAT
    99.35500,  2.40182, 0.89591, 0.64955, // ACMMCA
    97.85840,  2.40182, 0.89646, 0.65015, // ACMMCC
    99.21720,  2.40182, 0.90089, 0.65497, // ACMMCG
    97.85840,  2.40182, 0.89646, 0.65015, // ACMMCM
    99.09360,  2.40182, 0.87739, 0.62951, // ACMMCT
    101.93700, 2.43192, 0.92684, 0.68212, // ACMMGA
    99.34460,  2.43192, 0.94800, 0.70561, // ACMMGC
    100.73300, 2.43192, 0.90440, 0.65750, // ACMMGG
    99.34460,  2.43192, 0.94800, 0.70561, // ACMMGM
    100.16700, 2.43192, 1.03949, 0.81019, // ACMMGT
    99.35500,  2.40182, 0.89591, 0.64955, // ACMMMA
    97.85840,  2.40182, 0.89646, 0.65015, // ACMMMC
    99.21720,  2.40182, 0.90089, 0.65497, // ACMMMG
    97.85840,  2.40182, 0.89646, 0.65015, // ACMMMM
    99.09360,  2.40182, 0.87739, 0.62951, // ACMMMT
    98.28370,  2.44710, 0.97684, 0.73578, // ACMMTA
    96.48630,  2.44710, 1.13600, 0.92274, // ACMMTC
    97.66750,  2.44710, 0.99337, 0.75453, // ACMMTG
    96.48630,  2.44710, 1.13600, 0.92274, // ACMMTM
    96.88320,  2.44710, 1.19237, 0.99226, // ACMMTT
    98.01510,  2.77408, 1.30411, 1.02284, // ACMTAA
    95.25880,  2.77408, 1.37864, 1.11177, // ACMTAC
    97.16900,  2.77408, 1.05253, 0.74164, // ACMTAG
    95.25880,  2.77408, 1.37864, 1.11177, // ACMTAM
    96.13040,  2.77408, 1.36413, 1.09426, // ACMTAT
    91.24900,  3.16210, 1.71292, 1.55628, // ACMTCA
    90.14290,  3.16210, 1.86109, 1.76252, // ACMTCC
    91.39750,  3.16210, 1.62527, 1.43836, // ACMTCG
    90.14290,  3.16210, 1.86109, 1.76252, // ACMTCM
    91.26810,  3.16210, 1.78964, 1.66200, // ACMTCT
    95.41650,  2.63830, 1.26908, 0.98862, // ACMTGA
    92.07510,  2.63830, 1.41720, 1.16665, // ACMTGC
    94.01390,  2.63830, 1.29676, 1.02113, // ACMTGG
    92.07510,  2.63830, 1.41720, 1.16665, // ACMTGM
    93.17520,  2.63830, 1.44582, 1.20217, // ACMTGT
    91.24900,  3.16210, 1.71292, 1.55628, // ACMTMA
    90.14290,  3.16210, 1.86109, 1.76252, // ACMTMC
    91.39750,  3.16210, 1.62527, 1.43836, // ACMTMG
    90.14290,  3.16210, 1.86109, 1.76252, // ACMTMM
    91.26810,  3.16210, 1.78964, 1.66200, // ACMTMT
    86.81060,  2.91311, 1.71890, 1.48956, // ACMTTA
    83.39010,  2.91311, 1.70994, 1.47793, // ACMTTC
    85.81540,  2.91311, 1.56463, 1.29360, // ACMTTG
    83.39010,  2.91311, 1.70994, 1.47793, // ACMTTM
    84.69540,  2.91311, 1.58306, 1.31652, // ACMTTT
    109.78300, 2.34971, 1.78499, 1.34274, // ACTAAA
    107.24400, 2.55557, 1.77711, 1.33385, // ACTAAC
    108.82800, 2.38834, 1.82866, 1.39231, // ACTAAG
    108.45200, 2.59718, 1.77711, 1.33385, // ACTAAM
    108.93000, 2.64054, 1.98981, 1.58036, // ACTAAT
    105.25900, 2.30570, 1.53278, 1.06551, // ACTACA
    104.75700, 2.44284, 1.69406, 1.23803, // ACTACC
    104.60200, 2.58253, 1.50915, 1.04096, // ACTACG
    105.93300, 2.42999, 1.69406, 1.23803, // ACTACM
    106.05900, 2.54810, 1.81195, 1.36948, // ACTACT
    107.86200, 2.27067, 1.83695, 1.55379, // ACTAGA
    105.70100, 2.45496, 1.61234, 1.27770, // ACTAGC
    106.73600, 2.30273, 1.53484, 1.18670, // ACTAGG
    106.95900, 2.38882, 1.61234, 1.27770, // ACTAGM
    106.96000, 2.55356, 1.60849, 1.27313, // ACTAGT
    104.91300, 3.58253, 1.53278, 1.06551, // ACTAMA
    104.37800, 3.58253, 1.69406, 1.23803, // ACTAMC
    112.46700, 2.37281, 1.50915, 1.04096, // ACTAMG
    104.37800, 3.58253, 1.69406, 1.23803, // ACTAMM
    105.78100, 3.58253, 1.81195, 1.36948, // ACTAMT
    105.22900, 2.61914, 1.91641, 1.40773, // ACTATA
    105.69100, 2.80131, 2.02830, 1.53279, // ACTATC
    104.99800, 2.68636, 1.87534, 1.36271, // ACTATG
    106.06400, 2.80881, 2.02830, 1.53279, // ACTATM
    107.23000, 2.94545, 2.25275, 1.79414, // ACTATT
    115.57300, 2.34555, 1.81878, 1.59109, // ACTCAA
    113.01100, 2.37488, 1.69623, 1.43303, // ACTCAC
    114.80200, 2.33073, 1.82823, 1.60352, // ACTCAG
    114.51100, 2.14004, 1.69623, 1.43303, // ACTCAM
    114.24600, 2.37100, 1.94818, 1.76388, // ACTCAT
    114.11300, 2.38294, 1.82491, 1.51937, // ACTCCA
    112.29700, 2.37200, 1.74480, 1.42044, // ACTCCC
    113.64900, 2.47041, 1.84678, 1.54677, // ACTCCG
    113.52000, 2.26124, 1.74480, 1.42044, // ACTCCM
    113.50500, 2.39035, 1.95376, 1.68309, // ACTCCT
    113.92000, 2.49891, 1.79737, 1.48675, // ACTCGA
    111.63700, 2.49891, 1.85362, 1.55710, // ACTCGC
    114.07000, 2.49891, 1.98026, 1.71936, // ACTCGG
    111.63700, 3.49891, 1.85362, 1.55710, // ACTCGM
    113.36900, 2.49891, 2.00748, 1.75494, // ACTCGT
    113.71500, 3.47041, 1.82491, 1.51937, // ACTCMA
    111.72700, 3.47041, 1.74480, 1.42044, // ACTCMC
    115.48500, 2.15020, 1.84678, 1.54677, // ACTCMG
    111.72700, 3.47041, 1.74480, 1.42044, // ACTCMM
    112.87600, 3.47041, 1.95376, 1.68309, // ACTCMT
    113.71800, 2.42523, 1.82568, 1.50849, // ACTCTA
    112.53000, 2.52100, 1.99910, 1.72846, // ACTCTC
    113.40300, 2.50815, 1.94200, 1.65494, // ACTCTG
    113.55400, 2.46574, 1.99910, 1.72846, // ACTCTM
    113.74900, 2.59421, 2.31939, 2.16006, // ACTCTT
    104.85900, 2.37043, 1.75335, 1.35976, // ACTGAA
    101.53700, 2.12286, 1.64828, 1.23939, // ACTGAC
    104.40800, 2.28515, 1.72515, 1.32709, // ACTGAG
    103.02600, 2.35379, 1.64828, 1.23939, // ACTGAM
    103.08900, 2.43721, 1.82084, 1.43903, // ACTGAT
    103.43200, 2.30692, 1.57276, 1.10065, // ACTGCA
    102.40000, 2.39926, 1.62044, 1.15109, // ACTGCC
    102.89100, 2.56584, 1.64559, 1.17798, // ACTGCG
    103.14400, 2.38000, 1.62044, 1.15109, // ACTGCM
    104.55700, 2.49972, 1.76473, 1.30821, // ACTGCT
    105.34800, 2.23986, 1.75997, 1.38612, // ACTGGA
    103.86500, 2.50392, 1.64528, 1.25286, // ACTGGC
    104.80900, 2.20888, 1.73224, 1.35349, // ACTGGG
    103.87100, 2.35243, 1.64528, 1.25286, // ACTGGM
    105.04900, 2.52154, 1.86940, 1.51738, // ACTGGT
    102.97600, 3.56584, 1.57276, 1.10065, // ACTGMA
    102.18700, 3.56584, 1.62044, 1.15109, // ACTGMC
    108.74300, 2.48401, 1.64559, 1.17798, // ACTGMG
    102.18700, 3.56584, 1.62044, 1.15109, // ACTGMM
    103.88800, 3.56584, 1.76473, 1.30821, // ACTGMT
    103.58800, 2.63846, 1.91169, 1.39461, // ACTGTA
    103.23200, 2.71686, 2.04214, 1.53976, // ACTGTC
    103.38100, 2.74016, 1.91520, 1.39845, // ACTGTG
    103.58500, 2.86438, 2.04214, 1.53976, // ACTGTM
    105.46400, 2.96733, 2.23889, 1.76757, // ACTGTT
    115.09200, 3.42793, 1.81878, 1.59109, // ACTMAA
    112.57800, 3.42793, 1.69623, 1.43303, // ACTMAC
    114.35000, 3.42793, 1.82823, 1.60352, // ACTMAG
    112.57800, 3.42793, 1.69623, 1.43303, // ACTMAM
    113.79400, 3.42793, 1.94818, 1.76388, // ACTMAT
    113.71500, 3.47041, 1.82491, 1.51937, // ACTMCA
    111.72700, 3.47041, 1.74480, 1.42044, // ACTMCC
    113.64900, 3.47041, 1.84678, 1.54677, // ACTMCG
    111.72700, 3.47041, 1.74480, 1.42044, // ACTMCM
    112.87600, 3.47041, 1.95376, 1.68309, // ACTMCT
    109.33400, 2.29212, 1.79737, 1.48675, // ACTMGA
    107.14500, 2.45813, 1.85362, 1.55710, // ACTMGC
    108.48400, 2.24137, 1.98026, 1.71936, // ACTMGG
    108.33400, 2.48398, 1.85362, 1.55710, // ACTMGM
    108.43500, 2.49412, 2.00748, 1.75494, // ACTMGT
    113.71500, 3.47041, 1.82491, 1.51937, // ACTMMA
    111.72700, 3.47041, 1.74480, 1.42044, // ACTMMC
    113.64900, 3.47041, 1.84678, 1.54677, // ACTMMG
    111.72700, 3.47041, 1.74480, 1.42044, // ACTMMM
    112.87600, 3.47041, 1.95376, 1.68309, // ACTMMT
    113.72600, 3.72204, 1.82568, 1.50849, // ACTMTA
    111.73300, 3.72204, 1.99910, 1.72846, // ACTMTC
    113.20400, 3.72204, 1.94200, 1.65494, // ACTMTG
    111.73300, 3.72204, 1.99910, 1.72846, // ACTMTM
    112.99600, 3.72204, 2.31939, 2.16006, // ACTMTT
    104.19200, 1.62580, 1.35553, 0.99397, // ACTTAA
    99.91900,  1.55130, 1.25649, 0.88704, // ACTTAC
    103.96100, 1.80822, 1.31032, 0.94465, // ACTTAG
    101.00500, 1.71694, 1.25649, 0.88704, // ACTTAM
    101.89900, 1.54951, 1.34247, 0.97963, // ACTTAT
    99.14580,  1.86639, 1.45211, 1.11851, // ACTTCA
    96.82950,  1.76953, 1.44106, 1.10576, // ACTTCC
    98.39870,  1.79286, 1.42524, 1.08761, // ACTTCG
    97.79270,  1.74843, 1.44106, 1.10576, // ACTTCM
    98.56330,  1.74151, 1.42580, 1.08825, // ACTTCT
    102.59800, 1.73316, 1.27431, 0.94949, // ACTTGA
    97.82180,  1.56435, 1.25882, 0.93224, // ACTTGC
    101.50200, 1.66124, 1.36817, 1.05630, // ACTTGG
    99.36560,  1.69955, 1.25882, 0.93224, // ACTTGM
    99.67910,  1.59024, 1.30757, 0.98691, // ACTTGT
    98.45000,  2.79285, 1.45211, 1.11851, // ACTTMA
    96.66330,  2.79285, 1.44106, 1.10576, // ACTTMC
    103.01900, 1.99369, 1.42524, 1.08761, // ACTTMG
    96.66330,  2.79285, 1.44106, 1.10576, // ACTTMM
    98.02970,  2.79285, 1.42580, 1.08825, // ACTTMT
    94.39440,  1.81229, 1.23750, 0.84797, // ACTTTA
    91.71020,  1.90912, 1.20209, 0.81183, // ACTTTC
    93.75920,  1.98399, 1.22881, 0.83906, // ACTTTG
    93.62700,  1.73961, 1.20209, 0.81183, // ACTTTM
    94.07210,  1.79575, 1.23664, 0.84708, // ACTTTT
    80.77460,  1.61601, 1.08810, 0.72343, // AGAAAA
    78.03040,  1.25223, 1.04629, 0.68214, // AGAAAC
    78.03940,  1.25195, 1.07330, 0.70873, // AGAAAG
    79.39170,  1.35538, 1.04629, 0.68214, // AGAAAM
    77.39670,  1.38754, 1.05928, 0.69488, // AGAAAT
    69.43410,  2.06004, 1.55604, 1.21560, // AGAACA
    68.39880,  1.94311, 1.55865, 1.21866, // AGAACC
    69.54790,  2.18144, 1.56447, 1.22549, // AGAACG
    69.02990,  2.04847, 1.55865, 1.21866, // AGAACM
    69.51490,  1.97967, 1.42127, 1.06115, // AGAACT
    76.70920,  1.30552, 1.10042, 0.74694, // AGAAGA
    77.34400,  1.20208, 0.97460, 0.62257, // AGAAGC
    76.53900,  1.23122, 1.09712, 0.74359, // AGAAGG
    76.45110,  1.77630, 0.97460, 0.62257, // AGAAGM
    76.86510,  1.08468, 1.00034, 0.64740, // AGAAGT
    69.45980,  3.18144, 1.55604, 1.21560, // AGAAMA
    68.24820,  3.18144, 1.55865, 1.21866, // AGAAMC
    74.10360,  1.47749, 1.56447, 1.22549, // AGAAMG
    68.24820,  3.18144, 1.55865, 1.21866, // AGAAMM
    69.47770,  3.18144, 1.42127, 1.06115, // AGAAMT
    69.68920,  1.80245, 1.32986, 0.97283, // AGAATA
    68.05090,  1.59128, 1.23796, 0.87374, // AGAATC
    68.87000,  1.72914, 1.25434, 0.89114, // AGAATG
    69.07190,  1.65810, 1.23796, 0.87374, // AGAATM
    69.37080,  1.46764, 1.13737, 0.76944, // AGAATT
    84.59880,  2.68649, 1.78971, 1.65509, // AGACAA
    83.26610,  2.46321, 1.60586, 1.40672, // AGACAC
    83.78870,  2.45872, 1.63755, 1.44857, // AGACAG
    84.11280,  2.48536, 1.60586, 1.40672, // AGACAM
    84.01460,  2.56035, 1.56983, 1.35965, // AGACAT
    80.60550,  2.34811, 1.83649, 1.83058, // AGACCA
    79.58990,  2.27263, 1.62768, 1.52742, // AGACCC
    80.75780,  2.19720, 1.78318, 1.75145, // AGACCG
    80.36080,  2.37529, 1.62768, 1.52742, // AGACCM
    80.55980,  2.37363, 1.72728, 1.66974, // AGACCT
    83.75340,  1.98191, 1.62836, 1.52196, // AGACGA
    82.32480,  1.98191, 1.50701, 1.35505, // AGACGC
    82.85510,  1.98191, 1.60290, 1.48640, // AGACGG
    82.32480,  2.98191, 1.50701, 1.35505, // AGACGM
    82.61910,  1.98191, 1.57058, 1.44168, // AGACGT
    80.71250,  3.19720, 1.83649, 1.83058, // AGACMA
    79.65680,  3.19720, 1.62768, 1.52742, // AGACMC
    81.52470,  2.18436, 1.78318, 1.75145, // AGACMG
    79.65680,  3.19720, 1.62768, 1.52742, // AGACMM
    80.84230,  3.19720, 1.72728, 1.66974, // AGACMT
    80.11000,  2.06602, 1.50382, 1.39992, // AGACTA
    78.73050,  1.73290, 1.41325, 1.27537, // AGACTC
    79.69940,  1.99796, 1.57927, 1.50658, // AGACTG
    79.66600,  1.96581, 1.41325, 1.27537, // AGACTM
    79.29250,  1.71056, 1.19775, 0.99508, // AGACTT
    79.95630,  1.24451, 1.00120, 0.70704, // AGAGAA
    75.98190,  1.25919, 1.16624, 0.88889, // AGAGAC
    78.76720,  1.21596, 1.03765, 0.74601, // AGAGAG
    76.72750,  1.70369, 1.16624, 0.88889, // AGAGAM
    77.09820,  1.25486, 1.16682, 0.88955, // AGAGAT
    72.79900,  1.46411, 1.21805, 0.86020, // AGAGCA
    71.51790,  1.50821, 1.18467, 0.82508, // AGAGCC
    72.79340,  1.59099, 1.12391, 0.76242, // AGAGCG
    72.20920,  1.52980, 1.18467, 0.82508, // AGAGCM
    72.61680,  1.41546, 1.10945, 0.74776, // AGAGCT
    79.15500,  1.21860, 0.98338, 0.67166, // AGAGGA
    77.01930,  1.13649, 1.08303, 0.77629, // AGAGGC
    78.55130,  1.14321, 0.97898, 0.66715, // AGAGGG
    77.69610,  0.99677, 1.08303, 0.77629, // AGAGGM
    77.54070,  1.18432, 1.08809, 0.78174, // AGAGGT
    72.78260,  2.59098, 1.21805, 0.86020, // AGAGMA
    71.45410,  2.59098, 1.18467, 0.82508, // AGAGMC
    76.57130,  1.16291, 1.12391, 0.76242, // AGAGMG
    71.45410,  2.59098, 1.18467, 0.82508, // AGAGMM
    72.84450,  2.59098, 1.10945, 0.74776, // AGAGMT
    72.36470,  1.15386, 0.94952, 0.63402, // AGAGTA
    71.01430,  1.32956, 1.12795, 0.82087, // AGAGTC
    72.25030,  1.13848, 1.04330, 0.73022, // AGAGTG
    72.10410,  1.11941, 1.12795, 0.82087, // AGAGTM
    72.11180,  1.36813, 1.21518, 0.91792, // AGAGTT
    84.95640,  3.48599, 1.78971, 1.65509, // AGAMAA
    83.40050,  3.48599, 1.60586, 1.40672, // AGAMAC
    84.01270,  3.48599, 1.63755, 1.44857, // AGAMAG
    83.40050,  3.48599, 1.60586, 1.40672, // AGAMAM
    84.30640,  3.48599, 1.56983, 1.35965, // AGAMAT
    80.71250,  3.19720, 1.83649, 1.83058, // AGAMCA
    79.65680,  3.19720, 1.62768, 1.52742, // AGAMCC
    80.75780,  3.19720, 1.78318, 1.75145, // AGAMCG
    79.65680,  3.19720, 1.62768, 1.52742, // AGAMCM
    80.84230,  3.19720, 1.72728, 1.66974, // AGAMCT
    81.89060,  1.72411, 1.62836, 1.52196, // AGAMGA
    80.32840,  1.44426, 1.50701, 1.35505, // AGAMGC
    80.86340,  1.54584, 1.60290, 1.48640, // AGAMGG
    80.90710,  1.49181, 1.50701, 1.35505, // AGAMGM
    80.53870,  1.46788, 1.57058, 1.44168, // AGAMGT
    80.71250,  3.19720, 1.83649, 1.83058, // AGAMMA
    79.65680,  3.19720, 1.62768, 1.52742, // AGAMMC
    80.75780,  3.19720, 1.78318, 1.75145, // AGAMMG
    79.65680,  3.19720, 1.62768, 1.52742, // AGAMMM
    80.84230,  3.19720, 1.72728, 1.66974, // AGAMMT
    80.39250,  2.92945, 1.50382, 1.39992, // AGAMTA
    79.03190,  2.92945, 1.41325, 1.27537, // AGAMTC
    79.92160,  2.92945, 1.57927, 1.50658, // AGAMTG
    79.03190,  2.92945, 1.41325, 1.27537, // AGAMTM
    79.45910,  2.92945, 1.19775, 0.99508, // AGAMTT
    77.55740,  2.31104, 2.01072, 2.00564, // AGATAA
    75.60150,  2.19926, 1.89705, 1.83800, // AGATAC
    77.15460,  2.00974, 1.87561, 1.80693, // AGATAG
    76.02580,  2.35391, 1.89705, 1.83800, // AGATAM
    76.81870,  2.54810, 2.05402, 2.07077, // AGATAT
    71.34860,  2.35209, 2.29287, 2.70016, // AGATCA
    70.57040,  2.46993, 2.46108, 3.00267, // AGATCC
    71.48610,  2.70241, 2.29132, 2.69741, // AGATCG
    71.91080,  2.83580, 2.46108, 3.00267, // AGATCM
    71.61110,  2.71319, 2.62501, 3.30762, // AGATCT
    75.35540,  1.98275, 1.82321, 1.91469, // AGATGA
    72.88750,  1.89292, 1.87844, 2.00234, // AGATGC
    74.42240,  1.91668, 1.80686, 1.88899, // AGATGG
    73.95440,  2.28022, 1.87844, 2.00234, // AGATGM
    73.72110,  2.12711, 2.01780, 2.22926, // AGATGT
    71.31330,  3.70241, 2.29287, 2.70016, // AGATMA
    70.60060,  3.70241, 2.46108, 3.00267, // AGATMC
    73.36280,  1.89418, 2.29132, 2.69741, // AGATMG
    70.60060,  3.70241, 2.46108, 3.00267, // AGATMM
    72.29040,  3.70241, 2.62501, 3.30762, // AGATMT
    66.78600,  2.16310, 1.96410, 2.00682, // AGATTA
    64.31710,  1.97007, 1.82858, 1.80274, // AGATTC
    66.10640,  2.00653, 1.79933, 1.75965, // AGATTG
    65.92490,  2.22005, 1.82858, 1.80274, // AGATTM
    65.72070,  1.97184, 1.73700, 1.66902, // AGATTT
    104.60000, 2.49448, 1.79959, 1.48473, // AGCAAA
    102.65300, 2.11453, 1.92923, 1.64802, // AGCAAC
    104.90100, 2.11776, 1.84686, 1.54362, // AGCAAG
    103.78800, 2.01367, 1.92923, 1.64802, // AGCAAM
    103.64200, 2.30144, 1.99074, 1.72747, // AGCAAT
    96.74290,  1.59302, 1.22791, 0.85677, // AGCACA
    95.14010,  1.68527, 1.27845, 0.91020, // AGCACC
    96.42330,  1.72978, 1.25632, 0.88667, // AGCACG
    96.29170,  1.63783, 1.27845, 0.91020, // AGCACM
    97.01480,  1.83794, 1.45681, 1.10718, // AGCACT
    102.47100, 2.26135, 1.78239, 1.43898, // AGCAGA
    99.71470,  2.27212, 1.74636, 1.39557, // AGCAGC
    102.46100, 2.18402, 1.78162, 1.43805, // AGCAGG
    101.35900, 2.02192, 1.74636, 1.39557, // AGCAGM
    101.72400, 2.34716, 1.99085, 1.69867, // AGCAGT
    96.59160,  2.72979, 1.22791, 0.85677, // AGCAMA
    95.09220,  2.72979, 1.27845, 0.91020, // AGCAMC
    102.28400, 1.70084, 1.25632, 0.88667, // AGCAMG
    95.09220,  2.72979, 1.27845, 0.91020, // AGCAMM
    96.68610,  2.72979, 1.45681, 1.10718, // AGCAMT
    96.59520,  1.81252, 1.45939, 1.12231, // AGCATA
    94.67910,  2.20807, 1.62236, 1.31545, // AGCATC
    95.85660,  1.85685, 1.46005, 1.12307, // AGCATG
    96.25840,  1.92608, 1.62236, 1.31545, // AGCATM
    96.85290,  2.29043, 1.75748, 1.48317, // AGCATT
    106.74100, 1.93088, 1.47199, 1.26140, // AGCCAA
    103.87700, 1.99623, 1.52091, 1.32481, // AGCCAC
    105.72500, 2.00389, 1.49291, 1.28839, // AGCCAG
    104.64500, 1.88585, 1.52091, 1.32481, // AGCCAM
    105.61800, 2.14093, 1.69935, 1.56467, // AGCCAT
    103.37500, 1.76724, 1.35848, 1.13096, // AGCCCA
    100.99700, 1.64718, 1.26908, 1.02118, // AGCCCC
    102.46000, 1.76337, 1.33123, 1.09709, // AGCCCG
    101.84900, 1.51586, 1.26908, 1.02118, // AGCCCM
    102.09100, 1.85523, 1.45288, 1.25086, // AGCCCT
    104.92200, 1.93464, 1.56010, 1.39917, // AGCCGA
    102.30700, 1.93464, 1.55199, 1.38828, // AGCCGC
    103.88500, 1.93464, 1.55128, 1.38734, // AGCCGG
    102.30700, 2.93465, 1.55199, 1.38828, // AGCCGM
    103.44200, 1.93464, 1.70782, 1.60254, // AGCCGT
    102.61300, 2.76337, 1.35848, 1.13096, // AGCCMA
    100.70000, 2.76337, 1.26908, 1.02118, // AGCCMC
    103.82200, 1.77196, 1.33123, 1.09709, // AGCCMG
    100.70000, 2.76337, 1.26908, 1.02118, // AGCCMM
    101.79700, 2.76337, 1.45288, 1.25086, // AGCCMT
    101.66100, 1.86415, 1.38877, 1.05146, // AGCCTA
    99.62160,  1.94847, 1.52241, 1.20682, // AGCCTC
    100.44800, 2.03929, 1.46404, 1.13809, // AGCCTG
    100.86500, 1.69132, 1.52241, 1.20682, // AGCCTM
    100.52200, 2.00480, 1.67194, 1.38892, // AGCCTT
    101.13800, 2.24827, 1.89422, 1.65705, // AGCGAA
    96.90530,  2.24827, 1.75699, 1.48028, // AGCGAC
    100.51200, 2.24827, 1.70520, 1.41531, // AGCGAG
    96.90530,  3.24827, 1.75699, 1.48028, // AGCGAM
    98.86570,  2.24827, 1.95617, 1.73899, // AGCGAT
    95.93570,  1.86985, 1.45399, 1.10557, // AGCGCA
    94.43320,  1.86985, 1.44846, 1.09927, // AGCGCC
    95.92200,  1.86985, 1.45659, 1.10854, // AGCGCG
    94.43320,  2.86985, 1.44846, 1.09927, // AGCGCM
    96.01060,  1.86985, 1.61437, 1.29345, // AGCGCT
    100.98800, 2.16392, 1.91151, 1.70788, // AGCGGA
    98.00530,  2.16392, 1.72836, 1.46840, // AGCGGC
    100.15100, 2.16392, 1.76206, 1.51155, // AGCGGG
    98.00530,  3.16392, 1.72836, 1.46840, // AGCGGM
    99.63020,  2.16392, 1.90136, 1.69429, // AGCGGT
    95.93570,  2.86985, 1.45399, 1.10557, // AGCGMA
    94.43320,  2.86985, 1.44846, 1.09927, // AGCGMC
    95.92200,  2.86985, 1.45659, 1.10854, // AGCGMG
    94.43320,  2.86985, 1.44846, 1.09927, // AGCGMM
    96.01060,  2.86985, 1.61437, 1.29345, // AGCGMT
    95.73930,  2.46720, 1.74294, 1.44600, // AGCGTA
    94.22060,  2.46720, 1.81332, 1.53445, // AGCGTC
    95.20070,  2.46720, 1.81042, 1.53078, // AGCGTG
    94.22060,  3.46720, 1.81332, 1.53445, // AGCGTM
    96.12240,  2.46720, 2.09402, 1.90421, // AGCGTT
    106.23400, 2.96521, 1.47199, 1.26140, // AGCMAA
    103.37800, 2.96521, 1.52091, 1.32481, // AGCMAC
    105.35500, 2.96521, 1.49291, 1.28839, // AGCMAG
    103.37800, 2.96521, 1.52091, 1.32481, // AGCMAM
    104.70700, 2.96521, 1.69935, 1.56467, // AGCMAT
    102.61300, 2.76337, 1.35848, 1.13096, // AGCMCA
    100.70000, 2.76337, 1.26908, 1.02118, // AGCMCC
    102.46000, 2.76337, 1.33123, 1.09709, // AGCMCG
    100.70000, 2.76337, 1.26908, 1.02118, // AGCMCM
    101.79700, 2.76337, 1.45288, 1.25086, // AGCMCT
    102.09000, 1.99159, 1.56010, 1.39917, // AGCMGA
    98.86000,  1.98308, 1.55199, 1.38828, // AGCMGC
    100.51900, 1.68728, 1.55128, 1.38734, // AGCMGG
    100.16000, 1.82867, 1.55199, 1.38828, // AGCMGM
    99.97250,  1.98889, 1.70782, 1.60254, // AGCMGT
    102.61300, 2.76337, 1.35848, 1.13096, // AGCMMA
    100.70000, 2.76337, 1.26908, 1.02118, // AGCMMC
    102.46000, 2.76337, 1.33123, 1.09709, // AGCMMG
    100.70000, 2.76337, 1.26908, 1.02118, // AGCMMM
    101.79700, 2.76337, 1.45288, 1.25086, // AGCMMT
    101.54900, 2.85738, 1.38877, 1.05146, // AGCMTA
    99.30650,  2.85738, 1.52241, 1.20682, // AGCMTC
    100.90200, 2.85738, 1.46404, 1.13809, // AGCMTG
    99.30650,  2.85738, 1.52241, 1.20682, // AGCMTM
    100.45000, 2.85738, 1.67194, 1.38892, // AGCMTT
    99.90660,  1.89508, 1.67024, 1.36156, // AGCTAA
    96.57760,  1.64768, 1.47981, 1.13548, // AGCTAC
    98.75210,  1.73906, 1.58090, 1.25379, // AGCTAG
    97.57010,  1.79843, 1.47981, 1.13548, // AGCTAM
    97.95410,  1.74843, 1.53013, 1.19388, // AGCTAT
    91.83340,  1.86754, 1.67351, 1.42475, // AGCTCA
    90.09440,  1.84587, 1.74461, 1.51651, // AGCTCC
    91.84240,  2.01289, 1.60820, 1.34218, // AGCTCG
    90.86750,  1.92756, 1.74461, 1.51651, // AGCTCM
    91.51970,  1.90933, 1.71803, 1.48199, // AGCTCT
    96.96340,  1.79939, 1.51234, 1.20029, // AGCTGA
    93.39320,  1.51241, 1.37694, 1.04276, // AGCTGC
    96.54960,  1.94102, 1.47689, 1.15834, // AGCTGG
    94.86450,  1.59743, 1.37694, 1.04276, // AGCTGM
    94.68060,  1.57437, 1.45287, 1.13019, // AGCTGT
    91.71820,  3.01289, 1.67351, 1.42475, // AGCTMA
    89.79100,  3.01289, 1.74461, 1.51651, // AGCTMC
    94.59900,  1.59461, 1.60820, 1.34218, // AGCTMG
    89.79100,  3.01289, 1.74461, 1.51651, // AGCTMM
    91.37890,  3.01289, 1.71803, 1.48199, // AGCTMT
    87.36060,  1.59013, 1.41299, 1.10879, // AGCTTA
    84.53990,  1.42968, 1.24603, 0.91820, // AGCTTC
    86.83210,  1.52974, 1.29414, 0.97188, // AGCTTG
    85.98210,  1.40852, 1.24603, 0.91820, // AGCTTM
    85.97230,  1.43018, 1.10217, 0.76386, // AGCTTT
    72.52090,  1.59596, 1.22764, 0.79505, // AGGAAA
    70.19790,  1.61369, 1.35461, 0.92154, // AGGAAC
    71.16560,  1.69404, 1.29147, 0.85786, // AGGAAG
    71.42260,  1.88742, 1.35461, 0.92154, // AGGAAM
    70.81960,  1.68063, 1.32620, 0.89270, // AGGAAT
    61.33360,  2.19474, 1.84078, 1.43995, // AGGACA
    59.97460,  2.05489, 1.92056, 1.53457, // AGGACC
    60.98800,  2.32589, 1.93153, 1.54774, // AGGACG
    61.10450,  2.11987, 1.92056, 1.53457, // AGGACM
    61.43000,  2.07663, 1.71638, 1.29648, // AGGACT
    70.63940,  1.64640, 1.29856, 0.93297, // AGGAGA
    69.93990,  1.78331, 1.23683, 0.86724, // AGGAGC
    69.52290,  1.73361, 1.36777, 1.00855, // AGGAGG
    69.03330,  1.53062, 1.23683, 0.86724, // AGGAGM
    69.80190,  1.92556, 1.21569, 0.84511, // AGGAGT
    60.91350,  3.32589, 1.84078, 1.43995, // AGGAMA
    60.08860,  3.32589, 1.92056, 1.53457, // AGGAMC
    66.56770,  1.90654, 1.93153, 1.54774, // AGGAMG
    60.08860,  3.32589, 1.92056, 1.53457, // AGGAMM
    61.03660,  3.32589, 1.71638, 1.29648, // AGGAMT
    61.78440,  1.96490, 1.73037, 1.42446, // AGGATA
    60.53880,  1.71651, 1.55428, 1.21266, // AGGATC
    61.12020,  1.91372, 1.60928, 1.27759, // AGGATG
    61.59370,  1.88278, 1.55428, 1.21266, // AGGATM
    61.35540,  1.50254, 1.28752, 0.91427, // AGGATT
    83.44640,  2.22558, 1.60776, 1.39868, // AGGCAA
    81.43600,  1.92531, 1.32000, 1.04051, // AGGCAC
    82.50950,  1.96410, 1.56674, 1.34549, // AGGCAG
    82.32890,  2.08878, 1.32000, 1.04051, // AGGCAM
    82.58060,  2.06234, 1.36378, 1.09271, // AGGCAT
    79.10000,  1.90818, 1.60340, 1.43590, // AGGCCA
    77.96350,  1.73195, 1.55874, 1.37633, // AGGCCC
    79.11550,  1.71082, 1.59689, 1.42717, // AGGCCG
    78.94340,  1.97090, 1.55874, 1.37633, // AGGCCM
    78.97110,  1.85852, 1.57318, 1.39549, // AGGCCT
    82.02140,  1.57550, 1.41475, 1.17987, // AGGCGA
    79.62120,  1.57550, 1.31215, 1.05387, // AGGCGC
    80.90570,  1.57550, 1.38856, 1.14727, // AGGCGG
    79.62120,  2.57551, 1.31215, 1.05387, // AGGCGM
    80.29600,  1.57550, 1.33347, 1.07967, // AGGCGT
    79.25190,  2.71082, 1.60340, 1.43590, // AGGCMA
    78.17080,  2.71082, 1.55874, 1.37633, // AGGCMC
    80.39750,  1.69061, 1.59689, 1.42717, // AGGCMG
    78.17080,  2.71082, 1.55874, 1.37633, // AGGCMM
    79.05210,  2.71082, 1.57318, 1.39549, // AGGCMT
    78.27730,  1.66094, 1.42184, 1.16218, // AGGCTA
    76.39650,  1.35542, 1.29132, 1.00588, // AGGCTC
    77.33560,  1.63247, 1.36160, 1.08912, // AGGCTG
    77.53390,  1.54527, 1.29132, 1.00588, // AGGCTM
    77.22330,  1.39363, 1.09321, 0.78353, // AGGCTT
    75.28270,  1.35562, 1.10878, 0.72685, // AGGGAA
    70.86340,  1.64710, 1.37412, 1.00281, // AGGGAC
    74.11740,  1.49957, 1.08724, 0.70578, // AGGGAG
    72.08810,  1.59464, 1.37412, 1.00281, // AGGGAM
    72.37920,  1.67239, 1.29842, 0.92109, // AGGGAT
    68.60520,  1.73964, 1.44825, 1.06143, // AGGGCA
    67.19310,  1.71806, 1.46362, 1.07838, // AGGGCC
    68.28450,  1.76808, 1.41889, 1.02932, // AGGGCG
    68.07420,  1.80891, 1.46362, 1.07838, // AGGGCM
    68.40860,  1.66522, 1.37433, 0.98121, // AGGGCT
    75.61360,  1.11496, 1.02860, 0.69371, // AGGGGA
    73.54700,  1.33171, 1.10436, 0.77175, // AGGGGC
    75.05510,  1.11029, 0.95862, 0.62413, // AGGGGG
    74.39280,  1.18606, 1.10436, 0.77175, // AGGGGM
    74.12850,  1.40144, 1.10899, 0.77661, // AGGGGT
    68.22510,  2.76808, 1.44825, 1.06143, // AGGGMA
    66.74490,  2.76808, 1.46362, 1.07838, // AGGGMC
    72.89370,  1.25263, 1.41889, 1.02932, // AGGGMG
    66.74490,  2.76808, 1.46362, 1.07838, // AGGGMM
    68.07770,  2.76808, 1.37433, 0.98121, // AGGGMT
    68.31910,  1.40148, 1.13416, 0.81137, // AGGGTA
    66.82080,  1.60396, 1.24686, 0.93526, // AGGGTC
    67.54070,  1.33267, 1.12686, 0.80355, // AGGGTG
    67.03170,  1.33557, 1.24686, 0.93526, // AGGGTM
    67.84770,  1.66168, 1.35120, 1.05508, // AGGGTT
    83.65620,  2.89614, 1.60776, 1.39868, // AGGMAA
    81.50610,  2.89614, 1.32000, 1.04051, // AGGMAC
    82.56150,  2.89614, 1.56674, 1.34549, // AGGMAG
    81.50610,  2.89614, 1.32000, 1.04051, // AGGMAM
    82.51460,  2.89614, 1.36378, 1.09271, // AGGMAT
    79.25190,  2.71082, 1.60340, 1.43590, // AGGMCA
    78.17080,  2.71082, 1.55874, 1.37633, // AGGMCC
    79.11550,  2.71082, 1.59689, 1.42717, // AGGMCG
    78.17080,  2.71082, 1.55874, 1.37633, // AGGMCM
    79.05210,  2.71082, 1.57318, 1.39549, // AGGMCT
    78.58270,  1.31119, 1.41475, 1.17987, // AGGMGA
    76.36130,  1.20973, 1.31215, 1.05387, // AGGMGC
    76.72900,  1.03279, 1.38856, 1.14727, // AGGMGG
    76.97750,  1.19214, 1.31215, 1.05387, // AGGMGM
    76.33300,  1.01059, 1.33347, 1.07967, // AGGMGT
    79.25190,  2.71082, 1.60340, 1.43590, // AGGMMA
    78.17080,  2.71082, 1.55874, 1.37633, // AGGMMC
    79.11550,  2.71082, 1.59689, 1.42717, // AGGMMG
    78.17080,  2.71082, 1.55874, 1.37633, // AGGMMM
    79.05210,  2.71082, 1.57318, 1.39549, // AGGMMT
    78.40380,  2.47431, 1.42184, 1.16218, // AGGMTA
    76.46660,  2.47431, 1.29132, 1.00588, // AGGMTC
    77.72340,  2.47431, 1.36160, 1.08912, // AGGMTG
    76.46660,  2.47431, 1.29132, 1.00588, // AGGMTM
    77.21190,  2.47431, 1.09321, 0.78353, // AGGMTT
    74.12460,  2.14320, 1.94833, 1.80553, // AGGTAA
    70.49650,  2.10188, 1.70329, 1.47585, // AGGTAC
    73.03780,  2.08012, 1.90831, 1.75019, // AGGTAG
    72.63350,  2.80740, 1.70329, 1.47585, // AGGTAM
    72.15500,  2.62528, 1.90007, 1.73887, // AGGTAT
    66.23130,  2.16460, 2.00760, 2.12192, // AGGTCA
    65.18130,  2.31619, 2.16087, 2.36949, // AGGTCC
    66.18780,  2.69199, 1.91538, 1.97741, // AGGTCG
    66.29010,  2.61292, 2.16087, 2.36949, // AGGTCM
    66.23210,  2.47389, 2.21170, 2.45359, // AGGTCT
    69.82080,  1.88434, 1.71347, 1.59926, // AGGTGA
    66.99640,  1.83637, 1.77829, 1.69087, // AGGTGC
    68.75190,  1.66582, 1.62552, 1.47772, // AGGTGG
    68.47880,  2.22438, 1.77829, 1.69087, // AGGTGM
    67.87910,  1.90866, 1.79891, 1.72035, // AGGTGT
    66.21880,  3.69199, 2.00760, 2.12192, // AGGTMA
    65.06780,  3.69199, 2.16087, 2.36949, // AGGTMC
    68.75110,  1.80898, 1.91538, 1.97741, // AGGTMG
    65.06780,  3.69199, 2.16087, 2.36949, // AGGTMM
    66.28900,  3.69199, 2.21170, 2.45359, // AGGTMT
    62.32690,  2.45145, 2.07388, 1.97327, // AGGTTA
    59.61540,  2.35111, 2.30480, 2.31187, // AGGTTC
    61.74900,  2.46466, 2.01714, 1.89285, // AGGTTG
    61.70210,  2.65663, 2.30480, 2.31187, // AGGTTM
    60.87770,  2.36181, 2.00778, 1.87969, // AGGTTT
    105.59100, 3.21768, 1.79959, 1.48473, // AGMAAA
    102.46000, 3.21768, 1.92923, 1.64802, // AGMAAC
    104.51200, 3.21768, 1.84686, 1.54362, // AGMAAG
    102.46000, 3.21768, 1.92923, 1.64802, // AGMAAM
    103.80200, 3.21768, 1.99074, 1.72747, // AGMAAT
    96.59160,  2.72979, 1.22791, 0.85677, // AGMACA
    95.09220,  2.72979, 1.27845, 0.91020, // AGMACC
    96.42330,  2.72979, 1.25632, 0.88667, // AGMACG
    95.09220,  2.72979, 1.27845, 0.91020, // AGMACM
    96.68610,  2.72979, 1.45681, 1.10718, // AGMACT
    102.73200, 3.29394, 1.78239, 1.43898, // AGMAGA
    99.73500,  3.29394, 1.74636, 1.39557, // AGMAGC
    101.49300, 3.29394, 1.78162, 1.43805, // AGMAGG
    99.73500,  3.29394, 1.74636, 1.39557, // AGMAGM
    101.01800, 3.29394, 1.99085, 1.69867, // AGMAGT
    96.59160,  2.72979, 1.22791, 0.85677, // AGMAMA
    95.09220,  2.72979, 1.27845, 0.91020, // AGMAMC
    96.42330,  2.72979, 1.25632, 0.88667, // AGMAMG
    95.09220,  2.72979, 1.27845, 0.91020, // AGMAMM
    96.68610,  2.72979, 1.45681, 1.10718, // AGMAMT
    96.08070,  3.05386, 1.45939, 1.12231, // AGMATA
    94.57680,  3.05386, 1.62236, 1.31545, // AGMATC
    95.46810,  3.05386, 1.46005, 1.12307, // AGMATG
    94.57680,  3.05386, 1.62236, 1.31545, // AGMATM
    96.14380,  3.05386, 1.75748, 1.48317, // AGMATT
    106.23400, 2.96521, 1.47199, 1.26140, // AGMCAA
    103.37800, 2.96521, 1.52091, 1.32481, // AGMCAC
    105.35500, 2.96521, 1.49291, 1.28839, // AGMCAG
    103.37800, 2.96521, 1.52091, 1.32481, // AGMCAM
    104.70700, 2.96521, 1.69935, 1.56467, // AGMCAT
    102.61300, 2.76337, 1.35848, 1.13096, // AGMCCA
    100.70000, 2.76337, 1.26908, 1.02118, // AGMCCC
    102.46000, 2.76337, 1.33123, 1.09709, // AGMCCG
    100.70000, 2.76337, 1.26908, 1.02118, // AGMCCM
    101.79700, 2.76337, 1.45288, 1.25086, // AGMCCT
    104.92200, 2.93465, 1.56010, 1.39917, // AGMCGA
    102.30700, 2.93465, 1.55199, 1.38828, // AGMCGC
    103.88500, 2.93465, 1.55128, 1.38734, // AGMCGG
    102.30700, 2.93465, 1.55199, 1.38828, // AGMCGM
    103.44200, 2.93465, 1.70782, 1.60254, // AGMCGT
    102.61300, 2.76337, 1.35848, 1.13096, // AGMCMA
    100.70000, 2.76337, 1.26908, 1.02118, // AGMCMC
    102.46000, 2.76337, 1.33123, 1.09709, // AGMCMG
    100.70000, 2.76337, 1.26908, 1.02118, // AGMCMM
    101.79700, 2.76337, 1.45288, 1.25086, // AGMCMT
    101.54900, 2.85738, 1.38877, 1.05146, // AGMCTA
    99.30650,  2.85738, 1.52241, 1.20682, // AGMCTC
    100.90200, 2.85738, 1.46404, 1.13809, // AGMCTG
    99.30650,  2.85738, 1.52241, 1.20682, // AGMCTM
    100.45000, 2.85738, 1.67194, 1.38892, // AGMCTT
    100.04000, 2.29615, 1.89422, 1.65705, // AGMGAA
    95.77460,  2.16006, 1.75699, 1.48028, // AGMGAC
    99.09840,  2.23036, 1.70520, 1.41531, // AGMGAG
    97.50860,  2.11486, 1.75699, 1.48028, // AGMGAM
    98.11540,  2.27711, 1.95617, 1.73899, // AGMGAT
    93.94850,  2.08005, 1.45399, 1.10557, // AGMGCA
    93.08270,  1.93330, 1.44846, 1.09927, // AGMGCC
    95.92200,  2.86985, 1.45659, 1.10854, // AGMGCG
    93.18040,  2.24886, 1.44846, 1.09927, // AGMGCM
    94.07790,  2.13571, 1.61437, 1.29345, // AGMGCT
    99.97490,  2.37484, 1.91151, 1.70788, // AGMGGA
    97.09250,  2.22815, 1.72836, 1.46840, // AGMGGC
    99.27370,  2.20992, 1.76206, 1.51155, // AGMGGG
    98.19400,  2.15555, 1.72836, 1.46840, // AGMGGM
    98.60860,  2.35086, 1.90136, 1.69429, // AGMGGT
    95.93570,  2.86985, 1.45399, 1.10557, // AGMGMA
    94.43320,  2.86985, 1.44846, 1.09927, // AGMGMC
    98.47700,  1.86923, 1.45659, 1.10854, // AGMGMG
    94.43320,  2.86985, 1.44846, 1.09927, // AGMGMM
    96.01060,  2.86985, 1.61437, 1.29345, // AGMGMT
    93.95860,  2.28932, 1.74294, 1.44600, // AGMGTA
    92.71590,  2.40892, 1.81332, 1.53445, // AGMGTC
    93.98100,  2.25071, 1.81042, 1.53078, // AGMGTG
    93.41720,  2.42235, 1.81332, 1.53445, // AGMGTM
    94.68310,  2.35481, 2.09402, 1.90421, // AGMGTT
    106.23400, 2.96521, 1.47199, 1.26140, // AGMMAA
    103.37800, 2.96521, 1.52091, 1.32481, // AGMMAC
    105.35500, 2.96521, 1.49291, 1.28839, // AGMMAG
    103.37800, 2.96521, 1.52091, 1.32481, // AGMMAM
    104.70700, 2.96521, 1.69935, 1.56467, // AGMMAT
    102.61300, 2.76337, 1.35848, 1.13096, // AGMMCA
    100.70000, 2.76337, 1.26908, 1.02118, // AGMMCC
    102.46000, 2.76337, 1.33123, 1.09709, // AGMMCG
    100.70000, 2.76337, 1.26908, 1.02118, // AGMMCM
    101.79700, 2.76337, 1.45288, 1.25086, // AGMMCT
    104.92200, 2.93465, 1.56010, 1.39917, // AGMMGA
    102.30700, 2.93465, 1.55199, 1.38828, // AGMMGC
    103.88500, 2.93465, 1.55128, 1.38734, // AGMMGG
    102.30700, 2.93465, 1.55199, 1.38828, // AGMMGM
    103.44200, 2.93465, 1.70782, 1.60254, // AGMMGT
    102.61300, 2.76337, 1.35848, 1.13096, // AGMMMA
    100.70000, 2.76337, 1.26908, 1.02118, // AGMMMC
    102.46000, 2.76337, 1.33123, 1.09709, // AGMMMG
    100.70000, 2.76337, 1.26908, 1.02118, // AGMMMM
    101.79700, 2.76337, 1.45288, 1.25086, // AGMMMT
    101.54900, 2.85738, 1.38877, 1.05146, // AGMMTA
    99.30650,  2.85738, 1.52241, 1.20682, // AGMMTC
    100.90200, 2.85738, 1.46404, 1.13809, // AGMMTG
    99.30650,  2.85738, 1.52241, 1.20682, // AGMMTM
    100.45000, 2.85738, 1.67194, 1.38892, // AGMMTT
    99.63620,  2.80194, 1.67024, 1.36156, // AGMTAA
    96.45280,  2.80194, 1.47981, 1.13548, // AGMTAC
    98.46220,  2.80194, 1.58090, 1.25379, // AGMTAG
    96.45280,  2.80194, 1.47981, 1.13548, // AGMTAM
    97.76010,  2.80194, 1.53013, 1.19388, // AGMTAT
    91.71820,  3.01289, 1.67351, 1.42475, // AGMTCA
    89.79100,  3.01289, 1.74461, 1.51651, // AGMTCC
    91.84240,  3.01289, 1.60820, 1.34218, // AGMTCG
    89.79100,  3.01289, 1.74461, 1.51651, // AGMTCM
    91.37890,  3.01289, 1.71803, 1.48199, // AGMTCT
    96.88070,  2.67840, 1.51234, 1.20029, // AGMTGA
    93.46390,  2.67840, 1.37694, 1.04276, // AGMTGC
    95.56790,  2.67840, 1.47689, 1.15834, // AGMTGG
    93.46390,  2.67840, 1.37694, 1.04276, // AGMTGM
    94.55180,  2.67840, 1.45287, 1.13019, // AGMTGT
    91.71820,  3.01289, 1.67351, 1.42475, // AGMTMA
    89.79100,  3.01289, 1.74461, 1.51651, // AGMTMC
    91.84240,  3.01289, 1.60820, 1.34218, // AGMTMG
    89.79100,  3.01289, 1.74461, 1.51651, // AGMTMM
    91.37890,  3.01289, 1.71803, 1.48199, // AGMTMT
    87.36690,  2.51841, 1.41299, 1.10879, // AGMTTA
    84.42880,  2.51841, 1.24603, 0.91820, // AGMTTC
    86.85300,  2.51841, 1.29414, 0.97188, // AGMTTG
    84.42880,  2.51841, 1.24603, 0.91820, // AGMTTM
    85.66450,  2.51841, 1.10217, 0.76386, // AGMTTT
    107.15900, 2.90615, 2.47623, 2.00239, // AGTAAA
    104.94400, 2.85407, 2.48563, 2.01380, // AGTAAC
    106.30900, 2.80102, 2.45718, 1.97934, // AGTAAG
    106.63800, 2.84848, 2.48563, 2.01380, // AGTAAM
    107.29000, 2.94223, 2.62386, 2.18411, // AGTAAT
    100.75600, 2.65012, 2.08182, 1.59872, // AGTACA
    100.55700, 2.79735, 2.22721, 1.76909, // AGTACC
    100.47600, 2.66382, 2.06477, 1.57912, // AGTACG
    101.92000, 2.73676, 2.22721, 1.76909, // AGTACM
    102.18700, 2.88721, 2.15706, 1.68617, // AGTACT
    105.00800, 2.76526, 2.33354, 1.93877, // AGTAGA
    103.14100, 2.75369, 2.30493, 1.90322, // AGTAGC
    104.01400, 2.55472, 2.17747, 1.74755, // AGTAGG
    104.39000, 2.72730, 2.30493, 1.90322, // AGTAGM
    105.27300, 2.80991, 2.62265, 2.31000, // AGTAGT
    100.41600, 3.66382, 2.08182, 1.59872, // AGTAMA
    100.23200, 3.66382, 2.22721, 1.76909, // AGTAMC
    110.61700, 2.92742, 2.06477, 1.57912, // AGTAMG
    100.23200, 3.66382, 2.22721, 1.76909, // AGTAMM
    102.00700, 3.66382, 2.15706, 1.68617, // AGTAMT
    102.46200, 3.00910, 2.51775, 2.00755, // AGTATA
    103.63900, 3.12978, 2.62019, 2.13132, // AGTATC
    102.27900, 3.03665, 2.63292, 2.14686, // AGTATG
    104.11000, 3.17270, 2.62019, 2.13132, // AGTATM
    105.78200, 3.28528, 2.94594, 2.54087, // AGTATT
    116.02500, 2.99999, 2.71665, 2.89390, // AGTCAA
    113.86300, 2.87483, 2.60991, 2.72502, // AGTCAC
    115.66000, 2.90829, 2.59796, 2.70633, // AGTCAG
    116.27500, 2.52105, 2.60991, 2.72502, // AGTCAM
    114.77800, 2.95695, 2.71690, 2.89428, // AGTCAT
    114.36900, 3.08391, 2.50562, 2.55218, // AGTCCA
    112.29600, 3.06201, 2.44614, 2.46185, // AGTCCC
    114.07300, 2.92671, 2.68237, 2.82694, // AGTCCG
    114.14000, 2.57621, 2.44614, 2.46185, // AGTCCM
    113.79500, 2.91180, 2.51036, 2.55943, // AGTCCT
    115.51000, 2.90558, 2.77885, 2.95622, // AGTCGA
    112.77400, 2.90558, 2.53831, 2.58082, // AGTCGC
    114.52700, 2.90558, 2.71568, 2.85600, // AGTCGG
    112.77400, 3.90558, 2.53831, 2.58082, // AGTCGM
    114.77200, 2.90558, 2.74253, 2.89845, // AGTCGT
    114.16700, 3.92671, 2.50562, 2.55218, // AGTCMA
    112.41100, 3.92671, 2.44614, 2.46185, // AGTCMC
    116.64400, 2.49364, 2.68237, 2.82694, // AGTCMG
    112.41100, 3.92671, 2.44614, 2.46185, // AGTCMM
    113.52300, 3.92671, 2.51036, 2.55943, // AGTCMT
    114.52700, 2.95288, 3.10125, 3.31683, // AGTCTA
    113.38100, 2.91993, 2.68468, 2.67151, // AGTCTC
    114.34600, 2.94566, 2.58891, 2.52984, // AGTCTG
    114.75300, 2.77804, 2.68468, 2.67151, // AGTCTM
    114.55700, 3.04440, 2.94227, 3.06507, // AGTCTT
    105.13800, 2.79432, 2.47043, 2.23788, // AGTGAA
    101.39600, 2.55169, 2.55404, 2.35245, // AGTGAC
    103.93400, 2.75239, 2.39201, 2.13218, // AGTGAG
    103.36400, 2.56105, 2.55404, 2.35245, // AGTGAM
    103.56200, 2.74482, 2.70593, 2.56540, // AGTGAT
    102.97200, 2.91308, 2.24864, 1.91265, // AGTGCA
    102.16600, 2.90089, 2.26848, 1.93801, // AGTGCC
    102.89100, 2.92533, 2.37726, 2.07907, // AGTGCG
    103.43000, 2.70793, 2.26848, 1.93801, // AGTGCM
    104.52800, 3.00590, 2.53149, 2.28464, // AGTGCT
    105.35400, 2.87573, 2.45545, 2.24024, // AGTGGA
    103.32800, 2.67077, 2.44236, 2.22234, // AGTGGC
    104.77900, 2.79860, 2.31369, 2.04906, // AGTGGG
    104.65100, 2.55929, 2.44236, 2.22234, // AGTGGM
    105.93800, 2.78346, 2.67063, 2.54107, // AGTGGT
    102.74800, 3.92533, 2.24864, 1.91265, // AGTGMA
    102.19900, 3.92533, 2.26848, 1.93801, // AGTGMC
    111.53700, 2.81976, 2.37726, 2.07907, // AGTGMG
    102.19900, 3.92533, 2.26848, 1.93801, // AGTGMM
    104.54500, 3.92533, 2.53149, 2.28464, // AGTGMT
    105.42400, 3.15046, 2.78922, 2.54947, // AGTGTA
    106.02600, 3.15145, 2.88082, 2.67608, // AGTGTC
    105.54900, 3.06645, 2.73224, 2.47174, // AGTGTG
    106.15400, 3.17963, 2.88082, 2.67608, // AGTGTM
    108.34100, 3.33034, 3.12153, 3.01841, // AGTGTT
    115.74500, 3.78871, 2.71665, 2.89390, // AGTMAA
    113.45600, 3.78871, 2.60991, 2.72502, // AGTMAC
    115.02900, 3.78871, 2.59796, 2.70633, // AGTMAG
    113.45600, 3.78871, 2.60991, 2.72502, // AGTMAM
    114.52900, 3.78871, 2.71690, 2.89428, // AGTMAT
    114.16700, 3.92671, 2.50562, 2.55218, // AGTMCA
    112.41100, 3.92671, 2.44614, 2.46185, // AGTMCC
    114.07300, 3.92671, 2.68237, 2.82694, // AGTMCG
    112.41100, 3.92671, 2.44614, 2.46185, // AGTMCM
    113.52300, 3.92671, 2.51036, 2.55943, // AGTMCT
    108.83800, 2.96981, 2.77885, 2.95622, // AGTMGA
    106.82000, 2.85414, 2.53831, 2.58082, // AGTMGC
    107.68400, 2.74188, 2.71568, 2.85600, // AGTMGG
    108.33000, 2.85112, 2.53831, 2.58082, // AGTMGM
    108.32000, 2.87306, 2.74253, 2.89845, // AGTMGT
    114.16700, 3.92671, 2.50562, 2.55218, // AGTMMA
    112.41100, 3.92671, 2.44614, 2.46185, // AGTMMC
    114.07300, 3.92671, 2.68237, 2.82694, // AGTMMG
    112.41100, 3.92671, 2.44614, 2.46185, // AGTMMM
    113.52300, 3.92671, 2.51036, 2.55943, // AGTMMT
    113.82400, 4.02381, 3.10125, 3.31683, // AGTMTA
    113.10500, 4.02381, 2.68468, 2.67151, // AGTMTC
    114.17100, 4.02381, 2.58891, 2.52984, // AGTMTG
    113.10500, 4.02381, 2.68468, 2.67151, // AGTMTM
    114.23700, 4.02381, 2.94227, 3.06507, // AGTMTT
    103.71500, 2.78722, 2.07483, 2.08587, // AGTTAA
    100.66700, 2.65153, 1.81180, 1.70207, // AGTTAC
    103.61400, 2.60412, 1.86412, 1.77632, // AGTTAG
    101.11800, 2.47443, 1.81180, 1.70207, // AGTTAM
    102.51800, 2.19340, 1.83710, 1.73785, // AGTTAT
    97.94960,  2.35193, 2.19881, 2.08101, // AGTTCA
    95.58250,  2.18540, 2.26075, 2.16956, // AGTTCC
    97.34270,  2.21733, 2.18418, 2.06027, // AGTTCG
    96.88390,  1.97343, 2.26075, 2.16956, // AGTTCM
    97.31480,  2.16986, 2.11452, 1.96250, // AGTTCT
    102.13000, 2.34570, 1.90687, 1.75635, // AGTTGA
    97.31420,  2.01457, 1.86502, 1.69886, // AGTTGC
    100.91000, 2.86258, 1.90940, 1.75985, // AGTTGG
    99.14410,  1.83873, 1.86502, 1.69886, // AGTTGM
    98.94700,  2.32143, 1.88914, 1.73192, // AGTTGT
    97.29690,  3.21733, 2.19881, 2.08101, // AGTTMA
    95.36010,  3.21733, 2.26075, 2.16956, // AGTTMC
    101.53300, 1.93063, 2.18418, 2.06027, // AGTTMG
    95.36010,  3.21733, 2.26075, 2.16956, // AGTTMM
    97.15650,  3.21733, 2.11452, 1.96250, // AGTTMT
    93.75080,  1.87116, 1.91003, 1.76480, // AGTTTA
    91.27770,  1.71102, 1.73059, 1.52205, // AGTTTC
    92.71670,  1.86590, 1.88580, 1.73132, // AGTTTG
    93.10590,  1.68307, 1.73059, 1.52205, // AGTTTM
    93.71320,  1.83505, 1.53996, 1.27762, // AGTTTT
    92.71180,  3.70987, 1.59825, 1.19732, // AMAAAA
    89.28400,  3.70987, 1.81392, 1.44767, // AMAAAC
    91.93550,  3.70987, 1.69696, 1.30994, // AMAAAG
    89.28400,  3.70987, 1.81392, 1.44767, // AMAAAM
    90.32440,  3.70987, 1.85900, 1.50197, // AMAAAT
    83.13680,  3.16112, 1.51177, 1.20984, // AMAACA
    82.21650,  3.16112, 1.54321, 1.24778, // AMAACC
    83.18670,  3.16112, 1.54582, 1.25095, // AMAACG
    82.21650,  3.16112, 1.54321, 1.24778, // AMAACM
    82.76920,  3.16112, 1.49251, 1.18680, // AMAACT
    91.01500,  3.81796, 1.60187, 1.24600, // AMAAGA
    88.61560,  3.81796, 1.71362, 1.37863, // AMAAGC
    90.15780,  3.81796, 1.61613, 1.26266, // AMAAGG
    88.61560,  3.81796, 1.71362, 1.37863, // AMAAGM
    88.88450,  3.81796, 1.87238, 1.57459, // AMAAGT
    83.13680,  3.16112, 1.51177, 1.20984, // AMAAMA
    82.21650,  3.16112, 1.54321, 1.24778, // AMAAMC
    83.18670,  3.16112, 1.54582, 1.25095, // AMAAMG
    82.21650,  3.16112, 1.54321, 1.24778, // AMAAMM
    82.76920,  3.16112, 1.49251, 1.18680, // AMAAMT
    82.18980,  3.24890, 1.64608, 1.39736, // AMAATA
    80.65210,  3.24890, 1.97752, 1.83998, // AMAATC
    81.65630,  3.24890, 1.76664, 1.55366, // AMAATG
    80.65210,  3.24890, 1.97752, 1.83998, // AMAATM
    81.09840,  3.24890, 2.00648, 1.88054, // AMAATT
    94.96910,  2.22481, 1.05715, 0.80493, // AMACAA
    92.34910,  2.22481, 1.06102, 0.80936, // AMACAC
    94.13070,  2.22481, 1.04348, 0.78937, // AMACAG
    92.34910,  2.22481, 1.06102, 0.80936, // AMACAM
    93.39940,  2.22481, 0.97070, 0.70824, // AMACAT
    91.76720,  2.29044, 1.19393, 0.96929, // AMACCA
    90.32110,  2.29044, 1.15189, 0.91854, // AMACCC
    91.49950,  2.29044, 1.10979, 0.86865, // AMACCG
    90.32110,  2.29044, 1.15189, 0.91854, // AMACCM
    91.20300,  2.29044, 1.12044, 0.88119, // AMACCT
    94.08570,  2.39798, 1.00994, 0.73926, // AMACGA
    91.54810,  2.39798, 1.08903, 0.82779, // AMACGC
    93.27920,  2.39798, 1.05984, 0.79473, // AMACGG
    91.54810,  2.39798, 1.08903, 0.82779, // AMACGM
    92.34440,  2.39798, 1.07297, 0.80954, // AMACGT
    91.76720,  2.29044, 1.19393, 0.96929, // AMACMA
    90.32110,  2.29044, 1.15189, 0.91854, // AMACMC
    91.49950,  2.29044, 1.10979, 0.86865, // AMACMG
    90.32110,  2.29044, 1.15189, 0.91854, // AMACMM
    91.20300,  2.29044, 1.12044, 0.88119, // AMACMT
    90.18550,  2.56257, 1.12931, 0.82924, // AMACTA
    87.93340,  2.56257, 1.12747, 0.82720, // AMACTC
    89.42720,  2.56257, 1.12618, 0.82579, // AMACTG
    87.93340,  2.56257, 1.12747, 0.82720, // AMACTM
    89.02180,  2.56257, 1.09133, 0.78776, // AMACTT
    88.19820,  3.60118, 1.64646, 1.26460, // AMAGAA
    82.73940,  3.60118, 1.92949, 1.60431, // AMAGAC
    87.04280,  3.60118, 1.73953, 1.37333, // AMAGAG
    82.73940,  3.60118, 1.92949, 1.60431, // AMAGAM
    84.61650,  3.60118, 1.92427, 1.59781, // AMAGAT
    83.01820,  3.08236, 1.42329, 1.19082, // AMAGCA
    81.45030,  3.08236, 1.43105, 1.20057, // AMAGCC
    82.88880,  3.08236, 1.40829, 1.17205, // AMAGCG
    81.45030,  3.08236, 1.43105, 1.20057, // AMAGCM
    82.80060,  3.08236, 1.52849, 1.32526, // AMAGCT
    87.98540,  3.50147, 1.58798, 1.26495, // AMAGGA
    83.96550,  3.50147, 1.73486, 1.44444, // AMAGGC
    86.51230,  3.50147, 1.55245, 1.22273, // AMAGGG
    83.96550,  3.50147, 1.73486, 1.44444, // AMAGGM
    85.46980,  3.50147, 1.82887, 1.56343, // AMAGGT
    83.01820,  3.08236, 1.42329, 1.19082, // AMAGMA
    81.45030,  3.08236, 1.43105, 1.20057, // AMAGMC
    82.88880,  3.08236, 1.40829, 1.17205, // AMAGMG
    81.45030,  3.08236, 1.43105, 1.20057, // AMAGMM
    82.80060,  3.08236, 1.52849, 1.32526, // AMAGMT
    80.35500,  3.40625, 1.91397, 1.72963, // AMAGTA
    76.81660,  3.40625, 2.09972, 1.98743, // AMAGTC
    78.88420,  3.40625, 1.89939, 1.70991, // AMAGTG
    76.81660,  3.40625, 2.09972, 1.98743, // AMAGTM
    78.04570,  3.40625, 2.06168, 1.93366, // AMAGTT
    94.96910,  2.22481, 1.05715, 0.80493, // AMAMAA
    92.34910,  2.22481, 1.06102, 0.80936, // AMAMAC
    94.13070,  2.22481, 1.04348, 0.78937, // AMAMAG
    92.34910,  2.22481, 1.06102, 0.80936, // AMAMAM
    93.39940,  2.22481, 0.97070, 0.70824, // AMAMAT
    91.76720,  2.29044, 1.19393, 0.96929, // AMAMCA
    90.32110,  2.29044, 1.15189, 0.91854, // AMAMCC
    91.49950,  2.29044, 1.10979, 0.86865, // AMAMCG
    90.32110,  2.29044, 1.15189, 0.91854, // AMAMCM
    91.20300,  2.29044, 1.12044, 0.88119, // AMAMCT
    94.08570,  2.39798, 1.00994, 0.73926, // AMAMGA
    91.54810,  2.39798, 1.08903, 0.82779, // AMAMGC
    93.27920,  2.39798, 1.05984, 0.79473, // AMAMGG
    91.54810,  2.39798, 1.08903, 0.82779, // AMAMGM
    92.34440,  2.39798, 1.07297, 0.80954, // AMAMGT
    91.76720,  2.29044, 1.19393, 0.96929, // AMAMMA
    90.32110,  2.29044, 1.15189, 0.91854, // AMAMMC
    91.49950,  2.29044, 1.10979, 0.86865, // AMAMMG
    90.32110,  2.29044, 1.15189, 0.91854, // AMAMMM
    91.20300,  2.29044, 1.12044, 0.88119, // AMAMMT
    90.18550,  2.56257, 1.12931, 0.82924, // AMAMTA
    87.93340,  2.56257, 1.12747, 0.82720, // AMAMTC
    89.42720,  2.56257, 1.12618, 0.82579, // AMAMTG
    87.93340,  2.56257, 1.12747, 0.82720, // AMAMTM
    89.02180,  2.56257, 1.09133, 0.78776, // AMAMTT
    89.15220,  2.95200, 1.62646, 1.31493, // AMATAA
    85.54060,  2.95200, 1.69418, 1.39791, // AMATAC
    88.49880,  2.95200, 1.54416, 1.21641, // AMATAG
    85.54060,  2.95200, 1.69418, 1.39791, // AMATAM
    87.29000,  2.95200, 1.68679, 1.38878, // AMATAT
    82.32280,  3.22932, 2.14178, 2.16515, // AMATCA
    80.65780,  3.22932, 2.23229, 2.30384, // AMATCC
    82.23760,  3.22932, 2.00829, 1.96592, // AMATCG
    80.65780,  3.22932, 2.23229, 2.30384, // AMATCM
    81.80180,  3.22932, 2.09412, 2.09328, // AMATCT
    86.59770,  3.18506, 1.56195, 1.23545, // AMATGA
    82.80010,  3.18506, 1.79534, 1.52245, // AMATGC
    85.50660,  3.18506, 1.74257, 1.45582, // AMATGG
    82.80010,  3.18506, 1.79534, 1.52245, // AMATGM
    83.96140,  3.18506, 1.83102, 1.56805, // AMATGT
    82.32280,  3.22932, 2.14178, 2.16515, // AMATMA
    80.65780,  3.22932, 2.23229, 2.30384, // AMATMC
    82.23760,  3.22932, 2.00829, 1.96592, // AMATMG
    80.65780,  3.22932, 2.23229, 2.30384, // AMATMM
    81.80180,  3.22932, 2.09412, 2.09328, // AMATMT
    76.51900,  3.43089, 2.12527, 1.99846, // AMATTA
    73.42810,  3.43089, 2.23765, 2.15905, // AMATTC
    75.79690,  3.43089, 2.19356, 2.09555, // AMATTG
    73.42810,  3.43089, 2.23765, 2.15905, // AMATTM
    74.69010,  3.43089, 2.04004, 1.87946, // AMATTT
    103.97700, 2.63085, 1.23603, 1.02003, // AMCAAA
    101.47400, 2.63085, 1.45536, 1.30325, // AMCAAC
    103.08300, 2.63085, 1.29590, 1.09503, // AMCAAG
    101.47400, 2.63085, 1.45536, 1.30325, // AMCAAM
    102.38000, 2.63085, 1.49323, 1.35444, // AMCAAT
    96.35810,  2.46322, 1.05360, 0.74833, // AMCACA
    95.03570,  2.46322, 1.10929, 0.80845, // AMCACC
    96.08640,  2.46322, 1.09260, 0.79027, // AMCACG
    95.03570,  2.46322, 1.10929, 0.80845, // AMCACM
    95.95960,  2.46322, 1.15982, 0.86431, // AMCACT
    101.89800, 2.84203, 1.39859, 1.15660, // AMCAGA
    99.59450,  2.84203, 1.44253, 1.21154, // AMCAGC
    100.71000, 2.84203, 1.39727, 1.15497, // AMCAGG
    99.59450,  2.84203, 1.44253, 1.21154, // AMCAGM
    100.27000, 2.84203, 1.59888, 1.41375, // AMCAGT
    96.35810,  2.46322, 1.05360, 0.74833, // AMCAMA
    95.03570,  2.46322, 1.10929, 0.80845, // AMCAMC
    96.08640,  2.46322, 1.09260, 0.79027, // AMCAMG
    95.03570,  2.46322, 1.10929, 0.80845, // AMCAMM
    95.95960,  2.46322, 1.15982, 0.86431, // AMCAMT
    95.04160,  2.79927, 1.30899, 0.96780, // AMCATA
    93.32730,  2.79927, 1.50756, 1.19616, // AMCATC
    94.58490,  2.79927, 1.39952, 1.06991, // AMCATG
    93.32730,  2.79927, 1.50756, 1.19616, // AMCATM
    94.37830,  2.79927, 1.64925, 1.36870, // AMCATT
    103.40400, 2.44981, 0.87226, 0.61194, // AMCCAA
    100.50300, 2.44981, 0.96606, 0.71325, // AMCCAC
    102.49000, 2.44981, 0.89341, 0.63433, // AMCCAG
    100.50300, 2.44981, 0.96606, 0.71325, // AMCCAM
    101.52700, 2.44981, 1.06095, 0.82088, // AMCCAT
    99.35500,  2.40182, 0.89591, 0.64955, // AMCCCA
    97.85840,  2.40182, 0.89646, 0.65015, // AMCCCC
    99.21720,  2.40182, 0.90089, 0.65497, // AMCCCG
    97.85840,  2.40182, 0.89646, 0.65015, // AMCCCM
    99.09360,  2.40182, 0.87739, 0.62951, // AMCCCT
    101.93700, 2.43192, 0.92684, 0.68212, // AMCCGA
    99.34460,  2.43192, 0.94800, 0.70561, // AMCCGC
    100.73300, 2.43192, 0.90440, 0.65750, // AMCCGG
    99.34460,  2.43192, 0.94800, 0.70561, // AMCCGM
    100.16700, 2.43192, 1.03949, 0.81019, // AMCCGT
    99.35500,  2.40182, 0.89591, 0.64955, // AMCCMA
    97.85840,  2.40182, 0.89646, 0.65015, // AMCCMC
    99.21720,  2.40182, 0.90089, 0.65497, // AMCCMG
    97.85840,  2.40182, 0.89646, 0.65015, // AMCCMM
    99.09360,  2.40182, 0.87739, 0.62951, // AMCCMT
    98.28370,  2.44710, 0.97684, 0.73578, // AMCCTA
    96.48630,  2.44710, 1.13600, 0.92274, // AMCCTC
    97.66750,  2.44710, 0.99337, 0.75453, // AMCCTG
    96.48630,  2.44710, 1.13600, 0.92274, // AMCCTM
    96.88320,  2.44710, 1.19237, 0.99226, // AMCCTT
    99.07040,  2.82025, 1.36048, 1.11248, // AMCGAA
    94.17780,  2.82025, 1.59801, 1.41621, // AMCGAC
    97.84920,  2.82025, 1.41891, 1.18491, // AMCGAG
    94.17780,  2.82025, 1.59801, 1.41621, // AMCGAM
    95.99360,  2.82025, 1.67392, 1.51830, // AMCGAT
    94.48170,  2.58093, 1.12144, 0.78696, // AMCGCA
    92.97800,  2.58093, 1.21249, 0.88471, // AMCGCC
    94.16970,  2.58093, 1.18195, 0.85150, // AMCGCG
    92.97800,  2.58093, 1.21249, 0.88471, // AMCGCM
    94.08150,  2.58093, 1.21946, 0.89235, // AMCGCT
    98.80270,  2.74812, 1.35912, 1.15166, // AMCGGA
    95.47870,  2.74812, 1.45115, 1.27059, // AMCGGC
    97.57450,  2.74812, 1.36074, 1.15372, // AMCGGG
    95.47870,  2.74812, 1.45115, 1.27059, // AMCGGM
    96.85990,  2.74812, 1.51550, 1.35604, // AMCGGT
    94.48170,  2.58093, 1.12144, 0.78696, // AMCGMA
    92.97800,  2.58093, 1.21249, 0.88471, // AMCGMC
    94.16970,  2.58093, 1.18195, 0.85150, // AMCGMG
    92.97800,  2.58093, 1.21249, 0.88471, // AMCGMM
    94.08150,  2.58093, 1.21946, 0.89235, // AMCGMT
    92.62400,  3.13467, 1.44073, 1.19456, // AMCGTA
    90.17100,  3.13467, 1.67103, 1.49214, // AMCGTC
    91.68520,  3.13467, 1.59610, 1.39291, // AMCGTG
    90.17100,  3.13467, 1.67103, 1.49214, // AMCGTM
    91.56360,  3.13467, 1.79201, 1.65708, // AMCGTT
    103.40400, 2.44981, 0.87226, 0.61194, // AMCMAA
    100.50300, 2.44981, 0.96606, 0.71325, // AMCMAC
    102.49000, 2.44981, 0.89341, 0.63433, // AMCMAG
    100.50300, 2.44981, 0.96606, 0.71325, // AMCMAM
    101.52700, 2.44981, 1.06095, 0.82088, // AMCMAT
    99.35500,  2.40182, 0.89591, 0.64955, // AMCMCA
    97.85840,  2.40182, 0.89646, 0.65015, // AMCMCC
    99.21720,  2.40182, 0.90089, 0.65497, // AMCMCG
    97.85840,  2.40182, 0.89646, 0.65015, // AMCMCM
    99.09360,  2.40182, 0.87739, 0.62951, // AMCMCT
    101.93700, 2.43192, 0.92684, 0.68212, // AMCMGA
    99.34460,  2.43192, 0.94800, 0.70561, // AMCMGC
    100.73300, 2.43192, 0.90440, 0.65750, // AMCMGG
    99.34460,  2.43192, 0.94800, 0.70561, // AMCMGM
    100.16700, 2.43192, 1.03949, 0.81019, // AMCMGT
    99.35500,  2.40182, 0.89591, 0.64955, // AMCMMA
    97.85840,  2.40182, 0.89646, 0.65015, // AMCMMC
    99.21720,  2.40182, 0.90089, 0.65497, // AMCMMG
    97.85840,  2.40182, 0.89646, 0.65015, // AMCMMM
    99.09360,  2.40182, 0.87739, 0.62951, // AMCMMT
    98.28370,  2.44710, 0.97684, 0.73578, // AMCMTA
    96.48630,  2.44710, 1.13600, 0.92274, // AMCMTC
    97.66750,  2.44710, 0.99337, 0.75453, // AMCMTG
    96.48630,  2.44710, 1.13600, 0.92274, // AMCMTM
    96.88320,  2.44710, 1.19237, 0.99226, // AMCMTT
    98.01510,  2.77408, 1.30411, 1.02284, // AMCTAA
    95.25880,  2.77408, 1.37864, 1.11177, // AMCTAC
    97.16900,  2.77408, 1.05253, 0.74164, // AMCTAG
    95.25880,  2.77408, 1.37864, 1.11177, // AMCTAM
    96.13040,  2.77408, 1.36413, 1.09426, // AMCTAT
    91.24900,  3.16210, 1.71292, 1.55628, // AMCTCA
    90.14290,  3.16210, 1.86109, 1.76252, // AMCTCC
    91.39750,  3.16210, 1.62527, 1.43836, // AMCTCG
    90.14290,  3.16210, 1.86109, 1.76252, // AMCTCM
    91.26810,  3.16210, 1.78964, 1.66200, // AMCTCT
    95.41650,  2.63830, 1.26908, 0.98862, // AMCTGA
    92.07510,  2.63830, 1.41720, 1.16665, // AMCTGC
    94.01390,  2.63830, 1.29676, 1.02113, // AMCTGG
    92.07510,  2.63830, 1.41720, 1.16665, // AMCTGM
    93.17520,  2.63830, 1.44582, 1.20217, // AMCTGT
    91.24900,  3.16210, 1.71292, 1.55628, // AMCTMA
    90.14290,  3.16210, 1.86109, 1.76252, // AMCTMC
    91.39750,  3.16210, 1.62527, 1.43836, // AMCTMG
    90.14290,  3.16210, 1.86109, 1.76252, // AMCTMM
    91.26810,  3.16210, 1.78964, 1.66200, // AMCTMT
    86.81060,  2.91311, 1.71890, 1.48956, // AMCTTA
    83.39010,  2.91311, 1.70994, 1.47793, // AMCTTC
    85.81540,  2.91311, 1.56463, 1.29360, // AMCTTG
    83.39010,  2.91311, 1.70994, 1.47793, // AMCTTM
    84.69540,  2.91311, 1.58306, 1.31652, // AMCTTT
    84.15780,  2.13730, 1.56701, 1.16930, // AMGAAA
    80.20460,  2.16843, 1.88115, 1.53799, // AMGAAC
    82.34610,  2.08670, 1.70811, 1.33074, // AMGAAG
    80.31970,  1.90287, 1.88115, 1.53799, // AMGAAM
    81.15770,  2.35480, 1.92702, 1.59459, // AMGAAT
    74.35130,  1.86803, 1.81079, 1.55061, // AMGACA
    73.39670,  1.93199, 1.65631, 1.35647, // AMGACC
    75.24330,  3.00332, 1.71221, 1.42572, // AMGACG
    73.70470,  1.90926, 1.65631, 1.35647, // AMGACM
    74.08250,  1.76233, 1.66362, 1.36547, // AMGACT
    82.75770,  1.95917, 1.51583, 1.11615, // AMGAGA
    80.62020,  2.39190, 1.78219, 1.42291, // AMGAGC
    81.63410,  1.92269, 1.51980, 1.12053, // AMGAGG
    80.54740,  1.81383, 1.78219, 1.42291, // AMGAGM
    80.80280,  2.62371, 1.83001, 1.48055, // AMGAGT
    75.36940,  3.00332, 1.81079, 1.55061, // AMGAMA
    74.19600,  3.00332, 1.65631, 1.35647, // AMGAMC
    78.26150,  2.08395, 1.71221, 1.42572, // AMGAMG
    74.19600,  3.00332, 1.65631, 1.35647, // AMGAMM
    75.11470,  3.00332, 1.66362, 1.36547, // AMGAMT
    72.92740,  1.66069, 1.56104, 1.41982, // AMGATA
    72.60430,  2.30490, 1.80103, 1.75951, // AMGATC
    72.47720,  1.77312, 1.56900, 1.43069, // AMGATG
    71.65490,  1.42806, 1.80103, 1.75951, // AMGATM
    72.38870,  2.06484, 1.93844, 1.96467, // AMGATT
    87.12370,  1.25801, 1.25549, 0.98457, // AMGCAA
    84.90570,  1.26572, 1.18231, 0.89975, // AMGCAC
    86.94200,  1.64247, 1.23807, 0.96416, // AMGCAG
    85.51720,  1.26712, 1.18231, 0.89975, // AMGCAM
    85.86510,  1.28264, 1.16388, 0.87880, // AMGCAT
    82.97910,  1.04439, 1.41921, 1.19162, // AMGCCA
    82.45760,  1.23567, 1.44306, 1.22178, // AMGCCC
    86.59410,  2.52741, 1.41830, 1.19047, // AMGCCG
    82.80030,  1.26929, 1.44306, 1.22178, // AMGCCM
    84.09390,  1.75765, 1.36153, 1.11971, // AMGCCT
    88.86570,  2.58181, 1.28360, 0.98233, // AMGCGA
    85.75870,  2.58181, 1.35971, 1.07099, // AMGCGC
    87.67580,  2.58181, 1.34631, 1.05519, // AMGCGG
    85.75870,  2.58181, 1.35971, 1.07099, // AMGCGM
    86.47440,  2.58181, 1.34700, 1.05600, // AMGCGT
    86.80540,  2.52741, 1.41921, 1.19162, // AMGCMA
    85.42380,  2.52741, 1.44306, 1.22178, // AMGCMC
    83.67760,  1.06927, 1.41830, 1.19047, // AMGCMG
    85.42380,  2.52741, 1.44306, 1.22178, // AMGCMM
    86.47020,  2.52741, 1.36153, 1.11971, // AMGCMT
    81.94870,  1.34159, 1.41851, 1.12330, // AMGCTA
    79.92320,  1.37284, 1.39000, 1.08960, // AMGCTC
    81.63210,  1.77340, 1.39386, 1.09415, // AMGCTG
    80.70450,  1.32085, 1.39000, 1.08960, // AMGCTM
    80.36440,  1.26730, 1.18494, 0.85761, // AMGCTT
    82.26780,  2.49942, 1.76816, 1.37549, // AMGGAA
    76.37140,  2.27005, 2.03723, 1.70112, // AMGGAC
    80.44940,  2.43000, 1.82726, 1.44503, // AMGGAG
    77.87070,  2.19997, 2.03723, 1.70112, // AMGGAM
    78.13200,  2.37148, 2.01232, 1.67002, // AMGGAT
    78.16100,  1.91568, 1.49247, 1.20792, // AMGGCA
    76.56290,  1.75787, 1.50469, 1.22280, // AMGGCC
    79.78450,  3.03186, 1.48313, 1.19660, // AMGGCG
    77.07830,  1.86342, 1.50469, 1.22280, // AMGGCM
    77.96730,  2.13520, 1.50224, 1.21981, // AMGGCT
    83.69480,  2.25239, 1.63675, 1.21654, // AMGGGA
    81.16370,  2.47976, 1.81525, 1.42087, // AMGGGC
    82.58260,  2.23679, 1.59567, 1.17102, // AMGGGG
    81.28620,  2.32764, 1.81525, 1.42087, // AMGGGM
    82.02250,  2.53183, 1.85320, 1.46566, // AMGGGT
    79.99910,  3.03186, 1.49247, 1.20792, // AMGGMA
    78.44170,  3.03186, 1.50469, 1.22280, // AMGGMC
    79.99090,  1.64679, 1.48313, 1.19660, // AMGGMG
    78.44170,  3.03186, 1.50469, 1.22280, // AMGGMM
    79.70290,  3.03186, 1.50224, 1.21981, // AMGGMT
    76.61960,  1.99350, 1.84312, 1.56160, // AMGGTA
    73.92960,  2.34096, 2.00903, 1.77714, // AMGGTC
    75.62370,  2.24981, 1.92296, 1.66416, // AMGGTG
    74.69120,  2.00244, 2.00903, 1.77714, // AMGGTM
    74.93890,  2.73403, 1.99035, 1.75241, // AMGGTT
    90.72400,  2.44291, 1.25549, 0.98457, // AMGMAA
    87.92700,  2.44291, 1.18231, 0.89975, // AMGMAC
    89.69160,  2.44291, 1.23807, 0.96416, // AMGMAG
    87.92700,  2.44291, 1.18231, 0.89975, // AMGMAM
    88.90820,  2.44291, 1.16388, 0.87880, // AMGMAT
    86.80540,  2.52741, 1.41921, 1.19162, // AMGMCA
    85.42380,  2.52741, 1.44306, 1.22178, // AMGMCC
    86.59410,  2.52741, 1.41830, 1.19047, // AMGMCG
    85.42380,  2.52741, 1.44306, 1.22178, // AMGMCM
    86.47020,  2.52741, 1.36153, 1.11971, // AMGMCT
    85.83480,  1.99589, 1.28360, 0.98233, // AMGMGA
    82.49870,  1.69176, 1.35971, 1.07099, // AMGMGC
    84.90660,  2.27394, 1.34631, 1.05519, // AMGMGG
    83.37320,  1.67386, 1.35971, 1.07099, // AMGMGM
    82.90510,  1.76316, 1.34700, 1.05600, // AMGMGT
    86.80540,  2.52741, 1.41921, 1.19162, // AMGMMA
    85.42380,  2.52741, 1.44306, 1.22178, // AMGMMC
    86.59410,  2.52741, 1.41830, 1.19047, // AMGMMG
    85.42380,  2.52741, 1.44306, 1.22178, // AMGMMM
    86.47020,  2.52741, 1.36153, 1.11971, // AMGMMT
    85.13680,  2.56040, 1.41851, 1.12330, // AMGMTA
    82.69010,  2.56040, 1.39000, 1.08960, // AMGMTC
    84.23900,  2.56040, 1.39386, 1.09415, // AMGMTG
    82.69010,  2.56040, 1.39000, 1.08960, // AMGMTM
    83.58710,  2.56040, 1.18494, 0.85761, // AMGMTT
    79.39490,  1.72463, 1.80972, 1.57415, // AMGTAA
    75.63040,  1.59406, 1.93097, 1.73496, // AMGTAC
    78.48570,  1.76534, 1.89084, 1.68117, // AMGTAG
    76.87960,  1.82983, 1.93097, 1.73496, // AMGTAM
    77.13840,  1.68605, 2.00056, 1.82960, // AMGTAT
    73.27330,  1.85214, 2.48078, 2.86185, // AMGTCA
    72.00580,  1.80222, 2.54924, 2.98112, // AMGTCC
    75.21510,  3.49565, 2.45874, 2.82380, // AMGTCG
    72.40770,  1.94764, 2.54924, 2.98112, // AMGTCM
    72.97370,  1.84268, 2.47828, 2.85752, // AMGTCT
    76.77490,  1.92691, 1.90469, 1.77665, // AMGTGA
    73.11180,  1.94814, 2.14974, 2.13031, // AMGTGC
    75.47950,  1.94435, 2.11114, 2.07319, // AMGTGG
    75.09470,  2.52342, 2.14974, 2.13031, // AMGTGM
    74.14520,  1.87611, 2.24488, 2.27328, // AMGTGT
    75.28890,  3.49565, 2.48078, 2.86185, // AMGTMA
    73.78890,  3.49565, 2.54924, 2.98112, // AMGTMC
    75.97890,  2.19888, 2.45874, 2.82380, // AMGTMG
    73.78890,  3.49565, 2.54924, 2.98112, // AMGTMM
    75.03440,  3.49565, 2.47828, 2.85752, // AMGTMT
    68.18140,  1.99144, 2.39457, 2.49536, // AMGTTA
    64.86680,  1.94703, 2.51962, 2.69335, // AMGTTC
    67.65150,  2.13364, 2.44079, 2.56795, // AMGTTG
    67.18220,  2.60932, 2.51962, 2.69335, // AMGTTM
    66.90070,  2.31829, 2.40054, 2.50469, // AMGTTT
    103.97700, 2.63085, 1.23603, 1.02003, // AMMAAA
    101.47400, 2.63085, 1.45536, 1.30325, // AMMAAC
    103.08300, 2.63085, 1.29590, 1.09503, // AMMAAG
    101.47400, 2.63085, 1.45536, 1.30325, // AMMAAM
    102.38000, 2.63085, 1.49323, 1.35444, // AMMAAT
    96.35810,  2.46322, 1.05360, 0.74833, // AMMACA
    95.03570,  2.46322, 1.10929, 0.80845, // AMMACC
    96.08640,  2.46322, 1.09260, 0.79027, // AMMACG
    95.03570,  2.46322, 1.10929, 0.80845, // AMMACM
    95.95960,  2.46322, 1.15982, 0.86431, // AMMACT
    101.89800, 2.84203, 1.39859, 1.15660, // AMMAGA
    99.59450,  2.84203, 1.44253, 1.21154, // AMMAGC
    100.71000, 2.84203, 1.39727, 1.15497, // AMMAGG
    99.59450,  2.84203, 1.44253, 1.21154, // AMMAGM
    100.27000, 2.84203, 1.59888, 1.41375, // AMMAGT
    96.35810,  2.46322, 1.05360, 0.74833, // AMMAMA
    95.03570,  2.46322, 1.10929, 0.80845, // AMMAMC
    96.08640,  2.46322, 1.09260, 0.79027, // AMMAMG
    95.03570,  2.46322, 1.10929, 0.80845, // AMMAMM
    95.95960,  2.46322, 1.15982, 0.86431, // AMMAMT
    95.04160,  2.79927, 1.30899, 0.96780, // AMMATA
    93.32730,  2.79927, 1.50756, 1.19616, // AMMATC
    94.58490,  2.79927, 1.39952, 1.06991, // AMMATG
    93.32730,  2.79927, 1.50756, 1.19616, // AMMATM
    94.37830,  2.79927, 1.64925, 1.36870, // AMMATT
    103.40400, 2.44981, 0.87226, 0.61194, // AMMCAA
    100.50300, 2.44981, 0.96606, 0.71325, // AMMCAC
    102.49000, 2.44981, 0.89341, 0.63433, // AMMCAG
    100.50300, 2.44981, 0.96606, 0.71325, // AMMCAM
    101.52700, 2.44981, 1.06095, 0.82088, // AMMCAT
    99.35500,  2.40182, 0.89591, 0.64955, // AMMCCA
    97.85840,  2.40182, 0.89646, 0.65015, // AMMCCC
    99.21720,  2.40182, 0.90089, 0.65497, // AMMCCG
    97.85840,  2.40182, 0.89646, 0.65015, // AMMCCM
    99.09360,  2.40182, 0.87739, 0.62951, // AMMCCT
    101.93700, 2.43192, 0.92684, 0.68212, // AMMCGA
    99.34460,  2.43192, 0.94800, 0.70561, // AMMCGC
    100.73300, 2.43192, 0.90440, 0.65750, // AMMCGG
    99.34460,  2.43192, 0.94800, 0.70561, // AMMCGM
    100.16700, 2.43192, 1.03949, 0.81019, // AMMCGT
    99.35500,  2.40182, 0.89591, 0.64955, // AMMCMA
    97.85840,  2.40182, 0.89646, 0.65015, // AMMCMC
    99.21720,  2.40182, 0.90089, 0.65497, // AMMCMG
    97.85840,  2.40182, 0.89646, 0.65015, // AMMCMM
    99.09360,  2.40182, 0.87739, 0.62951, // AMMCMT
    98.28370,  2.44710, 0.97684, 0.73578, // AMMCTA
    96.48630,  2.44710, 1.13600, 0.92274, // AMMCTC
    97.66750,  2.44710, 0.99337, 0.75453, // AMMCTG
    96.48630,  2.44710, 1.13600, 0.92274, // AMMCTM
    96.88320,  2.44710, 1.19237, 0.99226, // AMMCTT
    99.07040,  2.82025, 1.36048, 1.11248, // AMMGAA
    94.17780,  2.82025, 1.59801, 1.41621, // AMMGAC
    97.84920,  2.82025, 1.41891, 1.18491, // AMMGAG
    94.17780,  2.82025, 1.59801, 1.41621, // AMMGAM
    95.99360,  2.82025, 1.67392, 1.51830, // AMMGAT
    94.48170,  2.58093, 1.12144, 0.78696, // AMMGCA
    92.97800,  2.58093, 1.21249, 0.88471, // AMMGCC
    94.16970,  2.58093, 1.18195, 0.85150, // AMMGCG
    92.97800,  2.58093, 1.21249, 0.88471, // AMMGCM
    94.08150,  2.58093, 1.21946, 0.89235, // AMMGCT
    98.80270,  2.74812, 1.35912, 1.15166, // AMMGGA
    95.47870,  2.74812, 1.45115, 1.27059, // AMMGGC
    97.57450,  2.74812, 1.36074, 1.15372, // AMMGGG
    95.47870,  2.74812, 1.45115, 1.27059, // AMMGGM
    96.85990,  2.74812, 1.51550, 1.35604, // AMMGGT
    94.48170,  2.58093, 1.12144, 0.78696, // AMMGMA
    92.97800,  2.58093, 1.21249, 0.88471, // AMMGMC
    94.16970,  2.58093, 1.18195, 0.85150, // AMMGMG
    92.97800,  2.58093, 1.21249, 0.88471, // AMMGMM
    94.08150,  2.58093, 1.21946, 0.89235, // AMMGMT
    92.62400,  3.13467, 1.44073, 1.19456, // AMMGTA
    90.17100,  3.13467, 1.67103, 1.49214, // AMMGTC
    91.68520,  3.13467, 1.59610, 1.39291, // AMMGTG
    90.17100,  3.13467, 1.67103, 1.49214, // AMMGTM
    91.56360,  3.13467, 1.79201, 1.65708, // AMMGTT
    103.40400, 2.44981, 0.87226, 0.61194, // AMMMAA
    100.50300, 2.44981, 0.96606, 0.71325, // AMMMAC
    102.49000, 2.44981, 0.89341, 0.63433, // AMMMAG
    100.50300, 2.44981, 0.96606, 0.71325, // AMMMAM
    101.52700, 2.44981, 1.06095, 0.82088, // AMMMAT
    99.35500,  2.40182, 0.89591, 0.64955, // AMMMCA
    97.85840,  2.40182, 0.89646, 0.65015, // AMMMCC
    99.21720,  2.40182, 0.90089, 0.65497, // AMMMCG
    97.85840,  2.40182, 0.89646, 0.65015, // AMMMCM
    99.09360,  2.40182, 0.87739, 0.62951, // AMMMCT
    101.93700, 2.43192, 0.92684, 0.68212, // AMMMGA
    99.34460,  2.43192, 0.94800, 0.70561, // AMMMGC
    100.73300, 2.43192, 0.90440, 0.65750, // AMMMGG
    99.34460,  2.43192, 0.94800, 0.70561, // AMMMGM
    100.16700, 2.43192, 1.03949, 0.81019, // AMMMGT
    99.35500,  2.40182, 0.89591, 0.64955, // AMMMMA
    97.85840,  2.40182, 0.89646, 0.65015, // AMMMMC
    99.21720,  2.40182, 0.90089, 0.65497, // AMMMMG
    97.85840,  2.40182, 0.89646, 0.65015, // AMMMMM
    99.09360,  2.40182, 0.87739, 0.62951, // AMMMMT
    98.28370,  2.44710, 0.97684, 0.73578, // AMMMTA
    96.48630,  2.44710, 1.13600, 0.92274, // AMMMTC
    97.66750,  2.44710, 0.99337, 0.75453, // AMMMTG
    96.48630,  2.44710, 1.13600, 0.92274, // AMMMTM
    96.88320,  2.44710, 1.19237, 0.99226, // AMMMTT
    98.01510,  2.77408, 1.30411, 1.02284, // AMMTAA
    95.25880,  2.77408, 1.37864, 1.11177, // AMMTAC
    97.16900,  2.77408, 1.05253, 0.74164, // AMMTAG
    95.25880,  2.77408, 1.37864, 1.11177, // AMMTAM
    96.13040,  2.77408, 1.36413, 1.09426, // AMMTAT
    91.24900,  3.16210, 1.71292, 1.55628, // AMMTCA
    90.14290,  3.16210, 1.86109, 1.76252, // AMMTCC
    91.39750,  3.16210, 1.62527, 1.43836, // AMMTCG
    90.14290,  3.16210, 1.86109, 1.76252, // AMMTCM
    91.26810,  3.16210, 1.78964, 1.66200, // AMMTCT
    95.41650,  2.63830, 1.26908, 0.98862, // AMMTGA
    92.07510,  2.63830, 1.41720, 1.16665, // AMMTGC
    94.01390,  2.63830, 1.29676, 1.02113, // AMMTGG
    92.07510,  2.63830, 1.41720, 1.16665, // AMMTGM
    93.17520,  2.63830, 1.44582, 1.20217, // AMMTGT
    91.24900,  3.16210, 1.71292, 1.55628, // AMMTMA
    90.14290,  3.16210, 1.86109, 1.76252, // AMMTMC
    91.39750,  3.16210, 1.62527, 1.43836, // AMMTMG
    90.14290,  3.16210, 1.86109, 1.76252, // AMMTMM
    91.26810,  3.16210, 1.78964, 1.66200, // AMMTMT
    86.81060,  2.91311, 1.71890, 1.48956, // AMMTTA
    83.39010,  2.91311, 1.70994, 1.47793, // AMMTTC
    85.81540,  2.91311, 1.56463, 1.29360, // AMMTTG
    83.39010,  2.91311, 1.70994, 1.47793, // AMMTTM
    84.69540,  2.91311, 1.58306, 1.31652, // AMMTTT
    109.60500, 3.64098, 1.78499, 1.34274, // AMTAAA
    107.06700, 3.64098, 1.77711, 1.33385, // AMTAAC
    108.71700, 3.64098, 1.82866, 1.39231, // AMTAAG
    107.06700, 3.64098, 1.77711, 1.33385, // AMTAAM
    108.65000, 3.64098, 1.98981, 1.58036, // AMTAAT
    104.91300, 3.58253, 1.53278, 1.06551, // AMTACA
    104.37800, 3.58253, 1.69406, 1.23803, // AMTACC
    104.60200, 3.58253, 1.50915, 1.04096, // AMTACG
    104.37800, 3.58253, 1.69406, 1.23803, // AMTACM
    105.78100, 3.58253, 1.81195, 1.36948, // AMTACT
    107.84100, 3.51606, 1.83695, 1.55379, // AMTAGA
    105.68900, 3.51606, 1.61234, 1.27770, // AMTAGC
    106.36400, 3.51606, 1.53484, 1.18670, // AMTAGG
    105.68900, 3.51606, 1.61234, 1.27770, // AMTAGM
    107.10900, 3.51606, 1.60849, 1.27313, // AMTAGT
    104.91300, 3.58253, 1.53278, 1.06551, // AMTAMA
    104.37800, 3.58253, 1.69406, 1.23803, // AMTAMC
    104.60200, 3.58253, 1.50915, 1.04096, // AMTAMG
    104.37800, 3.58253, 1.69406, 1.23803, // AMTAMM
    105.78100, 3.58253, 1.81195, 1.36948, // AMTAMT
    104.96100, 4.03892, 1.91641, 1.40773, // AMTATA
    105.35600, 4.03892, 2.02830, 1.53279, // AMTATC
    104.56500, 4.03892, 1.87534, 1.36271, // AMTATG
    105.35600, 4.03892, 2.02830, 1.53279, // AMTATM
    106.93500, 4.03892, 2.25275, 1.79414, // AMTATT
    115.09200, 3.42793, 1.81878, 1.59109, // AMTCAA
    112.57800, 3.42793, 1.69623, 1.43303, // AMTCAC
    114.35000, 3.42793, 1.82823, 1.60352, // AMTCAG
    112.57800, 3.42793, 1.69623, 1.43303, // AMTCAM
    113.79400, 3.42793, 1.94818, 1.76388, // AMTCAT
    113.71500, 3.47041, 1.82491, 1.51937, // AMTCCA
    111.72700, 3.47041, 1.74480, 1.42044, // AMTCCC
    113.64900, 3.47041, 1.84678, 1.54677, // AMTCCG
    111.72700, 3.47041, 1.74480, 1.42044, // AMTCCM
    112.87600, 3.47041, 1.95376, 1.68309, // AMTCCT
    113.92000, 3.49891, 1.79737, 1.48675, // AMTCGA
    111.63700, 3.49891, 1.85362, 1.55710, // AMTCGC
    114.07000, 3.49891, 1.98026, 1.71936, // AMTCGG
    111.63700, 3.49891, 1.85362, 1.55710, // AMTCGM
    113.36900, 3.49891, 2.00748, 1.75494, // AMTCGT
    113.71500, 3.47041, 1.82491, 1.51937, // AMTCMA
    111.72700, 3.47041, 1.74480, 1.42044, // AMTCMC
    113.64900, 3.47041, 1.84678, 1.54677, // AMTCMG
    111.72700, 3.47041, 1.74480, 1.42044, // AMTCMM
    112.87600, 3.47041, 1.95376, 1.68309, // AMTCMT
    113.72600, 3.72204, 1.82568, 1.50849, // AMTCTA
    111.73300, 3.72204, 1.99910, 1.72846, // AMTCTC
    113.20400, 3.72204, 1.94200, 1.65494, // AMTCTG
    111.73300, 3.72204, 1.99910, 1.72846, // AMTCTM
    112.99600, 3.72204, 2.31939, 2.16006, // AMTCTT
    105.11100, 3.50593, 1.75335, 1.35976, // AMTGAA
    101.17500, 3.50593, 1.64828, 1.23939, // AMTGAC
    104.20200, 3.50593, 1.72515, 1.32709, // AMTGAG
    101.17500, 3.50593, 1.64828, 1.23939, // AMTGAM
    103.16400, 3.50593, 1.82084, 1.43903, // AMTGAT
    102.97600, 3.56584, 1.57276, 1.10065, // AMTGCA
    102.18700, 3.56584, 1.62044, 1.15109, // AMTGCC
    102.89100, 3.56584, 1.64559, 1.17798, // AMTGCG
    102.18700, 3.56584, 1.62044, 1.15109, // AMTGCM
    103.88800, 3.56584, 1.76473, 1.30821, // AMTGCT
    105.30300, 3.47346, 1.75997, 1.38612, // AMTGGA
    102.53300, 3.47346, 1.64528, 1.25286, // AMTGGC
    104.40800, 3.47346, 1.73224, 1.35349, // AMTGGG
    102.53300, 3.47346, 1.64528, 1.25286, // AMTGGM
    104.49900, 3.47346, 1.86940, 1.51738, // AMTGGT
    102.97600, 3.56584, 1.57276, 1.10065, // AMTGMA
    102.18700, 3.56584, 1.62044, 1.15109, // AMTGMC
    102.89100, 3.56584, 1.64559, 1.17798, // AMTGMG
    102.18700, 3.56584, 1.62044, 1.15109, // AMTGMM
    103.88800, 3.56584, 1.76473, 1.30821, // AMTGMT
    103.07900, 4.11250, 1.91169, 1.39461, // AMTGTA
    103.11900, 4.11250, 2.04214, 1.53976, // AMTGTC
    103.12700, 4.11250, 1.91520, 1.39845, // AMTGTG
    103.11900, 4.11250, 2.04214, 1.53976, // AMTGTM
    104.95900, 4.11250, 2.23889, 1.76757, // AMTGTT
    115.09200, 3.42793, 1.81878, 1.59109, // AMTMAA
    112.57800, 3.42793, 1.69623, 1.43303, // AMTMAC
    114.35000, 3.42793, 1.82823, 1.60352, // AMTMAG
    112.57800, 3.42793, 1.69623, 1.43303, // AMTMAM
    113.79400, 3.42793, 1.94818, 1.76388, // AMTMAT
    113.71500, 3.47041, 1.82491, 1.51937, // AMTMCA
    111.72700, 3.47041, 1.74480, 1.42044, // AMTMCC
    113.64900, 3.47041, 1.84678, 1.54677, // AMTMCG
    111.72700, 3.47041, 1.74480, 1.42044, // AMTMCM
    112.87600, 3.47041, 1.95376, 1.68309, // AMTMCT
    113.92000, 3.49891, 1.79737, 1.48675, // AMTMGA
    111.63700, 3.49891, 1.85362, 1.55710, // AMTMGC
    114.07000, 3.49891, 1.98026, 1.71936, // AMTMGG
    111.63700, 3.49891, 1.85362, 1.55710, // AMTMGM
    113.36900, 3.49891, 2.00748, 1.75494, // AMTMGT
    113.71500, 3.47041, 1.82491, 1.51937, // AMTMMA
    111.72700, 3.47041, 1.74480, 1.42044, // AMTMMC
    113.64900, 3.47041, 1.84678, 1.54677, // AMTMMG
    111.72700, 3.47041, 1.74480, 1.42044, // AMTMMM
    112.87600, 3.47041, 1.95376, 1.68309, // AMTMMT
    113.72600, 3.72204, 1.82568, 1.50849, // AMTMTA
    111.73300, 3.72204, 1.99910, 1.72846, // AMTMTC
    113.20400, 3.72204, 1.94200, 1.65494, // AMTMTG
    111.73300, 3.72204, 1.99910, 1.72846, // AMTMTM
    112.99600, 3.72204, 2.31939, 2.16006, // AMTMTT
    104.24200, 2.86007, 1.35553, 0.99397, // AMTTAA
    100.05900, 2.86007, 1.25649, 0.88704, // AMTTAC
    103.66600, 2.86007, 1.31032, 0.94465, // AMTTAG
    100.05900, 2.86007, 1.25649, 0.88704, // AMTTAM
    101.76100, 2.86007, 1.34247, 0.97963, // AMTTAT
    98.45000,  2.79285, 1.45211, 1.11851, // AMTTCA
    96.66330,  2.79285, 1.44106, 1.10576, // AMTTCC
    98.39870,  2.79285, 1.42524, 1.08761, // AMTTCG
    96.66330,  2.79285, 1.44106, 1.10576, // AMTTCM
    98.02970,  2.79285, 1.42580, 1.08825, // AMTTCT
    102.26200, 2.88803, 1.27431, 0.94949, // AMTTGA
    98.11490,  2.88803, 1.25882, 0.93224, // AMTTGC
    101.32600, 2.88803, 1.36817, 1.05630, // AMTTGG
    98.11490,  2.88803, 1.25882, 0.93224, // AMTTGM
    99.81330,  2.88803, 1.30757, 0.98691, // AMTTGT
    98.45000,  2.79285, 1.45211, 1.11851, // AMTTMA
    96.66330,  2.79285, 1.44106, 1.10576, // AMTTMC
    98.39870,  2.79285, 1.42524, 1.08761, // AMTTMG
    96.66330,  2.79285, 1.44106, 1.10576, // AMTTMM
    98.02970,  2.79285, 1.42580, 1.08825, // AMTTMT
    94.60810,  2.89728, 1.23750, 0.84797, // AMTTTA
    92.09170,  2.89728, 1.20209, 0.81183, // AMTTTC
    93.95160,  2.89728, 1.22881, 0.83906, // AMTTTG
    92.09170,  2.89728, 1.20209, 0.81183, // AMTTTM
    94.31760,  2.89728, 1.23664, 0.84708, // AMTTTT
    92.99920,  2.57817, 1.57687, 1.22726, // ATAAAA
    91.93660,  2.73706, 1.89717, 1.61958, // ATAAAC
    94.09070,  2.67750, 1.62758, 1.28693, // ATAAAG
    91.32650,  2.63663, 1.89717, 1.61958, // ATAAAM
    92.17970,  2.83225, 1.88984, 1.61020, // ATAAAT
    85.23130,  2.39130, 1.69423, 1.46061, // ATAACA
    83.84470,  2.33237, 1.61840, 1.36365, // ATAACC
    84.38440,  2.30963, 1.64764, 1.40078, // ATAACG
    84.22150,  2.26129, 1.61840, 1.36365, // ATAACM
    84.75690,  2.54027, 1.57393, 1.30784, // ATAACT
    92.91490,  2.35804, 1.68310, 1.33125, // ATAAGA
    90.91790,  2.83068, 1.83482, 1.51526, // ATAAGC
    91.82490,  2.48717, 1.67035, 1.31616, // ATAAGG
    90.49900,  2.29382, 1.83482, 1.51526, // ATAAGM
    91.37840,  2.95606, 1.90871, 1.60770, // ATAAGT
    84.68290,  3.30963, 1.69423, 1.46061, // ATAAMA
    83.48880,  3.30963, 1.61840, 1.36365, // ATAAMC
    89.63480,  2.54915, 1.64764, 1.40078, // ATAAMG
    83.48880,  3.30963, 1.61840, 1.36365, // ATAAMM
    84.23410,  3.30963, 1.57393, 1.30784, // ATAAMT
    84.97320,  2.78268, 1.78173, 1.57934, // ATAATA
    83.37300,  2.70853, 2.15061, 2.09438, // ATAATC
    84.09420,  2.35155, 1.93051, 1.78124, // ATAATG
    84.41630,  2.59980, 2.15061, 2.09438, // ATAATM
    83.98440,  3.10225, 2.17920, 2.13629, // ATAATT
    95.63480,  1.37602, 1.26931, 1.02022, // ATACAA
    93.66500,  1.31675, 1.17206, 0.90524, // ATACAC
    94.52270,  1.30201, 1.17395, 0.90743, // ATACAG
    94.35080,  1.49046, 1.17206, 0.90524, // ATACAM
    94.38990,  1.30429, 1.13887, 0.86706, // ATACAT
    92.45450,  1.44271, 1.40558, 1.12366, // ATACCA
    91.83310,  1.50117, 1.37829, 1.09109, // ATACCC
    92.20570,  1.41774, 1.38253, 1.09614, // ATACCG
    92.39020,  1.59696, 1.37829, 1.09109, // ATACCM
    92.51020,  1.43075, 1.30744, 1.00806, // ATACCT
    94.14070,  1.34132, 1.21208, 0.90475, // ATACGA
    92.09590,  1.34132, 1.21240, 0.90510, // ATACGC
    93.18780,  1.34132, 1.23697, 0.93276, // ATACGG
    92.09590,  2.34132, 1.21240, 0.90510, // ATACGM
    92.64420,  1.34132, 1.20091, 0.89227, // ATACGT
    92.38950,  2.41774, 1.40558, 1.12366, // ATACMA
    91.19000,  2.41774, 1.37829, 1.09109, // ATACMC
    92.78190,  1.45966, 1.38253, 1.09614, // ATACMG
    91.19000,  2.41774, 1.37829, 1.09109, // ATACMM
    91.98580,  2.41774, 1.30744, 1.00806, // ATACMT
    91.70460,  1.59222, 1.41339, 1.17198, // ATACTA
    89.83520,  1.52996, 1.26653, 0.99414, // ATACTC
    90.82040,  1.52737, 1.35891, 1.10486, // ATACTG
    90.73590,  1.61810, 1.26653, 0.99414, // ATACTM
    90.81820,  1.63251, 1.37557, 1.12525, // ATACTT
    92.17390,  2.43650, 1.60769, 1.32628, // ATAGAA
    87.44610,  2.63871, 1.83709, 1.62004, // ATAGAC
    90.92230,  2.52521, 1.64517, 1.37293, // ATAGAG
    88.51390,  2.71239, 1.83709, 1.62004, // ATAGAM
    89.01770,  2.61821, 2.05073, 1.91070, // ATAGAT
    87.30920,  2.55132, 1.56045, 1.37884, // ATAGCA
    86.19680,  2.28485, 1.78523, 1.68725, // ATAGCC
    86.45720,  2.28919, 1.60563, 1.43915, // ATAGCG
    86.83260,  2.37959, 1.78523, 1.68725, // ATAGCM
    86.49920,  2.37359, 1.71304, 1.58595, // ATAGCT
    91.82930,  2.31171, 1.55603, 1.25638, // ATAGGA
    89.05180,  2.70573, 1.79962, 1.56266, // ATAGGC
    90.73180,  2.43013, 1.60771, 1.31949, // ATAGGG
    89.31490,  2.58998, 1.79962, 1.56266, // ATAGGM
    90.09680,  2.64480, 1.86582, 1.64968, // ATAGGT
    86.51260,  3.28919, 1.56045, 1.37884, // ATAGMA
    85.46650,  3.28919, 1.78523, 1.68725, // ATAGMC
    89.46030,  2.37875, 1.60563, 1.43915, // ATAGMG
    85.46650,  3.28919, 1.78523, 1.68725, // ATAGMM
    85.95470,  3.28919, 1.71304, 1.58595, // ATAGMT
    85.34130,  2.62401, 1.96004, 1.87494, // ATAGTA
    83.00640,  2.99592, 2.03688, 1.98627, // ATAGTC
    84.34280,  2.78393, 1.94091, 1.84756, // ATAGTG
    83.78090,  2.84488, 2.03688, 1.98627, // ATAGTM
    84.22430,  3.04540, 2.28158, 2.35475, // ATAGTT
    95.13490,  2.32011, 1.26931, 1.02022, // ATAMAA
    93.12880,  2.32011, 1.17206, 0.90524, // ATAMAC
    94.32320,  2.32011, 1.17395, 0.90743, // ATAMAG
    93.12880,  2.32011, 1.17206, 0.90524, // ATAMAM
    94.09220,  2.32011, 1.13887, 0.86706, // ATAMAT
    92.38950,  2.41774, 1.40558, 1.12366, // ATAMCA
    91.19000,  2.41774, 1.37829, 1.09109, // ATAMCC
    92.20570,  2.41774, 1.38253, 1.09614, // ATAMCG
    91.19000,  2.41774, 1.37829, 1.09109, // ATAMCM
    91.98580,  2.41774, 1.30744, 1.00806, // ATAMCT
    96.70020,  2.19793, 1.21208, 0.90475, // ATAMGA
    95.19800,  2.70967, 1.21240, 0.90510, // ATAMGC
    95.70050,  2.29261, 1.23697, 0.93276, // ATAMGG
    95.31400,  2.31238, 1.21240, 0.90510, // ATAMGM
    95.94590,  2.15383, 1.20091, 0.89227, // ATAMGT
    92.38950,  2.41774, 1.40558, 1.12366, // ATAMMA
    91.19000,  2.41774, 1.37829, 1.09109, // ATAMMC
    92.20570,  2.41774, 1.38253, 1.09614, // ATAMMG
    91.19000,  2.41774, 1.37829, 1.09109, // ATAMMM
    91.98580,  2.41774, 1.30744, 1.00806, // ATAMMT
    91.12710,  2.52174, 1.41339, 1.17198, // ATAMTA
    89.44630,  2.52174, 1.26653, 0.99414, // ATAMTC
    90.62900,  2.52174, 1.35891, 1.10486, // ATAMTG
    89.44630,  2.52174, 1.26653, 0.99414, // ATAMTM
    90.24400,  2.52174, 1.37557, 1.12525, // ATAMTT
    90.78690,  1.90175, 1.80275, 1.66447, // ATATAA
    87.94980,  2.05548, 1.83923, 1.71525, // ATATAC
    89.98300,  1.92506, 1.71064, 1.53855, // ATATAG
    88.92590,  2.32678, 1.83923, 1.71525, // ATATAM
    89.22440,  2.09677, 1.86705, 1.75431, // ATATAT
    85.89850,  2.51351, 2.35953, 2.49103, // ATATCA
    85.14660,  2.62386, 2.38703, 2.53470, // ATATCC
    85.37970,  2.46291, 2.24608, 2.31355, // ATATCG
    85.72540,  2.86179, 2.38703, 2.53470, // ATATCM
    86.06300,  2.53106, 2.31993, 2.42857, // ATATCT
    88.81380,  2.12952, 1.78046, 1.56483, // ATATGA
    85.73230,  2.32681, 2.06531, 1.95500, // ATATGC
    87.84340,  2.28266, 1.90620, 1.73349, // ATATGG
    86.70710,  2.41043, 2.06531, 1.95500, // ATATGM
    86.69020,  2.28101, 2.04659, 1.92847, // ATATGT
    85.68590,  3.46291, 2.35953, 2.49103, // ATATMA
    84.45940,  3.46291, 2.38703, 2.53470, // ATATMC
    87.73330,  2.38669, 2.24608, 2.31355, // ATATMG
    84.45940,  3.46291, 2.38703, 2.53470, // ATATMM
    85.28440,  3.46291, 2.31993, 2.42857, // ATATMT
    81.26430,  2.86154, 2.52054, 2.58799, // ATATTA
    78.17330,  2.94898, 2.48029, 2.52626, // ATATTC
    80.51750,  2.89001, 2.41837, 2.43225, // ATATTG
    79.62000,  3.00923, 2.48029, 2.52626, // ATATTM
    79.82140,  2.97799, 2.33927, 2.31390, // ATATTT
    106.15900, 2.38985, 1.65247, 1.41304, // ATCAAA
    102.60300, 2.34462, 1.83989, 1.66013, // ATCAAC
    104.24300, 2.20069, 1.68580, 1.45600, // ATCAAG
    103.25900, 2.21074, 1.83989, 1.66013, // ATCAAM
    105.05400, 2.74697, 1.95166, 1.81367, // ATCAAT
    98.60580,  2.15306, 1.67386, 1.39585, // ATCACA
    97.69250,  2.49415, 1.67189, 1.39338, // ATCACC
    97.77410,  2.10126, 1.61007, 1.31682, // ATCACG
    97.81590,  2.46720, 1.67189, 1.39338, // ATCACM
    98.14550,  2.43343, 1.74883, 1.49067, // ATCACT
    104.21500, 2.45202, 1.84262, 1.64081, // ATCAGA
    102.62500, 2.84470, 1.94509, 1.77956, // ATCAGC
    101.67300, 2.67570, 1.87611, 1.68574, // ATCAGG
    101.83500, 2.55753, 1.94509, 1.77956, // ATCAGM
    102.61800, 2.74051, 2.05478, 1.93220, // ATCAGT
    98.08030,  3.10126, 1.67386, 1.39585, // ATCAMA
    96.85750,  3.10126, 1.67189, 1.39338, // ATCAMC
    101.75200, 2.32785, 1.61007, 1.31682, // ATCAMG
    96.85750,  3.10126, 1.67189, 1.39338, // ATCAMM
    97.54660,  3.10126, 1.74883, 1.49067, // ATCAMT
    97.08410,  2.49146, 1.82769, 1.59362, // ATCATA
    95.34700,  2.81781, 1.97392, 1.78865, // ATCATC
    96.58420,  2.57934, 1.85877, 1.63444, // ATCATG
    96.22330,  2.73994, 1.97392, 1.78865, // ATCATM
    96.27310,  2.82551, 2.16147, 2.04953, // ATCATT
    104.49300, 1.57706, 1.39898, 1.21060, // ATCCAA
    101.74000, 1.91250, 1.41598, 1.23273, // ATCCAC
    102.27700, 1.90441, 1.27224, 1.04987, // ATCCAG
    102.43400, 1.72254, 1.41598, 1.23273, // ATCCAM
    102.73100, 1.90430, 1.52237, 1.37424, // ATCCAT
    101.33700, 1.88867, 1.21153, 0.98116, // ATCCCA
    100.45100, 1.81125, 1.36765, 1.17679, // ATCCCC
    100.56300, 1.60868, 1.29170, 1.08014, // ATCCCG
    100.84600, 1.78160, 1.36765, 1.17679, // ATCCCM
    100.60600, 1.69670, 1.37158, 1.18186, // ATCCCT
    102.72000, 1.82797, 1.45262, 1.21134, // ATCCGA
    100.44500, 1.82797, 1.50033, 1.27151, // ATCCGC
    101.86600, 1.82797, 1.43010, 1.18329, // ATCCGG
    100.44500, 2.82797, 1.50033, 1.27151, // ATCCGM
    101.15000, 1.82797, 1.57180, 1.36343, // ATCCGT
    100.29800, 2.60867, 1.21153, 0.98116, // ATCCMA
    99.31160,  2.60867, 1.36765, 1.17679, // ATCCMC
    101.85900, 1.81426, 1.29170, 1.08014, // ATCCMG
    99.31160,  2.60867, 1.36765, 1.17679, // ATCCMM
    100.02900, 2.60867, 1.37158, 1.18186, // ATCCMT
    100.13000, 1.88060, 1.42880, 1.21337, // ATCCTA
    98.08450,  2.16543, 1.73183, 1.61918, // ATCCTC
    99.13740,  2.00709, 1.57036, 1.39809, // ATCCTG
    98.53990,  2.09558, 1.73183, 1.61918, // ATCCTM
    98.88240,  2.25241, 1.80432, 1.72190, // ATCCTT
    101.68100, 2.59137, 1.86527, 1.68529, // ATCGAA
    97.17920,  2.59137, 2.09177, 2.00140, // ATCGAC
    100.62100, 2.59137, 1.94115, 1.78917, // ATCGAG
    97.17920,  3.59137, 2.09177, 2.00140, // ATCGAM
    98.73660,  2.59137, 2.21640, 2.18291, // ATCGAT
    97.05360,  2.48599, 1.76431, 1.56306, // ATCGCA
    95.50120,  2.48599, 1.81133, 1.62596, // ATCGCC
    96.57900,  2.48599, 1.80250, 1.61408, // ATCGCG
    95.50120,  3.48599, 1.81133, 1.62596, // ATCGCM
    96.38580,  2.48599, 1.84527, 1.67187, // ATCGCT
    101.51900, 2.51279, 1.87047, 1.66318, // ATCGGA
    98.27160,  2.51279, 2.00178, 1.84135, // ATCGGC
    100.40600, 2.51279, 1.81423, 1.58873, // ATCGGG
    98.27160,  3.51279, 2.00178, 1.84135, // ATCGGM
    99.59240,  2.51279, 2.05045, 1.90890, // ATCGGT
    97.05360,  3.48599, 1.76431, 1.56306, // ATCGMA
    95.50120,  3.48599, 1.81133, 1.62596, // ATCGMC
    96.57900,  3.48599, 1.80250, 1.61408, // ATCGMG
    95.50120,  3.48599, 1.81133, 1.62596, // ATCGMM
    96.38580,  3.48599, 1.84527, 1.67187, // ATCGMT
    94.49650,  5.74004, 1.91023, 1.75033, // ATCGTA
    91.78190,  5.74004, 1.79642, 1.59626, // ATCGTC
    91.98360,  5.74004, 1.77468, 1.56738, // ATCGTG
    91.78190,  6.74004, 1.79642, 1.59626, // ATCGTM
    93.43770,  5.74004, 1.92510, 1.77080, // ATCGTT
    103.89800, 2.63253, 1.39898, 1.21060, // ATCMAA
    101.14800, 2.63253, 1.41598, 1.23273, // ATCMAC
    103.06700, 2.63253, 1.27224, 1.04987, // ATCMAG
    101.14800, 2.63253, 1.41598, 1.23273, // ATCMAM
    102.18600, 2.63253, 1.52237, 1.37424, // ATCMAT
    100.29800, 2.60867, 1.21153, 0.98116, // ATCMCA
    99.31160,  2.60867, 1.36765, 1.17679, // ATCMCC
    100.56300, 2.60867, 1.29170, 1.08014, // ATCMCG
    99.31160,  2.60867, 1.36765, 1.17679, // ATCMCM
    100.02900, 2.60867, 1.37158, 1.18186, // ATCMCT
    103.45500, 2.47865, 1.45262, 1.21134, // ATCMGA
    100.63400, 2.86764, 1.50033, 1.27151, // ATCMGC
    102.33500, 2.43500, 1.43010, 1.18329, // ATCMGG
    101.79800, 2.52093, 1.50033, 1.27151, // ATCMGM
    101.62400, 2.93767, 1.57180, 1.36343, // ATCMGT
    100.29800, 2.60867, 1.21153, 0.98116, // ATCMMA
    99.31160,  2.60867, 1.36765, 1.17679, // ATCMMC
    100.56300, 2.60867, 1.29170, 1.08014, // ATCMMG
    99.31160,  2.60867, 1.36765, 1.17679, // ATCMMM
    100.02900, 2.60867, 1.37158, 1.18186, // ATCMMT
    99.78580,  2.91501, 1.42880, 1.21337, // ATCMTA
    97.38260,  2.91501, 1.73183, 1.61918, // ATCMTC
    98.70790,  2.91501, 1.57036, 1.39809, // ATCMTG
    97.38260,  2.91501, 1.73183, 1.61918, // ATCMTM
    98.40450,  2.91501, 1.80432, 1.72190, // ATCMTT
    99.75770,  1.95287, 1.66516, 1.33296, // ATCTAA
    96.81040,  2.10597, 1.75171, 1.43822, // ATCTAC
    98.87940,  1.97042, 1.39076, 1.01745, // ATCTAG
    97.32620,  2.01570, 1.75171, 1.43822, // ATCTAM
    97.95410,  2.06311, 1.82747, 1.53253, // ATCTAT
    94.39410,  2.34531, 2.10394, 2.08340, // ATCTCA
    93.09640,  2.43305, 2.15828, 2.16463, // ATCTCC
    93.57150,  2.32915, 2.17261, 2.18622, // ATCTCG
    93.48700,  2.50184, 2.15828, 2.16463, // ATCTCM
    93.91150,  2.33922, 2.20544, 2.23597, // ATCTCT
    97.90110,  2.23262, 1.75945, 1.53289, // ATCTGA
    94.47430,  2.34736, 1.87688, 1.68888, // ATCTGC
    96.74900,  2.33669, 1.73864, 1.50577, // ATCTGG
    95.70750,  2.37413, 1.87688, 1.68888, // ATCTGM
    95.58910,  2.30373, 1.77508, 1.55335, // ATCTGT
    94.03530,  3.32915, 2.10394, 2.08340, // ATCTMA
    92.34260,  3.32915, 2.15828, 2.16463, // ATCTMC
    95.91540,  2.43739, 2.17261, 2.18622, // ATCTMG
    92.34260,  3.32915, 2.15828, 2.16463, // ATCTMM
    93.39220,  3.32915, 2.20544, 2.23597, // ATCTMT
    89.79570,  2.71915, 2.19609, 2.14834, // ATCTTA
    87.03740,  2.79570, 2.19508, 2.14685, // ATCTTC
    88.97560,  2.68554, 2.06379, 1.95715, // ATCTTG
    88.25850,  2.89599, 2.19508, 2.14685, // ATCTTM
    87.54170,  2.50281, 2.01551, 1.88888, // ATCTTT
    87.69430,  3.21014, 1.94009, 1.52771, // ATGAAA
    84.90640,  3.43194, 2.23581, 1.88999, // ATGAAC
    86.29460,  3.41237, 1.94070, 1.52842, // ATGAAG
    84.26570,  3.18113, 2.23581, 1.88999, // ATGAAM
    86.05780,  3.57926, 2.29793, 1.96929, // ATGAAT
    77.59010,  2.66595, 1.92617, 1.69681, // ATGACA
    76.45850,  2.49499, 1.79974, 1.53251, // ATGACC
    76.73400,  2.53545, 1.83494, 1.57769, // ATGACG
    76.81580,  2.47925, 1.79974, 1.53251, // ATGACM
    77.00590,  2.42981, 1.72531, 1.43843, // ATGACT
    87.18920,  3.13914, 1.93846, 1.61228, // ATGAGA
    85.16140,  3.47977, 2.08360, 1.79671, // ATGAGC
    86.00960,  3.16072, 1.91537, 1.58356, // ATGAGG
    84.25790,  2.98121, 2.08360, 1.79671, // ATGAGM
    85.54510,  3.65766, 2.10524, 1.82477, // ATGAGT
    76.96090,  3.53545, 1.92617, 1.69681, // ATGAMA
    75.72700,  3.53545, 1.79974, 1.53251, // ATGAMC
    83.49180,  3.03871, 1.83494, 1.57769, // ATGAMG
    75.72700,  3.53545, 1.79974, 1.53251, // ATGAMM
    76.55040,  3.53545, 1.72531, 1.43843, // ATGAMT
    77.50370,  2.98915, 1.86362, 1.63677, // ATGATA
    76.47770,  2.92257, 2.36282, 2.33668, // ATGATC
    76.52170,  2.68104, 2.07416, 1.92183, // ATGATG
    77.01950,  2.73797, 2.36282, 2.33668, // ATGATM
    76.31490,  3.29488, 2.34114, 2.30459, // ATGATT
    93.39180,  1.47250, 1.34877, 1.09880, // ATGCAA
    91.27020,  1.51485, 1.25738, 0.98904, // ATGCAC
    92.15600,  1.52707, 1.34120, 1.08956, // ATGCAG
    92.16840,  1.68561, 1.25738, 0.98904, // ATGCAM
    91.99400,  1.44987, 1.16966, 0.88737, // ATGCAT
    91.32580,  2.02070, 1.55970, 1.30892, // ATGCCA
    89.39570,  1.79843, 1.52228, 1.26210, // ATGCCC
    89.89600,  1.66578, 1.52564, 1.26628, // ATGCCG
    89.83910,  1.85198, 1.52228, 1.26210, // ATGCCM
    89.88690,  1.60314, 1.45460, 1.17887, // ATGCCT
    92.06640,  1.67467, 1.34368, 0.99526, // ATGCGA
    89.53870,  1.67467, 1.42637, 1.08853, // ATGCGC
    90.86740,  1.67467, 1.40515, 1.06433, // ATGCGG
    89.53870,  2.67467, 1.42637, 1.08853, // ATGCGM
    90.16700,  1.67467, 1.43259, 1.09566, // ATGCGT
    90.04480,  2.66578, 1.55970, 1.30892, // ATGCMA
    88.88190,  2.66578, 1.52228, 1.26210, // ATGCMC
    90.66910,  1.34048, 1.52564, 1.26628, // ATGCMG
    88.88190,  2.66578, 1.52228, 1.26210, // ATGCMM
    89.61950,  2.66578, 1.45460, 1.17887, // ATGCMT
    89.20970,  1.90286, 1.56710, 1.36488, // ATGCTA
    87.10230,  1.84631, 1.51788, 1.30108, // ATGCTC
    87.48290,  1.94219, 1.48527, 1.25938, // ATGCTG
    87.95190,  1.93515, 1.51788, 1.30108, // ATGCTM
    88.52250,  1.94178, 1.62826, 1.44555, // ATGCTT
    90.68350,  3.22799, 2.06265, 1.86345, // ATGGAA
    85.92580,  3.49839, 2.32679, 2.23263, // ATGGAC
    89.23890,  3.24565, 2.13094, 1.95675, // ATGGAG
    86.61960,  3.52661, 2.32679, 2.23263, // ATGGAM
    87.54000,  3.53274, 2.41193, 2.35627, // ATGGAT
    85.88730,  2.97280, 1.83263, 1.59952, // ATGGCA
    84.76600,  2.86155, 1.90176, 1.69087, // ATGGCC
    84.98440,  2.83028, 1.89036, 1.67570, // ATGGCG
    85.23800,  2.97861, 1.90176, 1.69087, // ATGGCM
    85.07950,  2.97379, 1.97253, 1.78613, // ATGGCT
    91.05960,  2.95367, 1.92663, 1.61767, // ATGGGA
    88.72110,  3.25377, 2.06233, 1.79156, // ATGGGC
    89.84800,  3.09381, 1.85858, 1.53273, // ATGGGG
    88.17960,  3.46056, 2.06233, 1.79156, // ATGGGM
    89.53940,  3.24682, 2.09716, 1.83714, // ATGGGT
    85.04500,  3.83028, 1.83263, 1.59952, // ATGGMA
    83.76940,  3.83028, 1.90176, 1.69087, // ATGGMC
    88.40210,  2.95700, 1.89036, 1.67570, // ATGGMG
    83.76940,  3.83028, 1.90176, 1.69087, // ATGGMM
    84.45680,  3.83028, 1.97253, 1.78613, // ATGGMT
    84.69910,  3.39737, 2.22849, 1.94542, // ATGGTA
    82.72120,  3.73192, 2.38819, 2.15824, // ATGGTC
    84.30160,  3.69820, 2.28373, 2.01820, // ATGGTG
    82.95420,  3.50568, 2.38819, 2.15824, // ATGGTM
    84.21410,  3.86275, 2.54805, 2.37852, // ATGGTT
    93.31500,  2.48208, 1.34877, 1.09880, // ATGMAA
    90.83250,  2.48208, 1.25738, 0.98904, // ATGMAC
    92.34620,  2.48208, 1.34120, 1.08956, // ATGMAG
    90.83250,  2.48208, 1.25738, 0.98904, // ATGMAM
    91.74390,  2.48208, 1.16966, 0.88737, // ATGMAT
    90.04480,  2.66578, 1.55970, 1.30892, // ATGMCA
    88.88190,  2.66578, 1.52228, 1.26210, // ATGMCC
    89.89600,  2.66578, 1.52564, 1.26628, // ATGMCG
    88.88190,  2.66578, 1.52228, 1.26210, // ATGMCM
    89.61950,  2.66578, 1.45460, 1.17887, // ATGMCT
    93.54220,  2.71636, 1.34368, 0.99526, // ATGMGA
    91.68130,  3.21345, 1.42637, 1.08853, // ATGMGC
    92.14720,  2.75252, 1.40515, 1.06433, // ATGMGG
    91.66640,  2.78125, 1.42637, 1.08853, // ATGMGM
    92.11880,  3.36794, 1.43259, 1.09566, // ATGMGT
    90.04480,  2.66578, 1.55970, 1.30892, // ATGMMA
    88.88190,  2.66578, 1.52228, 1.26210, // ATGMMC
    89.89600,  2.66578, 1.52564, 1.26628, // ATGMMG
    88.88190,  2.66578, 1.52228, 1.26210, // ATGMMM
    89.61950,  2.66578, 1.45460, 1.17887, // ATGMMT
    88.72250,  2.83904, 1.56710, 1.36488, // ATGMTA
    86.64500,  2.83904, 1.51788, 1.30108, // ATGMTC
    87.88990,  2.83904, 1.48527, 1.25938, // ATGMTG
    86.64500,  2.83904, 1.51788, 1.30108, // ATGMTM
    87.77670,  2.83904, 1.62826, 1.44555, // ATGMTT
    88.06010,  2.21890, 1.90357, 1.62371, // ATGTAA
    84.62520,  2.33336, 2.03149, 1.79010, // ATGTAC
    86.93940,  2.30072, 2.03863, 1.79954, // ATGTAG
    85.82940,  2.58419, 2.03149, 1.79010, // ATGTAM
    85.96000,  2.32825, 2.03236, 1.79125, // ATGTAT
    83.34880,  2.90915, 2.63435, 2.66436, // ATGTCA
    82.10740,  2.92010, 2.70387, 2.77052, // ATGTCC
    82.42730,  2.83133, 2.50788, 2.47481, // ATGTCG
    82.67970,  3.20759, 2.70387, 2.77052, // ATGTCM
    82.88930,  2.78558, 2.48871, 2.44649, // ATGTCT
    85.65530,  2.63571, 2.18261, 1.99618, // ATGTGA
    81.99290,  2.83259, 2.38566, 2.28111, // ATGTGC
    84.15150,  2.71047, 2.26082, 2.10442, // ATGTGG
    83.14330,  2.87275, 2.38566, 2.28111, // ATGTGM
    83.05660,  2.76485, 2.28172, 2.13367, // ATGTGT
    82.69950,  3.83133, 2.63435, 2.66436, // ATGTMA
    81.50010,  3.83133, 2.70387, 2.77052, // ATGTMC
    85.30340,  2.85033, 2.50788, 2.47481, // ATGTMG
    81.50010,  3.83133, 2.70387, 2.77052, // ATGTMM
    82.13820,  3.83133, 2.48871, 2.44649, // ATGTMT
    78.60500,  3.21599, 2.69835, 2.77889, // ATGTTA
    75.35270,  3.30705, 2.72964, 2.82737, // ATGTTC
    77.31260,  3.09094, 2.69614, 2.77547, // ATGTTG
    77.05280,  3.39422, 2.72964, 2.82737, // ATGTTM
    76.97690,  3.41525, 2.55241, 2.55652, // ATGTTT
    104.69500, 3.08052, 1.65247, 1.41304, // ATMAAA
    102.35300, 3.08052, 1.83989, 1.66013, // ATMAAC
    103.83200, 3.08052, 1.68580, 1.45600, // ATMAAG
    102.35300, 3.08052, 1.83989, 1.66013, // ATMAAM
    103.25100, 3.08052, 1.95166, 1.81367, // ATMAAT
    98.08030,  3.10126, 1.67386, 1.39585, // ATMACA
    96.85750,  3.10126, 1.67189, 1.39338, // ATMACC
    97.77410,  3.10126, 1.61007, 1.31682, // ATMACG
    96.85750,  3.10126, 1.67189, 1.39338, // ATMACM
    97.54660,  3.10126, 1.74883, 1.49067, // ATMACT
    103.14700, 3.40061, 1.84262, 1.64081, // ATMAGA
    101.17100, 3.40061, 1.94509, 1.77956, // ATMAGC
    102.15400, 3.40061, 1.87611, 1.68574, // ATMAGG
    101.17100, 3.40061, 1.94509, 1.77956, // ATMAGM
    101.83100, 3.40061, 2.05478, 1.93220, // ATMAGT
    98.08030,  3.10126, 1.67386, 1.39585, // ATMAMA
    96.85750,  3.10126, 1.67189, 1.39338, // ATMAMC
    97.77410,  3.10126, 1.61007, 1.31682, // ATMAMG
    96.85750,  3.10126, 1.67189, 1.39338, // ATMAMM
    97.54660,  3.10126, 1.74883, 1.49067, // ATMAMT
    96.41120,  3.47361, 1.82769, 1.59362, // ATMATA
    94.93350,  3.47361, 1.97392, 1.78865, // ATMATC
    95.74030,  3.47361, 1.85877, 1.63444, // ATMATG
    94.93350,  3.47361, 1.97392, 1.78865, // ATMATM
    95.84220,  3.47361, 2.16147, 2.04953, // ATMATT
    103.89800, 2.63253, 1.39898, 1.21060, // ATMCAA
    101.14800, 2.63253, 1.41598, 1.23273, // ATMCAC
    103.06700, 2.63253, 1.27224, 1.04987, // ATMCAG
    101.14800, 2.63253, 1.41598, 1.23273, // ATMCAM
    102.18600, 2.63253, 1.52237, 1.37424, // ATMCAT
    100.29800, 2.60867, 1.21153, 0.98116, // ATMCCA
    99.31160,  2.60867, 1.36765, 1.17679, // ATMCCC
    100.56300, 2.60867, 1.29170, 1.08014, // ATMCCG
    99.31160,  2.60867, 1.36765, 1.17679, // ATMCCM
    100.02900, 2.60867, 1.37158, 1.18186, // ATMCCT
    102.72000, 2.82797, 1.45262, 1.21134, // ATMCGA
    100.44500, 2.82797, 1.50033, 1.27151, // ATMCGC
    101.86600, 2.82797, 1.43010, 1.18329, // ATMCGG
    100.44500, 2.82797, 1.50033, 1.27151, // ATMCGM
    101.15000, 2.82797, 1.57180, 1.36343, // ATMCGT
    100.29800, 2.60867, 1.21153, 0.98116, // ATMCMA
    99.31160,  2.60867, 1.36765, 1.17679, // ATMCMC
    100.56300, 2.60867, 1.29170, 1.08014, // ATMCMG
    99.31160,  2.60867, 1.36765, 1.17679, // ATMCMM
    100.02900, 2.60867, 1.37158, 1.18186, // ATMCMT
    99.78580,  2.91501, 1.42880, 1.21337, // ATMCTA
    97.38260,  2.91501, 1.73183, 1.61918, // ATMCTC
    98.70790,  2.91501, 1.57036, 1.39809, // ATMCTG
    97.38260,  2.91501, 1.73183, 1.61918, // ATMCTM
    98.40450,  2.91501, 1.80432, 1.72190, // ATMCTT
    100.07200, 2.13555, 1.86527, 1.68529, // ATMGAA
    95.30510,  2.23168, 2.09177, 2.00140, // ATMGAC
    98.92470,  2.14803, 1.94115, 1.78917, // ATMGAG
    97.10500,  2.39941, 2.09177, 2.00140, // ATMGAM
    97.87150,  2.41994, 2.21640, 2.18291, // ATMGAT
    94.71500,  2.19337, 1.76431, 1.56306, // ATMGCA
    92.47300,  2.21151, 1.81133, 1.62596, // ATMGCC
    96.57900,  3.48599, 1.80250, 1.61408, // ATMGCG
    93.74220,  2.51456, 1.81133, 1.62596, // ATMGCM
    95.07460,  2.63992, 1.84527, 1.67187, // ATMGCT
    99.49040,  2.08404, 1.87047, 1.66318, // ATMGGA
    96.36810,  2.17096, 2.00178, 1.84135, // ATMGGC
    98.40730,  2.00355, 1.81423, 1.58873, // ATMGGG
    97.45800,  2.21706, 2.00178, 1.84135, // ATMGGM
    97.67150,  2.18917, 2.05045, 1.90890, // ATMGGT
    97.05360,  3.48599, 1.76431, 1.56306, // ATMGMA
    95.50120,  3.48599, 1.81133, 1.62596, // ATMGMC
    97.80750,  1.94476, 1.80250, 1.61408, // ATMGMG
    95.50120,  3.48599, 1.81133, 1.62596, // ATMGMM
    96.38580,  3.48599, 1.84527, 1.67187, // ATMGMT
    93.95280,  2.54132, 1.91023, 1.75033, // ATMGTA
    92.32680,  2.88976, 1.79642, 1.59626, // ATMGTC
    93.18670,  2.53641, 1.77468, 1.56738, // ATMGTG
    92.77890,  2.70186, 1.79642, 1.59626, // ATMGTM
    93.69550,  2.84188, 1.92510, 1.77080, // ATMGTT
    103.89800, 2.63253, 1.39898, 1.21060, // ATMMAA
    101.14800, 2.63253, 1.41598, 1.23273, // ATMMAC
    103.06700, 2.63253, 1.27224, 1.04987, // ATMMAG
    101.14800, 2.63253, 1.41598, 1.23273, // ATMMAM
    102.18600, 2.63253, 1.52237, 1.37424, // ATMMAT
    100.29800, 2.60867, 1.21153, 0.98116, // ATMMCA
    99.31160,  2.60867, 1.36765, 1.17679, // ATMMCC
    100.56300, 2.60867, 1.29170, 1.08014, // ATMMCG
    99.31160,  2.60867, 1.36765, 1.17679, // ATMMCM
    100.02900, 2.60867, 1.37158, 1.18186, // ATMMCT
    102.72000, 2.82797, 1.45262, 1.21134, // ATMMGA
    100.44500, 2.82797, 1.50033, 1.27151, // ATMMGC
    101.86600, 2.82797, 1.43010, 1.18329, // ATMMGG
    100.44500, 2.82797, 1.50033, 1.27151, // ATMMGM
    101.15000, 2.82797, 1.57180, 1.36343, // ATMMGT
    100.29800, 2.60867, 1.21153, 0.98116, // ATMMMA
    99.31160,  2.60867, 1.36765, 1.17679, // ATMMMC
    100.56300, 2.60867, 1.29170, 1.08014, // ATMMMG
    99.31160,  2.60867, 1.36765, 1.17679, // ATMMMM
    100.02900, 2.60867, 1.37158, 1.18186, // ATMMMT
    99.78580,  2.91501, 1.42880, 1.21337, // ATMMTA
    97.38260,  2.91501, 1.73183, 1.61918, // ATMMTC
    98.70790,  2.91501, 1.57036, 1.39809, // ATMMTG
    97.38260,  2.91501, 1.73183, 1.61918, // ATMMTM
    98.40450,  2.91501, 1.80432, 1.72190, // ATMMTT
    99.33430,  2.86097, 1.66516, 1.33296, // ATMTAA
    96.19990,  2.86097, 1.75171, 1.43822, // ATMTAC
    97.96390,  2.86097, 1.39076, 1.01745, // ATMTAG
    96.19990,  2.86097, 1.75171, 1.43822, // ATMTAM
    97.50570,  2.86097, 1.82747, 1.53253, // ATMTAT
    94.03530,  3.32915, 2.10394, 2.08340, // ATMTCA
    92.34260,  3.32915, 2.15828, 2.16463, // ATMTCC
    93.57150,  3.32915, 2.17261, 2.18622, // ATMTCG
    92.34260,  3.32915, 2.15828, 2.16463, // ATMTCM
    93.39220,  3.32915, 2.20544, 2.23597, // ATMTCT
    97.20680,  3.03287, 1.75945, 1.53289, // ATMTGA
    93.97880,  3.03287, 1.87688, 1.68888, // ATMTGC
    95.94910,  3.03287, 1.73864, 1.50577, // ATMTGG
    93.97880,  3.03287, 1.87688, 1.68888, // ATMTGM
    94.73560,  3.03287, 1.77508, 1.55335, // ATMTGT
    94.03530,  3.32915, 2.10394, 2.08340, // ATMTMA
    92.34260,  3.32915, 2.15828, 2.16463, // ATMTMC
    93.57150,  3.32915, 2.17261, 2.18622, // ATMTMG
    92.34260,  3.32915, 2.15828, 2.16463, // ATMTMM
    93.39220,  3.32915, 2.20544, 2.23597, // ATMTMT
    89.23130,  3.37489, 2.19609, 2.14834, // ATMTTA
    85.81330,  3.37489, 2.19508, 2.14685, // ATMTTC
    88.27380,  3.37489, 2.06379, 1.95715, // ATMTTG
    85.81330,  3.37489, 2.19508, 2.14685, // ATMTTM
    87.03910,  3.37489, 2.01551, 1.88888, // ATMTTT
    109.06100, 2.21363, 1.49912, 1.18194, // ATTAAA
    105.48600, 2.02810, 1.69367, 1.41934, // ATTAAC
    107.20200, 1.84867, 1.48251, 1.16235, // ATTAAG
    106.60700, 2.22116, 1.69367, 1.41934, // ATTAAM
    108.55700, 2.57812, 1.73534, 1.47203, // ATTAAT
    101.97200, 1.73704, 1.29689, 0.91929, // ATTACA
    101.75400, 1.92633, 1.36380, 0.99134, // ATTACC
    101.72500, 2.01953, 1.30086, 0.92351, // ATTACG
    102.15900, 2.29895, 1.36380, 0.99134, // ATTACM
    103.12800, 2.02704, 1.41064, 1.04285, // ATTACT
    105.53100, 1.75373, 1.45085, 1.13485, // ATTAGA
    103.91000, 1.89732, 1.63101, 1.35266, // ATTAGC
    104.78400, 1.76662, 1.53412, 1.23394, // ATTAGG
    104.65700, 1.94457, 1.63101, 1.35266, // ATTAGM
    105.08200, 2.00863, 1.66538, 1.39565, // ATTAGT
    101.74600, 3.01953, 1.29689, 0.91929, // ATTAMA
    101.08200, 3.01953, 1.36380, 0.99134, // ATTAMC
    108.68800, 1.93078, 1.30086, 0.92351, // ATTAMG
    101.08200, 3.01953, 1.36380, 0.99134, // ATTAMM
    102.42600, 3.01953, 1.41064, 1.04285, // ATTAMT
    101.53000, 2.11986, 1.63863, 1.31333, // ATTATA
    101.72400, 2.45557, 1.81357, 1.52916, // ATTATC
    101.48600, 2.17163, 1.68931, 1.37473, // ATTATG
    102.22500, 2.51234, 1.81357, 1.52916, // ATTATM
    104.87300, 3.00576, 1.87435, 1.60667, // ATTATT
    110.72500, 1.95184, 1.36739, 1.04675, // ATTCAA
    108.55800, 2.00266, 1.36330, 1.04206, // ATTCAC
    110.97500, 2.09302, 1.36114, 1.03959, // ATTCAG
    110.39200, 1.86593, 1.36330, 1.04206, // ATTCAM
    109.51500, 2.03841, 1.47687, 1.17494, // ATTCAT
    108.59300, 1.94963, 1.32279, 0.97959, // ATTCCA
    107.19600, 1.96737, 1.31956, 0.97601, // ATTCCC
    108.53600, 2.17160, 1.40284, 1.06985, // ATTCCG
    108.44200, 1.88303, 1.31956, 0.97601, // ATTCCM
    108.05300, 2.03121, 1.42231, 1.09219, // ATTCCT
    109.85900, 2.22607, 1.48296, 1.18243, // ATTCGA
    107.72700, 2.22607, 1.38529, 1.06757, // ATTCGC
    109.51200, 2.22607, 1.41513, 1.10224, // ATTCGG
    107.72700, 3.22607, 1.38529, 1.06757, // ATTCGM
    109.12700, 2.22607, 1.53435, 1.24442, // ATTCGT
    108.67900, 3.17160, 1.32279, 0.97959, // ATTCMA
    107.19500, 3.17160, 1.31956, 0.97601, // ATTCMC
    110.27100, 1.90765, 1.40284, 1.06985, // ATTCMG
    107.19500, 3.17160, 1.31956, 0.97601, // ATTCMM
    108.09900, 3.17160, 1.42231, 1.09219, // ATTCMT
    109.16800, 1.98484, 1.40010, 1.08926, // ATTCTA
    108.02100, 2.00133, 1.49782, 1.20527, // ATTCTC
    108.33400, 2.12433, 1.48496, 1.18978, // ATTCTG
    108.91000, 1.90721, 1.49782, 1.20527, // ATTCTM
    108.75200, 2.13405, 1.62198, 1.35820, // ATTCTT
    103.17300, 1.87605, 1.50205, 1.29065, // ATTGAA
    100.22600, 1.94148, 1.71114, 1.56931, // ATTGAC
    102.85100, 1.69370, 1.54398, 1.34507, // ATTGAG
    101.77000, 1.73548, 1.71114, 1.56931, // ATTGAM
    101.26300, 1.96452, 1.85854, 1.77640, // ATTGAT
    101.75100, 1.80625, 1.27285, 0.93220, // ATTGCA
    99.86280,  1.96473, 1.33577, 1.00216, // ATTGCC
    100.77600, 2.06630, 1.32718, 0.99251, // ATTGCG
    101.63600, 1.67349, 1.33577, 1.00216, // ATTGCM
    103.00800, 2.20969, 1.37337, 1.04477, // ATTGCT
    103.69600, 1.65331, 1.42783, 1.19349, // ATTGGA
    101.61500, 1.67238, 1.51760, 1.30780, // ATTGGC
    103.01600, 1.61586, 1.41716, 1.18014, // ATTGGG
    102.32400, 1.80087, 1.51760, 1.30780, // ATTGGM
    103.01200, 1.72444, 1.55136, 1.35167, // ATTGGT
    100.77900, 3.06630, 1.27285, 0.93220, // ATTGMA
    100.02700, 3.06630, 1.33577, 1.00216, // ATTGMC
    106.52000, 1.80450, 1.32718, 0.99251, // ATTGMG
    100.02700, 3.06630, 1.33577, 1.00216, // ATTGMM
    101.46200, 3.06630, 1.37337, 1.04477, // ATTGMT
    101.04200, 1.98972, 1.54362, 1.21925, // ATTGTA
    100.83700, 2.24341, 1.72791, 1.44398, // ATTGTC
    100.92000, 2.20516, 1.65559, 1.35428, // ATTGTG
    101.52700, 2.54059, 1.72791, 1.44398, // ATTGTM
    102.52300, 2.48966, 1.79033, 1.52293, // ATTGTT
    110.52100, 3.19857, 1.36739, 1.04675, // ATTMAA
    108.65800, 3.19857, 1.36330, 1.04206, // ATTMAC
    109.93900, 3.19857, 1.36114, 1.03959, // ATTMAG
    108.65800, 3.19857, 1.36330, 1.04206, // ATTMAM
    109.55700, 3.19857, 1.47687, 1.17494, // ATTMAT
    108.67900, 3.17160, 1.32279, 0.97959, // ATTMCA
    107.19500, 3.17160, 1.31956, 0.97601, // ATTMCC
    108.53600, 3.17160, 1.40284, 1.06985, // ATTMCG
    107.19500, 3.17160, 1.31956, 0.97601, // ATTMCM
    108.09900, 3.17160, 1.42231, 1.09219, // ATTMCT
    106.88000, 1.56296, 1.48296, 1.18243, // ATTMGA
    105.38600, 1.71354, 1.38529, 1.06757, // ATTMGC
    106.10200, 1.52181, 1.41513, 1.10224, // ATTMGG
    106.39100, 1.66432, 1.38529, 1.06757, // ATTMGM
    106.47000, 1.73609, 1.53435, 1.24442, // ATTMGT
    108.67900, 3.17160, 1.32279, 0.97959, // ATTMMA
    107.19500, 3.17160, 1.31956, 0.97601, // ATTMMC
    108.53600, 3.17160, 1.40284, 1.06985, // ATTMMG
    107.19500, 3.17160, 1.31956, 0.97601, // ATTMMM
    108.09900, 3.17160, 1.42231, 1.09219, // ATTMMT
    108.88100, 3.26458, 1.40010, 1.08926, // ATTMTA
    107.89300, 3.26458, 1.49782, 1.20527, // ATTMTC
    108.54900, 3.26458, 1.48496, 1.18978, // ATTMTG
    107.89300, 3.26458, 1.49782, 1.20527, // ATTMTM
    108.75400, 3.26458, 1.62198, 1.35820, // ATTMTT
    101.44200, 1.43769, 1.08220, 0.75389, // ATTTAA
    98.32770,  1.24449, 1.02777, 0.69774, // ATTTAC
    100.65300, 1.32782, 1.05562, 0.72630, // ATTTAG
    99.45790,  1.33243, 1.02777, 0.69774, // ATTTAM
    99.00840,  1.32507, 1.05025, 0.72076, // ATTTAT
    95.18860,  1.72900, 1.19292, 0.90778, // ATTTCA
    94.35960,  1.28077, 1.15448, 0.86426, // ATTTCC
    96.04810,  1.56977, 1.16328, 0.87417, // ATTTCG
    95.75810,  1.35906, 1.15448, 0.86426, // ATTTCM
    95.47660,  1.38200, 1.17229, 0.88434, // ATTTCT
    100.67300, 1.38542, 1.00007, 0.67717, // ATTTGA
    96.94920,  1.22406, 1.03624, 0.71424, // ATTTGC
    98.82850,  1.26241, 1.01909, 0.69658, // ATTTGG
    97.65400,  1.31011, 1.03624, 0.71424, // ATTTGM
    97.41140,  1.29017, 1.04119, 0.71937, // ATTTGT
    96.27520,  2.56977, 1.19292, 0.90778, // ATTTMA
    94.48900,  2.56977, 1.15448, 0.86426, // ATTTMC
    99.83060,  1.26864, 1.16328, 0.87417, // ATTTMG
    94.48900,  2.56977, 1.15448, 0.86426, // ATTTMM
    95.89440,  2.56977, 1.17229, 0.88434, // ATTTMT
    93.15820,  1.37987, 1.13431, 0.81682, // ATTTTA
    90.04230,  1.43117, 1.10265, 0.78287, // ATTTTC
    92.41120,  1.45429, 1.03481, 0.71174, // ATTTTG
    91.91130,  1.31927, 1.10265, 0.78287, // ATTTTM
    92.80920,  2.41950, 1.03607, 0.71304, // ATTTTT
    85.98350,  1.42365, 1.05997, 0.72794, // CAAAAA
    84.04500,  1.62396, 1.28891, 0.97609, // CAAAAC
    86.13760,  1.62828, 1.11668, 0.78713, // CAAAAG
    84.15560,  1.50391, 1.28891, 0.97609, // CAAAAM
    83.81990,  1.81617, 1.31591, 1.00692, // CAAAAT
    75.98680,  1.80838, 1.51867, 1.18640, // CAAACA
    75.40170,  1.69933, 1.45700, 1.11487, // CAAACC
    76.67240,  1.70501, 1.49200, 1.15528, // CAAACG
    75.95160,  1.74682, 1.45700, 1.11487, // CAAACM
    76.51900,  1.70858, 1.34246, 0.98603, // CAAACT
    85.06040,  1.61142, 1.07529, 0.74838, // CAAAGA
    83.11750,  1.55115, 1.18789, 0.86895, // CAAAGC
    83.94300,  1.56818, 1.10766, 0.78242, // CAAAGG
    82.81850,  1.21726, 1.18789, 0.86895, // CAAAGM
    82.59900,  1.71906, 1.27292, 0.96390, // CAAAGT
    76.43800,  2.70501, 1.51867, 1.18640, // CAAAMA
    75.27270,  2.70501, 1.45700, 1.11487, // CAAAMC
    80.44910,  1.57457, 1.49200, 1.15528, // CAAAMG
    75.27270,  2.70501, 1.45700, 1.11487, // CAAAMM
    76.34600,  2.70501, 1.34246, 0.98603, // CAAAMT
    75.97000,  1.70436, 1.32188, 1.05318, // CAAATA
    73.74140,  1.45346, 1.20799, 0.92005, // CAAATC
    74.81470,  1.52960, 1.24307, 0.96041, // CAAATG
    75.22340,  1.44407, 1.20799, 0.92005, // CAAATM
    75.37530,  1.64694, 1.48202, 1.25025, // CAAATT
    89.69020,  1.84990, 1.41203, 1.25157, // CAACAA
    87.28260,  1.52425, 1.21474, 0.99866, // CAACAC
    87.74470,  1.63574, 1.30512, 1.11216, // CAACAG
    87.94960,  1.73226, 1.21474, 0.99866, // CAACAM
    87.60240,  1.65765, 1.20035, 0.98096, // CAACAT
    85.02670,  1.52822, 1.49102, 1.42202, // CAACCA
    84.26170,  1.39754, 1.36144, 1.24073, // CAACCC
    85.35100,  1.83613, 1.41150, 1.30978, // CAACCG
    85.04160,  1.55663, 1.36144, 1.24073, // CAACCM
    85.18600,  1.47834, 1.34393, 1.21687, // CAACCT
    87.97780,  1.55453, 1.17444, 0.99030, // CAACGA
    86.21400,  1.55453, 1.13424, 0.93990, // CAACGC
    87.10740,  1.55453, 1.21104, 1.03696, // CAACGG
    86.21400,  2.55453, 1.13424, 0.93990, // CAACGM
    86.72750,  1.55453, 1.14390, 0.95193, // CAACGT
    85.41710,  2.83613, 1.49102, 1.42202, // CAACMA
    84.15070,  2.83613, 1.36144, 1.24073, // CAACMC
    85.64510,  1.25046, 1.41150, 1.30978, // CAACMG
    84.15070,  2.83613, 1.36144, 1.24073, // CAACMM
    85.06310,  2.83613, 1.34393, 1.21687, // CAACMT
    84.54520,  1.32982, 1.21695, 1.04253, // CAACTA
    82.54430,  1.07795, 1.13613, 0.94042, // CAACTC
    83.57370,  1.29508, 1.21996, 1.04640, // CAACTG
    83.73470,  1.24080, 1.13613, 0.94042, // CAACTM
    83.28380,  1.05885, 0.92727, 0.69341, // CAACTT
    85.05600,  1.37536, 1.09305, 0.79133, // CAAGAA
    80.37070,  1.67544, 1.41330, 1.16345, // CAAGAC
    83.89760,  1.53797, 1.15152, 0.85567, // CAAGAG
    81.78190,  1.62428, 1.41330, 1.16345, // CAAGAM
    81.82290,  1.68479, 1.35624, 1.09371, // CAAGAT
    78.69520,  1.50030, 1.19950, 0.91657, // CAAGCA
    77.31430,  1.49183, 1.15734, 0.86867, // CAAGCC
    78.57010,  1.42710, 1.12321, 0.83052, // CAAGCG
    77.85740,  1.45519, 1.15734, 0.86867, // CAAGCM
    78.06220,  1.28273, 1.05215, 0.75298, // CAAGCT
    83.82410,  1.30885, 1.08415, 0.78258, // CAAGGA
    80.91210,  1.49679, 1.20464, 0.91660, // CAAGGC
    82.89410,  1.35783, 1.04511, 0.74070, // CAAGGG
    81.64770,  1.40760, 1.20464, 0.91660, // CAAGGM
    81.95200,  1.54146, 1.26150, 0.98226, // CAAGGT
    78.59100,  2.42710, 1.19950, 0.91657, // CAAGMA
    77.31480,  2.42710, 1.15734, 0.86867, // CAAGMC
    81.87780,  1.41406, 1.12321, 0.83052, // CAAGMG
    77.31480,  2.42710, 1.15734, 0.86867, // CAAGMM
    78.12360,  2.42710, 1.05215, 0.75298, // CAAGMT
    77.12820,  1.27570, 1.19640, 0.93765, // CAAGTA
    74.63760,  1.59418, 1.49101, 1.30450, // CAAGTC
    76.32190,  1.44150, 1.27961, 1.03715, // CAAGTG
    75.95670,  1.49695, 1.49101, 1.30450, // CAAGTM
    75.81210,  1.75498, 1.46633, 1.27225, // CAAGTT
    89.08820,  2.96203, 1.41203, 1.25157, // CAAMAA
    87.22140,  2.96203, 1.21474, 0.99866, // CAAMAC
    88.18540,  2.96203, 1.30512, 1.11216, // CAAMAG
    87.22140,  2.96203, 1.21474, 0.99866, // CAAMAM
    88.05050,  2.96203, 1.20035, 0.98096, // CAAMAT
    85.41710,  2.83613, 1.49102, 1.42202, // CAAMCA
    84.15070,  2.83613, 1.36144, 1.24073, // CAAMCC
    85.35100,  2.83613, 1.41150, 1.30978, // CAAMCG
    84.15070,  2.83613, 1.36144, 1.24073, // CAAMCM
    85.06310,  2.83613, 1.34393, 1.21687, // CAAMCT
    85.89220,  1.30804, 1.17444, 0.99030, // CAAMGA
    84.39360,  1.29280, 1.13424, 0.93990, // CAAMGC
    83.97210,  1.10525, 1.21104, 1.03696, // CAAMGG
    84.33410,  1.17417, 1.13424, 0.93990, // CAAMGM
    83.75190,  1.10783, 1.14390, 0.95193, // CAAMGT
    85.41710,  2.83613, 1.49102, 1.42202, // CAAMMA
    84.15070,  2.83613, 1.36144, 1.24073, // CAAMMC
    85.35100,  2.83613, 1.41150, 1.30978, // CAAMMG
    84.15070,  2.83613, 1.36144, 1.24073, // CAAMMM
    85.06310,  2.83613, 1.34393, 1.21687, // CAAMMT
    84.42250,  2.58806, 1.21695, 1.04253, // CAAMTA
    82.48830,  2.58806, 1.13613, 0.94042, // CAAMTC
    83.96410,  2.58806, 1.21996, 1.04640, // CAAMTG
    82.48830,  2.58806, 1.13613, 0.94042, // CAAMTM
    83.09860,  2.58806, 0.92727, 0.69341, // CAAMTT
    82.53860,  1.55345, 1.58228, 1.50907, // CAATAA
    80.01000,  1.48431, 1.43132, 1.29833, // CAATAC
    81.78550,  1.37692, 1.49654, 1.38808, // CAATAG
    80.65140,  1.74292, 1.43132, 1.29833, // CAATAM
    81.51490,  1.67154, 1.56482, 1.48416, // CAATAT
    74.79470,  1.97367, 1.78694, 2.05373, // CAATCA
    74.62010,  2.05506, 1.87371, 2.20511, // CAATCC
    75.96250,  2.22770, 1.80635, 2.08727, // CAATCG
    75.26860,  2.17270, 1.87371, 2.20511, // CAATCM
    75.25690,  2.05419, 1.87779, 2.21231, // CAATCT
    79.20560,  1.51974, 1.36550, 1.24621, // CAATGA
    77.75930,  1.65521, 1.49675, 1.43014, // CAATGC
    79.39180,  1.54637, 1.43102, 1.33697, // CAATGG
    78.15120,  1.70391, 1.49675, 1.43014, // CAATGM
    77.78080,  1.60282, 1.54090, 1.49388, // CAATGT
    75.75470,  3.22770, 1.78694, 2.05373, // CAATMA
    74.65850,  3.22770, 1.87371, 2.20511, // CAATMC
    78.14550,  1.52281, 1.80635, 2.08727, // CAATMG
    74.65850,  3.22770, 1.87371, 2.20511, // CAATMM
    75.74440,  3.22770, 1.87779, 2.21231, // CAATMT
    70.47730,  2.10922, 1.93329, 1.95732, // CAATTA
    67.78220,  2.07069, 2.04579, 2.13063, // CAATTC
    69.80550,  2.10327, 1.95478, 1.99004, // CAATTG
    69.43140,  2.36550, 2.04579, 2.13063, // CAATTM
    68.89760,  1.99123, 1.83616, 1.81167, // CAATTT
    106.30300, 1.99469, 1.62999, 1.33064, // CACAAA
    103.55000, 1.94475, 1.56215, 1.24844, // CACAAC
    105.36700, 1.94732, 1.59723, 1.29073, // CACAAG
    104.68400, 1.71882, 1.56215, 1.24844, // CACAAM
    105.02800, 2.00855, 1.81583, 1.56457, // CACAAT
    98.18960,  1.40336, 1.07277, 0.69823, // CACACA
    96.86690,  1.43483, 1.09403, 0.71909, // CACACC
    97.96050,  1.74166, 1.08854, 0.71368, // CACACG
    97.77590,  1.39946, 1.09403, 0.71909, // CACACM
    97.95670,  1.53938, 1.20908, 0.83545, // CACACT
    103.69600, 1.92094, 1.63204, 1.32031, // CACAGA
    101.12100, 1.93529, 1.50958, 1.17454, // CACAGC
    102.70100, 1.87312, 1.50902, 1.17388, // CACAGG
    102.20500, 1.77546, 1.50958, 1.17454, // CACAGM
    102.03300, 2.11215, 1.64728, 1.33886, // CACAGT
    98.03620,  2.74166, 1.07277, 0.69823, // CACAMA
    96.60370,  2.74166, 1.09403, 0.71909, // CACAMC
    102.49100, 1.62408, 1.08854, 0.71368, // CACAMG
    96.60370,  2.74166, 1.09403, 0.71909, // CACAMM
    97.86230,  2.74166, 1.20908, 0.83545, // CACAMT
    97.64900,  1.57797, 1.21455, 0.88930, // CACATA
    96.02130,  1.70517, 1.32818, 1.01698, // CACATC
    97.10960,  1.64397, 1.26461, 0.94485, // CACATG
    97.26320,  1.63327, 1.32818, 1.01698, // CACATM
    97.18600,  1.88015, 1.56600, 1.30200, // CACATT
    106.25900, 1.90716, 1.27184, 1.02419, // CACCAA
    103.64900, 1.88615, 1.31919, 1.08192, // CACCAC
    108.53300, 2.98798, 1.28414, 1.03909, // CACCAG
    105.14600, 1.64146, 1.31919, 1.08192, // CACCAM
    106.33400, 2.13523, 1.51882, 1.33658, // CACCAT
    103.26700, 1.90035, 1.11442, 0.84757, // CACCCA
    100.98000, 1.49769, 1.08234, 0.81124, // CACCCC
    102.63400, 1.75332, 1.10970, 0.84219, // CACCCG
    102.03600, 1.39501, 1.08234, 0.81124, // CACCCM
    101.95800, 1.72093, 1.20743, 0.95586, // CACCCT
    105.21100, 1.90443, 1.40549, 1.14491, // CACCGA
    102.64800, 1.90443, 1.30003, 1.01849, // CACCGC
    104.20800, 1.90443, 1.29735, 1.01535, // CACCGG
    102.64800, 2.90443, 1.30003, 1.01849, // CACCGM
    103.62300, 1.90443, 1.40655, 1.14621, // CACCGT
    102.70800, 2.75332, 1.11442, 0.84757, // CACCMA
    100.91400, 2.75332, 1.08234, 0.81124, // CACCMC
    103.05500, 1.75609, 1.10970, 0.84219, // CACCMG
    100.91400, 2.75332, 1.08234, 0.81124, // CACCMM
    101.95500, 2.75332, 1.20743, 0.95586, // CACCMT
    101.62900, 1.65751, 1.30625, 0.97574, // CACCTA
    99.71330,  1.62052, 1.25861, 0.92285, // CACCTC
    100.62200, 1.75646, 1.28755, 0.95487, // CACCTG
    100.76000, 1.57089, 1.25861, 0.92285, // CACCTM
    100.24700, 1.80814, 1.45893, 1.15173, // CACCTT
    101.11500, 2.24722, 1.49862, 1.19211, // CACGAA
    96.29780,  2.24722, 1.40991, 1.08784, // CACGAC
    99.93820,  2.24722, 1.46548, 1.15279, // CACGAG
    96.29780,  3.24722, 1.40991, 1.08784, // CACGAM
    98.37730,  2.24722, 1.62384, 1.34460, // CACGAT
    95.92710,  2.02317, 1.13880, 0.78694, // CACGCA
    94.73980,  2.02317, 1.18990, 0.84049, // CACGCC
    96.11710,  2.02317, 1.19548, 0.84641, // CACGCG
    94.73980,  3.02317, 1.18990, 0.84049, // CACGCM
    96.04510,  2.02317, 1.30168, 0.96167, // CACGCT
    100.66800, 2.22668, 1.61661, 1.32867, // CACGGA
    97.29490,  2.22668, 1.41218, 1.08478, // CACGGC
    99.66140,  2.22668, 1.49791, 1.18505, // CACGGG
    97.29490,  3.22668, 1.41218, 1.08478, // CACGGM
    98.94610,  2.22668, 1.62796, 1.34268, // CACGGT
    95.92710,  3.02317, 1.13880, 0.78694, // CACGMA
    94.73980,  3.02317, 1.18990, 0.84049, // CACGMC
    96.11710,  3.02317, 1.19548, 0.84641, // CACGMG
    94.73980,  3.02317, 1.18990, 0.84049, // CACGMM
    96.04510,  3.02317, 1.30168, 0.96167, // CACGMT
    95.22340,  2.64556, 1.33682, 0.96556, // CACGTA
    93.11050,  2.64556, 1.50426, 1.15255, // CACGTC
    94.24710,  2.64556, 1.47420, 1.11817, // CACGTG
    93.11050,  3.64556, 1.50426, 1.15255, // CACGTM
    94.85050,  2.64556, 1.68166, 1.36232, // CACGTT
    106.62500, 2.88264, 1.27184, 1.02419, // CACMAA
    103.98400, 2.88264, 1.31919, 1.08192, // CACMAC
    105.85800, 2.88264, 1.28414, 1.03909, // CACMAG
    103.98400, 2.88264, 1.31919, 1.08192, // CACMAM
    105.24100, 2.88264, 1.51882, 1.33658, // CACMAT
    102.70800, 2.75332, 1.11442, 0.84757, // CACMCA
    100.91400, 2.75332, 1.08234, 0.81124, // CACMCC
    102.63400, 2.75332, 1.10970, 0.84219, // CACMCG
    100.91400, 2.75332, 1.08234, 0.81124, // CACMCM
    101.95500, 2.75332, 1.20743, 0.95586, // CACMCT
    101.55500, 1.63879, 1.40549, 1.14491, // CACMGA
    98.61920,  1.67197, 1.30003, 1.01849, // CACMGC
    100.67300, 1.49752, 1.29735, 1.01535, // CACMGG
    100.14500, 1.57741, 1.30003, 1.01849, // CACMGM
    99.91900,  1.67838, 1.40655, 1.14621, // CACMGT
    102.70800, 2.75332, 1.11442, 0.84757, // CACMMA
    100.91400, 2.75332, 1.08234, 0.81124, // CACMMC
    102.63400, 2.75332, 1.10970, 0.84219, // CACMMG
    100.91400, 2.75332, 1.08234, 0.81124, // CACMMM
    101.95500, 2.75332, 1.20743, 0.95586, // CACMMT
    101.25200, 2.83098, 1.30625, 0.97574, // CACMTA
    99.67360,  2.83098, 1.25861, 0.92285, // CACMTC
    100.98000, 2.83098, 1.28755, 0.95487, // CACMTG
    99.67360,  2.83098, 1.25861, 0.92285, // CACMTM
    100.43600, 2.83098, 1.45893, 1.15173, // CACMTT
    100.14600, 1.93661, 1.40820, 0.98708, // CACTAA
    96.86600,  1.67882, 1.28716, 0.86259, // CACTAC
    99.14680,  1.80601, 1.34421, 0.92057, // CACTAG
    97.70400,  2.19482, 1.28716, 0.86259, // CACTAM
    98.13850,  1.87851, 1.38804, 0.96596, // CACTAT
    92.08390,  1.80225, 1.49377, 1.30937, // CACTCA
    90.69130,  1.79851, 1.61647, 1.47398, // CACTCC
    92.25000,  1.86136, 1.59089, 1.43913, // CACTCG
    91.46310,  1.91502, 1.61647, 1.47398, // CACTCM
    91.89720,  1.90898, 1.59545, 1.44532, // CACTCT
    96.98270,  1.73195, 1.33084, 0.99967, // CACTGA
    93.42160,  1.46974, 1.24836, 0.90819, // CACTGC
    95.62870,  1.68820, 1.27417, 0.93650, // CACTGG
    94.89080,  1.61920, 1.24836, 0.90819, // CACTGM
    94.51580,  1.58580, 1.29863, 0.96360, // CACTGT
    92.08680,  2.86136, 1.49377, 1.30937, // CACTMA
    90.53940,  2.86136, 1.61647, 1.47398, // CACTMC
    94.62270,  1.58960, 1.59089, 1.43913, // CACTMG
    90.53940,  2.86136, 1.61647, 1.47398, // CACTMM
    92.04480,  2.86136, 1.59545, 1.44532, // CACTMT
    87.15270,  1.44712, 1.20250, 0.87086, // CACTTA
    84.54360,  1.46324, 1.24603, 0.91857, // CACTTC
    86.61540,  1.38172, 1.13987, 0.80372, // CACTTG
    86.19140,  1.42114, 1.24603, 0.91857, // CACTTM
    85.43770,  1.32366, 1.07642, 0.73755, // CACTTT
    77.55110,  1.80267, 1.26788, 0.85510, // CAGAAA
    75.78220,  1.66915, 1.56883, 1.17696, // CAGAAC
    77.19330,  2.06214, 1.33389, 0.92273, // CAGAAG
    76.33010,  1.55564, 1.56883, 1.17696, // CAGAAM
    75.69190,  1.78891, 1.54722, 1.15273, // CAGAAT
    67.27490,  2.05657, 1.86316, 1.48913, // CAGACA
    66.36940,  1.94708, 1.87829, 1.50730, // CAGACC
    67.34780,  2.14481, 1.84872, 1.47185, // CAGACG
    67.15580,  2.06350, 1.87829, 1.50730, // CAGACM
    67.27150,  1.92767, 1.81787, 1.43517, // CAGACT
    76.27480,  1.63973, 1.31162, 0.97512, // CAGAGA
    75.06360,  1.56551, 1.40216, 1.07781, // CAGAGC
    75.11860,  1.69542, 1.37510, 1.04677, // CAGAGG
    74.36390,  1.40980, 1.40216, 1.07781, // CAGAGM
    75.06860,  1.66317, 1.42244, 1.10128, // CAGAGT
    67.34890,  3.14481, 1.86316, 1.48913, // CAGAMA
    66.32630,  3.14481, 1.87829, 1.50730, // CAGAMC
    72.13260,  1.84993, 1.84872, 1.47185, // CAGAMG
    66.32630,  3.14481, 1.87829, 1.50730, // CAGAMM
    67.48380,  3.14481, 1.81787, 1.43517, // CAGAMT
    67.48600,  1.89008, 1.66387, 1.45195, // CAGATA
    66.08600,  1.64479, 1.44213, 1.17160, // CAGATC
    66.60410,  1.80266, 1.55884, 1.31666, // CAGATG
    67.01490,  1.78567, 1.44213, 1.17160, // CAGATM
    67.07050,  1.57662, 1.54524, 1.29948, // CAGATT
    88.75260,  2.86843, 1.50906, 1.26558, // CAGCAA
    84.43760,  1.51057, 1.23853, 0.94100, // CAGCAC
    87.97060,  2.70777, 1.37079, 1.09569, // CAGCAG
    85.39080,  1.65646, 1.23853, 0.94100, // CAGCAM
    83.95270,  1.84788, 1.25884, 0.96424, // CAGCAT
    83.92640,  2.28599, 1.52389, 1.37369, // CAGCCA
    80.88600,  1.33232, 1.38859, 1.19487, // CAGCCC
    82.26860,  1.70618, 1.49104, 1.32951, // CAGCCG
    82.08730,  1.49303, 1.38859, 1.19487, // CAGCCM
    81.91090,  1.40494, 1.39173, 1.19893, // CAGCCT
    84.75760,  1.56014, 1.31804, 1.09096, // CAGCGA
    82.01970,  1.56014, 1.18841, 0.93404, // CAGCGC
    83.55610,  1.56014, 1.33545, 1.11265, // CAGCGG
    82.01970,  2.56014, 1.18841, 0.93404, // CAGCGM
    82.52780,  1.56014, 1.24375, 1.00004, // CAGCGT
    82.36200,  2.70618, 1.52389, 1.37369, // CAGCMA
    80.84540,  2.70618, 1.38859, 1.19487, // CAGCMC
    82.82470,  1.28169, 1.49104, 1.32951, // CAGCMG
    80.84540,  2.70618, 1.38859, 1.19487, // CAGCMM
    81.92860,  2.70618, 1.39173, 1.19893, // CAGCMT
    80.87330,  1.35880, 1.26239, 0.99727, // CAGCTA
    79.03070,  1.27073, 1.11691, 0.82993, // CAGCTC
    80.07710,  1.34692, 1.17465, 0.89511, // CAGCTG
    80.22230,  1.26520, 1.11691, 0.82993, // CAGCTM
    80.60950,  1.36971, 0.99619, 0.69909, // CAGCTT
    76.93730,  1.92360, 1.30527, 0.91775, // CAGGAA
    73.03030,  1.80956, 1.71768, 1.38543, // CAGGAC
    76.77670,  1.80158, 1.40180, 1.02141, // CAGGAG
    74.34630,  1.79866, 1.71768, 1.38543, // CAGGAM
    74.07100,  1.88337, 1.65844, 1.31439, // CAGGAT
    71.63440,  1.70623, 1.50261, 1.23637, // CAGGCA
    70.80420,  1.65031, 1.48910, 1.21974, // CAGGCC
    71.92390,  1.67690, 1.44127, 1.16143, // CAGGCG
    71.34670,  1.71225, 1.48910, 1.21974, // CAGGCM
    71.51530,  1.52233, 1.31434, 1.01143, // CAGGCT
    78.20350,  1.46590, 1.18922, 0.86688, // CAGGGA
    75.58110,  1.59706, 1.37318, 1.07562, // CAGGGC
    77.38400,  1.50701, 1.16083, 0.83602, // CAGGGG
    76.08050,  1.42139, 1.37318, 1.07562, // CAGGGM
    76.00880,  1.76521, 1.40579, 1.11416, // CAGGGT
    72.08290,  2.67690, 1.50261, 1.23637, // CAGGMA
    70.63450,  2.67690, 1.48910, 1.21974, // CAGGMC
    75.53310,  1.44475, 1.44127, 1.16143, // CAGGMG
    70.63450,  2.67690, 1.48910, 1.21974, // CAGGMM
    71.68840,  2.67690, 1.31434, 1.01143, // CAGGMT
    71.65740,  1.55059, 1.26601, 0.93187, // CAGGTA
    69.56660,  1.64298, 1.56124, 1.27615, // CAGGTC
    70.53270,  1.45611, 1.32927, 1.00258, // CAGGTG
    70.52110,  1.57150, 1.56124, 1.27615, // CAGGTM
    71.23540,  2.02706, 1.54529, 1.25665, // CAGGTT
    86.69550,  2.87706, 1.50906, 1.26558, // CAGMAA
    84.42120,  2.87706, 1.23853, 0.94100, // CAGMAC
    85.65230,  2.87706, 1.37079, 1.09569, // CAGMAG
    84.42120,  2.87706, 1.23853, 0.94100, // CAGMAM
    85.15600,  2.87706, 1.25884, 0.96424, // CAGMAT
    82.36200,  2.70618, 1.52389, 1.37369, // CAGMCA
    80.84540,  2.70618, 1.38859, 1.19487, // CAGMCC
    82.26860,  2.70618, 1.49104, 1.32951, // CAGMCG
    80.84540,  2.70618, 1.38859, 1.19487, // CAGMCM
    81.92860,  2.70618, 1.39173, 1.19893, // CAGMCT
    81.11060,  1.15458, 1.31804, 1.09096, // CAGMGA
    79.25490,  1.46342, 1.18841, 0.93404, // CAGMGC
    80.83190,  1.50099, 1.33545, 1.11265, // CAGMGG
    79.98270,  1.34228, 1.18841, 0.93404, // CAGMGM
    79.99740,  1.60823, 1.24375, 1.00004, // CAGMGT
    82.36200,  2.70618, 1.52389, 1.37369, // CAGMMA
    80.84540,  2.70618, 1.38859, 1.19487, // CAGMMC
    82.26860,  2.70618, 1.49104, 1.32951, // CAGMMG
    80.84540,  2.70618, 1.38859, 1.19487, // CAGMMM
    81.92860,  2.70618, 1.39173, 1.19893, // CAGMMT
    80.88730,  2.42173, 1.26239, 0.99727, // CAGMTA
    79.24280,  2.42173, 1.11691, 0.82993, // CAGMTC
    80.43950,  2.42173, 1.17465, 0.89511, // CAGMTG
    79.24280,  2.42173, 1.11691, 0.82993, // CAGMTM
    79.90990,  2.42173, 0.99619, 0.69909, // CAGMTT
    77.14670,  1.80674, 1.59630, 1.43276, // CAGTAA
    73.73930,  1.99090, 1.68936, 1.55986, // CAGTAC
    75.81340,  1.69847, 1.60659, 1.44663, // CAGTAG
    74.92510,  2.24353, 1.68936, 1.55986, // CAGTAM
    74.87760,  2.03575, 1.75277, 1.64850, // CAGTAT
    69.47230,  2.29426, 2.22866, 2.44113, // CAGTCA
    68.20030,  2.36462, 2.23318, 2.44855, // CAGTCC
    69.27580,  2.42133, 2.08051, 2.20180, // CAGTCG
    69.47170,  2.64040, 2.23318, 2.44855, // CAGTCM
    69.21620,  2.47902, 2.22287, 2.43162, // CAGTCT
    73.00570,  1.84600, 1.67817, 1.64669, // CAGTGA
    69.98340,  1.98579, 2.00998, 2.15846, // CAGTGC
    72.04390,  1.94424, 1.72222, 1.71194, // CAGTGG
    71.58680,  2.32889, 2.00998, 2.15846, // CAGTGM
    70.91140,  2.07966, 1.92874, 2.02894, // CAGTGT
    69.06540,  3.42133, 2.22866, 2.44113, // CAGTMA
    67.81110,  3.42133, 2.23318, 2.44855, // CAGTMC
    72.22370,  1.94413, 2.08051, 2.20180, // CAGTMG
    67.81110,  3.42133, 2.23318, 2.44855, // CAGTMM
    68.96240,  3.42133, 2.22287, 2.43162, // CAGTMT
    64.39100,  2.29479, 2.28022, 2.27826, // CAGTTA
    61.19080,  2.12268, 2.24417, 2.22444, // CAGTTC
    63.58950,  2.24684, 2.17729, 2.12576, // CAGTTG
    63.50360,  2.59511, 2.24417, 2.22444, // CAGTTM
    62.12220,  2.11513, 2.17873, 2.12787, // CAGTTT
    106.55700, 3.19132, 1.62999, 1.33064, // CAMAAA
    103.61400, 3.19132, 1.56215, 1.24844, // CAMAAC
    105.30100, 3.19132, 1.59723, 1.29073, // CAMAAG
    103.61400, 3.19132, 1.56215, 1.24844, // CAMAAM
    104.86100, 3.19132, 1.81583, 1.56457, // CAMAAT
    98.03620,  2.74166, 1.07277, 0.69823, // CAMACA
    96.60370,  2.74166, 1.09403, 0.71909, // CAMACC
    97.96050,  2.74166, 1.08854, 0.71368, // CAMACG
    96.60370,  2.74166, 1.09403, 0.71909, // CAMACM
    97.86230,  2.74166, 1.20908, 0.83545, // CAMACT
    103.50100, 3.28987, 1.63204, 1.32031, // CAMAGA
    100.77100, 3.28987, 1.50958, 1.17454, // CAMAGC
    102.44900, 3.28987, 1.50902, 1.17388, // CAMAGG
    100.77100, 3.28987, 1.50958, 1.17454, // CAMAGM
    101.99000, 3.28987, 1.64728, 1.33886, // CAMAGT
    98.03620,  2.74166, 1.07277, 0.69823, // CAMAMA
    96.60370,  2.74166, 1.09403, 0.71909, // CAMAMC
    97.96050,  2.74166, 1.08854, 0.71368, // CAMAMG
    96.60370,  2.74166, 1.09403, 0.71909, // CAMAMM
    97.86230,  2.74166, 1.20908, 0.83545, // CAMAMT
    97.52790,  3.17760, 1.21455, 0.88930, // CAMATA
    95.87150,  3.17760, 1.32818, 1.01698, // CAMATC
    96.93890,  3.17760, 1.26461, 0.94485, // CAMATG
    95.87150,  3.17760, 1.32818, 1.01698, // CAMATM
    96.99620,  3.17760, 1.56600, 1.30200, // CAMATT
    106.62500, 2.88264, 1.27184, 1.02419, // CAMCAA
    103.98400, 2.88264, 1.31919, 1.08192, // CAMCAC
    105.85800, 2.88264, 1.28414, 1.03909, // CAMCAG
    103.98400, 2.88264, 1.31919, 1.08192, // CAMCAM
    105.24100, 2.88264, 1.51882, 1.33658, // CAMCAT
    102.70800, 2.75332, 1.11442, 0.84757, // CAMCCA
    100.91400, 2.75332, 1.08234, 0.81124, // CAMCCC
    102.63400, 2.75332, 1.10970, 0.84219, // CAMCCG
    100.91400, 2.75332, 1.08234, 0.81124, // CAMCCM
    101.95500, 2.75332, 1.20743, 0.95586, // CAMCCT
    105.21100, 2.90443, 1.40549, 1.14491, // CAMCGA
    102.64800, 2.90443, 1.30003, 1.01849, // CAMCGC
    104.20800, 2.90443, 1.29735, 1.01535, // CAMCGG
    102.64800, 2.90443, 1.30003, 1.01849, // CAMCGM
    103.62300, 2.90443, 1.40655, 1.14621, // CAMCGT
    102.70800, 2.75332, 1.11442, 0.84757, // CAMCMA
    100.91400, 2.75332, 1.08234, 0.81124, // CAMCMC
    102.63400, 2.75332, 1.10970, 0.84219, // CAMCMG
    100.91400, 2.75332, 1.08234, 0.81124, // CAMCMM
    101.95500, 2.75332, 1.20743, 0.95586, // CAMCMT
    101.25200, 2.83098, 1.30625, 0.97574, // CAMCTA
    99.67360,  2.83098, 1.25861, 0.92285, // CAMCTC
    100.98000, 2.83098, 1.28755, 0.95487, // CAMCTG
    99.67360,  2.83098, 1.25861, 0.92285, // CAMCTM
    100.43600, 2.83098, 1.45893, 1.15173, // CAMCTT
    101.73900, 1.89747, 1.49862, 1.19211, // CAMGAA
    97.01200,  1.79610, 1.40991, 1.08784, // CAMGAC
    100.52100, 1.83581, 1.46548, 1.15279, // CAMGAG
    98.94430,  1.80062, 1.40991, 1.08784, // CAMGAM
    98.73530,  2.06840, 1.62384, 1.34460, // CAMGAT
    95.46040,  1.85914, 1.13880, 0.78694, // CAMGCA
    93.96610,  1.85387, 1.18990, 0.84049, // CAMGCC
    96.11710,  3.02317, 1.19548, 0.84641, // CAMGCG
    95.24330,  1.62107, 1.18990, 0.84049, // CAMGCM
    95.61430,  1.90594, 1.30168, 0.96167, // CAMGCT
    101.57600, 1.91781, 1.61661, 1.32867, // CAMGGA
    98.07230,  1.79631, 1.41218, 1.08478, // CAMGGC
    100.56200, 1.83551, 1.49791, 1.18505, // CAMGGG
    99.47970,  1.83595, 1.41218, 1.08478, // CAMGGM
    99.39190,  2.16305, 1.62796, 1.34268, // CAMGGT
    95.92710,  3.02317, 1.13880, 0.78694, // CAMGMA
    94.73980,  3.02317, 1.18990, 0.84049, // CAMGMC
    99.44280,  1.76078, 1.19548, 0.84641, // CAMGMG
    94.73980,  3.02317, 1.18990, 0.84049, // CAMGMM
    96.04510,  3.02317, 1.30168, 0.96167, // CAMGMT
    95.38120,  1.86038, 1.33682, 0.96556, // CAMGTA
    93.58160,  1.96609, 1.50426, 1.15255, // CAMGTC
    94.87120,  1.87627, 1.47420, 1.11817, // CAMGTG
    94.61230,  1.98922, 1.50426, 1.15255, // CAMGTM
    94.69840,  2.28284, 1.68166, 1.36232, // CAMGTT
    106.62500, 2.88264, 1.27184, 1.02419, // CAMMAA
    103.98400, 2.88264, 1.31919, 1.08192, // CAMMAC
    105.85800, 2.88264, 1.28414, 1.03909, // CAMMAG
    103.98400, 2.88264, 1.31919, 1.08192, // CAMMAM
    105.24100, 2.88264, 1.51882, 1.33658, // CAMMAT
    102.70800, 2.75332, 1.11442, 0.84757, // CAMMCA
    100.91400, 2.75332, 1.08234, 0.81124, // CAMMCC
    102.63400, 2.75332, 1.10970, 0.84219, // CAMMCG
    100.91400, 2.75332, 1.08234, 0.81124, // CAMMCM
    101.95500, 2.75332, 1.20743, 0.95586, // CAMMCT
    105.21100, 2.90443, 1.40549, 1.14491, // CAMMGA
    102.64800, 2.90443, 1.30003, 1.01849, // CAMMGC
    104.20800, 2.90443, 1.29735, 1.01535, // CAMMGG
    102.64800, 2.90443, 1.30003, 1.01849, // CAMMGM
    103.62300, 2.90443, 1.40655, 1.14621, // CAMMGT
    102.70800, 2.75332, 1.11442, 0.84757, // CAMMMA
    100.91400, 2.75332, 1.08234, 0.81124, // CAMMMC
```
