## Supplementary material for "GPU Accelerated Adaptive Banded Event Alignment for Rapid Comparative Nanopore Signal Analysis": f5c-v0.2-beta-release: nanopolish_index.c

```
//---------------------------------------------------------
// Copyright 2017 Ontario Institute for Cancer Research
// Written by Jared Simpson
//
// nanopolish index - build an index from FASTA/FASTQ file
// and the associated signal-level data
//
//---------------------------------------------------------
//

//
// Getopt
//

#include <iostream>
#include <fstream>
#include <sstream>
#include <getopt.h>
#include <vector>
#include <dirent.h>

#include "nanopolish_read_db.h"
#include "fast5lite.h"

// ref: http://stackoverflow.com/a/612176/717706
// return true if the given name is a directory
bool is_directory(const std::string& file_name){
    auto dir = opendir(file_name.c_str());
    if(not dir) {
        return false;
    }
    closedir(dir);
    return true;
}

// Split a string into parts based on the delimiter (nanopolish_common.cpp)
std::vector<std::string> split(std::string in, char delimiter);

std::vector< std::string > list_directory(const std::string& file_name)
{
    std::vector< std::string > res;
    DIR* dir;
    struct dirent *ent;

    dir = opendir(file_name.c_str());
    if(not dir) {
        return res;
    }
    while((ent = readdir(dir)) != nullptr) {
        res.push_back(ent->d_name);
    }
    closedir(dir);
    return res;
}


static const char *INDEX_USAGE_MESSAGE =
"Usage: f5c index [OPTIONS] -d nanopore_raw_file_directory reads.fastq\n"
"Build an index mapping from basecalled reads to the signals measured by the sequencer\n"
"f5c index is equivalent to nanopolish index by Jared Simpson\n"
"\n"
"  -h, --help                           display this help and exit\n"
"  -v, --verbose                        display verbose output\n"
"  -d, --directory                      path to the directory containing the raw ONT signal files. This option can be given multiple times.\n"
"  -s, --sequencing-summary             the sequencing summary file from albacore, providing this option will make indexing much faster\n"
"  -f, --summary-fofn                   file containing the paths to the sequencing summary files (one per line)\n"
"\n\n"
;

namespace opt
{
    static unsigned int verbose = 0;
    static std::vector<std::string> raw_file_directories;
    static std::string reads_file;
    static std::vector<std::string> sequencing_summary_files;
    static std::string sequencing_summary_fofn;
}
//static std::ostream* os_p;

void index_file_from_map(ReadDB& read_db, const std::string& fn, const std::map<std::string, std::string>& fast5_to_read_name_map)
{
    //PROFILE_FUNC("index_file_from_map")

    // Check if this fast5 file is in the map
    size_t last_dir_pos = fn.find_last_of("/");
    std::string fast5_basename = last_dir_pos == std::string::npos ? fn : fn.substr(last_dir_pos + 1);

    const auto& iter = fast5_to_read_name_map.find(fast5_basename);
    if(iter != fast5_to_read_name_map.end()) {
        if(read_db.has_read(iter->second)) {
            read_db.add_signal_path(iter->second.c_str(), fn);
        }
    } else {
        if(opt::verbose > 0) {
            fprintf(stderr, "Could not find read %s in sequencing summary file\n", fn.c_str());
        }
    }
} // process_file

void index_file_from_fast5(ReadDB& read_db, const std::string& fn, const std::map<std::string, std::string>& fast5_to_read_name_map)
{
    //PROFILE_FUNC("index_file_from_fast5")

    char* fast5_path =(char*)malloc(fn.size()+1);
    strcpy(fast5_path, fn.c_str());

    hid_t hdf5_file = fast5_open(fast5_path);
    if(hdf5_file < 0) {
        fprintf(stderr, "could not open fast5 file: %s\n", fast5_path);
    }

    std::string read_id = fast5_get_read_id(hdf5_file);
    if(read_id != "") {
        read_db.add_signal_path(read_id, fn);
    }
    free(fast5_path);

    fast5_close(hdf5_file);
} // process_file

void index_path(ReadDB& read_db, const std::string& path, const std::map<std::string, std::string>& fast5_to_read_name_map)
{
    fprintf(stderr, "[readdb] indexing %s\n", path.c_str());
    if (is_directory(path)) {
        auto dir_list = list_directory(path);
        for (const auto& fn : dir_list) {
            if(fn == "." or fn == "..") {
                continue;
            }

            std::string full_fn = path + "/" + fn;
            if(is_directory(full_fn)) {
                // recurse
                index_path(read_db, full_fn, fast5_to_read_name_map);
            } else if (full_fn.find(".fast5") !=  std::string::npos) {
                if(!fast5_to_read_name_map.empty()) {
                    index_file_from_map(read_db, full_fn, fast5_to_read_name_map);
                } else {
                    index_file_from_fast5(read_db, full_fn, fast5_to_read_name_map);
                }
            }
        }
    }
} // process_path

// read sequencing summary files from the fofn and add them to the list
void process_summary_fofn()
{
    if(opt::sequencing_summary_fofn.empty()) {
        return;
    }

    // open
    std::ifstream in_file(opt::sequencing_summary_fofn.c_str());
    if(in_file.bad()) {
        fprintf(stderr, "error: could not file %s\n", opt::sequencing_summary_fofn.c_str());
        exit(EXIT_FAILURE);
    }

    // read
    std::string filename;
    while(getline(in_file, filename)) {
        opt::sequencing_summary_files.push_back(filename);
    }
}

void exit_bad_header(const std::string& str, const std::string& filename)
{
    fprintf(stderr, "Could not find %s column in the header of %s\n", str.c_str(), filename.c_str());
    exit(EXIT_FAILURE);
}

//
void parse_sequencing_summary(const std::string& filename, std::map<std::string, std::string>& out_map)
{
    // open
    std::ifstream in_file(filename.c_str());
    if(in_file.bad()) {
        fprintf(stderr, "error: could not file %s\n", filename.c_str());
        exit(EXIT_FAILURE);
    }

    // read header to get the column index of the read and file name
    std::string header;
    getline(in_file, header);
    std::vector<std::string> fields = split(header, '\t');

    const std::string READ_NAME_STR = "read_id";
    const std::string FILENAME_STR = "filename";
    size_t filename_idx = SIZE_MAX;
    size_t read_name_idx = SIZE_MAX;

    for(size_t i = 0; i < fields.size(); ++i) {
        if(fields[i] == READ_NAME_STR) {
            read_name_idx = i;
        }

        if(fields[i] == FILENAME_STR) {
            filename_idx = i;
        }
    }

    if(filename_idx == SIZE_MAX) {
        exit_bad_header(FILENAME_STR, filename);
    }

    if(read_name_idx == SIZE_MAX) {
        exit_bad_header(READ_NAME_STR, filename);
    }

    // read records and add to map
    std::string line;
    while(getline(in_file, line)) {
        fields = split(line, '\t');
        std::string fast5_filename = fields[filename_idx];
        std::string read_name = fields[read_name_idx];
        out_map[fast5_filename] = read_name;
    }
}

static const char* shortopts = "vhd:f:s:";

enum {
    OPT_HELP = 1,
    OPT_VERSION,
    OPT_LOG_LEVEL,
};

static const struct option longopts[] = {
    { "help",                      no_argument,       NULL, OPT_HELP },
    { "version",                   no_argument,       NULL, OPT_VERSION },
    { "verbose",                   no_argument,       NULL, 'v' },
    { "directory",                 required_argument, NULL, 'd' },
    { "sequencing-summary-file",   required_argument, NULL, 's' },
    { "summary-fofn",              required_argument, NULL, 'f' },
    { NULL, 0, NULL, 0 }
};

void parse_index_options(int argc, char** argv)
{
    bool die = false;
    //std::vector< std::string> log_level;
    for (int c; (c = getopt_long(argc, argv, shortopts, longopts, NULL)) != -1;) {
        std::istringstream arg(optarg != NULL ? optarg : "");
        switch (c) {
            case OPT_HELP:
                std::cout << INDEX_USAGE_MESSAGE;
                exit(EXIT_SUCCESS);
            // case OPT_VERSION:
            //     std::cout << INDEX_VERSION_MESSAGE;
            //     exit(EXIT_SUCCESS);
            // case OPT_LOG_LEVEL:
            //     log_level.push_back(arg.str());
            //     break;
            case 'v': opt::verbose++; break;
            case 's': opt::sequencing_summary_files.push_back(arg.str()); break;
            case 'd': opt::raw_file_directories.push_back(arg.str()); break;
            case 'f': arg >> opt::sequencing_summary_fofn; break;
        }
    }

    // set log levels
    // auto default_level = (int)logger::warning + opt::verbose;
    // logger::Logger::set_default_level(default_level);
    // logger::Logger::set_levels_from_options(log_level, &std::clog);

    if (argc - optind < 1) {
        std::cerr << "[f5c index]  not enough arguments\n";
        die = true;
    }

    if (argc - optind > 1) {
        std::cerr << "[f5c index] too many arguments\n";
        die = true;
    }

    if (die)
    {
        std::cout << "\n" << INDEX_USAGE_MESSAGE;
        exit(EXIT_FAILURE);
    }

    opt::reads_file = argv[optind++];
}

int index_main(int argc, char** argv)
{
    parse_index_options(argc, argv);


    // Read a map from fast5 files to read name from the sequencing summary files (if any)
    process_summary_fofn();
    std::map<std::string, std::string> fast5_to_read_name_map;
    for(const auto& ss_filename : opt::sequencing_summary_files) {
        if(opt::verbose > 2) {
            fprintf(stderr, "summary: %s\n", ss_filename.c_str());
        }
        parse_sequencing_summary(ss_filename, fast5_to_read_name_map);
    }

    // import read names, and possibly fast5 paths, from the fasta/fastq file
    ReadDB read_db;
    read_db.build(opt::reads_file);
    bool all_reads_have_paths = read_db.check_signal_paths();

    // if the input fastq did not contain a complete set of paths
    // use the fofn/directory provided to augment the index
    if(!all_reads_have_paths) {

        for(const auto& dir_name : opt::raw_file_directories) {
            index_path(read_db, dir_name, fast5_to_read_name_map);
        }
    }

    size_t num_with_path = read_db.get_num_reads_with_path();
    if(num_with_path == 0) {
        fprintf(stderr, "Error: no fast5 files found\n");
        exit(EXIT_FAILURE);
    } else {
        read_db.print_stats();
        read_db.save();
    }
    return 0;
}
```
