## Supplementary material for "GPU Accelerated Adaptive Banded Event Alignment for Rapid Comparative Nanopore Signal Analysis": f5c-v0.2-beta-release: nanopolish_read_db.c

```
//---------------------------------------------------------
// Copyright 2017 Ontario Institute for Cancer Research
// Written by Jared Simpson
//---------------------------------------------------------
//
// nanopolish_read_db -- database of reads and their
// associated signal data
//
#include <assert.h>
#include <fstream>
#include <iostream>
#include <ostream>
#include <sys/stat.h>
#include <vector>
#include <zlib.h>
//#include "nanopolish_common.h"
#include "nanopolish_read_db.h"
#include <htslib/bgzf.h>
#include <htslib/faidx.h>
#include <htslib/kseq.h>
#include <map>

#define READ_DB_SUFFIX ".readdb"
#define GZIPPED_READS_SUFFIX ".index"

// Tell KSEQ what functions to use to open/read files
KSEQ_INIT(gzFile, gzread)

//
ReadDB::ReadDB() : m_fai(NULL) {}

// Split a string into parts based on the delimiter
std::vector<std::string> split(std::string in, char delimiter) {
    std::vector<std::string> out;
    size_t lastPos = 0;
    size_t pos = in.find_first_of(delimiter);

    while (pos != std::string::npos) {
        out.push_back(in.substr(lastPos, pos - lastPos));
        lastPos = pos + 1;
        pos = in.find_first_of(delimiter, lastPos);
    }
    out.push_back(in.substr(lastPos));
    return out;
}

//
void ReadDB::build(const std::string& input_reads_filename) {
    // generate output filename
    m_indexed_reads_filename = input_reads_filename + GZIPPED_READS_SUFFIX;

    // Populate database with read names and convert the fastq
    // input into fasta for faidx
    import_reads(input_reads_filename, m_indexed_reads_filename);

    // build faidx
    int ret = fai_build(m_indexed_reads_filename.c_str());
    if (ret != 0) {
        fprintf(stderr, "Error running faidx_build on %s\n",
                m_indexed_reads_filename.c_str());
        exit(EXIT_FAILURE);
    }

    m_fai = NULL;
}

//
void ReadDB::load(const std::string& input_reads_filename) {
    // generate input filenames
    m_indexed_reads_filename = input_reads_filename + GZIPPED_READS_SUFFIX;
    std::string in_filename = m_indexed_reads_filename + READ_DB_SUFFIX;

    //
    std::ifstream in_file(in_filename.c_str());
    bool success = false;
    if (in_file.good()) {
        // read the database
        std::string line;
        while (getline(in_file, line)) {
            std::vector<std::string> fields = split(line, '\t');

            std::string name = "";
            std::string path = "";
            if (fields.size() == 2) {
                name = fields[0];
                path = fields[1];
                m_data[name].signal_data_path = path;
            }
        }

        // load faidx
        m_fai = fai_load3(m_indexed_reads_filename.c_str(), NULL, NULL, 0);
        if (m_fai != NULL) {
            success = true;
        }
    }

    if (!success) {
        fprintf(stderr,
                "error: could not load the index files for input file %s\n",
                input_reads_filename.c_str());
        fprintf(
            stderr,
            "Please run f5c (or nanopolish) index on your reads (see documentation)\n");
        exit(EXIT_FAILURE);
    }
}

ReadDB::~ReadDB() {
    if (m_fai != NULL) {
        fai_destroy(m_fai);
    }
}

//
void ReadDB::import_reads(const std::string& input_filename,
                          const std::string& out_fasta_filename) {
    // Open readers
    FILE* read_fp = fopen(input_filename.c_str(), "r");
    if (read_fp == NULL) {
        fprintf(stderr, "error: could not open %s for read\n",
                input_filename.c_str());
        exit(EXIT_FAILURE);
    }

    gzFile gz_read_fp = gzdopen(fileno(read_fp), "r");
    if (gz_read_fp == NULL) {
        fprintf(stderr, "error: could not open %s using gzdopen\n",
                input_filename.c_str());
        exit(EXIT_FAILURE);
    }

    // Open writers
    FILE* write_fp = fopen(out_fasta_filename.c_str(), "w");
    if (write_fp == NULL) {
        fprintf(stderr, "error: could not open %s for write\n",
                out_fasta_filename.c_str());
        exit(EXIT_FAILURE);
    }

    BGZF* bgzf_write_fp = bgzf_dopen(fileno(write_fp), "w");
    if (bgzf_write_fp == NULL) {
        fprintf(stderr, "error: could not open %s for bgzipped write\n",
                out_fasta_filename.c_str());
        exit(EXIT_FAILURE);
    }

    // read input sequences, add to DB and convert to fasta
    int ret = 0;
    kseq_t* seq = kseq_init(gz_read_fp);
    while ((ret = kseq_read(seq)) >= 0) {
        // Check for a path to the fast5 file in the comment of the read
        std::string path = "";
        if (seq->comment.l > 0) {
            // This splitting code implicitly handles both the 2 and 3 field
            // fasta format that poretools will output. The FAST5 path
            // is always the last field.
            std::vector<std::string> fields = split(seq->comment.s, ' ');
            path = fields.back();

            // as a sanity check we require the path name to end in ".fast5"
            if (path.substr(path.length() - 6) != ".fast5") {
                path = "";
            }
        }

        // sanity check that the read does not exist in the database
        auto iter = m_data.find(seq->name.s);
        if (iter != m_data.end()) {
            fprintf(stderr,
                    "Error: duplicate read name %s found in fasta file\n",
                    seq->name.s);
            exit(EXIT_FAILURE);
        }

        // add path
        add_signal_path(seq->name.s, path);

        // write sequence in gzipped fasta for fai indexing later
        std::string out_record;
        out_record += ">";
        out_record += seq->name.s;
        out_record += "\n";
        out_record += seq->seq.s;
        out_record += "\n";
        size_t write_length =
            bgzf_write(bgzf_write_fp, out_record.c_str(), out_record.length());
        if (write_length != out_record.length()) {
            fprintf(stderr, "error in bgzf_write, aborting\n");
            exit(EXIT_FAILURE);
        }
    }

    // check for abnormal exit conditions
    if (ret <= -2) {
        fprintf(stderr,
                "kseq_read returned %d indicating an error with the input file "
                "%s\n",
                ret, input_filename.c_str());
        exit(EXIT_FAILURE);
    }

    // cleanup
    kseq_destroy(seq);

    gzclose(gz_read_fp);
    fclose(read_fp);

    bgzf_close(bgzf_write_fp);
    fclose(write_fp);
}

//
void ReadDB::add_signal_path(const std::string& read_id,
                             const std::string& path) {
    m_data[read_id].signal_data_path = path;
}

bool ReadDB::has_read(const std::string& read_id) const {
    const auto& iter = m_data.find(read_id);
    return iter != m_data.end();
}

//
std::string ReadDB::get_signal_path(const std::string& read_id) const {
    const auto& iter = m_data.find(read_id);
    if (iter == m_data.end()) {
        return "";
    } else {
        return iter->second.signal_data_path;
    }
}

//
std::string ReadDB::get_read_sequence(const std::string& read_id) const {
    assert(m_fai != NULL);

    int length;
    char* seq;

    // this call is not threadsafe
    seq = fai_fetch(m_fai, read_id.c_str(), &length);

    if (seq == NULL) {
        return "";
    }

    std::string out(seq);
    free(seq);
    return out;
}

//
void ReadDB::save() const {
    std::string out_filename = m_indexed_reads_filename + READ_DB_SUFFIX;

    std::ofstream out_file(out_filename.c_str());

    for (const auto& iter : m_data) {
        const ReadDBData& entry = iter.second;
        out_file << iter.first << "\t" << entry.signal_data_path << "\n";
    }
}

//
size_t ReadDB::get_num_reads_with_path() const {
    size_t num_reads_with_path = 0;
    for (const auto& iter : m_data) {
        if (iter.second.signal_data_path != "") {
            num_reads_with_path += 1;
        }
    }
    return num_reads_with_path;
}

bool ReadDB::check_signal_paths() const {
    size_t num_reads_with_path = get_num_reads_with_path();
    return num_reads_with_path == m_data.size();
}

//
void ReadDB::print_stats() const {
    size_t num_reads_with_path = 0;
    for (const auto& iter : m_data) {
        num_reads_with_path += iter.second.signal_data_path != "";
    }
    fprintf(stderr,
            "[readdb] num reads: %zu, num reads with path to fast5: %zu\n",
            m_data.size(), num_reads_with_path);
}
```
