## Supplementary material for "GPU Accelerated Adaptive Banded Event Alignment for Rapid Comparative Nanopore Signal Analysis": f5c-v0.2-beta-release: nanopolish_read_db.h

```
//---------------------------------------------------------
// Copyright 2017 Ontario Institute for Cancer Research
// Written by Jared Simpson
//---------------------------------------------------------
//
// nanopolish_read_db -- database of reads and their
// associated signal data
//
#ifndef NANOPOLISH_READ_DB
#define NANOPOLISH_READ_DB

#include <map>
#include <string>

#include <htslib/faidx.h>

struct ReadDBData {
    // path to the signal-level data for this read
    std::string signal_data_path;
};

class ReadDB {
  public:
    ReadDB();
    ~ReadDB();

    //
    // I/O
    //

    //  construct the database from an input reads file
    void build(const std::string& reads_filename);

    // save the database to disk
    void save() const;

    // restore the database from disk
    void load(const std::string& reads_filename);

    //
    // Data Access
    //

    // set the signal path for the given read
    void add_signal_path(const std::string& read_id, const std::string& path);

    // returns the path to the signal data for the given read
    std::string get_signal_path(const std::string& read_id) const;

    // returns true if a read with this ID is in the DB
    bool has_read(const std::string& read_id) const;

    // returns the basecalled sequence for the given read
    std::string get_read_sequence(const std::string& read_id) const;

    // returns the number of reads in the database
    size_t get_num_reads() const { return m_data.size(); }

    //
    // Summaries and sanity checks
    //

    // return the number of reads with a fast5 file
    size_t get_num_reads_with_path() const;

    // returns true if all reads in the database have paths to their
    // signal-level data
    bool check_signal_paths() const;

    // print some summary stats about the database
    void print_stats() const;

  private:
    //
    void import_reads(const std::string& input_filename,
                      const std::string& output_fasta_filename);

    // the filename of the indexed data, after converting to fasta
    std::string m_indexed_reads_filename;

    //
    std::map<std::string, ReadDBData> m_data;

    //
    faidx_t* m_fai;
};

#endif
```
